# Triphenyl phosphate is a selective PPARγ modulator that does not induce brite adipogenesis *in vitro* and *in vivo*

**DOI:** 10.1101/626390

**Authors:** Stephanie Kim, Nabil Rabhi, Benjamin C. Blum, Ryan Hekman, Kieran Wynne, Andrew Emili, Stephen Farmer, Jennifer J. Schlezinger

**Affiliations:** Boston University Superfund Research Program, Boston University, MA 02118 USA; Boston University School of Public Health, Department of Environmental Health, MA 02118 USA; Boston University School of Medicine, Department of Biochemistry, MA 02118 USA; Boston University School of Medicine, Center for Network Systems Biology, MA 02118 USA

**Keywords:** triphenyl phosphate, PPARγ, post-translational modification, selective PPARγ modulator

## Abstract

Triphenyl phosphate (TPhP) is an environmental PPARγ ligand, and growing evidence suggests that it is a metabolic disruptor. We have shown previously that the structurally similar ligand, tributyltin, does not induce brite adipocyte gene expression. Here, using *in vivo* and *in vitro* models, we tested the hypothesis that TPhP is a selective PPARγ ligand, which fails to induce brite adipogenesis. C57BL/6J male mice were fed either a low or very high fat diet for 13 weeks. From weeks 7-13, mice were injected intraperitoneally, daily, with vehicle, rosiglitazone (Rosi), or TPhP (10 mg/kg). Compared to Rosi, TPhP did not induce expression of browning-related genes (e.g. *Elovl3, Cidea, Acaa2, CoxIV*) in mature adipocytes isolated from inguinal adipose. To determine if this resulted from an effect directly on the adipocytes, 3T3-L1 cells and primary human preadipocytes were differentiated into adipocytes in the presence of Rosi or TPhP. Rosi, but not TPhP, induced expression of brite adipocyte genes, mitochondrial biogenesis and cellular respiration. Further, Rosi and TPhP induced distinct proteomes and phosphoproteomes; Rosi enriched more regulatory pathways related to fatty acid oxidation and mitochondrial proteins. We assessed the role of phosphorylation of PPARγ in these differences in 3T3-L1 cells. Only Rosi protected PPARγ from phosphorylation at Ser273. TPhP gained the ability to stimulate brite adipocyte gene expression in the presence of the CDK5 inhibitor and in 3T3-L1 cells expressing alanine at position 273. We conclude that TPhP is a selective PPARγ modulator that fails to protect PPARγ from phosphorylation at ser273.

## Introduction

Obesity is characterized by an increased adipose tissue mass driven by a chronic caloric excess (Schwartz et al. 2017). The increasing prevalence of obesity has been largely explained by sedentary lifestyle, changes in diet, genetic predisposition and aging. However, recent epidemiological studies suggested a strong contribution of environmental factors including chemicals. Indeed, a specific class of environmental compounds that interfere with metabolism and termed metabolic disrupting chemical (MDC) has been suggested to be contributing to the rapid rise in obesity (Heindel et al. 2017).

Organophosphate esters (OPEs) are member of the MDC class that are used extensively as flame retardants and plasticizers in consumer products from furniture to nail polish (Wang et al. 2019b; Young et al. 1998). Of a particular concern, 92% of American were tested positive for the presence of OPEs including triphenyl phosphate (TPhP) and tris(1,3-dichloro-2-propyl) phosphate in a large-scale urine samples analysis (Ospina et al. 2018). Moreover, studies evaluating the metabolic effects of TPhP during early life exposures showed evidence of impaired glucose homeostasis in adults following early life exposures to TPhP (Green et al. 2017; Patisaul et al. 2013; Wang et al. 2019a). This effect was associated with increased body weight, liver weight, lipid-related metabolites, and fat mass and suppressed pyruvate metabolism and tricarboxylic acid cycles in adults (Green et al. 2017; Wang et al. 2019a).

Studies have demonstrated the pro-adipogenic activity of TPhP in 3T3-L1 mouse preadipocytes, mouse bone marrow-derived multipotent cells (BMS2 cells) and human primary preadipocytes (Cano-Sancho et al. 2017; Pillai et al. 2014; Tung et al. 2017a; Tung et al. 2017b). Further, studies demonstrate that this effect is due to a direct action on peroxisome proliferator-activated receptor γ (PPARγ) (Pillai et al. 2014; Tung et al. 2017a), a master regulator of adipocytes differentiation and function (Lefterova et al. 2014; Ma et al. 2018).

At least two subtypes of adipocytes exist. White adipocytes are characterized by a single large lipid droplet play an important role in storing excess energy in forms of triglycerides (Ma et al. 2018). Brown adipocytes contain multiple, small lipid droplets, have a high density of mitochondria, a high capacity for fatty acid oxidation and play an important role in maintaining body temperature. The thermogenic activity of brown adipocytes is due mainly to the expression of uncoupling protein 1 (UCP1)(Ma et al. 2018). Recent works have demonstrated that thermogenic cells named beige/brite can emerge within the white depots upon cold challenge. Those cells express both white and brown adipocyte features (Chen et al. 2016; Ma et al. 2018). The ability of PPARγ to induce the differentiation and control the function of both white and brown adipocytes results from a fine tuning of its transcriptional repertoire by post-translational modification and recruitment of coregulators (Ahmadian et al. 2013; Ma et al. 2018).

PPARγ can be regulated by post-translational modifications such as phosphorylation, sumoylation and acetylation (Ahmadian et al. 2013). Phosphorylation of PPARγ on ser112 in the N-terminal A/B domain inhibits ligand binding, but reduced phosphorylation at ser112 is associated with increased adiposity (Rangwala et al. 2003; Shao et al. 1998). PPARγ also can be phosphorylated by cyclin-dependent kinase 5 (CDK5) on ser273 in the ligand-binding domain, which dysregulates the expression of genes involved in insulin-sensitization including adipsin and adiponectin (Choi et al. 2010). PPARγ ligands, such as rosiglitazone (an anti-diabetic therapeutic), that are capable of inhibiting phosphorylation at ser273 improve glycemic control and insulin sensitivity (Choi et al. 2010; Choi et al. 2011). These healthful metabolic effects can be mimicked by a CDK5 inhibitor, roscovitine (Wang et al. 2016). Thus, it has been suggested that PPARγ agonism is not a prerequisite for the beneficial effects of inhibited phosphorylation at ser273 (Choi et al. 2011). Moreover, ser273 of PPARγ is in close proximity with the RXRα DNA binding domain, implicating protein-protein interactions and resulting dynamics as a target for phosphorylation that can ultimately regulate the recruitment of coregulators (Chandra et al. 2008; Lemkul et al. 2015). Indeed, selective PPARγ activation is a strategy being used to design therapeutics that maximize insulin sensitization while minimizing adverse effects (Garcia-Vallve et al. 2015).

Here, we investigate the molecular mechanism of TPhP action on PPARy. Using, *in vivo* and *in vitro* (3T3-L1 cells and primary human pre-adipocytes) models, we demonstrate that unlike classical PPARy agonists like rosiglitazone, TPhP favors white adipogenesis. Proteomics and phosphoproteomics analysis further support those observations. This effect was due to the inability of TPhP to protect PPARy form it phosphorylation on Ser273. Overall, our results demonstrate that TPhP is a selective PPARγ modulator.

## Methods

### Chemicals

DMSO was purchased from American Bioanalytical (Natick, MA). Cell culture chemicals were from Sigma-Aldrich (St. Louis, MO). Rosiglitazone (cat. #71740, purity ≥ 98%) was from Cayman Chemical (Ann Arbor, MI). Roscovitine (#R7772, purity ≥ 98%) and triphenyl phosphate (cat. #241288, purity ≥ 99%) were from Sigma-Aldrich. All other reagents were from Thermo Fisher Scientific (Waltham, MA), unless noted.

### In vivo experiment

Six-week-old, male C57BL/6J mice (DIO, Stock No: 380050 and DIO Control, Stock No: 380056) were obtained from Jackson Laboratory (Bar Harbor, ME) and housed at 23°C in a 12 hr light/dark cycle. Experimental procedures were approved by the Boston University Institutional Animal Care and Use Committee and performed in an American Association for the Accreditation of Laboratory Animal Care accredited facility (Animal Welfare Assurance Number: A3316-01). Water and food were provided *ad libitum*. Mice were fed either a diet with 60% kcal% fat (high fat diet, D12492, Research Diets, New Brunswick, NJ) or 10% kcal% fat (low fat diet, D12450B, Research Diets) for a total of 13 weeks. Seven weeks after initiation of the diet, mice were injected intraperitoneally (ip), daily for 6 weeks with vehicle (composition: 50% saline, 45% PG400, and 5% TWEEN 80; volume was calculated depending on the mouse weight), rosiglitazone (Rosi, 10 mg/kg), or triphenyl phosphate (TPhP, 10 mg/kg). Mice were weighed daily. Body composition was measured by noninvasive quantitative MRI (EchoMRI700) before euthanasia. To isolate mature adipocytes (MA), inguinal white adipose tissue (IWAT) was minced, resuspended in 5 ml 1% collagenase type II in DMEM with 2.5% BSA and incubated, rocking for 40 min at 37°C. Samples were filtered through 100 and 40 μm strainers (BD Bioscience), and centrifuged for 10 min at 500 × rpm (pellet fraction were mature adipocytes). For immunohistochemistry, IWAT was fixed in 4% formalin, paraffin-embedded, sectioned (5 μm) and stained with hematoxylin and eosin (Rabhi et al. 2018). Adipocyte size and number were measured using ImageJ software (Schneider et al. 2012).

### Cell Culture

NIH 3T3-L1 (ATCC: CL-173, RRID:CVCL_0123, Lot # 63343749), 3T3-PPARγ WT and 3T3-PPARγ S273A cells (Qiang et al. 2012) were maintained in high-glucose DMEM with 10% calf serum, 100 U/ml penicillin, 100 μg/ml streptomycin, 0.25 μg/ml amphotericin B. All experiments with 3T3-L1 cells were conducted with passages 3-8. Cells were plated in maintenance medium and incubated for 3 days. “Naïve” cells were cultured in maintenance medium for the duration of an experiment. On day 0, differentiation was induced by replacing the maintenance medium with DMEM containing 10% fetal bovine serum (FBS, Sigma-Aldrich), 250 nM dexamethasone, 167 nM of 1 µg/ml human insulin, 0.5 mM IBMX, 100 U/ml penicillin, and 100 μg/ml streptomycin. On days 3 and 5 of differentiation, medium was replaced with adipocyte maintenance medium (DMEM, 10% FBS, 167 nM human insulin, 100 U/ml penicillin, 100 μg/ml streptomycin), and the cultures were dosed with vehicle (DMSO, 0.1% final concentration), Rosi (20uM), or TPhP (20uM). On Day 7 of differentiation, medium was replaced with adipocyte medium (DMEM, 10% FBS, 100 U/ml penicillin, 100 μg/ml streptomycin), and the cultures were re-dosed. For CDK5 inhibition experiments, cells were treated with Vh or roscovitine (Rosco, 20 µM) at the initiation of differentiation as well as on days 3, 5, and 7 of differentiation. For PPARγ antagonist experiments, cells were treated with Vh or T0070907 (20 µM) at the initiation of differentiation as well as on days 3, 4, 5 and 6 of differentiation. Following 10 days of differentiation and dosing, cells were harvested for cell number analyses (JANUS staining), gene expression, proteome expression, lipid accumulation, mitochondrial biogenesis, and cellular respiration analyses.

Primary, human, subcutaneous pre-adipocytes were obtained from the Boston Nutrition Obesity Research Center (Boston, MA). Pre-adipocytes were maintained in αMEM with 10% FBS, 100 U/ml penicillin, 100 μg/ml streptomycin, 0.25 μg/ml amphotericin B. Pre-adipocytes were plated in maintenance medium and incubated for 3 days. “Naïve” cells were cultured in maintenance medium for the duration of the experiment. On day 0, differentiation was induced by replacing the maintenance medium with DMEM/F12, 25 mM NaHCO3, 100 U/ml penicillin, 100 μg/ml streptomycin, 33 μM d-Biotin, 17 μM pantothenate, 100 nM dexamethasone, 100 nM human insulin, 0.5 mM IBMX, 2 nM T3, and 10 μg/ml transferrin, and the cultures were dosed with vehicle with vehicle (DMSO, 0.1% final concentration), Rosi (4μM), or TPhP (4μM). On day 3 of differentiation, medium was replaced with induction medium, and the cultures were re-dosed. On days 5, 7, 10, and 12 of differentiation, the medium was replaced with adipocyte medium (DMEM/F12, 25 mM NaHCO3, 100 U/ml penicillin, 100 μg/ml streptomycin, 33 μM d-Biotin, 17 μM pantothenate, 10 nM dexamethasone, 10 nM insulin, and 3% bovine calf serum) and the cultures were re-dosed. Following 14 days of differentiation and dosing, cells were harvested for cell number analyses (JANUS staining), gene expression, lipid accumulation, fatty acid uptake, mitochondrial biogenesis, and cellular respiration analyses.

### Reverse Transcriptase (RT)-qPCR

Total RNA was extracted and genomic DNA was removed using the Direct-zol 96-well MagBead RNA Kit (Zymo Research, Orange, CA). cDNA was synthesized from total RNA using the iScript™ Reverse Transcription System (Bio-Rad, Hercules, CA). All qPCR reactions were performed using PowerUp™ SYBR Green Master Mix (Thermo Fisher Scientific, Waltham, MA). The qPCR reactions were performed using a 7500 Fast Real-Time PCR System (Applied Biosystems, Carlsbad, CA): UDG activation (50°C for 2 min), polymerase activation (95°C for 2 min), 40 cycles of denaturation (95°C for 15 sec) and annealing (various temperatures for 15 sec), extension (72°C for 60 sec). The primer sequences and annealing temperatures are provided in Table S1. Relative gene expression was determined using the Pfaffl method to account for differential primer efficiencies (Pfaffl 2001), using the geometric mean of the Cq values for beta-2-microglobulin (*B2m*) and 18s ribosomal RNA (*Rn18s*) for mouse gene normalization and of ribosomal protein L27 (*RPL27*) and *B2M* for human gene normalization. For *in vivo* experiments, the average Cq value from the low fat vehicle group was used as the reference point. For *in vitro* experiments, the Cq value from experiment-specific naïve, undifferentiated cultures was used as the reference point. Data are reported as “Relative Expression.”

### Lipid Accumulation

3T3-L1 cells or human preadipocytes were plated in 24 well plates at 50,000 cells per well in 0.5 ml maintenance medium at initiation of the experiment. Medium was removed from the differentiated cells, and they were rinsed with PBS. The cells were then incubated with Nile Red (1 µg/ml in PBS) for 15 min in the dark. Fluorescence (λex= 485 nm, λem= 530 nm) was measured using a Synergy2 plate reader (BioTek Inc., Winooski, VT). The fluorescence in all control and experimental wells was normalized by subtracting the fluorescence measured in naïve (undifferentiated) cells. The naïve-corrected fluorescence in the experimental wells is reported as “Fluorescence (RFU).”

### Fatty Acid Uptake

3T3-L1 cells or human preadipocytes were plated in 96 well, black-sided plates at 10,000 cells per well in 0.2 ml maintenance medium at initiation of the experiment. Fatty acid uptake was measured by treating differentiated cells with 100 μL of Fatty Acid Dye Loading Solution (Sigma-Aldrich, MAK156). Following a 1 hr incubation, measurement of fluorescence intensity (λex= 485nm, λem= 530nm) was performed using a Synergy2 plate reader. The fluorescence in experimental wells was normalized by subtracting the fluorescence measured in naïve (undifferentiated) cells and reported as “Fluorescence (RFU).”

### Mitochondrial Biogenesis

3T3-L1 cells or human preadipocytes were plated in 24 well plates at 50,000 cells per well in 0.5 ml maintenance medium at initiation of the experiment. Mitochondrial biogenesis was measured in differentiated cells using the MitoBiogenesis In-Cell Elisa Colorimetric Kit, following the manufacturer’s protocol (Abcam). The expression of two mitochondrial proteins (COX1 and SDH) were measured simultaneously and normalized to the total protein content via JANUS staining. Absorbance (OD 600nm for COX1, OD 405nm for SDH, and OD 595nm for JANUS) was measured using a BioTek Synergy2 plate reader. The absorbance ratios of COX/SDH in experimental wells were divided by those in naïve (undifferentiated) cells, and the data are reported as “Relative Mitochondrial Protein Expression.”

### Oxygen Consumption

3T3-L1 cells or human preadipocytes were plated in Agilent Seahorse plates at a density of 50,000 cells per well in 0.5 ml maintenance medium at initiation of the experiment. Prior to all assays, cell media was changed to Seahorse XF Assay Medium without glucose (1mM sodium pyruvate, 1mM GlutaMax, pH 7.4) and incubated at 37°C in a non-CO2 incubator for 30 min. To measure mitochondrial respiration, the Agilent Seahorse XF96 Cell Mito Stress Test Analyzer (available at BUMC Analytical Instrumentation Core) was used, following the manufacturer’s standard protocol. The compounds and their concentrations used to determine oxygen consumption rate (OCR) included 1) 0.5 μM oligomycin, 1.0 μM carbonyl cyanide-p-trifluoromethoxyphenylhydrazone (FCCP) and 2 μM rotenone for 3T3-L1s; and 2) 5 μM oligomycin, 2.5 μM FCCP, and 10 μM rotenone for the primary human adipocytes.

### Immunoblot Analyses

3T3-L1 cells were plated in 6-well plates at 200,000 cells per well in 2 ml maintenance medium at initiation of the experiment. Cells were lysed in RIPA Buffer with PMSF (Cell Signaling Technology, Danvers, MA). Proteins (50 μg) were fractionated using SDS-PAGE on 10% Mini-PROTEAN TGX protein gels (Bio-Rad) and were transferred to nitrocellulose membranes (Bio-Rad). Following transfer, the membranes were blocked with 5% bovine serum albumin in phosphate-buffered saline-0.1% Tween 20 and probed with rabbit anti-PPARγ antibody (#2443S, Cell Signaling Technology) and phospho-ser273 PPARγ (bs-4888R, Bioss, Woburn, MA). Immunoreactive bands were detected using HRP-conjugated secondary antibodies (Bio-Rad) followed by enhanced chemiluminescence.

### Proteomic Analyses

3T3-L1 cells were plated in T75 flasks at 1,000,000 cells per well in 10 ml maintenance medium at initiation of the experiment. Cells were re-suspended in 8M Urea/50mM triethylammonium bicarbonate (TEAB), with phosphatase and protease inhibitors (Roche, Basel, CH) then sonicated (3×10 seconds) on ice. Samples were reduced for 30 minutes with 8 mM dithiothreitol and alkylated for 15 minutes with 20mM iodoacetamide at 30°C. The 8M urea solution was diluted to 1M with 50 mM TEAB, and samples were digested overnight at 37°C with 20 μg sequencing grade trypsin (90057, Thermo Fisher Scientific).

Prior to TMT (Tandem Mass Tag) labeling, peptides were extracted from each sample using c18 spin columns (Toptip, Glygen, Columbia, MD), and peptide concentrations were normalized to 100ug in 100ul of 100mM TEAB. Peptides were labelled with 0.8 mg of TMT label (TMT10plex™ Isobaric Label Reagent Set, Thermo Fisher Scientific). Labelled samples were pooled, and 95% was set aside for phospho-peptide enrichment using TiO2 (Titansphere Phos-TiO Bulk 10 um, GL Sciences, Tokyo, JP) (Cantin et al. 2007). The remaining 5% of labelled peptides and the phospho-peptide enriched samples were analyzed separately by mass spectrometry.

Samples were analyzed by a Q Exactive HFX mass spectrometer connected to Easy nLC 1200 reverse-phase chromatography system (Thermo Fisher Scientific). Mobile phase A was 0.1% formic acid and 2% acetonitrile, mobile phase B was 0.1% formic acid and 80% acetonitrile. Peptides were resuspended in 0.1% formic acid for loading. The samples were loaded onto a nano-trap column with mobile phase A, (75μm i.d. × 2 cm, Acclaim PepMap100 C18 3μm, 100Å, Thermo Fisher Scientific) and were separated over an EASY-Spray column, (50 cm × 75 μm ID, PepMap RSLC C18, Thermo Fisher Scientific) by an increasing mobile phase B gradient over 180 minutes at a flow rate of 250 nL/min. The mass spectrometer was operated in positive ion mode with a capillary temperature of 300°C, and with a potential of 2100V applied to the frit. All data was acquired with the mass spectrometer operating in automatic data dependent switching mode. A high resolution (60,000) MS scan (350-1500 m/z) was performed using the Q Exactive to select the 10 most intense ions prior to MS/MS analysis using HCD (NCE 33, 45,000 resolution).

Resulting RAW files were searched using MaxQuant (version 1.6.0; www.coxdocs.org/doku.php?id=maxquant:start) under standard settings using the UniProt mouse database (downloaded October 2018, www.uniprot.org/taxonomy/10090) allowing for two missed trypsin cleavage sites and variable modifications of N-terminal acetylation and methionine oxidation (Tyanova et al. 2016). Additionally, protein phosphorylation at S, T, and Y residues were included as variable modifications in the phosphoproteomics data. Carbamidomethylation of cysteine residues was a fixed modification in the search. Candidate (phospho)peptides, proteins, and phosphorylation-site identifications were filtered based on a 1% false discovery rate threshold based on searching the reverse sequence database. Data are deposited and publicly available at the PRIDE archive. All phosphopeptide identifications in the MaxQuant evidence file had to meet a 0.7 probability cutoff.

The searched intensity data were filtered, normalized, and clustered using R [https://www.R-project.org/]. Feature filtering was performed to remove any feature in less than 70 percent of samples with 1627 proteins and 1066 phospho-sites passing the filter in the proteomics and phosphoproteomic data sets, respectively. The LIMMA R package was used for LOESS normalization and differential expression analysis (Ritchie et al. 2015). A combined ranked list for both sets was generated where duplicate gene entries were removed to keep the entry with the highest absolute rank value.

Gene Set Enrichment Analysis (GSEA) software from the Broad Institute (software. broadinstitute.org/GSEA)(version 3.0) was used in rank mode along with gene sets downloaded from the Bader Lab (Mouse_GOBP_AllPathways_no_GO_iea_October_01_2018_symbol.gmt) from http://baderlab.org/GeneSets) (Merico et al. 2010; Subramanian et al. 2005). GSEA results were visualized using the Enrichment Map app (Version 3.1) in Cytoscape (Version 3.6.1) and highly related pathways were grouped into a theme and labeled by AutoAnnotate (version 1.2). For the merged gene set analyses, we applied an enrichment P < 0.01 and FDR ≤ 0.1 cutoff and calculated overlap between gene set annotations using a combination of Jaccard and overlap coefficients with a cutoff of 0.375.

### Statistical Analyses and Publicly Available Data

All statistical analyses were performed in R (v 3.5.0) and Prism 7 (GraphPad Software, Inc., La Jolla, CA). Data are presented as means ± standard error (SE). For 3T3-L1 experiments, the biological replicate is an independently plated experiment. For human primary preadipocyte experiments, the biological replicate is a single individual’s preadipocytes (3 individuals in all). The qPCR data were log-transformed before statistical analyses. One-factor ANOVAs (Dunnett’s) were performed to analyze the qPCR and phenotypic data. The mass spectrometry proteomics data are available from PRIDE (http://www.proteomexchange.org) under accession number PXD012337.

## Results

### Effect of TPhP on adipocyte browning in vivo

We examined *in vivo* effects of 6-week exposure to Rosi and TPhP (10 mg/kg/day) on C57BL/6J male mice that were also fed either low fat diet (LFD) or high fat diet (HFD) for 13 weeks. Only a slight effect was observed on body weight gain in mice exposed to Rosi or TPhP fed a LFD (**Fig. S1a**). However, mice under HFD gained less weight following Rosi or TPhP exposure when compared to Vehicle (Vh) treated mice. This effect is due to changes in fat mass as no significant differences was observed in lean mass (**Figs. S1b**).

We next examined the inguinal adipose tissue (IWAT) morphology. H&E staining on both LFD and HFD fed mice exposed to Rosi or TPhP showed smaller adipocytes (**Fig. 1a**). Quantification of adipocyte number showed an increase in cell number under both LFD and HFD following Rosi or TPhP exposure (**Fig. 1b**). In contrast, we observed a decreased adipocyte size in IWAT form mice treated with Rosi or TPhP fed either diet (**Fig. 1b**). Because Rosi action through PPARy has been shown to induce a brown adipose-like phenotype in the inguinal depot, we measured mRNA expression of white and brite adipocyte marker genes (Qiang et al. 2012) in mature inguinal adipocytes. Rosi and TPhP did not change the expression of *Fabp4* but significantly induced expression of *Plin1* and *Cidec* in both diet groups (**Fig. 1c**). TPhP increased *Retn* expression in both diet groups, while Rosi increased its expression only in the HFD group (**Fig. 1c**). Rosi only significantly reduced *Wdnm1* expression in the LFD group (**Fig. 1c**). In the LFD fed mice, Rosi only significantly induced expression of known markers of brite adipocytes; while TPhP reduced expression of *Elovl3*, *Ucp1* and *Acaa2* (**Fig. 1d**). In the HFD fed mice, Rosi significantly induced expression of *Elovl3, Cidea, Acaa2,* and *CoxIV*; while TPhP significantly increased only *Acaa2* expression (**Fig. 1c**). Moreover, in the HFD group, TPhP significantly reduced expression of *Elovl3* and *Ucp1* (**Fig. 1d**). Hence, whereas both Rosi and TPhP are PPARγ agonists, only Rosi was able to induce thermogenic gene expression *in vivo,* suggesting that Rosi and TPhP may activate PPARγ in a different manner.

**Fig. 1.**
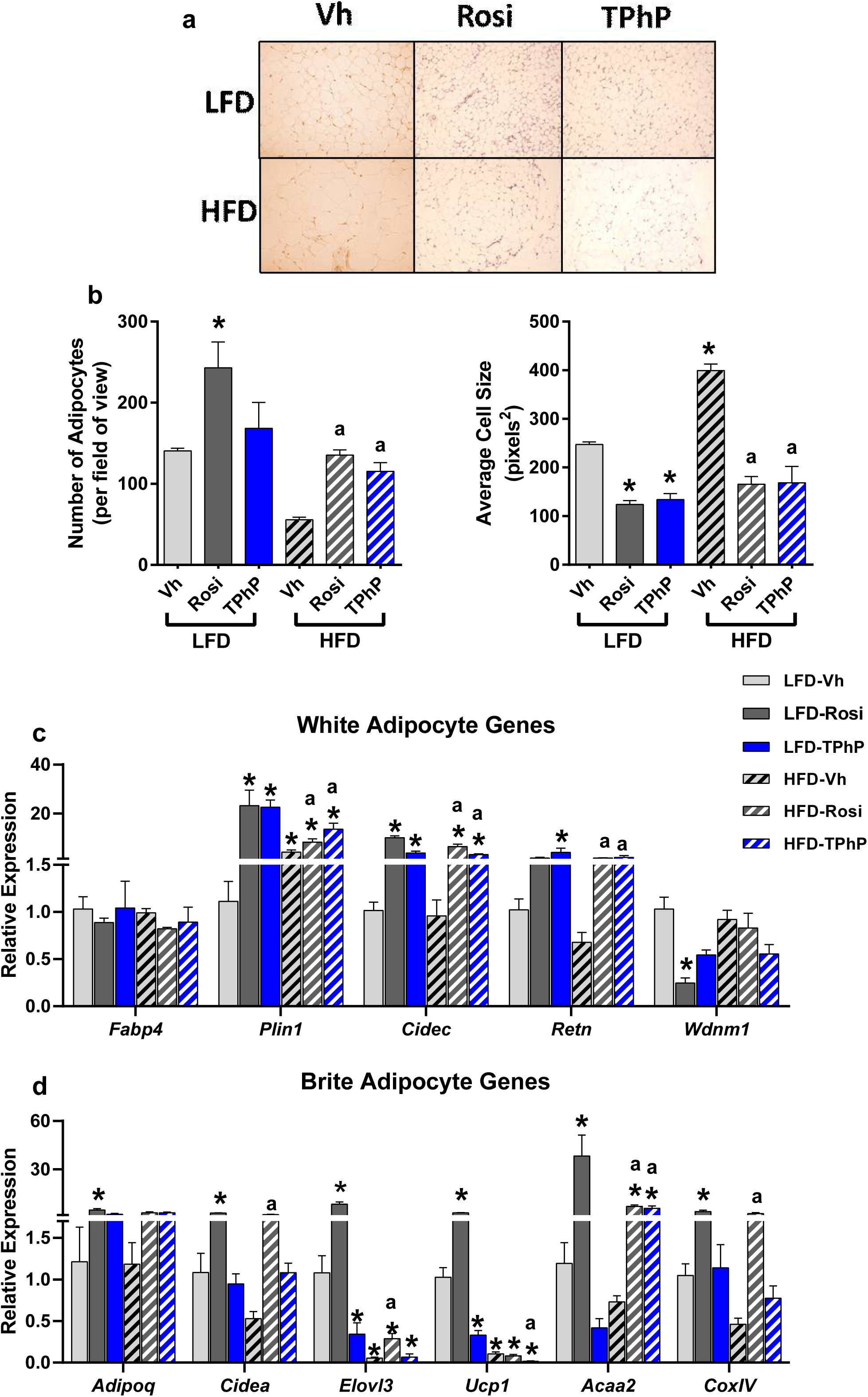
IWAT histology and gene expression in Rosi- and TPhP-treated mice. Six-week-old C57BL/6J male mice were fed either a diet with 10% kcal% fat (LFD) or 60% kcal% fat (HFD) for a total of 13 weeks. Seven weeks after initiation of the diet, mice were ip injected, daily for 6 weeks with vehicle, rosiglitazone (10 mg/kg), or triphenyl phosphate (10 mg/kg). Inguinal white adipose tissue (IWAT) was stained with hematoxylin and eosin. **a** Representative micrograph. **b** The number and size of adipocytes were measured using ImageJ. Mature adipocytes were isolated from IWAT by digestion, filtering and centrifugation. Adipocytes were analyzed for gene expression by RT-qPCR. **c** White adipocyte marker genes. **d** Brite adipocyte genes. Data are presented as mean ± SE (n=5). Statistically different from LFD Vh-treated (*p<0.05) or HFD Vh-treated (^a^p<0.05, ANOVA, Dunnett’s).

### Effect of TPhP on adipocyte browning in vitro

In order to examine if this effect is due to a direct action on adipocytes, we characterized the adipocytes induced by the two PPARγ ligands, Rosi and TPhP, in 3T3-L1 cells, a mouse-derived pre-adipocyte model. 3T3-L1 cells were differentiated using a standard hormone cocktail for 10 days. During differentiation, cells were treated with Vh (DMSO), Rosi (20 μM), or TPhP (20 μM). Rosi, but not TPhP, significantly increased cell number (**Fig. S2a**). Rosi and TPhP both significantly induced lipid accumulation (**Fig. 2a**). Rosi and TPhP induced *Plin1* mRNA expression in a PPARγ-dependent manner (**Fig. 2b**). Rosi, but not TPhP, induced expression of the brite adipocyte genes *Elovl3* and *Ucp1*, and this induction also was PPARγ-dependent (**Fig. 2b**). Only Rosi significantly induced mitochondrial biogenesis and cellular respiration (**Figs. 2c-d**).

**Fig. 2.**
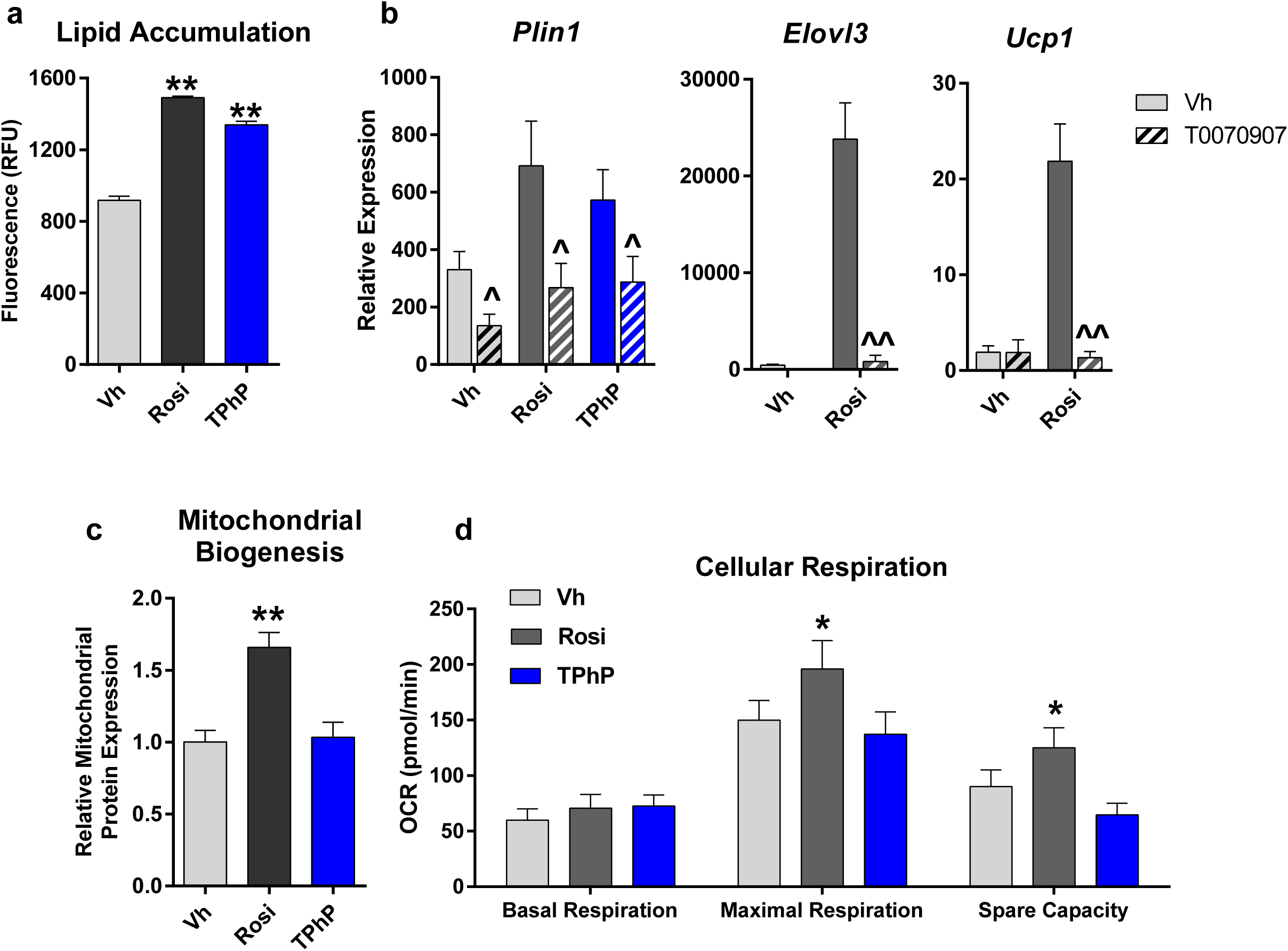
Functions of and gene expression in 3T3-L1 adipocytes differentiated with Rosi and TPhP. Confluent 3T3-L1 cells were differentiated using a standard hormone cocktail for 10 days. During differentiation, cells were treated with Vh (0.1% DMSO, final concentration), Rosi (20 μM), or TPhP (20 μM) in the presence or absence of the PPARγ antagonist T0070907 (20 µM). **a** Lipid accumulation was determined by Nile Red staining. **b** Gene expression was analyzed by RT-qPCR. **c** Mitochondrial biogenesis was analyzed by measuring mitochondria-specific proteins. **d** Cellular respiration was measured using the Seahorse assay. Data are presented as mean ± SE (n=3-6). Statistically different from Vh-treated (*p<0.05 or **p<0.01, ANOVA, Dunnett’s).

Thus, we performed proteomic analyses with differentiated and treated 3T3-L1s. The heatmap in **Fig. 3a** shows that Rosi-treated cells expresses a suite of protein that are distinct from Vh- and TPhP-treated cells. TPhP-treated cells cluster and have expression patterns more similar to Vh-treated cells (**Fig. 3a**). The list of protein intensities and results of the differential analysis are presented in Table S2. Rosi highly upregulated CD36, FABP4, and glycerol-3phosphate dehydrogenase 1 (GPD1)(**Fig. 3b**), which are involved in fatty acid metabolism (Coburn et al. 2000; Hertzel et al. 2006; Moustaid et al. 1996). TPhP highly upregulated calponin 2 (CNN2) and vasodilator stimulated phosphoprotein (VASP), which are actin-associated proteins (Galler et al. 2006; Wu and Jin 2008), SEC13, which is involved in vesicle budding from the endoplasmic reticulum (Salama et al. 1993), and epoxide hydrolase 1 (EPHX1), which is an enzyme that biotransforms epoxides to diols (Vaclavikova et al. 2015) (**Fig. 3b**).

**Fig. 3.**
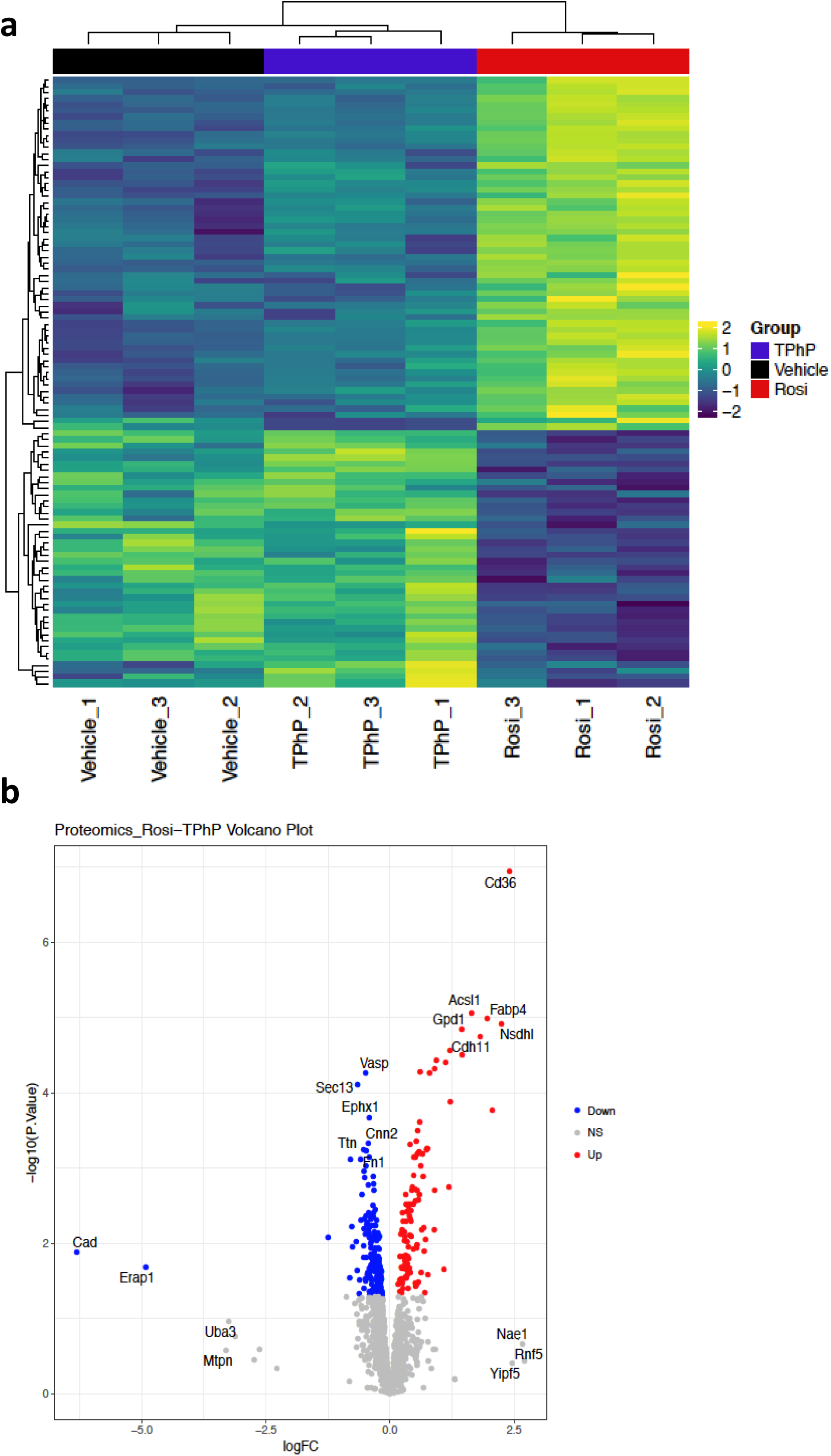
Proteomes of 3T3-L1 adipocytes differentiated with Rosi and TPhP. Confluent 3T3-L1 cells were differentiated as described in Fig. 3. The proteome was analyzed by precision quantitative nanoLC-tandem MS. **a** Heatmap and **(b)** volcano plot of top differentially expressed proteins between Rosi and TPhP.

Protein phosphorylation is one of the most common post-translational modifications through which protein function is regulated; therefore we also investigated the effects of Rosi and TPhP on the phospho-proteome. Rosi differentially regulated phosphorylation sites compared to Vh and TPhP (**Fig. 4a**). All phosphorylation sites and results of the differential analysis are provided in Table S3. Rosi highly upregulated the phosphoproteins, SDPR, which is involved in caveolae formation and function (Nassar and Parat 2015), acyl-CoA-binding domain-containing protein 5 (ACBD5), which participates in peroxisomal very-long chain fatty acid metabolism (Yagita et al. 2017), and glycogen synthase 1 (GYS1)(Pederson et al. 2005) (**Fig. 4b**). TPhP favored the expression of the phosphoproteins STEAP3 and erythrocyte membrane protein band 4.1 Like 3 (EPB41L3)(**Fig. 4b**), which are involved in secretion (Amzallag et al. 2004) and cell adhesion/spreading, respectively (Wang et al. 2014).

**Fig. 4.**
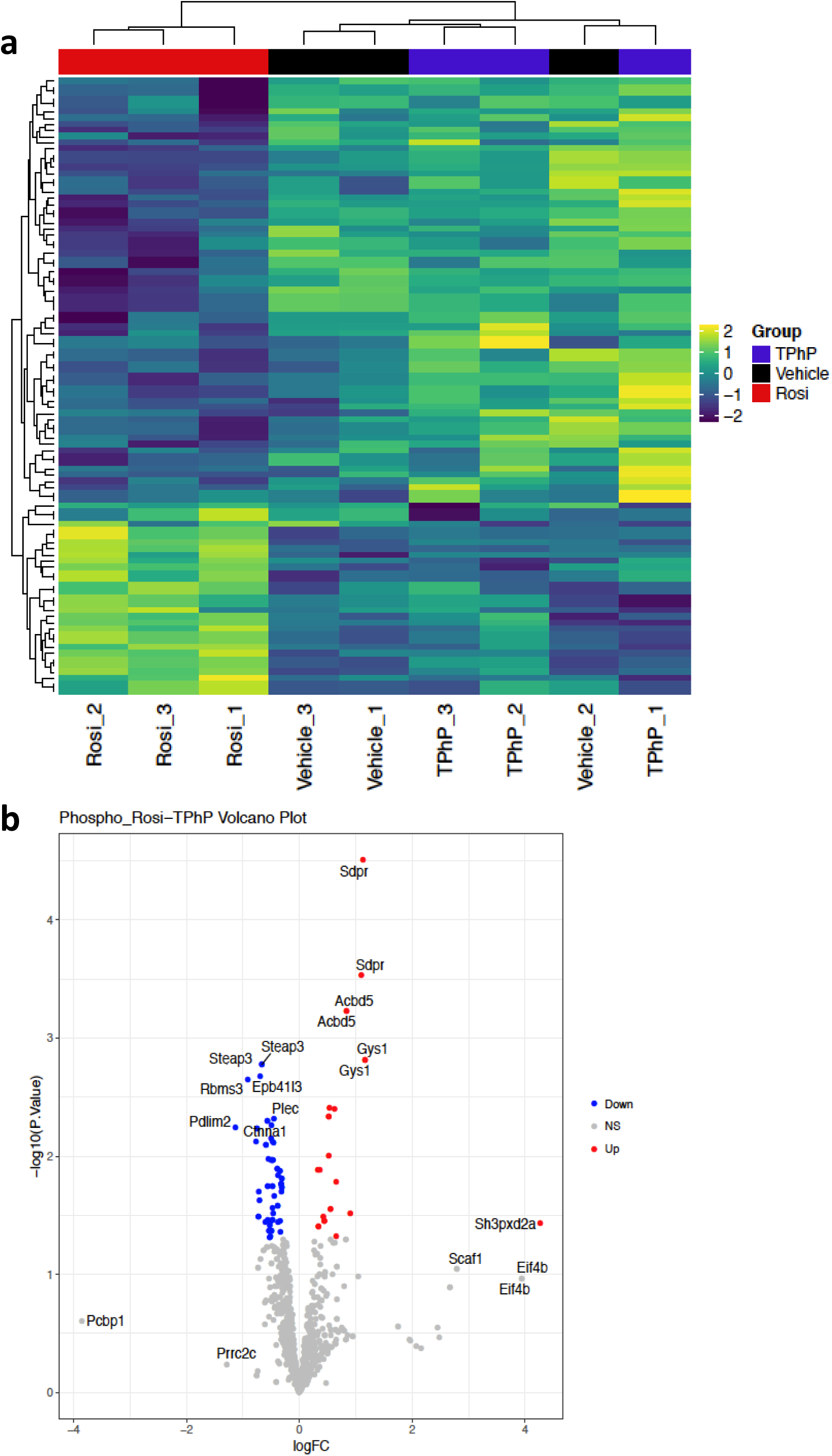
Phosphoproteomes of 3T3-L1 adipocytes differentiated with Rosi and TPhP. Confluent 3T3-L1 cells were differentiated as described in Fig. 3. Phospho-peptides were enriched using TiO2 and then analyzed by precision quantitative nanoLC-tandem MS. **a** Heatmap and **(b)** volcano plot of top differentially expressed phospho-proteins between Rosi and TPhP.

We created a combined differential protein and phosphoprotein ranked list to perform enrichment analysis in order to determine which pathways are enriched or shared by the two PPARγ ligands (**Fig. 5**). The complete enrichment results are provided in Table S4. **Fig. 5** shows the network representation of pathways, where nodes are pathways and edges are shared genes between pathways. Pathways (nodes) significantly enriched by Rosi (red colored nodes), significantly enriched by TPhP (purple colored nodes), or shared by both (nodes split with red and purple). In order to interrogate the role of PPARy in the pathway networks, nodes were highlighted green to indicate PPARγ is in the gene set, showing many pathway clusters relate to PPARy activity. Rosi regulated networks related to oxidation/metabolic processes, mitochondrial proteins, and tissue morphogenesis. TPhP regulated networks related to ubiquitination of proteasomes. Shared networks were related to oxidation/metabolic processes as well as regulation of lipid ketones. That Rosi regulated protein networks related to oxidation/metabolic and mitochondrial processes is consistent with the phenotypic differences in the adipocytes that Rosi and TPhP induced.

**Fig. 5.**
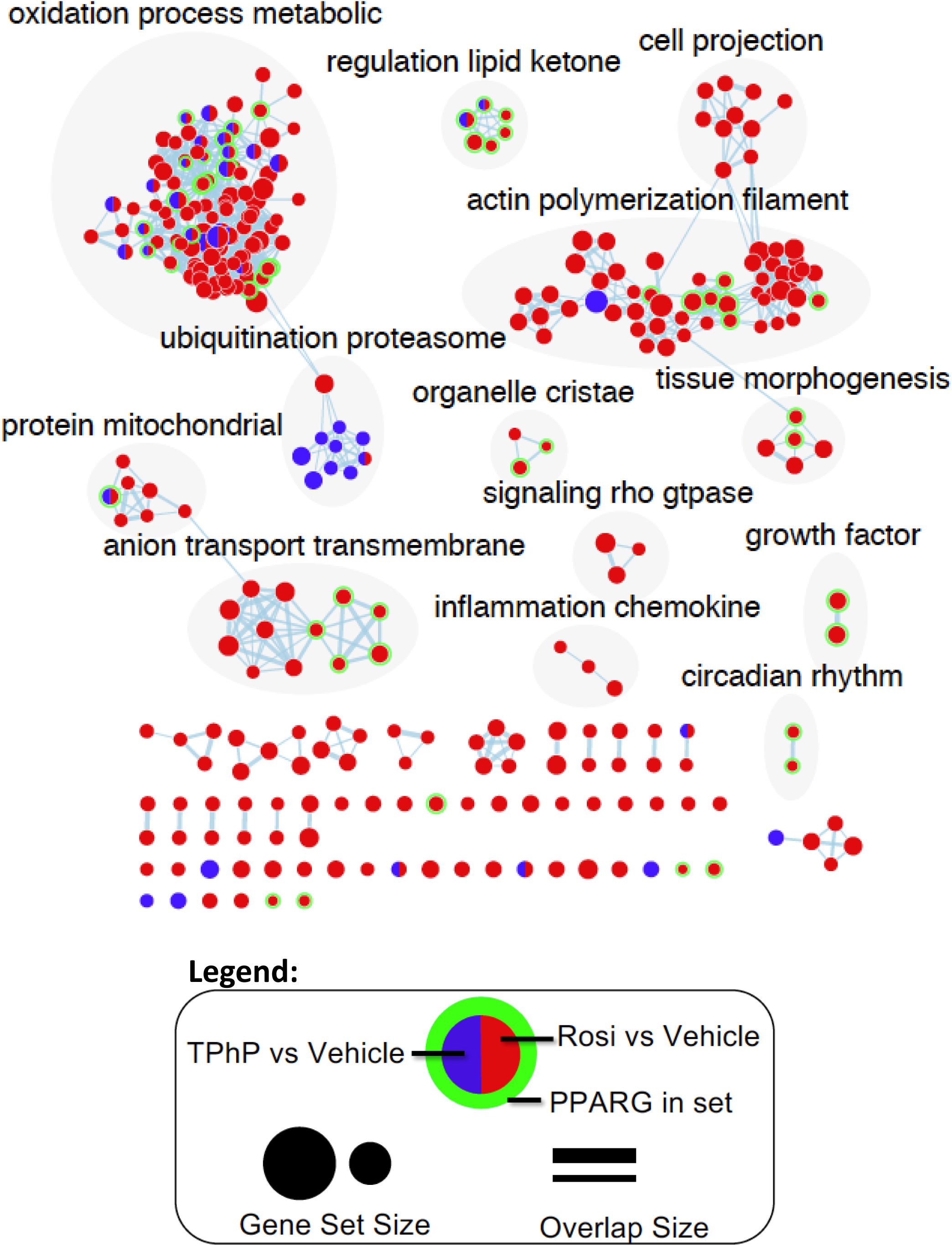
Network analyses of Rosi and TPhP-induced proteomes. Pathway themes are displayed above highlighted clusters. Nodes in red indicates enriched pathways in Rosi, blue indicates enriched pathways by TPhP, red/blue indicates shared pathways between Rosi and TPhP. Nodes with green surrounding indicate the presence of PPARγ in the pathway set.

### Effect of TPhP on PPARγ phosphorylation and its contribution to adipocyte differentiation

PPARγ is itself a phosphoprotein, and the phosphorylation status of S273 influences the target genes that are expressed (Choi et al. 2010; Choi et al. 2011). Therefore, we used immunoblotting and a mutant form of PPARγ to investigate how phosphorylation of PPARγ at ser273 differentiates the effects of Rosi and TPhP. 3T3-L1 cells were differentiated and treated as described above. Rosi significantly reduced phosphorylation of PPARγ at ser273 (**Fig. 6a**), which also has been shown by others (Banks et al. 2015; Choi et al. 2010). TPhP did not protect PPARγ from phosphorylation (**Fig. 6a**).

**Fig. 6.**
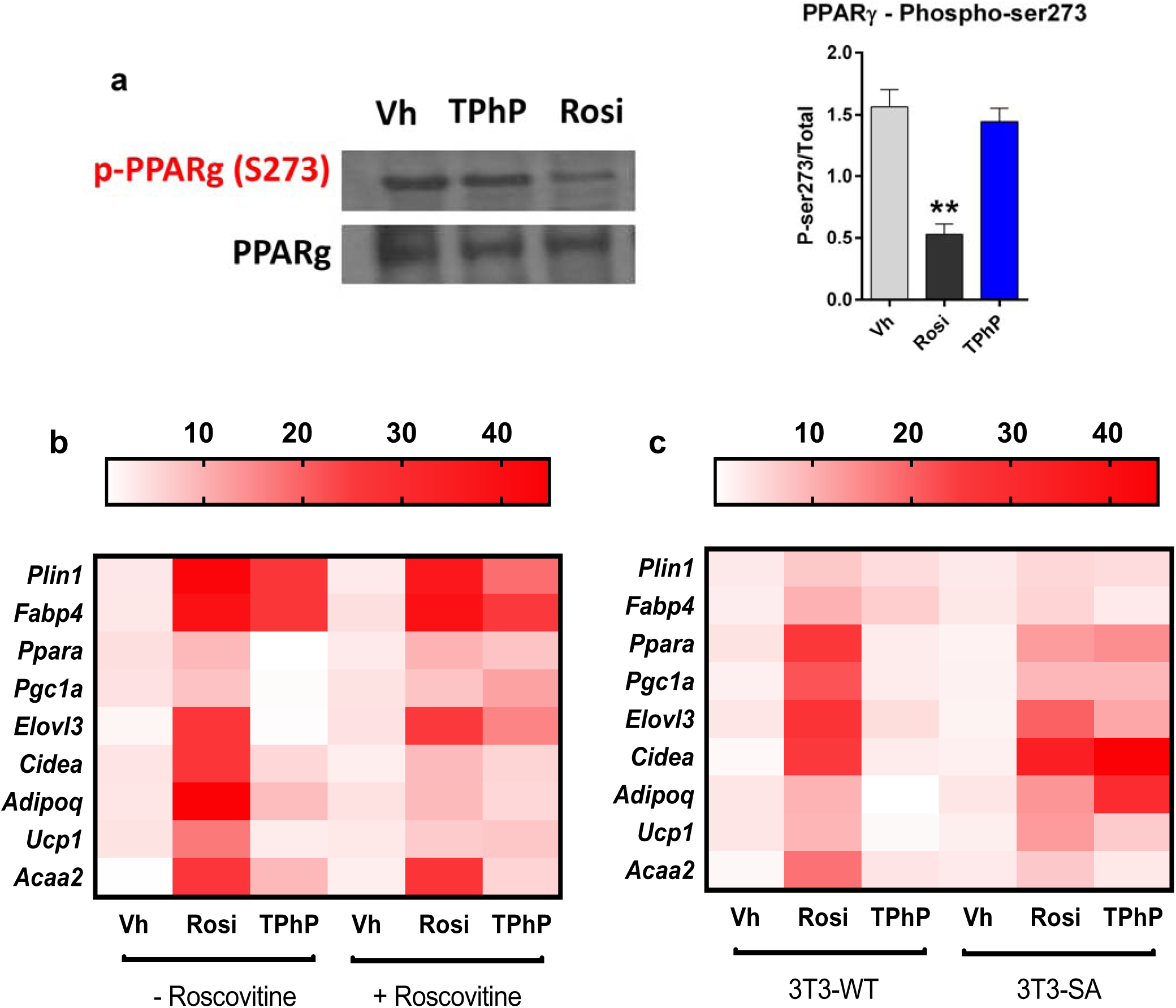
PPARγ phosphorylation and its effect on gene expression in mouse adipocytes differentiated with Rosi and TPhP. **a** Confluent 3T3-L1 cells were differentiated as described in Fig. 3 and phosphorylation of PPARγ at ser273 was determined relative to total PPARγ by immunoblot. **b** Confluent 3T3-L1 cells were differentiated as described in Fig. 3, in the presence or absence of roscovitine (Rosco, 20 μM). Gene expression was determined by RT-qPCR and presented as a heatmap of expression levels of white adipocyte and brite adipocyte marker genes. **c** Confluent 3T3 cells expressing wildtype PPARγ (3T3-WT) or PPARγ with alanine substituted for serine 273 (3T3-SA) were differentiated as described for 3T3-L1 cells in Fig. 3. Gene expression was determined by RT-qPCR and presented as a heatmap of expression levels of white and brite adipocyte marker genes. Data are presented as mean ± SE (n≥3). Statistically different from Vh-treated (*p<0.05 or **p<0.01, ANOVA, Dunnett’s).

CDK5 is known to phosphorylate PPARγ at ser273; therefore we also tested whether inhibition of CDK5 by roscovitine (Rosco) would allow TPhP to induce brite adipocyte gene expression. Treatment of 3T3-L1 cells with Rosco (20 μM), along with Rosi and TPhP, during differentiation modestly decreased lipid accumulation (**Fig. S3a**) and did not change white adipocyte marker gene expression (**Fig. 6b, Fig. S3b**). However, inhibition of Cdk5 by Rosco significantly increased the ability of TPhP to induce brite adipocyte gene expression (**Fig. 6b, Fig. S3c**).

Using Swiss 3T3 cells expressing wild-type PPARγ (3T3-WT) or PPARγ with alanine substitute for serine at position273 (3T3-SA), we examined whether a constitutively dephosphorylated PPARγ would enable TPhP to induce brite adipocyte gene expression. Lipid accumulation was less in 3T3-SA cells than in 3T3-WT cells treated with either Rosi or TPhP (**Fig. S4a**) and expression of white adipocyte genes was modestly but not significantly reduced (**Fig. 6c, Fig. S4b**). However, in 3T3-SA cells, TPhP had greater efficacy in inducing brite adipocyte genes, *Adipoq*, *Ucp1* and *Cidea*, in particular (**Fig. 6c, Fig. S4c**). In summary, when liganded with TPhP, PPARγ remain phosphorylated at ser273, which limits PPARγ’s ability to induce brite adipogenesis.

### Effect of TPhP on differentiation of human adipocytes

Finally, we determined if Rosi and TPhP induced differentiation of distinct adipocyte phenotypes in a human model. Human primary pre-adipocytes were differentiated with a standard human hormone cocktail for 14 days. During differentiation, cells were treated with Vh (DMSO), Rosi (4 μM), or TPhP (4 μM). Both Rosi and TPhP significantly increased cell number (**Fig. S2b**) and lipid accumulation (**Fig. 7a**). However, only Rosi significantly induced fatty acid uptake (**Fig. 7b**). Moreover, PPARy activation by Rosi lead to an increased mitochondrial biogenesis and increased spare capacity, although maximal respiration was not change. TPhP had no effect on mitochondrial biogenesis or cellular respiration didn’t affect the mitochondrial activity of the cells (**Figs. 7b-d**). At the gene expression level, both Rosi and TPhP significantly increased mRNA expression of white adipocyte marker genes including *PPARG*, *PLIN1, CIDEC,* and *FABP4* (**Fig. 7e**). While both Rosi and TPhP significantly induced the energy expenditure gene, *ACAA2*, only Rosi significantly increased expression of browning markers, *ELOVL3*, *CIDEA* and *UCP1* (**Fig. 7f**). These observations suggest TPhP selectively activates PPARγ’s transcriptional activities in human, as well as mouse, adipocytes.

**Fig. 7.**
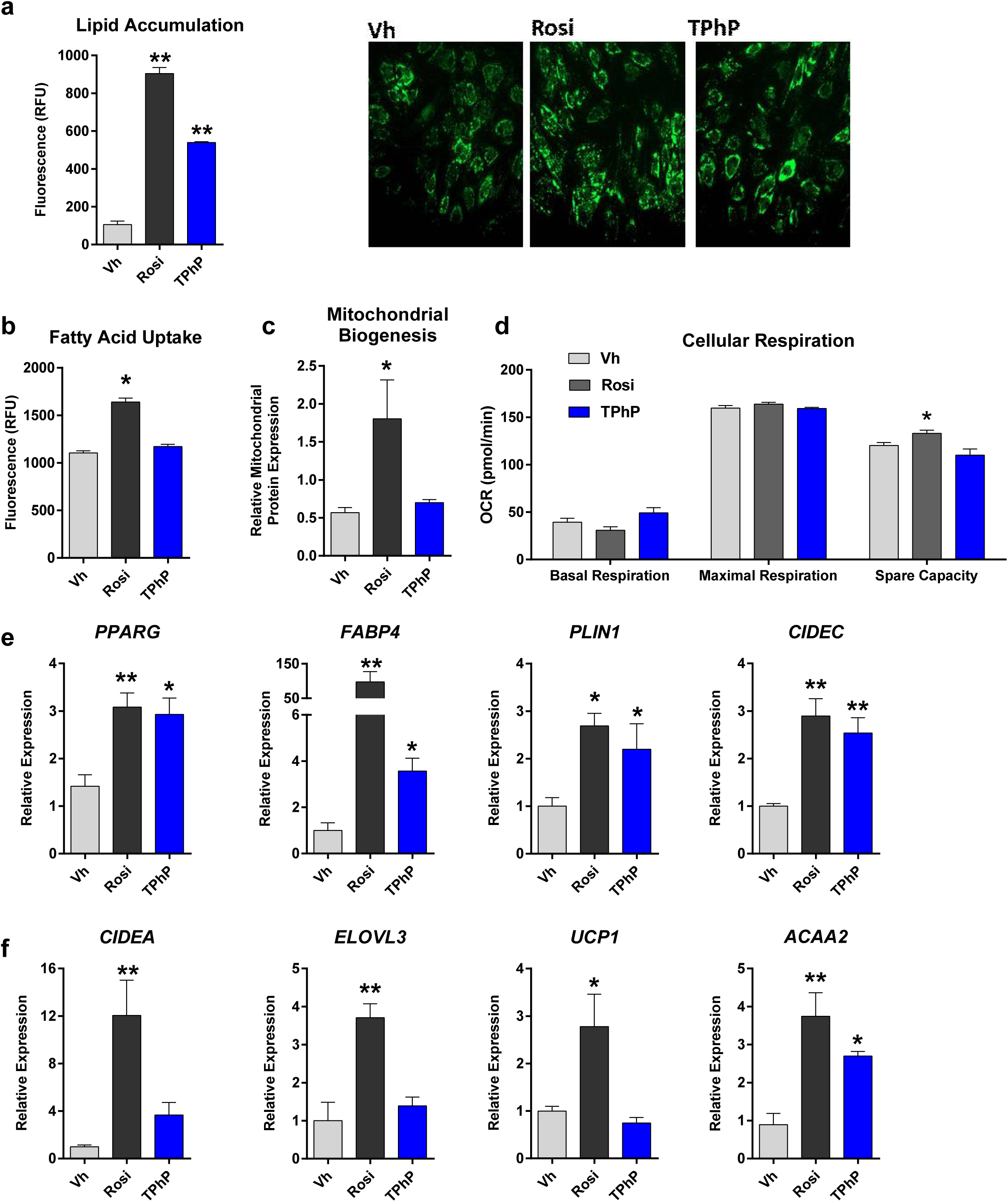
Functions of and gene expression in human adipocytes differentiated with Rosi and TPhP. Confluent primary human adipocytes were differentiated using a standard hormone cocktail for a total of 14 days. During differentiation, cells were treated with vehicle (0.1% DMSO, final concentration), Rosi (4 μM), or TPhP (4 μM). **a** Lipid accumulation was determined by Nile Red staining. **b** Fatty acid uptake was analyzed using a dodecanoic acid fluorescent fatty acid substrate. **c** Mitochondrial biogenesis was analyzed by measuring mitochondria-specific proteins. **d** Cellular respiration was measured using the Seahorse assay. Gene expression was analyzed by RT-qPCR. **e** White adipocyte marker genes. **f** Brite adipocyte marker genes. Data are presented as mean ± SE of adipocytes differentiated from 3 individuals. Statistically different from Vh-treated (*p<0.05 or **p<0.01, ANOVA, Dunnett’s).

## Discussion

The environmental metabolism disrupting chemical, TPhP, is a PPARγ ligand that can induce adipocyte differentiation (Cano-Sancho et al. 2017; Pillai et al. 2014; Tung et al. 2017a). Previous studies have shown using the well-studied and characterized preadipocyte model, 3T3-L1cells, that TPhP increases lipid accumulation and mRNA levels of adipocyte differentiation markers such as *Plin1* (Cano-Sancho et al. 2017; Tung et al. 2017a). However, no study has yet investigated the phenotype of the adipocyte that forms. Brite adipogenesis can impact the ratio of lipid-storing (typical of white adipocytes) to lipid-burning adipocytes (Rosenwald and Wolfrum 2014). Thus, skewing of adipogenesis toward white and away from brite adipogenesis could have consequences on healthful adipocyte function. Indeed, rodents that lack brite adipocytes develop obesity, with insulin resistance and hepatic steatosis (Cohen et al. 2014). Here, we examined the characteristics of TPhP-induced adipocytes in multiple models and examined the mechanism that differentiates TPhP-liganded-PPARγ from Rosi-liganded-PPARγ.

Inguinal adipose tissue is the largest adipose depot that can recruit brite adipocytes upon chronic cold exposure of mice, and mature adipocytes in this depot have the potential to transdifferentiate from cells with the typical characteristics of white adipocytes to characteristics of brown adipocytes (Rosenwald and Wolfrum 2014; Walden et al. 2012). The reduction of adipocyte size and reduction of adipocyte number in IWAT of TPhP-exposed mice suggests that TPhP induced adipocyte hyperplasia, rather than hypertrophy. However, TPhP failed to induce expression of *Cidea* and significantly reduced the expression of *Elovl3* and *Ucp1* in the mature adipocytes isolated from the IWAT. Thus, it appears that TPhP induces the differentiation of new, white adipocytes in IWAT. Lineage-tracing studies are needed to confirm this hypothesis.

We previously reported that the environmental PPARγ/RXR ligand tributyltin induces distinct transcriptional response and adipocyte phenotype distinct from that induced by rosiglitazone (Kim et al. 2018). Another environmental PPARγ ligand, mono-2-ethylhexyl phthalate (MEHP), modifies PPARγ coregulator recruitment distinctly from rosiglitazone, as well (Feige et al. 2007). MEHP binds to PPARγ in a configuration similar to rosiglitazone; however, MEHP is unable to stabilize helix 12 of PPARγ thus limiting its ability to release the corepressor NCOR. MEHP stimulates the recruitment of the coactivators MED1 and PGC1α but not of p300 and SRC1 (Feige et al. 2007). We have shown previously that TPhP interacts with the PPARγ ligand binding domain more like the partial ligand nTZDpa than rosiglitazone (Pillai et al. 2014). Here, we show that TPhP also acts as a selective PPARγ ligand.

Phosphorylation of PPARγ regulates the suite of genes it can transactivate. In obese mice, which have been fed a high-fat diet, Cdk5 becomes activated in adipose tissues and phosphorylates PPARγ at ser273 (Choi et al. 2010). This modification of PPARγ reduces expression of the insulin-sensitizing adipokine, adiponectin, and studies have suggested that Cdk5-mediated phosphorylation of PPARγ may be involved in the pathogenesis of insulin-resistance (Banks et al. 2015; Choi et al. 2010; Choi et al. 2011). The phosphorylation of PPARγ by Cdk5 is blocked by anti-diabetic PPARγ ligands, such as rosiglitazone, and by PPARγ-modifying compounds like roscovitine (Banks et al. 2015; Wang et al. 2016). The inhibition of phosphorylation of PPARγ at ser273 improves insulin sensitivity and recently has been linked to browning of adipose (Choi et al. 2010; Choi et al. 2011; Wang et al. 2016). In our *in vitro* studies with 3T3-L1 cells, we observed that in the presence of TPhP, PPARγ remains phosphorylated at ser273. Further, when PPARγ cannot be phosphorylated at ser273 (either by inhibition of CDK5 or mutation of ser273), TPhP was able to induce brite genes such as *Pgc1a*, *Elovl3*, and *Ucp1*. Interestingly, TPhP gained the ability to induce *Adipoq* expression in 3T3-SA cells; adiponectin is well known to increase insulin sensitivity (Maeda et al. 2002).

The likely link between the ligand-determined phosphorylation state of PPARγ and the resulting transcriptional repertoire is differential co-regulator recruitment. PPARγ ser273 phosphorylation is regulated by a complex of interacting cofactors and can have downstream effects on the diabetic gene program (Ma et al. 2018). For example, a PPARγ corepressor, NCoR (nuclear receptor corepressor 1), enhances Cdk5 activity on phosphorylating PPARγ ser273, and an *in vivo* study has found that compared to wild type mice fed a HFD, fat-specific NCoR-deficient mice on HFD were prone to obesity yet have enhanced insulin sensitivity (Li et al. 2011). Additionally, thyroid hormone receptor-associated protein 3 (THRAP3) can preferentially interact with PPARγ when ser273 is phosphorylated, and knockdown of *Thrap3* in mature adipocytes restored expression of several genes (i.e. *Adipoq*) dysregulated by Cdk5-mediated PPARγ phosphorylation, without altering adipogenesis (Choi et al. 2014). Hence, in future studies, we will validate the role of differential recruitment of coregulators upon ligand-activated PPARγ phosphorylation.

Differences in mRNA expression are associated with differences in the proteomes and functional characteristics of the 3T3-L1 adipocytes induced by rosiglitazone and TPhP. Rosiglitazone induced the differentiation of significantly more energetic adipocytes than TPhP. This most likely is a result of enhanced mitochondrial biogenesis. Moreover, our combined proteomic and phosphoproteomic analyses revealed that, compared to TPhP, Rosi enriched more regulatory networks related to oxidation/metabolic processes and mitochondrial proteins. Our data set also uncovered an array of phosphorylation changes on several key enzymes, such as SDPR and GYS1, involved in lipid and glucose homeostasis; and these key enzymes were only upregulated by the therapeutic PPARγ ligand, Rosi. Interestingly, TPhP appeared to favor expression of genes that interact with the cytoskeleton. Cytoskeletal rearrangement is a necessary step for fibroblasts to transform from a spread to round morphology, the first step in adipocyte differentiation (Mor-Yossef Moldovan et al. 2019). In addition, STEAP3 was identified in a recent proteomic analysis as being “adipocyte-associated” (Ye et al. 2011).

Like in mice *in vivo* and in 3T3-L1 cells, TPhP failed to induce brite adipocyte differentiation in primary human preadipocytes. In accordance with a previous study (Tung et al. 2017b), TPhP induced lipid accumulation and expression of white adipocyte genes (i.e. *Plin1* and *Cidec*). However, TPhP induced neither brite adipocyte genes such as *Elovl3* and *Ucp1* nor mitochondrial biogenesis or activity in the differentiated primary human adipocytes. Interesting, humans with a lower propensity to develop brite adipocytes are more likely to be obese/diabetic (Claussnitzer et al. 2015; Timmons and Pedersen 2009).

If TPhP, and structurally similar organophosphate esters, body burden in humans reaches a level to activate PPARγ is an important question. The dose we chose for *in vivo* TPhP exposure (10 mg/kg/day) was to match the dose for Rosi that was previously used to evaluate browning potential of the adipose tissue (Wang et al. 2016). Our *in vivo* dose is higher than the human average daily exposure (up to 5 µg/kg day). Exposure to TPhP and related organophosphate ester intake is estimated between 50-500 ng/kg/day for dietary intake (Zhao et al. 2019) and 2-8 ng/kg/day for dust intake (Zhao et al. 2019). Drinking water concentrations have been found to be as high as 700 ng/L in the United States; assuming that an average 70 kg person consumes the recommended 3 L of water per day, drinking water intake may add another 30 ng/kg/day exposure (Li et al. 2014). Last, dermal exposure through the use of nail polish may result in exposures as high as 5000 ng/kg./day (Tokumura et al. 2019). While our implemented dose is higher than the typical human exposure, it is lower than that in studies used to derive a non-regulatory reference dose for TPhP (70 mg/kg/day) (Li et al. 2018) as well as the recently set benchmark dose of 19 mg/kg/day (NTP 2018).

## Conclusions

Exposure to the metabolism disrupting chemical, TPhP, skews adipogenesis toward white adipocytes and away from brite adipocytes. TPhP fails to upregulate expression of genes that contribute to mitochondrial biogenesis and energy expenditure. This is in contrast to the therapeutic PPARγ ligand, rosiglitazone. The mechanism contributing to this difference is likely related to the fact that while rosiglitazone protects PPARγ from phosphorylation at ser273, TPhP fails to do so. Indeed, when PPARγ cannot be phosphorylated, TPhP gains the ability upregulate expression of brite adipocyte genes. Thus, we propose the novel conclusion that TPhP is a selective PPARγ modulator and the basis of that selective modulation is the failure to protect PPARγ from phosphorylation. We hypothesize that this mechanism of selective modulation may explain why a number of other environmental PPARγ ligands also fail to induce brite adipogenesis. Critical questions remain to be answered, including how exposure to TPhP at environmentally relevant doses in a human-like dietary context influences metabolic homeostasis and how changes in coregulatory recruitment link environmental ligand-induced differences in PPARγ coregulatory recruitment with specific gene repertoires.

## Funding

This work was supported by the National Institute of Environmental Health Sciences Superfund Research Program P42 ES007381 to JJS, the National Institute of Diabetes and Digestive and Kidney Diseases R01DK117161 to SF and the American Heart Association 17POST33660875 to NR.

## Ethics Statement

Animal studies were reviewed and approved by the Institutional Animal Care and Use Committee at Boston University and performed in an American Association for the Accreditation of Laboratory Animal Care accredited facility (Animal Welfare Assurance Number: A3316-01). All animals were treated humanely and with regard for alleviation of suffering.

## Conflict of Interest

The authors declare that they have no conflict of interest.

## Acknowledgements

This work was supported by the National Institute of Environmental Health Sciences Superfund Research Program P42 ES007381 (Jennifer Schlezinger), the National Institute of Diabetes and Digestive and Kidney Diseases R01 DK117161 and the American Heart Association 17POST33660875 (Nabil Rabhi).

