## Supplemental Material for "Triphenyl phosphate is a selective PPARγ modulator that does not induce brite adipogenesis *in vitro* and *in vivo*"

**Table of Contents**

| | | PAGE |
|---|---|---|
| **TABLE S1** | Mouse and human primer sequences for reverse transcriptase qPCR. | **2** |
| **TABLE S2** | Detailed list of significant differential proteins between chemical groups. | **3** |
| **TABLE S3** | Detailed list of significant differential phosphoproteins between chemical groups. | **3** |
| **TABLE S4** | Detailed list of network pathways. | **3** |
| **FIG S1** | Weight gain and fat mass in mice treated with Rosi and TPhP. | **4** |
| **FIG S2** | Effect of rosiglitazone and TPhP on cell number in differentiating adipocytes | **5** |
| **FIG S3** | Adipocyte differentiation and gene expression in roscovitine-treated 3T3 L1 cells. | **6** |
| **FIG S4** | Adipocyte differentiation and gene expression 3T3-WT and 3T3-SA cells. | **7** |

**Table S1. Mouse (M) and human (H) primer sequences for reverse transcriptase qPCR.**

| Gene Symbol | FORWARD | REVERSE | Annealing Temp. °C |
|---|---|---|---|
| ***M-Rn18s*** | GTAACCCGTTGAACCCCATT | CCATCCAATCGGTAGTAGCG | 55 |
| ***M-B2M*** | CTGCTACGTAACACAGTTCCACCC | CATGATGCTTGATCACATGTCTCG | 55 |
| ***M-Cidea*** | TGCTCTTCTGTATCGCCCAGT | GCCGTGTTAAGGAATCTGCTG | 55 |
| ***M-Cidec*** | AGGCCCTGTCGTGTTAGCAC | CATGATGCCTTTGCGAACCT | 55 |
| ***M-Elovl3*** | TCCGCGTTCTCATGTAGGTCT | GGACCTGATGCAACCCTATGA | 55 |
| ***M-CoxIV*** | TCACTGCGCTCGTTCTGATT | CAGCATTCGCTTGGTCTGC | 55 |
| ***M-Fabp4*** | AGCCCAACATGATCATCAGC | TTTCCATCCCACTTCTGCAC | 55 |
| ***M-Plin1*** | GGGACCTGTGAGTGCTTCC | GTATTGAAGAGCCGGGATCTTTT | 55 |
| ***M-Pgc1a*** | AACAAGCACTTCGGTCATCCCTG | TTACTGAAGTCGCCATCCCTTAG | 55 |
| ***M-Ppara*** | GCGTACGGCAATGGCTTTAT | GAACGGCTTCCTCAGGTTCTT | 55 |
| ***M-Pparg2*** | TGGGTGAAACTCTGGGAGATTC | AATTTCTTGTGAAGTGCTCATAGGC | 55 |
| ***M-Rip140*** | AGAACGCACATCAGGTGGCA | GATGGCCAGACACCCCTTTG | 55 |
| ***M-Adipoq*** | GCACTGGCAAGTTCTACTGCAA | GTAGGTGAAGAGAACGGCCTTGT | 55 |
| ***M-Ucp1*** | ACTGCCACACCTCCAGTCATT | CTTTGCCTCACTCAGGATTGG | 55 |
| ***M-Acaa2*** | TAACGAGGCTGGCTACTTCAA | AGGGGCGTGAAGTTATGTTTT | 55 |
| ***M-Retn*** | TCGTGGGACATTCGTGAAGA | GGGCTGCTGTCCAGTCTATCC | 55 |
| ***M-Wdnm1*** | TGTCTGTTTGGAGCTCTGTGACCA | AACAACTCCTTGTCCCACTCCACT | 55 |
| ***H-RPL27*** | GTGAAAGTGTATAACTACAATCACC | TCAAACTTGACCTTGGCCT | 58 |
| ***H-B2M*** | GCTATCCAGCGTACTCCAAAG | CACACGGCAGGCATACTC | 58 |
| ***H-PPARG*** | TCAGCTCCGTGGATCTCTCC | ACCCTTGCATCCTTCACAAG | 55 |
| ***H-CIDEA*** | GGCAGGTTCACGTGTGGATA | GAAACACAGTGTTTGGCTCAAGA | 60 |
| ***H-PLIN1*** | ATTGCTCTGAGCTGAAGGACACCA | AGCTCGAGTGTTGGCAGCAAATTC | 60 |
| ***H-FABP4*** | GCTTTTGTAGGTACCTGGAAACTT | ACACTGATGATCATGTTAGGTTTGG | 55 |
| ***H-ACAA2*** | TGTGGTTCTGGTTTTCAGTCC | GTGACTGCAGGGCATATTTGT | 55 |
| ***H-UCP1*** | GGGCTTCAGCGGCAAATCAGCT | CAATGAATACTGCCACTCCTC | 55 |
| ***H-ELOVL3*** | ATGTAGTTCTGCCCCACAGC | AAGGACATGAGGCCCTTTTT | 55 |
| ***H-CIDEC*** | GGGATACAGTGTTCATGGTCCT | TCAATCTTCTTGGCAGGCTTATG | 55 |

**Table S2. Detailed list of proteins differentially expressed between chemical groups.**

Attached separately as an Excel file.

**Table S3. Detailed list of phosphoproteins differentially expressed between chemical groups.**

Attached separately as an Excel file.

**Table S4. Detailed list of network pathways.**

Attached separately as an Excel file.

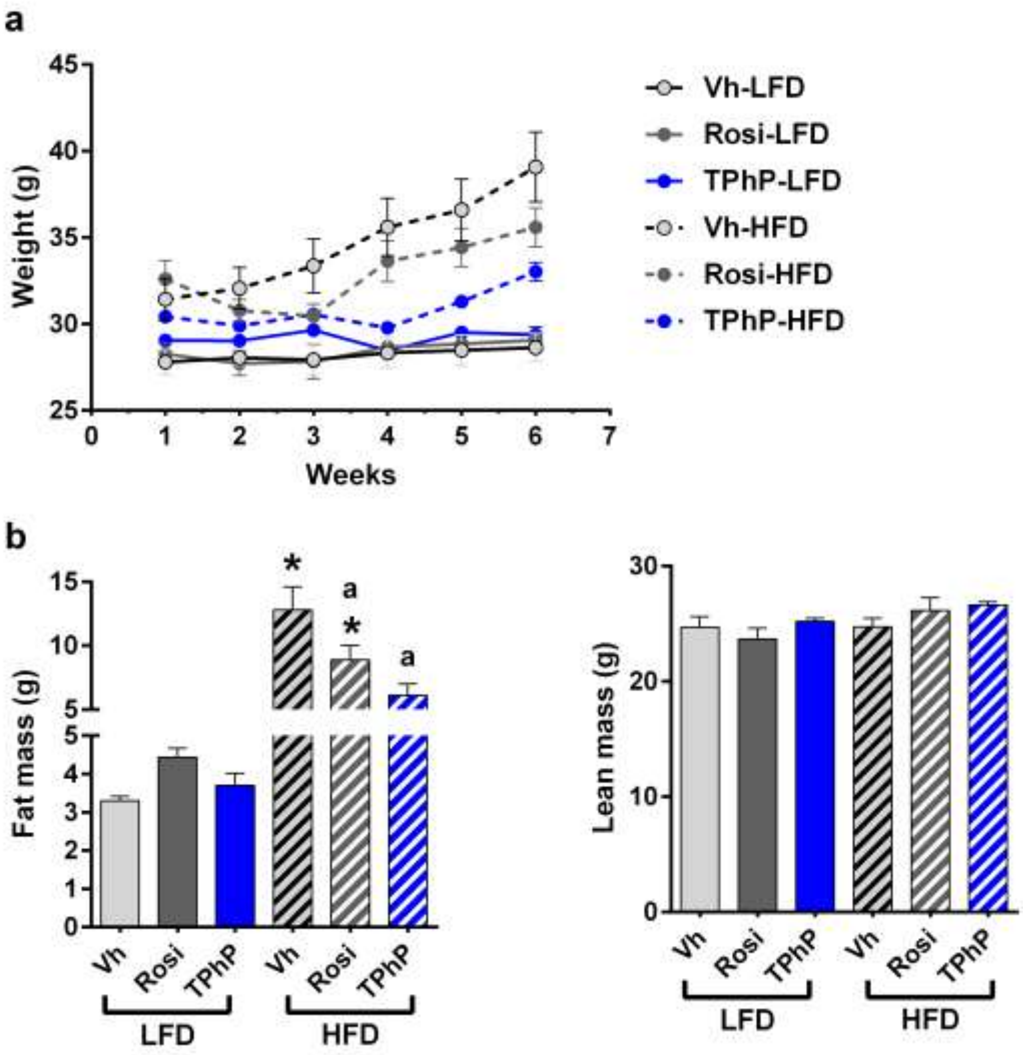

**Fig. S1.** Weight gain and fat mass in mice treated with Rosi and TPhP.
Six-week-old C57BL/6J male mice were either fed a diet with 10% kcal% fat (LFD) or 60% kcal% fat (HFD) for a total of 13 weeks. Seven weeks after initiation of the diet, mice were ip, daily for 6 weeks with vehicle, rosiglitazone (10 mg/kg), or triphenyl phosphate (10 mg/kg). **a** Weight gain. **b** Body composition at 6 weeks of treatment. Data are presented as mean + SE (n=5). Statistically different from LFD Vh-treated (*p<0.05) or HFD Vh-treated ([a]p<0.05, ANOVA, Dunnett's).

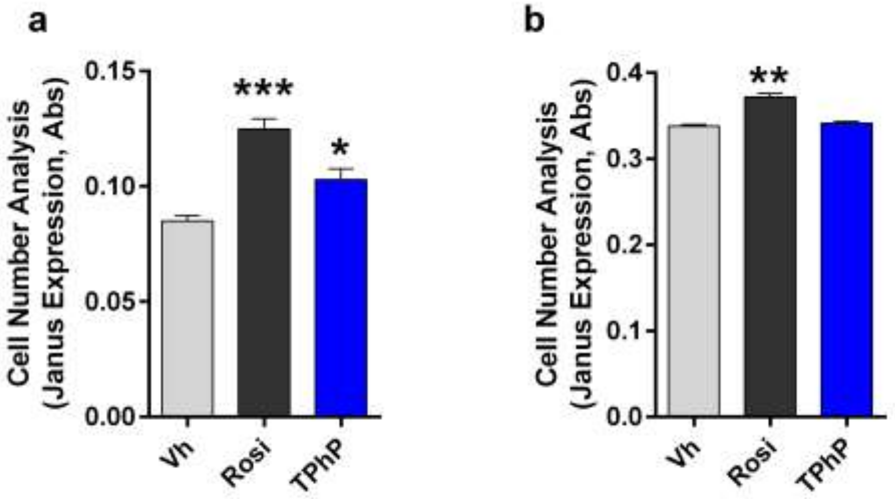

**Fig. S2.** Effect of rosiglitazone and TPhP on cell number in differentiating adipocytes.
**a** Confluent 3T3 L1 cells were differentiated using a standard hormone cocktail for 10 days. During differentiation, cells were treated with Vh (0.1% DMSO, final concentration), Rosi (20 μM), or TPhP (20 μM). On days 3, 5, and 7 of differentiation, the adipocyte maintenance medium was replaced and the cultures re-dosed. Cells were incubated for a total of 10 days of differentiation. **b** Confluent primary human adipocytes were differentiated using a standard hormone cocktail for a total of 14 days. During differentiation, cells were treated with vehicle (0.1% DMSO, final concentration), Rosi (4 μM), or TPhP (4 μM). On days 3, 5, 7, 10, and 12 of differentiation, the medium was replace and the cultures re-dosed. Cells were incubated for a total of 14 days of differentiation. Cultures were analyzed for relative cell density using JANUS staining. Data are presented as mean ± SE (n=8 for 3T3 L1 cells and n=3 for human adipocytes, each n is from pre-adipocytes from an individual). Statistically different from Vh-treated (*p<0.05, **p<0.01, ***p<0.001, ANOVA, Dunnett’s).

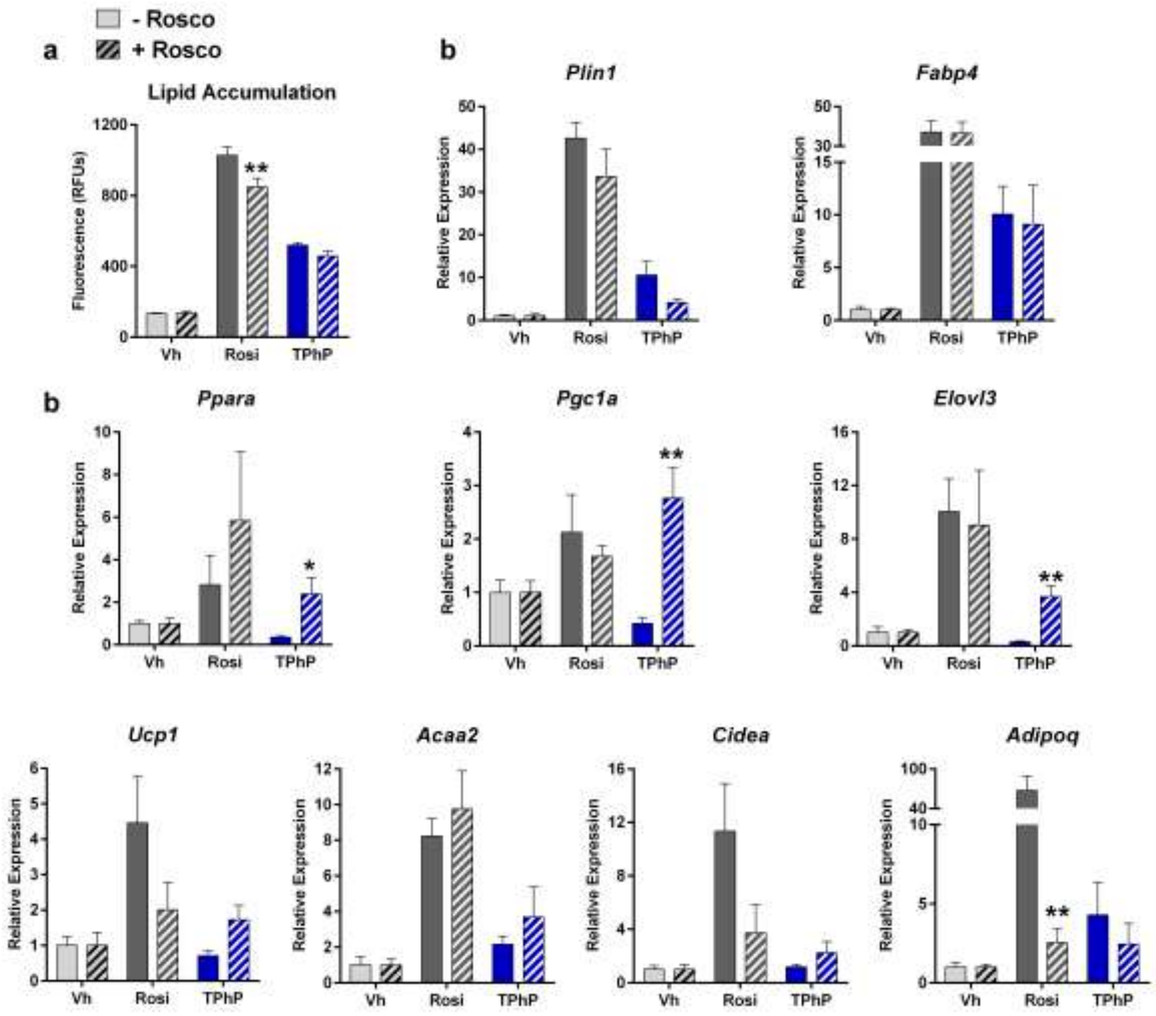

**Fig. S3.** Adipocyte differentiation and gene expression in roscovitine-treated 3T3 L1 cells. 3T3 L1 cells were differentiated using a standard hormone cocktail for 10 days. During differentiation, cells were treated with Vh (0.1% DMSO, final concentration), Rosi (20 μM), or TPhP (20 μM) in the presence or absence of roscovitine (Rosco, 20 μM). **a** Lipid accumulation was measured by Nile Red staining. Gene expression was determined by RT-qPCR analysis of white adipocyte (**b**) and brite adipocyte (**c**) marker genes. Data are presented as mean + SE (n=6). Statistically different from - Rosco (*p<0.05 or **p<0.01, 2-Factor ANOVA, Sidak's).

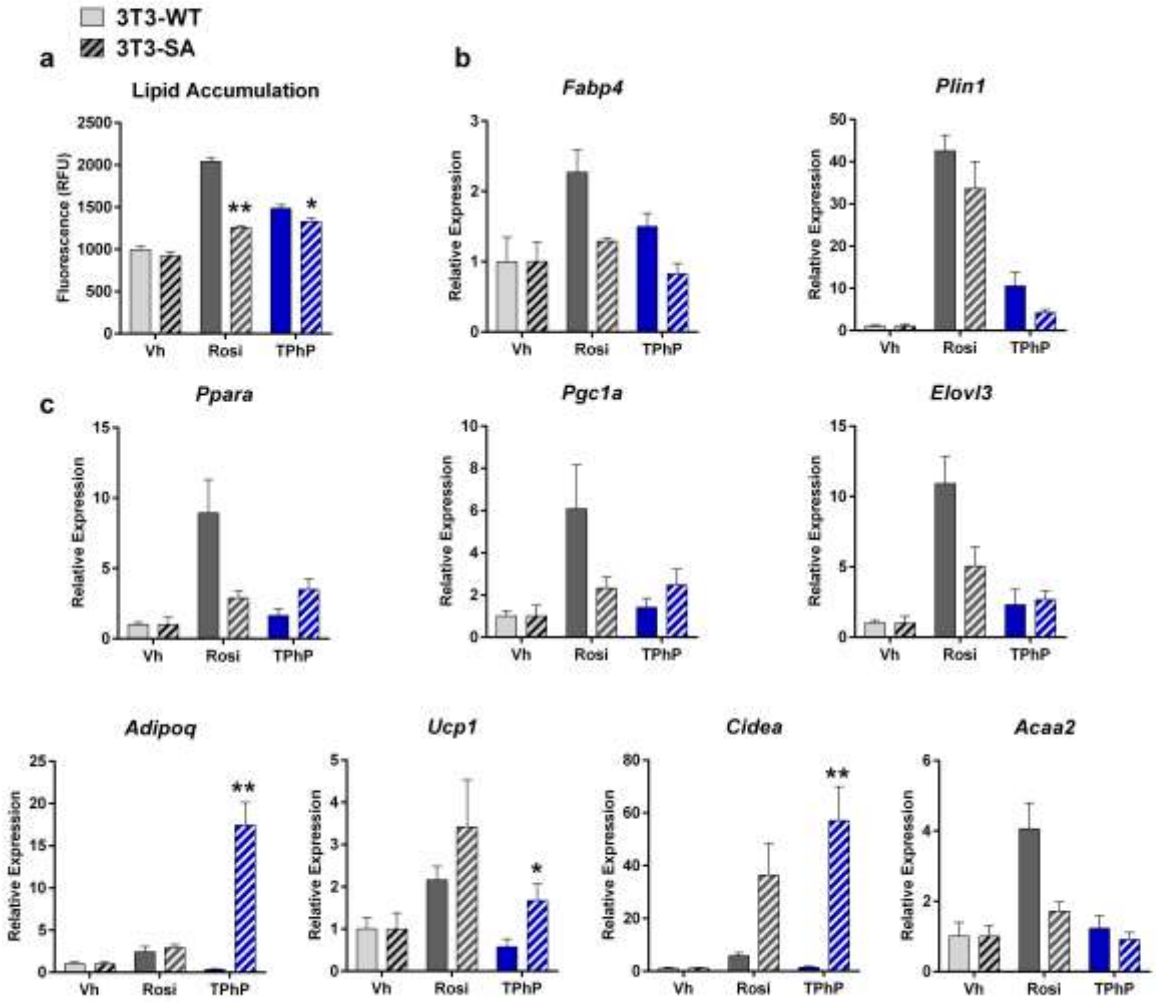

**Fig. S4.** Adipocyte differentiation and gene expression in 3T3-WT and 3T3-SA cells.
3T3 cells expressing WT PPARγ (3T3 WT) or PPARγ with alanine substituted for serine at position 273 (3T3-SA) were differentiated using a standard hormone cocktail for 10 days. During differentiation, cells were treated with Vh (0.1% DMSO, final concentration), Rosi (20 μM), or TPhP (20 μM). **a** Lipid accumulation was measured by Nile Red staining. Gene expression was determined by RT-qPCR analysis of white adipocyte (**b**) and brite adipocyte (**c**) marker genes. Data are presented as mean ± SE (n=5-6). Statistically different from 3T3-WT (*p<0.05 or **p<0.01, 2-Factor ANOVA, Sidak's).

| EM1_Name | EM1_Gene | EM1_Colou | EM1_Colou | EM1_ES..C |
|---|---|---|---|---|
| OXIDATION-REDUCTION PROCESS%GOBP%GO:0055114 | NOX4\|ME3 | 1 | 1 | 0.7015 |
| TRANSMEMBRANE TRANSPORT%GOBP%GO:0055085 | ZFAND2B\|I | 1 | 0.9361 | 0.6156 |
| FATTY ACID BETA-OXIDATION%GOBP%GO:0006635 | ACADM\|ET | 1 | 0.992 | 0.8335 |
| PLASMA MEMBRANE BOUNDED CELL PROJECTION ASSEMBLY%G | PARVA\|RA | -1 | -0.0106 | -0.623 |
| NUCLEOSIDE BISPHOSPHATE METABOLIC PROCESS%GOBP%GO:0 | ACOT7\|MP | 1 | 0.9881 | 0.8479 |
| ELECTRON TRANSPORT CHAIN%GOBP%GO:0022900 | ETFDH\|DLD | 1 | 0.9987 | 0.7045 |
| ORGANOPHOSPHATE BIOSYNTHETIC PROCESS%GOBP%GO:00904 | ADA\|PIK3C | 1 | 0.9944 | 0.6589 |
| REGULATION OF PROTEIN DEPOLYMERIZATION%GOBP%GO:1901 | WDR1\|TW | -0.986 | -0.488 | -0.5698 |
| RESPIRATORY ELECTRON TRANSPORT CHAIN%GOBP%GO:00229C | DLD\|PINK1 | 0.9947 | 0.9866 | 0.6997 |
| CARBOXYLIC ACID TRANSPORT%GOBP%GO:0046942 | PAK1\|KCNJ | 0.9963 | 0.7968 | 0.6911 |
| COENZYME METABOLIC PROCESS%GOBP%GO:0006732 | LDHA\|VNN | 1 | 0.9933 | 0.7112 |
| CITRIC ACID CYCLE (TCA CYCLE)%REACTOME%R-HSA-71403.2 | ME2\|DLD\| | 0.9981 | 0.9865 | 0.7506 |
| REGULATION OF CELL PROJECTION ORGANIZATION%GOBP%GO:( | PAK1\|GATA | -1 | 0.0975 | -0.4526 |
| CIRCADIAN RHYTHM%GOBP%GO:0007623 | NTRK3\|PPA | -0.9784 | -0.2929 | -0.5853 |
| BIOLOGICAL ADHESION%GOBP%GO:0022610 | PARVA\|CD | -1 | 0.5796 | -0.4775 |
| PURINE NUCLEOTIDE METABOLIC PROCESS%GOBP%GO:0006163 | MSH2\|NT5 | 1 | 0.9989 | 0.7065 |
| PURINE NUCLEOSIDE TRIPHOSPHATE METABOLIC PROCESS%GOB | AK3\|MSH2 | 1 | 0.9403 | 0.6319 |
| REGULATION OF LEUKOCYTE PROLIFERATION%GOBP%GO:00706 | CD209A\|AI | -0.9953 | -0.9841 | -0.6351 |
| REGULATION OF CELL MORPHOGENESIS%GOBP%GO:0022604 | PAK1\|CDH | -1 | 0.228 | -0.4841 |
| MITOCHONDRIAL MEMBRANE ORGANIZATION%GOBP%GO:0007 | BCL2L1\|SIV | 0.9966 | 0.8104 | 0.6855 |

| | | | | |
|---|---|---|---|---|
| REGULATION OF CYTOSOLIC CALCIUM ION CONCENTRATION%GC | CCL28\|TRP | 0.9769 | 0.6759 | 0.5795 |
| CELLULAR LIPID METABOLIC PROCESS%GOBP%GO:0044255 | LCN5\|PIK3( | 1 | 1 | 0.7073 |
| EPH-EPHRIN SIGNALING%REACTOME DATABASE ID RELEASE 66% | PAK1\|EPH/ | -1 | -0.4242 | -0.5692 |
| CELL-MATRIX ADHESION%GOBP%GO:0007160 | CTTN\|THY1 | -0.9978 | 0.0701 | -0.7183 |
| ERYTHROCYTE HOMEOSTASIS%GOBP%GO:0034101 | TAL1\|SOX6 | 0.993 | 0.4425 | 0.6531 |

| | | | | |
|---|---|---|---|---|
| MUSCLE CONTRACTION%REACTOME DATABASE ID RELEASE 66% | PAK1\|SCN2 | -1 | -0.9298 | -0.711 |
| PROTEIN HETEROOLIGOMERIZATION%GOBP%GO:0051291 | PARVA\|H2- | 0.6667 | 0.9962 | 0.3844 |
| NABA_MATRISOME_ASSOCIATED%MSIGDB_C2%NABA_MATRISC | IL17F\|CCL2 | -0.9973 | -0.4459 | -0.532 |
| SIGNALING BY NUCLEAR RECEPTORS%REACTOME DATABASE ID F | CDK9\|FOS\| | 1 | 0.9118 | 0.6 |

| | | | | |
|---|---|---|---|---|
| RESPONSE TO GROWTH FACTOR%GOBP%GO:0070848 | SOX6\|GAT/ | -1 | -0.9693 | -0.5219 |
| MUSCLE STRUCTURE DEVELOPMENT%GOBP%GO:0061061 | SOX6\|FLN( | -0.9972 | 0.5186 | -0.4997 |
| NUCLEOSIDE PHOSPHATE BIOSYNTHETIC PROCESS%GOBP%GO:1 | AK3\|ACOT | 1 | 0.9762 | 0.6815 |
| RIBONUCLEOSIDE TRIPHOSPHATE METABOLIC PROCESS%GOBP% | AK3\|MSH2 | 1 | 0.9553 | 0.6374 |
| PURINE NUCLEOSIDE MONOPHOSPHATE METABOLIC PROCESS%( | AK3\|MSH2 | 1 | 0.9641 | 0.6861 |

| | | | | |
|---|---|---|---|---|
| PURINE RIBONUCLEOSIDE TRIPHOSPHATE BIOSYNTHETIC PROCES | AK3\|ALDO/ | 0.9982 | 0.6386 | 0.6842 |
| MUSCLE CELL DIFFERENTIATION%GOBP%GO:0042692 | SOX6\|RAR/ | -0.9976 | 0.2806 | -0.522 |
| CARBOXYLIC ACID CATABOLIC PROCESS%GOBP%GO:0046395 | ACOT7\|CYI | 1 | 0.9904 | 0.6761 |
| DNA PACKAGING%GOBP%GO:0006323 | CDCA5\|HIF | -0.9833 | 0.3535 | -0.5881 |
| NABA_ECM_REGULATORS%MSIGDB_C2%NABA_ECM_REGULATO | SERPINB3D | -1 | -0.8102 | -0.7337 |

| | | | | |
|---|---|---|---|---|
| FATTY ACID CATABOLIC PROCESS%GOBP%GO:0009062 | CES1F\|ACC | 1 | 0.9987 | 0.8023 |
| LIPID OXIDATION%GOBP%GO:0034440 | ACADM\|ET | 1 | 1 | 0.7955 |
| PURINE RIBONUCLEOTIDE BIOSYNTHETIC PROCESS%GOBP%GO:0 | AK3\|ACOT | 1 | 0.9895 | 0.7636 |
| SULFUR COMPOUND METABOLIC PROCESS%GOBP%GO:0006790 | CHST7\|CHS | 1 | 0.9108 | 0.5889 |
| RESPIRATORY ELECTRON TRANSPORT%REACTOME DATABASE ID | ETFDH\|ND | 1 | 0.9924 | 0.736 |

| | | | | |
|---|---|---|---|---|
| CATION TRANSMEMBRANE TRANSPORT%GOBP%GO:0098655 | NALCN\|TRI | 0.995 | 0.9287 | 0.579 |
| HALLMARK_OXIDATIVE_PHOSPHORYLATION%MSIGDB_C2%HALL | LDHA\|SLC2 | 1 | 1 | 0.7349 |
| NEGATIVE REGULATION OF CELLULAR COMPONENT MOVEMENT | GATA3\|SEM | -0.992 | 0.6667 | -0.4932 |
| CRISTAE FORMATION%REACTOME DATABASE ID RELEASE 66%89 | CHCHD3\|A | 1 | 0.8661 | 0.7869 |

| | | | | |
|---|---|---|---|---|
| TISSUE MORPHOGENESIS%GOBP%GO:0048729 | NPHP3\|PAI | -1 | 0.0789 | -0.4891 |
| MONOCARBOXYLIC ACID METABOLIC PROCESS%GOBP%GO:0032 | LCN5\|PLAA | 1 | 0.9989 | 0.7321 |
| ORGANOPHOSPHATE METABOLIC PROCESS%GOBP%GO:0019637 | NT5C2\|AD/ | 1 | 1 | 0.6675 |
| NADH METABOLIC PROCESS%GOBP%GO:0006734 | SLC37A2\|L | 1 | 0.9063 | 0.8104 |
| HEDGEHOG LIGAND BIOGENESIS%REACTOME DATABASE ID RELE | PSMD2\|PS | 0.9763 | 0.9974 | 0.5399 |

| | | | | |
|---|---|---|---|---|
| MITOCHONDRIAL PROTEIN IMPORT%REACTOME%R-HSA-126802 | TOMM40\|( | 1 | 0.9948 | 0.8165 |
| POSITIVE REGULATION OF LIPID METABOLIC PROCESS%GOBP%G( | CTDNEP1\|I | 0.9912 | 0.9723 | 0.6643 |
| INORGANIC ION TRANSMEMBRANE TRANSPORT%GOBP%GO:009 | NALCN\|TRI | 1 | 0.9001 | 0.6234 |
| REGULATION OF LYMPHOCYTE PROLIFERATION%GOBP%GO:005( | CD209A\|AI | -0.9977 | -0.9835 | -0.6428 |
| NABA_MATRISOME%MSIGDB_C2%NABA_MATRISOME | CD209A\|LA | -1 | -0.9811 | -0.4897 |

| | | | | |
|---|---|---|---|---|
| CILIUM ASSEMBLY%GOBP%GO:0060271 | PARVA\|RAI | -0.9836 | -0.6137 | -0.5661 |
| THIOESTER METABOLIC PROCESS%GOBP%GO:0035383 | ACOT7\|MF | 1 | 0.9946 | 0.8629 |
| SIG_REGULATION_OF_THE_ACTIN_CYTOSKELETON_BY_RHO_GTI | PAK1\|FLNC | -0.9912 | 0.7241 | -0.6888 |
| PROTEIN LOCALIZATION TO MITOCHONDRION%GOBP%GO:0070! | DNAJC15\|T | 0.9966 | 0.9809 | 0.6536 |

| | | | | |
|---|---|---|---|---|
| TP53 REGULATES METABOLIC GENES%REACTOME DATABASE ID I | LAMTOR1\| | 0.9929 | 0.9819 | 0.5971 |
| MOVEMENT OF CELL OR SUBCELLULAR COMPONENT%GOBP%GO | PARVA\|GA | -1 | 0.1546 | -0.4521 |
| RIBONUCLEOSIDE TRIPHOSPHATE BIOSYNTHETIC PROCESS%GOBI | AK3\|CTPS2 | 1 | 0.6411 | 0.6597 |
| MITOCHONDRION ORGANIZATION%GOBP%GO:0007005 | COA3\|GGC | 1 | 0.9966 | 0.5885 |

| | | | | |
|---|---|---|---|---|
| INTERLEUKIN-1 SIGNALING%REACTOME%R-HSA-9020702.1 | PSMD2\|PS | 0.8855 | 0.9976 | 0.448 |
| ESTABLISHMENT OR MAINTENANCE OF CELL POLARITY%GOBP%( | PARVA\|NP | -0.9873 | 0.0073 | -0.5159 |
| ANTIBIOTIC METABOLIC PROCESS%GOBP%GO:0016999 | CAT\|HBA-X | 0.9984 | 0.9842 | 0.646 |
| AEROBIC RESPIRATION%GOBP%GO:0009060 | CAT\|FH1\|D | 1 | 0.9909 | 0.8133 |
| ATP BIOSYNTHETIC PROCESS%GOBP%GO:0006754 | AK3\|ALDO | 1 | 0.6297 | 0.7121 |

| | | | | |
|---|---|---|---|---|
| ANTIGEN PROCESSING-CROSS PRESENTATION%REACTOME DATA | PSMD2\|ST. | 0.968 | 1 | 0.4708 |
| REGULATION OF ACTIN FILAMENT-BASED PROCESS%GOBP%GO:( | PAK1\|TWF | -1 | 0.7773 | -0.5276 |
| CELL MORPHOGENESIS%GOBP%GO:0000902 | PARVA\|SO. | -1 | 0.8254 | -0.4415 |
| CARBOHYDRATE DERIVATIVE BIOSYNTHETIC PROCESS%GOBP%G( | ADA\|CHST | 1 | 0.9624 | 0.5849 |

| | | | | |
|---|---|---|---|---|
| REGULATION OF GENE EXPRESSION, EPIGENETIC%GOBP%GO:004 | CBX8\|RBM | -0.993 | -0.9384 | -0.5601 |
| TRICARBOXYLIC ACID METABOLIC PROCESS%GOBP%GO:0072350 | GM5424\|S | 1 | 0.9608 | 0.748 |
| ACTIN FILAMENT BUNDLE ASSEMBLY%GOBP%GO:0051017 | HSP90B1\|E | -1 | 0.0084 | -0.7511 |
| REGULATION OF EXTENT OF CELL GROWTH%GOBP%GO:0061387 | NTRK3\|PAI | -0.9843 | 0.3831 | -0.6292 |
| ESTABLISHMENT OF PROTEIN LOCALIZATION TO MITOCHONDRIC | DNAJC15\|T | 0.9914 | 0.987 | 0.6502 |

| | | | | |
|---|---|---|---|---|
| CELL JUNCTION ASSEMBLY%GOBP%GO:0034329 | CDH4\|CDH | -0.9975 | 0.2748 | -0.5827 |
| PID_CDC42_PATHWAY%MSIGDB_C2%PID_CDC42_PATHWAY | RASGRF1\|R | -0.9857 | 0.3818 | -0.6224 |
| MHC CLASS II ANTIGEN PRESENTATION%REACTOME%R-HSA-213 | CENPE\|CTS | -0.9848 | 0.25 | -0.5041 |
| SMALL MOLECULE CATABOLIC PROCESS%GOBP%GO:0044282 | INPP5B\|AD | 1 | 0.9548 | 0.5643 |
| ORGANIC ACID CATABOLIC PROCESS%GOBP%GO:0016054 | ACOT7\|CYP | 1 | 0.9893 | 0.6761 |

| | | | | |
|---|---|---|---|---|
| CYTOSKELETON ORGANIZATION%GOBP%GO:0007010 | PARVA\|PAI | -1 | -0.0286 | -0.562 |
| REGULATION OF ANATOMICAL STRUCTURE SIZE%GOBP%GO:009 | PAK1\|SNTA | -1 | 0.766 | -0.5016 |
| PROTEIN COMPLEX OLIGOMERIZATION%GOBP%GO:0051259 | PARVA\|AR | 1 | 0.9606 | 0.4673 |
| GLYCEROLIPID METABOLIC PROCESS%GOBP%GO:0046486 | INPP5B\|CA | 1 | 0.9974 | 0.7325 |
| CELL PROJECTION ASSEMBLY%GOBP%GO:0030031 | PARVA\|RAI | -1 | -0.0058 | -0.6312 |

| | | | | |
|---|---|---|---|---|
| AROMATIC COMPOUND BIOSYNTHETIC PROCESS%GOBP%GO:00 | CDK9\|GAT/ | 1 | 0.9594 | 0.5423 |
| REGULATION OF LIPID BIOSYNTHETIC PROCESS%GOBP%GO:0046 | CTDNEP1\|/ | 0.9945 | 0.9973 | 0.7229 |
| REGULATION OF SMALL MOLECULE METABOLIC PROCESS%GOBP | ME1\|PPAR | 1 | 0.8619 | 0.6625 |
| MONOCARBOXYLIC ACID CATABOLIC PROCESS%GOBP%GO:0072: | CES1F\|ACC | 1 | 0.9975 | 0.7831 |

| | | | | |
|---|---|---|---|---|
| PURINE RIBONUCLEOSIDE MONOPHOSPHATE METABOLIC PROCE | AK3\|MSH2 | 1 | 0.9676 | 0.6861 |
| HEART MORPHOGENESIS%GOBP%GO:0003007 | PARVA\|NP | -0.9975 | -0.0922 | -0.5988 |
| PURINE-CONTAINING COMPOUND METABOLIC PROCESS%GOBP% | MSH2\|NT5 | 1 | 0.9957 | 0.6931 |
| ACTIN FILAMENT-BASED MOVEMENT%GOBP%GO:0030048 | PARVA\|SN | -1 | -0.0199 | -0.7077 |
| NUCLEOSIDE PHOSPHATE METABOLIC PROCESS%GOBP%GO:000 | NT5C2\|AD | 1 | 1 | 0.694 |

| | | | | |
|---|---|---|---|---|
| REGULATION OF PLASMA MEMBRANE BOUNDED CELL PROJECTIO | PAK1\|GAT/ | -1 | 0.1081 | -0.4526 |
| PYRUVATE METABOLIC PROCESS%GOBP%GO:0006090 | ME1\|MPC2 | 0.9982 | 0.5591 | 0.7295 |
| REACTIVE OXYGEN SPECIES METABOLIC PROCESS%GOBP%GO:00 | CAT\|NOX4 | 0.9784 | 0.3934 | 0.6037 |
| RIBONUCLEOSIDE MONOPHOSPHATE BIOSYNTHETIC PROCESS%G | AK3\|UMPS | 1 | 0.8562 | 0.6803 |

| | | | | |
|---|---|---|---|---|
| MONOVALENT INORGANIC CATION TRANSPORT%GOBP%GO:001 | NALCN\|KCI | 0.9878 | 0.5962 | 0.6966 |
| REGULATION OF MONONUCLEAR CELL PROLIFERATION%GOBP%( | CD209A\|AI | -1 | -0.9705 | -0.6428 |
| ABC TRANSPORTER DISORDERS%REACTOME DATABASE ID RELEA | PSMD2\|PS | 0.9515 | 0.9938 | 0.5083 |
| COLLAGEN FORMATION%REACTOME%R-HSA-1474290.1 | PLOD1\|PLC | -0.9857 | 0.1598 | -0.5926 |
| MUSCLE ORGAN DEVELOPMENT%GOBP%GO:0007517 | FOS\|NKX2- | -0.9855 | 0.6568 | -0.5487 |

| | | | | |
|---|---|---|---|---|
| AMMONIUM ION METABOLIC PROCESS%GOBP%GO:0097164 | ODC1\|SPH | 1 | 0.7263 | 0.7902 |
| ACTIN CYTOSKELETON ORGANIZATION%GOBP%GO:0030036 | PARVA\|PAI | -1 | 0.0066 | -0.6371 |
| REGULATION OF SUPRAMOLECULAR FIBER ORGANIZATION%GOE | PAK1\|TWF | -1 | 0.7745 | -0.4877 |
| NUCLEOBASE-CONTAINING COMPOUND BIOSYNTHETIC PROCESS | CDK9\|CDK | 1 | 0.9636 | 0.5484 |

| | | | | |
|---|---|---|---|---|
| PURINE NUCLEOSIDE MONOPHOSPHATE BIOSYNTHETIC PROCESS | AK3\|HPRT\| | 1 | 0.837 | 0.685 |
| PROTEIN TETRAMERIZATION%GOBP%GO:0051262 | ME1\|CAT\| | 0.9983 | 0.9756 | 0.6111 |
| HALLMARK_PEROXISOME%MSIGDB_C2%HALLMARK_PEROXISOM | TTR\|MSH2 | 1 | 0.6495 | 0.7659 |
| ACTIN FILAMENT BUNDLE ORGANIZATION%GOBP%GO:0061572 | HSP90B1\|E | -1 | 0.0067 | -0.7511 |
| REGULATION OF AXONOGENESIS%GOBP%GO:0050770 | PAK1\|CDH | -0.9905 | 0.139 | -0.5718 |

| | | | | |
|---|---|---|---|---|
| SUPRAMOLECULAR FIBER ORGANIZATION%GOBP%GO:0097435 | HSP90B1\|T | -1 | -0.0423 | -0.5161 |
| REGULATION OF TUMOR NECROSIS FACTOR SUPERFAMILY CYTOI | RARA\|CCR: | 0.9782 | 0.5243 | 0.6592 |
| REGULATION OF T CELL PROLIFERATION%GOBP%GO:0042129 | CD209A\|C/ | -0.9954 | -0.9759 | -0.6702 |
| RIBONUCLEOSIDE MONOPHOSPHATE METABOLIC PROCESS%GOI | AK3\|MSH2 | 1 | 0.9676 | 0.6831 |

| | | | | |
|---|---|---|---|---|
| REGULATION OF ACTIN FILAMENT DEPOLYMERIZATION%GOBP%( | WDR1\|TW | -0.9929 | -0.5697 | -0.6246 |
| MITOCHONDRIAL FATTY ACID BETA-OXIDATION%REACTOME%R- | ACADM\|A( | 1 | 0.9755 | 0.8135 |
| REGULATION OF LIPID METABOLIC PROCESS%GOBP%GO:001921 | CTDNEP1\|I | 1 | 0.9976 | 0.6759 |
| RESPONSE TO INTERFERON-GAMMA%GOBP%GO:0034341 | CCL22\|STX | -0.9931 | 0.0388 | -0.6556 |
| PID_PDGFRB_PATHWAY%MSIGDB_C2%PID_PDGFRB_PATHWAY | FOS\|PAK1\| | -0.9975 | 0.8337 | -0.4959 |

| | | | | |
|---|---|---|---|---|
| DRUG METABOLIC PROCESS%GOBP%GO:0017144 | NT5C2\|AD/ | 1 | 0.9946 | 0.6055 |
| REGULATION OF PROTEIN COMPLEX DISASSEMBLY%GOBP%GO:0 | TNF\|WDR1 | -0.9974 | -0.881 | -0.5744 |
| PURINE RIBONUCLEOTIDE METABOLIC PROCESS%GOBP%GO:000 | MSH2\|NT5 | 1 | 0.9989 | 0.7306 |
| REGULATION OF TUMOR NECROSIS FACTOR PRODUCTION%GOBI | RARA\|CCR. | 0.983 | 0.5451 | 0.6592 |
| REGULATION OF PROTEIN POLYMERIZATION%GOBP%GO:003227 | PAK1\|TWF | -0.9975 | 0.9693 | -0.5224 |

| | | | | |
|---|---|---|---|---|
| REGULATION OF ACTIN FILAMENT ORGANIZATION%GOBP%GO:0 | PAK1\|TWF | -1 | 0.8994 | -0.5675 |
| PID_CXCR4_PATHWAY%MSIGDB_C2%PID_CXCR4_PATHWAY | PAK1\|PIK3 | -0.9812 | 0.1923 | -0.5903 |
| HALLMARK_BILE_ACID_METABOLISM%MSIGDB_C2%HALLMARK_ | TTR\|CAT\|A | 0.9876 | 0.5409 | 0.6889 |
| POSITIVE REGULATION OF ACTIN FILAMENT POLYMERIZATION%G | CCL24\|EVL | -0.9952 | 0.9363 | -0.6446 |

| | | | | |
|---|---|---|---|---|
| STRIATED MUSCLE CELL DIFFERENTIATION%GOBP%GO:0051146 | SOX6\|RAR/ | -0.995 | 0.3374 | -0.5475 |
| VEGFA-VEGFR2 PATHWAY%REACTOME DATABASE ID RELEASE 6( | PAK1\|CYFII | -0.9815 | 0.2984 | -0.6216 |
| NEGATIVE REGULATION OF LEUKOCYTE CELL-CELL ADHESION%G( | PPARA\|CCI | -0.9771 | -0.7483 | -0.6336 |
| SEMAPHORIN INTERACTIONS%REACTOME DATABASE ID RELEASI | PTPRC\|PAI | -0.9957 | 0.0253 | -0.7022 |

| | | | | |
|---|---|---|---|---|
| LOCALIZATION OF CELL%GOBP%GO:0051674 | PARVA\|GA | -1 | -0.1343 | -0.488 |
| ION TRANSPORT%GOBP%GO:0006811 | PAK1\|ADA | 1 | 0.7874 | 0.5158 |
| MITOCHONDRIAL BIOGENESIS%REACTOME%R-HSA-1592230.2 | PPARA\|ESF | 1 | 0.9712 | 0.7573 |
| MUSCLE CELL DEVELOPMENT%GOBP%GO:0055001 | KEL\|FLNC\| | -1 | -0.5187 | -0.7207 |
| INFLAMMATION MEDIATED BY CHEMOKINE AND CYTOKINE SIGN | CCL22\|PAK | -0.9975 | 0.7285 | -0.5664 |

| | | | | |
|---|---|---|---|---|
| CELLULAR RESPONSE TO GROWTH FACTOR STIMULUS%GOBP%G | SOX6\|GAT/ | -1 | -0.971 | -0.5216 |
| SIGNALING BY RHO GTPASES%REACTOME%R-HSA-194315.4 | DEPDC7\|P/ | -1 | 0.3414 | -0.5827 |
| RIBONUCLEOTIDE BIOSYNTHETIC PROCESS%GOBP%GO:0009260 | AK3\|ACOT | 1 | 0.9977 | 0.7433 |
| REGULATION OF CELLULAR KETONE METABOLIC PROCESS%GOBI | PPARA\|CO | 0.9982 | 0.9728 | 0.7423 |

| | | | | |
|---|---|---|---|---|
| POSITIVE REGULATION OF SUPRAMOLECULAR FIBER ORGANIZAT | PAK1\|CCL2 | -1 | 0.8106 | -0.6331 |
| RESPIRATORY ELECTRON TRANSPORT, ATP SYNTHESIS BY CHEMI( | ATP5S\|MT- | 1 | 0.9926 | 0.764 |
| CARBOXYLIC ACID METABOLIC PROCESS%GOBP%GO:0019752 | LCN5\|NOX | 1 | 0.999 | 0.5811 |

| | | | | |
|---|---|---|---|---|
| CELL DEVELOPMENT%GOBP%GO:0048468 | PARVA\|PAI | -1 | 0.6078 | -0.4387 |
| REGULATION OF CYTOSKELETON ORGANIZATION%GOBP%GO:00: | PAK1\|TWF | -0.9968 | 0.6641 | -0.4928 |
| GOLGI-TO-ER RETROGRADE TRANSPORT%REACTOME%R-HSA-88! | RAB6A\|CEI | -1 | 0.3015 | -0.538 |
| PROTEIN HOMOTETRAMERIZATION%GOBP%GO:0051289 | ACADM\|A( | 0.9983 | 0.9318 | 0.6752 |
| OXOACID METABOLIC PROCESS%GOBP%GO:0043436 | CHST7\|CHS | 1 | 1 | 0.5754 |

| | | | | |
|---|---|---|---|---|
| METABOLISM OF STEROIDS%REACTOME DATABASE ID RELEASE 6 | PPARA\|HSI | 1 | 0.7456 | 0.7103 |
| COFACTOR METABOLIC PROCESS%GOBP%GO:0051186 | GSTM2\|MI | 1 | 1 | 0.661 |
| AMEBOIDAL-TYPE CELL MIGRATION%GOBP%GO:0001667 | NANOS1\|S | -0.9811 | 0.1782 | -0.5549 |
| PID_RAC1_PATHWAY%MSIGDB_C2%PID_RAC1_PATHWAY | PAK1\|CYFII | -0.9799 | 0.2397 | -0.6341 |

| | | | | |
|---|---|---|---|---|
| PDGFR-BETA SIGNALING PATHWAY%PATHWAY INTERACTION DA | PAK1\|FOS\| | -0.9923 | 0.8374 | -0.4959 |
| STRIATED MUSCLE CELL DEVELOPMENT%GOBP%GO:0055002 | KEL\|FLNC\| | -1 | -0.3259 | -0.7419 |
| PLASMA MEMBRANE BOUNDED CELL PROJECTION ORGANIZATIC | PARVA\|GA | -1 | 0.4175 | -0.4512 |

| | | | | |
|---|---|---|---|---|
| CARBOHYDRATE BIOSYNTHETIC PROCESS%GOBP%GO:0016051 | ST3GAL4\|A | 0.9786 | 0.9797 | 0.6245 |
| RIBONUCLEOSIDE BISPHOSPHATE METABOLIC PROCESS%GOBP% | ACOT7\|MF | 1 | 0.996 | 0.8479 |
| PYRIDINE NUCLEOTIDE METABOLIC PROCESS%GOBP%GO:00193( | MPC2\|LDH | 1 | 0.9091 | 0.7274 |
| OXIDOREDUCTION COENZYME METABOLIC PROCESS%GOBP%GO | MPC2\|LDH | 1 | 0.8803 | 0.734 |
| RESPONSE TO TOXIC SUBSTANCE%GOBP%GO:0009636 | PPARA\|CA | 0.505 | 0.9988 | 0.3118 |

| | | | | |
|---|---|---|---|---|
| CELL CYCLE PROCESS%GOBP%GO:0022402 | MYB\|UHM | -0.997 | 0.0654 | -0.4266 |
| LIPID BIOSYNTHETIC PROCESS%GOBP%GO:0008610 | PIK3CA\|SR | 1 | 0.9611 | 0.7118 |
| DEFECTIVE CFTR CAUSES CYSTIC FIBROSIS%REACTOME DATABAS | PSMD2\|PS | 0.9642 | 0.995 | 0.5083 |
| MORPHOGENESIS OF AN EPITHELIUM%GOBP%GO:0002009 | NPHP3\|PA | -0.9973 | 0.3054 | -0.4788 |
| NUCLEOTIDE BIOSYNTHETIC PROCESS%GOBP%GO:0009165 | AK3\|ACOT | 1 | 0.9792 | 0.6733 |

| | | | | |
|---|---|---|---|---|
| RIBOSE PHOSPHATE METABOLIC PROCESS%GOBP%GO:0019693 | MSH2\|NT5 | 1 | 0.9978 | 0.7091 |
| HEAD DEVELOPMENT%GOBP%GO:0060322 | PAK1\|ARL1 | -1 | 0.5196 | -0.4479 |
| COFACTOR BIOSYNTHETIC PROCESS%GOBP%GO:0051188 | ACOT7\|MF | 1 | 0.7866 | 0.6669 |
| NICOTINAMIDE NUCLEOTIDE METABOLIC PROCESS%GOBP%GO:( | MPC2\|LDH | 1 | 0.9215 | 0.7274 |
| PROTEIN TARGETING TO MITOCHONDRION%GOBP%GO:000662€ | DNAJC15\|T | 0.9873 | 0.9481 | 0.6896 |

| | | | | |
|---|---|---|---|---|
| ER TO GOLGI ANTEROGRADE TRANSPORT%REACTOME DATABAS | SEC22A\|RA | -0.9974 | 0.3933 | -0.5601 |
| COENZYME BIOSYNTHETIC PROCESS%GOBP%GO:0009108 | ACOT7\|MF | 1 | 0.8316 | 0.6618 |
| CELL-SUBSTRATE ADHESION%GOBP%GO:0031589 | PARVA\|FNI | -1 | 0.0646 | -0.6596 |
| POSITIVE REGULATION OF PROTEIN POLYMERIZATION%GOBP%G | PAK1\|CCL2 | -0.9974 | 0.9328 | -0.6007 |
| ORGANIC ANION TRANSPORT%GOBP%GO:0015711 | PAK1\|KCNI | 0.993 | 0.7987 | 0.6713 |

| | | | | |
|---|---|---|---|---|
| HALLMARK_COAGULATION%MSIGDB_C2%HALLMARK_COAGULA | A2M\|C2\|LI | -1 | -0.5021 | -0.6585 |
| CELLULAR AMINO ACID CATABOLIC PROCESS%GOBP%GO:00090 | OTC\|TDO2 | 0.9751 | 0.9873 | 0.5976 |
| RIBONUCLEOTIDE METABOLIC PROCESS%GOBP%GO:0009259 | MSH2\|NT5 | 1 | 1 | 0.7244 |
| CELL MIGRATION%GOBP%GO:0016477 | PARVA\|GA | -1 | -0.1807 | -0.4945 |
| ION TRANSMEMBRANE TRANSPORT%GOBP%GO:0034220 | KCNN4\|KC | 1 | 0.8816 | 0.5827 |

| | | | | |
|---|---|---|---|---|
| MONOCARBOXYLIC ACID BIOSYNTHETIC PROCESS%GOBP%GO:00 | ACOT7\|LDI | 0.9825 | 0.7074 | 0.5906 |
| COPI-MEDIATED ANTEROGRADE TRANSPORT%REACTOME DATAI | TMED3\|RA | -0.9932 | 0.384 | -0.5891 |
| RESPONSE TO BACTERIUM%GOBP%GO:0009617 | MIR381\|M | 0.9861 | 0.6572 | 0.53 |
| MITOCHONDRIAL TRANSMEMBRANE TRANSPORT%GOBP%GO:19 | MPC2\|DNA | 1 | 0.9908 | 0.8237 |
| GLUCOSE METABOLIC PROCESS%GOBP%GO:0006006 | TNF\|PPAR/ | 0.9947 | 0.9946 | 0.6764 |

| | | | | |
|---|---|---|---|---|
| PURINE-CONTAINING COMPOUND BIOSYNTHETIC PROCESS%GOE | AK3\|ACOT | 1 | 0.9436 | 0.7203 |
| AMINO ACID AND DERIVATIVE METABOLISM%REACTOME%R-HS/ | TAT\|ALDH9 | 0.9986 | 0.9117 | 0.4485 |
| HALLMARK_FATTY_ACID_METABOLISM%MSIGDB_C2%HALLMAR | ME1\|PPAR | 1 | 1 | 0.7274 |

| | | | | |
|---|---|---|---|---|
| NUCLEOBASE-CONTAINING SMALL MOLECULE METABOLIC PROC | NT5C2\|AD/ | 1 | 0.999 | 0.6678 |
| SYNAPSE ORGANIZATION%GOBP%GO:0050808 | PAK1\|SNT/ | -0.9974 | 0.6351 | -0.573 |
| ORGANIC ACID METABOLIC PROCESS%GOBP%GO:0006082 | NOX4\|PTG | 1 | 0.999 | 0.5807 |
| REGULATION OF CELLULAR COMPONENT SIZE%GOBP%GO:0032[illegible] | PAK1\|CDH[illegible] | -1 | 0.7681 | -0.5089 |

| | | | | |
|---|---|---|---|---|
| PYRUVATE METABOLISM AND CITRIC ACID (TCA) CYCLE%REACTO | ME1\|MPC2 | 1 | 0.9543 | 0.7141 |
| ATP SYNTHESIS COUPLED ELECTRON TRANSPORT%GOBP%GO:00 | COX4I1\|DL | 1 | 0.9754 | 0.7382 |
| MITOCHONDRIAL TRANSPORT%GOBP%GO:0006839 | BCL2L1\|MI | 0.9984 | 0.9692 | 0.6917 |
| CELLULAR RESPIRATION%GOBP%GO:0045333 | CAT\|FASTK | 1 | 0.9988 | 0.7578 |
| PYRIDINE-CONTAINING COMPOUND METABOLIC PROCESS%GOB | MPC2\|LDH | 1 | 0.9099 | 0.7027 |

| | | | | |
|---|---|---|---|---|
| PURINE NUCLEOTIDE BIOSYNTHETIC PROCESS%GOBP%GO:00061 | AK3\|ACOT | 1 | 0.9336 | 0.729 |
| RHO GTPASE EFFECTORS%REACTOME DATABASE ID RELEASE 66% | PAK1\|AR\|N | -1 | 0.549 | -0.586 |
| ATP METABOLIC PROCESS%GOBP%GO:0046034 | AK3\|MSH2 | 1 | 0.9044 | 0.6899 |
| INTEGRIN SIGNALLING PATHWAY%PANTHER PATHWAY%P00034 | PARVA\|PIK | -1 | 0.9044 | -0.5956 |
| ENERGY DERIVATION BY OXIDATION OF ORGANIC COMPOUNDS% | PYGB\|CAT\| | 1 | 1 | 0.7179 |

| | | | | |
|---|---|---|---|---|
| NUCLEOTIDE METABOLIC PROCESS%GOBP%GO:0009117 | NT5C2\|AD/ | 1 | 0.9989 | 0.694 |
| PURINE RIBONUCLEOSIDE MONOPHOSPHATE BIOSYNTHETIC PRC | AK3\|HPRT\| | 1 | 0.8509 | 0.685 |
| ACTIN FILAMENT-BASED PROCESS%GOBP%GO:0030029 | PARVA\|PA | -1 | 0.0044 | -0.638 |
| CHROMATIN ASSEMBLY%GOBP%GO:0031497 | HIRA\|H1F0 | -0.9928 | 0.2959 | -0.6146 |
| HALLMARK_APICAL_JUNCTION%MSIGDB_C2%HALLMARK_APICA | PARVA\|CD. | -1 | 0.2694 | -0.59 |

| | | | | |
|---|---|---|---|---|
| COFACTOR CATABOLIC PROCESS%GOBP%GO:0051187 | ACOT7\|CAT | 1 | 0.971 | 0.7638 |
| ACTOMYOSIN STRUCTURE ORGANIZATION%GOBP%GO:0031032 | WDR1\|FRN | -1 | -0.797 | -0.7916 |
| HALLMARK_MTORC1_SIGNALING%MSIGDB_C2%HALLMARK_MT | MCM4\|ME | 1 | 0.9989 | 0.5796 |
| RIBOSE PHOSPHATE BIOSYNTHETIC PROCESS%GOBP%GO:00463S | AK3\|ACOT | 1 | 0.9774 | 0.7165 |

| | | | | |
|---|---|---|---|---|
| GENERATION OF PRECURSOR METABOLITES AND ENERGY%GOBF | MSH2\|CAT | 1 | 0.9955 | 0.6695 |
| CELLULAR LIPID CATABOLIC PROCESS%GOBP%GO:0044242 | ACOT7\|CYI | 1 | 0.9915 | 0.6804 |
| CELLULAR COMPONENT ASSEMBLY INVOLVED IN MORPHOGENES | WDR1\|NEU | -1 | -0.8163 | -0.8408 |
| REGULATION OF CELL PROJECTION ASSEMBLY%GOBP%GO:0060⁴ | DEF8\|MYO | -0.9857 | -0.4314 | -0.5417 |

| | | | | |
|---|---|---|---|---|
| NUCLEOSIDE MONOPHOSPHATE BIOSYNTHETIC PROCESS%GOBP | AK3\|UMPS | 1 | 0.8494 | 0.6803 |
| LIPID METABOLIC PROCESS%GOBP%GO:0006629 | NPHP3\|PIK | 1 | 1 | 0.7086 |
| MITOCHONDRIAL ATP SYNTHESIS COUPLED ELECTRON TRANSPO | COX4I1\|DL | 0.9983 | 0.9685 | 0.7382 |
| HALLMARK_MITOTIC_SPINDLE%MSIGDB_C2%HALLMARK_MITO | CENPE\|RFC | -1 | 0 | -0.6597 |

| | | | | |
|---|---|---|---|---|
| CYTOSKELETAL REGULATION BY RHO GTPASE%PANTHER PATHW/ | PAK1\|ACTE | -1 | 0.2886 | -0.6888 |
| REGULATION OF PLASMA MEMBRANE BOUNDED CELL PROJECTI( | DEF8\|MYO | -0.9879 | -0.3947 | -0.5417 |
| PROTON TRANSMEMBRANE TRANSPORT%GOBP%GO:1902600 | SLC36A3\|A | 0.9982 | 0.9609 | 0.7622 |
| ORGANIC CYCLIC COMPOUND BIOSYNTHETIC PROCESS%GOBP%( | CDK9\|GAT/ | 1 | 0.9672 | 0.5329 |

| | | | | |
|---|---|---|---|---|
| REGULATION OF ACTIN FILAMENT POLYMERIZATION%GOBP%GO | HAX1\|TWF | -1 | 0.9599 | -0.5547 |
| ANIMAL ORGAN MORPHOGENESIS%GOBP%GO:0009887 | PARVA\|GA | -1 | -0.0556 | -0.4355 |
| ESTABLISHMENT OF ORGANELLE LOCALIZATION%GOBP%GO:005 | SAPCD2\|SY | -0.9972 | 0.1877 | -0.4692 |
| PROTEIN FOLDING%GOBP%GO:0006457 | HSPE1\|DN/ | 1 | 0.4505 | 0.6599 |
| NUCLEOSIDE TRIPHOSPHATE METABOLIC PROCESS%GOBP%GO:( | AK3\|MSH2 | 1 | 0.9328 | 0.6194 |

| | | | | |
|---|---|---|---|---|
| CHAPERONE-MEDIATED PROTEIN FOLDING%GOBP%GO:0061077 | HSPE1\|HSF | 0.9947 | 0.8708 | 0.7029 |
| CELL MOTILITY%GOBP%GO:0048870 | PARVA\|GA | -1 | -0.1216 | -0.488 |
| TRANSPORT TO THE GOLGI AND SUBSEQUENT MODIFICATION%F | ST3GAL4\|S | -1 | 0.4535 | -0.5601 |

| | | | | |
|---|---|---|---|---|
| HALLMARK_EPITHELIAL_MESENCHYMAL_TRANSITION%MSIGDB_ | PLOD1\|PL( | -0.9971 | -0.8803 | -0.509 |
| CELLULAR COMPONENT MORPHOGENESIS%GOBP%GO:0032989 | PARVA\|GA | -1 | 0.8344 | -0.4267 |
| TCR SIGNALING%REACTOME DATABASE ID RELEASE 66%202403 | PAK1\|PIK3( | 0.6371 | 0.9975 | 0.3672 |
| CELL JUNCTION ORGANIZATION%GOBP%GO:0034330 | CDH4\|CDH | -1 | 0.2127 | -0.6129 |

| | | | | |
|---|---|---|---|---|
| HALLMARK_MYC_TARGETS_V1%MSIGDB_C2%HALLMARK_MYC_ | MCM4\|AB( | 0.8256 | 1 | 0.332 |
| PURINE NUCLEOSIDE TRIPHOSPHATE BIOSYNTHETIC PROCESS%G | AK3\|ALDO/ | 1 | 0.6324 | 0.6593 |
| POSITIVE REGULATION OF CYTOSOLIC CALCIUM ION CONCENTRA | CCL28\|TRP | 0.9947 | 0.6425 | 0.633 |
| RHYTHMIC PROCESS%GOBP%GO:0048511 | NTRK3\|PP/ | -0.9908 | -0.3093 | -0.5824 |

| | | | | |
|---|---|---|---|---|
| HALLMARK_CHOLESTEROL_HOMEOSTASIS%MSIGDB_C2%HALLN | ATF3\|FDPS | 0.9928 | 0.8435 | 0.6953 |
| ACYL-COA METABOLIC PROCESS%GOBP%GO:0006637 | ACOT7\|MF | 1 | 0.9936 | 0.8629 |
| EXTRACELLULAR MATRIX ORGANIZATION%REACTOME DATABASI | TTR\|OPTC\| | -1 | -0.8027 | -0.6044 |
| GLYCEROPHOSPHOLIPID BIOSYNTHESIS%REACTOME DATABASE I | ETNPPL\|CS | 0.9964 | 0.9732 | 0.711 |

| | | | | |
|---|---|---|---|---|
| REGULATION OF ACTIN CYTOSKELETON ORGANIZATION%GOBP% | PAK1\|TWF | -1 | 0.8562 | -0.542 |
| ER-PHAGOSOME PATHWAY%REACTOME%R-HSA-1236974.4 | PSMD2\|ST | 0.9921 | 1 | 0.5389 |
| PURINE RIBONUCLEOSIDE TRIPHOSPHATE METABOLIC PROCESS% | AK3\|MSH2 | 1 | 0.9516 | 0.6519 |
| INORGANIC CATION TRANSMEMBRANE TRANSPORT%GOBP%GO | NALCN\|TRI | 0.9937 | 0.9071 | 0.5904 |

| | | | | |
|---|---|---|---|---|
| REGULATION OF ACTIN FILAMENT LENGTH%GOBP%GO:0030832 | HAX1\|TWF | -1 | 0.8931 | -0.5665 |
| SMALL MOLECULE BIOSYNTHETIC PROCESS%GOBP%GO:0044283 | ADA\|HAS3 | 0.9955 | 0.9689 | 0.4737 |
| STEROID METABOLIC PROCESS%GOBP%GO:0008202 | CAT\|HSD17 | 0.9874 | 0.9587 | 0.7073 |
| HALLMARK_ADIPOGENESIS%MSIGDB_C2%HALLMARK_ADIPOGEI | ME1\|CAT\|S | 1 | 0.9967 | 0.799 |
| LIPID MODIFICATION%GOBP%GO:0030258 | ST3GAL4\|II | 1 | 0.9976 | 0.7922 |

| | | | | |
|---|---|---|---|---|
| ANATOMICAL STRUCTURE FORMATION INVOLVED IN MORPHOG | PARVA\|GA | -1 | 0.0474 | -0.4795 |
| POSITIVE REGULATION OF CYTOSKELETON ORGANIZATION%GOB | PAK1\|CCL2 | -1 | 0.5862 | -0.6182 |
| ANTIGEN PROCESSING: UBIQUITINATION & PROTEASOME DEGRA | FBXL4\|RCH | 0.5801 | 0.9977 | 0.3354 |

| | | | | |
|---|---|---|---|---|
| HETEROCYCLE BIOSYNTHETIC PROCESS%GOBP%GO:0018130 | CDK9\|CDK[illegible] | 1 | 0.9543 | 0.5238 |
| THE CITRIC ACID (TCA) CYCLE AND RESPIRATORY ELECTRON TRAN | ME1\|MPC[illegible] | 1 | 0.9988 | 0.7113 |
| NUCLEOSIDE MONOPHOSPHATE METABOLIC PROCESS%GOBP%G | AK3\|MSH2 | 1 | 0.9655 | 0.6811 |
| PURINE NUCLEOSIDE BISPHOSPHATE METABOLIC PROCESS%GOB | ACOT7\|MF | 1 | 0.9935 | 0.8479 |

| | | | | |
|---|---|---|---|---|
| PROTEIN LOCALIZATION%REACTOME DATABASE ID RELEASE 66% | CAT\|SLC25 | 1 | 0.9976 | 0.7911 |
| NEGATIVE REGULATION OF ACTIN FILAMENT DEPOLYMERIZATIOI | TWF2\|EPS8 | -0.9885 | -0.4015 | -0.6651 |
| CITRATE METABOLIC PROCESS%GOBP%GO:0006101 | SUCLG1\|FH | 1 | 0.9703 | 0.8337 |
| ERYTHROCYTE DIFFERENTIATION%GOBP%GO:0030218 | TAL1\|SOX6 | 0.9982 | 0.2011 | 0.7504 |
| REGULATION OF MITOCHONDRIAL MEMBRANE POTENTIAL%GOE | BCL2L1\|PIN | 0.8778 | 0.9973 | 0.5018 |

| | | | | |
|---|---|---|---|---|
| REGULATION OF REACTIVE OXYGEN SPECIES METABOLIC PROCES | RAB27A\|CI | 0.8322 | 1 | 0.4433 |
| CELL PROJECTION ORGANIZATION%GOBP%GO:0030030 | PARVA\|GA | -1 | 0.5463 | -0.4639 |
| ALPHA-AMINO ACID CATABOLIC PROCESS%GOBP%GO:1901606 | OTC\|TDO2 | 0.9915 | 0.9853 | 0.6349 |
| PHOSPHOLIPID METABOLIC PROCESS%GOBP%GO:0006644 | INPP5B\|PII | 0.9738 | 0.9789 | 0.6005 |
| ORGANIC ACID TRANSPORT%GOBP%GO:0015849 | PAK1\|KCNJ | 0.9929 | 0.7836 | 0.6911 |

| | | | | |
|---|---|---|---|---|
| NAD METABOLIC PROCESS%GOBP%GO:0019674 | SLC37A2\|L | 1 | 0.908 | 0.8154 |
| CDC42 SIGNALING EVENTS%PATHWAY INTERACTION DATABASE | PAK1\|RAS( | -0.986 | 0.4021 | -0.6224 |
| NEGATIVE REGULATION OF NEURON PROJECTION DEVELOPMEN | SEMA3F\|SI | -0.9848 | 0.744 | -0.5975 |
| RESPONSE TO WOUNDING%GOBP%GO:0009611 | PAK1\|PPAI | -1 | -0.8444 | -0.6158 |

| | | | | |
|---|---|---|---|---|
| CELLULAR RESPONSE TO INTERFERON-GAMMA%GOBP%GO:0071 | STX4A\|STA | -0.9977 | 0.0373 | -0.6556 |
| CELL PART MORPHOGENESIS%GOBP%GO:0032990 | PAK1\|GAT/ | -0.3507 | 0.9977 | -0.2678 |
| SIGNALING BY VEGF%REACTOME DATABASE ID RELEASE 66%194 | PAK1\|PIK3 | -0.9881 | 0.2999 | -0.6216 |
| FATTY ACID METABOLISM%REACTOME%R-HSA-8978868.4 | ACOT7\|AC | 1 | 1 | 0.7694 |
| REGULATION OF CELL SHAPE%GOBP%GO:0008360 | SPRR1B\|M | -0.9976 | -0.6771 | -0.5947 |

| | | | | |
|---|---|---|---|---|
| LOCOMOTION%GOBP%GO:0040011 | PARVA\|GA | -1 | 0.1149 | -0.4732 |
| ANGIOGENESIS%PANTHER PATHWAY%P00005 | FOS\|PAK1\| | -0.9869 | 0.2438 | -0.6283 |
| ANION TRANSPORT%GOBP%GO:0006820 | PAK1\|KCNI | 1 | 0.9042 | 0.6463 |
| REGULATION OF ACTIN POLYMERIZATION OR DEPOLYMERIZATIO | HAX1\|TWF | -1 | 0.906 | -0.5665 |
| NEGATIVE REGULATION OF PROTEIN COMPLEX DISASSEMBLY%G | TNF\|TWF2 | -0.9814 | -0.3007 | -0.6133 |

| | | | | |
|---|---|---|---|---|
| ORGANELLE BIOGENESIS AND MAINTENANCE%REACTOME%R-HS | RAB8A\|GM | 0.9905 | 0.8258 | 0.4921 |
| NUCLEOSIDE TRIPHOSPHATE BIOSYNTHETIC PROCESS%GOBP%G( | AK3\|CTPS2 | 1 | 0.6708 | 0.6423 |
| CARBOHYDRATE DERIVATIVE METABOLIC PROCESS%GOBP%GO:1 | ADA\|ALG2 | 1 | 0.9265 | 0.5596 |
| RHO GTPASES ACTIVATE PKNS%REACTOME%R-HSA-5625740.1 | PAK1\|AR\|F | -0.9931 | 0.0013 | -0.5716 |

| | | | | |
|---|---|---|---|---|
| ACTIN FILAMENT ORGANIZATION%GOBP%GO:0007015 | HSP90B1\|T | -1 | -0.7647 | -0.6624 |
| OXIDATIVE PHOSPHORYLATION%GOBP%GO:0006119 | MSH2\|DLD | 1 | 0.9721 | 0.7385 |
| NEUROMUSCULAR PROCESS%GOBP%GO:0050905 | DBN1\|SCN | -0.9976 | 0.8402 | -0.7009 |
| DICARBOXYLIC ACID METABOLIC PROCESS%GOBP%GO:0043648 | ME1\|DLD\| | 1 | 1 | 0.6471 |
| PROTEIN HOMOOLIGOMERIZATION%GOBP%GO:0051260 | THBS4\|CAT | 1 | 0.7321 | 0.5404 |
| WOUND HEALING%GOBP%GO:0042060 | PAK1\|PPAF | -1 | -0.8804 | -0.6926 |

| | | | | |
|---|---|---|---|---|
| REGULATION OF STEROID METABOLIC PROCESS%GOBP%GO:001! | ADORA2B\| | 0.9982 | 0.965 | 0.7685 |
| METABOLISM OF LIPIDS%REACTOME%R-HSA-556833.5 | PIK3CA\|HE | 1 | 0.9968 | 0.6619 |
| NEGATIVE REGULATION OF CELL PROJECTION ORGANIZATION%G | SEMA3F\|SE | -0.9977 | 0.4937 | -0.5646 |
| SULFUR COMPOUND BIOSYNTHETIC PROCESS%GOBP%GO:00442 | MPC2\|DLD | 0.9965 | 0.8333 | 0.7231 |
| FATTY ACID METABOLIC PROCESS%GOBP%GO:0006631 | CYP2D11\|F | 1 | 1 | 0.7716 |

| | | | | |
|---|---|---|---|---|
| PROTEASOME-MEDIATED UBIQUITIN-DEPENDENT PROTEIN CATA | HSP90B1\|F | 0.7763 | 0.9964 | 0.3752 |
| POSITIVE REGULATION OF ORGANELLE ORGANIZATION%GOBP%( | CDK9\|PAK1 | -1 | 0.2942 | -0.4411 |
| FATTY ACID OXIDATION%GOBP%GO:0019395 | ACADM\|ET | 1 | 1 | 0.7955 |
| CELL ADHESION%GOBP%GO:0007155 | PARVA\|CD | -1 | 0.4908 | -0.5327 |
| LIPID CATABOLIC PROCESS%GOBP%GO:0016042 | HSD17B11 | 1 | 0.9976 | 0.684 |

| | | | | |
|---|---|---|---|---|
| HORMONE-MEDIATED SIGNALING PATHWAY%GOBP%GO:00097! | TSHB\|PPAF | 0.9909 | 0.8491 | 0.6779 |
| REGULATION OF COLD-INDUCED THERMOGENESIS%GOBP%GO:0 | THRA\|PDG | 1 | 0.9755 | 0.8668 |

| EM1_ES..C | EM1_fdr_q | EM1_fdr_q | EM1_Form | EM1_GS_DESCR |
|---|---|---|---|---|
| 0.816 | 0 | 0.0031 | | oxidation-reduction process |
| 0.6842 | 0.0011 | 0.3643 | TRANSME | transmembrane transport |
| 0.8592 | 0 | 0.1268 | FATTY | fatty acid beta-oxidation |
| -0.3148 | 0.0018 | 1 | PLASMA | plasma membrane bounded cell projection assembly |
| 0.8507 | 0 | 0.122 | NUCLEOSI | nucleoside bisphosphate metabolic process |
| 0.8607 | 0.0023 | 0.0943 | ELECTRON | electron transport chain |
| 0.7416 | 0 | 0.1119 | ORGANOP | organophosphate biosynthetic process |
| -0.4671 | 0.0748 | 1 | REGULATI | regulation of protein depolymerization |
| 0.8544 | 0.0052 | 0.1293 | RESPIRAT | respiratory electron transport chain |
| 0.7436 | 0.0192 | 0.5442 | CARBOXYL | carboxylic acid transport |
| 0.7418 | 0 | 0.1316 | COENZYM | coenzyme metabolic process |
| 0.8736 | 0.0168 | 0.1592 | CITRIC | Citric acid cycle (TCA cycle) |
| 0.3964 | 0.0377 | 0.9523 | REGULATI | regulation of cell projection organization |
| -0.4346 | 0.0899 | 1 | CIRCADIA | circadian rhythm |
| 0.5508 | 0.0778 | 0.8141 | BIOLOGIC | biological adhesion |
| 0.7215 | 0 | 0.1285 | PURINE | purine nucleotide metabolic process |
| 0.692 | 4.00E-04 | 0.3136 | PURINE | purine nucleoside triphosphate metabolic process |
| -0.8086 | 0.0373 | 0.5998 | REGULATION OF LEUKOCYTE PROLIFERATION%GOBP%GO:0070663 | regulation of leukocyte proliferation |
| 0.434 | 0.029 | 0.9031 | REGULATION OF CELL MORPHOGENESIS%GOBP%GO:0022604 | regulation of cell morphogenesis |
| 0.7201 | 0.0074 | 0.568 | MITOCHONDRIAL MEMBRANE ORGANIZATION%GOBP%GO:0007006 | mitochondrial membrane organization |

| | | | | |
|---|---|---|---|---|
| 0.6552 | 0.0875 | 0.701 | REGULATION OF CYTOSOLIC CALCIUM ION CONCENTRATION%GOBP%GO:0051480 | regulation of cytosolic calcium ion concentration |
| 0.7999 | 0 | 0.0086 | CELLULAR LIPID METABOLIC PROCESS%GOBP%GO:0044255 | cellular lipid metabolic process |
| -0.4336 | 0.0398 | 1 | EPH-EPHRIN SIGNALING%REACTOME DATABASE ID RELEASE 66%2682334 | EPH-Ephrin signaling |
| 0.4108 | 0.0149 | 0.9903 | CELL-MATRIX ADHESION%GOBP%GO:0007160 | cell-matrix adhesion |
| 0.6108 | 0.0487 | 0.8381 | ERYTHROCYTE HOMEOSTASIS%GOBP%GO:0034101 | erythrocyte homeostasis |

| | | | | |
|---|---|---|---|---|
| -0.7111 | 0.0026 | 0.8336 | MUSCLE CONTRACTION%REACTOME DATABASE ID RELEASE 66%397014 | Muscle contraction |
| 0.8402 | 0.6523 | 0.0736 | PROTEIN HETEROOLIGOMERIZATION%GOBP%GO:0051291 | protein heterooligomerization |
| -0.4236 | 0.048 | 1 | NABA_MATRISOME_ASSOCIATED%MSIGDB_C2%NABA_MATRISOME_ASSOCIATED | NABA_MATRISOME_ASSOCIATED |
| 0.7418 | 0.0378 | 0.4048 | SIGNALING BY NUCLEAR RECEPTORS%REACTOME DATABASE ID RELEASE 66%9006931 | Signaling by Nuclear Receptors |

| | | | | |
|---|---|---|---|---|
| -0.5809 | 0.029 | 0.8636 | RESPONSE TO GROWTH FACTOR%GOBP%GO:0070848 | response to growth factor |
| 0.5227 | 0.0286 | 0.8316 | MUSCLE STRUCTURE DEVELOPMENT%GOBP%GO:0061061 | muscle structure development |
| 0.7236 | 0 | 0.2123 | NUCLEOSIDE PHOSPHATE BIOSYNTHETIC PROCESS%GOBP%GO:1901293 | nucleoside phosphate biosynthetic process |
| 0.7057 | 4.00E-04 | 0.2949 | RIBONUCLEOSIDE TRIPHOSPHATE METABOLIC PROCESS%GOBP%GO:0009199 | ribonucleoside triphosphate metabolic process |
| 0.7222 | 0 | 0.2425 | PURINE NUCLEOSIDE MONOPHOSPHATE METABOLIC PROCESS%GOBP%GO:0009126 | purine nucleoside monophosphate metabolic process |

| | | | | |
|---|---|---|---|---|
| 0.6379 | 0.0045 | 0.7399 | PURINE RIBONUCLEOSIDE TRIPHOSPHATE BIOSYNTHETIC PROCESS %GOBP% GO:0009206 | purine ribonucleoside triphosphate biosynthetic process |
| 0.4678 | 0.057 | 0.8976 | MUSCLE CELL DIFFERENTIATION% GOBP%GO:0042692 | muscle cell differentiation |
| 0.7589 | 0 | 0.1204 | CARBOXYLIC ACID CATABOLIC PROCESS %GOBP% GO:0046395 | carboxylic acid catabolic process |
| 0.5483 | 0.0533 | 0.8688 | DNA PACKAGING%GOBP %GO:0006323 | DNA packaging |
| -0.6357 | 0.0051 | 0.8465 | NABA_ECM_ REGULATORS%MSIGDB_ C2%NABA _ECM_ REGULATORS | NABA_ECM_REGULATORS |

| | | | | |
|---|---|---|---|---|
| 0.8468 | 0 | 0.0891 | FATTY ACID CATABOLIC PROCESS %GOBP% GO:0009062 | fatty acid catabolic process |
| 0.8639 | 0 | 0.0617 | LIPID OXIDATION%GOBP%GO:0034440 | lipid oxidation |
| 0.778 | 0 | 0.1241 | PURINE RIBONUCLEOTIDE BIOSYNTHETIC PROCESS %GOBP% GO:0009152 | purine ribonucleotide biosynthetic process |
| 0.6796 | 0.0052 | 0.3925 | SULFUR COMPOUND METABOLIC PROCESS %GOBP% GO:0006790 | sulfur compound metabolic process |
| 0.8214 | 3.00E-04 | 0.166 | RESPIRATORY ELECTRON TRANSPORT%REACTOME DATABASE ID RELEASE 66%611105 | Respiratory electron transport |

| | | | | |
|---|---|---|---|---|
| 0.7301 | 0.0381 | 0.3498 | CATION TRANSMEMBRANE TRANSPORT%GOBP%GO:0098655 | cation transmembrane transport |
| 0.8109 | 0 | 0.0031 | HALLMARK_ OXIDATIVE_ PHOSPHORYLATION%MSIGDB_ C2%HALLMARK_ OXIDATIVE_ PHOSPHORYLATION | HALLMARK_OXIDATIVE_PHOSPHORYLATION |
| 0.6194 | 0.0913 | 0.7205 | NEGATIVE REGULATION OF CELLULAR COMPONENT MOVEMENT%GOBP%GO:0051271 | negative regulation of cellular component movement |
| 0.7966 | 0.003 | 0.4962 | CRISTAE FORMATION%REACTOME DATABASE ID RELEASE 66%8949613 | Cristae formation |

| | | | | |
|---|---|---|---|---|
| 0.3999 | 0.0382 | 0.9575 | TISSUE MORPHOGENESIS%GOBP%GO:0048729 | tissue morphogenesis |
| 0.7802 | 0 | 0.0487 | MONOCARBOXYLIC ACID METABOLIC PROCESS%GOBP%GO:0032787 | monocarboxylic acid metabolic process |
| 0.7385 | 0 | 0.0645 | ORGANOPHOSPHATE METABOLIC PROCESS%GOBP%GO:0019637 | organophosphate metabolic process |
| 0.8119 | 0.0012 | 0.4228 | NADH METABOLIC PROCESS%GOBP%GO:0006734 | NADH metabolic process |
| 0.8262 | 0.1035 | 0.0906 | HEDGEHOG LIGAND BIOGENESIS%REACTOME DATABASE ID RELEASE 66%5358346 | Hedgehog ligand biogenesis |

| | | | | |
|---|---|---|---|---|
| 0.847 | 0 | 0.1325 | MITOCHONDRIAL PROTEIN IMPORT%REACTOME%R-HSA-1268020.3 | Mitochondrial protein import |
| 0.8372 | 0.0478 | 0.2569 | POSITIVE REGULATION OF LIPID METABOLIC PROCESS%GOBP%GO:0045834 | positive regulation of lipid metabolic process |
| 0.7127 | 0.0065 | 0.4277 | INORGANIC ION TRANSMEMBRANE TRANSPORT%GOBP%GO:0098660 | inorganic ion transmembrane transport |
| -0.813 | 0.0291 | 0.7361 | REGULATION OF LYMPHOCYTE PROLIFERATION%GOBP%GO:0050670 | regulation of lymphocyte proliferation |
| -0.5675 | 0.038 | 0.7617 | NABA_MATRISOME%MSIGDB_C2%NABA_MATRISOME | NABA_MATRISOME |

| | | | | |
|---|---|---|---|---|
| -0.5223 | 0.091 | 0.9553 | CILIUM ASSEMBLY %GOBP% GO:0060271 | cilium assembly |
| 0.8534 | 0 | 0.1301 | THIOESTER METABOLIC PROCESS %GOBP% GO:0035383 | thioester metabolic process |
| 0.7405 | 0.0406 | 0.6476 | SIG_REGULATION_OF_ THE_ACTIN_ CYTOSKELETON_BY_ RHO_ GTPASES%MSIGDB_ C2%SIG_ REGULATION_OF_THE_ ACTIN_ CYTOSKELETON_BY_ RHO_GTPASES | SIG_REGULATION_OF_THE_ACTIN_CYTOSKELETON_BY_RHO_GT |
| 0.8201 | 0.0233 | 0.1752 | PROTEIN LOCALIZATION TO MITOCHONDRION%GOBP%GO:0070585 | protein localization to mitochondrion |

| | | | | |
|---|---|---|---|---|
| 0.8181 | 0.0816 | 0.1661 | TP53 REGULATES METABOLIC GENES%REACTOME DATABASE ID RELEASE 66%5628897 | TP53 Regulates Metabolic Genes |
| 0.4137 | 0.0235 | 0.9067 | MOVEMENT OF CELL OR SUBCELLULAR COMPONENT%GOBP%GO:0006928 | movement of cell or subcellular component |
| 0.6343 | 0.0062 | 0.7321 | RIBONUCLEOSIDE TRIPHOSPHATE BIOSYNTHETIC PROCESS%GOBP%GO:0009201 | ribonucleoside triphosphate biosynthetic process |
| 0.7675 | 0.0016 | 0.0716 | MITOCHONDRION ORGANIZATION%GOBP%GO:0007005 | mitochondrion organization |

| | | | | |
|---|---|---|---|---|
| 0.8032 | 0.3386 | 0.0932 | INTERLEUKIN-1 SIGNALING%REACTOME%R-HSA-9020702.1 | Interleukin-1 signaling |
| 0.3321 | 0.071 | 0.9969 | ESTABLISHMENT OR MAINTENANCE OF CELL POLARITY%GOBP%GO:0007163 | establishment or maintenance of cell polarity |
| 0.7852 | 0.0024 | 0.1672 | ANTIBIOTIC METABOLIC PROCESS%GOBP%GO:0016999 | antibiotic metabolic process |
| 0.8358 | 0 | 0.1577 | AEROBIC RESPIRATION%GOBP%GO:0009060 | aerobic respiration |
| 0.6432 | 0.0016 | 0.7466 | ATP BIOSYNTHETIC PROCESS%GOBP%GO:0006754 | ATP biosynthetic process |

| | | | | |
|---|---|---|---|---|
| 0.8203 | 0.1807 | 0.0247 | ANTIGEN PROCESSING-CROSS PRESENTATION%REACTOME DATABASE ID RELEASE 66%1236975 | Antigen processing-Cross presentation |
| 0.6037 | 0.0147 | 0.6729 | REGULATION OF ACTIN FILAMENT-BASED PROCESS%GOBP%GO:0032970 | regulation of actin filament-based process |
| 0.6079 | 0.0823 | 0.6234 | CELL MORPHOGENESIS%GOBP%GO:0000902 | cell morphogenesis |
| 0.6797 | 0.0014 | 0.2853 | CARBOHYDRATE DERIVATIVE BIOSYNTHETIC PROCESS%GOBP%GO:1901137 | carbohydrate derivative biosynthetic process |

| | | | | |
|---|---|---|---|---|
| -0.6925 | 0.0708 | 0.9657 | REGULATION OF GENE EXPRESSION, EPIGENETIC%GOBP%GO:0040029 | regulation of gene expression, epigenetic |
| 0.8147 | 0.0019 | 0.2497 | TRICARBOXYLIC ACID METABOLIC PROCESS%GOBP%GO:0072350 | tricarboxylic acid metabolic process |
| 0.3107 | 0.0017 | 1 | ACTIN FILAMENT BUNDLE ASSEMBLY%GOBP%GO:0051017 | actin filament bundle assembly |
| 0.5884 | 0.079 | 0.8613 | REGULATION OF EXTENT OF CELL GROWTH%GOBP%GO:0061387 | regulation of extent of cell growth |
| 0.8405 | 0.0341 | 0.155 | ESTABLISHMENT OF PROTEIN LOCALIZATION TO MITOCHONDRION%GOBP%GO:0072655 | establishment of protein localization to mitochondrion |

| | | | | |
|---|---|---|---|---|
| 0.5002 | 0.0414 | 0.8994 | CELL JUNCTION ASSEMBLY %GOBP% GO:0034329 | cell junction assembly |
| 0.5864 | 0.0803 | 0.858 | PID_CDC42_ PATHWAY %MSIGDB_ C2%PID_CDC42_ PATHWAY | PID_CDC42_PATHWAY |
| 0.4772 | 0.0995 | 0.9034 | MHC CLASS II ANTIGEN PRESENTATION%REACTOME% R-HSA-2132295.3 | MHC class II antigen presentation |
| 0.6864 | 0.0064 | 0.3304 | SMALL MOLECULE CATABOLIC PROCESS %GOBP% GO:0044282 | small molecule catabolic process |
| 0.7589 | 0 | 0.1316 | ORGANIC ACID CATABOLIC PROCESS %GOBP% GO:0016054 | organic acid catabolic process |

| | | | | |
|---|---|---|---|---|
| -0.322 | 0 | 0.9964 | CYTOSKELETON ORGANIZATION%GOBP%GO:0007010 | cytoskeleton organization |
| 0.5741 | 0.0153 | 0.6946 | REGULATION OF ANATOMICAL STRUCTURE SIZE%GOBP%GO:0090066 | regulation of anatomical structure size |
| 0.6405 | 0.0496 | 0.3936 | PROTEIN COMPLEX OLIGOMERIZATION%GOBP%GO:0051259 | protein complex oligomerization |
| 0.8344 | 3.00E-04 | 0.0917 | GLYCEROLIPID METABOLIC PROCESS%GOBP%GO:0046486 | glycerolipid metabolic process |
| -0.3107 | 0.0019 | 1 | CELL PROJECTION ASSEMBLY%GOBP%GO:0030031 | cell projection assembly |

| | | | | |
|---|---|---|---|---|
| 0.6387 | 0.002 | 0.4243 | AROMATIC COMPOUND BIOSYNTHETIC PROCESS %GOBP% GO:0019438 | aromatic compound biosynthetic process |
| 0.8907 | 0.0096 | 0.0906 | REGULATION OF LIPID BIOSYNTHETIC PROCESS %GOBP% GO:0046890 | regulation of lipid biosynthetic process |
| 0.6711 | 2.00E-04 | 0.5096 | REGULATION OF SMALL MOLECULE METABOLIC PROCESS %GOBP% GO:0062012 | regulation of small molecule metabolic process |
| 0.8372 | 0 | 0.1302 | MONOCARBOXYLIC ACID CATABOLIC PROCESS %GOBP% GO:0072329 | monocarboxylic acid catabolic process |

| | | | | |
|---|---|---|---|---|
| 0.7222 | 0 | 0.2328 | PURINE RIBONUCLEOSIDE MONOPHOSPHATE METABOLIC PROCESS%GOBP%GO:0009167 | purine ribonucleoside monophosphate metabolic process |
| -0.3696 | 0.0215 | 1 | HEART MORPHOGENESIS%GOBP%GO:0003007 | heart morphogenesis |
| 0.7138 | 0 | 0.1604 | PURINE-CONTAINING COMPOUND METABOLIC PROCESS%GOBP%GO:0072521 | purine-containing compound metabolic process |
| -0.248 | 0.0282 | 1 | ACTIN FILAMENT-BASED MOVEMENT%GOBP%GO:0030048 | actin filament-based movement |
| 0.7318 | 0 | 0.0918 | NUCLEOSIDE PHOSPHATE METABOLIC PROCESS%GOBP%GO:0006753 | nucleoside phosphate metabolic process |

| | | | | |
|---|---|---|---|---|
| 0.3964 | 0.0381 | 0.9494 | REGULATION OF PLASMA MEMBRANE BOUNDED CELL PROJECTION ORGANIZATION%GOBP%GO:0120035 | regulation of plasma membrane bounded cell projection organiz |
| 0.6399 | 0.0036 | 0.7982 | PYRUVATE METABOLIC PROCESS%GOBP%GO:0006090 | pyruvate metabolic process |
| 0.5733 | 0.094 | 0.8573 | REACTIVE OXYGEN SPECIES METABOLIC PROCESS%GOBP%GO:0072593 | reactive oxygen species metabolic process |
| 0.7055 | 0.0017 | 0.4876 | RIBONUCLEOSIDE MONOPHOSPHATE BIOSYNTHETIC PROCESS%GOBP%GO:0009156 | ribonucleoside monophosphate biosynthetic process |

| | | | | |
|---|---|---|---|---|
| 0.6587 | 0.0158 | 0.7668 | MONOVALENT INORGANIC CATION TRANSPORT%GOBP%GO:0015672 | monovalent inorganic cation transport |
| -0.813 | 0.0343 | 0.6507 | REGULATION OF MONONUCLEAR CELL PROLIFERATION%GOBP%GO:0032944 | regulation of mononuclear cell proliferation |
| 0.805 | 0.1801 | 0.0926 | ABC TRANSPORTER DISORDERS%REACTOME DATABASE ID RELEASE 66%5619084 | ABC transporter disorders |
| 0.4428 | 0.0637 | 0.9578 | COLLAGEN FORMATION%REACTOME%R-HSA-1474290.1 | Collagen formation |
| 0.6476 | 0.0833 | 0.7201 | MUSCLE ORGAN DEVELOPMENT%GOBP%GO:0007517 | muscle organ development |

| | | | | |
|---|---|---|---|---|
| 0.729 | 0.0015 | 0.6585 | AMMONIUM ION METABOLIC PROCESS%GOBP%GO:0097164 | ammonium ion metabolic process |
| 0.3309 | 0 | 0.9895 | ACTIN CYTOSKELETON ORGANIZATION%GOBP%GO:0030036 | actin cytoskeleton organization |
| 0.6025 | 0.0408 | 0.6859 | REGULATION OF SUPRAMOLECULAR FIBER ORGANIZATION%GOBP%GO:1902903 | regulation of supramolecular fiber organization |
| 0.6556 | 0.0021 | 0.3362 | NUCLEOBASE-CONTAINING COMPOUND BIOSYNTHETIC PROCESS%GOBP%GO:0034654 | nucleobase-containing compound biosynthetic process |

| | | | | |
|---|---|---|---|---|
| 0.7069 | 0.0017 | 0.5291 | PURINE NUCLEOSIDE MONOPHOSPHATE BIOSYNTHETIC PROCESS %GOBP% GO:0009127 | purine nucleoside monophosphate biosynthetic process |
| 0.781 | 0.0141 | 0.167 | PROTEIN TETRAMERIZATION %GOBP% GO:0051262 | protein tetramerization |
| 0.6471 | 1.00E-04 | 0.7485 | HALLMARK_ PEROXISOME%MSIGDB_ C2%HALLMARK_ PEROXISOME | HALLMARK_PEROXISOME |
| 0.3107 | 0.0018 | 1 | ACTIN FILAMENT BUNDLE ORGANIZATION%GOBP%GO:0061572 | actin filament bundle organization |
| 0.4277 | 0.08 | 0.9667 | REGULATION OF AXONOGENESIS%GOBP%GO:0050770 | regulation of axonogenesis |

| | | | | |
|---|---|---|---|---|
| -0.3286 | 0.0064 | 1 | SUPRAMOLECULAR FIBER ORGANIZATION%GOBP%GO:0097435 | supramolecular fiber organization |
| 0.6482 | 0.0613 | 0.8039 | REGULATION OF TUMOR NECROSIS FACTOR SUPERFAMILY CYTOKINE PRODUCTION%GOBP%GO:1903555 | regulation of tumor necrosis factor superfamily cytokine product |
| -0.8265 | 0.0335 | 0.5764 | REGULATION OF T CELL PROLIFERATION%GOBP%GO:0042129 | regulation of T cell proliferation |
| 0.7213 | 0 | 0.2494 | RIBONUCLEOSIDE MONOPHOSPHATE METABOLIC PROCESS%GOBP%GO:0009161 | ribonucleoside monophosphate metabolic process |

| | | | | |
|---|---|---|---|---|
| -0.5201 | 0.0557 | 0.977 | REGULATION OF ACTIN FILAMENT DEPOLYMERIZATION%GOBP%GO:0030834 | regulation of actin filament depolymerization |
| 0.8343 | 3.00E-04 | 0.2539 | MITOCHONDRIAL FATTY ACID BETA-OXIDATION%REACTOME%R-HSA-77289.5 | Mitochondrial Fatty Acid Beta-Oxidation |
| 0.8291 | 4.00E-04 | 0.0705 | REGULATION OF LIPID METABOLIC PROCESS%GOBP%GO:0019216 | regulation of lipid metabolic process |
| 0.3812 | 0.0306 | 0.9963 | RESPONSE TO INTERFERON-GAMMA%GOBP%GO:0034341 | response to interferon-gamma |
| 0.6648 | 0.0952 | 0.5565 | PID_PDGFRB_ PATHWAY%MSIGDB_C2%PID_PDGFRB_ PATHWAY | PID_PDGFRB_PATHWAY |

| | | | | |
|---|---|---|---|---|
| 0.6962 | 0 | 0.1584 | DRUG METABOLIC PROCESS %GOBP% GO:0017144 | drug metabolic process |
| -0.6373 | 0.0454 | 0.7159 | REGULATION OF PROTEIN COMPLEX DISASSEMBLY%GOBP%GO:0043244 | regulation of protein complex disassembly |
| 0.7683 | 0 | 0.0679 | PURINE RIBONUCLEOTIDE METABOLIC PROCESS %GOBP% GO:0009150 | purine ribonucleotide metabolic process |
| 0.6482 | 0.0619 | 0.803 | REGULATION OF TUMOR NECROSIS FACTOR PRODUCTION%GOBP%GO:0032680 | regulation of tumor necrosis factor production |
| 0.7401 | 0.035 | 0.181 | REGULATION OF PROTEIN POLYMERIZATION% GOBP%GO:0032271 | regulation of protein polymerization |

| | | | | |
|---|---|---|---|---|
| 0.6834 | 0.0115 | 0.4635 | REGULATION OF ACTIN FILAMENT ORGANIZATION%GOBP%GO:0110053 | regulation of actin filament organization |
| 0.4852 | 0.0907 | 0.948 | PID_CXCR4_PATHWAY%MSIGDB_C2%PID_CXCR4_PATHWAY | PID_CXCR4_PATHWAY |
| 0.6688 | 0.0463 | 0.8002 | HALLMARK_BILE_ACID_METABOLISM%MSIGDB_C2%HALLMARK_BILE_ACID_METABOLISM | HALLMARK_BILE_ACID_METABOLISM |
| 0.7921 | 0.0285 | 0.3631 | POSITIVE REGULATION OF ACTIN FILAMENT POLYMERIZATION%GOBP%GO:0030838 | positive regulation of actin filament polymerization |

| | | | | |
|---|---|---|---|---|
| 0.5044 | 0.0408 | 0.8774 | STRIATED MUSCLE CELL DIFFERENTIATION%GOBP%GO:0051146 | striated muscle cell differentiation |
| 0.5458 | 0.0855 | 0.897 | VEGFA-VEGFR2 PATHWAY%REACTOME DATABASE ID RELEASE 66%4420097 | VEGFA-VEGFR2 Pathway |
| -0.6445 | 0.0901 | 0.9063 | NEGATIVE REGULATION OF LEUKOCYTE CELL-CELL ADHESION%GOBP%GO:1903038 | negative regulation of leukocyte cell-cell adhesion |
| 0.3401 | 0.0277 | 1 | SEMAPHORIN INTERACTIONS%REACTOME DATABASE ID RELEASE 66%373755 | Semaphorin interactions |

| | | | | |
|---|---|---|---|---|
| -0.345 | 0.017 | 0.9926 | LOCALIZATION OF CELL%GOBP%GO:0051674 | localization of cell |
| 0.5971 | 0.0208 | 0.6671 | ION TRANSPORT%GOBP%GO:0006811 | ion transport |
| 0.8078 | 5.00E-04 | 0.2064 | MITOCHONDRIAL BIOGENESIS%REACTOME%R-HSA-1592230.2 | Mitochondrial biogenesis |
| -0.456 | 7.00E-04 | 1 | MUSCLE CELL DEVELOPMENT%GOBP%GO:0055001 | muscle cell development |
| 0.6409 | 0.0287 | 0.687 | INFLAMMATION MEDIATED BY CHEMOKINE AND CYTOKINE SIGNALING PATHWAY%PANTHER PATHWAY%P00031 | Inflammation mediated by chemokine and cytokine signaling pat |

| | | | | |
|---|---|---|---|---|
| -0.5832 | 0.03 | 0.782 | CELLULAR RESPONSE TO GROWTH FACTOR STIMULUS %GOBP% GO:0071363 | cellular response to growth factor stimulus |
| 0.4767 | 9.00E-04 | 0.871 | SIGNALING BY RHO GTPASES% REACTOME%R-HSA-194315.4 | Signaling by Rho GTPases |
| 0.7876 | 0 | 0.0916 | RIBONUCLEOTIDE BIOSYNTHETIC PROCESS %GOBP% GO:0009260 | ribonucleotide biosynthetic process |
| 0.8436 | 0.0078 | 0.2707 | REGULATION OF CELLULAR KETONE METABOLIC PROCESS %GOBP% GO:0010565 | regulation of cellular ketone metabolic process |

| | | | | |
|---|---|---|---|---|
| 0.6697 | 0.0043 | 0.5694 | POSITIVE REGULATION OF SUPRAMOLECULAR FIBER ORGANIZATION%GOBP%GO:1902905 | positive regulation of supramolecular fiber organization |
| 0.8186 | 0 | 0.125 | RESPIRATORY ELECTRON TRANSPORT, ATP SYNTHESIS BY CHEMIOSMOTIC COUPLING, AND HEAT PRODUCTION BY UNCOUPLING PROTEINS.%REACTOME%R-HSA-163200.1 | Respiratory electron transport, ATP synthesis by chemiosmotic c |
| 0.6946 | 0 | 0.152 | CARBOXYLIC ACID METABOLIC PROCESS%GOBP%GO:0019752 | carboxylic acid metabolic process |

| | | | | |
|---|---|---|---|---|
| 0.5094 | 0.0195 | 0.8026 | CELL DEVELOPMENT%GOBP%GO:0048468 | cell development |
| 0.5475 | 0.017 | 0.7639 | REGULATION OF CYTOSKELETON ORGANIZATION%GOBP%GO:0051493 | regulation of cytoskeleton organization |
| 0.4976 | 0.0411 | 0.8914 | GOLGI-TO-ER RETROGRADE TRANSPORT%REACTOME%R-HSA-8856688.1 | Golgi-to-ER retrograde transport |
| 0.7854 | 0.0087 | 0.2896 | PROTEIN HOMOTETRAMERIZATION%GOBP%GO:0051289 | protein homotetramerization |
| 0.6827 | 0 | 0.1595 | OXOACID METABOLIC PROCESS%GOBP%GO:0043436 | oxoacid metabolic process |

| | | | | |
|---|---|---|---|---|
| 0.7262 | 0.0112 | 0.6183 | METABOLISM OF STEROIDS %REACTOME DATABASE ID RELEASE 66%8957322 | Metabolism of steroids |
| 0.727 | 0 | 0.1203 | COFACTOR METABOLIC PROCESS %GOBP%GO:0051186 | cofactor metabolic process |
| 0.462 | 0.0991 | 0.9543 | AMEBOIDAL-TYPE CELL MIGRATION%GOBP%GO:0001667 | ameboidal-type cell migration |
| 0.5127 | 0.0988 | 0.9392 | PID_RAC1_PATHWAY%MSIGDB_C2%PID_RAC1_PATHWAY | PID_RAC1_PATHWAY |

| | | | | |
|---|---|---|---|---|
| | | | PDGFR-BETA SIGNALING PATHWAY%PATHWAY INTERACTION DATABASE NCI-NATURE CURATED DATA%PDGFR-BETA SIGNALING PATHWAY | |
| 0.6648 | 0.0904 | 0.5555 | | PDGFR-beta signaling pathway |
| -0.4199 | 0.001 | 1 | STRIATED MUSCLE CELL DEVELOPMENT%GOBP%GO:0055002 | striated muscle cell development |
| 0.4851 | 0.0335 | 0.8397 | PLASMA MEMBRANE BOUNDED CELL PROJECTION ORGANIZATION%GOBP%GO:0120036 | plasma membrane bounded cell projection organization |

| | | | | |
|---|---|---|---|---|
| 0.8423 | 0.0878 | 0.2116 | CARBOHYDRATE BIOSYNTHETIC PROCESS %GOBP% GO:0016051 | carbohydrate biosynthetic process |
| 0.8507 | 0 | 0.1212 | RIBONUCLEOSIDE BISPHOSPHATE METABOLIC PROCESS %GOBP% GO:0033875 | ribonucleoside bisphosphate metabolic process |
| 0.7123 | 0 | 0.4034 | PYRIDINE NUCLEOTIDE METABOLIC PROCESS %GOBP% GO:0019362 | pyridine nucleotide metabolic process |
| 0.693 | 0 | 0.4817 | OXIDOREDUCTION COENZYME METABOLIC PROCESS %GOBP% GO:0006733 | oxidoreduction coenzyme metabolic process |
| 0.7945 | 0.8092 | 0.0765 | RESPONSE TO TOXIC SUBSTANCE%GOBP% GO:0009636 | response to toxic substance |

| | | | | |
|---|---|---|---|---|
| 0.3807 | 0.0711 | 0.9569 | CELL CYCLE PROCESS %GOBP% GO:0022402 | cell cycle process |
| 0.7901 | 0.0012 | 0.1892 | LIPID BIOSYNTHETIC PROCESS %GOBP% GO:0008610 | lipid biosynthetic process |
| 0.805 | 0.1747 | 0.0979 | DEFECTIVE CFTR CAUSES CYSTIC FIBROSIS% REACTOME DATABASE ID RELEASE 66%5678895 | Defective CFTR causes cystic fibrosis |
| 0.4624 | 0.065 | 0.8951 | MORPHOGENESIS OF AN EPITHELIUM%GOBP %GO:0002009 | morphogenesis of an epithelium |
| 0.7243 | 0 | 0.1762 | NUCLEOTIDE BIOSYNTHETIC PROCESS %GOBP% GO:0009165 | nucleotide biosynthetic process |

| | | | | |
|---|---|---|---|---|
| 0.7331 | 0 | 0.1149 | RIBOSE PHOSPHATE METABOLIC PROCESS %GOBP% GO:0019693 | ribose phosphate metabolic process |
| 0.5125 | 0.0554 | 0.8297 | HEAD DEVELOPMENT%GOBP%GO:0060322 | head development |
| 0.6463 | 4.00E-04 | 0.6233 | COFACTOR BIOSYNTHETIC PROCESS %GOBP% GO:0051188 | cofactor biosynthetic process |
| 0.7123 | 0 | 0.4233 | NICOTINAMIDE NUCLEOTIDE METABOLIC PROCESS %GOBP% GO:0046496 | nicotinamide nucleotide metabolic process |
| 0.8319 | 0.0293 | 0.2646 | PROTEIN TARGETING TO MITOCHONDRION%GOBP%GO:0006626 | protein targeting to mitochondrion |

| | | | | |
|---|---|---|---|---|
| 0.5312 | 0.0378 | 0.8602 | ER TO GOLGI ANTEROGRADE TRANSPORT%REACTOME DATABASE ID RELEASE 66%199977 | ER to Golgi Anterograde Transport |
| 0.6839 | 0.0012 | 0.5434 | COENZYME BIOSYNTHETIC PROCESS%GOBP%GO:0009108 | coenzyme biosynthetic process |
| 0.3904 | 0.0225 | 0.9889 | CELL-SUBSTRATE ADHESION%GOBP%GO:0031589 | cell-substrate adhesion |
| 0.7689 | 0.0291 | 0.3462 | POSITIVE REGULATION OF PROTEIN POLYMERIZATION%GOBP%GO:0032273 | positive regulation of protein polymerization |
| 0.7166 | 0.0169 | 0.5795 | ORGANIC ANION TRANSPORT%GOBP%GO:0015711 | organic anion transport |

| | | | | |
|---|---|---|---|---|
| -0.4754 | 0.0175 | 1 | HALLMARK_ COAGULATION%MSIGDB_ C2%HALLMARK_ COAGULATION | HALLMARK_COAGULATION |
| 0.8395 | 0.0937 | 0.1728 | CELLULAR AMINO ACID CATABOLIC PROCESS %GOBP% GO:0009063 | cellular amino acid catabolic process |
| 0.7745 | 0 | 0.0235 | RIBONUCLEOTIDE METABOLIC PROCESS %GOBP% GO:0009259 | ribonucleotide metabolic process |
| -0.352 | 0.0149 | 0.9993 | CELL MIGRATION%GOBP %GO:0016477 | cell migration |
| 0.6795 | 0.0135 | 0.4563 | ION TRANSMEMBRANE TRANSPORT%GOBP %GO:0034220 | ion transmembrane transport |

| | | | | |
|---|---|---|---|---|
| 0.6827 | 0.0881 | 0.6846 | MONOCARBOXYLIC ACID BIOSYNTHETIC PROCESS %GOBP% GO:0072330 | monocarboxylic acid biosynthetic process |
| 0.5273 | 0.0262 | 0.8666 | COPI-MEDIATED ANTEROGRADE TRANSPORT%REACTOME DATABASE ID RELEASE 66%6807878 | COPI-mediated anterograde transport |
| 0.5919 | 0.0649 | 0.7623 | RESPONSE TO BACTERIUM%GOBP%GO:0009617 | response to bacterium |
| 0.8613 | 2.00E-04 | 0.1596 | MITOCHONDRIAL TRANSMEMBRANE TRANSPORT%GOBP%GO:1990542 | mitochondrial transmembrane transport |
| 0.8479 | 0.0127 | 0.1608 | GLUCOSE METABOLIC PROCESS %GOBP% GO:0006006 | glucose metabolic process |

| | | | | |
|---|---|---|---|---|
| 0.719 | 0 | 0.2947 | PURINE-CONTAINING COMPOUND BIOSYNTHETIC PROCESS%GOBP%GO:0072522 | purine-containing compound biosynthetic process |
| 0.6057 | 0.0807 | 0.549 | AMINO ACID AND DERIVATIVE METABOLISM%REACTOME%R-HSA-71291.4 | Amino acid and derivative metabolism |
| 0.8301 | 0 | 0.0042 | HALLMARK_FATTY_ACID_METABOLISM%MSIGDB_C2%HALLMARK_FATTY_ACID_METABOLISM | HALLMARK_FATTY_ACID_METABOLISM |

| | | | | |
|---|---|---|---|---|
| 0.7141 | 0 | 0.1246 | NUCLEOBASE-CONTAINING SMALL MOLECULE METABOLIC PROCESS%GOBP%GO:0055086 | nucleobase-containing small molecule metabolic process |
| 0.6283 | 0.0344 | 0.7389 | SYNAPSE ORGANIZATION%GOBP%GO:0050808 | synapse organization |
| 0.6935 | 0 | 0.1327 | ORGANIC ACID METABOLIC PROCESS%GOBP%GO:0006082 | organic acid metabolic process |
| 0.5906 | 0.015 | 0.6865 | REGULATION OF CELLULAR COMPONENT SIZE%GOBP%GO:0032535 | regulation of cellular component size |

| | | | | |
|---|---|---|---|---|
| 0.7832 | 9.00E-04 | 0.2646 | PYRUVATE METABOLISM AND CITRIC ACID (TCA) CYCLE%REACTOME%R-HSA-71406.1 | Pyruvate metabolism and Citric Acid (TCA) cycle |
| 0.8392 | 0.0031 | 0.1831 | ATP SYNTHESIS COUPLED ELECTRON TRANSPORT%GOBP%GO:0042773 | ATP synthesis coupled electron transport |
| 0.7571 | 1.00E-04 | 0.1824 | MITOCHONDRIAL TRANSPORT%GOBP%GO:0006839 | mitochondrial transport |
| 0.824 | 0 | 0.0667 | CELLULAR RESPIRATION%GOBP%GO:0045333 | cellular respiration |
| 0.7054 | 0 | 0.4209 | PYRIDINE-CONTAINING COMPOUND METABOLIC PROCESS%GOBP%GO:0072524 | pyridine-containing compound metabolic process |

| | | | | |
|---|---|---|---|---|
| 0.7197 | 0 | 0.2882 | PURINE NUCLEOTIDE BIOSYNTHETIC PROCESS %GOBP%GO:0006164 | purine nucleotide biosynthetic process |
| 0.5323 | 0.0019 | 0.8235 | RHO GTPASE EFFECTORS%REACTOME DATABASE ID RELEASE 66%195258 | RHO GTPase Effectors |
| 0.6911 | 0 | 0.4284 | ATP METABOLIC PROCESS %GOBP%GO:0046034 | ATP metabolic process |
| 0.7018 | 0.0051 | 0.4205 | INTEGRIN SIGNALLING PATHWAY %PANTHER PATHWAY %P00034 | Integrin signalling pathway |
| 0.8355 | 0 | 0.0138 | ENERGY DERIVATION BY OXIDATION OF ORGANIC COMPOUNDS%GOBP%GO:0015980 | energy derivation by oxidation of organic compounds |

| | | | | |
|---|---|---|---|---|
| 0.7318 | 0 | 0.0842 | NUCLEOTIDE METABOLIC PROCESS %GOBP% GO:0009117 | nucleotide metabolic process |
| 0.7069 | 0.0013 | 0.5339 | PURINE RIBONUCLEOSIDE MONOPHOSPHATE BIOSYNTHETIC PROCESS %GOBP% GO:0009168 | purine ribonucleoside monophosphate biosynthetic process |
| 0.316 | 0 | 0.9966 | ACTIN FILAMENT-BASED PROCESS %GOBP% GO:0030029 | actin filament-based process |
| 0.5322 | 0.0504 | 0.898 | CHROMATIN ASSEMBLY %GOBP% GO:0031497 | chromatin assembly |
| 0.4883 | 0.019 | 0.8985 | HALLMARK_APICAL_JUNCTION %MSIGDB_C2%HALLMARK_APICAL_JUNCTION | HALLMARK_APICAL_JUNCTION |

| | | | | |
|---|---|---|---|---|
| 0.8453 | 0.0026 | 0.2486 | COFACTOR CATABOLIC PROCESS%GOBP%GO:0051187 | cofactor catabolic process |
| -0.6849 | 0.0019 | 0.7979 | ACTOMYOSIN STRUCTURE ORGANIZATION%GOBP%GO:0031032 | actomyosin structure organization |
| 0.7825 | 0.0019 | 0.0513 | HALLMARK_MTORC1_SIGNALING%MSIGDB_C2%HALLMARK_MTORC1_SIGNALING | HALLMARK_MTORC1_SIGNALING |
| 0.7332 | 0 | 0.1923 | RIBOSE PHOSPHATE BIOSYNTHETIC PROCESS%GOBP%GO:0046390 | ribose phosphate biosynthetic process |

| | | | | |
|---|---|---|---|---|
| 0.7717 | 0 | 0.0637 | GENERATION OF PRECURSOR METABOLITES AND ENERGY%GOBP%GO:0006091 | generation of precursor metabolites and energy |
| 0.7852 | 3.00E-04 | 0.1231 | CELLULAR LIPID CATABOLIC PROCESS%GOBP%GO:0044242 | cellular lipid catabolic process |
| -0.7429 | 0.0011 | 0.7214 | CELLULAR COMPONENT ASSEMBLY INVOLVED IN MORPHOGENESIS%GOBP%GO:0010927 | cellular component assembly involved in morphogenesis |
| -0.4353 | 0.082 | 1 | REGULATION OF CELL PROJECTION ASSEMBLY%GOBP%GO:0060491 | regulation of cell projection assembly |

| | | | | |
|---|---|---|---|---|
| 0.7055 | 0.0014 | 0.5147 | NUCLEOSIDE MONOPHOSPHATE BIOSYNTHETIC PROCESS %GOBP% GO:0009124 | nucleoside monophosphate biosynthetic process |
| 0.7799 | 0 | 0.0144 | LIPID METABOLIC PROCESS %GOBP% GO:0006629 | lipid metabolic process |
| 0.8392 | 0.004 | 0.2286 | MITOCHONDRIAL ATP SYNTHESIS COUPLED ELECTRON TRANSPORT%GOBP %GO:0042775 | mitochondrial ATP synthesis coupled electron transport |
| -0.2207 | 8.00E-04 | 1 | HALLMARK_MITOTIC_ SPINDLE%MSIGDB_ C2%HALLMARK_ MITOTIC_ SPINDLE | HALLMARK_MITOTIC_SPINDLE |

| | | | | |
|---|---|---|---|---|
| 0.5083 | 0.0062 | 0.8976 | CYTOSKELETAL REGULATION BY RHO GTPASE%PANTHER PATHWAY%P00016 | Cytoskeletal regulation by Rho GTPase |
| -0.4353 | 0.0959 | 1 | REGULATION OF PLASMA MEMBRANE BOUNDED CELL PROJECTION ASSEMBLY%GOBP%GO:0120032 | regulation of plasma membrane bounded cell projection assemb |
| 0.8501 | 0.0076 | 0.2759 | PROTON TRANSMEMBRANE TRANSPORT%GOBP%GO:1902600 | proton transmembrane transport |
| 0.6499 | 0.0023 | 0.3364 | ORGANIC CYCLIC COMPOUND BIOSYNTHETIC PROCESS%GOBP%GO:1901362 | organic cyclic compound biosynthetic process |

| | | | | |
|---|---|---|---|---|
| 0.7448 | 0.0306 | 0.2389 | REGULATION OF ACTIN FILAMENT POLYMERIZATION%GOBP%GO:0030833 | regulation of actin filament polymerization |
| -0.3474 | 0.0573 | 0.9965 | ANIMAL ORGAN MORPHOGENESIS%GOBP%GO:0009887 | animal organ morphogenesis |
| 0.4307 | 0.0822 | 0.9381 | ESTABLISHMENT OF ORGANELLE LOCALIZATION%GOBP%GO:0051656 | establishment of organelle localization |
| 0.54 | 0.0013 | 0.8397 | PROTEIN FOLDING%GOBP%GO:0006457 | protein folding |
| 0.6855 | 5.00E-04 | 0.3644 | NUCLEOSIDE TRIPHOSPHATE METABOLIC PROCESS%GOBP%GO:0009141 | nucleoside triphosphate metabolic process |

| | | | | |
|---|---|---|---|---|
| 0.7709 | 0.01 | 0.5176 | CHAPERONE-MEDIATED PROTEIN FOLDING %GOBP% GO:0061077 | chaperone-mediated protein folding |
| -0.345 | 0.0156 | 0.9956 | CELL MOTILITY %GOBP% GO:0048870 | cell motility |
| 0.5312 | 0.0333 | 0.843 | TRANSPORT TO THE GOLGI AND SUBSEQUENT MODIFICATION%REACTOME DATABASE ID RELEASE 66%948021 | Transport to the Golgi and subsequent modification |

| | | | | |
|---|---|---|---|---|
| -0.5326 | 0.0485 | 0.8908 | HALLMARK_ EPITHELIAL_ MESENCHYMAL_ TRANSITION%MSIGDB_ C2%HALLMARK_ EPITHELIAL_ MESENCHYMAL_ TRANSITION | HALLMARK_EPITHELIAL_MESENCHYMAL_TRANSITION |
| 0.5989 | 0.0829 | 0.6023 | CELLULAR COMPONENT MORPHOGENESIS%GOBP%GO:0032989 | cellular component morphogenesis |
| 0.818 | 0.6813 | 0.0853 | TCR SIGNALING%REACTOME DATABASE ID RELEASE 66%202403 | TCR signaling |
| 0.471 | 0.015 | 0.9372 | CELL JUNCTION ORGANIZATION%GOBP%GO:0034330 | cell junction organization |

| | | | | |
|---|---|---|---|---|
| 0.7234 | 0.5321 | 0.0909 | HALLMARK_MYC_TARGETS_V1%MSIGDB_C2%HALLMARK_MYC_TARGETS_V1 | HALLMARK_MYC_TARGETS_V1 |
| 0.6361 | 0.0101 | 0.7513 | PURINE NUCLEOSIDE TRIPHOSPHATE BIOSYNTHETIC PROCESS%GOBP%GO:0009145 | purine nucleoside triphosphate biosynthetic process |
| 0.6633 | 0.0468 | 0.7214 | POSITIVE REGULATION OF CYTOSOLIC CALCIUM ION CONCENTRATION%GOBP%GO:0007204 | positive regulation of cytosolic calcium ion concentration |
| -0.4183 | 0.0647 | 1 | RHYTHMIC PROCESS%GOBP%GO:0048511 | rhythmic process |

| | | | | |
|---|---|---|---|---|
| 0.7812 | 0.0388 | 0.5288 | HALLMARK_CHOLESTEROL_HOMEOSTASIS%MSIGDB_C2%HALLMARK_CHOLESTEROL_HOMEOSTASIS | HALLMARK_CHOLESTEROL_HOMEOSTASIS |
| 0.8534 | 0 | 0.1263 | ACYL-COA METABOLIC PROCESS%GOBP%GO:0006637 | acyl-CoA metabolic process |
| -0.4842 | 0.0018 | 0.8836 | EXTRACELLULAR MATRIX ORGANIZATION%REACTOME DATABASE ID RELEASE 66%1474244 | Extracellular matrix organization |
| 0.8407 | 0.0192 | 0.2428 | GLYCEROPHOSPHOLIPID BIOSYNTHESIS%REACTOME DATABASE ID RELEASE 66%1483206 | Glycerophospholipid biosynthesis |

| | | | | |
|---|---|---|---|---|
| 0.6389 | 0.0089 | 0.5518 | REGULATION OF ACTIN CYTOSKELETON ORGANIZATION%GOBP%GO:0032956 | regulation of actin cytoskeleton organization |
| 0.8198 | 0.0625 | 0.0733 | ER-PHAGOSOME PATHWAY%REACTOME%R-HSA-1236974.4 | ER-Phagosome pathway |
| 0.7076 | 3.00E-04 | 0.3282 | PURINE RIBONUCLEOSIDE TRIPHOSPHATE METABOLIC PROCESS%GOBP%GO:0009205 | purine ribonucleoside triphosphate metabolic process |
| 0.737 | 0.0397 | 0.3886 | INORGANIC CATION TRANSMEMBRANE TRANSPORT%GOBP%GO:0098662 | inorganic cation transmembrane transport |

| | | | | |
|---|---|---|---|---|
| 0.6939 | 0.016 | 0.4435 | REGULATION OF ACTIN FILAMENT LENGTH%GOBP%GO:0030832 | regulation of actin filament length |
| 0.6833 | 0.0819 | 0.2722 | SMALL MOLECULE BIOSYNTHETIC PROCESS%GOBP%GO:0044283 | small molecule biosynthetic process |
| 0.8431 | 0.0177 | 0.1968 | STEROID METABOLIC PROCESS%GOBP%GO:0008202 | steroid metabolic process |
| 0.7475 | 0 | 0.102 | HALLMARK_ADIPOGENESIS%MSIGDB_C2%HALLMARK_ADIPOGENESIS | HALLMARK_ADIPOGENESIS |
| 0.8459 | 0 | 0.0627 | LIPID MODIFICATION%GOBP%GO:0030258 | lipid modification |

| | | | | |
|---|---|---|---|---|
| 0.3745 | 0.0274 | 0.9618 | ANATOMICAL STRUCTURE FORMATION INVOLVED IN MORPHOGENESIS%GOBP%GO:0048646 | anatomical structure formation involved in morphogenesis |
| 0.5799 | 0.0061 | 0.8009 | POSITIVE REGULATION OF CYTOSKELETON ORGANIZATION%GOBP%GO:0051495 | positive regulation of cytoskeleton organization |
| 0.7706 | 0.7049 | 0.0876 | ANTIGEN PROCESSING: UBIQUITINATION & PROTEASOME DEGRADATION%REACTOME DATABASE ID RELEASE 66%983168 | Antigen processing: Ubiquitination & Proteasome degradation |

| | | | | |
|---|---|---|---|---|
| 0.6335 | 0.0061 | 0.4351 | HETEROCYCLE BIOSYNTHETIC PROCESS %GOBP% GO:0018130 | heterocycle biosynthetic process |
| 0.7908 | 0 | 0.0705 | THE CITRIC ACID (TCA) CYCLE AND RESPIRATORY ELECTRON TRANSPORT%REACTOME%R-HSA-1428517.1 | The citric acid (TCA) cycle and respiratory electron transport |
| 0.7204 | 0 | 0.2228 | NUCLEOSIDE MONOPHOSPHATE METABOLIC PROCESS %GOBP% GO:0009123 | nucleoside monophosphate metabolic process |
| 0.8507 | 0 | 0.1138 | PURINE NUCLEOSIDE BISPHOSPHATE METABOLIC PROCESS %GOBP% GO:0034032 | purine nucleoside bisphosphate metabolic process |

| | | | | |
|---|---|---|---|---|
| 0.8347 | 0 | 0.0712 | PROTEIN LOCALIZATION%REACTOME DATABASE ID RELEASE 66%9609507 | Protein localization |
| -0.4879 | 0.056 | 1 | NEGATIVE REGULATION OF ACTIN FILAMENT DEPOLYMERIZATION%GOBP%GO:0030835 | negative regulation of actin filament depolymerization |
| 0.8432 | 4.00E-04 | 0.287 | CITRATE METABOLIC PROCESS%GOBP%GO:0006101 | citrate metabolic process |
| 0.4962 | 0.0102 | 0.9543 | ERYTHROCYTE DIFFERENTIATION%GOBP%GO:0030218 | erythrocyte differentiation |
| 0.8701 | 0.3281 | 0.0914 | REGULATION OF MITOCHONDRIAL MEMBRANE POTENTIAL%GOBP%GO:0051881 | regulation of mitochondrial membrane potential |

| | | | | |
|---|---|---|---|---|
| 0.8768 | 0.4296 | 0.024 | REGULATION OF REACTIVE OXYGEN SPECIES METABOLIC PROCESS%GOBP%GO:2000377 | regulation of reactive oxygen species metabolic process |
| 0.5071 | 0.0225 | 0.8234 | CELL PROJECTION ORGANIZATION%GOBP%GO:0030030 | cell projection organization |
| 0.8452 | 0.0528 | 0.1555 | ALPHA-AMINO ACID CATABOLIC PROCESS%GOBP%GO:1901606 | alpha-amino acid catabolic process |
| 0.8228 | 0.0764 | 0.1963 | PHOSPHOLIPID METABOLIC PROCESS%GOBP%GO:0006644 | phospholipid metabolic process |
| 0.7436 | 0.0252 | 0.56 | ORGANIC ACID TRANSPORT%GOBP%GO:0015849 | organic acid transport |

| | | | | |
|---|---|---|---|---|
| 0.8057 | 9.00E-04 | 0.3827 | NAD METABOLIC PROCESS %GOBP% GO:0019674 | NAD metabolic process |
| 0.5864 | 0.0675 | 0.8602 | CDC42 SIGNALING EVENTS%PATHWAY INTERACTION DATABASE NCI-NATURE CURATED DATA%CDC42 SIGNALING EVENTS | CDC42 signaling events |
| 0.7038 | 0.0647 | 0.6241 | NEGATIVE REGULATION OF NEURON PROJECTION DEVELOPMENT%GOBP%GO:0010977 | negative regulation of neuron projection development |
| -0.5467 | 0.0044 | 0.8734 | RESPONSE TO WOUNDING%GOBP %GO:0009611 | response to wounding |

| | | | | |
|---|---|---|---|---|
| 0.3812 | 0.023 | 0.9972 | CELLULAR RESPONSE TO INTERFERON-GAMMA%GOBP%GO:0071346 | cellular response to interferon-gamma |
| 0.7755 | 0.78 | 0.0772 | CELL PART MORPHOGENESIS%GOBP%GO:0032990 | cell part morphogenesis |
| 0.5458 | 0.0572 | 0.8949 | SIGNALING BY VEGF%REACTOME DATABASE ID RELEASE 66%194138 | Signaling by VEGF |
| 0.8142 | 0 | 0.0786 | FATTY ACID METABOLISM%REACTOME%R-HSA-8978868.4 | Fatty acid metabolism |
| -0.5225 | 0.0385 | 0.9475 | REGULATION OF CELL SHAPE%GOBP%GO:0008360 | regulation of cell shape |

| | | | | |
|---|---|---|---|---|
| 0.3991 | 0.017 | 0.943 | LOCOMOTION%GOBP%GO:0040011 | locomotion |
| 0.5238 | 0.0876 | 0.9123 | ANGIOGENESIS%PANTHER PATHWAY%P00005 | Angiogenesis |
| 0.7228 | 0.003 | 0.4273 | ANION TRANSPORT%GOBP%GO:0006820 | anion transport |
| 0.6939 | 0.0177 | 0.4361 | REGULATION OF ACTIN POLYMERIZATION OR DEPOLYMERIZATION%GOBP%GO:0008064 | regulation of actin polymerization or depolymerization |
| -0.4355 | 0.09 | 1 | NEGATIVE REGULATION OF PROTEIN COMPLEX DISASSEMBLY%GOBP%GO:0043242 | negative regulation of protein complex disassembly |

| | | | | |
|---|---|---|---|---|
| 0.6342 | 0.0932 | 0.5763 | ORGANELLE BIOGENESIS AND MAINTENANCE%REACTOME%R-HSA-1852241.4 | Organelle biogenesis and maintenance |
| 0.6256 | 0.0063 | 0.7262 | NUCLEOSIDE TRIPHOSPHATE BIOSYNTHETIC PROCESS%GOBP%GO:0009142 | nucleoside triphosphate biosynthetic process |
| 0.6015 | 3.00E-04 | 0.5352 | CARBOHYDRATE DERIVATIVE METABOLIC PROCESS%GOBP%GO:1901135 | carbohydrate derivative metabolic process |

| | | | | |
|---|---|---|---|---|
| 0.2469 | 0.0777 | 1 | RHO GTPASES ACTIVATE PKNS%REACTOME%R-HSA-5625740.1 | RHO GTPases activate PKNs |

| | | | | |
|---|---|---|---|---|
| -0.461 | 0 | 0.9015 | ACTIN FILAMENT ORGANIZATION%GOBP%GO:0007015 | actin filament organization |
| 0.8211 | 0.002 | 0.2135 | OXIDATIVE PHOSPHORYLATION%GOBP%GO:0006119 | oxidative phosphorylation |
| 0.7941 | 0.0339 | 0.5308 | NEUROMUSCULAR PROCESS%GOBP%GO:0050905 | neuromuscular process |
| 0.8579 | 0.0051 | 0.0357 | DICARBOXYLIC ACID METABOLIC PROCESS%GOBP%GO:0043648 | dicarboxylic acid metabolic process |
| 0.5927 | 0.0204 | 0.6948 | PROTEIN HOMOOLIGOMERIZATION%GOBP%GO:0051260 | protein homooligomerization |
| -0.6057 | 0.0011 | 0.76 | WOUND HEALING%GOBP%GO:0042060 | wound healing |

| | | | | |
|---|---|---|---|---|
| 0.8515 | 0.0044 | 0.2186 | REGULATION OF STEROID METABOLIC PROCESS%GOBP%GO:0019218 | regulation of steroid metabolic process |
| 0.7147 | 0 | 0.1286 | METABOLISM OF LIPIDS%REACTOME%R-HSA-556833.5 | Metabolism of lipids |
| 0.5618 | 0.0416 | 0.8367 | NEGATIVE REGULATION OF CELL PROJECTION ORGANIZATION%GOBP%GO:0031345 | negative regulation of cell projection organization |
| 0.7557 | 0.0092 | 0.5294 | SULFUR COMPOUND BIOSYNTHETIC PROCESS%GOBP%GO:0044272 | sulfur compound biosynthetic process |
| 0.8359 | 0 | 0.0125 | FATTY ACID METABOLIC PROCESS%GOBP%GO:0006631 | fatty acid metabolic process |

| | | | | |
|---|---|---|---|---|
| 0.7826 | 0.534 | 0.0747 | PROTEASOME-MEDIATED UBIQUITIN-DEPENDENT PROTEIN CATABOLIC PROCESS%GOBP%GO:0043161 | proteasome-mediated ubiquitin-dependent protein catabolic prc |
| 0.4538 | 0.0588 | 0.8716 | POSITIVE REGULATION OF ORGANELLE ORGANIZATION%GOBP%GO:0010638 | positive regulation of organelle organization |
| 0.8639 | 0 | 0.0675 | FATTY ACID OXIDATION%GOBP%GO:0019395 | fatty acid oxidation |
| 0.5181 | 0.0163 | 0.8347 | CELL ADHESION%GOBP%GO:0007155 | cell adhesion |
| 0.7939 | 1.00E-04 | 0.0913 | LIPID CATABOLIC PROCESS%GOBP%GO:0016042 | lipid catabolic process |

| | | | | |
|---|---|---|---|---|
| 0.7713 | 0.0378 | 0.4875 | HORMONE-MEDIATED SIGNALING PATHWAY%GOBP%GO:0009755 | hormone-mediated signaling pathway |
| 0.863 | 0 | 0.1818 | REGULATION OF COLD-INDUCED THERMOGENESIS%GOBP%GO:0120161 | regulation of cold-induced thermogenesis |

| EM1_gs_si: | EM1_NES.. | EM1_NES.. | EM1_pvalu | EM1_pvalu | name | shared.name |
|---|---|---|---|---|---|---|
| 423 | 2.4327 | 1.6118 | 0 | 0 | OXIDATION | OXIDATION-REDUCTION PROCESS% |
| 626 | 1.9431 | 1.2971 | 0 | 0.0639 | TRANSMEN | TRANSMEMBRANE TRANSPORT%G |
| 50 | 2.1536 | 1.4314 | 0 | 0.008 | FATTY ACID | FATTY ACID BETA-OXIDATION%GO |
| 354 | -2.0587 | -0.7309 | 0 | 0.9894 | PLASMA M | PLASMA MEMBRANE BOUNDED C |
| 91 | 2.2486 | 1.429 | 0 | 0.0119 | NUCLEOSID | NUCLEOSIDE BISPHOSPHATE META |
| 68 | 1.885 | 1.4602 | 0 | 0.0013 | ELECTRON | ELECTRON TRANSPORT CHAIN%G( |
| 327 | 2.1673 | 1.4436 | 0 | 0.0056 | ORGANOPH | ORGANOPHOSPHATE BIOSYNTHET |
| 73 | -1.6448 | -0.9537 | 0.014 | 0.512 | REGULATIC | REGULATION OF PROTEIN DEPOLY |
| 66 | 1.8347 | 1.4238 | 0.0053 | 0.0134 | RESPIRATO | RESPIRATORY ELECTRON TRANSPC |
| 218 | 1.7296 | 1.2205 | 0.0037 | 0.2032 | CARBOXYLI | CARBOXYLIC ACID TRANSPORT%G |
| 253 | 2.3033 | 1.4207 | 0 | 0.0067 | COENZYME | COENZYME METABOLIC PROCESS% |
| 22 | 1.7416 | 1.3912 | 0.0019 | 0.0135 | CITRIC ACID | CITRIC ACID CYCLE (TCA CYCLE)%R |
| 754 | -1.741 | 0.7778 | 0 | 0.9025 | REGULATIC | REGULATION OF CELL PROJECTION |
| 125 | -1.6118 | -0.8386 | 0.0216 | 0.7071 | CIRCADIAN | CIRCADIAN RHYTHM%GOBP%GO:( |
| 599 | -1.6388 | 1.0291 | 0 | 0.4204 | BIOLOGICA | BIOLOGICAL ADHESION%GOBP%G |
| 322 | 2.3895 | 1.4234 | 0 | 0.0011 | PURINE NU | PURINE NUCLEOTIDE METABOLIC |
| 172 | 1.9958 | 1.3167 | 0 | 0.0597 | PURINE NU | PURINE NUCLEOSIDE TRIPHOSPHA |
| 234 | -1.7407 | -1.5697 | 0.0047 | 0.0159 | REGULATIC | REGULATION OF LEUKOCYTE PROL |
| 496 | -1.7849 | 0.8452 | 0 | 0.772 | REGULATIC | REGULATION OF CELL MORPHOGE |
| 93 | 1.8096 | 1.2039 | 0.0034 | 0.1896 | MITOCHON | MITOCHONDRIAL MEMBRANE OR( |

| | | | | | | |
|---|---|---|---|---|---|---|
| 328 | 1.5699 | 1.1262 | 0.0231 | 0.3241 | REGULATIC | REGULATION OF CYTOSOLIC CALCI |
| 729 | 2.3887 | 1.5626 | 0 | 0 | CELLULAR I | CELLULAR LIPID METABOLIC PROC |
| 84 | -1.7336 | -0.9253 | 0 | 0.5758 | EPH-EPHRI | EPH-EPHRIN SIGNALING%REACTOI |
| 101 | -1.8828 | 0.6536 | 0.0022 | 0.9299 | CELL-MATF | CELL-MATRIX ADHESION%GOBP% |
| 98 | 1.6372 | 0.9926 | 0.007 | 0.5575 | ERYTHROC | ERYTHROCYTE HOMEOSTASIS%GC |

| | | | | | | |
|---|---|---|---|---|---|---|
| 184 | -2.0187 | -1.3941 | 0 | 0.0702 | MUSCLE CO | MUSCLE CONTRACTION%REACTOM |
| 166 | 1.0811 | 1.4753 | 0.3333 | 0.0038 | PROTEIN H | PROTEIN HETEROOLIGOMERIZATIO |
| 627 | -1.7094 | -0.9385 | 0.0027 | 0.5541 | NABA_MAT | NABA_MATRISOME_ASSOCIATED% |
| 153 | 1.6651 | 1.2836 | 0 | 0.0882 | SIGNALING | SIGNALING BY NUCLEAR RECEPTOR |

| | | | | | | |
|---|---|---|---|---|---|---|
| 422 | -1.7809 | -1.3952 | 0 | 0.0307 | RESPONSE | RESPONSE TO GROWTH FACTOR% |
| 457 | -1.7831 | 0.9989 | 0.0028 | 0.4814 | MUSCLE ST | MUSCLE STRUCTURE DEVELOPME |
| 187 | 2.1536 | 1.3582 | 0 | 0.0238 | NUCLEOSIE | NUCLEOSIDE PHOSPHATE BIOSYNT |
| 167 | 1.9867 | 1.3228 | 0 | 0.0447 | RIBONUCLE | RIBONUCLEOSIDE TRIPHOSPHATE |
| 163 | 2.1402 | 1.3475 | 0 | 0.0359 | PURINE NU | PURINE NUCLEOSIDE MONOPHOS |

| | | | | | | |
|---|---|---|---|---|---|---|
| 66 | 1.8449 | 1.0995 | 0.0018 | 0.3614 | PURINE RIE | PURINE RIBONUCLEOSIDE TRIPHOS |
| 259 | -1.6828 | 0.8592 | 0.0024 | 0.7194 | MUSCLE CE | MUSCLE CELL DIFFERENTIATION% |
| 183 | 2.0876 | 1.4289 | 0 | 0.0096 | CARBOXYLI | CARBOXYLIC ACID CATABOLIC PRC |
| 142 | -1.6959 | 0.9128 | 0.0167 | 0.6465 | DNA PACK/ | DNA PACKAGING%GOBP%GO:000 |
| 213 | -1.9562 | -1.2255 | 0 | 0.1898 | NABA_ECM | NABA_ECM_REGULATORS%MSIGD |

| | | | | | | |
|---|---|---|---|---|---|---|
| 76 | 2.1677 | 1.4584 | 0 | 0.0013 | FATTY ACID | FATTY ACID CATABOLIC PROCESS% |
| 69 | 2.1334 | 1.4815 | 0 | 0 | LIPID OXIDA | LIPID OXIDATION%GOBP%GO:003 |
| 131 | 2.3247 | 1.4357 | 0 | 0.0105 | PURINE RIB | PURINE RIBONUCLEOTIDE BIOSYN |
| 249 | 1.8331 | 1.288 | 0 | 0.0892 | SULFUR CO | SULFUR COMPOUND METABOLIC |
| 91 | 2.0119 | 1.3982 | 0 | 0.0076 | RESPIRATO | RESPIRATORY ELECTRON TRANSPO |

| | | | | | | |
|---|---|---|---|---|---|---|
| 358 | 1.6655 | 1.3028 | 0.005 | 0.0713 | CATION TR | CATION TRANSMEMBRANE TRANS |
| 180 | 2.4751 | 1.593 | 0 | 0 | HALLMARK | HALLMARK_OXIDATIVE_PHOSPHO |
| 305 | -1.6091 | 1.1105 | 0.008 | 0.3333 | NEGATIVE | NEGATIVE REGULATION OF CELLU |
| 24 | 1.8684 | 1.2445 | 0 | 0.1339 | CRISTAE FC | CRISTAE FORMATION%REACTOME |

| | | | | | | |
|---|---|---|---|---|---|---|
| 621 | -1.7437 | 0.7493 | 0 | 0.9211 | TISSUE MO | TISSUE MORPHOGENESIS%GOBP% |
| 417 | 2.426 | 1.506 | 0 | 0.0011 | MONOCAR | MONOCARBOXYLIC ACID METABO |
| 711 | 2.4087 | 1.4838 | 0 | 0 | ORGANOPI | ORGANOPHOSPHATE METABOLIC |
| 24 | 1.939 | 1.2775 | 0 | 0.0937 | NADH MET | NADH METABOLIC PROCESS%GOB |
| 60 | 1.5471 | 1.4551 | 0.0237 | 0.0026 | HEDGEHOG | HEDGEHOG LIGAND BIOGENESIS% |

| | | | | | | |
|---|---|---|---|---|---|---|
| 59 | 2.1797 | 1.4165 | 0 | 0.0052 | MITOCHON | MITOCHONDRIAL PROTEIN IMPOR |
| 151 | 1.6396 | 1.3399 | 0.0088 | 0.0277 | POSITIVE R | POSITIVE REGULATION OF LIPID M |
| 376 | 1.8189 | 1.2735 | 0 | 0.0999 | INORGANIC | INORGANIC ION TRANSMEMBRAN |
| 223 | -1.7832 | -1.5801 | 0.0023 | 0.0165 | REGULATIC | REGULATION OF LYMPHOCYTE PR |
| 881 | -1.7455 | -1.3619 | 0 | 0.0189 | NABA_MA | NABA_MATRISOME%MSIGDB_C2% |

| | | | | | | |
|---|---|---|---|---|---|---|
| 264 | -1.6087 | -1.0323 | 0.0164 | 0.3863 | CILIUM ASS | CILIUM ASSEMBLY%GOBP%GO:00 |
| 76 | 2.1901 | 1.4207 | 0 | 0.0054 | THIOESTER | THIOESTER METABOLIC PROCESS% |
| 33 | -1.7279 | 1.1654 | 0.0088 | 0.2759 | SIG_REGUL | SIG_REGULATION_OF_THE_ACTIN |
| 69 | 1.711 | 1.3758 | 0.0034 | 0.0191 | PROTEIN L( | PROTEIN LOCALIZATION TO MITO( |

| | | | | | | |
|---|---|---|---|---|---|---|
| 75 | 1.5791 | 1.3822 | 0.0071 | 0.0181 | TP53 REGU | TP53 REGULATES METABOLIC GEN |
| 1288 | -1.8125 | 0.8401 | 0 | 0.8454 | MOVEMEN | MOVEMENT OF CELL OR SUBCELLU |
| 69 | 1.822 | 1.1035 | 0 | 0.3589 | RIBONUCLE | RIBONUCLEOSIDE TRIPHOSPHATE |
| 377 | 1.9147 | 1.4848 | 0 | 0.0034 | MITOCHON | MITOCHONDRION ORGANIZATION |

| | | | | | | |
|---|---|---|---|---|---|---|
| 95 | 1.3031 | 1.4564 | 0.1145 | 0.0024 | INTERLEUK | INTERLEUKIN-1 SIGNALING%REAC |
| 183 | -1.6518 | 0.6056 | 0.0127 | 0.9927 | ESTABLISHI | ESTABLISHMENT OR MAINTENANC |
| 104 | 1.8823 | 1.3939 | 0.0016 | 0.0158 | ANTIBIOTIC | ANTIBIOTIC METABOLIC PROCESS% |
| 59 | 2.161 | 1.4038 | 0 | 0.0091 | AEROBIC RI | AEROBIC RESPIRATION%GOBP%G |
| 62 | 1.9165 | 1.0919 | 0 | 0.3703 | ATP BIOSYI | ATP BIOSYNTHETIC PROCESS%GOI |

| | | | | | | |
|---|---|---|---|---|---|---|
| 91 | 1.4618 | 1.5313 | 0.032 | 0 | ANTIGEN P | ANTIGEN PROCESSING-CROSS PRE |
| 360 | -1.8811 | 1.1534 | 0 | 0.2227 | REGULATIC | REGULATION OF ACTIN FILAMENT- |
| 682 | -1.6259 | 1.1778 | 0 | 0.1746 | CELL MORF | CELL MORPHOGENESIS%GOBP%G( |
| 432 | 1.9241 | 1.3287 | 0 | 0.0376 | CARBOHYD | CARBOHYDRATE DERIVATIVE BIOS |

| | | | | | | |
|---|---|---|---|---|---|---|
| 227 | -1.6542 | -1.4228 | 0.007 | 0.0616 | REGULATIC | REGULATION OF GENE EXPRESSIOI |
| 34 | 1.9001 | 1.3441 | 0 | 0.0392 | TRICARBOX | TRICARBOXYLIC ACID METABOLIC |
| 66 | -2.0405 | 0.5127 | 0 | 0.9916 | ACTIN FILA | ACTIN FILAMENT BUNDLE ASSEME |
| 122 | -1.6361 | 0.9419 | 0.0157 | 0.6169 | REGULATIC | REGULATION OF EXTENT OF CELL ( |
| 63 | 1.6761 | 1.3877 | 0.0086 | 0.013 | ESTABLISHI | ESTABLISHMENT OF PROTEIN LOC/ |

| | | | | | | |
|---|---|---|---|---|---|---|
| 125 | -1.7242 | 0.8499 | 0.0025 | 0.7252 | CELL JUNCT | CELL JUNCTION ASSEMBLY%GOBP |
| 66 | -1.6324 | 0.9471 | 0.0143 | 0.6182 | PID_CDC42 | PID_CDC42_PATHWAY%MSIGDB_ |
| 96 | -1.593 | 0.8467 | 0.0152 | 0.75 | MHC CLASS | MHC CLASS II ANTIGEN PRESENTAT |
| 270 | 1.8195 | 1.31 | 0 | 0.0452 | SMALL MO | SMALL MOLECULE CATABOLIC PRO |
| 183 | 2.0945 | 1.4175 | 0 | 0.0107 | ORGANIC A | ORGANIC ACID CATABOLIC PROCES |

| | | | | | | |
|---|---|---|---|---|---|---|
| 974 | -2.2816 | -0.8749 | 0 | 0.9714 | CYTOSKELE | CYTOSKELETON ORGANIZATION%( |
| 523 | -1.8748 | 1.1319 | 0 | 0.234 | REGULATIC | REGULATION OF ANATOMICAL STI |
| 519 | 1.635 | 1.2875 | 0 | 0.0394 | PROTEIN C | PROTEIN COMPLEX OLIGOMERIZA |
| 262 | 2.0085 | 1.4593 | 0 | 0.0026 | GLYCEROLI | GLYCEROLIPID METABOLIC PROCE |
| 368 | -2.0529 | -0.7114 | 0 | 0.9942 | CELL PROJE | CELL PROJECTION ASSEMBLY%GOI |

| | | | | | | |
|---|---|---|---|---|---|---|
| 652 | 1.8919 | 1.2766 | 0 | 0.0406 | AROMATIC | AROMATIC COMPOUND BIOSYNTH |
| 161 | 1.7881 | 1.4588 | 0.0055 | 0.0027 | REGULATIC | REGULATION OF LIPID BIOSYNTHE |
| 329 | 2.0271 | 1.2404 | 0 | 0.1381 | REGULATIC | REGULATION OF SMALL MOLECUL |
| 91 | 2.119 | 1.4175 | 0 | 0.0025 | MONOCAR | MONOCARBOXYLIC ACID CATABOI |

| | | | | | | |
|---|---|---|---|---|---|---|
| 162 | 2.1179 | 1.3513 | 0 | 0.0324 | PURINE RIE | PURINE RIBONUCLEOSIDE MONOF |
| 268 | -1.825 | -0.783 | 0.0025 | 0.9078 | HEART MO | HEART MORPHOGENESIS%GOBP% |
| 359 | 2.3587 | 1.4039 | 0 | 0.0043 | PURINE-CO | PURINE-CONTAINING COMPOUND |
| 75 | -1.7886 | -0.4659 | 0 | 0.9801 | ACTIN FILA | ACTIN FILAMENT-BASED MOVEME |
| 428 | 2.418 | 1.4601 | 0 | 0 | NUCLEOSIE | NUCLEOSIDE PHOSPHATE METABC |

| | | | | | | |
|---|---|---|---|---|---|---|
| 745 | -1.7412 | 0.7849 | 0 | 0.8919 | REGULATIC | REGULATION OF PLASMA MEMBR |
| 60 | 1.8574 | 1.0507 | 0.0018 | 0.4409 | PYRUVATE | PYRUVATE METABOLIC PROCESS% |
| 103 | 1.5596 | 0.9471 | 0.0216 | 0.6066 | REACTIVE ( | REACTIVE OXYGEN SPECIES METAE |
| 81 | 1.9076 | 1.2508 | 0 | 0.1438 | RIBONUCLE | RIBONUCLEOSIDE MONOPHOSPHA |

| | | | | | | |
|---|---|---|---|---|---|---|
| 239 | 1.7473 | 1.0749 | 0.0122 | 0.4038 | MONOVAL | MONOVALENT INORGANIC CATIOI |
| 225 | -1.758 | -1.5755 | 0 | 0.0295 | REGULATIC | REGULATION OF MONONUCLEAR ( |
| 59 | 1.4571 | 1.4525 | 0.0485 | 0.0062 | ABC TRANS | ABC TRANSPORTER DISORDERS%R |
| 86 | -1.6702 | 0.7452 | 0.0143 | 0.8402 | COLLAGEN | COLLAGEN FORMATION%REACTOI |
| 260 | -1.6237 | 1.1138 | 0.0145 | 0.3432 | MUSCLE OI | MUSCLE ORGAN DEVELOPMENT% |

| | | | | | | |
|---|---|---|---|---|---|---|
| 136 | 1.9214 | 1.1605 | 0 | 0.2737 | AMMONIU | AMMONIUM ION METABOLIC PRC |
| 423 | -2.3853 | 0.6483 | 0 | 0.9934 | ACTIN CYT( | ACTIN CYTOSKELETON ORGANIZAT |
| 326 | -1.7293 | 1.1486 | 0 | 0.2255 | REGULATIC | REGULATION OF SUPRAMOLECUL/ |
| 582 | 1.8891 | 1.3072 | 0 | 0.0364 | NUCLEOBA | NUCLEOBASE-CONTAINING COMP |

| | | | | | | |
|---|---|---|---|---|---|---|
| 76 | 1.9091 | 1.2282 | 0 | 0.163 | PURINE NU | PURINE NUCLEOSIDE MONOPHOS |
| 154 | 1.7563 | 1.3961 | 0.0017 | 0.0244 | PROTEIN TI | PROTEIN TETRAMERIZATION%GOI |
| 102 | 2.0557 | 1.0929 | 0 | 0.3505 | HALLMARK | HALLMARK_PEROXISOME%MSIGD |
| 69 | -2.0489 | 0.5089 | 0 | 0.9933 | ACTIN FILA | ACTIN FILAMENT BUNDLE ORGANI |
| 200 | -1.6341 | 0.7277 | 0.0095 | 0.861 | REGULATIC | REGULATION OF AXONOGENESIS% |

| | | | | | | |
|---|---|---|---|---|---|---|
| 426 | -1.9419 | -0.8441 | 0 | 0.9577 | SUPRAMOL | SUPRAMOLECULAR FIBER ORGANI |
| 149 | 1.6129 | 1.0375 | 0.0218 | 0.4757 | REGULATIC | REGULATION OF TUMOR NECROSI |
| 168 | -1.7634 | -1.5338 | 0.0046 | 0.0241 | REGULATIC | REGULATION OF T CELL PROLIFERA |
| 167 | 2.136 | 1.3439 | 0 | 0.0324 | RIBONUCLE | RIBONUCLEOSIDE MONOPHOSPHA |

| | | | | | | |
|---|---|---|---|---|---|---|
| 38 | -1.6897 | -1.0036 | 0.0071 | 0.4303 | REGULATIC | REGULATION OF ACTIN FILAMENT |
| 37 | 2.0052 | 1.3415 | 0 | 0.0245 | MITOCHON | MITOCHONDRIAL FATTY ACID BET/ |
| 322 | 1.9844 | 1.4934 | 0 | 0.0024 | REGULATIC | REGULATION OF LIPID METABOLIC |
| 119 | -1.7732 | 0.6188 | 0.0069 | 0.9612 | RESPONSE | RESPONSE TO INTERFERON-GAMN |
| 127 | -1.5985 | 1.213 | 0.0025 | 0.1663 | PID_PDGFF | PID_PDGFRB_PATHWAY%MSIGDB |

| | | | | | | |
|---|---|---|---|---|---|---|
| 499 | 2.1159 | 1.3905 | 0 | 0.0054 | DRUG MET | DRUG METABOLIC PROCESS%GOB |
| 96 | -1.7147 | -1.3158 | 0.0026 | 0.119 | REGULATIC | REGULATION OF PROTEIN COMPLI |
| 305 | 2.4038 | 1.4927 | 0 | 0.0011 | PURINE RIE | PURINE RIBONUCLEOTIDE METAB( |
| 147 | 1.6112 | 1.0468 | 0.017 | 0.4549 | REGULATIC | REGULATION OF TUMOR NECROSI |
| 195 | -1.7533 | 1.3728 | 0.0025 | 0.0307 | REGULATIC | REGULATION OF PROTEIN POLYMI |

| | | | | | | |
|---|---|---|---|---|---|---|
| 237 | -1.9007 | 1.262 | 0 | 0.1006 | REGULATIC | REGULATION OF ACTIN FILAMENT |
| 99 | -1.6061 | 0.7874 | 0.0188 | 0.8077 | PID_CXCR4 | PID_CXCR4_PATHWAY%MSIGDB_( |
| 111 | 1.6438 | 1.0523 | 0.0124 | 0.4591 | HALLMARK | HALLMARK_BILE_ACID_METABOLI |
| 77 | -1.7905 | 1.2987 | 0.0048 | 0.0637 | POSITIVE R | POSITIVE REGULATION OF ACTIN F |

| | | | | | | |
|---|---|---|---|---|---|---|
| 207 | -1.7283 | 0.9013 | 0.005 | 0.6626 | STRIATED M | STRIATED MUSCLE CELL DIFFEREN |
| 90 | -1.6202 | 0.8735 | 0.0185 | 0.7016 | VEGFA-VEG | VEGFA-VEGFR2 PATHWAY%REACT |
| 128 | -1.6077 | -1.1887 | 0.0229 | 0.2517 | NEGATIVE I | NEGATIVE REGULATION OF LEUKO |
| 63 | -1.7968 | 0.5354 | 0.0043 | 0.9747 | SEMAPHOF | SEMAPHORIN INTERACTIONS%REA |

| | | | | | | |
|---|---|---|---|---|---|---|
| 909 | -1.8554 | -0.8795 | 0 | 0.8657 | LOCALIZAT | LOCALIZATION OF CELL%GOBP%G |
| 952 | 1.7212 | 1.1553 | 0 | 0.2126 | ION TRANS | ION TRANSPORT%GOBP%GO:0006 |
| 80 | 1.9741 | 1.3611 | 0 | 0.0288 | MITOCHON | MITOCHONDRIAL BIOGENESIS%RE |
| 155 | -2.1591 | -0.9631 | 0 | 0.4813 | MUSCLE CE | MUSCLE CELL DEVELOPMENT%GO |
| 179 | -1.7887 | 1.1418 | 0.0025 | 0.2715 | INFLAMMA | INFLAMMATION MEDIATED BY CH |

| | | | | | | |
|---|---|---|---|---|---|---|
| 406 | -1.7761 | -1.3628 | 0 | 0.029 | CELLULAR I | CELLULAR RESPONSE TO GROWTH |
| 360 | -2.1385 | 0.9222 | 0 | 0.6586 | SIGNALING | SIGNALING BY RHO GTPASES%REA |
| 139 | 2.2814 | 1.4542 | 0 | 0.0023 | RIBONUCLE | RIBONUCLEOTIDE BIOSYNTHETIC F |
| 117 | 1.8041 | 1.3337 | 0.0018 | 0.0272 | REGULATIC | REGULATION OF CELLULAR KETON |

| | | | | | | |
|---|---|---|---|---|---|---|
| 179 | -1.9735 | 1.2044 | 0 | 0.1894 | POSITIVE R | POSITIVE REGULATION OF SUPRAM |
| 107 | 2.1226 | 1.4301 | 0 | 0.0074 | RESPIRATO | RESPIRATORY ELECTRON TRANSPO |
| 713 | 2.1088 | 1.4087 | 0 | 0.001 | CARBOXYLI | CARBOXYLIC ACID METABOLIC PRO |

| | | | | | | |
|---|---|---|---|---|---|---|
| 1684 | -1.8351 | 1.0404 | 0 | 0.3922 | CELL DEVEI | CELL DEVELOPMENT%GOBP%GO:( |
| 496 | -1.8541 | 1.0776 | 0.0032 | 0.3359 | REGULATIC | REGULATION OF CYTOSKELETON C |
| 107 | -1.73 | 0.8813 | 0 | 0.6985 | GOLGI-TO-I | GOLGI-TO-ER RETROGRADE TRANS |
| 79 | 1.7966 | 1.3248 | 0.0017 | 0.0682 | PROTEIN H | PROTEIN HOMOTETRAMERIZATIOI |
| 753 | 2.1102 | 1.3905 | 0 | 0 | OXOACID N | OXOACID METABOLIC PROCESS%G |

| | | | | | | |
|---|---|---|---|---|---|---|
| 139 | 1.7758 | 1.1798 | 0 | 0.2544 | METABOLIS | METABOLISM OF STEROIDS%REAC |
| 402 | 2.2737 | 1.438 | 0 | 0 | COFACTOR | COFACTOR METABOLIC PROCESS% |
| 178 | -1.5916 | 0.7726 | 0.0189 | 0.8218 | AMEBOIDA | AMEBOIDAL-TYPE CELL MIGRATIO |
| 53 | -1.5928 | 0.8041 | 0.0201 | 0.7603 | PID_RAC1_ | PID_RAC1_PATHWAY%MSIGDB_C. |

| | | | | | | |
|---|---|---|---|---|---|---|
| 123 | -1.6083 | 1.2135 | 0.0077 | 0.1626 | PDGFR-BET | PDGFR-BETA SIGNALING PATHWAY |
| 142 | -2.1315 | -0.8584 | 0 | 0.6741 | STRIATED M | STRIATED MUSCLE CELL DEVELOPM |
| 1019 | -1.7608 | 0.9666 | 0 | 0.5825 | PLASMA M | PLASMA MEMBRANE BOUNDED C |

| | | | | | | |
|---|---|---|---|---|---|---|
| 84 | 1.5703 | 1.3587 | 0.0214 | 0.0203 | CARBOHYD | CARBOHYDRATE BIOSYNTHETIC PR |
| 91 | 2.2661 | 1.4276 | 0 | 0.004 | RIBONUCLE | RIBONUCLEOSIDE BISPHOSPHATE |
| 100 | 2.128 | 1.2847 | 0 | 0.0909 | PYRIDINE N | PYRIDINE NUCLEOTIDE METABOLIC |
| 115 | 2.2224 | 1.2537 | 0 | 0.1197 | OXIDOREDU | OXIDOREDUCTION COENZYME ME |
| 292 | 0.9641 | 1.472 | 0.495 | 0.0012 | RESPONSE | RESPONSE TO TOXIC SUBSTANCE% |

| | | | | | | |
|---|---|---|---|---|---|---|
| 721 | -1.6524 | 0.7607 | 0.003 | 0.9346 | CELL CYCLE | CELL CYCLE PROCESS%GOBP%GO:[illegible] |
| 353 | 1.9392 | 1.3688 | 0 | 0.0389 | LIPID BIOSY | LIPID BIOSYNTHETIC PROCESS%GO |
| 57 | 1.4668 | 1.4502 | 0.0358 | 0.005 | DEFECTIVE | DEFECTIVE CFTR CAUSES CYSTIC FI |
| 495 | -1.6658 | 0.8714 | 0.0027 | 0.6946 | MORPHOG | MORPHOGENESIS OF AN EPITHELI |
| 179 | 2.1419 | 1.3758 | 0 | 0.0208 | NUCLEOTIC | NUCLEOTIDE BIOSYNTHETIC PROC |

| | | | | | | |
|---|---|---|---|---|---|---|
| 330 | 2.4023 | 1.4408 | 0 | 0.0022 | RIBOSE PH( | RIBOSE PHOSPHATE METABOLIC P |
| 658 | -1.6915 | 1.0041 | 0 | 0.4804 | HEAD DEVE | HEAD DEVELOPMENT%GOBP%GO |
| 155 | 1.9959 | 1.1772 | 0 | 0.2134 | COFACTOR | COFACTOR BIOSYNTHETIC PROCES |
| 100 | 2.139 | 1.2771 | 0 | 0.0785 | NICOTINAM | NICOTINAMIDE NUCLEOTIDE MET/ |
| 52 | 1.6902 | 1.3365 | 0.0127 | 0.0519 | PROTEIN T/ | PROTEIN TARGETING TO MITOCH( |

| | | | | | | |
|---|---|---|---|---|---|---|
| 128 | -1.7383 | 0.9417 | 0.0026 | 0.6067 | ER TO GOL( | ER TO GOLGI ANTEROGRADE TRAN |
| 119 | 1.9345 | 1.221 | 0 | 0.1684 | COENZYME | COENZYME BIOSYNTHETIC PROCES |
| 149 | -1.8204 | 0.6482 | 0 | 0.9354 | CELL-SUBST | CELL-SUBSTRATE ADHESION%GOB |
| 109 | -1.7798 | 1.3041 | 0.0026 | 0.0672 | POSITIVE R | POSITIVE REGULATION OF PROTEI |
| 320 | 1.7416 | 1.1981 | 0.007 | 0.2013 | ORGANIC A | ORGANIC ANION TRANSPORT%GO |

| | | | | | | |
|---|---|---|---|---|---|---|
| 139 | -1.8483 | -0.9482 | 0 | 0.4979 | HALLMARK | HALLMARK_COAGULATION%MSIG |
| 80 | 1.5613 | 1.3801 | 0.0249 | 0.0127 | CELLULAR / | CELLULAR AMINO ACID CATABOLI( |
| 314 | 2.4376 | 1.5272 | 0 | 0 | RIBONUCLI | RIBONUCLEOTIDE METABOLIC PR( |
| 795 | -1.8741 | -0.8956 | 0 | 0.8193 | CELL MIGR/ | CELL MIGRATION%GOBP%GO:001 |
| 503 | 1.7601 | 1.2641 | 0 | 0.1184 | ION TRANS | ION TRANSMEMBRANE TRANSPOI |

| | | | | | | |
|---|---|---|---|---|---|---|
| 152 | 1.5686 | 1.1441 | 0.0175 | 0.2926 | MONOCAR | MONOCARBOXYLIC ACID BIOSYNT |
| 75 | -1.8031 | 0.9277 | 0.0068 | 0.616 | COPI-MEDI | COPI-MEDIATED ANTEROGRADE T |
| 822 | 1.6052 | 1.0783 | 0.0139 | 0.3428 | RESPONSE | RESPONSE TO BACTERIUM%GOBP |
| 64 | 2.0288 | 1.401 | 0 | 0.0092 | MITOCHON | MITOCHONDRIAL TRANSMEMBRA |
| 76 | 1.7661 | 1.3918 | 0.0053 | 0.0054 | GLUCOSE M | GLUCOSE METABOLIC PROCESS%G |

| | | | | | | |
|---|---|---|---|---|---|---|
| 145 | 2.1967 | 1.3226 | 0 | 0.0564 | PURINE-CO | PURINE-CONTAINING COMPOUND |
| 310 | 1.5815 | 1.2179 | 0.0014 | 0.0883 | AMINO ACI | AMINO ACID AND DERIVATIVE ME |
| 156 | 2.3251 | 1.5775 | 0 | 0 | HALLMARK | HALLMARK_FATTY_ACID_METABO |

| | | | | | | |
|---|---|---|---|---|---|---|
| 484 | 2.3399 | 1.4337 | 0 | 0.001 | NUCLEOBA | NUCLEOBASE-CONTAINING SMALL |
| 253 | -1.7554 | 1.101 | 0.0026 | 0.3649 | SYNAPSE O | SYNAPSE ORGANIZATION%GOBP% |
| 777 | 2.1359 | 1.4157 | 0 | 0.001 | ORGANIC A | ORGANIC ACID METABOLIC PROCE |
| 374 | -1.8777 | 1.1403 | 0 | 0.2319 | REGULATIO | REGULATION OF CELLULAR COMP |

| | | | | | | |
|---|---|---|---|---|---|---|
| 53 | 1.9512 | 1.336 | 0 | 0.0457 | PYRUVATE | PYRUVATE METABOLISM AND CITI |
| 46 | 1.8665 | 1.3719 | 0 | 0.0246 | ATP SYNTH | ATP SYNTHESIS COUPLED ELECTRC |
| 152 | 2.0505 | 1.3718 | 0.0016 | 0.0308 | MITOCHON | MITOCHONDRIAL TRANSPORT%GC |
| 120 | 2.2612 | 1.4916 | 0 | 0.0012 | CELLULAR I | CELLULAR RESPIRATION%GOBP%G |
| 104 | 2.0791 | 1.2787 | 0 | 0.0901 | PYRIDINE-C | PYRIDINE-CONTAINING COMPOUN |

| | | | | | | |
|---|---|---|---|---|---|---|
| 141 | 2.2015 | 1.3257 | 0 | 0.0664 | PURINE NU | PURINE NUCLEOTIDE BIOSYNTHET |
| 242 | -2.0605 | 1.0157 | 0 | 0.451 | RHO GTPAS | RHO GTPASE EFFECTORS%REACTO |
| 143 | 2.1043 | 1.2736 | 0 | 0.0956 | ATP METAE | ATP METABOLIC PROCESS%GOBP% |
| 162 | -1.9576 | 1.2785 | 0 | 0.0956 | INTEGRIN S | INTEGRIN SIGNALLING PATHWAY% |
| 170 | 2.1967 | 1.5444 | 0 | 0 | ENERGY DE | ENERGY DERIVATION BY OXIDATIC |

| | | | | | | |
|---|---|---|---|---|---|---|
| 416 | 2.4231 | 1.4661 | 0 | 0.0011 | NUCLEOTID | NUCLEOTIDE METABOLIC PROCESS |
| 76 | 1.9337 | 1.2266 | 0 | 0.1491 | PURINE RIE | PURINE RIBONUCLEOSIDE MONOF |
| 481 | -2.409 | 0.6216 | 0 | 0.9956 | ACTIN FILA | ACTIN FILAMENT-BASED PROCESS |
| 101 | -1.7018 | 0.8606 | 0.0072 | 0.7041 | CHROMATI | CHROMATIN ASSEMBLY%GOBP%G |
| 197 | -1.8384 | 0.8545 | 0 | 0.7306 | HALLMARK | HALLMARK_APICAL_JUNCTION%M |

| | | | | | | |
|---|---|---|---|---|---|---|
| 49 | 1.8759 | 1.343 | 0 | 0.029 | COFACTOR | COFACTOR CATABOLIC PROCESS% |
| 94 | -2.0491 | -1.2528 | 0 | 0.203 | ACTOMYOS | ACTOMYOSIN STRUCTURE ORGAN |
| 195 | 1.8979 | 1.5065 | 0 | 0.0011 | HALLMARK | HALLMARK_MTORC1_SIGNALING% |
| 146 | 2.1894 | 1.3675 | 0 | 0.0226 | RIBOSE PH( | RIBOSE PHOSPHATE BIOSYNTHETI( |

| | | | | | | |
|---|---|---|---|---|---|---|
| 233 | 2.1616 | 1.4817 | 0 | 0.0045 | GENERATIC | GENERATION OF PRECURSOR MET |
| 155 | 2.0082 | 1.4301 | 0 | 0.0085 | CELLULAR I | CELLULAR LIPID CATABOLIC PROCE |
| 92 | -2.1078 | -1.3229 | 0 | 0.1837 | CELLULAR ( | CELLULAR COMPONENT ASSEMBL |
| 183 | -1.6292 | -0.9054 | 0.0143 | 0.5686 | REGULATIC | REGULATION OF CELL PROJECTION |

| | | | | | | |
|---|---|---|---|---|---|---|
| 88 | 1.9269 | 1.2391 | 0 | 0.1506 | NUCLEOSID | NUCLEOSIDE MONOPHOSPHATE B |
| 894 | 2.4498 | 1.5507 | 0 | 0 | LIPID META | LIPID METABOLIC PROCESS%GOBF |
| 45 | 1.8522 | 1.3525 | 0.0017 | 0.0315 | MITOCHON | MITOCHONDRIAL ATP SYNTHESIS ( |
| 192 | -2.1521 | -0.5032 | 0 | 1 | HALLMARK | HALLMARK_MITOTIC_SPINDLE%M |

| | | | | | | |
|---|---|---|---|---|---|---|
| 63 | -1.9396 | 0.8604 | 0 | 0.7114 | CYTOSKELE | CYTOSKELETAL REGULATION BY RH |
| 181 | -1.5986 | -0.9132 | 0.0121 | 0.6053 | REGULATIC | REGULATION OF PLASMA MEMBR |
| 54 | 1.8074 | 1.3318 | 0.0018 | 0.0391 | PROTON TI | PROTON TRANSMEMBRANE TRAN |
| 736 | 1.8837 | 1.3069 | 0 | 0.0328 | ORGANIC C | ORGANIC CYCLIC COMPOUND BIO |

| | | | | | | |
|---|---|---|---|---|---|---|
| 142 | -1.7722 | 1.3486 | 0 | 0.0401 | REGULATIC | REGULATION OF ACTIN FILAMENT |
| 985 | -1.6832 | -0.8798 | 0 | 0.9444 | ANIMAL OI | ANIMAL ORGAN MORPHOGENESIS |
| 340 | -1.6278 | 0.8064 | 0.0028 | 0.8123 | ESTABLISHI | ESTABLISHMENT OF ORGANELLE L |
| 107 | 1.9318 | 0.9689 | 0 | 0.5495 | PROTEIN F( | PROTEIN FOLDING%GOBP%GO:00 |
| 190 | 1.9754 | 1.2982 | 0 | 0.0672 | NUCLEOSIE | NUCLEOSIDE TRIPHOSPHATE MET/ |

| | | | | | | |
|---|---|---|---|---|---|---|
| 51 | 1.784 | 1.2382 | 0.0053 | 0.1292 | CHAPERON | CHAPERONE-MEDIATED PROTEIN |
| 909 | -1.8666 | -0.8885 | 0 | 0.8784 | CELL MOTI | CELL MOTILITY%GOBP%GO:00488 |
| 157 | -1.7628 | 0.9625 | 0 | 0.5465 | TRANSPOR | TRANSPORT TO THE GOLGI AND SU |

| | | | | | | |
|---|---|---|---|---|---|---|
| 190 | -1.7069 | -1.2106 | 0.0029 | 0.1197 | HALLMARK | HALLMARK_EPITHELIAL_MESENCH |
| 778 | -1.626 | 1.186 | 0 | 0.1656 | CELLULAR ( | CELLULAR COMPONENT MORPHO |
| 102 | 1.0629 | 1.4647 | 0.3629 | 0.0025 | TCR SIGNAI | TCR SIGNALING%REACTOME DATA |
| 171 | -1.8716 | 0.8093 | 0 | 0.7873 | CELL JUNCT | CELL JUNCTION ORGANIZATION%( |

| | | | | | | |
|---|---|---|---|---|---|---|
| 185 | 1.167 | 1.4537 | 0.1744 | 0 | HALLMARK | HALLMARK_MYC_TARGETS_V1%M |
| 67 | 1.7831 | 1.0864 | 0 | 0.3676 | PURINE NU | PURINE NUCLEOSIDE TRIPHOSPHA |
| 279 | 1.6422 | 1.1128 | 0.0053 | 0.3575 | POSITIVE R | POSITIVE REGULATION OF CYTOSC |
| 184 | -1.6654 | -0.8446 | 0.0092 | 0.6907 | RHYTHMIC | RHYTHMIC PROCESS%GOBP%GO:( |

| | | | | | | |
|---|---|---|---|---|---|---|
| 72 | 1.662 | 1.2317 | 0.0072 | 0.1565 | HALLMARK | HALLMARK_CHOLESTEROL_HOME |
| 76 | 2.2383 | 1.4303 | 0 | 0.0064 | ACYL-COA I | ACYL-COA METABOLIC PROCESS% |
| 279 | -2.0721 | -1.1265 | 0 | 0.1973 | EXTRACELL | EXTRACELLULAR MATRIX ORGANIZ |
| 118 | 1.7289 | 1.3472 | 0.0036 | 0.0268 | GLYCEROPI | GLYCEROPHOSPHOLIPID BIOSYNTH |

| | | | | | | |
|---|---|---|---|---|---|---|
| 315 | -1.9186 | 1.2157 | 0 | 0.1438 | REGULATIC | REGULATION OF ACTIN CYTOSKELI |
| 77 | 1.6097 | 1.4874 | 0.0079 | 0 | ER-PHAGOS | ER-PHAGOSOME PATHWAY%REAC |
| 163 | 2.0128 | 1.3107 | 0 | 0.0484 | PURINE RIE | PURINE RIBONUCLEOSIDE TRIPHOS |
| 337 | 1.6592 | 1.2897 | 0.0063 | 0.0929 | INORGANI( | INORGANIC CATION TRANSMEMBI |

| | | | | | | |
|---|---|---|---|---|---|---|
| 164 | -1.8635 | 1.2673 | 0 | 0.1069 | REGULATIC | REGULATION OF ACTIN FILAMENT |
| 411 | 1.5794 | 1.3331 | 0.0045 | 0.0311 | SMALL MO | SMALL MOLECULE BIOSYNTHETIC |
| 177 | 1.7365 | 1.3647 | 0.0126 | 0.0413 | STEROID M | STEROID METABOLIC PROCESS%G |
| 195 | 2.6164 | 1.448 | 0 | 0.0033 | HALLMARK | HALLMARK_ADIPOGENESIS%MSIG |
| 163 | 2.2514 | 1.4835 | 0 | 0.0024 | LIPID MOD | LIPID MODIFICATION%GOBP%GO: |

| | | | | | | |
|---|---|---|---|---|---|---|
| 822 | -1.7964 | 0.7367 | 0 | 0.9526 | ANATOMIC | ANATOMICAL STRUCTURE FORMA |
| 202 | -1.9383 | 1.0466 | 0 | 0.4138 | POSITIVE R | POSITIVE REGULATION OF CYTOSK |
| 286 | 1.0462 | 1.4631 | 0.4199 | 0.0023 | ANTIGEN P | ANTIGEN PROCESSING: UBIQUITIN |

| | | | | | | |
|---|---|---|---|---|---|---|
| 636 | 1.8236 | 1.2711 | 0 | 0.0457 | HETEROCY( | HETEROCYCLE BIOSYNTHETIC PRO |
| 156 | 2.2238 | 1.4839 | 0 | 0.0012 | THE CITRIC | THE CITRIC ACID (TCA) CYCLE AND |
| 178 | 2.1174 | 1.3542 | 0 | 0.0345 | NUCLEOSID | NUCLEOSIDE MONOPHOSPHATE M |
| 91 | 2.2756 | 1.4417 | 0 | 0.0065 | PURINE NU | PURINE NUCLEOSIDE BISPHOSPHA |

| | | | | | | |
|---|---|---|---|---|---|---|
| 115 | 2.3011 | 1.4863 | 0 | 0.0024 | PROTEIN L( | PROTEIN LOCALIZATION%REACTOI |
| 26 | -1.6879 | -0.9008 | 0.0115 | 0.5985 | NEGATIVE I | NEGATIVE REGULATION OF ACTIN |
| 27 | 1.9867 | 1.327 | 0 | 0.0297 | CITRATE M | CITRATE METABOLIC PROCESS%G( |
| 87 | 1.7819 | 0.7721 | 0.0018 | 0.7989 | ERYTHROC' | ERYTHROCYTE DIFFERENTIATION% |
| 75 | 1.3151 | 1.4556 | 0.1222 | 0.0027 | REGULATIC | REGULATION OF MITOCHONDRIAL |

| | | | | | | |
|---|---|---|---|---|---|---|
| 190 | 1.2322 | 1.53 | 0.1678 | 0 | REGULATIC | REGULATION OF REACTIVE OXYGE |
| 1039 | -1.8181 | 1.0151 | 0 | 0.4537 | CELL PROJE | CELL PROJECTION ORGANIZATION' |
| 69 | 1.628 | 1.3878 | 0.0085 | 0.0147 | ALPHA-AM | ALPHA-AMINO ACID CATABOLIC PI |
| 257 | 1.5878 | 1.3646 | 0.0262 | 0.0211 | PHOSPHOL | PHOSPHOLIPID METABOLIC PROCE |
| 218 | 1.7038 | 1.2104 | 0.0071 | 0.2164 | ORGANIC A | ORGANIC ACID TRANSPORT%GOBI |

| | | | | | | |
|---|---|---|---|---|---|---|
| 37 | 1.9499 | 1.292 | 0 | 0.092 | NAD META | NAD METABOLIC PROCESS%GOBP |
| 66 | -1.6601 | 0.9403 | 0.014 | 0.5979 | CDC42 SIGI | CDC42 SIGNALING EVENTS%PATH\ |
| 171 | -1.6675 | 1.1779 | 0.0152 | 0.256 | NEGATIVE I | NEGATIVE REGULATION OF NEUR( |
| 334 | -1.9678 | -1.2074 | 0 | 0.1556 | RESPONSE | RESPONSE TO WOUNDING%GOBP |

| | | | | | | |
|---|---|---|---|---|---|---|
| 95 | -1.8151 | 0.6219 | 0.0023 | 0.9627 | CELLULAR I | CELLULAR RESPONSE TO INTERFEF |
| 473 | -0.9299 | 1.4708 | 0.6493 | 0.0023 | CELL PART | CELL PART MORPHOGENESIS%GOI |
| 99 | -1.6845 | 0.8711 | 0.0119 | 0.7001 | SIGNALING | SIGNALING BY VEGF%REACTOME I |
| 166 | 2.2757 | 1.4722 | 0 | 0 | FATTY ACIE | FATTY ACID METABOLISM%REACT |
| 133 | -1.7415 | -1.0763 | 0.0024 | 0.3229 | REGULATIC | REGULATION OF CELL SHAPE%GOI |

| | | | | | | |
|---|---|---|---|---|---|---|
| 1071 | -1.8525 | 0.7963 | 0 | 0.8851 | LOCOMOTI | LOCOMOTION%GOBP%GO:00400 |
| 153 | -1.6166 | 0.835 | 0.0131 | 0.7562 | ANGIOGEN | ANGIOGENESIS%PANTHER PATHW |
| 411 | 1.868 | 1.2749 | 0 | 0.0958 | ANION TRA | ANION TRANSPORT%GOBP%GO:0 |
| 161 | -1.8455 | 1.269 | 0 | 0.094 | REGULATIC | REGULATION OF ACTIN POLYMERI |
| 65 | -1.6129 | -0.828 | 0.0186 | 0.6993 | NEGATIVE | NEGATIVE REGULATION OF PROTE |

| | | | | | | |
|---|---|---|---|---|---|---|
| 258 | 1.5612 | 1.2007 | 0.0095 | 0.1742 | ORGANELL | ORGANELLE BIOGENESIS AND MAI |
| 80 | 1.8223 | 1.1075 | 0 | 0.3292 | NUCLEOSID | NUCLEOSIDE TRIPHOSPHATE BIOS |
| 783 | 2.0106 | 1.2249 | 0 | 0.0735 | CARBOHYD | CARBOHYDRATE DERIVATIVE MET |
| 52 | -1.64 | 0.4171 | 0.0069 | 0.9987 | RHO GTPAS | RHO GTPASES ACTIVATE PKNS%RE |

| | | | | | | |
|---|---|---|---|---|---|---|
| 212 | -2.2415 | -1.0981 | 0 | 0.2353 | ACTIN FILA | ACTIN FILAMENT ORGANIZATION% |
| 54 | 1.8954 | 1.3576 | 0 | 0.0279 | OXIDATIVE | OXIDATIVE PHOSPHORYLATION%G |
| 134 | -1.7577 | 1.2283 | 0.0024 | 0.1598 | NEUROMU | NEUROMUSCULAR PROCESS%GOE |
| 87 | 1.836 | 1.5172 | 0 | 0 | DICARBOXY | DICARBOXYLIC ACID METABOLIC P |
| 290 | 1.7234 | 1.1312 | 0 | 0.2679 | PROTEIN H | PROTEIN HOMOOLIGOMERIZATIO |
| 233 | -2.1323 | -1.2912 | 0 | 0.1196 | WOUND HI | WOUND HEALING%GOBP%GO:004 |

| | | | | | | |
|---|---|---|---|---|---|---|
| 100 | 1.846 | 1.3556 | 0.0018 | 0.035 | REGULATIC | REGULATION OF STEROID METABC |
| 687 | 2.2894 | 1.4224 | 0 | 0.0032 | METABOLIS | METABOLISM OF LIPIDS%REACTOI |
| 201 | -1.7249 | 0.9868 | 0.0023 | 0.5063 | NEGATIVE I | NEGATIVE REGULATION OF CELL P |
| 65 | 1.7919 | 1.2292 | 0.0035 | 0.1667 | SULFUR CO | SULFUR COMPOUND BIOSYNTHET |
| 277 | 2.3682 | 1.5503 | 0 | 0 | FATTY ACIE | FATTY ACID METABOLIC PROCESS% |

| | | | | | | |
|---|---|---|---|---|---|---|
| 270 | 1.1636 | 1.4707 | 0.2237 | 0.0036 | PROTEASO | PROTEASOME-MEDIATED UBIQUIT |
| 565 | -1.6789 | 0.9053 | 0 | 0.7058 | POSITIVE R | POSITIVE REGULATION OF ORGAN |
| 68 | 2.1342 | 1.4838 | 0 | 0 | FATTY ACID | FATTY ACID OXIDATION%GOBP%G |
| 588 | -1.86 | 0.9883 | 0 | 0.5092 | CELL ADHE | CELL ADHESION%GOBP%GO:0007 |
| 195 | 2.0543 | 1.4564 | 0 | 0.0024 | LIPID CATA | LIPID CATABOLIC PROCESS%GOBP |

| | | | | | | |
|---|---|---|---|---|---|---|
| 93 | 1.6655 | 1.2503 | 0.0091 | 0.1509 | HORMONE | HORMONE-MEDIATED SIGNALING |
| 141 | 2.1367 | 1.3717 | 0 | 0.0245 | REGULATIC | REGULATION OF COLD-INDUCED T |

%GOBP%GO:0055114
GOBP%GO:0055085
OBP%GO:0006635
ELL PROJECTION ASSEMBLY%GOBP%GO:0120031
ABOLIC PROCESS%GOBP%GO:0033865
OBP%GO:0022900
TIC PROCESS%GOBP%GO:0090407
MERIZATION%GOBP%GO:1901879
ORT CHAIN%GOBP%GO:0022904
OBP%GO:0046942
%GOBP%GO:0006732
EACTOME%R-HSA-71403.2
N ORGANIZATION%GOBP%GO:0031344
0007623
O:0022610
PROCESS%GOBP%GO:0006163
ATE METABOLIC PROCESS%GOBP%GO:0009144

LIFERATION%GOBP%GO:0070663

ENESIS%GOBP%GO:0022604

GANIZATION%GOBP%GO:0007006

UM ION CONCENTRATION%GOBP%GO:0051480

ESS%GOBP%GO:0044255

ME DATABASE ID RELEASE 66%2682334

GO:0007160

BP%GO:0034101

ME DATABASE ID RELEASE 66%397014

ON%GOBP%GO:0051291

%MSIGDB_C2%NABA_MATRISOME_ASSOCIATED

RS%REACTOME DATABASE ID RELEASE 66%9006931

GOBP%GO:0070848

NT%GOBP%GO:0061061

THETIC PROCESS%GOBP%GO:1901293

METABOLIC PROCESS%GOBP%GO:0009199

PHATE METABOLIC PROCESS%GOBP%GO:0009126

SPHATE BIOSYNTHETIC PROCESS%GOBP%GO:0009206

GOBP%GO:0042692

CESS%GOBP%GO:0046395

6323

B_C2%NABA_ECM_REGULATORS

%GOBP%GO:0009062

4440

THETIC PROCESS%GOBP%GO:0009152

PROCESS%GOBP%GO:0006790

ORT%REACTOME DATABASE ID RELEASE 66%611105

;PORT%GOBP%GO:0098655

)RYLATION%MSIGDB_C2%HALLMARK_OXIDATIVE_PHOSPHORYLATION

LAR COMPONENT MOVEMENT%GOBP%GO:0051271

: DATABASE ID RELEASE 66%8949613

;GO:0048729

LIC PROCESS%GOBP%GO:0032787

PROCESS%GOBP%GO:0019637

P%GO:0006734

REACTOME DATABASE ID RELEASE 66%5358346

ɹT%REACTOME%R-HSA-1268020.3

ɪETABOLIC PROCESS%GOBP%GO:0045834

E TRANSPORT%GOBP%GO:0098660

OLIFERATION%GOBP%GO:0050670

%NABA_MATRISOME

60271

%GOBP%GO:0035383

_CYTOSKELETON_BY_RHO_GTPASES%MSIGDB_C2%SIG_REGULATION_OF_THE_ACTIN_CYTOSKELETON_

CHONDRION%GOBP%GO:0070585

IES%REACTOME DATABASE ID RELEASE 66%5628897

JLAR COMPONENT%GOBP%GO:0006928

BIOSYNTHETIC PROCESS%GOBP%GO:0009201

I%GOBP%GO:0007005

TOME%R-HSA-9020702.1

CE OF CELL POLARITY%GOBP%GO:0007163

%GOBP%GO:0016999

D:0009060

3P%GO:0006754

SENTATION%REACTOME DATABASE ID RELEASE 66%1236975

-BASED PROCESS%GOBP%GO:0032970

O:0000902

YNTHETIC PROCESS%GOBP%GO:1901137

N, EPIGENETIC%GOBP%GO:0040029

PROCESS%GOBP%GO:0072350

3LY%GOBP%GO:0051017

GROWTH%GOBP%GO:0061387

ALIZATION TO MITOCHONDRION%GOBP%GO:0072655

%GO:0034329

C2%PID_CDC42_PATHWAY

TION%REACTOME%R-HSA-2132295.3

OCESS%GOBP%GO:0044282

SS%GOBP%GO:0016054

GOBP%GO:0007010

RUCTURE SIZE%GOBP%GO:0090066

TION%GOBP%GO:0051259

SS%GOBP%GO:0046486

BP%GO:0030031

HETIC PROCESS%GOBP%GO:0019438

TIC PROCESS%GOBP%GO:0046890

E METABOLIC PROCESS%GOBP%GO:0062012

LIC PROCESS%GOBP%GO:0072329

PHOSPHATE METABOLIC PROCESS%GOBP%GO:0009167

;GO:0003007

) METABOLIC PROCESS%GOBP%GO:0072521

ENT%GOBP%GO:0030048

OLIC PROCESS%GOBP%GO:0006753

ANE BOUNDED CELL PROJECTION ORGANIZATION%GOBP%GO:0120035

GOBP%GO:0006090

BOLIC PROCESS%GOBP%GO:0072593

ATE BIOSYNTHETIC PROCESS%GOBP%GO:0009156

N TRANSPORT%GOBP%GO:0015672

CELL PROLIFERATION%GOBP%GO:0032944

EACTOME DATABASE ID RELEASE 66%5619084

ME%R-HSA-1474290.1

GOBP%GO:0007517

)CESS%GOBP%GO:0097164

TION%GOBP%GO:0030036

AR FIBER ORGANIZATION%GOBP%GO:1902903

OUND BIOSYNTHETIC PROCESS%GOBP%GO:0034654

PHATE BIOSYNTHETIC PROCESS%GOBP%GO:0009127

3P%GO:0051262

)B_C2%HALLMARK_PEROXISOME

IZATION%GOBP%GO:0061572

;GOBP%GO:0050770

ZATION%GOBP%GO:0097435

S FACTOR SUPERFAMILY CYTOKINE PRODUCTION%GOBP%GO:1903555

ATION%GOBP%GO:0042129

ATE METABOLIC PROCESS%GOBP%GO:0009161

DEPOLYMERIZATION%GOBP%GO:0030834

A-OXIDATION%REACTOME%R-HSA-77289.5

: PROCESS%GOBP%GO:0019216

/IA%GOBP%GO:0034341

_C2%PID_PDGFRB_PATHWAY

P%GO:0017144

EX DISASSEMBLY%GOBP%GO:0043244

OLIC PROCESS%GOBP%GO:0009150

S FACTOR PRODUCTION%GOBP%GO:0032680

ERIZATION%GOBP%GO:0032271

ORGANIZATION%GOBP%GO:0110053

C2%PID_CXCR4_PATHWAY

ISM%MSIGDB_C2%HALLMARK_BILE_ACID_METABOLISM

FILAMENT POLYMERIZATION%GOBP%GO:0030838

TIATION%GOBP%GO:0051146

'OME DATABASE ID RELEASE 66%4420097

)CYTE CELL-CELL ADHESION%GOBP%GO:1903038

ACTOME DATABASE ID RELEASE 66%373755

O:0051674

5811

ACTOME%R-HSA-1592230.2

BP%GO:0055001

EMOKINE AND CYTOKINE SIGNALING PATHWAY%PANTHER PATHWAY%P00031

FACTOR STIMULUS%GOBP%GO:0071363

CTOME%R-HSA-194315.4

PROCESS%GOBP%GO:0009260

E METABOLIC PROCESS%GOBP%GO:0010565

MOLECULAR FIBER ORGANIZATION%GOBP%GO:1902905

ORT, ATP SYNTHESIS BY CHEMIOSMOTIC COUPLING, AND HEAT PRODUCTION BY UNCOUPLING PROTEIN

OCESS%GOBP%GO:0019752

)048468

)RGANIZATION%GOBP%GO:0051493

SPORT%REACTOME%R-HSA-8856688.1

N%GOBP%GO:0051289

GOBP%GO:0043436

:TOME DATABASE ID RELEASE 66%8957322

;GOBP%GO:0051186

N%GOBP%GO:0001667

2%PID_RAC1_PATHWAY

Y%PATHWAY INTERACTION DATABASE NCI-NATURE CURATED DATA%PDGFR-BETA SIGNALING PATHWA

MENT%GOBP%GO:0055002

ELL PROJECTION ORGANIZATION%GOBP%GO:0120036

ROCESS%GOBP%GO:0016051

METABOLIC PROCESS%GOBP%GO:0033875

C PROCESS%GOBP%GO:0019362

ETABOLIC PROCESS%GOBP%GO:0006733

%GOBP%GO:0009636

0022402

)BP%GO:0008610

BROSIS%REACTOME DATABASE ID RELEASE 66%5678895

UM%GOBP%GO:0002009

ESS%GOBP%GO:0009165

ROCESS%GOBP%GO:0019693

:0060322

S%GOBP%GO:0051188

ABOLIC PROCESS%GOBP%GO:0046496

NDRION%GOBP%GO:0006626

NSPORT%REACTOME DATABASE ID RELEASE 66%199977

SS%GOBP%GO:0009108

P%GO:0031589

N POLYMERIZATION%GOBP%GO:0032273

BP%GO:0015711

;DB_C2%HALLMARK_COAGULATION

C PROCESS%GOBP%GO:0009063

)CESS%GOBP%GO:0009259

6477

RT%GOBP%GO:0034220

HETIC PROCESS%GOBP%GO:0072330

RANSPORT%REACTOME DATABASE ID RELEASE 66%6807878

%GO:0009617

NE TRANSPORT%GOBP%GO:1990542

GOBP%GO:0006006

) BIOSYNTHETIC PROCESS%GOBP%GO:0072522

TABOLISM%REACTOME%R-HSA-71291.4

)LISM%MSIGDB_C2%HALLMARK_FATTY_ACID_METABOLISM

. MOLECULE METABOLIC PROCESS%GOBP%GO:0055086

;GO:0050808

ESS%GOBP%GO:0006082

ONENT SIZE%GOBP%GO:0032535

RIC ACID (TCA) CYCLE%REACTOME%R-HSA-71406.1

ON TRANSPORT%GOBP%GO:0042773

OBP%GO:0006839

GO:0045333

ND METABOLIC PROCESS%GOBP%GO:0072524

'IC PROCESS%GOBP%GO:0006164

)ME DATABASE ID RELEASE 66%195258

%GO:0046034

%PANTHER PATHWAY%P00034

)N OF ORGANIC COMPOUNDS%GOBP%GO:0015980

S%GOBP%GO:0009117

PHOSPHATE BIOSYNTHETIC PROCESS%GOBP%GO:0009168

%GOBP%GO:0030029

GO:0031497

MSIGDB_C2%HALLMARK_APICAL_JUNCTION

GOBP%GO:0051187

IZATION%GOBP%GO:0031032

%MSIGDB_C2%HALLMARK_MTORC1_SIGNALING

C PROCESS%GOBP%GO:0046390

'ABOLITES AND ENERGY%GOBP%GO:0006091

ESS%GOBP%GO:0044242

Y INVOLVED IN MORPHOGENESIS%GOBP%GO:0010927

N ASSEMBLY%GOBP%GO:0060491

3IOSYNTHETIC PROCESS%GOBP%GO:0009124

P%GO:0006629

COUPLED ELECTRON TRANSPORT%GOBP%GO:0042775

ISIGDB_C2%HALLMARK_MITOTIC_SPINDLE

HO GTPASE%PANTHER PATHWAY%P00016

ANE BOUNDED CELL PROJECTION ASSEMBLY%GOBP%GO:0120032

SPORT%GOBP%GO:1902600

SYNTHETIC PROCESS%GOBP%GO:1901362

POLYMERIZATION%GOBP%GO:0030833

S%GOBP%GO:0009887

OCALIZATION%GOBP%GO:0051656

06457

ABOLIC PROCESS%GOBP%GO:0009141

FOLDING%GOBP%GO:0061077

70

JBSEQUENT MODIFICATION%REACTOME DATABASE ID RELEASE 66%948021 ɪYMAL_TRANSITION%MSIGDB_C2%HALLMARK_EPITHELIAL_MESENCHYMAL_TRANSITION

GENESIS%GOBP%GO:0032989

ABASE ID RELEASE 66%202403

GOBP%GO:0034330

MSIGDB_C2%HALLMARK_MYC_TARGETS_V1

ATE BIOSYNTHETIC PROCESS%GOBP%GO:0009145

OLIC CALCIUM ION CONCENTRATION%GOBP%GO:0007204

0048511

:OSTASIS%MSIGDB_C2%HALLMARK_CHOLESTEROL_HOMEOSTASIS

GOBP%GO:0006637

ZATION%REACTOME DATABASE ID RELEASE 66%1474244

HESIS%REACTOME DATABASE ID RELEASE 66%1483206

:ETON ORGANIZATION%GOBP%GO:0032956

:TOME%R-HSA-1236974.4

SPHATE METABOLIC PROCESS%GOBP%GO:0009205

RANE TRANSPORT%GOBP%GO:0098662

LENGTH%GOBP%GO:0030832

PROCESS%GOBP%GO:0044283

OBP%GO:0008202

DB_C2%HALLMARK_ADIPOGENESIS

0030258

ATION INVOLVED IN MORPHOGENESIS%GOBP%GO:0048646

ELETON ORGANIZATION%GOBP%GO:0051495

IATION & PROTEASOME DEGRADATION%REACTOME DATABASE ID RELEASE 66%983168

CESS%GOBP%GO:0018130

RESPIRATORY ELECTRON TRANSPORT%REACTOME%R-HSA-1428517.1

METABOLIC PROCESS%GOBP%GO:0009123

TE METABOLIC PROCESS%GOBP%GO:0034032

ME DATABASE ID RELEASE 66%9609507

FILAMENT DEPOLYMERIZATION%GOBP%GO:0030835

OBP%GO:0006101

GOBP%GO:0030218

MEMBRANE POTENTIAL%GOBP%GO:0051881

N SPECIES METABOLIC PROCESS%GOBP%GO:2000377

%GOBP%GO:0030030

ROCESS%GOBP%GO:1901606

ESS%GOBP%GO:0006644

P%GO:0015849

%GO:0019674

WAY INTERACTION DATABASE NCI-NATURE CURATED DATA%CDC42 SIGNALING EVENTS

ON PROJECTION DEVELOPMENT%GOBP%GO:0010977

%GO:0009611

RON-GAMMA%GOBP%GO:0071346

BP%GO:0032990

DATABASE ID RELEASE 66%194138

OME%R-HSA-8978868.4

BP%GO:0008360

11

/AY%P00005

006820

ZATION OR DEPOLYMERIZATION%GOBP%GO:0008064

EIN COMPLEX DISASSEMBLY%GOBP%GO:0043242

NTENANCE%REACTOME%R-HSA-1852241.4

YNTHETIC PROCESS%GOBP%GO:0009142

ABOLIC PROCESS%GOBP%GO:1901135

EACTOME%R-HSA-5625740.1

%GOBP%GO:0007015

GOBP%GO:0006119

BP%GO:0050905

ROCESS%GOBP%GO:0043648

N%GOBP%GO:0051260

42060

OLIC PROCESS%GOBP%GO:0019218

ME%R-HSA-556833.5

ROJECTION ORGANIZATION%GOBP%GO:0031345

IC PROCESS%GOBP%GO:0044272

%GOBP%GO:0006631

TIN-DEPENDENT PROTEIN CATABOLIC PROCESS%GOBP%GO:0043161

ELLE ORGANIZATION%GOBP%GO:0010638

;O:0019395

155

%GO:0016042

PATHWAY%GOBP%GO:0009755

'HERMOGENESIS%GOBP%GO:0120161

_BY_RHO_GTPASES

S.%REACTOME%R-HSA-163200.1

.Y

Normalized Log2 Intensity, Row Z Score

| FEATURE | TPHP_1 | TPHP_2 | TPHP_3 | VEHICLE_1 | VEHICLE_2 | VEHICLE_3 | ROSI_1 | ROSI_2 | ROSI_3 | GENE |
|---|---|---|---|---|---|---|---|---|---|---|
| Cd36_Q08857 | -0.5 | -0.4 | -0.5 | -0.7 | -0.8 | -0.9 | 1.23 | 1.54 | 1.17 | Cd36 |
| Prps1_Q9D7G0 | -0.2 | -0.2 | -0.4 | -1.2 | -0.8 | -1 | 1.34 | 1.49 | 0.89 | Prps1 |
| Cdh11_P55288 | -0.2 | -0.3 | -0.4 | -1.1 | -0.6 | -1.1 | 1.45 | 1.34 | 0.97 | Cdh11 |
| Nsdhl_Q9R1J0 | -0.2 | -0.4 | -0.5 | -1.1 | -0.6 | -1 | 1.46 | 1.37 | 1 | Nsdhl |
| Acsl1_P41216 | -0.2 | -0.5 | -0.6 | -0.9 | -0.7 | -0.9 | 1.49 | 1.42 | 0.94 | Acsl1 |
| Fabp4_P04117 | -0.4 | -0.4 | -0.5 | -1.1 | -0.8 | -0.6 | 1.38 | 1.52 | 0.96 | Fabp4 |
| Fabp5_Q05816 | -0.2 | -0.4 | -0.4 | -1.2 | -0.9 | -0.8 | 1.52 | 1.37 | 0.88 | Fabp5 |
| Pc_Q05920 | -0.2 | -0.4 | -0.5 | -1.1 | -0.8 | -0.8 | 1.38 | 1.51 | 0.91 | Pc |
| Gpd1_P13707 | -0.3 | -0.3 | -0.7 | -1 | -0.7 | -0.7 | 1.54 | 1.37 | 0.96 | Gpd1 |
| Mpc2_Q9D023 | -0.2 | 0.11 | -0.2 | -1 | -1.3 | -0.9 | 1.12 | 1.58 | 0.77 | Mpc2 |
| Ppa1_Q9D819 | -0.4 | -0.4 | -0.6 | -0.8 | -1 | -0.7 | 1.31 | 1.64 | 0.9 | Ppa1 |
| Acadvl_P50544 | -0.3 | -0.4 | -0.8 | -0.8 | -0.9 | -0.7 | 1.5 | 1.3 | 1.09 | Acadvl |
| Grpel1_Q99LP6 | -0.5 | -0.5 | -0.7 | -1 | -0.4 | -0.9 | 1.44 | 1.51 | 0.92 | Grpel1 |
| Uqcrh_P99028 | -0.5 | -0.2 | -0.4 | -0.4 | -1.5 | -0.7 | 1.21 | 1.41 | 1.11 | Uqcrh |
| Adipoq_Q60994 | -0.3 | -0.4 | -0.6 | -0.7 | -0.7 | -1.1 | 1.7 | 1.34 | 0.74 | Adipoq |
| Glul_P15105 | -0.4 | 0.13 | 0.19 | -1.3 | -1.1 | -0.9 | 1.22 | 1.44 | 0.72 | Glul |
| Fech_P22315 | -0.1 | -0.4 | -0.5 | -0.8 | -0.7 | -1.2 | 1.56 | 1.47 | 0.65 | Fech |
| Etfa_Q99LC5 | -0.5 | -0.1 | 0.25 | -1.3 | -1.1 | -0.8 | 1.38 | 1.17 | 0.97 | Etfa |
| Marc2_Q922Q1 | -0.5 | -0.1 | -0.1 | -0.6 | -1.5 | -0.8 | 1.21 | 1.49 | 0.93 | Marc2 |
| Hspa9_P38647 | -0.8 | -0.1 | -0.2 | -0.9 | -1 | -0.8 | 1.15 | 1.45 | 1.22 | Hspa9 |
| Cs_Q9CZU6 | -0 | -0.5 | -0.5 | -0.9 | -1.1 | -0.7 | 1.46 | 1.51 | 0.76 | Cs |
| Idh3a_Q9D6R2 | -0.1 | -0.6 | -0.4 | -0.7 | -0.7 | -1.3 | 1.27 | 1.58 | 0.87 | Idh3a |
| Cyc1_Q9D0M3 | 0.43 | -0 | 0.14 | -1 | -0.8 | -1.8 | 1.2 | 1.06 | 0.72 | Cyc1 |
| Atp5c1_Q91VR2 | -0.1 | 0.02 | -0.3 | -0.8 | -1.2 | -1.2 | 1.23 | 1.36 | 1.01 | Atp5c1 |
| Hspd1_P63038 | 0.1 | -0.4 | -0.4 | -1.2 | -0.8 | -0.9 | 1.5 | 1.45 | 0.64 | Hspd1 |
| Fdps_Q920E5 | -0.6 | 0.03 | -0.1 | -0.5 | -1.6 | -0.8 | 1.26 | 1.3 | 0.99 | Fdps |
| Acadm_P45952 | -0.2 | -0.4 | -0.2 | -1.3 | -0.6 | -0.9 | 1.07 | 1.78 | 0.73 | Acadm |
| Mdh2_P08249 | -0.4 | -0.2 | -0.2 | -0.9 | -0.9 | -1 | 1.31 | 1.75 | 0.56 | Mdh2 |
| Ephx2_P34914 | -0.4 | -0.7 | -0.5 | -0.4 | -0.8 | -1 | 1.41 | 1.63 | 0.78 | Ephx2 |
| Hsd17b12_O70503 | -0.6 | -0.9 | -0.2 | -0.6 | -0.7 | -0.8 | 1.45 | 1.58 | 0.81 | Hsd17b12 |
| Prdx3_P20108 | -0.2 | -0.3 | 0.13 | -1 | -0.5 | -1.6 | 1.26 | 1.19 | 1.03 | Prdx3 |
| Phb_P67778 | -0.4 | -0.2 | 0.09 | -0.7 | -1.4 | -0.9 | 0.77 | 1.42 | 1.36 | Phb |
| Hadhb_Q99JY0 | 0.18 | -0.3 | -0.4 | -0.9 | -0.7 | -1.4 | 1.61 | 1.1 | 0.85 | Hadhb |
| Mgst1_Q91VS7 | -0.8 | -0.1 | -0 | -0.5 | -1.5 | -0.7 | 0.89 | 1.5 | 1.17 | Mgst1 |
| Dnaja4_Q9JMC3 | -1.1 | 0.25 | -0 | -1 | -0.9 | -0.8 | 1.24 | 0.98 | 1.34 | Dnaja4 |
| Coq9_Q8K1Z0 | -0.4 | -0.5 | -0.7 | -0.7 | -1.3 | -0.2 | 1.06 | 1.37 | 1.38 | Coq9 |
| Ttn_A2ASS6 | 0.83 | 0.25 | 0.47 | 0.56 | 0.79 | 0.88 | -0.7 | -1.7 | -1.5 | Ttn |
| Dbi_P31786 | 0.18 | -0.4 | -0.7 | -1.2 | -0.3 | -1.1 | 1.37 | 1.36 | 0.85 | Dbi |
| Slc16a1_P53986 | 0.32 | 0.01 | 0.06 | 1.26 | 0.59 | 1.16 | -1.8 | -0.6 | -1 | Slc16a1 |
| Ahcyl2_Q68FL4 | -1.1 | -0.3 | -0.1 | -0.4 | -1.1 | -0.7 | 1.03 | 1.37 | 1.34 | Ahcyl2 |
| Fn1_P11276 | 0.84 | 0.47 | 0.29 | 0.58 | 1.15 | 0.47 | -1.2 | -1.7 | -0.9 | Fn1 |
| Slc25a1_Q8JZU2 | -1.1 | -0.1 | -0.4 | -0.4 | -1.1 | -0.7 | 1.52 | 1.35 | 0.87 | Slc25a1 |

| | | | | | | | | | | |
|---|---|---|---|---|---|---|---|---|---|---|
| Decr1_Q9CQ62 | 0.41 | -0.5 | -0.1 | -0.7 | -0.8 | -1.6 | 1.32 | 1.26 | 0.74 | Decr1 |
| Slc25a10_Q9QZD8 | -0.7 | -0.2 | -0.3 | -0.3 | -0.8 | -1.3 | 1.56 | 1.42 | 0.67 | Slc25a10 |
| Bcap31_Q61335 | -0.3 | -0.1 | 0.05 | -1.6 | -1.1 | -0.5 | 1.07 | 1.44 | 0.91 | Bcap31 |
| Fam129b_Q8R1F1 | 0.36 | 0.07 | -0.1 | 0.52 | 1.37 | 1.15 | -1.7 | -0.6 | -1.1 | Fam129b |
| Plin1_Q8CGN5 | 0.34 | 0.35 | -0.5 | -0.6 | -1.4 | -1.3 | 1.23 | 1.25 | 0.6 | Plin1 |
| Prdx1_P35700 | -0.4 | 0.1 | -0.2 | -1 | -1 | -1.2 | 1.11 | 1.42 | 1.09 | Prdx1 |
| Atp5a1_Q03265 | 0.14 | -0.3 | -0.5 | -1.1 | -0.8 | -1 | 0.96 | 1.6 | 1.08 | Atp5a1 |
| Bcl2l13_P59017 | -0.4 | 0.07 | -0.3 | -0.3 | -1.8 | -0.6 | 1.13 | 1.22 | 1.07 | Bcl2l13 |
| Foxn4_Q8K3Q3 | 0.66 | 0.25 | 0.63 | 0.39 | 0.39 | 1.39 | -0.8 | -1.4 | -1.5 | Foxn4 |
| Ppic_P30412 | 0.98 | -0.3 | 0.47 | 0.94 | 0.85 | 0.74 | -1.3 | -1.2 | -1.2 | Ppic |
| Hcfc1_Q61191 | -0.6 | -0.4 | 0.3 | -1.6 | -0.8 | -0.3 | 0.93 | 1.2 | 1.31 | Hcfc1 |
| Plin4_O88492 | -0.1 | -0.9 | -1 | -1.1 | -0.3 | -0.4 | 1.2 | 1.56 | 0.98 | Plin4 |
| Opa1_P58281 | 0.46 | -0.6 | 0.43 | -1.1 | -0.9 | -1.3 | 1.69 | 0.46 | 0.77 | Opa1 |
| Caprin1_Q60865 | 0.43 | -0.3 | 0.24 | 0.86 | 0.7 | 1.42 | -0.7 | -0.9 | -1.8 | Caprin1 |
| Aldh18a1_Q9Z110 | 1.1 | -0 | -0.1 | 0.58 | 0.91 | 1.1 | -0.8 | -1.5 | -1.2 | Aldh18a1 |
| Aldoa_P05064 | 0.38 | -0.4 | -0.9 | -0.9 | -0.7 | -1 | 1.5 | 1.37 | 0.67 | Aldoa |
| Actn1_Q7TPR4 | 0.91 | 0.09 | 0.12 | 0.57 | 1.2 | 0.79 | -1.3 | -1.6 | -0.8 | Actn1 |
| Serpinc1_P32261 | 0.67 | -0.4 | -0.2 | 1.17 | 0.84 | 1.15 | -1.6 | -0.9 | -0.8 | Serpinc1 |
| Prkag1_O54950 | 0.07 | -0.8 | 0.59 | -1.3 | -1.3 | -0.5 | 1 | 1.22 | 0.96 | Prkag1 |
| Letm1_Q9Z2I0 | 0.7 | -0.6 | -0.2 | -0.6 | -0.9 | -1.5 | 0.77 | 1.46 | 0.95 | Letm1 |
| Dld_O08749 | -0.1 | 0.31 | -0.4 | -1.1 | -1.5 | -0.4 | 1.11 | 1.32 | 0.95 | Dld |
| Acadl_P51174 | 0.39 | -0.2 | -0.5 | -1.4 | -0.3 | -1.3 | 1.32 | 1.15 | 0.83 | Acadl |
| Acat1_Q8QZT1 | -0.2 | -0.1 | -0.1 | -1.4 | -0.6 | -1.1 | 1.38 | 1.3 | 0.9 | Acat1 |
| Slc25a4_P48962 | 0.33 | -0.4 | -0.7 | -1.1 | -0.5 | -1.2 | 1.33 | 1.35 | 0.88 | Slc25a4 |
| Taldo1_Q93092 | -0.1 | -1 | -0.1 | -0.6 | -0.4 | -1.4 | 1.3 | 1.46 | 0.84 | Taldo1 |
| Alad_P10518 | -0.3 | -0 | 0.17 | -0.6 | -1.1 | -1.4 | 1.87 | 0.95 | 0.37 | Alad |
| Tax1bp3_Q9DBG9 | -1.3 | 0.17 | -0.3 | -0.7 | -1.1 | -0.4 | 1.1 | 1.38 | 1.12 | Tax1bp3 |
| Ppil1_Q9D0W5 | 0.4 | -0.1 | -0.2 | 0.44 | 0.86 | 1.68 | -0.5 | -0.9 | -1.7 | Ppil1 |
| Pdhx_Q8BKZ9 | -0.2 | 0.37 | -0.5 | -0.6 | -1.5 | -0.9 | 0.58 | 1.67 | 1.05 | Pdhx |
| Nnt_Q61941 | 0.93 | 0.06 | 0.11 | 0.56 | 0.66 | 1.34 | -1 | -1.2 | -1.4 | Nnt |
| Timm44_O35857 | -0.2 | -0.5 | -0.4 | -1.4 | -1 | 0.02 | 1.51 | 0.92 | 1.14 | Timm44 |
| Sorcs2_Q9EPR5 | 1.09 | 0.53 | 0.41 | 0.14 | 0.93 | 0.56 | -1.7 | -1.5 | -0.5 | Sorcs2 |
| P4ha2_Q60716 | 1.14 | 0.79 | -0 | 0.71 | 0.25 | 0.83 | -1 | -1.1 | -1.6 | P4ha2 |
| Rps24_P62849 | 0.57 | 0.06 | 0.15 | 0.61 | 1.53 | 0.5 | -1.2 | -0.5 | -1.7 | Rps24 |
| Cnn2_Q08093 | 0.92 | 0.97 | 0.49 | 0.56 | 0.85 | 0.01 | -1.5 | -1.3 | -1 | Cnn2 |
| Myl1_P05977 | 1.27 | 0.63 | 0.2 | 0.4 | 0.37 | 0.82 | -1.3 | -1.7 | -0.8 | Myl1 |
| Calr_P14211 | -0.8 | 0.26 | 0.3 | -0.6 | -1.3 | -1.2 | 1.05 | 1.46 | 0.79 | Calr |
| Loxhd1_C8YR32 | 0.36 | -0.6 | -0.8 | -0.3 | -0.7 | -1.3 | 1.73 | 0.76 | 0.97 | Loxhd1 |
| Bcat2_O35855 | 0.86 | 0.36 | -0.5 | -0.4 | -1.2 | -1.7 | 1.04 | 1.01 | 0.52 | Bcat2 |
| Mrc2_Q64449 | -0.4 | 0.98 | 0.45 | 0.62 | 0.92 | 0.97 | -1.5 | -1.3 | -0.7 | Mrc2 |
| Man2b1_O09159 | 0.67 | -0.2 | -0.2 | 0.99 | 1.22 | 0.91 | -1.4 | -1.3 | -0.6 | Man2b1 |
| Otud6a_Q6IE21 | 0.96 | -0.8 | -0.3 | 0.86 | 0.85 | 1.36 | -1 | -0.7 | -1.3 | Otud6a |
| Cpt2_P52825 | 0.23 | -0.6 | -0.7 | -0.9 | -1.4 | -0.2 | 0.85 | 1.58 | 1.07 | Cpt2 |
| Immt_Q8CAQ8 | -1 | 0.14 | 0.32 | -0.6 | -1.6 | -0.5 | 1.07 | 1.25 | 0.95 | Immt |
| Atp5d_Q9D3D9 | -0.1 | -0.1 | -0.4 | -0.7 | -0.1 | -1.9 | 1.27 | 1.24 | 0.81 | Atp5d |
| Oxsr1_Q6P9R2 | -0.5 | 0.66 | 0.69 | 0.82 | 0.56 | 1.21 | -1.6 | -1.1 | -0.7 | Oxsr1 |
| Lpl_P11152 | -0.3 | -1.1 | -0.8 | -0.9 | -0.3 | -0.2 | 1.81 | 1.25 | 0.58 | Lpl |
| Dpysl3_Q62188 | 1.13 | -0.1 | 0.42 | 0.55 | 1.19 | 0.45 | -1.2 | -1.5 | -0.9 | Dpysl3 |
| Idh3g_P70404 | -0.6 | -1.1 | 0.08 | -0.6 | -1.2 | -0.2 | 1.11 | 1.55 | 0.95 | Idh3g |
| Tmpo_Q61029 | -0.4 | 0.41 | -0.1 | 0.66 | 0.65 | 1.87 | -0.9 | -0.8 | -1.3 | Tmpo |

| | | | | | | | | | | |
|---|---|---|---|---|---|---|---|---|---|---|
| Hadh_Q61425 | -0.4 | 0.57 | -0.3 | -1.4 | -0.8 | -0.8 | 1.49 | 1.36 | 0.3 | Hadh |
| Mtap_Q9CQ65 | -0.1 | -0.1 | 0.38 | 1.36 | 1.2 | 0.49 | -0.6 | -1.8 | -0.9 | Mtap |
| Suclg1_Q9WUM5 | -0.9 | 0.25 | -0.5 | -1.1 | -1 | -0.4 | 1.25 | 1.63 | 0.66 | Suclg1 |
| Marcks_P26645 | 0.87 | 0.9 | 0.66 | 0.72 | 0.67 | -0.1 | -1.3 | -1.6 | -0.8 | Marcks |
| Pdia4_P08003 | -0.1 | -0.2 | -0.1 | -1.3 | -0.7 | -1.2 | 1.44 | 1.37 | 0.71 | Pdia4 |
| Lasp1_Q61792 | 0.28 | 0.34 | 0.21 | 1 | 0.13 | 1.56 | -1.4 | -1.3 | -0.8 | Lasp1 |
| Flna_Q8BTM8 | 0.57 | 0.92 | 0.55 | 1 | 0.12 | 0.66 | -1.3 | -1.6 | -0.9 | Flna |
| Nudt9_Q8BVU5 | -1.3 | -0.1 | -0.4 | -0.3 | -1.1 | -0.4 | 0.72 | 1.57 | 1.3 | Nudt9 |
| Sfxn1_Q99JR1 | -0.5 | -0.9 | -1.1 | -0.7 | -0.3 | -0.2 | 1.27 | 1.82 | 0.55 | Sfxn1 |
| Pdha1_P35486 | -0.5 | -1.2 | -0.2 | -1 | -0.5 | -0.3 | 1.24 | 1.27 | 1.29 | Pdha1 |
| Timp2_P25785 | 1.09 | -0.6 | -0.2 | 0.68 | 1.29 | 0.87 | -0.8 | -0.9 | -1.4 | Timp2 |
| Hspa5_P20029 | -0.4 | -0.7 | -0.4 | -0.8 | -0.7 | -0.8 | 1.58 | 1.59 | 0.62 | Hspa5 |
| Crtap_Q9CYD3 | 0.63 | -0.4 | -0.5 | 1.06 | 1.04 | 1.18 | -0.9 | -1.6 | -0.5 | Crtap |
| Cyb5b_Q9CQX2 | -0.7 | 0.75 | 0.18 | -0.6 | -1.3 | -1.3 | 0.97 | 0.77 | 1.26 | Cyb5b |
| Lpp_Q8BFW7 | -0.4 | 0.83 | -0.4 | 1.4 | 0.44 | 1.17 | -1.2 | -1.4 | -0.6 | Lpp |
| Slc25a5_P51881 | 0.53 | -0.8 | -0.6 | -1.3 | -0.2 | -1 | 1.27 | 1.37 | 0.7 | Slc25a5 |
| Psmb6_Q60692 | -0.4 | -0.2 | -0.4 | -0.2 | -0.5 | -1.8 | 1.08 | 1.63 | 0.67 | Psmb6 |
| Tpp1_O89023 | 0.83 | 0.27 | 0.25 | 0.63 | 0.34 | 1.29 | -0.5 | -1.5 | -1.6 | Tpp1 |
| Kras_P32883 | 1.5 | 0.05 | -0.1 | 0.79 | 0.87 | 0.39 | -1 | -1.5 | -1 | Kras |
| Tpm1_P58771 | 0.28 | 0.86 | 0.58 | 0.96 | 0.71 | 0.19 | -1.5 | -1.7 | -0.3 | Tpm1 |
| Actg1_P63260 | 1.03 | -0.4 | -0.3 | 0.7 | 0.99 | 1.05 | -0.8 | -0.4 | -1.8 | Actg1 |
| Nomo1_Q6GQT9 | -0.5 | -0.1 | 0.25 | -1.5 | -1.2 | -0 | 0.63 | 0.64 | 1.82 | Nomo1 |
| Ephx1_Q9D379 | 1.09 | 1.07 | 0.91 | 0.2 | -0.1 | 0.51 | -1.3 | -1.3 | -1 | Ephx1 |
| Eftud2_O08810 | 0.8 | -0.6 | -0.1 | 1.12 | 1.41 | 0.51 | -0.8 | -1.2 | -1.2 | Eftud2 |
| Myof_Q69ZN7 | 1 | 0.1 | 0.3 | 0.17 | 1.44 | 0.56 | -1 | -1.6 | -1 | Myof |
| Tuba1b_P05213 | 0.32 | -0.3 | -0.7 | -0.7 | -0.5 | -1.5 | 0.55 | 1.79 | 0.96 | Tuba1b |
| Mgst3_Q9CPU4 | 0.05 | -1.2 | -0.3 | -1.5 | -0.4 | -0 | 1.26 | 1.25 | 0.89 | Mgst3 |
| Aldh3a1_P47739 | 0.73 | 0.33 | -0.1 | 0.24 | 0.46 | 1.65 | -0.7 | -1.7 | -1 | Aldh3a1 |
| Fasn_P19096 | 0.22 | -0.7 | -0.6 | -1 | -0.4 | -0.9 | 1.6 | 1.54 | 0.3 | Fasn |
| Csnk2a1_Q60737 | -0.4 | -1.3 | -0.1 | -0.4 | -0.6 | -0.8 | 1.85 | 1.03 | 0.7 | Csnk2a1 |
| Tspo_P50637 | -0.5 | 0.74 | 0.41 | -0.3 | -1.8 | -1.1 | 0.31 | 1.06 | 1.14 | Tspo |
| Prkar2b_P31324 | -0.7 | 0.27 | -0.1 | -0.8 | -1.4 | -0.4 | 1.77 | 1.17 | 0.28 | Prkar2b |
| S100a6_P14069 | -1.4 | 0.15 | 0.49 | -0.6 | -0.8 | -1 | 0.78 | 0.77 | 1.61 | S100a6 |
| Ass1_P16460 | 1.02 | 0.12 | -0 | 0.62 | 1.03 | 0.56 | -1.2 | -0.2 | -1.9 | Ass1 |
| Hprt1_P00493 | -0.8 | -0.1 | -0.5 | -0.3 | -0 | -1.6 | 1.32 | 1.5 | 0.55 | Hprt1 |
| Igfbp4_P47879 | -1.1 | -1.2 | -1.2 | 0.6 | -0.1 | -0.2 | 1.37 | 0.72 | 1.1 | Igfbp4 |
| Tpm3_P21107 | -0.6 | 0.41 | 0.26 | 1.35 | 0.83 | 0.97 | -1.6 | -0.6 | -1 | Tpm3 |
| Psmd14_O35593 | 0.05 | -0.3 | -1 | -0.8 | -0.1 | -1.4 | 1.35 | 1.3 | 0.87 | Psmd14 |
| Zyx_Q62523 | 1.12 | 0.4 | 1.09 | 0.26 | 0.96 | -0.3 | -1.2 | -1.4 | -0.9 | Zyx |
| Pdhb_Q9D051 | 0.4 | -0.6 | -0.4 | -1.7 | -0.4 | -0.4 | 1.41 | 1.4 | 0.31 | Pdhb |
| Hibadh_Q99L13 | 0.76 | -0.9 | -0.8 | -0.8 | -0.4 | -1.1 | 1.36 | 1.28 | 0.64 | Hibadh |
| Rpl27a_P14115 | 0.12 | -0.1 | -0.3 | 0.74 | 1.42 | 0.86 | -0.3 | -0.3 | -2 | Rpl27a |
| Fh_P97807 | 0.27 | -0.5 | -0.6 | -1.2 | -0.5 | -1 | 1.41 | 1.59 | 0.46 | Fh |
| Anxa1_P10107 | 1.38 | 0.09 | 0.15 | 0.29 | 1.37 | 0.19 | -1 | -1.4 | -1.1 | Anxa1 |
| Aldh9a1_Q9JLJ2 | -0.8 | -0.2 | -0.3 | -0.7 | -1.1 | -0.7 | 1.14 | 1.74 | 0.9 | Aldh9a1 |
| Vta1_Q9CR26 | 0.41 | 0.97 | -0.6 | 0.85 | 1.05 | 0.47 | -1.1 | -1.8 | -0.2 | Vta1 |
| Cat_P24270 | -0.9 | -0.2 | 0.01 | -1.3 | -0.6 | -0.5 | 1.65 | 1.41 | 0.41 | Cat |
| Ldhb_P16125 | 0.54 | -1 | -0.9 | -0.9 | -0.4 | -0.7 | 1.47 | 1.48 | 0.4 | Ldhb |
| Adh7_Q64437 | 0.41 | 1.03 | 0.72 | 0.55 | 0.06 | 0.98 | -1.7 | -1.2 | -0.9 | Adh7 |
| Rhoc_Q62159 | 1.26 | 0.45 | 0.29 | 0.41 | 0.52 | 0.84 | -1.1 | -1.7 | -0.9 | Rhoc |

| | | | | | | | | | | |
|---|---|---|---|---|---|---|---|---|---|---|
| Atp5b_P56480 | 0.74 | -0.5 | -0.8 | -1.1 | -0.8 | -1 | 1.36 | 1 | 1 | Atp5b |
| Dctn2_Q99KJ8 | 0.68 | 0.3 | 0.44 | 0.31 | 0.75 | 0.93 | -0 | -1.8 | -1.6 | Dctn2 |
| Actc1_P68033 | -0.2 | 1.19 | 0.83 | 0.44 | 0.86 | 0.4 | -0.7 | -1.8 | -1 | Actc1 |
| Snx3_O70492 | 0.2 | -0.4 | -0.5 | 0.98 | 1.85 | 0.49 | -0.7 | -1.5 | -0.3 | Snx3 |
| Cstb_Q62426 | 1.28 | -0.3 | 0.01 | 0.57 | 1.02 | 0.88 | -1.2 | -1.4 | -0.8 | Cstb |
| Col1a1_P11087 | -0.1 | 0.83 | 0.58 | 1.3 | 0.45 | 0.53 | -1.4 | -1.5 | -0.6 | Col1a1 |
| Atp5i_Q06185 | 0.83 | -0.9 | -0.4 | -1.3 | -0.3 | -1 | 1.22 | 1.34 | 0.51 | Atp5i |
| Eif4e_P63073 | 1.1 | -0 | 0.2 | -0.7 | -2 | -0.8 | 0.7 | 0.88 | 0.62 | Eif4e |
| Bgn_P28653 | 0.1 | 0.08 | -0.5 | 1.41 | 1.49 | 0.3 | -1.6 | -0.8 | -0.5 | Bgn |
| Cdc42_P60766 | 0.38 | 0.01 | 0.03 | 0.64 | 1.3 | 0.97 | -0.9 | -1.9 | -0.5 | Cdc42 |
| Lpcat3_Q91V01 | 0.1 | -0.9 | 0.89 | -0.3 | -1.1 | -1.4 | 1.21 | 0.39 | 1.17 | Lpcat3 |
| Rbmx2_Q8R0F5 | -0.3 | 0.22 | 1.17 | 0.46 | 0.18 | 1.55 | -0.7 | -1.5 | -1.1 | Rbmx2 |
| Pygb_Q8CI94 | 0.28 | 0.27 | 0.1 | 1.21 | 0.52 | 1.13 | -0.9 | -0.7 | -1.8 | Pygb |
| Sec13_Q9D1M0 | 1.07 | 0.8 | 1.53 | 0.03 | 0.11 | -0.3 | -1.4 | -0.8 | -1.1 | Sec13 |
| Thy1_P01831 | 0.71 | -0.7 | -0.2 | 0.76 | 1.55 | 0.86 | -1.5 | -0.9 | -0.7 | Thy1 |
| Tufm_Q8BFR5 | 0.46 | -0.5 | -0.2 | -1.3 | -0.4 | -1.4 | 1.43 | 1.3 | 0.47 | Tufm |
| Wdr1_O88342 | 1.01 | -0.3 | -0 | 0.41 | 0.94 | 1.29 | -0.7 | -1.7 | -1 | Wdr1 |
| Slc25a3_Q8VEM8 | 0.78 | -0.5 | -0.5 | -1.1 | 0.01 | -1.6 | 1.06 | 1.29 | 0.55 | Slc25a3 |
| Tagln_P37804 | 0.23 | 0.43 | 0.55 | 1.26 | -0.5 | 1.35 | -1 | -0.9 | -1.4 | Tagln |
| Ptges3_Q9R0Q7 | -0 | 0.88 | 0.47 | -1.7 | -1.7 | 0.01 | 0.64 | 0.54 | 0.82 | Ptges3 |
| Snx5_Q9D8U8 | -0.3 | -0.8 | 0.25 | 1.07 | 1.34 | 0.97 | -1.7 | -0.2 | -0.6 | Snx5 |
| Plxnb2_B2RXS4 | 1.28 | 0.53 | -0.1 | 1.06 | 0.6 | 0.1 | -1.8 | -1 | -0.7 | Plxnb2 |
| Pdxk_Q8K183 | -0.6 | 0.48 | 0.64 | 1.75 | 0.32 | 0.6 | -1.2 | -1.1 | -0.8 | Pdxk |
| Dlst_Q9D2G2 | -1.2 | 0.24 | 0.26 | -0.1 | -1.8 | -0.5 | 0.87 | 1.17 | 0.96 | Dlst |
| Pmm2_Q9Z2M7 | 1.24 | 0.16 | -0.5 | 0.37 | 0.78 | 1.08 | -1 | -1.8 | -0.4 | Pmm2 |
| Cttn_Q60598 | 0.87 | 0.12 | 0.1 | 0.2 | 1.5 | 0.79 | -1.1 | -1.4 | -1.1 | Cttn |
| Rps10_P63325 | -1.4 | -0 | 0.35 | -0.5 | -1.4 | -0.5 | 1.04 | 1.13 | 1.17 | Rps10 |
| Vps35_Q9EQH3 | 0.71 | -0.5 | 0.31 | 0.43 | 1.79 | 0.48 | -0.9 | -1 | -1.3 | Vps35 |
| Flnb_Q80X90 | 1.29 | -0 | -0.2 | 0.49 | 1.35 | 0.41 | -0.6 | -1.4 | -1.3 | Flnb |
| Myadm_O35682 | 0.4 | 0.73 | 0.66 | -1.8 | 0.16 | -1.7 | 0.19 | 0.58 | 0.72 | Myadm |
| Atp5f1_Q9CQQ7 | -1 | -0.3 | 0.21 | 0.09 | -1.5 | -0.7 | 0.56 | 1.19 | 1.51 | Atp5f1 |
| Glrx5_Q80Y14 | -1.4 | 0.04 | -0.2 | -0.5 | -1.4 | 0.19 | 1.01 | 0.99 | 1.32 | Glrx5 |
| Mcu_Q3UMR5 | 0.94 | -1.2 | 0.12 | 0.86 | 0.64 | 1.36 | -0.7 | -0.8 | -1.2 | Mcu |
| Capg_P24452 | 1.2 | -0 | -0 | 0.27 | 1.52 | 0.28 | -0.5 | -1.1 | -1.6 | Capg |
| Erap1_Q9EQH2 | 0.61 | 0.19 | 0.41 | 0.45 | 0.71 | 0.78 | -1.7 | -1.7 | 0.32 | Erap1 |
| Cacybp_Q9CXW3 | -1.3 | -0.7 | 0.48 | -1.2 | -0.3 | -0.2 | 1.33 | 0.86 | 1.13 | Cacybp |
| Txnl1_Q8CDN6 | 0.6 | -0.1 | -0.3 | 0.57 | 0.49 | 1.74 | -1.6 | -0.3 | -1.1 | Txnl1 |
| Dhx9_O70133 | 0.12 | 0.05 | -0.8 | 1.26 | 1.53 | 0.72 | -1.2 | -0.6 | -1 | Dhx9 |
| Sod1_P08228 | -1.4 | -0.3 | 0.65 | -0.8 | -1 | -0.3 | 0.92 | 0.91 | 1.41 | Sod1 |
| Arhgap1_Q5FWK3 | 0.73 | -0.8 | -0.9 | 0.62 | 1.09 | 1.44 | -0 | -1.1 | -1.2 | Arhgap1 |
| Gna13_P27601 | 0.96 | 0.29 | 0.83 | -0.2 | 0.65 | 0.88 | -0.3 | -1.2 | -1.9 | Gna13 |
| Flnc_Q8VHX6 | 1.45 | -0.1 | 0.36 | 0.37 | 1.06 | 0.32 | -1.3 | -1.5 | -0.6 | Flnc |
| Vdac3_Q60931 | 1.47 | 0.82 | 0.48 | 0 | 0.58 | 0.28 | -1.4 | -0.9 | -1.3 | Vdac3 |
| Scp2_P32020 | 0.09 | -0.3 | -0.4 | -1.7 | 0.47 | -1.1 | 1.01 | 1.44 | 0.52 | Scp2 |
| Kif5b_Q61768 | 0.95 | 0.4 | 0.69 | 0.01 | 1.21 | 0.17 | -0.7 | -2 | -0.7 | Kif5b |
| Cbr2_P08074 | 0.13 | -0.1 | 0.09 | -1 | -1.8 | -0.2 | 1.09 | 1.55 | 0.21 | Cbr2 |
| Ap2b1_Q9DBG3 | 0.87 | -0 | -0.1 | 0.61 | 1.14 | 0.98 | -1.4 | -0.5 | -1.5 | Ap2b1 |
| Epb41l2_O70318 | -0.1 | 0.88 | 0.51 | 0.59 | 0.46 | 1.25 | -1.7 | -0.9 | -1 | Epb41l2 |
| Myh10_Q61879 | 0.7 | 0.6 | 0.2 | 0.82 | 0.68 | 0.82 | -1.3 | -1.8 | -0.8 | Myh10 |
| Pdlim5_Q8CI51 | 0.94 | 0.32 | 0.32 | 0.74 | -0.3 | 1.43 | -0.8 | -1.5 | -1.2 | Pdlim5 |

| | | | | | | | | | | |
|---|---|---|---|---|---|---|---|---|---|---|
| Cops8_Q8VBV7 | -0.7 | 0.69 | 0.89 | -0.8 | -0.3 | -1.9 | 0.22 | 1.08 | 0.84 | Cops8 |
| Fbln2_P37889 | -0.6 | 0.74 | 0.73 | 1.27 | -0.2 | 1.16 | -1.3 | -1.2 | -0.5 | Fbln2 |
| Dlat_Q8BMF4 | -1.1 | -0.9 | -0.1 | -0.7 | -0.6 | -0.2 | 0.5 | 1.82 | 1.26 | Dlat |
| P4hb_P09103 | -0 | 0.46 | -0 | -1.5 | -1 | -1 | 1.12 | 1.41 | 0.56 | P4hb |
| Capzb_P47757 | 1.14 | 0.61 | 0.5 | -0.3 | 1.29 | 0.18 | -0.8 | -1.6 | -1 | Capzb |
| Fkbp10_Q61576 | 0.62 | -0.1 | 0.08 | -0.1 | -2 | -0.7 | 1.21 | 0.44 | 0.75 | Fkbp10 |
| Slc29a1_Q9JIM1 | -0.3 | -0.3 | 0.52 | -0.1 | 1.28 | 1.67 | -0.8 | -0.4 | -1.5 | Slc29a1 |
| Ctnna1_P26231 | 1.05 | 0.74 | 1.14 | 0.22 | -0 | 0.43 | -0.7 | -1.1 | -1.8 | Ctnna1 |
| Ezr_P26040 | 0.91 | 0.26 | -0 | -0.1 | 1.38 | 1 | -1.3 | -1.5 | -0.6 | Ezr |
| Rbm3_O89086 | -0.8 | 0.24 | 0.17 | 1.44 | 0.01 | 1.57 | -0.9 | -1.3 | -0.4 | Rbm3 |
| Sptbn1_Q62261 | 1.09 | 0.79 | 0.15 | 0.6 | 1 | -0.1 | -1.1 | -1.8 | -0.7 | Sptbn1 |
| Uqcrc1_Q9CZ13 | 0.48 | -0.9 | 0.25 | -0.5 | -1.2 | -1.1 | 1.68 | 1.07 | 0.21 | Uqcrc1 |
| Snrpa_Q62189 | 1.05 | 0.24 | -0.8 | 0.29 | 1.59 | 0.51 | -0.4 | -0.9 | -1.5 | Snrpa |
| Cd248_Q91V98 | 1.47 | 0.06 | 0.7 | -0.3 | 0.23 | 1.16 | -1 | -1.4 | -0.9 | Cd248 |
| Copg1_Q9QZE5 | 0.8 | 0.5 | -0.2 | 1.34 | 0.3 | 0.44 | -2 | -0.4 | -0.7 | Copg1 |
| Bag6_Q9Z1R2 | 0.91 | 0.1 | 0.17 | 0.79 | 0.26 | 0.85 | -0.9 | 0.03 | -2 | Bag6 |
| Oat_P29758 | 0.71 | -0.1 | 0.73 | -0.2 | 1.19 | 0.88 | -0.2 | -1.5 | -1.6 | Oat |
| Chchd4_Q8VEA4 | -1.8 | 0.18 | 0.28 | -0.8 | -0.8 | -0.1 | 1.08 | 1.02 | 1.02 | Chchd4 |
| Mybbp1a_Q7TPV4 | 1.11 | -0.6 | 1.12 | 0.41 | 0.17 | 1.05 | -0.7 | -1.1 | -1.5 | Mybbp1a |
| Lima1_Q9ERG0 | 0.95 | 0.32 | -0.5 | 0.44 | 1.52 | 0.4 | -1.7 | -0.9 | -0.5 | Lima1 |
| Mmp14_P53690 | 0.97 | 0.67 | 0.36 | 0.32 | 1.32 | 0.08 | -1.3 | -1.3 | -1.2 | Mmp14 |
| Eea1_Q8BL66 | 0.42 | -0.7 | 0.75 | 0.85 | 0.56 | 1.27 | -0.5 | -1.8 | -0.8 | Eea1 |
| Timm23_Q9WTQ8 | 0.49 | 0.57 | 0.48 | -1.8 | -1.7 | 0.38 | 0.43 | 0.65 | 0.51 | Timm23 |
| Sfxn3_Q91V61 | 1.29 | 0.18 | -1.1 | 0.48 | 0.64 | 1.33 | -0.8 | -1 | -1.1 | Sfxn3 |
| Capza1_P47753 | 1.21 | -0.2 | -0.5 | 0.1 | 1.32 | 1.03 | -0.7 | -0.7 | -1.5 | Capza1 |
| Plod3_Q9R0E1 | 0.3 | 0.07 | 0.73 | -0.2 | 0.48 | 1.84 | -0.8 | -0.8 | -1.5 | Plod3 |
| Vcam1_P29533 | 1.82 | 0.13 | 0.42 | 0.43 | 0.2 | 0.43 | -1.3 | -1.3 | -0.8 | Vcam1 |
| Ptges2_Q8BWM0 | 0.66 | 0.67 | 0.19 | -1.8 | -1.7 | 0.37 | 0.5 | 0.54 | 0.57 | Ptges2 |
| Chchd2_Q9D1L0 | -0.7 | 1.08 | 0.97 | -0.7 | -1.6 | -1.1 | 0.26 | 0.75 | 0.9 | Chchd2 |
| Fis1_Q9CQ92 | -0.5 | -0.7 | 0.92 | -1 | -1.1 | -0.6 | 0.29 | 1.69 | 0.96 | Fis1 |
| Arhgdia_Q99PT1 | 1.39 | -0.1 | -0.3 | 0.39 | 1.55 | 0.12 | -1.2 | -1.2 | -0.7 | Arhgdia |
| Sept7_O55131 | -0 | 1.08 | 0.65 | 0.84 | -0.2 | 1.16 | -1 | -1.5 | -1 | Sept7 |
| Copg2_Q9QXK3 | -0.3 | -0.1 | 0.05 | 0.66 | 2 | -0.2 | -0.7 | -1.3 | -0.4 | Copg2 |
| Smap_Q9R0P4 | 0.43 | 1.2 | 0.5 | 0.76 | 1 | -0.6 | -1.4 | -0.5 | -1.4 | Smap |
| Prdx5_P99029 | 0.93 | -0.4 | -0.6 | -1.4 | 0.04 | -1.3 | 1.25 | 1.11 | 0.38 | Prdx5 |
| Arpc5_Q9CPW4 | 0.5 | 0.33 | -0.3 | 0.37 | 0.89 | 1.06 | 0.42 | -1.8 | -1.5 | Arpc5 |
| G3bp2_P97379 | -1.4 | 0.9 | 0.28 | -1.2 | -0.7 | -0.6 | 0.62 | 1.46 | 0.63 | G3bp2 |
| Cad_B2RQC6 | 0.84 | 0.48 | 0.4 | 0.37 | 0.46 | 0.58 | -1.7 | -1.7 | 0.34 | Cad |
| Gsn_P13020 | -0.2 | 0.74 | 0.83 | 0.95 | 0.08 | 1.08 | -1.7 | -0.6 | -1.2 | Gsn |
| Ran_P62827 | 1.33 | -0.2 | 0.22 | 0.32 | 1.09 | 0.53 | -0.4 | -1.7 | -1.2 | Ran |
| Capn6_O35646 | 0.43 | -1 | 0.07 | 0.87 | 1.94 | 0.15 | -1.1 | -0.9 | -0.4 | Capn6 |
| Acaa2_Q8BWT1 | 1.16 | -0.8 | -0.5 | -1.3 | -0.5 | -0.9 | 1.42 | 0.84 | 0.57 | Acaa2 |
| Prep_Q9QUR6 | -0.3 | 0.24 | 0.73 | -1.2 | -1.5 | -0.3 | 0.21 | 0.33 | 1.81 | Prep |
| Atp5j_P97450 | -0.3 | -0.8 | -1 | -1.1 | 0.47 | -0.6 | 1.8 | 0.98 | 0.56 | Atp5j |
| Atp1a1_Q8VDN2 | 0.19 | 0.56 | -0 | 1.19 | 1.34 | 0.13 | -0.5 | -1.3 | -1.6 | Atp1a1 |
| Ehd4_Q9EQP2 | -1.1 | -0.2 | -1.1 | 0.92 | 0.38 | 2 | 0.07 | -0.6 | -0.4 | Ehd4 |
| Tceb2_P62869 | 0.19 | -0.2 | -0.5 | 0.11 | -0.9 | -1.5 | -0 | 1.92 | 0.93 | Tceb2 |
| Myl6_Q60605 | 1.5 | 0.34 | 0.26 | -0.3 | 1.29 | 0.16 | -1.1 | -1.4 | -0.8 | Myl6 |
| Tubb5_P99024 | 1.59 | 0.29 | -0.2 | 0.46 | 1.16 | 0.05 | -0.9 | -1.4 | -1.1 | Tubb5 |
| Pak2_Q8CIN4 | 1.24 | -0.7 | -0.2 | 0.09 | 1.15 | 1.15 | -0.5 | -0.5 | -1.6 | Pak2 |

| | | | | | | | | | | |
|---|---|---|---|---|---|---|---|---|---|---|
| Pip4k2c_Q91XU3 | -0.9 | 0.62 | -0.6 | 0.74 | 2 | 0.29 | -0.8 | -0.5 | -0.9 | Pip4k2c |
| Me1_P06801 | -0.4 | 0.94 | -0.4 | -1.5 | 0.42 | -1.5 | 0.89 | 1.08 | 0.48 | Me1 |
| Vwa5a_Q99KC8 | 1.43 | 0.71 | 0.16 | -0.3 | 1 | 0.48 | -1.3 | -0.8 | -1.3 | Vwa5a |
| Prkar1a_Q9DBC7 | 0.93 | 0.68 | 0.43 | 0.33 | 0.01 | 0.79 | 0.18 | -1.5 | -1.9 | Prkar1a |
| Hexb_P20060 | 1.49 | -0 | 0.09 | -0.1 | 1.44 | 0.29 | -0.7 | -1.3 | -1.2 | Hexb |
| Hmgb2_P30681 | -0.1 | -0.5 | -0.6 | 0.41 | 1.7 | 0.94 | 0.58 | -0.9 | -1.5 | Hmgb2 |
| Nnmt_O55239 | -0.7 | 0.23 | 0.45 | 0.24 | -1.5 | -1.5 | 0.42 | 1.24 | 1.07 | Nnmt |
| Atp5o_Q9DB20 | 0.2 | 0.02 | -0.1 | -2 | 0.24 | -0.9 | 0.82 | 1.54 | 0.18 | Atp5o |
| Cbr3_Q8K354 | 0.72 | -0.6 | -0.4 | -1.6 | 0.24 | -1.1 | 1.53 | 0.8 | 0.39 | Cbr3 |
| Hnrnpf_Q9Z2X1 | 1.25 | -0.6 | -0.2 | 0.34 | 1.62 | 0.57 | -0.8 | -1 | -1.2 | Hnrnpf |
| Rpl24_Q8BP67 | -0.2 | -0.3 | -0.3 | -0.3 | -1.1 | -1 | 0.29 | 2 | 0.65 | Rpl24 |
| Dnaja2_Q9QYJ0 | -0.6 | -1.1 | -0.7 | 1.41 | 0.46 | 1.57 | -1.2 | -0.1 | 0.17 | Dnaja2 |
| Aacs_Q9D2R0 | 0.31 | 0.51 | 0.37 | 0.36 | -1.9 | -1.4 | 0.61 | 0.05 | 1.13 | Aacs |
| Lsm4_Q9QXA5 | -0 | -1.2 | 0.59 | 0.79 | 0.48 | 1.7 | -0.3 | -0.7 | -1.4 | Lsm4 |
| Vasp_P70460 | 1.1 | 1.27 | 1.34 | -0.1 | -0.1 | -0.5 | -1 | -1.1 | -1 | Vasp |
| Dpep1_P31428 | 0.25 | 0.5 | 0.95 | 0.68 | -0.4 | 1.28 | -1.7 | -0.3 | -1.2 | Dpep1 |
| Rcn3_Q8BH97 | -0.2 | 0.43 | 1.22 | 1.23 | -0.5 | 0.97 | -0.9 | -1.5 | -0.7 | Rcn3 |
| Ywhab_Q9CQV8 | 0.92 | 0.02 | -1 | -1.1 | -0.3 | -1.2 | 0.7 | 1.59 | 0.44 | Ywhab |
| Sptan1_P16546 | 0.61 | 1.22 | 0.65 | 0.98 | 0.29 | -0.1 | -0.9 | -1.6 | -1.1 | Sptan1 |
| Shmt2_Q9CZN7 | 0.6 | 0.14 | 1.15 | -1.5 | -1 | -1.3 | 0.14 | 1.04 | 0.65 | Shmt2 |
| Vcp_Q01853 | -0.7 | 0.55 | 0.72 | -0.5 | -2 | -0.5 | 0.58 | 1.01 | 0.89 | Vcp |
| Apmap_Q9D7N9 | 0.69 | -0.3 | -0.1 | -2 | -0 | -0.6 | 1.51 | 0.23 | 0.64 | Apmap |
| Hist1h1c_P15864 | 1.57 | -0.4 | 0.33 | 0.44 | 0.78 | 0.7 | -1.2 | -1.3 | -0.8 | Hist1h1c |
| Rpl18a_P62717 | 1.79 | 0.07 | -0.1 | 0.08 | 0.86 | 0.54 | -0.7 | -1.3 | -1.3 | Rpl18a |
| Acss2_Q9QXG4 | 0.46 | 0.5 | 0.42 | 0.54 | -1.8 | -1.7 | 0.43 | 0.5 | 0.66 | Acss2 |
| Lrrc59_Q922Q8 | -1.5 | 0.25 | 0.26 | -0.1 | -1.5 | -0.4 | 0.33 | 1.37 | 1.19 | Lrrc59 |
| Acadsb_Q9DBL1 | 0.86 | 0.95 | -0.3 | -1.7 | -0.8 | -0.6 | 1.02 | 1.06 | -0.5 | Acadsb |
| Mapk1_P63085 | 1.66 | -0.2 | -0.6 | 0.14 | 0.78 | 1.14 | -0.7 | -1.1 | -1.1 | Mapk1 |
| Eif3k_Q9DBZ5 | 0.21 | -1.6 | -0.5 | 1.08 | 1.78 | 0.27 | -0 | -0.5 | -0.8 | Eif3k |
| Acly_Q91V92 | -0.4 | 0.38 | 0.32 | -1.8 | -0.9 | -0 | 1.2 | 1.42 | -0.2 | Acly |
| Dbn1_Q9QXS6 | -0.7 | 0.74 | 0.69 | 0.72 | -0.2 | 1.56 | -1.5 | -0.3 | -1 | Dbn1 |
| Cyb5r3_Q9DCN2 | 1.3 | 0.31 | 0.06 | -0.1 | 1.44 | 0.37 | -0.8 | -1.2 | -1.4 | Cyb5r3 |
| Hspa1a_Q61696 | 0.85 | 0.45 | 1.25 | 0.82 | 0.57 | -0.6 | -1.6 | -1 | -0.7 | Hspa1a |
| Snrpb_P27048 | 1.26 | 0.76 | 0.47 | -0.3 | 1.27 | -0 | -1.1 | -1 | -1.3 | Snrpb |
| Ppa2_Q91VM9 | -0 | 0.04 | -0.6 | 0.45 | -1.2 | -1.5 | 1.11 | 0.18 | 1.54 | Ppa2 |
| Rps16_P14131 | 1.16 | 0.34 | -0.1 | 1.05 | 0.62 | 0.44 | -1.9 | -0.8 | -0.9 | Rps16 |
| Arf6_P62331 | 0.17 | 1.08 | 1.11 | 1.12 | -0.1 | -0 | -1.7 | -0.8 | -0.9 | Arf6 |
| Ctsb_P10605 | 0.63 | 0.5 | -0.5 | 0.56 | -0.2 | 1.91 | -0.6 | -1.4 | -0.9 | Ctsb |
| Srsf2_Q62093 | -0.9 | -0.1 | 0.59 | 1.64 | 0.58 | 0.73 | -1.7 | -0.2 | -0.7 | Srsf2 |
| Rab22a_P35285 | -0.3 | -0.9 | 0.04 | 1.68 | 1.07 | 0.22 | -0.7 | -1.6 | 0.4 | Rab22a |
| Col8a1_Q00780 | 0.68 | 0.79 | -0.2 | 0.03 | 1.61 | 0.18 | -1.9 | -0.7 | -0.5 | Col8a1 |
| Actn4_P57780 | 1.03 | 0.5 | 0.46 | 0.34 | 0.84 | 0.64 | -1 | -1.8 | -1.1 | Actn4 |
| Cox6c_Q9CPQ1 | 0.4 | -1 | -1.1 | -1.2 | 0.19 | -0.5 | 1.46 | 0.97 | 0.83 | Cox6c |
| Clybl_Q8R4N0 | 0.26 | -0.7 | 0.05 | 0.23 | -2 | -0.5 | 1.45 | 0.36 | 0.84 | Clybl |
| Adi1_Q99JT9 | 1.33 | -0.5 | 0.23 | 0.51 | 1.03 | 0.38 | -0.4 | -0.7 | -2 | Adi1 |
| Farsa_Q8C0C7 | 1.23 | 0.4 | 0.24 | -0.8 | 0.82 | 1.27 | -1.2 | -1 | -0.9 | Farsa |
| Hist1h1b_P43276 | 1.37 | -0.5 | 1.02 | 0.03 | 0.97 | 0.35 | -0.8 | -1.5 | -1 | Hist1h1b |
| Grn_P28798 | -1.9 | 0.27 | 0.01 | -0.2 | -1.1 | -0.1 | 0.93 | 0.96 | 1.14 | Grn |
| Esyt1_Q3U7R1 | 0.05 | -1.2 | -0.8 | -0.3 | -0.1 | -0.9 | 2 | 0.5 | 0.72 | Esyt1 |
| Vcan_Q62059 | -0.2 | -0.3 | -0.3 | -0.2 | 2 | 0.91 | -1.4 | -0.8 | 0.22 | Vcan |

| | | | | | | | | | | |
|---|---|---|---|---|---|---|---|---|---|---|
| Mtco2_P00405 | -1 | 0.56 | 0.64 | 0.14 | -1.6 | -1.1 | 0.22 | 0.89 | 1.25 | Mtco2 |
| Tnpo1_Q8BFY9 | -0.6 | 0.03 | -0.1 | 0.33 | 1.89 | 0.56 | 0.45 | -1 | -1.6 | Tnpo1 |
| Gstm2_P15626 | 0.16 | 0.81 | 0.18 | 0.55 | 1.86 | -0.6 | -1.4 | -0.8 | -0.8 | Gstm2 |
| Pls3_Q99K51 | 1.14 | 0.06 | 0.16 | 1.24 | 1.13 | -0.4 | -1 | -1.3 | -1 | Pls3 |
| Ap3d1_O54774 | 0.89 | 1.34 | 0.51 | -0.3 | -1.5 | -1.5 | -0.3 | 0.34 | 0.56 | Ap3d1 |
| Mars_Q68FL6 | 0.28 | -0.3 | -1.3 | 0.71 | 1.95 | 0.38 | -0.1 | -1.2 | -0.4 | Mars |
| Ndufa4_Q62425 | 0.96 | -0.1 | -0.3 | -1.2 | -0 | -1.8 | 0.99 | 0.75 | 0.77 | Ndufa4 |
| Akr7a2_Q8CG76 | 0.06 | 0.41 | -0 | -0.3 | -0.1 | -2 | 0.86 | 1.28 | 0.09 | Akr7a2 |
| Ddx6_P54823 | 0.09 | -0 | 0.18 | -0.5 | -2 | -0 | 0.49 | 1.78 | 0.07 | Ddx6 |
| Plin3_Q9DBG5 | 0.65 | 0.77 | 0.1 | 0.44 | 1.24 | 0.36 | -1.2 | -1.9 | -0.5 | Plin3 |
| Hspa8_P63017 | 0.48 | -0.1 | -0.5 | -1.2 | -0.1 | -1.4 | 1.49 | 1.38 | -0 | Hspa8 |
| Trnt1_Q8K1J6 | -1.1 | 0.58 | -0.5 | -0.5 | -0.6 | -1 | 0.05 | 1.56 | 1.48 | Trnt1 |
| Snx6_Q6P8X1 | -1 | -0.1 | 1.26 | 0.49 | 0.86 | 1.1 | -1.7 | -0.5 | -0.5 | Snx6 |
| Canx_P35564 | -1.2 | 0.61 | 0.83 | -0.3 | -1.7 | -0.5 | 0.61 | 1.14 | 0.62 | Canx |
| Dhodh_O35435 | 0.6 | 0.49 | 0.4 | -1.8 | 0.58 | -1.7 | 0.44 | 0.51 | 0.5 | Dhodh |
| Lrrc47_Q505F5 | -1.1 | 0.9 | 0.83 | -0.7 | -1.8 | -0.3 | 0.74 | 0.28 | 1.05 | Lrrc47 |
| Rrbp1_Q99PL5 | -0.2 | 0.81 | 0.85 | 1.17 | -0.3 | 1.05 | -1.4 | -1.1 | -0.8 | Rrbp1 |
| Eci1_P42125 | 1.52 | -0.1 | -0.2 | -1.1 | -0.8 | -1.2 | 0.96 | 1.14 | -0.2 | Eci1 |
| Dhrs4_Q99LB2 | 0.9 | -0.2 | 0.66 | -0.6 | -0.5 | -1.9 | 1.59 | -0.2 | 0.19 | Dhrs4 |
| Sqrdl_Q9R112 | -0 | -0.1 | 0.39 | -0.5 | 0.01 | -2 | 1.24 | -0 | 1.18 | Sqrdl |
| Cndp2_Q9D1A2 | 1.5 | -0.1 | -0.1 | -0.1 | 1.49 | 0.45 | -0.7 | -1.1 | -1.2 | Cndp2 |
| Aars_Q8BGQ7 | 1.33 | -0.4 | -0.4 | 0.13 | 1.47 | 0.63 | -0.3 | -1.6 | -0.8 | Aars |
| Pfn2_Q9JJV2 | 0.92 | 1.01 | -0.1 | -0.4 | 0.9 | 0.74 | -1.7 | -1.3 | -0.1 | Pfn2 |
| Sdha_Q8K2B3 | 0.58 | 0.68 | 0.26 | -0.1 | -1 | -2 | 0.62 | 0.9 | 0.3 | Sdha |
| Scpep1_Q920A5 | 1.11 | 0.06 | 1.23 | 0.3 | 0.17 | 0.64 | -1.4 | -0.5 | -1.5 | Scpep1 |
| Sf3a3_Q9D554 | 1.18 | 1.23 | -0.4 | -0.4 | 0.45 | 1.11 | -1 | -1 | -1.2 | Sf3a3 |
| Rbms1_Q91W59 | 1.58 | 0.59 | -0.4 | 0.45 | 0.91 | -0.1 | -0.3 | -1.5 | -1.3 | Rbms1 |
| Suclg2_Q9Z2I8 | 0.57 | 0.27 | -0 | -1.4 | 0.48 | -2 | 0.5 | 0.77 | 0.8 | Suclg2 |
| Tln1_P26039 | 0.61 | 0.53 | 0.74 | 1.17 | -0.3 | 0.84 | -1.4 | -1.4 | -0.7 | Tln1 |
| Eef1g_Q9D8N0 | 0.55 | 0.39 | 0.29 | -0.5 | -1.3 | -1.9 | 1.08 | 0.54 | 0.76 | Eef1g |
| Ndufc2_Q9CQ54 | -0.1 | 0.57 | 0.21 | -1.2 | 0.54 | -2 | 0.19 | 1.16 | 0.68 | Ndufc2 |
| Apeh_Q8R146 | 0.49 | 0.59 | -0.6 | -0.2 | 2 | 0.34 | -0.8 | -1.3 | -0.6 | Apeh |
| Aak1_Q3UHJ0 | 1.24 | -0 | -0.2 | -0 | 1.12 | 0.91 | -0.1 | -1.7 | -1.2 | Aak1 |
| Galk1_Q9R0N0 | 0.22 | -0.9 | -0.1 | -0.3 | 1.18 | 1.85 | 0.11 | -1 | -1 | Galk1 |
| Srsf3_P84104 | 0.06 | -0.2 | -0.1 | -0.1 | 0.48 | 2 | -0.4 | -1.6 | -0.3 | Srsf3 |
| Manf_Q9CXI5 | 0.96 | 0.23 | -1.5 | 0.36 | 1.59 | 0.59 | -1.1 | -0.6 | -0.6 | Manf |
| Rps19_Q9CZX8 | 0.11 | 0.21 | -0.2 | 0.18 | -1.9 | -0.9 | -0.1 | 1.25 | 1.36 | Rps19 |
| Sh3gl1_Q62419 | -0.7 | -0.4 | 1.16 | 0.21 | 1.4 | 1 | -1.1 | -0.3 | -1.3 | Sh3gl1 |
| Ivd_Q9JHI5 | -0.2 | 0.49 | -0.5 | -1.3 | -0.9 | -0.5 | 1.2 | 1.83 | -0.2 | Ivd |
| Ilf3_Q9Z1X4 | 1.18 | 0.48 | -0.7 | -0 | 1.56 | 0.56 | -1.2 | -1 | -0.9 | Ilf3 |
| Postn_Q62009 | 1.69 | -0.1 | -0.2 | 0 | 1.16 | 0.55 | -0.8 | -1.5 | -0.8 | Postn |
| Gpi_P06745 | -1.2 | -0.1 | -0.2 | -0.3 | -1.5 | 0.09 | 0.57 | 1.29 | 1.42 | Gpi |
| Ifi30_Q9ESY9 | 0.02 | -0.8 | -0.7 | 0.5 | -0.1 | -1.8 | 0.9 | 1.5 | 0.55 | Ifi30 |
| Prpsap1_Q9D0M1 | -0.9 | -0.7 | 0.23 | -0.1 | 1.24 | 1.85 | -0.2 | -1.2 | -0.1 | Prpsap1 |
| Psmd12_Q9D8W5 | 0.33 | -0.3 | -1 | -1 | -1.3 | 0.37 | 1.33 | 1.56 | -0.1 | Psmd12 |
| Rps29_P62274 | 0.36 | 0.52 | 0.72 | -1.8 | 0.58 | -1.7 | 0.5 | 0.54 | 0.28 | Rps29 |
| Ube2n_P61089 | 0.73 | 1.1 | 0.84 | -0.3 | 0.91 | 0.33 | -0.9 | -1 | -1.6 | Ube2n |
| Leprel4_Q8K2B0 | 0.47 | 1.15 | -1 | -0.2 | 1.07 | 1.19 | -0.9 | -1.3 | -0.3 | Leprel4 |
| Arhgdib_Q61599 | 1.49 | -0.2 | 0.21 | -0.1 | 1.39 | 0.26 | -0.6 | -1.6 | -0.9 | Arhgdib |
| Ptn_P63089 | 0.51 | 1.16 | 1.02 | 1.03 | -0.6 | 0.02 | -1.4 | -1.4 | -0.4 | Ptn |

| | | | | | | | | | | |
|---|---|---|---|---|---|---|---|---|---|---|
| Maob_Q8BW75 | -0.6 | 0.35 | -0.1 | -2 | -0.7 | 0.26 | 1.45 | 0.24 | 0.97 | Maob |
| Aldh6a1_Q9EQ20 | 0.88 | -0.3 | -0.8 | -1.6 | 0.01 | -0.7 | 1.58 | 0.94 | 0.02 | Aldh6a1 |
| Timm13_P62075 | -1.2 | 1.19 | -0.5 | -0 | -1.5 | -0.5 | 0.8 | 0.79 | 0.98 | Timm13 |
| Denr_Q9CQJ6 | -0.9 | -0.3 | -0.1 | -0.3 | -1.8 | 0.4 | 0.88 | 1.58 | 0.5 | Denr |
| Tmem43_Q9DBS1 | -0.4 | -0.3 | -1 | -1.1 | 0.45 | -1 | 1.14 | 1.59 | 0.66 | Tmem43 |
| Stx7_O70439 | 1.01 | 1.35 | -1 | 0.69 | 0.38 | 0.46 | -0.8 | -1.6 | -0.6 | Stx7 |
| Ndufs8_Q8K3J1 | 0.37 | 0 | 0.43 | 0.19 | -2 | -0.4 | 0.69 | 0.57 | 0.62 | Ndufs8 |
| Cox7a2_P48771 | 1.89 | 0.2 | 0.1 | 0.61 | 0.19 | 0.05 | -0.6 | -0.7 | -1.8 | Cox7a2 |
| _C0HKD9 | 0.57 | 0.56 | 0.68 | 0.56 | -1.8 | -1.7 | 0.21 | 0.32 | 0.59 | |
| Bsg_P18572 | -0.2 | 0.84 | 1.05 | -1.1 | -0.6 | -1.5 | -0.1 | 0.05 | 1.57 | Bsg |
| Uap1l1_Q3TW96 | 0.32 | -0.8 | 0.09 | 0.39 | 1.23 | 1.34 | -0.7 | -0.1 | -1.8 | Uap1l1 |
| Arsb_P50429 | -0.5 | 0.57 | 0.93 | 1.75 | -0 | 0.44 | -0.9 | -1.1 | -1.1 | Arsb |
| Lamp2_P17047 | 1.13 | 0.41 | 0.59 | 0.18 | 1.5 | -0.6 | -1.1 | -1.3 | -0.9 | Lamp2 |
| Nsfl1c_Q9CZ44 | 1.66 | -0.1 | -0.7 | -0.1 | 1.19 | 0.76 | -0.8 | -1.2 | -0.7 | Nsfl1c |
| Sept11_Q8C1B7 | 1.03 | -0.5 | 1.08 | 0.58 | 1.19 | -0.3 | -0.4 | -1.4 | -1.2 | Sept11 |
| Myl12b_Q3THE2 | 1.53 | 0.22 | 0.71 | 0.92 | 0.56 | -0.8 | -1.1 | -1.1 | -0.9 | Myl12b |
| Actb_P60710 | 1.74 | 0.09 | -0.1 | 0.07 | 1.23 | 0.06 | -0.9 | -1.4 | -0.8 | Actb |
| Mdh1_P14152 | -0.4 | 0.52 | -1.1 | -0.9 | -1.1 | 0.09 | 1.48 | 1.47 | -0 | Mdh1 |
| Palm_Q9Z0P4 | 0.07 | -1.7 | 0.59 | -1.4 | 0.02 | -0.3 | 1.29 | 0.57 | 0.87 | Palm |
| Lmnb1_P14733 | 0.77 | 0.91 | 0.02 | 0.26 | -0.3 | 1.53 | -0.4 | -1.6 | -1.2 | Lmnb1 |
| Ppp3ca_P63328 | 1.05 | 0.45 | -1.1 | 0.02 | 1.12 | 0.95 | -1.7 | -0.6 | -0.2 | Ppp3ca |
| Pafah1b3_Q61205 | -0.3 | 1.05 | 1.08 | 0.34 | 0.12 | 0.9 | -1.3 | -0.1 | -1.7 | Pafah1b3 |
| Tomm70a_Q9CZW5 | -0 | 0.85 | -0.9 | -1.8 | -0.4 | -0.3 | 1.59 | 0.63 | 0.31 | Tomm70a |
| Cox4i1_P19783 | 0.93 | -0.4 | -1.2 | -1.2 | 0.4 | -1.2 | 0.95 | 1.07 | 0.67 | Cox4i1 |
| Pebp1_P70296 | -0.1 | 1.48 | 0 | 0.02 | 1.48 | 0.26 | -0.8 | -1.1 | -1.3 | Pebp1 |
| Ftl1_P29391 | -0 | -0.5 | 0.61 | 0.29 | 1.94 | 0.06 | 0.08 | -1.7 | -0.7 | Ftl1 |
| Reep5_Q60870 | -1.4 | 0.31 | 0.38 | -0.4 | -1.7 | 0.27 | 0.46 | 1.18 | 0.96 | Reep5 |
| Acbd3_Q8BMP6 | -0.9 | -0.2 | -0.6 | 1.83 | 1.21 | -0 | -0.6 | 0.49 | -1.2 | Acbd3 |
| Snrpe_P62305 | 0.25 | -0.8 | -1.1 | 0.31 | 2 | 0.51 | -0.3 | -0.9 | -0.1 | Snrpe |
| Dync1li2_Q6PDL0 | 0.76 | -0.5 | -1.3 | 0.74 | 1.92 | 0.12 | -0.3 | -0.5 | -0.9 | Dync1li2 |
| Got2_P05202 | 0.71 | 0.99 | -0.1 | -1.5 | 0.19 | -1.8 | 0.38 | 0.9 | 0.19 | Got2 |
| Psmd13_Q9WVJ2 | -0.1 | -0.5 | 1.32 | 0.09 | -1.7 | -1 | 1.43 | 0.17 | 0.29 | Psmd13 |
| Nln_Q91YP2 | 0.51 | 0.57 | -1.5 | 0.54 | 1.3 | 0.67 | 0.23 | -1.2 | -1.1 | Nln |
| Cpq_Q9WVJ3 | 0.67 | 0.44 | -1.4 | 0.57 | 1.01 | 0.95 | -0.1 | -1.7 | -0.4 | Cpq |
| D10Jhu81e_Q9D172 | -1.1 | -0.5 | 0.57 | 0.14 | -1.9 | -0.1 | 0.79 | 1.15 | 0.89 | D10Jhu81e |
| Vat1_Q62465 | 1.71 | -0.4 | -0.2 | -0.2 | 1.43 | 0.41 | -0.8 | -1.1 | -0.9 | Vat1 |
| Cd44_P15379 | 0.09 | -0.8 | -0.6 | 0.83 | 0.44 | 1.55 | -0.2 | 0.58 | -1.8 | Cd44 |
| Abhd5_Q9DBL9 | 0.74 | 0.2 | 0.29 | -2 | -0.1 | -0 | 0.65 | 0.56 | 0.28 | Abhd5 |
| Asns_Q61024 | 0.29 | 0.92 | -0.1 | 0.02 | -1.3 | -1.7 | 1.36 | -0.3 | 0.79 | Asns |
| Elavl1_P70372 | 1.06 | -1.2 | -0.9 | 1.63 | 1.03 | 0.1 | -0.7 | -0.5 | -0.6 | Elavl1 |
| Ddost_O54734 | 0.61 | -0.6 | -0.6 | 0.66 | 1.5 | 0.45 | 0.18 | -0.2 | -2 | Ddost |
| Plscr3_Q9JIZ9 | 1.01 | -1 | -0.2 | -0.4 | 1.55 | 1.18 | -0.3 | -0.6 | -1.2 | Plscr3 |
| Dnm1l_Q8K1M6 | -0.4 | 0.12 | 0.34 | -0.3 | -2 | 0.13 | 0.41 | 0.65 | 1.31 | Dnm1l |
| Acads_Q07417 | 0.05 | -0.2 | -0.5 | -0.5 | 0.67 | -2 | 0.79 | 1.24 | 0.56 | Acads |
| Arf5_P84084 | -0.1 | 0.48 | 1.15 | 0.28 | -0.2 | 1.46 | -1.5 | -1.3 | -0.3 | Arf5 |
| Sdhb_Q9CQA3 | -0.2 | 0.51 | 0.95 | -0.6 | -2 | -0.4 | 1.27 | 0.83 | -0.4 | Sdhb |
| Flot2_Q60634 | -1.3 | 0.62 | 0.79 | -0.3 | -1.6 | -0.5 | 0.56 | 0.36 | 1.38 | Flot2 |
| Islr_Q6GU68 | -0.6 | 0.88 | 0.97 | -1.6 | -0.3 | -0.8 | 0.51 | 1.46 | -0.5 | Islr |
| Slc1a5_P51912 | -0.4 | -0.1 | 0.56 | -0.2 | -2 | -0.1 | -0 | 1.71 | 0.56 | Slc1a5 |
| Serpinb6_Q60854 | 1.54 | 0 | 1.09 | -0.2 | 0.72 | 0.16 | -1 | -1.5 | -0.8 | Serpinb6 |

| | | | | | | | | | | |
|---|---|---|---|---|---|---|---|---|---|---|
| Gla_P51569 | -0.4 | 0.21 | 0.72 | 0.12 | 0.81 | 1.18 | -2 | 0.12 | -0.6 | Gla |
| Pgm1_Q9D0F9 | 1.66 | -0.3 | -0.5 | -1.2 | -0.7 | -1 | 1.34 | 0.59 | 0.03 | Pgm1 |
| Hadha_Q8BMS1 | 0.98 | -0.5 | -0.9 | -1.6 | 0.41 | -1 | 1.23 | 0.79 | 0.55 | Hadha |
| Ganab_Q8BHN3 | -1.3 | 0.36 | 0.56 | -0.7 | -1.4 | -0.5 | 0.71 | 1.12 | 1.17 | Ganab |
| Hmgb1_P63158 | -0.3 | 1.21 | 0.39 | 0.98 | -0.5 | 1.19 | -1.6 | -0.9 | -0.4 | Hmgb1 |
| Ubtf_P25976 | 0.89 | -0.3 | -0.6 | 0.31 | 1.46 | 0.44 | 0.58 | -1.5 | -1.3 | Ubtf |
| Chchd3_Q9CRB9 | -1.7 | 0.36 | 0.71 | -0.5 | -1.3 | -0.1 | 0.48 | 1.09 | 0.97 | Chchd3 |
| Rps13_P62301 | -1.5 | 0.13 | -0 | 0.44 | -1.1 | -1 | 1.15 | 1.22 | 0.69 | Rps13 |
| Pdlim2_Q8R1G6 | 1.61 | -1 | -0.9 | 1 | 0.75 | 0.65 | -0.7 | -1 | -0.4 | Pdlim2 |
| Psmd9_Q9CR00 | 0.23 | 0.7 | 0.22 | 0.48 | 1.53 | -0.4 | 0.27 | -1.4 | -1.6 | Psmd9 |
| Spg20_Q8R1X6 | 1.73 | 0.97 | 0.32 | 0.03 | 0.06 | -0.1 | -1.5 | -1.3 | -0.2 | Spg20 |
| Cdkn2c_Q60772 | -0.1 | -1.1 | 0.01 | 0.1 | 0.96 | 1.65 | 0.13 | 0.09 | -1.8 | Cdkn2c |
| Pgd_Q9DCD0 | 0.61 | -0.4 | 0.37 | -0.1 | -0 | -2 | 0.93 | 0.69 | 0.39 | Pgd |
| Hnrnpd_Q60668 | 1.09 | 0.3 | 0.92 | 0.66 | 0.44 | 0.27 | -0.7 | -1.9 | -1.1 | Hnrnpd |
| Gstt1_Q64471 | 0.91 | 0.64 | -0.3 | 0.37 | 1.84 | -0.7 | -1.2 | -0.9 | -0.7 | Gstt1 |
| Lrp1_Q91ZX7 | 0.11 | 1.46 | 0.29 | 0.49 | 0.6 | 0.73 | -1.3 | -1.4 | -1 | Lrp1 |
| Lamp1_P11438 | -1 | 0.06 | -0.2 | -0.1 | -0.4 | -1.6 | 0.5 | 1.36 | 1.43 | Lamp1 |
| Pdcd6ip_Q9WU78 | 1.94 | -0.2 | -0.6 | 0.25 | 1.12 | 0.18 | -0.9 | -0.8 | -1 | Pdcd6ip |
| Arl1_P61211 | -0.7 | 0.31 | 1.39 | 0.23 | -1.5 | -1.4 | 1.03 | 0.14 | 0.47 | Arl1 |
| Gpx1_P11352 | 1.63 | 0.04 | -0.2 | -0.5 | 1.47 | 0.38 | -0.7 | -1.2 | -1 | Gpx1 |
| Ppp1cb_P62141 | -0.8 | 0.37 | 0.84 | 1.42 | -0.7 | 1.23 | -0.6 | -0.3 | -1.4 | Ppp1cb |
| Cnn3_Q9DAW9 | -1 | 0.87 | 1.3 | 0.88 | 0.3 | 0.45 | -1.6 | -1 | -0.2 | Cnn3 |
| Fermt2_Q8CIB5 | 1.05 | -0.1 | -0.7 | 0.75 | 0.87 | 0.76 | -1 | 0.28 | -1.8 | Fermt2 |
| Dynll1_P63168 | 0.48 | 1.05 | 1.05 | 0.61 | -0.3 | 0.51 | -1.9 | -1.1 | -0.5 | Dynll1 |
| Ncam1_P13595 | 0.11 | 1.21 | 1.43 | 0.35 | 0.19 | -0.1 | -1.7 | -1.2 | -0.4 | Ncam1 |
| Eci2_Q9WUR2 | 0.33 | 0.28 | -0.4 | -1.6 | 0.31 | -1 | 1.77 | 0.76 | -0.4 | Eci2 |
| Carkd_Q9CZ42 | 0.77 | 0.82 | -0.2 | 1.04 | -0.6 | 1.02 | -0.6 | -0.4 | -1.9 | Carkd |
| Trap1_Q9CQN1 | -1.4 | 0.83 | 0.57 | 0.28 | -1.1 | -1.5 | 0.52 | 0.98 | 0.72 | Trap1 |
| Igf2bp3_Q9CPN8 | 0.36 | 0.09 | -0.1 | 1.11 | 1.14 | 0.78 | -1.9 | -1 | -0.5 | Igf2bp3 |
| Ybx1_P62960 | -1.7 | 0.7 | 0.57 | 0.18 | -1.4 | -0.7 | 0.85 | 0.65 | 0.84 | Ybx1 |
| Sephs1_Q8BH69 | 0.01 | 0.1 | 0.43 | 0.12 | 0.89 | 0.81 | -2 | 0.4 | -0.3 | Sephs1 |
| Tbc1d9b_Q5SVR0 | 0.06 | -1 | 0.73 | 0.8 | -0.2 | 1.7 | -0.6 | 0.1 | -1.6 | Tbc1d9b |
| Timm17b_Q9Z0V7 | 0.47 | -1 | 1.6 | 0.69 | 0.73 | 0.14 | -0.1 | -1.2 | -1.3 | Timm17b |
| Uba3_Q8C878 | 0.28 | 0.45 | 0.21 | 0.29 | 0.62 | 0.67 | -2 | 0.27 | -0.2 | Uba3 |
| Dynlt1_P51807 | 0.2 | 0.98 | 0.89 | -1.6 | -1.3 | -0.2 | 0.96 | -0.7 | 0.76 | Dynlt1 |
| Nmt1_O70310 | 1.06 | 1.45 | -0.8 | -1.3 | -0.3 | -1.3 | 0.22 | 0.55 | 0.41 | Nmt1 |
| Lsm3_P62311 | 1.83 | -0.4 | 0.68 | -1.1 | -1.1 | -0.7 | -0.2 | 1.05 | -0 | Lsm3 |
| Man2b2_O54782 | 1.66 | -0.9 | 0.47 | 0.07 | 1 | 0.47 | -1.4 | -0.6 | -0.6 | Man2b2 |
| Tubb6_Q922F4 | 1.29 | 1.12 | -0.2 | 0.78 | 0.28 | -0.2 | -0.5 | -1.9 | -0.7 | Tubb6 |
| Sec62_Q8BU14 | 0.37 | 0.7 | -0.7 | -0.6 | -1.9 | -0.1 | 1.52 | -0.1 | 0.76 | Sec62 |
| Gpd2_Q64521 | -1.3 | 0.31 | 0.65 | -0.3 | -1.9 | 0.15 | 0.29 | 0.76 | 1.24 | Gpd2 |
| Fstl1_Q62356 | 1.01 | -0.4 | 0.63 | 0.8 | -0.8 | 1.51 | -0.6 | -1.4 | -0.8 | Fstl1 |
| Gusb_P12265 | 1.21 | 0.37 | 0.34 | 0.07 | 0.59 | 0.99 | -0.7 | -1.8 | -1.1 | Gusb |
| Tapbp_Q9R233 | -0.4 | 0.67 | 1.34 | -2 | -0.2 | -0.4 | 0.36 | 0.55 | 0.23 | Tapbp |
| Ogdh_Q60597 | 0.34 | -1.5 | 0.29 | -1 | -1.3 | 0.43 | 0.88 | 0.68 | 1.19 | Ogdh |
| Cnpy4_Q8BQ47 | 1.58 | -1.3 | -0.4 | 0.63 | 1.22 | 0.36 | -0.4 | -0.6 | -1 | Cnpy4 |
| Sept2_P42208 | 1.04 | -0.1 | 0.6 | 0.46 | 1.52 | -0.7 | -0.2 | -1.4 | -1.2 | Sept2 |
| Zadh2_Q8BGC4 | 0.34 | -0.4 | -0.5 | -1.6 | 0.28 | -0.6 | 1.19 | 1.71 | -0.4 | Zadh2 |
| Acyp1_P56376 | -0.7 | -0.5 | 1.18 | 0.42 | -1.6 | -1 | 1.08 | 0.26 | 0.91 | Acyp1 |
| Cap1_P40124 | 0.08 | -0.9 | 0.13 | 0.7 | 2 | -0.1 | -0.5 | -0.5 | -1.1 | Cap1 |

| | | | | | | | | | | |
|---|---|---|---|---|---|---|---|---|---|---|
| Stx12_Q9ER00 | -0.2 | -0.3 | 0.45 | 1.01 | -1.4 | -1.6 | 1.11 | 0.32 | 0.71 | Stx12 |
| Idh1_O88844 | -0.9 | 0.38 | 0.51 | -0.2 | -2 | -0.1 | -0.1 | 1.06 | 1.29 | Idh1 |
| Ctsa_P16675 | 1.69 | 0.05 | -0.6 | -0.2 | 1.48 | 0.19 | -0.7 | -0.7 | -1.2 | Ctsa |
| Gstt3_Q99L20 | 0.14 | -1 | 0.1 | -1.5 | 0.5 | -0.7 | 1.89 | 0.44 | 0.2 | Gstt3 |
| S100a4_P07091 | -0.6 | 1.02 | 1.13 | 1.02 | -0.7 | 0.85 | -1.1 | -1.3 | -0.3 | S100a4 |
| Galns_Q571E4 | -0 | 0.42 | 0.34 | 1.71 | -0.3 | 0.47 | 0.33 | -1.6 | -1.3 | Galns |
| Ak2_Q9WTP6 | 1.05 | -0.3 | -0.2 | -0.7 | -0.1 | -1.7 | 1.72 | 0.57 | -0.4 | Ak2 |
| Palld_Q9ET54 | -0.2 | 0.05 | 0.04 | 1.58 | -1 | 1.5 | -0.1 | -1.4 | -0.6 | Palld |
| Htatsf1_Q8BGC0 | -1.4 | 1.11 | 0.31 | -0.2 | -1.3 | -0.5 | -0.2 | 1.27 | 1.01 | Htatsf1 |
| Tkt_P40142 | 1.04 | -0.5 | -0.6 | -1 | -0.1 | -1.3 | 1.67 | 0.93 | -0.2 | Tkt |
| Ube2l3_P68037 | 0.65 | 1.68 | -0 | 1.11 | -0.3 | -0 | -1.4 | -0.8 | -1 | Ube2l3 |
| Ndufb11_O09111 | 0 | -1.4 | 0.39 | 0.06 | -0.4 | -1.2 | 0.96 | 1.81 | -0.2 | Ndufb11 |
| Foxk1_P42128 | -1.2 | 0.68 | 0.8 | 1.46 | -0.5 | 1.08 | -0.6 | -0.7 | -1 | Foxk1 |
| Epb41l1_Q9Z2H5 | -1.5 | 0.47 | 0.51 | 0.1 | -1.9 | -0 | 0.46 | 0.85 | 0.96 | Epb41l1 |
| Dpysl2_O08553 | 1.08 | 0.67 | 0.66 | 1.41 | -0.6 | 0.03 | -1.1 | -1.3 | -0.8 | Dpysl2 |
| Ywhaz_P63101 | 0.4 | 1 | 0.01 | 1.36 | 0.21 | 0.43 | -1.8 | -1 | -0.5 | Ywhaz |
| Khsrp_Q3U0V1 | 1.68 | -0 | -0.1 | -0.1 | 1.28 | 0.11 | -0.8 | -1.6 | -0.4 | Khsrp |
| Gdi2_Q61598 | 1.21 | -0.1 | -0.5 | 0.29 | 1.9 | -0.4 | -0.4 | -0.7 | -1.3 | Gdi2 |
| Basp1_Q91XV3 | -1.3 | 0.86 | 0.85 | -0.2 | -1.7 | -0.5 | 0.33 | 0.4 | 1.2 | Basp1 |
| Aldh3a2_P47740 | 0.77 | 0.46 | -0.7 | -0.2 | 2 | -0.1 | -0.2 | -1 | -1.1 | Aldh3a2 |
| Kars_Q99MN1 | 0.17 | -0 | -1 | -1.6 | -0.1 | -0.6 | 1.09 | 1.66 | 0.4 | Kars |
| Slc3a2_P10852 | 1.38 | 0.17 | 0.13 | -0.7 | 1.52 | 0.35 | -0.4 | -1.3 | -1.1 | Slc3a2 |
| Etfb_Q9DCW4 | 1.65 | -0.3 | -0.9 | -1.3 | -0.8 | -0.2 | 1.36 | 0.22 | 0.29 | Etfb |
| Slc39a7_Q31125 | 0.51 | 0.51 | 0.88 | -0.3 | -2 | -0.6 | -0.5 | 0.14 | 1.37 | Slc39a7 |
| Adk_P55264 | 0.81 | -0.1 | -0.3 | 0.43 | 0.19 | 1.3 | 0.69 | -1.3 | -1.8 | Adk |
| Mut_P16332 | -1.4 | -0 | -1 | 0.11 | -0.9 | 0.37 | 0 | 1.42 | 1.42 | Mut |
| Arpc2_Q9CVB6 | 0.72 | -1.4 | 0.53 | -0.1 | 1.44 | 0.84 | -0.3 | -0.2 | -1.5 | Arpc2 |
| Cxcl12_P40224 | 0.76 | -0.6 | -0.2 | 0.25 | 2 | -0.4 | -0.4 | -0.8 | -0.9 | Cxcl12 |
| Eno1_P17182 | 1.46 | -0.2 | -0.4 | -0.1 | 1.67 | 0.23 | -0.3 | -1.3 | -1 | Eno1 |
| Rpl7a_P12970 | 1.02 | 0.85 | -0.2 | -0.1 | 1.43 | -0 | -0.6 | -0.5 | -1.9 | Rpl7a |
| Tars_Q9D0R2 | 0.87 | 0.33 | 1.15 | 0.17 | -0.5 | 1.2 | -1.7 | -0.6 | -1 | Tars |
| Psmd7_P26516 | 1.17 | -1.4 | -0.6 | -0.6 | -0.3 | -0.9 | 1.24 | 1.16 | 0.27 | Psmd7 |
| Pnp_P23492 | -0.4 | 0.82 | 0.57 | 1.18 | -0.8 | 1.21 | -1.7 | -0.7 | -0.2 | Pnp |
| Skp1_Q9WTX5 | 1.05 | 1.51 | -0.9 | -0.2 | 0.74 | 0.7 | -0.8 | -1 | -1 | Skp1 |
| Hspg2_Q05793 | -0.5 | 1.24 | 0.56 | 1.11 | -0.4 | 0.72 | -1.6 | -1.2 | 0.01 | Hspg2 |
| Bclaf1_Q8K019 | -0.3 | 0.76 | 0.87 | 1.06 | -0.7 | 0.99 | -1.8 | -0.1 | -0.8 | Bclaf1 |
| Actbl2_Q8BFZ3 | 0.58 | 1.39 | -0.4 | -1.1 | -0.5 | -1.1 | -0 | -0.4 | 1.64 | Actbl2 |
| Cyfip1_Q7TMB8 | 0.64 | -0.3 | -0.1 | 0.8 | 1.99 | -0.9 | -1.3 | -0.4 | -0.4 | Cyfip1 |
| Emd_O08579 | 1.14 | -0.3 | -0.5 | -0.5 | 0.98 | 1.57 | -0.2 | -0.7 | -1.5 | Emd |
| Nid1_P10493 | 1.26 | -0.1 | -0.8 | 0.38 | 1.24 | 0.24 | 0.24 | -0.5 | -1.9 | Nid1 |
| Col5a1_O88207 | -0.1 | 1.53 | 0.12 | 0.19 | -0.3 | 1.52 | -1.3 | -0.5 | -1.1 | Col5a1 |
| Ndufb10_Q9DCS9 | -1.4 | 0.24 | 1.04 | 0.57 | -1.1 | -1.4 | 0.88 | 0.54 | 0.6 | Ndufb10 |
| Syncrip_Q7TMK9 | 1.01 | 1.21 | -0.5 | -0.3 | -1.6 | -1.1 | 0.13 | 1.15 | 0.07 | Syncrip |
| Hyou1_Q9JKR6 | -0.7 | 0.6 | 0.82 | 0.32 | -2 | -0.8 | 0.43 | 1.18 | 0.22 | Hyou1 |
| Rad23a_P54726 | 0.66 | 0 | -1.3 | -1.1 | 0.43 | -1 | -0.3 | 1.41 | 1.18 | Rad23a |
| Enah_Q03173 | -0.7 | 1.44 | 0.62 | 0.38 | 0.99 | -0.1 | 0.01 | -0.8 | -1.8 | Enah |
| Prdx6_O08709 | 1.42 | 0.02 | 0.21 | -0.8 | 0.68 | 1.28 | -0.7 | -0.6 | -1.6 | Prdx6 |
| Ubqln1_Q8R317 | -0.2 | 0.94 | 0.2 | -0.1 | 1.24 | 0.35 | -2 | 0.58 | -0.9 | Ubqln1 |
| Abhd12_Q99LR1 | 0.08 | -0.9 | 0.97 | 0.96 | 0.41 | 0.54 | -1 | 0.73 | -1.9 | Abhd12 |
| Snd1_Q78PY7 | 1.75 | -0.9 | -0.5 | 0.79 | 0.71 | 0.54 | -0.6 | -0.5 | -1.4 | Snd1 |

| | | | | | | | | | | |
|---|---|---|---|---|---|---|---|---|---|---|
| Eif3i_Q9QZD9 | -0.7 | 0.9 | 1.51 | -0.5 | -1.9 | -0.4 | 0.3 | 0.25 | 0.52 | Eif3i |
| Psmg1_Q9JK23 | -1.5 | 0.72 | -0.8 | -0.3 | -1.4 | 0.65 | 0.9 | 0.82 | 0.88 | Psmg1 |
| Igf2bp2_Q5SF07 | 0.9 | 0.99 | -0.7 | 0.22 | 0.58 | 0.83 | 0.22 | -1.6 | -1.4 | Igf2bp2 |
| Msn_P26041 | 0.22 | 0.26 | 1.04 | 1.33 | -0.2 | 0.45 | -2 | -0.8 | -0.3 | Msn |
| Sec31a_Q3UPL0 | -0.6 | 0.06 | -0.2 | 0.07 | 1.06 | 1.91 | -1.4 | -0.9 | 0.05 | Sec31a |
| Ugdh_O70475 | 1 | 0.15 | 0.42 | -0.1 | 1.79 | -0.5 | -1.6 | -0.8 | -0.3 | Ugdh |
| Ddb1_Q3U1J4 | -0.2 | 1.08 | 1.41 | -1.1 | -1.3 | -0.5 | -0.8 | 0.63 | 0.78 | Ddb1 |
| P4ha1_Q60715 | 1.73 | -0.2 | 0.33 | -0.1 | 1.42 | -0.3 | -0.8 | -1.1 | -0.9 | P4ha1 |
| Gmps_Q3THK7 | 1.13 | 0.53 | -1.8 | 1.19 | 0.69 | 0.2 | -0.7 | -0.6 | -0.7 | Gmps |
| Phb2_O35129 | -0.8 | 0.42 | -1 | -0.9 | -1.1 | 0.4 | 0.2 | 1.35 | 1.45 | Phb2 |
| Cotl1_Q9CQI6 | 0.92 | 0.74 | -0.4 | 0.88 | 1.32 | -0.8 | -1.4 | -1.2 | -0.1 | Cotl1 |
| Pck2_Q8BH04 | 1.81 | -0.2 | -0.3 | 0.52 | 1.31 | -0.5 | -0.6 | -1 | -1 | Pck2 |
| Park7_Q99LX0 | -0.1 | -0.3 | 1.37 | 0.62 | 0.13 | 1.42 | -0.6 | -1.2 | -1.4 | Park7 |
| Nqo2_Q9JI75 | 1.03 | -0.9 | 1.06 | -0.2 | 1.65 | -0.1 | -1.1 | -0.6 | -0.7 | Nqo2 |
| Ppfibp1_Q8C8U0 | 1.1 | -0.3 | -1.2 | -0.5 | 1.74 | 0.92 | -0.3 | -0.8 | -0.7 | Ppfibp1 |
| Asna1_O54984 | -0.5 | 1.29 | -0.8 | -0.2 | 0.99 | 1.25 | -1.1 | -1.2 | 0.33 | Asna1 |
| Galk2_Q68FH4 | 1.3 | 0.42 | -0.9 | -0.6 | 1.14 | 1.06 | -0.5 | -1.5 | -0.5 | Galk2 |
| Hsp90b1_P08113 | 1.29 | -0.2 | -0.4 | -0.9 | 0.16 | -1.9 | 1.16 | 0.28 | 0.53 | Hsp90b1 |
| Lsm8_Q6ZWM4 | -0.5 | 0.7 | -0.2 | -0.5 | -2 | 0.52 | 1.53 | 0.65 | -0.2 | Lsm8 |
| Otub1_Q7TQI3 | 1.96 | 0.04 | -0.8 | 0.69 | 0.57 | 0.16 | -0.6 | -1.4 | -0.6 | Otub1 |
| Atox1_O08997 | 1.1 | 0.49 | -0.6 | -0.1 | 0.92 | 0.77 | 0.47 | -1.5 | -1.5 | Atox1 |
| Pcna_P17918 | -0.5 | 0 | 1.18 | 0.79 | -0.3 | 1.27 | -1.7 | -1 | 0.31 | Pcna |
| Gas1_Q01721 | -0.5 | 1.73 | 1.14 | -0.5 | 0.57 | 0.37 | -0.8 | -1.1 | -1 | Gas1 |
| Pgam1_Q9DBJ1 | 1.14 | -0.4 | -0.8 | 0.37 | 1.99 | 0.01 | -0.7 | -0.9 | -0.7 | Pgam1 |
| Psma4_Q9R1P0 | 0.85 | -0.7 | -0.4 | -0.7 | 0.84 | -2 | 0.69 | 0.8 | 0.6 | Psma4 |
| Picalm_Q7M6Y3 | -1.4 | 0.48 | 0.54 | 0.21 | -1.9 | -0.1 | 0.08 | 0.95 | 1.06 | Picalm |
| Mtch2_Q791V5 | -0.1 | -1.5 | -0.2 | 0.07 | 0.72 | -1.6 | 1.06 | 0.54 | 1 | Mtch2 |
| Rpl17_Q9CPR4 | -0.1 | 1 | 0.32 | 0.56 | -2 | -0.9 | 0.45 | 0.77 | 0.16 | Rpl17 |
| Pcolce_Q61398 | 1.29 | -0.1 | -0.5 | -0.1 | 1.18 | 0.64 | -0.6 | -2 | 0.14 | Pcolce |
| Ptges_Q9JM51 | -1.6 | 0.45 | 0.57 | 0.05 | -1.8 | 0.11 | 0.22 | 0.99 | 0.93 | Ptges |
| Adprhl2_Q8CG72 | -1.8 | 0.65 | -0.1 | 1.71 | 0.61 | 0.16 | -0.4 | -1 | 0.12 | Adprhl2 |
| _Q8BWQ6 | 1.15 | -0.2 | -0.2 | 0.68 | -2 | -0.7 | 0.62 | 0.53 | 0.35 | |
| Tardbp_Q921F2 | -0.8 | 0.03 | -0.2 | 1.71 | -0.1 | 1.38 | -0.9 | -1.3 | 0.14 | Tardbp |
| Tns3_Q5SSZ5 | 0.75 | 1.16 | 0.67 | 1 | -0.7 | 0.15 | -0.3 | -1.8 | -1 | Tns3 |
| Mogs_Q80UM7 | 1.96 | 0.62 | -0.4 | -0.2 | 0.16 | 0.79 | -0.8 | -1.3 | -0.8 | Mogs |
| Psme2_P97372 | 1.54 | -1 | -0.2 | -0.2 | 0.72 | 1.3 | -0.9 | -1.2 | 0.01 | Psme2 |
| Akr1a1_Q9JII6 | 1.57 | -0.1 | -1 | 0.22 | 1.48 | 0.09 | -0.2 | -0.8 | -1.3 | Akr1a1 |
| Hrsp12_P52760 | 1.2 | 0.64 | -0.2 | -0.8 | 1.7 | 0.13 | -0.8 | -0.7 | -1.2 | Hrsp12 |
| Stat5b_P42232 | -1.6 | 0.33 | 0.75 | 0.73 | 1.76 | -0.2 | -0.4 | -0.6 | -0.7 | Stat5b |
| Igbp1_Q61249 | -1.7 | 0.57 | 0.86 | -0.1 | -1.1 | -0.7 | -0.2 | 1.31 | 0.9 | Igbp1 |
| Atp5h_Q9DCX2 | 0.83 | -0.2 | -1.2 | -1.6 | 0.34 | -0.5 | 1.51 | 0.8 | 0.02 | Atp5h |
| Hnrnpk_P61979 | -0.5 | 1.23 | 1.35 | 0.54 | 0.03 | 0.68 | -1.2 | -1 | -1 | Hnrnpk |
| Spryd4_Q91WK1 | 1.57 | -0.5 | -0.3 | -0.1 | 1.67 | -0.2 | -1 | -0 | -1.1 | Spryd4 |
| Psma6_Q9QUM9 | 2 | 0.15 | -0.3 | -1 | -0.4 | -1.5 | 0.34 | 0.04 | 0.44 | Psma6 |
| Lamtor1_Q9CQ22 | 0.32 | 0.08 | 0.64 | 0.29 | 0.01 | -2 | 0.29 | 0.48 | 0.51 | Lamtor1 |
| Csde1_Q91W50 | -1.8 | 0.69 | 0.72 | 0.19 | -1.5 | -0.4 | 0.52 | 0.76 | 0.77 | Csde1 |
| Rpl18_P35980 | 1.12 | 0.06 | -0.3 | -1.4 | -1.1 | -0.2 | 1.72 | 0.6 | -0.6 | Rpl18 |
| Nap1l1_P28656 | 0.7 | -0.2 | 0.83 | -2 | -0.3 | -0.2 | 1.54 | -0.2 | -0.1 | Nap1l1 |
| Ube2h_P62257 | 1.49 | -0.3 | -0.7 | 0.25 | 1.11 | 0.36 | 0.47 | -1.3 | -1.3 | Ube2h |
| Gtf2i_Q9ESZ8 | 0.54 | 0.14 | -0.5 | 0.3 | 1.73 | -0.2 | -0.3 | 0.31 | -2 | Gtf2i |

| | | | | | | | | | | |
|---|---|---|---|---|---|---|---|---|---|---|
| Glb1_P23780 | 1.59 | 0.34 | -0.8 | -0.4 | 1.1 | 0.67 | -0.2 | -1.5 | -0.8 | Glb1 |
| Usp9x_P70398 | 1.96 | -0.2 | -0.4 | 0.32 | 1.26 | -0.5 | -0.7 | -1 | -0.7 | Usp9x |
| Nucb1_Q02819 | -0.7 | 1.08 | -0.9 | -0.8 | -1.3 | -0.1 | 1.15 | 1.42 | 0.07 | Nucb1 |
| Hist1h1d_P43277 | 1.63 | -0.1 | 0.62 | -1 | 1.16 | 0.39 | -0.7 | -0.8 | -1.1 | Hist1h1d |
| Ubr4_A2AN08 | 2 | -0.4 | -0.2 | -0.2 | 1.16 | 0.06 | -0.8 | -1 | -0.6 | Ubr4 |
| Pcbp1_P60335 | 1.57 | 0.4 | 0.01 | -0.7 | 1 | 0.55 | -0.2 | -1.6 | -0.9 | Pcbp1 |
| Gspt1_Q8R050 | -0.1 | 0.97 | 1.04 | -1.3 | -1.4 | 0.13 | -0.6 | 1.32 | -0 | Gspt1 |
| Sept6_Q9R1T4 | 1.9 | -0.6 | -0.5 | 0.03 | 0.33 | 1.02 | -0.2 | -0.6 | -1.5 | Sept6 |
| Crk_Q64010 | 0.39 | 0.39 | 0.04 | 0.34 | 1.11 | 0.08 | 0.44 | -0.3 | -2 | Crk |
| Snx9_Q91VH2 | -0.5 | 1.64 | 0.1 | -0.8 | 0.08 | -1.8 | 0.93 | -0.2 | 0.53 | Snx9 |
| Rpl13_P47963 | -1.3 | 1.02 | 0.16 | 0.6 | -1.7 | -0.7 | 0.36 | 0.68 | 0.91 | Rpl13 |
| Gba_P17439 | -1.7 | -0.5 | 0.72 | 0.12 | 1.29 | 1.2 | -0.7 | -0.7 | 0.29 | Gba |
| Eprs_Q8CGC7 | 0.88 | 0.56 | 0.22 | 0.52 | 0.74 | 0.26 | 0.2 | -2 | -1.4 | Eprs |
| Aifm1_Q9Z0X1 | -1.8 | 0.83 | 0.61 | 0.17 | -1.5 | -0.3 | 0.45 | 0.89 | 0.66 | Aifm1 |
| Farsb_Q9WUA2 | 1.9 | -0.1 | 0.18 | 0.73 | 0.08 | 0.04 | -0.1 | -1.6 | -1.1 | Farsb |
| Stk3_Q9JI10 | -0.2 | -1 | 0.67 | -0.9 | 0.02 | -0.8 | 1.01 | 1.88 | -0.6 | Stk3 |
| Bckdha_P50136 | -0.5 | 1.35 | -1.4 | 0.28 | -0.7 | -1.1 | 0.1 | 0.72 | 1.27 | Bckdha |
| Sfn_O70456 | 0.61 | -0.3 | -1.1 | -0.5 | 0.46 | -1.6 | -0.1 | 1.13 | 1.44 | Sfn |
| Gstm1_P10649 | 1.7 | 0.08 | -0.3 | 0.22 | 1.36 | -0.5 | -0.8 | -1.5 | -0.3 | Gstm1 |
| Oxct1_Q9D0K2 | -0.4 | 0.97 | 0.88 | -2 | -0.9 | -0.1 | 0.48 | 0.85 | 0.27 | Oxct1 |
| Acp1_Q9D358 | 0.52 | 0.32 | 0.61 | 1.17 | 1.14 | -1.2 | -1.6 | -0.3 | -0.6 | Acp1 |
| Mthfd1l_Q3V3R1 | 1.66 | 0.92 | -0.3 | 0.25 | 1.04 | -0.8 | -1.1 | -0.7 | -0.9 | Mthfd1l |
| Tsfm_Q9CZR8 | -1.1 | 0.82 | -1 | 0.16 | 0.23 | 1.99 | -1 | 0.01 | -0.1 | Tsfm |
| Uggt1_Q6P5E4 | 1.72 | -0.7 | 0.46 | -0.6 | -0.1 | -1.8 | 0.57 | 0.53 | -0 | Uggt1 |
| Iars2_Q8BIJ6 | 1.01 | -1.6 | 0.35 | 0.07 | -0.8 | -0.9 | 1.58 | 0.21 | 0.2 | Iars2 |
| Hist1h2bp_Q8CGP2 | 1.94 | -0.4 | 0.14 | -0.2 | 1.03 | 0.14 | -0.3 | -1.3 | -1.1 | Hist1h2bp |
| Psmc3_O88685 | -1.2 | 0.84 | -0.5 | 0.05 | 0.36 | -1.8 | 0.56 | 0.39 | 1.32 | Psmc3 |
| Sh3bgrl3_Q91VW3 | -0.3 | 0.46 | 0.15 | 1.21 | -0.7 | 1.22 | -1.5 | -1.3 | 0.71 | Sh3bgrl3 |
| L3hypdh_Q9CXA2 | -0.4 | 0.44 | 0.38 | -2 | 0.15 | 0.14 | -0.1 | 0.9 | 0.93 | L3hypdh |
| Arf1_P84078 | 0.58 | 0.01 | 1.7 | 0.48 | 0.87 | -0.8 | -1.5 | -0.7 | -0.7 | Arf1 |
| Ddx1_Q91VR5 | -1 | 1.06 | -0.4 | -0.7 | -1.5 | 0.42 | 0.77 | 1.49 | -0.2 | Ddx1 |
| Cops3_O88543 | 1.37 | -0.6 | -0.3 | 0.46 | 1.85 | -0.7 | -0.9 | -0.7 | -0.4 | Cops3 |
| Mfsd1_Q9DC37 | -1.1 | 0.11 | 0.53 | 0.8 | -2 | -0.3 | 0.29 | 0.92 | 0.82 | Mfsd1 |
| Gng12_Q9DAS9 | -0.3 | -0 | 1.02 | 1.94 | 0.09 | 0.05 | -1.5 | -0.7 | -0.6 | Gng12 |
| Clca3a1_Q9QX15 | -0.9 | 1 | 0.76 | -1.8 | 0.2 | -0.7 | 0.57 | -0.4 | 1.21 | Clca3a1 |
| Rpl35a_O55142 | -0.1 | 0.04 | -0.5 | 0.92 | 2 | -1 | -0 | -0.8 | -0.8 | Rpl35a |
| Copz1_P61924 | 1.68 | -1.1 | -0.5 | -0.1 | 1.51 | 0.41 | -0.4 | -0.7 | -0.7 | Copz1 |
| Anxa5_P48036 | 2 | 0.16 | -0.2 | -0.4 | 0.86 | 0.21 | -0.6 | -1 | -1 | Anxa5 |
| Rps21_Q9CQR2 | 0.87 | -0.2 | -0.2 | 1.87 | 0.86 | -0.7 | -0.7 | -0.5 | -1.3 | Rps21 |
| Cand1_Q6ZQ38 | 0.24 | 0.88 | 0.58 | -0.5 | 1.54 | 0.59 | -0.8 | -1.4 | -1.2 | Cand1 |
| Rpl19_P84099 | 1.12 | -0.1 | 1.46 | -0.1 | 1.14 | -0.6 | -1.3 | -0.8 | -0.8 | Rpl19 |
| Luc7l2_Q7TNC4 | -1.4 | 1.06 | 0.47 | -0.4 | -1.7 | 0.03 | 0.16 | 1.04 | 0.73 | Luc7l2 |
| Psmd1_Q3TXS7 | -0.4 | 0.53 | -0.3 | -1.5 | 0.88 | -1.3 | 1.14 | 1.17 | -0.2 | Psmd1 |
| Cox7c_P17665 | -1.5 | 0.68 | 0.47 | 0.15 | -1.6 | -0.1 | -0.1 | 0.9 | 1.21 | Cox7c |
| Mlec_Q6ZQI3 | 1.62 | -0.1 | 0.41 | 0.52 | 0.62 | 0.3 | -0.6 | -1.3 | -1.5 | Mlec |
| Nhp2l1_Q9D0T1 | 1.72 | -0.6 | -0.7 | -0.4 | 1.42 | 0.49 | -0.2 | -1.1 | -0.8 | Nhp2l1 |
| Ppib_P24369 | 0.82 | 0.39 | 0.37 | -0.5 | 1.77 | -0.2 | -0.6 | -0.2 | -1.8 | Ppib |
| Atp5j2_P56135 | 0.28 | 0.19 | -1 | -2 | 0.06 | 0.57 | 0.75 | 0.84 | 0.51 | Atp5j2 |
| Lrpprc_Q6PB66 | 1.25 | 0.38 | -0.4 | -0.9 | 0 | -1.7 | 1.33 | 0.6 | -0.5 | Lrpprc |
| Matr3_Q8K310 | 1.81 | -0.5 | -1.5 | 0.54 | 0.53 | 0.81 | -0.3 | -0.7 | -0.8 | Matr3 |

| | | | | | | | | | | |
|---|---|---|---|---|---|---|---|---|---|---|
| Dpp3_Q99KK7 | 1.72 | -0.4 | 1.17 | 0.2 | 0.52 | -0.1 | -1 | -1 | -1.2 | Dpp3 |
| Mrpl12_Q9DB15 | -0.1 | -1.5 | 0.5 | -0.9 | 0.06 | -0.4 | 0.36 | 2 | -0.1 | Mrpl12 |
| Ccar1_Q8CH18 | 1.35 | 0.14 | 1.39 | -1.7 | 0.14 | -1.1 | -0.1 | -0 | -0.1 | Ccar1 |
| Gorasp2_Q99JX3 | 0.56 | -1.2 | 1.62 | -0.3 | -0.3 | -1.7 | 0.46 | 0.2 | 0.62 | Gorasp2 |
| Atp6v1h_Q8BVE3 | 1.09 | -0.4 | 0.59 | -0.3 | 1.89 | -0.2 | -0.6 | -0.9 | -1.2 | Atp6v1h |
| Mat2a_Q3THS6 | 1.21 | -1.4 | -0.2 | -0.2 | 0.62 | 1.54 | 0.24 | -1 | -0.9 | Mat2a |
| Myh9_Q8VDD5 | -0.4 | 1.24 | 1.11 | 1.09 | -0.5 | 0.26 | -1.5 | -1.1 | -0.1 | Myh9 |
| Psmd4_O35226 | -1 | -0.1 | 0.22 | -1.2 | -1.3 | 1.14 | 0.09 | 0.89 | 1.28 | Psmd4 |
| Ehd2_Q8BH64 | 0.61 | -0.2 | 0.18 | -1.2 | -0.2 | -1.1 | 1.94 | 0.76 | -0.8 | Ehd2 |
| Nars_Q8BP47 | -0.8 | 0.91 | 0.72 | 0.19 | -2 | -0.1 | 0.05 | 0.5 | 0.74 | Nars |
| Anxa4_P97429 | 1.25 | -0.3 | -0.2 | -1.1 | -1.5 | -0.4 | 0.14 | 1.57 | 0.52 | Anxa4 |
| Psmc4_P54775 | 0.26 | 0.8 | 1.42 | -1.1 | 0.93 | 0.57 | -1 | -0.7 | -1.1 | Psmc4 |
| Uqcr10_Q8R1I1 | 0.64 | -0.5 | -0.7 | -0.2 | 1.05 | 1.24 | 0.73 | -0.4 | -1.9 | Uqcr10 |
| Psip1_Q99JF8 | 0.91 | -0.1 | -1.2 | -0.7 | 1.3 | 1.42 | -0.4 | 0.01 | -1.1 | Psip1 |
| Myh14_Q6URW6 | 1.06 | -0.2 | -0.1 | 0.92 | 1.68 | -1.2 | -0.3 | -0.9 | -0.9 | Myh14 |
| Pafah1b2_Q61206 | 0.86 | 0.63 | 1.33 | 0.15 | -0.8 | 0.91 | -0.5 | -1.4 | -1.1 | Pafah1b2 |
| Lypla2_Q9WTL7 | 1.18 | -1.9 | 0.71 | 0.58 | 0.31 | 0.92 | -0.4 | -0.9 | -0.5 | Lypla2 |
| Hspa4_Q61316 | -1.2 | 0.01 | 0.64 | 0.08 | -2 | 0.06 | 0.89 | 0.63 | 0.91 | Hspa4 |
| Sod2_P09671 | 1.5 | -0.7 | -0.3 | -0.3 | 1.36 | 0.61 | -0.4 | -0 | -1.6 | Sod2 |
| Plin2_P43883 | 1.3 | 1.68 | -0.5 | -0.1 | 0.14 | 0.26 | -0.7 | -1.6 | -0.5 | Plin2 |
| Dcakd_Q8BHC4 | -2 | 0.82 | -0.3 | 0.26 | -1.1 | -0.1 | 0.55 | 1.02 | 0.8 | Dcakd |
| Dync1li1_Q8R1Q8 | 1.49 | -0.3 | 0.02 | 0.15 | 1.43 | -0.4 | -1.6 | -0.9 | 0.11 | Dync1li1 |
| Ythdf1_P59326 | -0.1 | -0.8 | -1.2 | 0.06 | 0.6 | 1.87 | 0.99 | -0.6 | -0.9 | Ythdf1 |
| Hebp1_Q9R257 | 1.63 | -0.1 | 0.12 | -0.9 | 1.22 | 0.64 | -0.6 | -1.3 | -0.7 | Hebp1 |
| Ywhae_P62259 | 1.95 | -0.4 | -0.4 | -0.1 | 1.36 | -0.1 | -0.4 | -1.1 | -0.8 | Ywhae |
| Txndc17_Q9CQM5 | 0.21 | 0.69 | 1.49 | -1.3 | -0.9 | -0.4 | -1.2 | 0.98 | 0.42 | Txndc17 |
| Ccdc186_Q8C9S4 | 0.23 | 0.34 | 0.64 | 0.42 | -2 | -0 | 0.34 | 0.25 | 0.43 | Ccdc186 |
| Znf512_Q69Z99 | -0.5 | 0.38 | 1.3 | -1.6 | -1.2 | 0.57 | 0.26 | -0.4 | 1.18 | Znf512 |
| Lgals9_O08573 | 0.48 | 0.84 | 1.05 | 1.25 | -1.2 | 0.16 | -0.4 | -0.7 | -1.6 | Lgals9 |
| Capn2_O08529 | -1.5 | 1.13 | 0.71 | -0.4 | -1.6 | 0.08 | 0.11 | 0.76 | 0.74 | Capn2 |
| Surf4_Q64310 | 1.14 | 0.7 | 0.21 | 0.14 | -0.1 | 0.51 | 0.72 | -1.7 | -1.5 | Surf4 |
| _Q9D1K7 | 1.35 | -0.2 | -1.1 | -0.4 | 1.01 | 1.1 | 0.41 | -1 | -1.2 | |
| Rars_Q9D0I9 | 0.71 | -0.3 | -0.1 | 2 | 0.16 | 0.11 | -1.6 | -0.2 | -0.8 | Rars |
| Col6a1_Q04857 | -1.6 | 0.57 | 0.67 | -0 | -1.8 | 0.26 | 0.62 | 0.65 | 0.68 | Col6a1 |
| Cst3_P21460 | 1.1 | 0.4 | -1 | 1.58 | -0 | 0.14 | -0.2 | -0.3 | -1.8 | Cst3 |
| C1qtnf5_Q8K479 | 0.67 | -0.2 | 0.6 | 1.73 | 0.03 | -0.6 | -0.4 | -1.9 | 0.07 | C1qtnf5 |
| Ldha_P06151 | 1.76 | -0.5 | -0 | 0.41 | 1.2 | -0.4 | -0.2 | -1.4 | -0.9 | Ldha |
| Arpc1b_Q9WV32 | 1.64 | 0.68 | 0.34 | -0.5 | -0.4 | 1 | -0.2 | -1.2 | -1.4 | Arpc1b |
| Pds5b_Q4VA53 | 0.54 | 0.14 | -0.1 | -0.1 | 0.81 | 0.63 | -2 | 0.26 | 0.3 | Pds5b |
| Set_Q9EQU5 | 0.38 | 0.62 | 0.82 | 0.82 | 0.21 | -0.2 | -2 | -0.5 | 0.19 | Set |
| Ube2v2_Q9D2M8 | -2 | 1.15 | 0.2 | -0.5 | -0.5 | -0.4 | 1.3 | 0.42 | 0.3 | Ube2v2 |
| Hnrnph1_O35737 | 0.1 | 0.5 | 0.44 | -0.2 | -2 | -0 | -0.2 | 0.27 | 1.44 | Hnrnph1 |
| Sec61b_Q9CQS8 | 0.79 | -0.1 | -1.4 | -0.2 | 1.35 | 0.86 | 0.77 | -1.1 | -1 | Sec61b |
| Abhd14b_Q8VCR7 | -1.2 | 1.1 | 0.24 | 1.16 | -0.6 | 1.23 | -1.3 | -0.5 | -0.1 | Abhd14b |
| H3f3a_P84244 | 1.94 | -0.3 | -0 | -0.2 | 1.37 | -0.3 | -0.5 | -0.9 | -1 | H3f3a |
| Steap3_Q8CI59 | 1.76 | 0.24 | -0.2 | -0.3 | 1.3 | 0.05 | -0.5 | -0.9 | -1.4 | Steap3 |
| Rpl7_P14148 | 0.86 | 1.21 | -0.4 | -0.7 | 1.26 | 0.66 | -0.7 | -0.7 | -1.5 | Rpl7 |
| Taco1_Q8K0Z7 | 0.66 | 1.28 | 0.16 | -0.7 | -0 | -1.8 | 0.43 | 1.02 | -1 | Taco1 |
| Nudt2_P56380 | -1.9 | 0.06 | 0.83 | -0.7 | -1 | 0.56 | 0.62 | 0.53 | 1.05 | Nudt2 |
| Gpx8_Q9D7B7 | 1.79 | -1.1 | 0.45 | -0.1 | 0.69 | 0.43 | 0.11 | -1.1 | -1.3 | Gpx8 |

| | | | | | | | | | | |
|---|---|---|---|---|---|---|---|---|---|---|
| Npc1_O35604 | 0.23 | 0.56 | 0.14 | 0.4 | -2 | 0.14 | 0.55 | 0.24 | 0.38 | Npc1 |
| Cltb_Q6IRU5 | -1.8 | 0.71 | 0.59 | 0.28 | 0.57 | 1.24 | -0.5 | -1.2 | 0.18 | Cltb |
| Mre11a_Q61216 | -1.1 | 1.34 | 0.95 | -0.5 | -1.7 | -0.1 | -0.1 | 0.9 | 0.26 | Mre11a |
| Snap23_O09044 | 0.39 | 0.76 | 0.16 | 0.83 | -0.8 | 0.97 | -1.9 | -1 | 0.6 | Snap23 |
| Ptpn11_P35235 | 1.27 | 0.08 | -0.6 | 1.07 | 0.94 | -0.6 | 0.09 | -0.4 | -1.9 | Ptpn11 |
| Ostc_Q78XF5 | -0.6 | 1.01 | -1.2 | 0.58 | -0.1 | -1.7 | 1.23 | 0.25 | 0.59 | Ostc |
| Sh3bp1_P55194 | -0.3 | 0.49 | -0.4 | -0.4 | 1 | 1.11 | 0.85 | -2 | -0.3 | Sh3bp1 |
| Erc1_Q99MI1 | -1.4 | 0.49 | 0.56 | -0.5 | -1.8 | 0.8 | 0.55 | 1.05 | 0.23 | Erc1 |
| Rnh1_Q91VI7 | 1.3 | 0.49 | -1.3 | 0 | 1.62 | 0.08 | -0.9 | -0.8 | -0.5 | Rnh1 |
| Sumo2_P61957 | -0.4 | 0.23 | 1.2 | 0.29 | 1.44 | -0.4 | -1.7 | -1 | 0.33 | Sumo2 |
| Nedd8_P29595 | -0.5 | 0.76 | 0.88 | -2 | -0 | -0 | -0.6 | 0.36 | 1.26 | Nedd8 |
| Clcn7_O70496 | -1.2 | 0.37 | 1.64 | -0.5 | -0.2 | -1.4 | -0.3 | 0.79 | 0.86 | Clcn7 |
| Chchd6_Q91VN4 | -2 | 0.53 | 0.38 | 0.06 | -0.6 | -0.5 | 0.67 | 1.04 | 0.62 | Chchd6 |
| Tmed10_Q9D1D4 | 1.64 | -1 | -0.4 | -0.3 | 1.59 | 0.21 | -0.1 | -0.6 | -1.1 | Tmed10 |
| Sec23a_Q01405 | -1.7 | 0.38 | -0.4 | 0.16 | 1.82 | 0.81 | 0.07 | -0.3 | -0.7 | Sec23a |
| Mapre1_Q61166 | 1.04 | 0.03 | 0.54 | -1.4 | 0.9 | 1.18 | -1.1 | -1.2 | -0 | Mapre1 |
| Fbxo6_Q9QZN4 | 0.57 | -0.4 | 0.56 | -0.1 | 0.08 | -2 | 1.62 | 0.29 | -0.6 | Fbxo6 |
| Ttll12_Q3UDE2 | -0.1 | 0.62 | -0.7 | -0.6 | -0.1 | 2 | -0.6 | -0.4 | -0.6 | Ttll12 |
| Etfdh_Q921G7 | 1.15 | -1.8 | -0 | -1 | 0.33 | -0.6 | 1.26 | 0.58 | 0.15 | Etfdh |
| Rbm39_Q8VH51 | 0.91 | -0.1 | -0.1 | -0.1 | 0.83 | 0.6 | 1.05 | -1.6 | -1.6 | Rbm39 |
| Creg1_O88668 | -1.2 | 0.99 | 0.42 | 0.14 | -1.8 | -0.1 | -0 | 0.24 | 1.4 | Creg1 |
| Pdcd4_Q61823 | -1.1 | 0.62 | 0.18 | -0.1 | 0.6 | 1.34 | -0.7 | -1.7 | 0.91 | Pdcd4 |
| Sord_Q64442 | -1.7 | 1.05 | 0.19 | 0.27 | -1.5 | -0.2 | 0.95 | 0.1 | 0.84 | Sord |
| Mylk_Q6PDN3 | -0.2 | 0.67 | 0.12 | -0.4 | 0.07 | 1.68 | -1.4 | -1.3 | 0.85 | Mylk |
| Anxa3_O35639 | 1.28 | 0.61 | -0.4 | -0.3 | 0.82 | 0.25 | -0.1 | 0.08 | -2 | Anxa3 |
| Rab21_P35282 | 0.67 | -1.2 | -1.1 | 0.31 | 2 | -0 | -0.2 | -0.5 | -0 | Rab21 |
| Hist1h1e_P43274 | 1.92 | -0.7 | 0.52 | 0.24 | 0.03 | 0.64 | -0.3 | -1.6 | -0.7 | Hist1h1e |
| Trmt112_Q9DCG9 | 0.46 | -1.9 | -0.2 | 0.84 | 0.6 | 0.84 | 0.68 | 0.09 | -1.4 | Trmt112 |
| Rab2a_P53994 | -1.4 | 0.76 | 0.43 | 0.17 | -2 | 0.22 | 0.66 | 0.55 | 0.62 | Rab2a |
| Scrn1_Q9CZC8 | 0.4 | 1.33 | 0.46 | 0.03 | -0.5 | 0.76 | -0.2 | -2 | -0.1 | Scrn1 |
| Rab3il1_Q8VDV3 | -0.5 | 1.8 | -0.9 | -0.1 | -1 | -1 | 1.25 | 0.47 | -0.1 | Rab3il1 |
| Psmc1_P62192 | 0.38 | 0.53 | -0.9 | 0.47 | 0.4 | 1.41 | 0.56 | -1.3 | -1.5 | Psmc1 |
| Cbx3_P23198 | 1.23 | 0.6 | 1.24 | 0.29 | 0.57 | -1 | -0.9 | -1.4 | -0.6 | Cbx3 |
| Pgk1_P09411 | 1.57 | 0.12 | 0.47 | 0.47 | 0.46 | 0.2 | -0.2 | -1.5 | -1.6 | Pgk1 |
| Cfl1_P18760 | -0.5 | 1.44 | 0.34 | 1.31 | -0.8 | 0.63 | -0.8 | -1.5 | -0.1 | Cfl1 |
| Rps3_P62908 | 1.93 | 0.08 | -1 | -0.2 | 1.26 | 0.07 | -0.8 | -0.5 | -0.8 | Rps3 |
| Mtx1_P47802 | 1.48 | -0.8 | -1.1 | -0.2 | 0.62 | 1.43 | -0.4 | -1.1 | 0.15 | Mtx1 |
| Trappc6b_Q9D289 | -1.2 | 0.42 | 0.5 | 0.62 | -2 | -0.1 | 0.09 | 0.35 | 1.3 | Trappc6b |
| Dync1h1_Q9JHU4 | 2 | -0.4 | -0.6 | -0 | 1.21 | 0.08 | -0.4 | -0.9 | -1 | Dync1h1 |
| Safb_D3YXK2 | 0.94 | 0.59 | 0.45 | -0.6 | -2 | 0.52 | -0.2 | -0 | 0.67 | Safb |
| Slc25a11_Q9CR62 | 0.58 | 0.04 | 0.67 | -0.1 | 0.27 | -2 | -0 | 0.19 | 0.9 | Slc25a11 |
| Sugt1_Q9CX34 | 0.43 | 1.17 | 1.08 | 0.23 | 0.89 | -0.8 | -1.4 | -1.3 | -0.2 | Sugt1 |
| Pabpc1_P29341 | 1.28 | -1.1 | -0.2 | 0.59 | 1.57 | -0.3 | 0.21 | -0.9 | -1.2 | Pabpc1 |
| Cox5a_P12787 | 0.39 | -0.6 | -0.1 | -2 | -0.4 | 1.1 | 1.2 | 0.43 | 0.12 | Cox5a |
| Prnp_P04925 | -1.6 | -0.7 | -0.8 | 1.54 | 0.18 | 1.02 | -0.7 | 0.74 | 0.2 | Prnp |
| Iqgap1_Q9JKF1 | 1.1 | 0.4 | 0.31 | -0.1 | 1.78 | -1.1 | -0.7 | -0.5 | -1.2 | Iqgap1 |
| Sept9_Q80UG5 | -1.1 | 1.01 | 0.91 | 0.83 | -0.4 | 0.83 | -1.5 | -1 | 0.43 | Sept9 |
| Tmem165_P52875 | 0.99 | -0.2 | 1.14 | -0.1 | -0.8 | 1.29 | -0.3 | -1.8 | -0.3 | Tmem165 |
| Thbs1_P35441 | 1.96 | 0 | -0.1 | 0.28 | 0.52 | -0.2 | -1 | -1.6 | 0.19 | Thbs1 |
| Pycrl_Q9DCC4 | 0.9 | 0.85 | 0.76 | 0.96 | 0.24 | -1.2 | -1.7 | -0.7 | -0.1 | Pycrl |

| | | | | | | | | | | |
|---|---|---|---|---|---|---|---|---|---|---|
| Xpnpep2_B1AVD1 | 0.88 | 0.91 | -1.9 | 0.68 | -0.1 | 1.01 | -0.3 | -0.1 | -1.1 | Xpnpep2 |
| Gls_D3Z7P3 | -0 | -0.8 | 1.34 | 1.65 | 0.74 | -1 | -0.9 | -0.5 | -0.5 | Gls |
| Rps20_P60867 | 1.35 | -0.2 | -0.8 | -1 | 0.84 | -1.7 | 0.7 | 0.58 | 0.26 | Rps20 |
| Ppp2r1a_Q76MZ3 | -1.6 | 1.62 | 0.27 | -0.7 | -0.5 | -0.7 | 0.95 | 0.71 | -0 | Ppp2r1a |
| Vbp1_P61759 | -1.4 | 0.46 | 0.92 | 0.13 | -2 | 0.3 | 0.32 | 0.41 | 0.84 | Vbp1 |
| Ncl_P09405 | -1.5 | 0.97 | 0.41 | 0.26 | -1.6 | -0.2 | -0.2 | 0.74 | 1.19 | Ncl |
| Dnm2_P39054 | 0.84 | 0.11 | -1 | 0.16 | 2 | -0.5 | -0.4 | -1.1 | -0.4 | Dnm2 |
| Eef1b_O70251 | 1.65 | 0.39 | -0.1 | 0.05 | 0.64 | 0.54 | -1.2 | -1.7 | -0.3 | Eef1b |
| Itgav_P43406 | -0.7 | 0.82 | 1 | 1.39 | -1.1 | 0.74 | -0.5 | -1.3 | -0.3 | Itgav |
| Tspan6_O70401 | 1.28 | 1.37 | -1 | 0.5 | 0.5 | -0.1 | -1.5 | -0.3 | -0.7 | Tspan6 |
| Srp68_Q8BMA6 | -0.6 | 1.32 | 0.3 | 0.39 | 0.37 | 0.31 | -2 | -0.2 | 0.4 | Srp68 |
| Dclk1_Q9JLM8 | 1.12 | 1.04 | 0.4 | -0.3 | 1.38 | -1 | -0.6 | -0.8 | -1.2 | Dclk1 |
| Nudt21_Q9CQF3 | 0.6 | 0.41 | -0.2 | -0 | -0.7 | 1.85 | -0.2 | 0.11 | -1.9 | Nudt21 |
| Sec31b_Q3TZ89 | 1.33 | -1 | 1.33 | -0.5 | 1.1 | 0.06 | -1.1 | -0.5 | -0.7 | Sec31b |
| Nagk_Q9QZ08 | 0.57 | -2 | 0.15 | 0.4 | 1.42 | 0.37 | -0.4 | 0.43 | -0.8 | Nagk |
| Hnrnpa2b1_O88569 | -1.6 | 0.39 | 0.47 | 0.35 | -1.8 | 0.29 | 0.16 | 0.74 | 1 | Hnrnpa2b1 |
| Atic_Q9CWJ9 | 1.72 | -0.3 | -0.6 | 0.07 | 1.61 | -0.5 | -0.5 | -0.3 | -1.2 | Atic |
| Lsm12_Q9D0R8 | -0.7 | -0.8 | -0.1 | 1.2 | -1 | 2 | -0.5 | -0.1 | -0.1 | Lsm12 |
| Arpc4_P59999 | 1.79 | 0.99 | -0.4 | -0.7 | 0.48 | 0.42 | -0.4 | -1.4 | -0.6 | Arpc4 |
| Psmc2_P46471 | 0.73 | 0.01 | 1.48 | 0.78 | 0.63 | -1.2 | -0.8 | -0.2 | -1.5 | Psmc2 |
| Srsf7_Q8BL97 | 1.19 | -0.2 | -0.4 | 0.51 | 1.44 | -0.6 | -1.9 | 0.09 | -0.1 | Srsf7 |
| Oplah_Q8K010 | 0.98 | 0.22 | -1 | -0.7 | 0.62 | 1.57 | 0.2 | -1.6 | -0.2 | Oplah |
| Apip_Q9WVQ5 | -1 | 0.21 | 0.58 | 0.55 | -2 | 0.1 | 1.24 | -0.2 | 0.59 | Apip |
| Idh2_P54071 | 1.36 | -0 | 1.59 | -0.6 | 0.21 | 0.28 | -0.4 | -1 | -1.4 | Idh2 |
| Eif1ax_Q8BMJ3 | -0 | -0.2 | 0.37 | 1.6 | 0.7 | -0.6 | 0.11 | 0.15 | -2 | Eif1ax |
| Psmb5_O55234 | -0.6 | -1.3 | 1.29 | -0.9 | 0.38 | -0.9 | 1.4 | -0.2 | 0.77 | Psmb5 |
| Sri_Q6P069 | -0.5 | 1.09 | 1.02 | 0.7 | -0.6 | 0.74 | -1.8 | -0.9 | 0.28 | Sri |
| Prkaca_P05132 | 0.11 | -0.6 | -1 | 0.27 | -0.3 | 2 | -0 | -0.9 | 0.08 | Prkaca |
| Adh5_P28474 | 1.12 | -0.2 | 0.97 | 0.17 | -1.1 | 1.51 | -0.9 | -1 | -0.5 | Adh5 |
| Coro1b_Q9WUM3 | -1.2 | 0.01 | 0.61 | 0.78 | -1.2 | -1 | 1.15 | 1.24 | -0.3 | Coro1b |
| Casp8_O89110 | -0.1 | -1.2 | -0 | 0.94 | 1.17 | -0 | 0.32 | 0.81 | -1.9 | Casp8 |
| Comt_O88587 | 1.25 | 0.66 | 0.18 | -1.4 | 1.23 | 0.53 | -0.7 | -0.6 | -1.1 | Comt |
| Snrnp70_Q62376 | -1.1 | 0.33 | 0.77 | -0.1 | 2 | -0.3 | -0.8 | -0.3 | -0.7 | Snrnp70 |
| Srm_Q64674 | -0.5 | 1.35 | -0.7 | 1.13 | -1 | 1.41 | -1 | -0.4 | -0.3 | Srm |
| Dhrs1_Q99L04 | 0.13 | 0.19 | 0.6 | 0.68 | -0.6 | 0.94 | -2 | 0.46 | -0 | Dhrs1 |
| Cyb5a_P56395 | 1.49 | -0.3 | -1.3 | -0.9 | 0.4 | -1.1 | 1.26 | -0.1 | 0.52 | Cyb5a |
| Uba5_Q8VE47 | 0.98 | 0.54 | -1.4 | -0.7 | -0.3 | -0.6 | 1.53 | 0.82 | -0.9 | Uba5 |
| Eno3_P21550 | 0.42 | 0.9 | -1.9 | 0.81 | 0.69 | 0.35 | 0.58 | -0.8 | -1 | Eno3 |
| Tcp1_P11983 | 1.98 | 0.31 | -0.5 | 0.05 | 1.01 | -0.5 | -0.1 | -1.1 | -1.1 | Tcp1 |
| Acot7_Q91V12 | 1.4 | -0.3 | -0.6 | -0.3 | -0.8 | -0.7 | -0.7 | 1.93 | 0.17 | Acot7 |
| Wwc2_Q6NXJ0 | -0 | 0.41 | 0.47 | 0.28 | -2 | 0.49 | 0.29 | 0.29 | 0.44 | Wwc2 |
| Nhp2_Q9CRB2 | -1.4 | 0.27 | 0.5 | -1.8 | -0.1 | 0.75 | 1.22 | 0.11 | 0.48 | Nhp2 |
| Vdac1_Q60932 | 1.3 | 1.4 | 1.12 | -0.9 | -0.7 | -1.2 | -0.2 | -0.5 | -0.3 | Vdac1 |
| Aldh2_P47738 | 1.06 | 0.73 | -0.6 | -1.2 | 0.07 | -1.3 | 1.15 | 0.91 | -0.8 | Aldh2 |
| Dtymk_P97930 | -1 | 0.17 | 0.26 | 1.24 | 1.69 | -1.1 | -0.3 | -1.1 | 0.09 | Dtymk |
| Erh_P84089 | 1.29 | -0.5 | -0.7 | -0.2 | 1.68 | 0.05 | -0.1 | -1.7 | 0.04 | Erh |
| Pgm2_Q7TSV4 | 1 | 1.42 | 0.19 | -1.6 | -0.1 | -0.8 | -0.8 | -0.3 | 0.97 | Pgm2 |
| Glipr2_Q9CYL5 | 1.64 | -1.1 | 0.38 | -1.6 | 0.22 | -0.5 | 0.68 | -0.2 | 0.59 | Glipr2 |
| Aimp1_P31230 | 1.94 | -0.3 | 0.21 | -0.2 | 1.1 | -0.4 | -1.1 | -1.2 | -0 | Aimp1 |
| Ap2a1_P17426 | 0.47 | -1.1 | 0.98 | 0.03 | 0.61 | 0.7 | 0.46 | -0 | -2 | Ap2a1 |

| | | | | | | | | | | |
|---|---|---|---|---|---|---|---|---|---|---|
| Isoc1_Q91V64 | -0.8 | 0.12 | -0.4 | -0.5 | 1.16 | 1.36 | 1.15 | -1.4 | -0.6 | Isoc1 |
| Fkbp1a_P26883 | 1.35 | 0.36 | -1.4 | -0.5 | 1.28 | 0.63 | 0.06 | -0.6 | -1.2 | Fkbp1a |
| Dars_Q922B2 | 1.66 | -0.8 | -0.8 | 0.28 | 1.63 | -0.4 | -0.2 | -0.4 | -1 | Dars |
| Sar1b_Q9CQC9 | 0.98 | 0.3 | -0.5 | -1.8 | 1.1 | -1.2 | 0.2 | 0.03 | 0.87 | Sar1b |
| Gpc4_P51655 | 1.61 | -1.4 | 0.09 | -0.1 | 1.55 | -0 | -0.5 | -0.5 | -0.6 | Gpc4 |
| Slirp_Q9D8T7 | -0.4 | -0.6 | -0.2 | -0.1 | -1.8 | 1.14 | -0.2 | 0.81 | 1.44 | Slirp |
| Fth1_P09528 | 1.31 | -0.3 | 0.83 | -2 | -0 | -0.3 | 0.78 | 0.41 | -0.6 | Fth1 |
| Eif4h_Q9WUK2 | -1.2 | 1.04 | 0.79 | -0 | -1.3 | -0.6 | -0.8 | 0.9 | 1.18 | Eif4h |
| Qars_Q8BML9 | -2 | 0.27 | 0.96 | -0.3 | -0.4 | -0.4 | -0.2 | 1.43 | 0.65 | Qars |
| Cask_O70589 | -1.1 | 0.72 | 0.74 | -0.2 | 0.49 | 0.97 | 0.11 | -2 | 0.33 | Cask |
| Cope_O89079 | 0.15 | -0.2 | 0.86 | -0.7 | -0.6 | 2 | -0.5 | -0.7 | -0.6 | Cope |
| Uqcrfs1_Q9CR68 | -0.7 | 0.84 | -0.9 | -0.3 | -1.4 | 0.65 | 1.35 | 1.11 | -0.6 | Uqcrfs1 |
| Tfam_P40630 | 0.21 | 0.54 | 0.09 | -1 | 0.77 | 1.27 | 0.2 | 0.04 | -2 | Tfam |
| Aip_O08915 | -1.3 | 0.11 | 0.85 | -1 | 1.16 | 1.53 | -0.5 | -0.7 | -0.1 | Aip |
| Acad9_Q8JZN5 | 1.74 | -0.7 | -1.3 | -0.2 | -1.3 | 0.14 | 0.58 | 0.78 | 0.27 | Acad9 |
| Cd63_P41731 | 0.52 | 1.37 | 0.95 | 0.45 | -1.6 | -1.3 | -0.2 | -0.6 | 0.35 | Cd63 |
| Rmdn3_Q3UJU9 | 0.37 | 0.34 | 0.28 | -2 | 0.34 | 0.44 | 0.24 | 0.33 | 0.32 | Rmdn3 |
| Actr3_Q99JY9 | 0.84 | -0.3 | -2 | -1 | -0.3 | 0.51 | 0.74 | 0.85 | 0.67 | Actr3 |
| Rbm25_B2RY56 | -1.6 | -0.4 | 0.66 | 0.52 | -1.5 | 0.22 | -0.1 | 0.9 | 1.27 | Rbm25 |
| Bag3_Q9JLV1 | -0.8 | -0.3 | -2 | 0.39 | 1.57 | 0.66 | 0.03 | -0.1 | 0.57 | Bag3 |
| Ttyh3_Q6P5F7 | -1.5 | -0.5 | -0.1 | 1.74 | -0.3 | 0.92 | -0.8 | -0.3 | 0.86 | Ttyh3 |
| Aldh7a1_Q9DBF1 | 0.37 | 0.87 | 1.67 | -0.4 | -0.1 | 0.61 | -1.7 | -0.8 | -0.6 | Aldh7a1 |
| Qdpr_Q8BVI4 | 1.15 | -0.3 | -0.4 | -0.8 | 1.03 | -1.9 | 0.42 | 0.93 | -0.1 | Qdpr |
| Itga5_P11688 | -1.9 | 1.24 | 0.24 | 0.88 | 0.28 | 0.58 | -1.2 | 0.29 | -0.5 | Itga5 |
| Cnpy2_Q9QXT0 | 1.81 | -0.7 | -1.1 | 0.76 | 0.7 | 0.06 | -0.7 | 0.34 | -1.2 | Cnpy2 |
| Ppp2ca_P63330 | -1.3 | 0.65 | 0.57 | 0.6 | -2 | 0.12 | 0.28 | 0.56 | 0.61 | Ppp2ca |
| H2afj_Q8R1M2 | -0.4 | 0.73 | 1.51 | 0.6 | 0.57 | -0.6 | 0.29 | -1 | -1.7 | H2afj |
| Clic1_Q9Z1Q5 | -1.1 | 0.33 | 0.59 | 0.18 | -2 | 0.48 | 0.26 | 1.14 | 0.29 | Clic1 |
| Glo1_Q9CPU0 | 0.59 | 1.14 | -0.2 | -1.1 | 1.66 | 0.05 | -1 | -0 | -1.2 | Glo1 |
| Mif_P34884 | 1.88 | -0 | -0 | 0.3 | 1.12 | -0.7 | -1.3 | -1 | -0.3 | Mif |
| Got1_P05201 | 1.7 | 0.18 | -0.8 | 0.83 | 1.03 | -0.8 | -1.1 | -0.9 | -0.1 | Got1 |
| Htra2_Q9JIY5 | 0.88 | -0.9 | -0.6 | 1.06 | 0.32 | 0.32 | 0.65 | -2 | 0.28 | Htra2 |
| Sdpr_Q63918 | -1.3 | 0.64 | 0.65 | 0.48 | -2 | 0.21 | 0.26 | 0.53 | 0.62 | Sdpr |
| Cdkn2a_P51480 | 0.74 | 1.8 | -0.4 | 1.03 | 0.06 | -0.9 | -1.2 | -0.4 | -0.7 | Cdkn2a |
| Eif3b_Q8JZQ9 | 1.64 | -0.9 | -1.1 | 0.32 | 0.34 | 0.99 | 0.59 | -1 | -0.9 | Eif3b |
| Arsa_P50428 | 1.58 | 0.07 | 0.55 | -0.9 | 1.43 | -0.2 | -0.7 | -1.1 | -0.7 | Arsa |
| Phpt1_Q9DAK9 | -1.2 | 0.76 | 0.43 | 1.66 | -0.8 | 0.67 | -0.2 | -1.4 | 0.03 | Phpt1 |
| Tecr_Q9CY27 | -0.2 | 0.1 | 0.08 | 0.86 | -2 | 0.21 | 0.31 | 0.76 | 0.43 | Tecr |
| Gns_Q8BFR4 | 0.71 | 0.77 | 0.79 | -0.7 | 1.33 | -0 | -1.7 | -1 | -0.2 | Gns |
| Gnao1_P18872 | -0.7 | 1.18 | -0.4 | -1.1 | 1.19 | 1.43 | -0.9 | -0.6 | -0 | Gnao1 |
| Akr1b1_P45376 | -0.4 | -0.1 | -1.7 | -1.3 | 0.43 | 0.05 | 1.09 | 1.1 | 0.81 | Akr1b1 |
| Cct3_P80318 | 1.77 | -0.5 | 0.4 | 1.1 | 0.56 | -0.9 | -0.6 | -0.8 | -1 | Cct3 |
| Ap2a2_P17427 | -0 | 0.89 | -0 | -1 | 1.77 | 0.59 | 0 | -0.7 | -1.5 | Ap2a2 |
| Ilkap_Q8R0F6 | 0.05 | -0.2 | 0.88 | 0.89 | -1.4 | 1.56 | -0.2 | -1.4 | -0.2 | Ilkap |
| Sparc_P07214 | 1.65 | 0.52 | -1 | -0.2 | 0.75 | 0.26 | -1 | -1.5 | 0.45 | Sparc |
| Stip1_Q60864 | -1.6 | 0.65 | 0.15 | 0.64 | -1.7 | -0 | 0.15 | 0.89 | 0.89 | Stip1 |
| Tuba1a_P68369 | 1.25 | 1.22 | 0.49 | -0.1 | 0.43 | -0.8 | -1.6 | -1.1 | 0.24 | Tuba1a |
| Paics_Q9DCL9 | 0.01 | 1.2 | -0.4 | 1.58 | -1.1 | 0.64 | -1 | -1 | 0.18 | Paics |
| Hdlbp_Q8VDJ3 | 1.78 | 0.8 | -0.4 | -0.4 | 1.07 | -0.1 | -1.2 | -0.8 | -0.7 | Hdlbp |
| Ugp2_Q91ZJ5 | 1.5 | -0.8 | -0.1 | -0.6 | 1.76 | 0.14 | -0.2 | -0.7 | -1 | Ugp2 |

| | | | | | | | | | | |
|---|---|---|---|---|---|---|---|---|---|---|
| Pea15_Q62048 | -0.7 | 0.7 | 0.95 | 1.01 | -1 | -1.9 | 0.42 | -0.2 | 0.64 | Pea15 |
| Arl8a_Q8VEH3 | 1.53 | -0.7 | 0.08 | -1.4 | 0.86 | -1.3 | 0.73 | 0.23 | 0.03 | Arl8a |
| Psme1_P97371 | -1.5 | 0.33 | 0.54 | 0.47 | -1.6 | -0.1 | -0.1 | 0.7 | 1.35 | Psme1 |
| Rpl30_P62889 | 1.74 | -0.6 | -0.5 | -0.2 | 1.45 | -0 | -0 | -1.4 | -0.4 | Rpl30 |
| Bag5_Q8CI32 | 1.07 | -1 | 0.31 | -1 | 0.22 | -0.9 | -1.2 | 1.13 | 1.22 | Bag5 |
| Prpf6_Q91YR7 | -1.6 | 0.79 | 0.82 | 0.46 | -1.8 | -0.1 | 0.26 | 0.47 | 0.65 | Prpf6 |
| Rpl22_P67984 | 1.02 | -0.1 | -0.9 | -1.1 | 1.55 | 0.96 | 0.25 | -1.1 | -0.6 | Rpl22 |
| Ilf2_Q9CXY6 | -0.6 | 0 | 1.32 | -0.5 | 0.69 | 0.78 | 0.64 | -2 | -0.3 | Ilf2 |
| Arl6ip5_Q8R5J9 | -1.1 | 1.28 | 0.6 | -0.5 | -0.2 | 1.7 | -0 | -1 | -0.7 | Arl6ip5 |
| Cops4_O88544 | 1.44 | -0.7 | -0.1 | 0.04 | 1.83 | -0.8 | -0.9 | -0.8 | -0 | Cops4 |
| Gstm5_P48774 | 1.17 | 0.34 | -0.5 | 0.04 | 1.79 | -1 | -1.4 | -0.4 | -0.1 | Gstm5 |
| Hs1bp3_Q3TC93 | -1.1 | 0.26 | 1.88 | -0.8 | 0.2 | -1.3 | 0.81 | 0.37 | -0.3 | Hs1bp3 |
| Gfpt1_P47856 | 0.01 | -0.3 | -1.5 | -0.1 | 1.23 | 1.52 | -1 | -0.6 | 0.7 | Gfpt1 |
| _Q91V76 | -1.1 | -1.1 | -1.2 | 0.2 | -0.1 | 0.73 | 0.5 | 1.77 | 0.32 | |
| Yars_Q91WQ3 | 1.48 | -1.2 | 0.22 | -0.7 | 0.75 | 1.35 | -0.3 | -0.6 | -0.9 | Yars |
| Dync1i2_O88487 | 0.03 | 1.77 | 0.05 | 0.5 | -0.9 | -1.8 | 0.67 | -0.1 | -0.2 | Dync1i2 |
| Emilin1_Q99K41 | -0.3 | -2 | 0.87 | -0 | 1.68 | 0.39 | 0.11 | -0.5 | -0.3 | Emilin1 |
| Hist1h4a_P62806 | 1.75 | -0.2 | 0.08 | -0.4 | 1.47 | -0.6 | -0.1 | -0.7 | -1.3 | Hist1h4a |
| Akt1_P31750 | -0.6 | 1.73 | 0.98 | -1.2 | 0.14 | -1.4 | -0.4 | 0.37 | 0.3 | Akt1 |
| Rpl6_P47911 | 1.22 | 0.46 | 0.65 | -0.3 | 1.16 | -0.2 | -0.4 | -0.7 | -1.9 | Rpl6 |
| Wars_P32921 | 1.47 | -0.9 | -0.4 | -0.6 | 1.9 | 0 | -0.2 | -0.7 | -0.6 | Wars |
| Erp29_P57759 | 1.56 | 0.6 | -0.3 | -1.6 | -0.6 | -0.1 | 0.85 | 0.62 | -1 | Erp29 |
| Adprh_P54923 | 1.29 | -1 | 1.01 | 0.24 | 0.89 | -0.5 | -1.2 | 0.5 | -1.3 | Adprh |
| Eef1a1_P10126 | 0.78 | -0.5 | -1 | -0.2 | 0.47 | -1.8 | -0.1 | 1.06 | 1.24 | Eef1a1 |
| Rab5a_Q9CQD1 | -2 | 0.45 | 0.82 | 0.06 | -0.9 | 0.05 | 0.21 | 0.64 | 0.93 | Rab5a |
| Gnai2_P08752 | 1.18 | 0.24 | -1.2 | -0.4 | 2 | -0.3 | -0.4 | -0.4 | -0.6 | Gnai2 |
| Rab7a_P51150 | 1.65 | -1 | -1.1 | -0.1 | 1.24 | 0.53 | 0.31 | -0.8 | -0.7 | Rab7a |
| Pkn1_P70268 | -0.8 | -0.9 | -0 | 0.26 | -1.8 | 1.16 | 0.34 | 0.55 | 1.21 | Pkn1 |
| Eif3h_Q91WK2 | 0.18 | 0.48 | 0.42 | 0.43 | 0.38 | -2 | -0 | 0.3 | 0.47 | Eif3h |
| Cops2_P61202 | 1.7 | 0.93 | 0.57 | -1.2 | -0.2 | 0.58 | -1 | -0.9 | -0.5 | Cops2 |
| Scarb2_O35114 | -1.6 | 0.07 | -0.5 | -0.9 | 0.58 | -0.3 | 1.72 | 1.04 | -0.2 | Scarb2 |
| Anp32e_P97822 | -1 | 1.25 | 0.42 | 0.4 | -2 | -0 | 0.83 | -0.3 | 0.48 | Anp32e |
| Eif4a3_Q91VC3 | 0.76 | 0.04 | -1.9 | 1.24 | 0.8 | -0.2 | 0.16 | -1.3 | 0.3 | Eif4a3 |
| Plbd2_Q3TCN2 | 1.8 | 0.23 | 0.2 | 1.18 | 0 | -1 | -0.5 | -0.7 | -1.2 | Plbd2 |
| Flot1_O08917 | -1.9 | 1.36 | 0.15 | -0.1 | -0.6 | -0.4 | -0.3 | 0.89 | 0.97 | Flot1 |
| Abcf1_Q6P542 | -0.4 | 0.51 | 1.51 | -0 | 0.1 | 0.39 | -2 | 0.74 | -0.8 | Abcf1 |
| Psmb7_P70195 | -1.2 | 0.37 | -0.1 | 0.26 | -1.9 | 0.79 | 0.42 | 1.35 | 0.05 | Psmb7 |
| Vps4b_P46467 | -0.1 | -2 | -0.2 | 1.53 | 0.05 | 0.73 | 0.18 | 0.54 | -0.6 | Vps4b |
| Rnpep_Q8VCT3 | -1.8 | 0.86 | 0.44 | 0.78 | -0.6 | 1.38 | -0.5 | -0.9 | 0.26 | Rnpep |
| Rps8_P62242 | 0.91 | 0.76 | -0.8 | 0.27 | 1.43 | -0.5 | 0.39 | -0.7 | -1.7 | Rps8 |
| Pycr2_Q922Q4 | 1.61 | -1.2 | -0.6 | -0.6 | 1.33 | 0.69 | 0.16 | -0.5 | -0.9 | Pycr2 |
| Glod4_Q9CPV4 | -0.2 | 0.06 | 1.13 | -1.9 | -0.7 | 0.42 | 0.51 | 1.31 | -0.6 | Glod4 |
| Echdc2_Q3TLP5 | -0.8 | -1.3 | 0.51 | -0.5 | 0.75 | -0.8 | -0.3 | 1.93 | 0.48 | Echdc2 |
| Ranbp2_Q9ERU9 | -1 | 1.52 | 0.47 | 0.52 | -0.5 | -1.8 | 0.64 | -0.2 | 0.41 | Ranbp2 |
| Fam136a_Q9CR98 | -0.1 | 0.68 | -0.3 | -0.1 | 2 | -0.9 | -0.4 | 0.04 | -1.2 | Fam136a |
| Sh3glb1_Q9JK48 | -1.7 | 0.39 | 0.34 | 0.49 | -1.7 | 0.44 | 0.14 | 0.63 | 1 | Sh3glb1 |
| Pttg1ip_Q8R143 | -1.2 | 1.38 | 1.23 | -0 | -1.4 | -0.5 | -0.2 | -0 | 0.82 | Pttg1ip |
| Lmna_P48678 | -1.3 | 0.75 | 0.73 | 0.26 | -2 | 0.28 | 0.27 | 0.38 | 0.72 | Lmna |
| Rab10_P61027 | -0.2 | -0.8 | -0.6 | 0.19 | 0.81 | -1.5 | 1.87 | -0.5 | 0.65 | Rab10 |
| Rbm10_Q99KG3 | -1.6 | -0.6 | -0.3 | 0.06 | -0.5 | 0.62 | 1.43 | 1.4 | -0.5 | Rbm10 |

| | | | | | | | | | | |
|---|---|---|---|---|---|---|---|---|---|---|
| Srp14_P16254 | -0.2 | 1.43 | 0.31 | 1.54 | -0.9 | -0.1 | -0.7 | 0.1 | -1.5 | Srp14 |
| Srsf5_O35326 | -1.3 | 1.09 | 1.13 | -0.6 | -1.5 | 0.4 | -0.3 | 0.87 | 0.29 | Srsf5 |
| Tcof1_O08784 | 1 | -1.5 | 0.11 | 0.05 | 1.02 | 0.48 | 0.24 | -1.8 | 0.4 | Tcof1 |
| Ccdc22_Q9JIG7 | -1 | 0.89 | 0.59 | 0.37 | -2 | 0.16 | -0.4 | 0.46 | 0.99 | Ccdc22 |
| Eif3g_Q9Z1D1 | -1.2 | 0.63 | 0.97 | 0.23 | -2 | 0.3 | -0.1 | 0.36 | 0.89 | Eif3g |
| FAM120A_Q6A0A9 | -1 | -0.1 | 0.62 | 0.17 | 0.55 | 1.41 | 0.58 | -0.2 | -2 | FAM120A |
| Col4a1_P02463 | -1.3 | -0 | 0.87 | 0.66 | -1.7 | -0.2 | 0.23 | -0.1 | 1.48 | Col4a1 |
| Sgta_Q8BJU0 | 2 | -1 | 0.01 | -1.2 | -0.6 | -0.5 | 0.92 | 0.06 | 0.12 | Sgta |
| Hsd17b4_P51660 | 0.09 | 0.17 | -0.9 | -0.2 | 0.96 | -1.8 | 1.69 | -0.1 | 0.13 | Hsd17b4 |
| Pcbp2_Q61990 | 1.96 | -0.1 | -0.1 | -0.3 | 1.27 | -0.6 | -0.2 | -0.7 | -1.2 | Pcbp2 |
| Serpinh1_P19324 | 0.26 | 0.57 | 1.39 | 0.49 | -1.7 | -1.3 | -0.6 | 0.3 | 0.6 | Serpinh1 |
| Adra1b_P97717 | 1.25 | -1.1 | -1.2 | 0.31 | 1.25 | 0.23 | -0.1 | 0.67 | -1.3 | Adra1b |
| Eif2s2_Q99L45 | -1.4 | 0.84 | 0.8 | -0.3 | -1.8 | 0.68 | 0.83 | -0.2 | 0.51 | Eif2s2 |
| Psat1_Q99K85 | 1.52 | -0.9 | -0.5 | -0.1 | 1.88 | -0.4 | -0.2 | -0.5 | -0.8 | Psat1 |
| Ctps2_P70303 | -0.8 | 0.92 | -1.1 | -0.8 | 1.15 | -1.2 | 0.26 | 1.32 | 0.26 | Ctps2 |
| Psmd2_Q8VDM4 | 1.84 | 0.37 | 0.07 | -1.3 | 0.27 | -1.5 | -0.4 | 0.16 | 0.51 | Psmd2 |
| Tmem55a_Q9CZX7 | 0.16 | 0.77 | 0.12 | 0.74 | 0.09 | -2 | 0.11 | 0.22 | 0.36 | Tmem55a |
| Bphl_Q8R164 | 0.85 | 0.22 | 0.04 | -1.3 | 0.25 | -0.9 | 1.38 | 0.86 | -1.4 | Bphl |
| Tsta3_P23591 | 0.06 | 1.24 | -0.3 | -0.2 | -1.2 | -0.6 | 1.81 | -1.2 | 0.26 | Tsta3 |
| Rpl23_P62830 | 1.66 | -0.9 | -0.9 | 0.56 | -0.1 | 1.13 | -0.7 | 0.39 | -1.1 | Rpl23 |
| Hnrnpu_Q8VEK3 | 2 | 0.11 | -0.4 | -0.1 | 0.08 | 0.38 | -0.5 | -1.1 | -0.9 | Hnrnpu |
| Snrnp200_Q6P4T2 | 0.91 | -0.7 | -0.9 | -0.8 | 1.18 | -1.5 | 0.91 | 0.95 | -0.1 | Snrnp200 |
| Vim_P20152 | -0.1 | 0.91 | 0.75 | 0.24 | -0.8 | 1.4 | -1.6 | -1.2 | 0.36 | Vim |
| Itpa_Q9D892 | 0.35 | -0.8 | -0.7 | 0.1 | -2 | 1.16 | 0.66 | 1.1 | 0.03 | Itpa |
| Ltbp3_Q61810 | -1.6 | 0.59 | 0.64 | 0.19 | -1.9 | 0.62 | 0.27 | 0.49 | 0.67 | Ltbp3 |
| Cct6a_P80317 | 1.2 | 0.44 | -1 | -0.1 | 1.81 | -0.6 | -0.2 | -0.3 | -1.2 | Cct6a |
| Ppp1r12a_Q9DBR7 | -0.7 | 0.66 | 0.47 | 0.95 | -1.2 | 1.43 | -1.2 | -0.9 | 0.49 | Ppp1r12a |
| Mfge8_P21956 | -1.2 | -0.7 | 0.5 | 1.25 | 0.62 | 0.15 | -1 | 1.37 | -1 | Mfge8 |
| Col5a2_Q3U962 | -1.1 | 0.52 | 0.5 | 1.42 | -1.1 | 1.1 | -1.3 | -0.2 | 0.2 | Col5a2 |
| Gnb2l1_P68040 | 2 | -0.5 | -0.1 | 0.31 | 0.75 | -0.5 | -0.8 | -0.8 | -0.6 | Gnb2l1 |
| Bub3_Q9WVA3 | 0.07 | -0.2 | 0.78 | -0.8 | -0.5 | 2 | -0.4 | -0.6 | -0.6 | Bub3 |
| Napa_Q9DB05 | 1.4 | 1.31 | -1.1 | -0.3 | -1 | -0.7 | 1.07 | -0.7 | -0 | Napa |
| Dnajb11_Q99KV1 | 1.48 | 0.63 | -1 | -0.8 | 1.52 | -0 | -0.1 | -0.8 | -0.9 | Dnajb11 |
| Asph_Q8BSY0 | -1 | 0.74 | 1.1 | 1 | -1.5 | -1.1 | -0.1 | 0.47 | 0.52 | Asph |
| Ranbp1_P34022 | 1.21 | 1.57 | -0.4 | -0.9 | -0.7 | -1.1 | 1.01 | -0.6 | -0.1 | Ranbp1 |
| Aldh1l2_Q8K009 | 1.96 | -0.4 | 0.1 | -0.8 | 1.15 | 0.03 | -0.1 | -1.2 | -0.8 | Aldh1l2 |
| Fam98b_Q80VD1 | 0.64 | 0.11 | -0.2 | -0.4 | -1.1 | -0 | 1.94 | 0.46 | -1.5 | Fam98b |
| Ap3s1_Q9DCR2 | -0.6 | 1.17 | 0.58 | 0.83 | -2 | -0.6 | 0.31 | 0.58 | -0.2 | Ap3s1 |
| Rab8a_P55258 | 1.85 | 0.55 | -0.7 | -1 | 0.61 | 0.69 | -0.2 | -0.6 | -1.3 | Rab8a |
| Atp5g1_Q9CR84 | -1.7 | 0.67 | 0.59 | 0.37 | -1.8 | 0.37 | 0.05 | 0.53 | 0.85 | Atp5g1 |
| Psmb3_Q9R1P1 | -1.5 | 0.32 | 0.13 | 0.19 | -1.8 | 0.51 | 0.4 | 0.88 | 0.95 | Psmb3 |
| M6pr_P24668 | 0.28 | -0.9 | -0.6 | -1.1 | 1.35 | 1.76 | 0.22 | -0.3 | -0.7 | M6pr |
| Ap3m1_Q9JKC8 | 0.79 | -0.3 | 1.7 | 0.24 | 0.6 | -0.9 | -0.9 | 0.24 | -1.6 | Ap3m1 |
| Psma1_Q9R1P4 | -1.2 | 1.07 | 0.6 | -0.1 | -1.5 | -0.1 | -0.6 | 1.41 | 0.51 | Psma1 |
| Pccb_Q99MN9 | -0.1 | -0.8 | -0.6 | -0.4 | -1.1 | 1.09 | 1.77 | -0.7 | 0.87 | Pccb |
| Prkcdbp_Q91VJ2 | 0.62 | 0.82 | -0.2 | 0.3 | 0.48 | -0.1 | 0.82 | -0.3 | -2 | Prkcdbp |
| Hspa2_P17156 | 1.1 | 1 | -2 | 0.49 | 0.75 | -0 | -0.5 | -0.8 | 0.02 | Hspa2 |
| Exoc4_O35382 | 0.99 | 0.12 | 0.3 | 0.38 | 0.91 | -0.9 | 1.01 | -1.2 | -1.7 | Exoc4 |
| Sucla2_Q9Z2I9 | 1.16 | -1.1 | -0.3 | -1.4 | 0.97 | -0.8 | 0.75 | 1.05 | -0.3 | Sucla2 |
| Hdgfrp2_Q3UMU9 | -1.9 | 0.21 | 0.61 | 0.52 | -1.6 | 0.47 | 0.46 | 0.59 | 0.63 | Hdgfrp2 |

| | | | | | | | | | | |
|---|---|---|---|---|---|---|---|---|---|---|
| Jagn1_Q5XKN4 | -1.7 | 0.45 | 0.29 | 0.69 | -1.8 | 0.38 | 0.42 | 0.58 | 0.69 | Jagn1 |
| Srrm1_Q52KI8 | -0.1 | -1.5 | 0.8 | 1.08 | 1.27 | -0.8 | -1.1 | 0.64 | -0.4 | Srrm1 |
| Cdc5l_Q6A068 | 0.48 | -1.2 | 0.46 | 1.89 | -0.7 | 0.21 | -0.7 | 0.63 | -1 | Cdc5l |
| Shank3_Q4ACU6 | -1.3 | 0.52 | 0.41 | 0.66 | -2 | 0.34 | -0.1 | 0.7 | 0.83 | Shank3 |
| Lnpep_Q8C129 | 0.55 | 2 | 0.08 | -0.6 | -1.3 | -0.5 | -0.9 | 0.64 | -0 | Lnpep |
| Slc7a5_Q9Z127 | -0.2 | 1.14 | -1.3 | -1 | 1.13 | 1.34 | 0.18 | -0.7 | -0.5 | Slc7a5 |
| Cav1_P49817 | 0.66 | 1.3 | -0.1 | -0.9 | -0.1 | 1.42 | 0 | -0.8 | -1.5 | Cav1 |
| Rab11a_P62492 | -1.7 | 0.64 | 0.82 | 0.19 | -1.7 | 0.43 | 0.11 | 0.34 | 0.88 | Rab11a |
| Eif4a2_P10630 | 0.73 | -1.1 | 0.46 | -0.4 | 2 | -0.5 | -0.5 | -0.7 | -0.1 | Eif4a2 |
| Samm50_Q8BGH2 | -1.4 | 0.73 | 0.79 | 0.14 | -0.6 | -0.9 | -0.8 | 0.31 | 1.71 | Samm50 |
| _Q8BH86 | 1.05 | 0.1 | -0.1 | -0.1 | -1.9 | 0.24 | 1.54 | 0.17 | -0.9 | |
| Snrpa1_P57784 | 0.22 | 0.43 | -0.9 | 0.72 | 1.93 | -1.3 | -0.9 | 0.1 | -0.2 | Snrpa1 |
| Eif2a_Q8BJW6 | 0.51 | -0.4 | -0.5 | 0.18 | 1.03 | 0.17 | 0.73 | 0.58 | -2 | Eif2a |
| Mvp_Q9EQK5 | 1.03 | -0.8 | -0.4 | -1.1 | 1.84 | 0.76 | -0.3 | -0.1 | -0.9 | Mvp |
| Slc9a3r2_Q9JHL1 | 0 | 0.8 | -0 | -0.3 | 0.31 | 0.83 | -2 | 0.33 | 0.58 | Slc9a3r2 |
| Dap_Q91XC8 | -1.5 | 0.85 | 0.72 | 0.48 | -1.9 | 0.17 | 0.02 | 0.45 | 0.71 | Dap |
| Rtcb_Q99LF4 | 1.74 | -0.3 | -0.4 | -1.3 | 1.13 | 0.81 | -0.3 | -0.9 | -0.5 | Rtcb |
| Tpi1_P17751 | -1.7 | 0.18 | 0.15 | 0.3 | -1.8 | 0.86 | 0.86 | 0.59 | 0.47 | Tpi1 |
| Inf2_Q0GNC1 | 1.07 | -0.2 | 0.03 | 1.7 | -0.6 | -0.4 | 0.34 | -1.8 | -0.1 | Inf2 |
| H2afv_Q3THW5 | 1.04 | 0.06 | 0.15 | -0.8 | 1.07 | -2 | 0.68 | 0.29 | -0.4 | H2afv |
| Vamp3_P63024 | -1.9 | 0.56 | 0.7 | -0.1 | 0.23 | 1.37 | -0.9 | 0.66 | -0.6 | Vamp3 |
| Cct7_P80313 | 1.8 | -1 | -0.2 | 0.17 | 1.47 | -0.7 | -0.1 | -0.9 | -0.6 | Cct7 |
| Arap1_Q4LDD4 | -0.9 | 1.21 | 1.32 | 1.07 | -0.8 | 0.03 | -1.2 | -0.9 | 0.13 | Arap1 |
| Ddrgk1_Q80WW9 | -1.7 | 0.72 | 0.76 | 0.56 | -1.7 | 0.09 | 0.04 | 0.35 | 0.87 | Ddrgk1 |
| Nsf_P46460 | 1.41 | -0.6 | -0.8 | 0.57 | 1.61 | -1 | -1 | 0.13 | -0.4 | Nsf |
| Acot13_Q9CQR4 | 0.09 | 0.55 | 1.19 | -1.9 | 0.76 | -0.9 | 0.84 | 0.1 | -0.7 | Acot13 |
| Usp39_Q3TIX9 | 0.08 | 0.54 | 0.03 | 0.48 | -2 | 0.65 | 0.2 | 0.15 | 0.47 | Usp39 |
| Fam114a1_Q9D281 | 2 | 0.5 | -0.7 | -0.2 | -1.2 | -0.6 | 0.36 | 0.36 | -0.7 | Fam114a1 |
| Stxbp3_Q60770 | 0.05 | 0.51 | 1.39 | 0.37 | 1.15 | -1.5 | -0.4 | -0.2 | -1.4 | Stxbp3 |
| Copa_Q8CIE6 | 1.6 | 0.38 | -0.3 | -0.7 | 1.63 | -0.7 | -0.4 | -0.5 | -1.1 | Copa |
| Mthfd1_Q922D8 | 1.96 | -0.8 | -0.9 | 1.05 | 0.36 | -0.1 | -0.3 | -1.1 | -0.1 | Mthfd1 |
| Psmb4_P99026 | 0.01 | 0.29 | 0.46 | 0.68 | -0.1 | -2 | -0.2 | 1.51 | -0.5 | Psmb4 |
| Dbnl_Q62418 | -1.7 | 0.34 | 0.77 | 1.28 | -0.9 | 1.16 | -0.2 | -0.8 | 0.04 | Dbnl |
| Cdc37_Q61081 | -1.7 | 1.08 | 0.77 | 0.29 | -1.3 | -0.2 | -0.4 | 0.7 | 0.85 | Cdc37 |
| Tbca_P48428 | 1.19 | -0.7 | -0.5 | -0.5 | 1.1 | 0.63 | 1.08 | -1.4 | -0.9 | Tbca |
| Ptgr2_Q8VDQ1 | -2 | 0.07 | 0.74 | 0.9 | -0.8 | -0.6 | 1.01 | 0.26 | 0.5 | Ptgr2 |
| Naca_Q60817 | -1.3 | 0.25 | 0.58 | 0.05 | -1.9 | 0.87 | -0.3 | 0.65 | 1.04 | Naca |
| Psma7_Q9Z2U0 | 1.44 | -0.7 | -1.3 | -0.6 | 0.93 | -1.3 | 0.21 | 0.5 | 0.78 | Psma7 |
| Psap_Q61207 | -0.4 | 0.21 | -0.1 | -1.8 | 0.82 | -0.6 | 1.87 | 0.12 | -0.1 | Psap |
| Mapk3_Q63844 | 1.73 | 0.21 | 0.64 | -1.8 | -0.6 | 0.17 | -0.7 | -0.3 | 0.62 | Mapk3 |
| Uba2_Q9Z1F9 | 0.23 | 0.39 | -1.7 | 0.59 | 0.65 | 0.39 | 0.52 | 0.72 | -1.8 | Uba2 |
| Rtn3_Q9ES97 | -0.5 | -0.3 | -0.3 | -0.3 | -0.2 | 2 | 0.64 | -1.3 | 0.02 | Rtn3 |
| Rplp2_P99027 | 0.29 | 1.11 | -0.8 | -0.5 | 0.84 | -2 | -0.3 | 0.58 | 0.8 | Rplp2 |
| Csad_Q9DBE0 | -1.1 | 1.15 | 0.3 | 0.79 | -1.6 | -0.5 | -0.7 | 0.72 | 0.93 | Csad |
| Fkbp4_P30416 | 1.5 | -0.1 | 0.25 | 0.6 | 1.08 | -1.2 | -1.3 | -0.9 | 0.17 | Fkbp4 |
| Rbm14_Q8C2Q3 | 0.13 | -0.7 | 1.17 | -0.5 | 1.78 | -0.4 | -0.2 | -1.6 | 0.26 | Rbm14 |
| Slc44a2_Q8BY89 | 1.54 | 0.4 | -0.5 | -1.4 | 1.04 | 0.79 | -0.2 | -0.6 | -1.1 | Slc44a2 |
| Map1lc3b_Q9CQV6 | 1.45 | -0.1 | -0.5 | 0.61 | 0.05 | -2 | 0.17 | -0.1 | 0.61 | Map1lc3b |
| Dazap1_Q9JII5 | 1.12 | 0.19 | -0.9 | -0.6 | 0.39 | -1.2 | 1.76 | 0.12 | -0.8 | Dazap1 |
| Ssr4_Q62186 | -0.2 | -1.2 | -0.3 | 1.74 | -1 | -1 | 0.34 | 0.83 | 0.8 | Ssr4 |

| | | | | | | | | | | |
|---|---|---|---|---|---|---|---|---|---|---|
| Ddah2_Q99LD8 | -1.2 | 0.79 | 0.76 | 1.11 | -1.3 | 1.17 | -0.8 | -0.7 | 0.14 | Ddah2 |
| U2af1_Q9D883 | 0.79 | 1.01 | 0.49 | 0.8 | -1.8 | 0.77 | -0.6 | -1.1 | -0.4 | U2af1 |
| Trpv3_Q8K424 | -2 | -0.2 | 0.65 | 0.15 | 1.38 | 0.37 | 0.5 | -0.2 | -0.4 | Trpv3 |
| Atp6v1g1_Q9CR51 | 1.1 | -0.8 | -0.7 | 0.21 | 1.77 | -0.7 | 0.59 | -0.3 | -1.3 | Atp6v1g1 |
| Abracl_Q4KML4 | -1.4 | -0.7 | 0.91 | 0.85 | -0.8 | -0.6 | 1.71 | 0.3 | -0.3 | Abracl |
| Sec61a1_P61620 | -1.1 | 0.39 | 0.95 | 0.49 | -2 | 0.18 | 0.31 | -0.4 | 1.13 | Sec61a1 |
| Bid_P70444 | 1.2 | 0.34 | 0.22 | 0.58 | 0.89 | -1.4 | 0.51 | -1.6 | -0.8 | Bid |
| Gnl1_P36916 | -1.6 | -0.6 | 0.91 | -0.4 | 0.81 | 1.29 | -0.5 | -0.9 | 0.95 | Gnl1 |
| Btf3l4_Q9CQH7 | -0.5 | -0.4 | 0.77 | -2 | 0.31 | 0.88 | -0.3 | 0.73 | 0.74 | Btf3l4 |
| Rab5b_P61021 | -1.3 | 0.34 | 1.11 | 1.59 | 0.57 | -1.1 | 0.01 | -0.9 | -0.4 | Rab5b |
| Dek_Q7TNV0 | 0.52 | -0.3 | 1.16 | -0.1 | -0.7 | 1.19 | 0.66 | -0.5 | -1.9 | Dek |
| Rpl34_Q9D1R9 | -1.4 | 0.71 | 0.53 | 0.83 | -2 | 0.12 | 0.1 | 0.34 | 0.74 | Rpl34 |
| Rbfox2_Q8BP71 | -0.7 | 1.39 | 0.52 | 0.2 | -0.7 | 0.99 | 0.81 | -1.2 | -1.3 | Rbfox2 |
| Ap3b1_Q9Z1T1 | 0.89 | 0.55 | 0.94 | 1.41 | -0.8 | -0.8 | -1.1 | 0.14 | -1.2 | Ap3b1 |
| Ergic1_Q9DC16 | 0.4 | -1.6 | 0.32 | -0.1 | 1.16 | 0.63 | 0.96 | -1.6 | -0.2 | Ergic1 |
| Fscn1_Q61553 | -1 | 1.34 | 0.86 | 0.84 | -1.2 | 0.84 | -0.9 | -0.9 | 0.12 | Fscn1 |
| Rap1b_Q99JI6 | 0.46 | 2 | 0.79 | -0.4 | -1.3 | -0.8 | 0.23 | -0.7 | -0.3 | Rap1b |
| Ap1g1_P22892 | -0.9 | 1.38 | 1.03 | 0.16 | -1.7 | -0.2 | -0.6 | -0.1 | 0.92 | Ap1g1 |
| Rbm8a_Q9CWZ3 | 1.05 | 0.18 | 0.47 | 0.29 | -0.8 | 0.62 | -2 | 0.77 | -0.4 | Rbm8a |
| Atp6v1a_P50516 | 1.24 | -1.2 | 0.25 | 1.61 | -0.8 | 0.5 | -0.1 | -1.2 | -0.3 | Atp6v1a |
| Arf4_P61750 | 1.2 | -1 | 0.33 | 1.23 | 0.25 | -0.6 | 0.57 | -0.2 | -1.8 | Arf4 |
| Impact_O55091 | -1.8 | 0.26 | 0.98 | 1.25 | -0.7 | 0.85 | -0.4 | 0.35 | -0.7 | Impact |
| Gli3_Q61602 | -0.8 | 0.79 | 1.24 | 1.01 | -1.4 | 0.76 | -1 | -0.9 | 0.19 | Gli3 |
| Rbmxl1_Q91VM5 | -1.5 | 0.4 | 0.76 | 0.49 | -1.9 | 0.43 | -0.1 | 0.46 | 0.93 | Rbmxl1 |
| Ywhaq_P68254 | 1.64 | -0.2 | -0.1 | 0.19 | 1.49 | -1.3 | -1.1 | -0.2 | -0.5 | Ywhaq |
| Efemp2_Q9WVJ9 | -1.2 | -1 | 1.43 | 1.62 | 0.46 | -0.4 | -0.4 | -0.7 | 0.05 | Efemp2 |
| Ywhah_P68510 | 1.83 | -0.1 | -1 | -0.2 | 1.36 | -0.4 | -0.9 | -0.8 | 0.25 | Ywhah |
| Sf1_Q64213 | 0.83 | 1.27 | 0.5 | -0.3 | -0.8 | -1.1 | 0.92 | -1.6 | 0.28 | Sf1 |
| Numa1_E9Q7G0 | -2 | 0.24 | 0.72 | 0.26 | -1.3 | 0.56 | 0.09 | 0.51 | 0.96 | Numa1 |
| Pdxdc1_Q99K01 | 0.51 | -1.4 | -0.5 | 0.52 | 1.71 | -0.5 | 0.41 | 0.52 | -1.2 | Pdxdc1 |
| Smarca5_Q91ZW3 | -0.5 | 0.03 | 1.76 | 0.6 | -0.3 | -2 | 0.5 | 0.12 | -0.2 | Smarca5 |
| Gdap2_Q9DBL2 | 0.11 | 0.39 | 0.52 | 0.46 | 0.63 | -2 | 0.01 | 0.26 | 0.24 | Gdap2 |
| Pdap1_Q3UHX2 | -1.7 | 0.97 | 0.44 | 0.5 | -1.6 | 0.11 | -0.2 | 0.41 | 1.04 | Pdap1 |
| Sec22b_O08547 | -1.8 | 0.63 | 0.64 | 0.48 | -1.6 | 0.31 | -0 | 0.48 | 0.91 | Sec22b |
| S100a11_P50543 | -1.7 | 0.89 | 0.75 | 0.46 | -1.6 | 0.17 | -0.1 | 0.5 | 0.71 | S100a11 |
| Zc3h4_Q6ZPZ3 | -1.6 | 0.86 | 0.75 | 0.32 | -1.8 | 0.42 | 0.14 | 0.12 | 0.78 | Zc3h4 |
| Anxa6_P14824 | 1.82 | -0.2 | -0.5 | -0.2 | 1.53 | -0.8 | -0.1 | -0.5 | -1.1 | Anxa6 |
| Lrpap1_P55302 | -1.5 | 1.2 | 0.55 | -0.3 | -0.4 | -0.7 | 1.08 | 1.08 | -1 | Lrpap1 |
| Hmgn2_P09602 | -1.5 | 0.84 | 0.81 | 0.45 | -1.9 | 0.32 | -0 | 0.51 | 0.49 | Hmgn2 |
| Alcam_Q61490 | -0.2 | 0.68 | 0.27 | 1.08 | -1.6 | 1.19 | -0.9 | 0.56 | -1.1 | Alcam |
| Paxbp1_P58501 | -0.1 | -0.8 | 0.66 | 0.63 | 0.15 | -1.8 | 0.11 | 1.74 | -0.6 | Paxbp1 |
| Rab5c_P35278 | -1.6 | 0.54 | 1.13 | 0.59 | -1.6 | -0.1 | -0.1 | 0.91 | 0.22 | Rab5c |
| Igf2r_Q07113 | -1.7 | 0.49 | -0.4 | 0.74 | -1.6 | 0.59 | 0.12 | 0.5 | 1.14 | Igf2r |
| Fus_P56959 | 1.93 | -0.1 | -1.4 | -0.2 | 0.93 | 0.11 | -0.8 | -0.9 | 0.34 | Fus |
| Lamtor3_O88653 | 0.81 | 0.55 | -1.8 | -0.3 | 0.15 | 1.5 | -0.5 | 0.54 | -0.9 | Lamtor3 |
| Plod1_Q9R0E2 | -2 | 0.55 | 1.04 | 0.5 | -1.2 | -0.3 | 0.5 | 0.83 | 0.04 | Plod1 |
| Rab1b_Q9D1G1 | -0 | -1.5 | 0.64 | -1.7 | 1.19 | -0.2 | 0.88 | 0.54 | 0.11 | Rab1b |
| Cops6_O88545 | -0.1 | 0.55 | 0.04 | -1.4 | 0.26 | -0.2 | 0.97 | -1.5 | 1.5 | Cops6 |
| Rab14_Q91V41 | 0.71 | 1.04 | -1.9 | 0.8 | 0.73 | -0.3 | 0.51 | -0.8 | -0.8 | Rab14 |
| Edc4_Q3UJB9 | -1.4 | 0.63 | -1.5 | 0.52 | 0.55 | 0.9 | 1.09 | 0.14 | -1 | Edc4 |

| | | | | | | | | | | |
|---|---|---|---|---|---|---|---|---|---|---|
| Anp32a_O35381 | -0.7 | 0.98 | 0.91 | 0.5 | -2 | 0.1 | -0.2 | 0.34 | 0.34 | Anp32a |
| Srsf1_Q6PDM2 | -1 | 0.9 | -0.1 | 0.77 | -1.1 | 1.6 | 0.11 | -1.3 | 0.18 | Srsf1 |
| Echs1_Q8BH95 | 2 | -0.9 | 0.47 | -0.6 | -0.4 | -1 | 0.14 | 0.43 | -0.4 | Echs1 |
| Coa6_Q8BGD8 | 0.35 | 0.22 | -0.4 | -0.6 | 1.47 | -2 | -0.1 | 0.94 | 0.23 | Coa6 |
| Glg1_Q61543 | -0.7 | 1.36 | 0.2 | 0.06 | -0.5 | 1.24 | -1.7 | -0.6 | 0.69 | Glg1 |
| Psmd6_Q99JI4 | 1.46 | -0.4 | 1.24 | -2 | -0.4 | 0.29 | -0.2 | -0.1 | 0.03 | Psmd6 |
| Psmc5_P62196 | -0.6 | 0.58 | 0.69 | 0.42 | -2 | 0.64 | 0.24 | 0.22 | 0.3 | Psmc5 |
| Rpl3_P27659 | 1.3 | 0.18 | -1.2 | -0.9 | 0.74 | -1.4 | 0.88 | 0.76 | -0.3 | Rpl3 |
| Gapdh_P16858 | 1.02 | -0.9 | -1.8 | -0.7 | 0.64 | -0.3 | 1.33 | 0.52 | 0.13 | Gapdh |
| Map4_P27546 | -0.6 | 0.8 | 1.31 | 0.48 | -1.2 | 1.05 | -1.3 | -0.8 | 0.31 | Map4 |
| Tnpo3_Q6P2B1 | 0.23 | -1 | -1.6 | 0.89 | 0.44 | -1 | 0.32 | 0.18 | 1.52 | Tnpo3 |
| Pafah1b1_P63005 | 1.48 | 0.66 | -0.9 | -0 | 1.47 | -1.1 | -0.6 | -0.1 | -0.9 | Pafah1b1 |
| Hsp90aa1_P07901 | 1.6 | -0 | -0.5 | 0.11 | 1.25 | -0.8 | 0.53 | -0.6 | -1.5 | Hsp90aa1 |
| Epdr1_Q99M71 | 1.36 | 0.96 | 0.37 | -0.7 | -0.3 | 0.49 | 0.39 | -0.7 | -1.9 | Epdr1 |
| Tmx3_Q8BXZ1 | 1.8 | -0.8 | 1.34 | -0.1 | -0.5 | 0.14 | -0.5 | -1.4 | -0.1 | Tmx3 |
| Cnpy3_Q9DAU1 | -0.9 | -1 | 0.75 | 0.24 | 1.17 | 0.14 | 0.39 | 0.97 | -1.8 | Cnpy3 |
| Fubp1_Q91WJ8 | -0.4 | 0.71 | -0.3 | 0.32 | -2 | 0.85 | -0.5 | 0.98 | 0.55 | Fubp1 |
| Rpsa_P14206 | 1.75 | -0.1 | 0.74 | -1.7 | 0.38 | -0.8 | 0.24 | 0.25 | -0.8 | Rpsa |
| Ndufa12_Q7TMF3 | 0.03 | 0.02 | 0 | 0.11 | 0.05 | 0.82 | 1.25 | 0.11 | -2 | Ndufa12 |
| Gsr_P47791 | 1.15 | 0.09 | 0.38 | -0.8 | 0.68 | -1.7 | 1.36 | -0.8 | -0.4 | Gsr |
| Gmds_Q8K0C9 | 0.21 | 0.32 | 1.07 | 1.14 | -1.6 | 0.71 | 0.12 | -1.4 | -0.6 | Gmds |
| Sept8_Q8CHH9 | -1.5 | 0.63 | 0.6 | 0.39 | -1.9 | 0.62 | -0.1 | 0.34 | 0.88 | Sept8 |
| Vcl_Q64727 | -0.2 | 1.33 | 0.78 | 0.84 | -1.5 | 0.97 | -0.7 | -0.8 | -0.7 | Vcl |
| Eef2_P58252 | 1.56 | 0.38 | -0.5 | 0.07 | 0.14 | -2 | 0.46 | 0.13 | -0 | Eef2 |
| Ubqln2_Q9QZM0 | 0.63 | -1.5 | -0.3 | -0.6 | 1.42 | -1.3 | 0.64 | 0.93 | 0.02 | Ubqln2 |
| Dnpep_Q9Z2W0 | 0.72 | 0.6 | -1.1 | -1.1 | 1.03 | 1.07 | 0.64 | -1.4 | -0.5 | Dnpep |
| Atp2a2_O55143 | 0.62 | 0.25 | -1.7 | -0.7 | 0.65 | 1.77 | -0.2 | 0.06 | -0.7 | Atp2a2 |
| Smarcc2_Q6PDG5 | -2 | 0.91 | 0.81 | 0.09 | 0.06 | 1.08 | -0.6 | -0.4 | 0.25 | Smarcc2 |
| Tln2_Q71LX4 | -1.3 | 0.64 | 0.76 | 0.71 | -2 | 0.23 | -0.2 | 0.33 | 0.8 | Tln2 |
| H2-D1_P01899 | 1 | 0.9 | 1.74 | -0.3 | -1.4 | -0.7 | -0.2 | -0.4 | -0.5 | H2-D1 |
| Akr1b8_P45377 | -0.1 | -1.2 | -0.7 | 0.98 | -0.9 | -0.5 | 0.64 | 1.92 | -0.2 | Akr1b8 |
| Rps27a_P62983 | 1.65 | -0.7 | -0.5 | -0.1 | 1.68 | -0.8 | 0.2 | -0.7 | -0.7 | Rps27a |
| Tpt1_P63028 | -1.8 | 0.76 | 0.68 | 0.63 | -1.6 | 0.16 | -0 | 0.44 | 0.76 | Tpt1 |
| Akap12_Q9WTQ5 | 0.87 | 0.69 | 0.88 | 0.03 | 0.63 | -1.1 | 0.58 | -1.9 | -0.7 | Akap12 |
| Pard3_Q99NH2 | -1.8 | 0.91 | 0.6 | 0.42 | -1.6 | 0.33 | -0.1 | 0.49 | 0.72 | Pard3 |
| Ehbp1_Q69ZW3 | -0.9 | 1.4 | 1.31 | -0.4 | 0.09 | -1.5 | -0.1 | 0.77 | -0.8 | Ehbp1 |
| Eif3e_P60229 | 1.37 | 0.3 | -1 | -1 | 1.82 | -0.1 | -0.4 | -0.3 | -0.6 | Eif3e |
| Rps3a_P97351 | 1.35 | -0.9 | 1.21 | 0.12 | -1.5 | -1 | 0.04 | -0.3 | 0.88 | Rps3a |
| Txndc5_Q91W90 | 2 | -0.7 | 0.52 | -0.2 | 0.65 | -0.6 | -0.8 | -0.9 | -0.2 | Txndc5 |
| Bax_Q07813 | 0.42 | 1.01 | 0.53 | -0.9 | -0.6 | 1.35 | -1.3 | -1.2 | 0.66 | Bax |
| Rhoa_Q9QUI0 | -0.1 | 0.62 | 0.34 | -0.5 | -1.1 | 2 | 0.19 | -0.3 | -1.2 | Rhoa |
| Mcts1_Q9DB27 | 0.39 | 0.16 | 0.38 | 0.6 | 0.62 | -2 | -0 | 0.09 | 0.37 | Mcts1 |
| Rpl37a_P61514 | -0.1 | -0.3 | 0.74 | -0.1 | -2 | 0.87 | 0.94 | -0.8 | 0.78 | Rpl37a |
| Rps7_P62082 | -0.8 | 0.51 | 0.47 | 0.61 | -2 | 0.51 | -0.3 | 0.14 | 1.11 | Rps7 |
| Txndc12_Q9CQU0 | 0.74 | 0.08 | -0.8 | 1.08 | -0.3 | -1.9 | 0.72 | 1.04 | -0.7 | Txndc12 |
| Nans_Q99J77 | 1.71 | -0 | -1 | -0 | 0.17 | 0.52 | -1.9 | 0.15 | 0.38 | Nans |
| A2m_Q61838 | 1.54 | -0.4 | -1 | -0.8 | 1.43 | 0.27 | -1 | -0.6 | 0.54 | A2m |
| Npm1_Q61937 | 2 | 0.76 | 0.62 | -0.1 | -0.3 | -0.4 | -0.3 | -1.3 | -1 | Npm1 |
| Pla2g15_Q8VEB4 | -0.1 | 1.83 | 0.91 | -1.3 | -1 | 0.03 | -0.7 | -0.3 | 0.68 | Pla2g15 |
| Rps15a_P62245 | 0.47 | 1.55 | -0.2 | -0.8 | -1.3 | 0.11 | 0.26 | 1.18 | -1.2 | Rps15a |

| | | | | | | | | | | |
|---|---|---|---|---|---|---|---|---|---|---|
| Ipo5_Q8BKC5 | 0.18 | 0.77 | 1.13 | 0.71 | 0.66 | -1.5 | -0.7 | 0.26 | -1.5 | Ipo5 |
| Pdia5_Q921X9 | 1.18 | -0.1 | 0.44 | 0.33 | 0.79 | -1 | 1 | -1 | -1.6 | Pdia5 |
| Aco2_Q99KI0 | 1.57 | 0.34 | -1.5 | -0.7 | -0.5 | -0.9 | 0.03 | 1.32 | 0.22 | Aco2 |
| Chmp1a_Q921W0 | -1.4 | 0.85 | 0.56 | 0.53 | -2 | 0.49 | -0.1 | 0.41 | 0.66 | Chmp1a |
| Cryz_P47199 | -0.3 | -0.5 | 1.08 | 1.34 | -1.4 | 1.09 | -1.2 | -0.1 | -0 | Cryz |
| Pawr_Q925B0 | -1.7 | 1.14 | 0.5 | 0.68 | -1.4 | -0.2 | -0.4 | 0.4 | 0.96 | Pawr |
| Plcd1_Q8R3B1 | -1.2 | 0.71 | -0.8 | 0.3 | 1.1 | 0.23 | -1.1 | -0.7 | 1.51 | Plcd1 |
| Ap2m1_P84091 | -0.1 | -0.6 | 0.96 | 2 | -0.2 | -0.3 | -1.4 | 0.04 | -0.4 | Ap2m1 |
| Asah1_Q9WV54 | 0.88 | 1.21 | 0.72 | 0.63 | -1.1 | -0.3 | 0.05 | -0.3 | -1.9 | Asah1 |
| Gmfb_Q9CQI3 | 1.6 | 0.13 | -0.3 | -0.3 | 0.17 | 0.73 | 0.38 | -0.3 | -2 | Gmfb |
| Cdh13_Q9WTR5 | 1.32 | 0.93 | -0.8 | -1 | -0.1 | -0.5 | 1.26 | 0.24 | -1.3 | Cdh13 |
| Fip1l1_Q9D824 | 0.75 | -1.4 | -0.2 | -0.4 | 1 | -1.1 | 1.35 | -0.8 | 0.81 | Fip1l1 |
| Gstz1_Q9WVL0 | 0.42 | -0.3 | -1.4 | -0.3 | 1.55 | 0.24 | 0.5 | 0.79 | -1.6 | Gstz1 |
| Smc3_Q9CW03 | 0.15 | 0.38 | 0.13 | 0.59 | -2 | 0.76 | 0.26 | -0 | 0.32 | Smc3 |
| Pofut2_Q8VHI3 | -0.3 | 1.04 | 0.06 | -0.8 | 1.92 | -0.6 | 0.46 | -0.5 | -1.3 | Pofut2 |
| Lxn_P70202 | 1.03 | -0.5 | 0.98 | -1.3 | -1.6 | 1.04 | 0.59 | -0.3 | -0 | Lxn |
| Ilk_O55222 | -0.8 | 0.99 | 0.73 | 0.6 | -2 | 0.29 | -0.2 | 0.38 | 0.29 | Ilk |
| Eif4a1_P60843 | 1.73 | -0.5 | -0.1 | -0.6 | 1.72 | -0.8 | -0.5 | -0.2 | -0.7 | Eif4a1 |
| Trpv2_Q9WTR1 | -0.2 | 0.23 | 0.12 | 1.64 | -1.8 | 0.99 | 0.29 | -0.9 | -0.4 | Trpv2 |
| Rbm7_Q9CQT2 | 0.01 | -1 | -0.2 | -0.6 | 1.51 | -1.4 | 0.06 | 0.02 | 1.56 | Rbm7 |
| Npepps_Q11011 | 1.48 | -0.4 | -0.7 | 1.13 | 0.71 | -0.6 | 0.34 | -1.7 | -0.2 | Npepps |
| Stom_P54116 | 2 | -0.7 | -0 | -0.8 | 0.37 | 0.71 | -0.6 | -1.3 | 0.21 | Stom |
| Uqcrc2_Q9DB77 | 1.88 | -0.3 | -0.3 | -1.3 | 0.86 | -1.2 | 0.57 | 0.14 | -0.4 | Uqcrc2 |
| Tceb1_P83940 | 1.24 | -1.2 | -0.1 | -0.1 | 1.6 | -0.3 | -0.4 | 0.67 | -1.3 | Tceb1 |
| Dstn_Q9R0P5 | 0.74 | 1.24 | -0.3 | 0.46 | 1.26 | -1.6 | -1.2 | -0.2 | -0.5 | Dstn |
| Atp6v1b2_P62814 | 1.19 | -0.6 | -0.1 | -1.6 | 1.27 | -0.9 | -0 | 1.07 | -0.3 | Atp6v1b2 |
| Plekho2_Q8K124 | 1.2 | 0.92 | 1 | 0.45 | -1.3 | -0.1 | -1 | 0.17 | -1.3 | Plekho2 |
| Adpgk_Q8VDL4 | 2 | -0.3 | -0.4 | -1.1 | 0.62 | 0.62 | 0.18 | -0.6 | -1.2 | Adpgk |
| Rpl28_P41105 | -1.5 | 0.74 | 0.79 | 0.55 | -1.8 | 0.25 | 0.43 | 0.81 | -0.3 | Rpl28 |
| Col6a2_Q02788 | -1 | -0.4 | 0.09 | 0.78 | -0.9 | 2 | 0.05 | 0.11 | -0.9 | Col6a2 |
| Gnpda1_O88958 | 0.84 | -0 | 0.02 | 0.68 | 0.8 | -1 | -2 | 0.89 | -0.1 | Gnpda1 |
| Coro1c_Q9WUM4 | -1 | 0.25 | 1.35 | 1.48 | -1.1 | 0.26 | -0.9 | -0.8 | 0.45 | Coro1c |
| Eif3d_O70194 | -1.3 | 0.48 | 0.41 | 0.81 | -1.8 | 0.3 | -0.6 | 0.58 | 1.1 | Eif3d |
| Plp2_Q9R1Q7 | 0.6 | 0.45 | 0.75 | 0.26 | -1.7 | 1.47 | -0.3 | -1.2 | -0.3 | Plp2 |
| Por_P37040 | 1.68 | -1.4 | -0.1 | -1.1 | 0.54 | -0.4 | 0.7 | 0.78 | -0.6 | Por |
| Fkbp3_Q62446 | 1.55 | -0.1 | -0.7 | -0 | 1.42 | -0.8 | -0.4 | 0.51 | -1.4 | Fkbp3 |
| Lemd2_Q6DVA0 | -1.4 | -0.4 | 0.44 | 0.75 | -1.8 | 0.88 | 0.15 | 0.39 | 0.98 | Lemd2 |
| Amacr_O09174 | 1.42 | 0.82 | -0.1 | -1.4 | -0.4 | -0.1 | -1.5 | 0.55 | 0.71 | Amacr |
| Esd_Q9R0P3 | 0.51 | 1.03 | 0.87 | -1.3 | 1.39 | -0.4 | -0.3 | -0.4 | -1.3 | Esd |
| Atp6v1c1_Q9Z1G3 | 1.32 | 0.14 | -0.5 | -1.4 | 1.3 | -1.3 | 0.63 | -0.2 | 0.05 | Atp6v1c1 |
| Khdrbs1_Q60749 | -1.8 | 0.51 | 0.54 | 0.64 | -1.7 | 0.57 | 0.22 | 0.52 | 0.5 | Khdrbs1 |
| Isoc2a_P85094 | -0.8 | -0.4 | -0.2 | -0.5 | -0.2 | 0.55 | 0.54 | 2 | -1.2 | Isoc2a |
| Tmlhe_Q91ZE0 | 1.48 | 0.6 | -0.9 | -0.7 | 1.29 | -0.3 | 0.35 | -0.4 | -1.5 | Tmlhe |
| Erlin2_Q8BFZ9 | -0.8 | 0.08 | -0.6 | 0.14 | 1.3 | -1.8 | 1.36 | -0.2 | 0.46 | Erlin2 |
| Brk1_Q91VR8 | -0.4 | -0 | 0.31 | 1.17 | -2 | 0.38 | 0.17 | 0.69 | 0.07 | Brk1 |
| Amdhd2_Q8JZV7 | -0.2 | 2 | 0.68 | 0.57 | -0.9 | -0.2 | -1.2 | -0.2 | -0.6 | Amdhd2 |
| Luc7l3_Q5SUF2 | -1.7 | 0.74 | 1.08 | -0.1 | -1.5 | 0.66 | -0.1 | 0.03 | 0.83 | Luc7l3 |
| Cct8_P42932 | 1.58 | 1.37 | -0.5 | -0.1 | 0.08 | 0.54 | -0.8 | -0.9 | -1.3 | Cct8 |
| Anxa2_P07356 | 1.96 | -0.7 | -0.4 | -0.6 | 1.51 | -0.3 | -0.3 | -0.6 | -0.5 | Anxa2 |
| Ostm1_Q8BGT0 | 0.42 | -0.1 | 0.24 | -2 | 1.16 | -0.2 | -0.1 | 1.27 | -0.6 | Ostm1 |

| | | | | | | | | | | |
|---|---|---|---|---|---|---|---|---|---|---|
| Hn1_P97825 | 0.29 | -0.2 | -0.2 | -0.1 | 0.2 | -0.9 | 2 | 0.34 | -1.5 | Hn1 |
| Kpna3_O35344 | -0.7 | -0.5 | 0.76 | 0.46 | -1.6 | 0.46 | -0.8 | 0.08 | 1.78 | Kpna3 |
| Anp32b_Q9EST5 | -0.1 | 0.85 | 1.11 | 0.73 | 0.95 | -1.8 | -0.1 | -0.9 | -0.7 | Anp32b |
| Eif4g1_Q6NZJ6 | 0.5 | 1.11 | -0.8 | -1 | -1.3 | 0.93 | 0.23 | 1.23 | -0.9 | Eif4g1 |
| Snrpd2_P62317 | -2 | 1.27 | 0.54 | 0.09 | -0 | -0.8 | 0.7 | 0.67 | -0.3 | Snrpd2 |
| Actr1a_P61164 | -1.6 | 1 | 0.61 | 0.66 | -1.5 | -0.1 | -0.5 | 0.39 | 1 | Actr1a |
| Tomm40_Q9QYA2 | -1.7 | 1.71 | 0.63 | 0.69 | -0.4 | 0.32 | -0.9 | -0 | -0.2 | Tomm40 |
| Rnf5_O35445 | 0.4 | -1.8 | 0.44 | 0.62 | -1.7 | 0.74 | 0.28 | 0.49 | 0.52 | Rnf5 |
| Cpne1_Q8C166 | 1.02 | 1.18 | 0.36 | -1.7 | -0.5 | 0.19 | 0.87 | -0.2 | -1.2 | Cpne1 |
| Eif2s1_Q6ZWX6 | 1.72 | 0.4 | -0.5 | -1.5 | 0.87 | -0.9 | 0.59 | -0.5 | -0.1 | Eif2s1 |
| Ckap5_A2AGT5 | 1.1 | -1 | -0.5 | -1.2 | 1.57 | 0.76 | 0.15 | 0.15 | -0.9 | Ckap5 |
| Gsta4_P24472 | 1.63 | -1 | 0.21 | -0.2 | 1.26 | -0.5 | 0.56 | -0.6 | -1.3 | Gsta4 |
| Apoo_Q9DCZ4 | -0.4 | 1.52 | 0.66 | -0.4 | -1.5 | 0.22 | 1.22 | -0.4 | -0.9 | Apoo |
| Hp1bp3_Q3TEA8 | 1.79 | 0.48 | 0.97 | -1.2 | 0.53 | -0.3 | -0.9 | -0.9 | -0.5 | Hp1bp3 |
| Calu_O35887 | -0.1 | 1.52 | 1.31 | -0.6 | -0.1 | -1.7 | 0.52 | -0.6 | -0.2 | Calu |
| Ckap4_Q8BMK4 | -0.4 | 1.25 | 1.55 | 0.54 | -1.2 | 0.22 | -1.2 | -0.9 | 0.05 | Ckap4 |
| Impa1_O55023 | 0.79 | 1.9 | -1.4 | -0.2 | -1 | -0.3 | -0.5 | 0.19 | 0.52 | Impa1 |
| Eif6_O55135 | 1.31 | -1.8 | -0.7 | -0.5 | 1.13 | 0.77 | 0.39 | -0.7 | 0.03 | Eif6 |
| Pgrmc2_Q80UU9 | 0.9 | -1.3 | -1.6 | -0.8 | 0.86 | 0.23 | 0.18 | 1.06 | 0.5 | Pgrmc2 |
| Siae_P70665 | 0.7 | -0.1 | 0.4 | 0.56 | 0.62 | -2 | 0.18 | 0.53 | -0.4 | Siae |
| Gart_Q64737 | 1.42 | -0.6 | -1.8 | 0.78 | -0.6 | 1.11 | -0 | -0.3 | -0 | Gart |
| Eef1d_P57776 | 1.22 | 0.74 | -0.8 | -1.9 | 0.96 | -0.6 | 0.56 | -0.3 | 0.18 | Eef1d |
| Kpnb1_P70168 | -1.3 | 1.29 | 0.76 | 0.49 | -1.4 | -0.4 | -0.6 | 0.06 | 1.12 | Kpnb1 |
| Apex1_P28352 | -0.9 | 0.49 | 0.99 | 1.37 | -1.8 | -0.7 | 0.46 | 0.29 | -0.2 | Apex1 |
| Dpp7_Q9ET22 | 1.55 | 0.3 | -0.1 | 0.68 | 0.83 | -1.5 | 0.27 | -1.1 | -0.9 | Dpp7 |
| Spr_Q64105 | 1.78 | -0.8 | 0.49 | -1.2 | 0.17 | 1.19 | -0.3 | -0.8 | -0.5 | Spr |
| Mp68_P56379 | 0.03 | -1 | -0.3 | -1.3 | 1 | 0.19 | 1.86 | 0.28 | -0.7 | Mp68 |
| Cltc_Q68FD5 | -1.4 | 1.04 | 0.71 | 0.5 | -2 | 0.12 | 0.39 | 0.12 | 0.47 | Cltc |
| Snx12_O70493 | -1.5 | -0.2 | 0.73 | 1.09 | -1.8 | 0.32 | 0.13 | 0.46 | 0.71 | Snx12 |
| Yipf5_Q9EQQ2 | 0.41 | 0.47 | -1.8 | 0.82 | -1.7 | 0.51 | 0.1 | 0.5 | 0.64 | Yipf5 |
| Blvra_Q9CY64 | 0.79 | -1.1 | -0.5 | -1 | 1.22 | 1.32 | 0.06 | 0.45 | -1.2 | Blvra |
| Larp1_Q6ZQ58 | -1.8 | 0.26 | 0.09 | 1.35 | -0.2 | -1.2 | -0.1 | 0.75 | 0.91 | Larp1 |
| Itpr3_P70227 | -1.3 | 1.59 | 1.06 | -0.3 | -0.5 | 0.9 | -0.5 | -1.1 | 0.14 | Itpr3 |
| Pdia3_P27773 | 2 | 0.11 | -0.5 | -0.8 | 1.2 | -0.3 | -0 | -0.8 | -1 | Pdia3 |
| Hspe1_Q64433 | 1.76 | -1.1 | -0.6 | -1.3 | 0.71 | -0.4 | 0.82 | 0.47 | -0.4 | Hspe1 |
| Dkc1_Q9ESX5 | -0.8 | 0.62 | 1.65 | 0.56 | -0.4 | -1.7 | 0.53 | 0.29 | -0.8 | Dkc1 |
| Tpp2_Q64514 | 1.14 | 0.83 | -1.2 | 0.46 | 1.11 | -1.1 | 0.28 | -1.3 | -0.2 | Tpp2 |
| Glrx_Q9QUH0 | 1.81 | 0.37 | -0.6 | -1.4 | 0.77 | 0.67 | -0.1 | -0.8 | -0.8 | Glrx |
| Tpr_F6ZDS4 | -0.4 | 0.19 | -0 | 0.58 | -1 | 1.32 | -2 | 0.54 | 0.74 | Tpr |
| Pdcd5_P56812 | -1.8 | 0.86 | 0.67 | 0.65 | -1.5 | 0.17 | -0.2 | 0.52 | 0.68 | Pdcd5 |
| Clpp_O88696 | 1.63 | -0.1 | -0.9 | -0.4 | 1.62 | -0.7 | -1.2 | -0.1 | 0.04 | Clpp |
| Vps26a_P40336 | 1.55 | 0.65 | -0.7 | -1 | 0.25 | 0.9 | -1.1 | -1 | 0.6 | Vps26a |
| Lap3_Q9CPY7 | 1.76 | -1.3 | -0.5 | -0.6 | 0.93 | -1.2 | 0.67 | 0.06 | 0.13 | Lap3 |
| Dlg1_Q811D0 | -1.4 | 0.62 | 0.81 | -1.3 | -0.8 | 1.3 | 0.87 | -0.1 | 0.07 | Dlg1 |
| Rcn2_Q8BP92 | 0.27 | -0.5 | 1.18 | -1.6 | 1.28 | 0.61 | 0.24 | -1.3 | -0.2 | Rcn2 |
| S100a10_P08207 | -0.8 | 1.53 | -0.3 | 0.23 | -0.3 | 0.82 | -0.6 | -1.6 | 1.04 | S100a10 |
| Myo1b_P46735 | 0.02 | -0.8 | 1.6 | 0.61 | -1.4 | -0.5 | -1.1 | 0.91 | 0.56 | Myo1b |
| Rpl9_P51410 | 1.88 | 0 | -1 | -0.7 | -0.1 | -0.6 | 1.45 | -0.6 | -0.4 | Rpl9 |
| Anxa11_P97384 | -0.7 | 0.84 | 1.21 | -0.8 | 1.62 | -0.7 | -1.3 | -0.3 | 0.01 | Anxa11 |
| Alyref_O08583 | -1 | -0.5 | 1.28 | -0.3 | -0.3 | 1.64 | -1.2 | -0.4 | 0.81 | Alyref |

| | | | | | | | | | | |
|---|---|---|---|---|---|---|---|---|---|---|
| Marcksl1_P28667 | 0.78 | 0.78 | 1.24 | 0.44 | -0.1 | -1.1 | 0.08 | -1.9 | -0.1 | Marcksl1 |
| Rpl12_P35979 | -1.8 | -0.4 | 1.17 | 1.4 | -0.6 | 0.54 | -0.8 | 0.18 | 0.25 | Rpl12 |
| Ano6_Q6P9J9 | -0.8 | 1.99 | 0.34 | -0.7 | -0.6 | -0.5 | 1.19 | -0.9 | -0 | Ano6 |
| Rps26_P62855 | -1.5 | 0.45 | 1.08 | 1.22 | -1.1 | 0.69 | 0.47 | -0.9 | -0.4 | Rps26 |
| Blvrb_Q923D2 | 0.08 | 0.61 | -0.1 | 0.16 | 1.9 | -1.5 | 0.47 | -0.6 | -1 | Blvrb |
| Nop58_Q6DFW4 | -0.1 | 1.32 | 0.64 | -0.7 | -0.1 | -0.9 | 0.38 | -1.7 | 1.18 | Nop58 |
| Mtch1_Q791T5 | -1.9 | 0.65 | 0.73 | 0.6 | -1.4 | 0.32 | -0.2 | 0.47 | 0.8 | Mtch1 |
| Ubxn1_Q922Y1 | 1.13 | 0.22 | -2 | -0.2 | 1.48 | -0.1 | -0.1 | 0.15 | -0.6 | Ubxn1 |
| Tubb2b_Q9CWF2 | -0.1 | -1.4 | 1.66 | -0.2 | 0.75 | -1.4 | 0.27 | -0.2 | 0.71 | Tubb2b |
| Ube2i_P63280 | 1.59 | -0.2 | -0.6 | -1.2 | 1.25 | 0.33 | 0.61 | -0.9 | -0.9 | Ube2i |
| Pitrm1_Q8K411 | 0.55 | 1.73 | 0.41 | -2 | -0.3 | 0.29 | -0.3 | -0.2 | -0.1 | Pitrm1 |
| Cops7a_Q9CZ04 | 0.96 | 0.14 | -1.7 | -1 | 0.75 | 1.32 | 0.2 | -0.9 | 0.2 | Cops7a |
| Srp54_P14576 | -1.7 | 0.65 | 0.74 | 0.64 | -1.7 | 0.4 | -0.2 | 0.18 | 0.92 | Srp54 |
| Atf7_Q8R0S1 | -0.9 | 0.2 | 0.69 | 0.62 | -2 | 0.87 | -0.1 | 0.28 | 0.58 | Atf7 |
| Dnm1_P39053 | -1.7 | 0.79 | 1.02 | 1.35 | -0.6 | 0.09 | -0.8 | -0.6 | 0.47 | Dnm1 |
| Igf2bp1_O88477 | 0.5 | 0.78 | 0.55 | -1.1 | 1.2 | -0.3 | -1.9 | -0.3 | 0.57 | Igf2bp1 |
| Clta_O08585 | -0.9 | 0.89 | 0.69 | 0.43 | -2 | 0.55 | -0 | -0.4 | 0.91 | Clta |
| Gnb2_P62880 | -1.4 | 0.5 | -0.8 | 1.71 | 0.67 | -0.6 | -1.1 | 0.37 | 0.53 | Gnb2 |
| Nradd_Q8CJ26 | -1.8 | 1.1 | -0.6 | 0.57 | -1.3 | 0.6 | 0.45 | 0.05 | 0.89 | Nradd |
| Rdx_P26043 | 0.55 | 0.67 | -1.4 | -0.8 | 0.05 | 1.94 | -0.7 | -0.4 | 0.17 | Rdx |
| Ube2v1_Q9CZY3 | -1.5 | 0.71 | 0.93 | 0.48 | -0.5 | 0.7 | -1.7 | 0.83 | -0 | Ube2v1 |
| Ddah1_Q9CWS0 | -1.9 | 1.25 | 0.78 | 0.84 | -0.7 | 0.57 | -0.7 | -0.4 | 0.28 | Ddah1 |
| Tmem263_Q9DAM7 | -0.5 | 1.5 | -1.5 | -0.7 | 0.73 | -0.5 | 0.04 | 1.3 | -0.3 | Tmem263 |
| Actr2_P61161 | 1.1 | -1 | -0.9 | -0.1 | 1.33 | 0.01 | 1.27 | -0.5 | -1.1 | Actr2 |
| Aoc3_O70423 | 0.76 | 0.34 | 0.44 | 1.18 | -2 | -0.4 | 0.47 | -0.3 | -0.2 | Aoc3 |
| F3_P20352 | -0.1 | 0.61 | 0.7 | 0.98 | -1.6 | -0.7 | 1.15 | -1.3 | 0.27 | F3 |
| Mpst_Q99J99 | 1.53 | 0.53 | -0.9 | -0.1 | 1.41 | -1.1 | 0.01 | -0.2 | -1.1 | Mpst |
| Sap18_O55128 | -0.3 | 0.45 | -0.1 | 0.6 | -2 | 0.77 | 0.95 | 0.66 | -0.9 | Sap18 |
| Rpl5_P47962 | -0.1 | 2 | -0 | -1.4 | -0.9 | 0.16 | 0.45 | -0.2 | -0.1 | Rpl5 |
| Adsl_P54822 | 1.26 | 0.92 | -0.5 | -0.9 | 0.48 | 0.36 | -0.3 | 0.57 | -1.9 | Adsl |
| Gnas_P63094 | 1.55 | -1 | -1.2 | -0.3 | -0.1 | 1.64 | -0.5 | -0.3 | 0.23 | Gnas |
| Rps11_P62281 | 1.52 | -0.2 | -1.2 | -0.1 | 1.52 | -0.6 | -0.1 | 0.38 | -1.1 | Rps11 |
| Rps2_P25444 | 1.54 | 0.04 | -0.2 | 0.35 | 1.26 | -1.1 | -0.4 | 0.08 | -1.6 | Rps2 |
| Sltm_Q8CH25 | -1.7 | 0.57 | 0.86 | 0.42 | -1.7 | 0.71 | 0.21 | 0.17 | 0.5 | Sltm |
| Rps27_Q6ZWU9 | 1.79 | -0.4 | -1 | -0.5 | 1.19 | -0.1 | -0.5 | 0.64 | -1.2 | Rps27 |
| Parva_Q9EPC1 | 0.69 | 0.86 | 0.22 | -1 | -1.1 | 1.9 | -0.3 | -0.5 | -0.7 | Parva |
| Rab1A_P62821 | 0.73 | 0.34 | 1.34 | -1.2 | 0.88 | -1.7 | -0.1 | 0.31 | -0.6 | Rab1A |
| Eif2s3x_Q9Z0N1 | 1.09 | -0.8 | -1 | 0.15 | 1.11 | -1.8 | 0.85 | 0.4 | -0.1 | Eif2s3x |
| Lnp_Q7TQ95 | -0.2 | 1.28 | 0.57 | 0.19 | 0.73 | -1 | 0.99 | -1 | -1.6 | Lnp |
| Map2k1_P31938 | 0.1 | 0.1 | 0.79 | -1 | 1.89 | -0.7 | -1.5 | 0.2 | 0.09 | Map2k1 |
| U2af2_P26369 | -1.5 | 0.57 | 0.54 | -0 | -1.7 | 1.21 | 0.43 | 0.71 | -0.2 | U2af2 |
| Rpl36a_P83882 | -1.7 | 1.19 | 0.34 | 0.66 | -1.5 | 0.21 | -0.4 | 0.61 | 0.58 | Rpl36a |
| Capn1_O35350 | 0.48 | 0.69 | 1.1 | 0.01 | -2 | 0.3 | -0 | -1.1 | 0.61 | Capn1 |
| Ufd1l_P70362 | -1.7 | -0.2 | -0.3 | 1.03 | 0.77 | -1.3 | 1.26 | 0.32 | 0.09 | Ufd1l |
| Rac1_P63001 | 1.87 | 0.16 | 0.29 | -1.4 | 0.96 | -0.1 | -0.1 | -0.9 | -0.9 | Rac1 |
| Maoa_Q64133 | 1.49 | -1.2 | -0.7 | -0.9 | 0.88 | 0.97 | 0.37 | -1.1 | 0.17 | Maoa |
| Rpl31_P62900 | 1.64 | -0 | -0.8 | -1 | 0.6 | 0.92 | 0.29 | -0.1 | -1.5 | Rpl31 |
| Lepre1_Q3V1T4 | -0.7 | 1.1 | 0.92 | 0.92 | -1.9 | -0.3 | -0.5 | -0.1 | 0.64 | Lepre1 |
| Naga_Q9QWR8 | 1.7 | -0.5 | 0.72 | 0.16 | 1.04 | -1.5 | -0.7 | -0.6 | -0.4 | Naga |
| Nedd4l_Q8CFI0 | 0.54 | 0.34 | 0.51 | -0.2 | -0.6 | 0.8 | 0.07 | -2 | 0.9 | Nedd4l |

| | | | | | | | | | | |
|---|---|---|---|---|---|---|---|---|---|---|
| Rps9_Q6ZWN5 | -1.3 | 0.85 | 0.58 | 0.92 | -1.8 | 0.1 | -0.6 | 0.55 | 0.69 | Rps9 |
| Uchl5_Q9WUP7 | 0.42 | -0.4 | -0.3 | -0.4 | 0.73 | -0.9 | 0.65 | 1.78 | -1.6 | Uchl5 |
| Sf3a1_Q8K4Z5 | 2 | 0.05 | -1.2 | -0.3 | 0.85 | -0.4 | -0.6 | -0.4 | -0.2 | Sf3a1 |
| Qki_Q9QYS9 | 0.92 | 0.96 | -0.2 | -1.5 | -0.6 | 0.63 | 1.31 | -1.1 | -0.3 | Qki |
| _Q6PIU9 | 0.05 | 0.81 | -1 | -0.1 | -1.3 | 0.76 | -1.3 | 1.39 | 0.68 | |
| Cct5_P80316 | 1.95 | -0.6 | -0.4 | -0.5 | 1.2 | -0.4 | 0.36 | -0.3 | -1.3 | Cct5 |
| Uba1_Q02053 | 1.49 | 0.46 | -0.1 | 0.63 | 0.63 | -0.8 | -1.8 | -0.8 | 0.35 | Uba1 |
| Pfkp_Q9WUA3 | -1.1 | 0.54 | -2 | -0.3 | 0.6 | 0.41 | 1.09 | -0 | 0.81 | Pfkp |
| Pcyox1_Q9CQF9 | 1.46 | -0.9 | 0.95 | 0.39 | 0.82 | -1.2 | 0.28 | -0.6 | -1.1 | Pcyox1 |
| Snw1_Q9CSN1 | 1.12 | 0.7 | 0.87 | -1.3 | 0.24 | 0.28 | -1.9 | 0.14 | -0.1 | Snw1 |
| Slc25a24_Q8BMD8 | -0.9 | 0.5 | 1.8 | 0.68 | -1.3 | 0.67 | -1 | -0.2 | -0.2 | Slc25a24 |
| Nfkb2_Q9WTK5 | -1.8 | 0.78 | 1.09 | 0.52 | -1.4 | 0.18 | -0 | -0 | 0.73 | Nfkb2 |
| Metap2_O08663 | -0.3 | 0.55 | 0.35 | 0.34 | 0.45 | -0.4 | 0.67 | -2 | 0.79 | Metap2 |
| Rsu1_Q01730 | -1.4 | 1.38 | -0.1 | 1.42 | 0.28 | -0.9 | 0.06 | -1 | 0.29 | Rsu1 |
| Hsp90ab1_P11499 | 1.71 | -0.4 | -0.6 | -1 | 1.16 | -1.2 | 0.57 | 0.31 | -0.5 | Hsp90ab1 |
| Lonp1_Q8CGK3 | 1.2 | -1.1 | -1.2 | -0.5 | 0.34 | 1.35 | 1 | -0.3 | -0.9 | Lonp1 |
| Arpc3_Q9JM76 | -0.7 | 0.77 | 0.48 | 0.17 | -2 | 1.12 | -0 | 0.01 | 0.42 | Arpc3 |
| Svil_Q8K4L3 | 1.63 | -0.3 | 0.08 | -1.5 | 1.54 | -0.1 | -0.7 | -0.3 | -0.4 | Svil |
| Gpc1_Q9QZF2 | 1.68 | 0.03 | -0.7 | -0.4 | 1.55 | -1 | 0.1 | -0.2 | -1 | Gpc1 |
| Hsd17b10_O08756 | 0.06 | -1.2 | 0.93 | -1.2 | 0.65 | -0.3 | 1.81 | -0.1 | -0.6 | Hsd17b10 |
| Ptma_P26350 | -1.5 | 1.22 | 0.92 | 1.08 | -1.3 | 0.55 | -0.5 | -0.4 | -0.1 | Ptma |
| Scfd1_Q8BRF7 | -1.8 | 0.56 | 0.67 | 0.83 | -1.7 | 0.42 | 0.39 | -0.1 | 0.6 | Scfd1 |
| Acp2_P24638 | -1.2 | -0.5 | -1.5 | -0.5 | 0.28 | 1.35 | 0.64 | 1.22 | 0.21 | Acp2 |
| Nucks1_Q80XU3 | -1.1 | 1.67 | -0 | 0.17 | 0.27 | -1.4 | -0.8 | 0.1 | 1.13 | Nucks1 |
| Ctnnb1_Q02248 | -0.1 | 0.84 | 0.51 | 1 | -1.9 | 0.94 | -0.8 | -0.9 | 0.39 | Ctnnb1 |
| Hnrnpul2_Q00PI9 | 1.25 | -0.2 | -1 | 0.33 | 0.75 | -1.8 | 1.01 | 0.23 | -0.5 | Hnrnpul2 |
| Gnb3_Q61011 | -1.7 | 0.74 | 0.67 | 0.51 | -1.7 | 0.7 | 0.24 | 0.03 | 0.54 | Gnb3 |
| Map1s_Q8C052 | -1.8 | 1.18 | -0.3 | 0.87 | -0.4 | -0.6 | -0.4 | 1.31 | 0.22 | Map1s |
| Hdac1_O09106 | 0.75 | 0.82 | 0.16 | 0.14 | 0.49 | -2 | 1.07 | -0.5 | -0.8 | Hdac1 |
| Rcn1_Q05186 | 0.74 | 1.88 | -0.1 | -0.2 | 0.01 | -1.9 | -0.2 | -0.4 | 0.25 | Rcn1 |
| Atad1_Q9D5T0 | 0.84 | -1.5 | 1.17 | -1.5 | 1.09 | -0.5 | 0.14 | 0.35 | -0.1 | Atad1 |
| Rbbp4_Q60972 | -2 | 0.68 | 1.71 | -0.5 | -0 | -0.3 | -0.3 | 0.4 | 0.37 | Rbbp4 |
| Lamtor5_Q9D1L9 | 1.4 | -1.2 | 0.14 | -1.1 | 1.24 | -1 | 0.5 | 0.58 | -0.6 | Lamtor5 |
| Gpr124_Q91ZV8 | -2 | 0.57 | 0.51 | 0.47 | -1.2 | 0.66 | 0.07 | 0.29 | 0.78 | Gpr124 |
| Hexa_P29416 | 1.98 | -0.3 | -0.6 | -0.4 | 1.4 | -0.9 | -0.3 | -0.8 | -0.2 | Hexa |
| Aldh1a7_O35945 | -1.3 | -0.1 | 1.24 | -0.8 | 0.73 | -0.5 | 0.8 | 1.13 | -1.2 | Aldh1a7 |
| Cct4_P80315 | 1.31 | -0.2 | -0.4 | -0.2 | 1.88 | -1.3 | 0.11 | -0.3 | -0.8 | Cct4 |
| Tubb4b_P68372 | 1.76 | -0.8 | -0.3 | -0.7 | 1.54 | -0.4 | 0.25 | -0.4 | -0.9 | Tubb4b |
| Ssb_P32067 | -1.9 | 1.01 | 0.59 | 0.67 | -1.4 | 0.22 | -0.2 | 0.42 | 0.59 | Ssb |
| Hint2_Q9D0S9 | 2 | -0.8 | -1.1 | 0.09 | 0.22 | 0.19 | -0.1 | 0.46 | -1.1 | Hint2 |
| Dynlrb1_P62627 | 0.53 | 0.94 | 0.71 | -1 | 0.29 | 0.13 | 1.06 | -0.8 | -1.9 | Dynlrb1 |
| Tmed2_Q9R0Q3 | 0.96 | 0.62 | -0.3 | 1.66 | -1 | -0.7 | -1.1 | 0.69 | -0.9 | Tmed2 |
| Gulp1_Q8K2A1 | 1.23 | -0.4 | -1.5 | -0.2 | -1 | 0.78 | -0.9 | 0.84 | 1.07 | Gulp1 |
| Rps28_P62858 | -1.7 | 0.16 | 1.45 | 1.04 | -1.1 | 0.73 | -0.1 | 0.01 | -0.5 | Rps28 |
| Cdv3_Q4VAA2 | 1.67 | -1.2 | 0.14 | -0.7 | 1.41 | -0.3 | 0.39 | -0.7 | -0.6 | Cdv3 |
| Col4a2_P08122 | -0.1 | -0.7 | 1.37 | 0.54 | -1.1 | 1.17 | -0.6 | 0.75 | -1.3 | Col4a2 |
| Atp2b1_G5E829 | -1.1 | 1.76 | 0.62 | -0.2 | -0.5 | 0.63 | 0.33 | 0.03 | -1.6 | Atp2b1 |
| Sars_P26638 | 1.09 | -1.2 | -0.4 | -1.9 | 0.73 | 0.76 | 0.34 | -0.1 | 0.64 | Sars |
| Stub1_Q9WUD1 | -1 | 1.13 | 0.73 | 1.59 | -1.2 | -0.2 | -0.4 | -1 | 0.31 | Stub1 |
| Plec_Q9QXS1 | -1 | 0.8 | 0.99 | 0.56 | -1.2 | 1.03 | -1.5 | -0.2 | 0.51 | Plec |

| | | | | | | | | | | |
|---|---|---|---|---|---|---|---|---|---|---|
| Hist2h2ac_Q64523 | 2 | 0.14 | 0.54 | -0.3 | 0.43 | -0.9 | 0.02 | -0.6 | -1.5 | Hist2h2ac |
| Xpo1_Q6P5F9 | 1.4 | 0.64 | 0.31 | 0.09 | 0.47 | -2 | -0.4 | -0.2 | -0 | Xpo1 |
| Stt3a_P46978 | -0.6 | 0.72 | 1.4 | 0.23 | -1.8 | 0.16 | -0.8 | -0.4 | 1.06 | Stt3a |
| Pdgfrb_P05622 | -0.3 | 0.42 | 0.95 | 1.3 | -2 | -0.5 | -0.7 | 0.37 | 0.47 | Pdgfrb |
| Sppl2a_Q9JJF9 | -2 | 0.3 | 0.55 | 0.9 | -1.2 | 0.37 | 0.06 | 0.54 | 0.62 | Sppl2a |
| Cmpk1_Q9DBP5 | -0.3 | -1.3 | -0.1 | -0.7 | 1.62 | -0.9 | 0.62 | 1.33 | -0.2 | Cmpk1 |
| Sgcd_P82347 | -1.4 | 0.83 | 0.87 | 0.54 | -1.9 | 0.61 | 0.08 | -0.1 | 0.48 | Sgcd |
| Ak1_Q9R0Y5 | 1.29 | -0.1 | -0 | 1.01 | 0.65 | -1.6 | -0.7 | -1.2 | 0.64 | Ak1 |
| Hmgn1_P18608 | -1.7 | 0.76 | 0.69 | 1.85 | -0.7 | -0.4 | -0.1 | -0.3 | -0.2 | Hmgn1 |
| Ndufa9_Q9DC69 | 1.21 | -0.6 | 0.42 | 1.07 | -1.6 | 0.72 | -0.6 | 0.46 | -1.1 | Ndufa9 |
| Fhl2_O70433 | -1.5 | 0.81 | -0.2 | -1.2 | 0.73 | 0.35 | 1.66 | -0.2 | -0.4 | Fhl2 |
| Dhx15_O35286 | 1.6 | 0.28 | -0.3 | -1.4 | 1.28 | -1.3 | -0.1 | -0.1 | 0.03 | Dhx15 |
| Fah_P35505 | 1.53 | 0.94 | -1 | -0.7 | -1.2 | 0.54 | 0.62 | 0.3 | -1 | Fah |
| Csnk2a2_O54833 | -2 | -0.4 | 0.77 | 0.94 | -0.3 | -0.3 | -0.3 | 0.58 | 1.15 | Csnk2a2 |
| Snrpd3_P62320 | 0.79 | -0.9 | -0.6 | -1.5 | 0.82 | 1.66 | -0 | 0.34 | -0.6 | Snrpd3 |
| Sfpq_Q8VIJ6 | 1.53 | 1.45 | -0.9 | 0.26 | -0 | -0.4 | -1.5 | 0.03 | -0.4 | Sfpq |
| Myg1_Q9JK81 | -1.1 | -0.1 | 0.65 | 1.37 | -1.6 | 1.09 | -0.8 | 0.45 | 0.03 | Myg1 |
| Sh3bgrl_Q9JJU8 | 1.04 | -1.4 | -0.2 | -0.1 | 1.96 | -0.9 | 0.13 | -0.5 | -0.1 | Sh3bgrl |
| Dctn1_O08788 | -2 | 1.06 | 0.4 | 0.64 | -1.1 | 0.17 | 0.52 | -0.1 | 0.49 | Dctn1 |
| Ncstn_P57716 | 0.22 | 0.32 | -0.9 | 1.12 | 0.13 | -0.4 | -1.8 | -0.1 | 1.5 | Ncstn |
| Eif3c_Q8R1B4 | 0.59 | 0.15 | -0.6 | -0.3 | 1.35 | -1.8 | 0.54 | 1 | -0.8 | Eif3c |
| Ndufs1_Q91VD9 | -1.1 | 1.16 | 0.62 | -1.5 | -0.6 | 1.19 | 0.63 | 0.32 | -0.6 | Ndufs1 |
| Ktn1_Q61595 | -0.8 | 0.45 | 0.33 | -1.6 | -0.8 | 1.82 | 0.06 | -0.2 | 0.74 | Ktn1 |
| Hmox1_P14901 | -1.2 | 0.81 | 1.03 | 0.82 | 0.04 | -1.8 | -0.2 | 0.84 | -0.3 | Hmox1 |
| Hnrnpc_Q9Z204 | -1.1 | 0.7 | 0.74 | 0.77 | -1.6 | 0.07 | -1.1 | 0.67 | 0.89 | Hnrnpc |
| Pfkl_P12382 | -1.5 | 0.85 | 0.66 | 0.56 | -1.3 | 1.36 | -0.7 | -0.4 | 0.4 | Pfkl |
| Acin1_Q9JIX8 | -1.9 | 0.57 | 0.89 | 0.27 | -1.4 | 0.76 | 0.35 | -0.2 | 0.67 | Acin1 |
| Ctnnd1_P30999 | -1.4 | 0.95 | 1.22 | 0.77 | -0.8 | 0.25 | -1.4 | -0.2 | 0.58 | Ctnnd1 |
| Pwp2_Q8BU03 | 1.36 | -0.5 | -0.9 | 0.31 | 0.98 | -1.9 | 0 | 0.56 | 0.13 | Pwp2 |
| Ap1s1_P61967 | 1.51 | -1.3 | 1.66 | -0.8 | -0.5 | -0.1 | -0.6 | 0.08 | 0.08 | Ap1s1 |
| Prmt1_Q9JIF0 | -2 | 0.58 | 0.7 | 0.75 | -0.2 | 0.44 | -0.6 | -0.6 | 1.05 | Prmt1 |
| Xpnpep1_Q6P1B1 | 1.31 | 0.31 | -0 | -0.3 | -2 | 0.97 | -0.9 | 0.34 | 0.28 | Xpnpep1 |
| Add1_Q9QYC0 | -1.4 | 1.3 | 1.05 | 0.51 | -1.3 | 0.83 | -0.7 | -0.6 | 0.18 | Add1 |
| Lmod2_Q3UHZ5 | -1.1 | 1.14 | 0.2 | 0.61 | -0.4 | -0.9 | -1.5 | 0.99 | 0.94 | Lmod2 |
| G3bp1_P97855 | -1.8 | -0.6 | 0.86 | 1.19 | -1.2 | 0.25 | 0.69 | 0.09 | 0.52 | G3bp1 |
| Arl3_Q9WUL7 | -1.4 | 0.89 | 0.69 | 0.75 | -1.5 | 0.07 | -0.5 | -0.2 | 1.27 | Arl3 |
| Huwe1_Q7TMY8 | -1.8 | 0.09 | 1.04 | -0.1 | -1.3 | 1.2 | 0.19 | -0.1 | 0.8 | Huwe1 |
| Rpl15_Q9CZM2 | -0.6 | 0.58 | 0.74 | 1.07 | -1.5 | -0.6 | -1.3 | 0.45 | 1.11 | Rpl15 |
| Cyb5r1_Q9DB73 | -0.6 | 1.28 | 1.33 | 0.72 | -0.7 | -1.6 | -0.3 | -0.6 | 0.5 | Cyb5r1 |
| Uchl3_Q9JKB1 | -0.3 | -1 | 1.9 | -0.4 | -1.3 | 0.86 | 0.62 | -0.5 | 0.17 | Uchl3 |
| Mydgf_Q9CPT4 | -1.6 | 0.64 | 1.05 | -0.7 | -0.1 | 0.1 | -1.2 | 1.2 | 0.63 | Mydgf |
| Gstp1_P19157 | 2 | -0.4 | -1.2 | -0.6 | 1.01 | -0.1 | -0.4 | 0.04 | -0.5 | Gstp1 |
| Pcca_Q91ZA3 | -0.3 | 1.01 | -1.2 | 0.59 | 0.5 | -1.4 | -0.9 | 1.42 | 0.25 | Pcca |
| Asl_Q91YI0 | -1.2 | 0.27 | 0.48 | 0.8 | -1.7 | 0.55 | -0.9 | 0.4 | 1.24 | Asl |
| Sub1_P11031 | 0.93 | 0.43 | -1.7 | -0.9 | 1.17 | -0.7 | 1.15 | -0.3 | -0.1 | Sub1 |
| Psmd11_Q8BG32 | 1.5 | -0.8 | -1.1 | -0.1 | 0.56 | -0.8 | -0.2 | 1.58 | -0.6 | Psmd11 |
| Ptpn23_Q6PB44 | -0.8 | 0.11 | 0.76 | -0.2 | -0.4 | 1.11 | -0.6 | -1.6 | 1.57 | Ptpn23 |
| Crkl_P47941 | 1.05 | -1.9 | 0.53 | 0.02 | -1.2 | 0.82 | -0.1 | -0.1 | 0.92 | Crkl |
| Vdac2_Q60930 | 1.39 | 1.41 | 0.59 | -0.3 | -0 | -1.7 | -0.5 | -0.3 | -0.5 | Vdac2 |
| Rps14_P62264 | 0.45 | -0.3 | -1.9 | 0.88 | 0 | 0.62 | 0.2 | 1.23 | -1.1 | Rps14 |

| | | | | | | | | | | |
|---|---|---|---|---|---|---|---|---|---|---|
| Bak1_O08734 | -2 | -0.4 | 0.56 | -0.5 | 0.69 | 0.28 | 1.42 | 0.43 | -0.4 | Bak1 |
| Fkbp9_Q9Z247 | -1.3 | 1.21 | 0.81 | 0.62 | -1.8 | 0.3 | -0.3 | 0 | 0.5 | Fkbp9 |
| Rps5_P97461 | -0.2 | 1.2 | -1.4 | 0.74 | -1.8 | 0.61 | 0.09 | 0.32 | 0.43 | Rps5 |
| Psmc6_P62334 | 0.66 | -1 | 0.19 | -0.4 | -1.3 | 1.25 | 0.97 | 0.83 | -1.2 | Psmc6 |
| Txn_P10639 | -1.2 | 1.56 | 0.76 | 0.47 | -1.7 | 0.12 | -0.4 | 0.13 | 0.33 | Txn |
| Sntb2_Q61235 | -1.1 | -1.5 | 1.19 | -0.3 | 1.66 | -0.1 | 0.34 | 0.15 | -0.3 | Sntb2 |
| Pgls_Q9CQ60 | -0.9 | 0.21 | -0.4 | -0.6 | 2 | -0.6 | 0.89 | -0.6 | -0.3 | Pgls |
| Actr1b_Q8R5C5 | -1.9 | 0.54 | 0.27 | 1.17 | -1.1 | -0 | -0.4 | 0.5 | 0.95 | Actr1b |
| Myo1c_Q9WTI7 | 2 | -0.3 | 0.33 | -1.4 | 0.4 | 0.49 | -0.4 | -1.1 | 0 | Myo1c |
| Cox6b1_P56391 | 0.51 | -0.3 | -2 | -0.5 | 0.19 | 0.82 | 1.39 | 0.22 | -0.2 | Cox6b1 |
| Map7d1_A2AJI0 | 1.86 | 0.92 | -1.6 | -0.2 | 0.14 | -0 | -0.9 | -0.5 | 0.3 | Map7d1 |
| Hnrnpl_Q8R081 | -1.7 | 0.93 | 0.87 | 0.58 | -1.4 | 0.28 | -0.5 | 0.12 | 0.86 | Hnrnpl |
| Tagln2_Q9WVA4 | -1 | 1.41 | 1.04 | 0.36 | -0.9 | 0.37 | -1.1 | -1 | 0.8 | Tagln2 |
| Ak3_Q9WTP7 | 0.55 | -1 | 0.35 | -1.6 | 0.23 | 1.97 | -0.3 | -0.1 | -0.1 | Ak3 |
| Dynll2_Q9D0M5 | -0.6 | -0.1 | -1 | -0.1 | 0.05 | 0.37 | 2 | 0.33 | -1.3 | Dynll2 |
| Rpl14_Q9CR57 | 1.46 | -1.4 | -0.9 | -0.3 | 1.18 | -0.9 | 0.66 | 0.41 | -0.1 | Rpl14 |
| Nup93_Q8BJ71 | 1.35 | -0.6 | -0.9 | 0.28 | -1.4 | 1.68 | -0.5 | -0.1 | 0.08 | Nup93 |
| Rhog_P84096 | -1.6 | 0.68 | 0.45 | 0.8 | -1.7 | 0.67 | -0 | -0 | 0.82 | Rhog |
| Cul7_Q8VE73 | -1.6 | -0.1 | 1.16 | 0.93 | -1.6 | 0.44 | -0.2 | 0.56 | 0.36 | Cul7 |
| Ide_Q9JHR7 | -0 | 0.45 | -0.1 | -1.2 | 0.92 | 0.56 | -0.5 | 1.5 | -1.6 | Ide |
| Psma5_Q9Z2U1 | 0.59 | -0.3 | -1.6 | -0.5 | 1.17 | 0.55 | 0.67 | -1.5 | 0.8 | Psma5 |
| Gars_Q9CZD3 | 1.04 | 0.23 | -0.8 | 1.3 | -1.9 | 0.9 | -0.4 | -0.2 | -0.2 | Gars |
| Srsf6_Q3TWW8 | -1.9 | 0.81 | -0.3 | 1.2 | -0.9 | 0.9 | 0.12 | -0.4 | 0.52 | Srsf6 |
| Clip2_Q9Z0H8 | -1.1 | -1.7 | 0.52 | -0.1 | -0.1 | 0.89 | 1.56 | -0.5 | 0.45 | Clip2 |
| Tmem106b_Q80X71 | 0.03 | 0.98 | 0.02 | -0.5 | 1.8 | -1.2 | -1.4 | 0.09 | 0.25 | Tmem106b |
| Lox_P28301 | -1.8 | 1.22 | 0.66 | 0.62 | -1.2 | 0.97 | -0.4 | -0.3 | 0.17 | Lox |
| Acot9_Q9R0X4 | -1 | -0.8 | -0.3 | 1.23 | 0.46 | -1.1 | -0.2 | -0.2 | 1.83 | Acot9 |
| Vars_Q9Z1Q9 | 0.44 | -1.7 | 1.83 | -0.1 | -0.8 | 0.11 | -0.7 | 0.41 | 0.49 | Vars |
| Cbr1_P48758 | 0.3 | 0.27 | -0.3 | -1.5 | 0.89 | -0.1 | 1.38 | 0.58 | -1.6 | Cbr1 |
| Hsd17b11_Q9EQ06 | 1.27 | -0.6 | 0.57 | -0.4 | 0.81 | -1.5 | -0.2 | 1.15 | -1.1 | Hsd17b11 |
| Aga_Q64191 | 0.41 | 0.4 | -1.1 | 0.12 | -1.4 | 1.93 | -0.6 | -0.3 | 0.53 | Aga |
| Ddx5_Q61656 | -1.4 | 0.94 | 0.85 | 0.86 | -1.3 | 0.77 | -0.9 | -0.4 | 0.62 | Ddx5 |
| Sgpl1_Q8R0X7 | 1.02 | -1.5 | 1.2 | 0.97 | -0.4 | -1.4 | -0.1 | -0 | 0.27 | Sgpl1 |
| Nenf_Q9CQ45 | 1.23 | 0.11 | 0.84 | -1.5 | 0.58 | 0.23 | -0.1 | -1.7 | 0.32 | Nenf |
| Sun2_Q8BJS4 | -1.1 | 1.48 | 0.97 | 0.65 | -0.3 | -1.5 | -0.8 | 0.58 | 0.01 | Sun2 |
| Pa2g4_P50580 | 1.98 | -0.2 | -0.6 | -0 | 0.69 | -1.7 | -0 | 0.26 | -0.3 | Pa2g4 |
| Vps29_Q9QZ88 | 0.11 | -0.9 | -0.3 | -1.2 | 1.94 | 0.31 | 1.12 | -0.5 | -0.6 | Vps29 |
| Pura_P42669 | -0.7 | 1.42 | 0.26 | 1.21 | -0.6 | -1.6 | 0.45 | -0.8 | 0.36 | Pura |
| Slc25a12_Q8BH59 | 1.4 | 0.04 | -0.2 | 1.66 | -0.2 | -1.6 | -0.7 | -0.2 | -0.2 | Slc25a12 |
| Erp44_Q9D1Q6 | 0.73 | -0.7 | 1.23 | -0.9 | 1.75 | -1 | -0.1 | -0.7 | -0.3 | Erp44 |
| Colgalt1_Q8K297 | 0.28 | -1 | 1.27 | 1.07 | 0.88 | -1.8 | -0 | -0.2 | -0.4 | Colgalt1 |
| Ssr3_Q9DCF9 | 1 | -1.9 | 1.39 | -0.3 | 0.19 | 0.33 | 0.55 | -0.9 | -0.3 | Ssr3 |
| Lsm2_O35900 | 1.58 | -0.3 | 0.84 | 0.93 | -0.7 | -1 | -0.3 | 0.39 | -1.5 | Lsm2 |
| Ctsd_P18242 | 1.4 | -1.2 | -1.5 | -0.7 | 0.86 | 0.11 | 0.96 | 0.19 | -0 | Ctsd |
| Rab18_P35293 | -1.6 | 0.79 | 1.36 | 0.21 | -1.5 | 0.58 | -0 | -0.2 | 0.4 | Rab18 |
| Col3a1_P08121 | 1.82 | 0.95 | -1.3 | -0.2 | 0.77 | -0.5 | -0.8 | 0.04 | -0.7 | Col3a1 |
| Fbl_P35550 | -0.4 | 0.17 | 2 | 0.43 | -1.1 | -0.8 | -0 | 0.58 | -1 | Fbl |
| Gbas_O55126 | 1.32 | -1.2 | 1.21 | -0.8 | 0.31 | -0.6 | 0.39 | -1.2 | 0.68 | Gbas |
| Scamp3_O35609 | -2 | 0.23 | 0.77 | 0.26 | -0.1 | 0.83 | -1 | 0.13 | 1 | Scamp3 |
| Hnrnpa3_Q8BG05 | -1.4 | 0.98 | 0.97 | 0.32 | -1.7 | 0.65 | -0.2 | -0.4 | 0.76 | Hnrnpa3 |

| | | | | | | | | | | |
|---|---|---|---|---|---|---|---|---|---|---|
| Farp1_F8VPU2 | -0.9 | 0.31 | 0.02 | 0.31 | 1.25 | -0.8 | 0.21 | 1.34 | -1.7 | Farp1 |
| Bola1_Q9D8S9 | -1.4 | 0.9 | 1.41 | 0.64 | -1.4 | 0.7 | -0.6 | 0.02 | -0.3 | Bola1 |
| Ptms_Q9D0J8 | -1.8 | 0.8 | 0.58 | 0.64 | -1.5 | 0.68 | -0.1 | 0.13 | 0.67 | Ptms |
| Ruvbl1_P60122 | 1.17 | -1.2 | -0.7 | 0.84 | 1.09 | -1.1 | 0.9 | -0.1 | -0.9 | Ruvbl1 |
| Rpl36_P47964 | 1.02 | 1.73 | -0.3 | -0.1 | -0.3 | -1.2 | -1.5 | 0.45 | 0.17 | Rpl36 |
| Cox5b_P19536 | 1.63 | -1.6 | -0.9 | -0.6 | 0.95 | -0.3 | 0.61 | 0.02 | 0.29 | Cox5b |
| Serbp1_Q9CY58 | -2 | 1.12 | 0.6 | 0.47 | -0.7 | 0.92 | -0.3 | -0.1 | 0.2 | Serbp1 |
| Hnrnpa1_P49312 | -0.7 | 0.39 | 1.73 | -0 | 1.18 | -1.3 | -1.2 | 0.1 | -0.3 | Hnrnpa1 |
| Phgdh_Q61753 | 1.59 | -0 | -0.7 | -1.3 | 1.68 | -0.2 | -0.3 | -0.4 | -0.3 | Phgdh |
| Bzw1_Q9CQC6 | 1.81 | 0.17 | 0.4 | -0 | 0.15 | -1.7 | -0.3 | 0.61 | -1.1 | Bzw1 |
| Mtpn_P62774 | 0.66 | 0.5 | 0.39 | 0.44 | 0.67 | -1.5 | 0.51 | -2 | 0.31 | Mtpn |
| Ddx3x_Q62167 | -1.9 | 0.06 | 0.73 | 0.58 | -1.3 | 0.78 | 0.67 | 0.78 | -0.4 | Ddx3x |
| Diablo_Q9JIQ3 | -1.8 | 1.14 | 0.67 | 0.13 | -1.2 | 0.63 | -0.7 | 0.64 | 0.51 | Diablo |
| Tmco1_Q921L3 | -0.1 | -1.8 | -1.1 | 1.12 | -0.3 | 1.02 | 0.35 | 0.93 | -0 | Tmco1 |
| Iars_Q8BU30 | 2 | -0.6 | 0.03 | -0.5 | 0.32 | -0.2 | -0.9 | 0.75 | -1 | Iars |
| H2afy_Q9QZQ8 | -2 | -0.4 | 1.26 | -0.1 | 0.08 | 0.2 | 1.3 | -0.3 | 0.1 | H2afy |
| Tpd52l2_Q9CYZ2 | 0.21 | 0.78 | 0.35 | -0.5 | -1.7 | 1.14 | 1.24 | -0.7 | -0.7 | Tpd52l2 |
| Esyt2_Q3TZZ7 | -1 | -1.4 | 0.35 | 1.4 | -0.2 | 0.24 | 1.22 | 0.48 | -1 | Esyt2 |
| Gdi1_P50396 | 1.71 | 0.71 | 1.06 | -0.6 | 0.27 | -1.2 | -0.2 | -1 | -0.8 | Gdi1 |
| Pgrmc1_O55022 | 1.61 | -0.4 | 1.05 | -0.2 | 0.61 | -1.9 | -0.3 | -0.3 | -0.2 | Pgrmc1 |
| G6pdx_Q00612 | 0.18 | -1.6 | -0.7 | 0.12 | 1.51 | -1 | 0.47 | 1.19 | -0.2 | G6pdx |
| Stmn1_P54227 | -0.7 | 1.2 | 0.66 | 0.59 | -2 | 0.47 | -0.7 | -0.1 | 0.59 | Stmn1 |
| Puf60_Q3UEB3 | 1.03 | -0.6 | -1.5 | -0.8 | 1.49 | -0.6 | 1.04 | 0.05 | -0.2 | Puf60 |
| Rbbp9_O88851 | 1.99 | -0.1 | -0.9 | -1.3 | 0.24 | 0.99 | -0.5 | 0.11 | -0.5 | Rbbp9 |
| Rpl8_P62918 | 1.38 | -1.2 | -0.4 | -0.6 | 1.77 | -0.6 | 0.05 | 0.33 | -0.8 | Rpl8 |
| Chmp4b_Q9D8B3 | 0.1 | 1.03 | -1.1 | 0.88 | 1.28 | -1.8 | 0.07 | -0.5 | -0 | Chmp4b |
| Hdgf_P51859 | 1.81 | -0.7 | -0.2 | -0.7 | 1.58 | -1 | -0.4 | -0 | -0.4 | Hdgf |
| Afap1_Q80YS6 | 2 | 0.51 | -0.3 | -0.7 | 0.53 | -0.8 | -0.3 | -0.9 | -0.4 | Afap1 |
| Rragc_Q99K70 | 0.87 | -0.1 | -1.3 | 0.86 | 0.91 | -1.9 | 0.46 | 0 | 0.21 | Rragc |
| Fxr1_Q61584 | -0.8 | 0.59 | -0.8 | -0.1 | 1.95 | -1 | -0.6 | -0.4 | 1.07 | Fxr1 |
| Cisd1_Q91WS0 | -0.3 | 0.53 | -0.7 | -0.9 | 1.5 | -0.8 | 1.06 | 0.86 | -1.3 | Cisd1 |
| Ero1l_Q8R180 | -0.2 | 0.59 | 1.64 | 0.35 | 0.02 | -1.8 | -0.9 | -0.5 | 0.75 | Ero1l |
| Arl8b_Q9CQW2 | -1.2 | 1.95 | 0 | -0.5 | -0.9 | 0.58 | -0.1 | 0.89 | -0.7 | Arl8b |
| Ppp2r4_P58389 | 1.48 | -0.4 | -0.5 | -1.5 | 1.38 | -0.6 | -0.7 | 0.07 | 0.68 | Ppp2r4 |
| Vapa_Q9WV55 | -0 | 0.53 | 1.35 | -0.4 | 0.94 | -0.8 | 0.64 | -0.3 | -1.9 | Vapa |
| Serpinf1_P97298 | 0.83 | 0.26 | -0.2 | 1 | 0.58 | -1.6 | 0.44 | 0.42 | -1.7 | Serpinf1 |
| Aprt_P08030 | -1.8 | 1.26 | 0.8 | 0.22 | -1.4 | 0.64 | -0.3 | 0.29 | 0.23 | Aprt |
| Srrt_Q99MR6 | -1.3 | 0.96 | 0.45 | 0.89 | -1.7 | 0.35 | -0.6 | -0 | 0.99 | Srrt |
| Anxa7_Q07076 | 1.37 | -0.4 | -0.3 | 1.09 | 0.02 | -0.9 | -0.9 | -1.2 | 1.25 | Anxa7 |
| Atp5l_Q9CPQ8 | 1.36 | -0.6 | -1.3 | 0.75 | 0.5 | -1.4 | 0.14 | 1.06 | -0.5 | Atp5l |
| Ehd1_Q9WVK4 | 0.03 | 0.27 | -0.4 | 0.26 | -1.7 | 2 | -0.5 | 0.46 | -0.5 | Ehd1 |
| Usmg5_Q78IK2 | 1.17 | -0.3 | 0.55 | -1.7 | -0.2 | 0.86 | 1.08 | -0.2 | -1.2 | Usmg5 |
| Upf1_Q9EPU0 | 1.17 | -0.5 | 0.64 | 1.33 | -1.8 | 0.19 | 0.18 | -0.3 | -0.9 | Upf1 |
| Purb_O35295 | 0.64 | 0.56 | 0.17 | 1.78 | -0.4 | -1.6 | -0.3 | -1 | 0.22 | Purb |
| Ptrf_O54724 | 1.27 | 0.15 | -0.9 | 0.23 | 0.94 | -1.9 | 0.57 | 0.4 | -0.8 | Ptrf |
| Rpn2_Q9DBG6 | -0 | 0.96 | -0.7 | -0.7 | 1.35 | -1.2 | -0.9 | 1.37 | -0.1 | Rpn2 |
| Psma2_P49722 | 0.84 | -0.9 | -1.1 | 1.95 | -0.4 | -0.6 | -0.1 | -0.5 | 0.82 | Psma2 |
| Apool_Q78IK4 | 0.08 | -0.3 | -1.2 | 1.34 | -0.7 | -0.4 | -1 | 1.65 | 0.5 | Apool |
| Hist1h2bc_Q6ZWY9 | -1.2 | 0.56 | 0.68 | 0.67 | -1.7 | 0.69 | -0.9 | 0.34 | 0.93 | Hist1h2bc |
| Ssbp1_Q9CYR0 | 1.55 | 0.91 | 0.46 | -1.5 | 0.11 | -0.3 | 0.3 | -0.1 | -1.4 | Ssbp1 |

| | | | | | | | | | | |
|---|---|---|---|---|---|---|---|---|---|---|
| Ndufs3_Q9DCT2 | 0.25 | -1.8 | -0.9 | -0.4 | 0.91 | 0.98 | 0.9 | 0.79 | -0.7 | Ndufs3 |
| Ptk7_Q8BKG3 | 0.32 | -0.8 | -1.1 | 1.58 | 0.03 | -0.5 | -1.3 | 1.11 | 0.63 | Ptk7 |
| Ctsl_P06797 | 1.06 | -0.5 | 1.2 | 0.65 | 0.03 | -1.2 | 0.9 | -1.4 | -0.7 | Ctsl |
| Sept5_Q9Z2Q6 | -0.1 | 1.17 | 0.2 | 0.07 | -2 | 1.18 | 0.3 | -0.5 | -0.1 | Sept5 |
| Usp14_Q9JMA1 | 0.33 | -0.1 | 1.05 | 0.79 | 0.49 | -2 | -0.7 | 0.59 | -0.2 | Usp14 |
| Casp3_P70677 | -0.4 | 1.1 | 0.85 | 1.14 | -1.9 | 0.26 | -0 | -1.1 | -0 | Casp3 |
| Wbp11_Q923D5 | -0.4 | 0.75 | 0.96 | 1.49 | -0.3 | -1.5 | 0.33 | -0 | -1.3 | Wbp11 |
| Col1a2_Q01149 | -1.2 | 1.25 | 0.85 | 1.01 | -1.5 | 0.34 | -0.9 | -0.4 | 0.44 | Col1a2 |
| Rtn4_Q99P72 | -1.8 | 1.63 | 1.18 | -0.3 | -0.3 | 0.52 | -0.5 | -0.4 | -0.1 | Rtn4 |
| Mtdh_Q80WJ7 | 1.04 | -1.8 | 0.92 | -0.9 | 1.01 | -0.6 | -0.5 | 0.7 | 0.06 | Mtdh |
| _Q9CQE8 | 0.98 | -0.7 | -0.1 | -1.4 | -0.8 | 1.68 | -0.3 | 0.98 | -0.3 | |
| Rpl23a_P62751 | 1.55 | -0 | -1.1 | -0.5 | 0.92 | -0 | 1.12 | -1.3 | -0.6 | Rpl23a |
| Atl3_Q91YH5 | -0.6 | -0.5 | 1.89 | -1 | -0.2 | 0.42 | 0.62 | 0.74 | -1.2 | Atl3 |
| Tsc22d3_Q9Z2S7 | 0.34 | -0.1 | -2 | -0.2 | 0.46 | 1.15 | 0.56 | 0.05 | 0.2 | Tsc22d3 |
| Tpm4_Q6IRU2 | -1.1 | 1.15 | 1.16 | 0.89 | -1.4 | 0.26 | -1 | -0.4 | 0.42 | Tpm4 |
| Chd4_Q6PDQ2 | 1.53 | -0 | 0.28 | -2 | 1.02 | -0.2 | -0 | -0 | -0.5 | Chd4 |
| Sacm1l_Q9EP69 | 1.94 | 0.45 | -0.3 | 0.61 | -1.5 | 0.17 | -0.8 | 0.19 | -0.8 | Sacm1l |
| Ppt1_O88531 | -0.9 | 1.54 | 0.52 | -0.2 | 0.21 | -1 | 0.03 | 1.2 | -1.4 | Ppt1 |
| Ndufa7_Q9Z1P6 | 1.06 | 0.12 | -1.1 | 1.19 | -1.7 | 0.8 | 0.56 | -0.5 | -0.4 | Ndufa7 |
| Ewsr1_Q61545 | 1 | 0.07 | 0.69 | -1.7 | 1.4 | -0.3 | -1.1 | -0.4 | 0.35 | Ewsr1 |
| Lars_Q8BMJ2 | -1.2 | -0.2 | -0.2 | 1.28 | 0.71 | -0.8 | -1.4 | 0.7 | 1.12 | Lars |
| Eif3l_Q8QZY1 | 1.68 | -0.7 | 0.13 | 0.23 | 0.81 | -1.9 | -0.5 | 0.12 | 0.16 | Eif3l |
| Ola1_Q9CZ30 | -0.1 | 1.88 | -0 | 0.62 | -1.6 | 0.52 | -0.9 | 0.14 | -0.6 | Ola1 |
| Eif4b_Q8BGD9 | -1.5 | 0.7 | 0.51 | 0.81 | -1.7 | 0.7 | -0.5 | 0.12 | 0.84 | Eif4b |
| Rpl10_Q6ZWV3 | 1.46 | -0.7 | -1.1 | -0 | 1.19 | -1.4 | 0.68 | -0.6 | 0.43 | Rpl10 |
| Epb41l3_Q9WV92 | -1.3 | 1.35 | 1.37 | 0.58 | -0.6 | -1 | -0.5 | -0.5 | 0.6 | Epb41l3 |
| Glud1_P26443 | 1.37 | 0.23 | -0.4 | -1.8 | 1.09 | -0.3 | 0.18 | 0.62 | -1 | Glud1 |
| Tor1aip1_Q921T2 | 1.06 | 1.07 | -0.5 | 1 | -0.8 | -0.7 | -0.1 | 0.65 | -1.7 | Tor1aip1 |
| Dak_Q8VC30 | 0.57 | 0.42 | 0.32 | -1.8 | 0.8 | 0.61 | -1.7 | 0.34 | 0.42 | Dak |
| Cct2_P80314 | 1.69 | -0.6 | -0.3 | -1.4 | 0.92 | -0.3 | 1.1 | -0.5 | -0.6 | Cct2 |
| Rpn1_Q91YQ5 | -1.8 | 1.06 | 0.83 | 0.93 | -1.3 | -0.1 | -0.3 | -0.1 | 0.69 | Rpn1 |
| Rps12_P63323 | 0.56 | 0.15 | 0.22 | -0.6 | 1.28 | -1.7 | 1.35 | -0.5 | -0.7 | Rps12 |
| Atp6v0d1_P51863 | 0.67 | -0.3 | -0.3 | -0.6 | 1.83 | -1.6 | -0.1 | -0.4 | 0.87 | Atp6v0d1 |
| Pfdn6_Q03958 | -0.8 | -0.3 | -1.4 | 1.42 | -0.6 | 0.14 | -0.1 | 1.71 | -0.1 | Pfdn6 |
| Dctn4_Q8CBY8 | 0.39 | 1.3 | 0.38 | 0.87 | -0.1 | -2 | -0.3 | -0.8 | 0.33 | Dctn4 |
| Eif3f_Q9DCH4 | -2 | 0.81 | 0.25 | 0.7 | -1.3 | 0.72 | -0.1 | 0.09 | 0.82 | Eif3f |
| Lgals3_P16110 | -1.2 | 0.86 | 0.63 | -0.2 | -0.8 | 1.24 | -1.5 | 0.02 | 0.98 | Lgals3 |
| Trim23_Q8BGX0 | -1.3 | -0.2 | 1.12 | 1.77 | -0.6 | -0.7 | -0.6 | 0.82 | -0.4 | Trim23 |
| Cast_P51125 | -0.6 | 0.81 | 0.55 | -0.2 | 0.12 | 0.05 | -2 | 0.04 | 1.38 | Cast |
| Xdh_Q00519 | -1.7 | 0.58 | 1.37 | -1.4 | 0.47 | 0.41 | -0.3 | 0.64 | -0.1 | Xdh |
| Golph3_Q9CRA5 | 1.2 | 0.06 | 0.07 | 0.54 | 0.67 | -1.6 | 1.05 | -1.3 | -0.7 | Golph3 |
| Hars_Q61035 | 1.36 | 0.36 | -0.5 | -0.9 | 0.99 | -0.4 | -1.6 | 1.03 | -0.3 | Hars |
| Camk2d_Q6PHZ2 | 0.96 | 0.17 | -1.4 | -0 | 1.6 | -1.1 | 0.75 | -0.1 | -0.9 | Camk2d |
| Raly_Q64012 | -0.7 | 0.33 | 0.16 | 1.63 | -1.3 | 0.12 | -0.7 | 1.31 | -0.8 | Raly |
| Rps6_P62754 | 0.14 | 0.1 | -1 | 1.94 | -1.5 | 0.45 | -0.8 | 0.11 | 0.51 | Rps6 |
| Chmp2b_Q8BJF9 | 2 | -0.7 | 0.14 | 0.5 | -1.1 | 0.09 | -0.2 | 0.19 | -1.1 | Chmp2b |
| Tm9sf2_P58021 | -1.5 | 0.33 | 0.2 | 1.65 | -1.3 | 0.47 | 0.17 | 0.67 | -0.7 | Tm9sf2 |
| Hnrnpa0_Q9CX86 | -0.2 | 1.2 | 1 | 0.22 | -1.9 | 0.95 | -0 | -0.9 | -0.3 | Hnrnpa0 |
| Rab6a_P35279 | -0.7 | 0.02 | -0.9 | -1.3 | 1.11 | 1.29 | 1.22 | 0.05 | -0.8 | Rab6a |
| Snrpf_P62307 | -0.1 | -0.4 | -0.5 | 1.08 | -0.5 | 0.16 | -1.9 | 0.62 | 1.44 | Snrpf |

| | | | | | | | | | | |
|---|---|---|---|---|---|---|---|---|---|---|
| Snrpg_P62309 | -0.9 | 1.21 | 0.18 | 0.37 | -1.7 | 1.4 | -0.8 | 0.14 | 0.13 | Snrpg |
| Ppia_P17742 | 1.68 | -0.7 | -0.8 | -0.9 | 1.67 | -0.5 | 0.03 | -0.2 | -0.2 | Ppia |
| Lgals1_P16045 | 1.29 | -0.3 | 0.96 | -0.6 | 0.93 | -1.8 | -0.5 | -0.5 | 0.56 | Lgals1 |
| Sypl1_O09117 | -0.3 | 0.91 | 1.12 | -0.3 | -1.9 | 0.99 | 0.12 | -1.1 | 0.36 | Sypl1 |
| Rpl27_P61358 | 2 | -0.7 | 0.37 | -0.1 | -0.5 | -0.7 | 0.65 | -0.1 | -1.2 | Rpl27 |
| Sf3b3_Q921M3 | 1.02 | 1.05 | -0.4 | 1.18 | -1.9 | -0.4 | -0.7 | 0.06 | 0.07 | Sf3b3 |
| Cul5_Q9D5V5 | 0.58 | 0.61 | 0.16 | -1.7 | 0.34 | 0.44 | 0.66 | 0.69 | -1.8 | Cul5 |
| Ppap2b_Q99JY8 | 1.13 | -1.4 | -0.9 | 1.53 | -1.1 | 0.45 | 0.42 | 0.05 | -0.2 | Ppap2b |
| Nop56_Q9D6Z1 | -1.5 | -0.3 | 0.7 | 0.79 | -1.4 | 1.48 | -0.4 | 0.25 | 0.41 | Nop56 |
| Tmed4_Q8R1V4 | 1.05 | -0.4 | 0.7 | 0.92 | 0.58 | -1.9 | -1.2 | 0.13 | 0.11 | Tmed4 |
| Atp6v0c_P63082 | -0.2 | -0.5 | -0.5 | 1.74 | 0.31 | -1.7 | -0.2 | 1.2 | -0.1 | Atp6v0c |
| Dhrs7_Q9CXR1 | 0.77 | -0.7 | 1.1 | 0.06 | -1.1 | 0.76 | -1.6 | -0.4 | 1.12 | Dhrs7 |
| Acaa1a_Q921H8 | 1.41 | -1.8 | -0.6 | 0.52 | 0.63 | -1 | 0.63 | 0.55 | -0.4 | Acaa1a |
| Atp6v0a1_Q9Z1G4 | 1.06 | -0.3 | -1.4 | -1.1 | 0.65 | 1.21 | 0.51 | 0.46 | -1 | Atp6v0a1 |
| Pkm_P52480 | 1.81 | -0.5 | -0.3 | -1 | 1.48 | -0.6 | 0.28 | -0.5 | -0.7 | Pkm |
| Mical1_Q8VDP3 | -1.6 | 0.95 | 0.2 | 0.61 | -1.6 | 0.94 | 0.58 | 0.3 | -0.3 | Mical1 |
| Mxra7_Q9CZH7 | -1.5 | 0.03 | 1.07 | 1.25 | -1.4 | 0.12 | -0.3 | -0.1 | 0.88 | Mxra7 |
| Hmga2_P52927 | -1.9 | 0.77 | 0.69 | 0.59 | -1.5 | 0.84 | 0.02 | 0.1 | 0.35 | Hmga2 |
| Atp6v1e1_P50518 | 0.01 | -0.5 | 0 | -1.5 | 1.77 | -0.4 | -0.2 | 1.35 | -0.5 | Atp6v1e1 |
| Copb2_O55029 | -0 | 0.65 | -0.3 | -0.4 | 1.07 | -0.6 | 1.65 | -0.4 | -1.7 | Copb2 |
| Pdia6_Q922R8 | 1.78 | -1.3 | -0.8 | -0.3 | 0.62 | -0.5 | 1.15 | 0.07 | -0.8 | Pdia6 |
| Rpl29_P47915 | -1.7 | 1.06 | 0.81 | 0.87 | -1.4 | 0.68 | -0.5 | 0.05 | 0.11 | Rpl29 |
| Gcat_O88986 | 0.78 | -0 | 0.23 | -0.3 | -1.3 | 1.38 | -0.6 | 1.19 | -1.4 | Gcat |
| Rps17_P63276 | -1.3 | 0.14 | -0.5 | 0.1 | -1 | 1.45 | -0.8 | 0.48 | 1.42 | Rps17 |
| Capza2_P47754 | 2 | -0.9 | -0.6 | -0.6 | 0.78 | -0.1 | 0.8 | -0.4 | -0.9 | Capza2 |
| Kpna4_O35343 | -1.6 | 0.38 | 0.93 | 1.44 | -1.4 | -0.1 | -0.2 | 0.23 | 0.4 | Kpna4 |
| Prosc_Q9Z2Y8 | 0.76 | -1.8 | -0.7 | -0.8 | 1.63 | 0.25 | 0.57 | -0.2 | 0.26 | Prosc |
| Acot2_Q9QYR9 | 1.75 | 0.43 | -0.4 | -1.1 | 1.12 | -1.1 | 0.39 | -0.3 | -0.8 | Acot2 |
| Ubxn4_Q8VCH8 | 0.27 | -2 | 0.64 | 0.19 | -0.4 | 0.79 | 0.22 | 0.48 | 0.34 | Ubxn4 |
| Ruvbl2_Q9WTM5 | 0.87 | 0.49 | -0.8 | -0.7 | 1.5 | -1.3 | -0.9 | 1.03 | -0.1 | Ruvbl2 |
| Tmx1_Q8VBT0 | 0.88 | -0.8 | -1 | -1.1 | 1.32 | -0 | 1.5 | -0.3 | -0.6 | Tmx1 |
| Ap1b1_O35643 | -1.5 | 0.81 | 0.17 | 1.48 | -1.4 | 0.48 | 0.55 | -0.5 | -0 | Ap1b1 |
| Nutf2_P61971 | 1.33 | 0.13 | -1.2 | -1.3 | 0.46 | 0.92 | -0.9 | 1.03 | -0.5 | Nutf2 |
| Top1_Q04750 | 0.21 | -0.1 | 1.46 | -0.6 | 1.4 | -1.8 | 0.08 | -0.2 | -0.4 | Top1 |
| Map1a_Q9QYR6 | 0.17 | 1.67 | -0.3 | 0.28 | -0.6 | -0.2 | 0.93 | -0 | -1.9 | Map1a |
| Usp5_P56399 | -0 | 0.63 | 1.17 | -0.6 | -0.2 | -0.4 | -1 | -1.2 | 1.75 | Usp5 |
| Rpl4_Q9D8E6 | 1.44 | -0.1 | -0.2 | 1.09 | 0.16 | -1.6 | -0.6 | 0.82 | -1.1 | Rpl4 |
| Pcnp_Q6P8I4 | -0.9 | 0.62 | 1.26 | 0.52 | -1.8 | 1.01 | 0.39 | -0.6 | -0.5 | Pcnp |
| Gaa_P70699 | -0.9 | -0.5 | 1.81 | 0.25 | 0.73 | -1.5 | -0.6 | 0.69 | 0.03 | Gaa |
| Gnb1_P62874 | -0.1 | -0.3 | 1.63 | 0.89 | 0.25 | -1.5 | -0.6 | -1.1 | 0.76 | Gnb1 |
| Rps15_P62843 | 1.55 | -0.1 | 0.91 | -0.1 | 0.85 | -1.7 | -0 | -0.4 | -1 | Rps15 |
| Ddx17_Q501J6 | 0.98 | 0.57 | 0.74 | 0.67 | 0.56 | -2 | -0.9 | -0.3 | -0.2 | Ddx17 |
| Ube2m_P61082 | -2 | 0.3 | 0.29 | 0.68 | 0.09 | -0.2 | -0.7 | 1.05 | 0.79 | Ube2m |
| H1f0_P10922 | -1.4 | 0.83 | 1.09 | 0.79 | -1.7 | 0.4 | -0.6 | 0.13 | 0.42 | H1f0 |
| Rpl32_P62911 | -0.7 | -0.8 | -1.4 | 1.66 | -0.6 | 0.56 | -0.3 | 0.42 | 1.13 | Rpl32 |
| Rpl26_P61255 | 1.22 | -1.3 | -1.3 | -0.4 | 1.21 | 0.18 | 0.26 | 0.88 | -0.7 | Rpl26 |
| Copb1_Q9JIF7 | -0.1 | 1.18 | 2 | -0.5 | -0.6 | -0.3 | -0.5 | -1.3 | 0.02 | Copb1 |
| Rplp0_P14869 | 2 | 0.09 | 0.09 | -0.7 | -0 | -0.7 | 0.36 | 0.09 | -1.4 | Rplp0 |
| Rps18_P62270 | 1.58 | -1.2 | -0.7 | -0.3 | 1.58 | -0.9 | 0.38 | -0.4 | -0.1 | Rps18 |
| Hnrnpll_Q921F4 | 0.47 | 1.34 | 1.45 | -0.1 | -1.4 | -0.4 | -0.3 | 0.14 | -1.3 | Hnrnpll |

| | | | | | | | | | | |
|---|---|---|---|---|---|---|---|---|---|---|
| Lims1_Q99JW4 | 2 | -0.6 | -0.6 | -0.5 | 0.16 | -0.1 | 0.34 | -0.1 | -1 | Lims1 |
| Impdh2_P24547 | 0.74 | 0.22 | 0.12 | 1.43 | 0.42 | -2 | 0.2 | -0.7 | -0.3 | Impdh2 |
| Arcn1_Q5XJY5 | -1 | 0.46 | 0.64 | 1.18 | -2 | 0.59 | -0 | 0.39 | -0.1 | Arcn1 |
| Prdx2_Q61171 | 1.98 | -0.5 | -0.6 | -0.7 | 1.39 | -0.9 | -0.3 | -0.5 | 0.05 | Prdx2 |
| Tcea1_P10711 | 0.67 | 0.45 | -0.8 | 0.12 | 1.45 | -1.5 | -0.5 | 1.07 | -1 | Tcea1 |
| C3_P01027 | -0.3 | 0.07 | -0 | 0.18 | -2 | 1.81 | 0.35 | 0.11 | -0.1 | C3 |
| Cuta_Q9CQ89 | 0.72 | -2 | -0.6 | -0.4 | 1.2 | 0.39 | 0.57 | 0.07 | 0.23 | Cuta |
| Pitpnb_P53811 | 0.46 | -0.4 | -0.7 | -0.6 | 2 | -1 | -0.1 | -0.6 | 0.85 | Pitpnb |
| Atpif1_O35143 | -0.9 | 0.74 | 0.42 | 0.83 | -2 | 0.85 | -0.6 | -0.1 | 0.79 | Atpif1 |
| Arfgap1_Q9EPJ9 | -1.9 | 0.6 | 0.47 | 0.93 | 0.73 | -1.4 | 0.4 | -0 | 0.26 | Arfgap1 |
| Nae1_Q8VBW6 | -2 | 0.32 | 0.5 | 0.31 | 0.02 | 0.4 | 0.32 | 0.36 | 0.41 | Nae1 |
| Glrx3_Q9CQM9 | 0.07 | 0.72 | 0.53 | 1.15 | -1.9 | -0.1 | -1.1 | 0.93 | -0.3 | Glrx3 |
| Katnal2_Q9D3R6 | 1.49 | -1.2 | -1.5 | 0.12 | 1.12 | -0.4 | 0.71 | -0.2 | -0.1 | Katnal2 |
| Rpl13a_P19253 | 0.16 | 1.18 | -0.1 | 0.77 | -1.9 | 0.3 | -0.8 | 1.05 | -0.7 | Rpl13a |
| Sqstm1_Q64337 | 1.44 | 0.63 | 0.64 | 0.42 | 0.01 | -1.9 | -0.8 | -0.8 | 0.31 | Sqstm1 |
| Nptn_P97300 | 1.84 | 0.91 | 0.1 | -1.2 | 0.78 | -0.9 | -0.8 | -0.6 | -0.2 | Nptn |
| Ywhag_P61982 | 1.68 | -0.1 | -1.4 | -0.8 | 1.05 | -0.1 | -0.8 | -0.2 | 0.76 | Ywhag |
| Agk_Q9ESW4 | 0.39 | 0.34 | -0.6 | 0.43 | 0.99 | -1.3 | 1.65 | -1 | -1 | Agk |
| Ipo4_Q8VI75 | -1.6 | 0.89 | 0.5 | 1.04 | -1.5 | 0.72 | -0.6 | -0 | 0.56 | Ipo4 |
| Rps4x_P62702 | 1.84 | 0.92 | -0.6 | -0.9 | 0.86 | -1.2 | -0.1 | -0.3 | -0.5 | Rps4x |
| Rpl10a_P53026 | 1.11 | -0.6 | -0 | -1.7 | 1.69 | -0 | 0.49 | -0.4 | -0.5 | Rpl10a |
| Hnrnpab_Q99020 | 0.84 | 0.83 | 0.23 | 0.15 | 0.47 | -1.8 | -1.2 | -0.6 | 1.1 | Hnrnpab |
| Rpl11_Q9CXW4 | 2 | -0.4 | -1.4 | -0.5 | 0.3 | -0.1 | -0.6 | -0.2 | 0.91 | Rpl11 |
| Psma3_O70435 | -1.3 | 0.77 | 0.35 | 0.57 | -1.5 | 0.79 | -1.1 | 0.28 | 1.1 | Psma3 |
| Eif3a_P23116 | 0.63 | 0.06 | 0.85 | 1.1 | -2 | 0.53 | -0.6 | 0.13 | -0.5 | Eif3a |
| Ggh_Q9Z0L8 | -0.6 | -0.4 | 1.81 | -0.1 | -1.6 | 1.01 | 0.64 | -0.4 | -0.4 | Ggh |
| Nit2_Q9JHW2 | -0.4 | -0.6 | 0.54 | -1 | 0.72 | 0.73 | -1.7 | 0.37 | 1.38 | Nit2 |
| Rpl35_Q6ZWV7 | 0.48 | -0.8 | 0.45 | -1.4 | 0.68 | 0.44 | 1.71 | -0.4 | -1.2 | Rpl35 |
| Arfgap2_Q99K28 | -2 | 0.32 | 0.55 | 0.88 | -1.2 | 0.76 | 0.01 | 0.03 | 0.73 | Arfgap2 |
| Rpia_P47968 | 0.18 | 0.47 | 0.17 | -1.5 | -0.8 | 1.67 | -0 | -1.1 | 0.91 | Rpia |
| Fkbp7_O54998 | 0.58 | 1.04 | 0.88 | 1.06 | -1.5 | -0.9 | 0.12 | -1.2 | -0 | Fkbp7 |
| Hk1_P17710 | 2 | -0.4 | 0.82 | -1.2 | 0.06 | 0.04 | -0.3 | -0.9 | -0.2 | Hk1 |
| Prpf8_Q99PV0 | 1.19 | -0.1 | 1.12 | -1.8 | -0.2 | 0.98 | -0.5 | 0.14 | -0.9 | Prpf8 |
| Rps23_P62267 | 1.63 | -0.1 | -0.7 | 1.46 | -1 | -0.6 | 0.55 | -0.3 | -0.9 | Rps23 |
| Vamp8_O70404 | -1.5 | -0.7 | 1.24 | -0.2 | 0.43 | -0.1 | -0.4 | 1.73 | -0.5 | Vamp8 |
| Rraga_Q80X95 | 0.73 | -0.1 | -0.7 | -1.3 | 1.12 | -0 | 1.72 | -0.7 | -0.9 | Rraga |
| Atp6v1d_P57746 | -0.2 | 1.28 | 1.03 | 0.35 | 0.41 | -1.9 | -0.9 | -0.5 | 0.45 | Atp6v1d |
| Banf1_O54962 | 0.56 | 1.16 | -0.1 | 0.89 | -1.4 | -0.4 | -1.6 | 0.17 | 0.79 | Banf1 |
| Tra2b_P62996 | -2 | 1.3 | 0.32 | -0.1 | -0.6 | 0.76 | -0.4 | -0 | 0.83 | Tra2b |
| Sarnp_Q9D1J3 | 1.78 | -1.2 | -0.4 | -0.5 | 1.38 | -0.8 | -0.3 | 0.4 | -0.4 | Sarnp |
| Rpl38_Q9JJI8 | 0.08 | 0.2 | -0.4 | -2 | 0.93 | 0.97 | 0.5 | 0.8 | -1.1 | Rpl38 |
| Strap_Q9Z1Z2 | 0.28 | -1.3 | -1.1 | 0.45 | 0.71 | 0.02 | 1.85 | 0.02 | -1 | Strap |
| Gcn1l1_E9PVA8 | 1.27 | 0.56 | 0.21 | 0.62 | 0.2 | -1.7 | 0.11 | 0.32 | -1.6 | Gcn1l1 |
| Psmd8_Q9CX56 | 1.03 | 0.02 | 0.91 | 0.04 | -2 | 0.92 | 0.06 | -0.9 | 0.05 | Psmd8 |
| Pgp_Q8CHP8 | -1.9 | -0.1 | 0.15 | 0.51 | -0.5 | 0.73 | -1.1 | 1.01 | 1.12 | Pgp |
| Twf1_Q91YR1 | 1.16 | 0.86 | -0.5 | 0.71 | 0.74 | -2 | -0.4 | 0 | -0.5 | Twf1 |
| Nme1_P15532 | 1.04 | -0.6 | 0.75 | -0.4 | -0.8 | 0.49 | 0.37 | 1.1 | -1.9 | Nme1 |
| Ppid_Q9CR16 | 0.91 | -0.2 | 0.53 | -0 | -2 | 1.59 | -0.1 | -0.5 | -0.2 | Ppid |
| Mt1_P02802 | -2 | 0.79 | 0.76 | 0.41 | -0.8 | 0.64 | -0.1 | 0.41 | 0.18 | Mt1 |
| Clic4_Q9QYB1 | 0.74 | 1.11 | -1.1 | 1.51 | -0.5 | -1.2 | 0.55 | -0.3 | -0.7 | Clic4 |

| | | | | | | | | | | |
|---|---|---|---|---|---|---|---|---|---|---|
| Tmem109_Q3UBX0 | -2 | 1.52 | 0.33 | 0.38 | 0.49 | -1 | -0.1 | -0.2 | 0.56 | Tmem109 |
| Hint1_P70349 | 1.27 | -0.5 | 1.37 | -1.4 | 0 | 0.14 | 0.42 | 0.12 | -1.5 | Hint1 |
| Ddx39b_Q9Z1N5 | -0.8 | 1.28 | 1.13 | 0.66 | -1.7 | 0.45 | -0.7 | -0.4 | 0.19 | Ddx39b |
| Tnks1bp1_P58871 | 0.58 | 0.89 | 0.01 | -0.7 | -1 | 1.1 | -0.3 | -1.7 | 1.09 | Tnks1bp1 |
| Crat_P47934 | 0.8 | -1.4 | -1.3 | -0.1 | 1.16 | -0 | -0.1 | 1.45 | -0.5 | Crat |
| Ubap2l_Q80X50 | -1.5 | 0.66 | 0.4 | 1.09 | -1.6 | 0.61 | -0.6 | 0.29 | 0.67 | Ubap2l |
| Snx1_Q9WV80 | -0.7 | -1.8 | 0.92 | -0.2 | -0.5 | 1.63 | 0.72 | 0.06 | -0.1 | Snx1 |
| Ahcy_P50247 | 0.36 | 1.07 | 0.35 | -1.9 | 1.37 | -0.2 | 0.31 | -0.5 | -0.8 | Ahcy |
| Nudc_O35685 | 0.36 | -0.4 | 0.81 | 0.6 | 1.15 | -2 | -0.5 | 0.17 | 0.05 | Nudc |
| Nrd1_Q8BHG1 | -0.2 | 0.47 | -0.9 | 1.51 | -1.9 | 0.83 | 0.11 | -0.4 | 0.49 | Nrd1 |
| Uso1_Q9Z1Z0 | 1.95 | -0.6 | -0.3 | 0.08 | 0.6 | -1 | 0.95 | -1 | -0.7 | Uso1 |
| Lipe_P54310 | 0.9 | -2 | 0.09 | -0.6 | 1.25 | -0.2 | 0.16 | 0.92 | -0.4 | Lipe |
| Psmd3_P14685 | -1.7 | 1.22 | -0.2 | 0.94 | -1 | 0.27 | -0.5 | -0.1 | 1.12 | Psmd3 |
| Itgb1_P09055 | 0.68 | 1.94 | -0.2 | -1.4 | 0.48 | -0.5 | 0.46 | -0.8 | -0.7 | Itgb1 |
| Nqo1_Q64669 | 1.86 | 0.54 | -0 | -1.3 | 0.15 | 0.05 | 0.72 | -0.8 | -1.2 | Nqo1 |
| Lta4h_P24527 | 1.54 | 0.83 | -0.4 | 0.33 | 0.64 | -1.8 | -0 | -0.2 | -0.9 | Lta4h |
| Thop1_Q8C1A5 | 1.63 | 0.17 | -0.1 | -1.8 | 0.8 | 0.31 | -1 | 0.37 | -0.4 | Thop1 |
| Lrp8_Q924X6 | 0.66 | -1.7 | -0.5 | 0.59 | 0.73 | -0.6 | 1.23 | -1 | 0.69 | Lrp8 |
| Uba6_Q8C7R4 | -1.2 | 1.81 | 0.82 | -1.3 | -0 | 0.74 | -0.4 | -0.4 | -0.1 | Uba6 |
| Cpt1a_P97742 | 1.54 | -1.5 | 0.06 | 0.25 | 1.16 | -1.3 | 0.17 | -0.2 | -0.1 | Cpt1a |
| P2rx4_Q9JJX6 | -1.8 | 0.21 | 0.17 | 1.21 | -1.2 | 0.83 | 0.61 | -0.6 | 0.56 | P2rx4 |
| Rps25_P62852 | 0.72 | 1.06 | -1 | 0.36 | -1.2 | 0.59 | 0.65 | 0.5 | -1.7 | Rps25 |
| Nampt_Q99KQ4 | 0.57 | -0.4 | -1.4 | 0.26 | 1.57 | -1.1 | 1.16 | 0.01 | -0.7 | Nampt |
| Stx4_P70452 | -1.9 | -0.8 | 0.01 | 1.64 | 0.09 | -0.3 | 0.25 | 0.01 | 0.99 | Stx4 |
| Eif5a2_Q8BGY2 | 1.74 | 0.02 | 0.65 | 0.4 | 0.3 | -1.8 | 0.18 | -0.8 | -0.7 | Eif5a2 |
| Csnk2b_P67871 | -0.5 | 1.26 | 1.12 | 0.36 | -1.6 | 0.41 | -1.4 | 0.21 | 0.12 | Csnk2b |
| Npc2_Q9Z0J0 | 1.53 | 0.09 | -0.3 | -1.3 | 1.5 | -0.7 | 0.4 | -0.2 | -0.9 | Npc2 |
| Sun1_Q9D666 | 0.49 | 0.57 | -2 | 0.45 | 0.12 | -0 | 0.14 | -0.6 | 1.21 | Sun1 |
| Vapb_Q9QY76 | 1.42 | -0.2 | -0.8 | -1.5 | 1.61 | -0.2 | 0.05 | 0.18 | -0.5 | Vapb |
| Srrm2_Q8BTI8 | -0.2 | 1.22 | 1.05 | -0.1 | 0.96 | -2 | -0.3 | 0.08 | -0.7 | Srrm2 |
| Ndufs6_P52503 | -0.7 | 0.5 | -1.4 | 0.54 | 1.36 | -1.2 | -0.4 | -0 | 1.28 | Ndufs6 |
| Serpinb1a_Q9D154 | 2 | 0.13 | -0.7 | -0.3 | -0.6 | 0.26 | 0.27 | 0.44 | -1.6 | Serpinb1a |
| Tm9sf3_Q9ET30 | -0.5 | 1.48 | 1.19 | 0.81 | -1.1 | -0.7 | 0.48 | -0.5 | -1.1 | Tm9sf3 |
| Kbp_Q6ZPU9 | -0.8 | 0.98 | -1.4 | -1.5 | 1.14 | 0.88 | 0.07 | 0.32 | 0.26 | Kbp |
| Apoa1bp_Q8K4Z3 | 0.69 | 0.73 | 1.5 | 0.61 | -1 | -1.1 | -1.4 | -0.3 | 0.26 | Apoa1bp |
| Nedd4_P46935 | 1.17 | 0.87 | 1.02 | 0.3 | -0.8 | -1 | -1.6 | 0.59 | -0.6 | Nedd4 |
| Aimp2_Q8R010 | 0.6 | 1.07 | 0.26 | -0.3 | -1.9 | 1.35 | -0.7 | -0.5 | 0.14 | Aimp2 |
| Snx2_Q9CWK8 | 0.18 | 0.88 | 1.7 | -1 | -1.2 | 0.9 | -0.1 | -0.9 | -0.4 | Snx2 |
| Gsto1_O09131 | -1.3 | 0.88 | 1.33 | 0.81 | -0.9 | -0.4 | -1.1 | -0.2 | 0.93 | Gsto1 |
| Cul4b_A2A432 | -1.8 | 1.15 | -0.5 | -0.4 | 0.15 | 0.9 | 0.75 | -1 | 0.78 | Cul4b |
| Cand2_Q6ZQ73 | -1 | 0.52 | 0.61 | 0.9 | -1.9 | 0.81 | -0.6 | -0.4 | 0.98 | Cand2 |
| St13_Q99L47 | 1.87 | 0.31 | -0.4 | -0.9 | 1.02 | -1 | 0.39 | -0.2 | -1.1 | St13 |
| Ppp2r5c_Q60996 | 0.52 | -0.3 | -1.4 | 0.27 | 1.69 | -1.3 | -0.6 | 0.62 | 0.53 | Ppp2r5c |
| Rai14_Q9EP71 | -1.7 | 0.81 | 0.67 | 0.36 | -0.7 | 0.57 | -1.4 | 0.59 | 0.84 | Rai14 |
| Tmod3_Q9JHJ0 | -0.4 | -0.7 | 1.11 | -1.8 | 1.15 | 0.61 | 0.05 | 0.74 | -0.7 | Tmod3 |
| Cd81_P35762 | -0.3 | 0.54 | -1.3 | 0.61 | 1.55 | -1.7 | -0.1 | 0.43 | 0.28 | Cd81 |
| Psmd5_Q8BJY1 | 1.17 | -1.4 | 0.67 | -0.5 | -1.3 | 1.46 | 0.24 | -0.4 | 0.04 | Psmd5 |
| Eva1b_Q8K2Y3 | -0 | -1.9 | -0.5 | 0.79 | -0.9 | 1.27 | 0.84 | -0.2 | 0.67 | Eva1b |
| Lypla1_P97823 | 0.18 | -2 | 0.66 | -0.1 | 0.71 | 0.17 | 0.3 | 1.07 | -0.7 | Lypla1 |
| Vac14_Q80WQ2 | 0.35 | -0.2 | 1.27 | -0.6 | -0.7 | 0.62 | 1.27 | -1.8 | -0.1 | Vac14 |

| | | | | | | | | | | |
|---|---|---|---|---|---|---|---|---|---|---|
| Stoml2_Q99JB2 | -0.7 | -0.3 | -0 | -0 | 0.93 | -0.5 | -1.8 | 1.72 | 0.58 | Stoml2 |
| Lman1_Q9D0F3 | -2 | 0.14 | 1.08 | 0.28 | -0.7 | 0.93 | -0.5 | 0.25 | 0.7 | Lman1 |
| H2-L_P01897 | 1.01 | -0.1 | -0.2 | -0.2 | -1.8 | 1.62 | -0.8 | -0.2 | 0.69 | H2-L |
| Pfdn2_O70591 | 0.54 | -1.2 | 1.3 | -0.9 | -0 | 0.62 | 1.05 | -1.5 | 0.1 | Pfdn2 |
| Ufm1_P61961 | -0.8 | -0.5 | -0.6 | -1 | -0.4 | 2 | 0.34 | 0.21 | 0.45 | Ufm1 |
| Vamp7_P70280 | 0.47 | -1 | 0.95 | 0.5 | 1.24 | -2 | 0.16 | -0.1 | -0.3 | Vamp7 |
| Nono_Q99K48 | -0.1 | 0.38 | 2 | -0.2 | -1.4 | 0.52 | 0.24 | -0.9 | -0.5 | Nono |
| Scamp2_Q9ERN0 | -1.7 | 0.56 | 0.76 | 1.01 | -1.1 | 0.31 | -0.7 | -0.3 | 1.16 | Scamp2 |
| Rpl21_O09167 | 0.72 | -0.8 | 0.19 | -0.4 | 1.1 | -0.8 | 1.76 | -0.7 | -1.1 | Rpl21 |
| Acat2_Q8CAY6 | 1.7 | 1.74 | -0.8 | -0.5 | -0.7 | -0.1 | -0.2 | -0.5 | -0.5 | Acat2 |
| Mccc2_Q3ULD5 | 1.38 | -1 | 0.6 | 0.14 | 0.34 | -1 | 1.37 | -1 | -0.8 | Mccc2 |
| Ptbp1_P17225 | -0.2 | 1.36 | 0.93 | 0.54 | -0.8 | -0.8 | -1.8 | 0.33 | 0.47 | Ptbp1 |
| Nme2_Q01768 | 1.79 | 0.61 | -1.3 | 0.11 | 0.56 | -1.2 | -0.6 | 0.51 | -0.4 | Nme2 |
| Prdx4_O08807 | 1.43 | -1 | -0.4 | -0.8 | 1.32 | -0.6 | 1.01 | 0 | -1.1 | Prdx4 |
| Trim28_Q62318 | 2 | -0.2 | -0.2 | -0.3 | 0.83 | -1.4 | 0.02 | -0.6 | -0.3 | Trim28 |
| Ndufv2_Q9D6J6 | -1.1 | 0.18 | -0.4 | 0.36 | -1.4 | 1.76 | -0.5 | 1.02 | 0.12 | Ndufv2 |
| Rpl22l1_Q9D7S7 | -0.8 | 0.46 | -0.8 | 1.4 | 0.99 | -1.8 | 0.1 | 0.57 | -0.1 | Rpl22l1 |
| Hsd17b8_P50171 | 1.56 | -0.4 | -1.8 | 0.38 | 0.95 | -1 | -0.2 | 0.3 | 0.18 | Hsd17b8 |
| Psmb1_O09061 | 1.79 | -1.3 | 0.63 | -0.3 | -0.2 | -0 | 1.07 | -0.9 | -0.7 | Psmb1 |
| Mrps36_Q9CQX8 | 0.93 | -0.6 | -0.1 | -1 | 1.27 | -0.5 | 1.29 | -1.5 | 0.13 | Mrps36 |
| Gnaq_P21279 | 1.64 | 1.29 | 0.77 | -0.7 | -0.4 | -0.7 | -1.3 | -0.1 | -0.5 | Gnaq |
| Fkbp5_Q64378 | -1.4 | 0.54 | 0.12 | 1.04 | -1.7 | 1.04 | -0.3 | -0.1 | 0.78 | Fkbp5 |
| Fau_P62862 | 0.54 | 0.54 | -0.6 | 1.41 | -0.1 | -1.5 | 0.26 | 0.88 | -1.4 | Fau |
| Mob1a_Q921Y0 | 0.29 | 1.4 | 1 | 0.86 | -0.6 | -1.6 | -0.5 | -1 | 0.17 | Mob1a |
| Ostf1_Q62422 | 0.11 | 0.57 | 1.74 | -1.7 | 0.53 | -0.1 | -0.9 | 0.44 | -0.8 | Ostf1 |
| Hagh_Q99KB8 | 0.06 | -1.8 | 1.04 | 0.86 | 0.86 | -1.3 | -0.3 | 0.28 | 0.34 | Hagh |
| Prkcsh_O08795 | 1.19 | -0.1 | 0.84 | -0.4 | 1.27 | -1.8 | -0.1 | 0.05 | -0.9 | Prkcsh |
| Nolc1_E9Q5C9 | -0.3 | 0.28 | -0.3 | -0.7 | -0.3 | 1.2 | 1.88 | -0.3 | -1.4 | Nolc1 |
| Rnmt_Q9D0L8 | -1.5 | 1.06 | 0.24 | -0.6 | -0.2 | 0.9 | 1.41 | -0.2 | -1.1 | Rnmt |
| Rplp1_P47955 | -0.1 | 1.18 | -1.6 | 0.7 | 0.84 | -1.3 | -0.7 | 0.1 | 0.84 | Rplp1 |
| Mmp2_P33434 | -1.6 | 0.38 | 0.79 | 1.12 | -1.6 | 0.66 | -0.5 | 0.03 | 0.63 | Mmp2 |
| Dcps_Q9DAR7 | -1.4 | 1.22 | 0.32 | 0.98 | -1.7 | 0.63 | -0.3 | -0.2 | 0.42 | Dcps |
| Dnaja1_P63037 | 1.12 | 0.63 | -1.6 | 0.29 | -0.5 | 0.1 | 1.06 | -1.5 | 0.33 | Dnaja1 |
| Blmh_Q8R016 | -1.6 | 1.08 | 0.96 | 0.86 | -1.4 | 0.34 | -0.5 | -0.1 | 0.43 | Blmh |
| Hnrnpm_Q9D0E1 | -1.5 | 0.54 | 0.81 | 1.15 | -1.5 | 0.45 | -0.7 | 0.2 | 0.6 | Hnrnpm |

| PROTEIN | UNIPROT | TPhP-Vehicle P.VALUE | TPhP-Vehicle LOGFC |
|---|---|---|---|
| Platelet glycoprotein 4 | Q08857 | 0.015176 | 0.411171 |
| Ribose-phosphate pyrophosphokinase 1;Ribose-phosphate pyro | Q9D7G0 | 0.003903 | 0.688735 |
| Cadherin-11 | P55288 | 0.005979 | 0.737173 |
| Sterol-4-alpha-carboxylate 3-dehydrogenase, decarboxylating | Q9R1J0 | 0.027774 | 0.623084 |
| Long-chain-fatty-acid--CoA ligase 1 | P41216 | 0.033833 | 0.414464 |
| Fatty acid-binding protein, adipocyte | P04117 | 0.052424 | 0.450378 |
| Fatty acid-binding protein, epidermal | Q05816 | 0.014628 | 0.423357 |
| Pyruvate carboxylase, mitochondrial | Q05920 | 0.020473 | 0.322759 |
| Glycerol-3-phosphate dehydrogenase [NAD(+)], cytoplasmic | P13707 | 0.091661 | 0.291175 |
| Mitochondrial pyruvate carrier 2 | Q9D023 | 0.002008 | 0.604222 |
| Inorganic pyrophosphatase | Q9D819 | 0.136211 | 0.168017 |
| Very long-chain specific acyl-CoA dehydrogenase, mitochondrial | P50544 | 0.20329 | 0.106723 |
| GrpE protein homolog 1, mitochondrial | Q99LP6 | 0.378411 | 0.104529 |
| Cytochrome b-c1 complex subunit 6, mitochondrial | P99028 | 0.083903 | 0.609172 |
| Adiponectin | Q60994 | 0.107914 | 0.317411 |
| Glutamine synthetase | P15105 | 0.004649 | 0.389196 |
| Ferrochelatase, mitochondrial | P22315 | 0.074284 | 0.278719 |
| Electron transfer flavoprotein subunit alpha, mitochondrial | Q99LC5 | 0.016417 | 0.234324 |
| Mitochondrial amidoxime reducing component 2 | Q922Q1 | 0.045372 | 0.230473 |
| Stress-70 protein, mitochondrial | P38647 | 0.138145 | 0.118048 |
| Citrate synthase, mitochondrial | Q9CZU6 | 0.10872 | 0.162149 |
| Isocitrate dehydrogenase [NAD] subunit alpha, mitochondrial | Q9D6R2 | 0.122533 | 0.165149 |
| Cytochrome c1, heme protein, mitochondrial | Q9D0M3 | 0.002661 | 0.418095 |
| ATP synthase subunit gamma, mitochondrial | Q91VR2 | 0.023574 | 0.175856 |
| 60 kDa heat shock protein, mitochondrial | P63038 | 0.046048 | 0.22656 |
| Farnesyl pyrophosphate synthase | Q920E5 | 0.049138 | 0.28096 |
| Medium-chain specific acyl-CoA dehydrogenase, mitochondrial | P45952 | 0.06267 | 0.240035 |
| Malate dehydrogenase, mitochondrial | P08249 | 0.065223 | 0.209148 |
| Bifunctional epoxide hydrolase 2;Cytosolic epoxide hydrolase 2;I | P34914 | 0.507896 | 0.064215 |
| Very-long-chain 3-oxoacyl-CoA reductase | O70503 | 0.69191 | 0.038999 |
| Thioredoxin-dependent peroxide reductase, mitochondrial | P20108 | 0.025407 | 0.231556 |
| Prohibitin | P67778 | 0.036904 | 0.207661 |
| Trifunctional enzyme subunit beta, mitochondrial;3-ketoacyl-Co | Q99JY0 | 0.044546 | 0.214476 |
| Microsomal glutathione S-transferase 1 | Q91VS7 | 0.101808 | 0.364235 |
| DnaJ homolog subfamily A member 4 | Q9JMC3 | 0.108377 | 0.466275 |
| Ubiquinone biosynthesis protein COQ9, mitochondrial | Q8K1Z0 | 0.574128 | 0.05368 |
| Titin | A2ASS6 | 0.498889 | -0.06817 |
| Acyl-CoA-binding protein | P31786 | 0.116743 | 0.234685 |
| Monocarboxylate transporter 1 | P53986 | 0.036309 | -0.30322 |
| Putative adenosylhomocysteinase 3;Putative adenosylhomocyst | Q68FL4 | 0.536351 | 0.077502 |
| Fibronectin;Anastellin | P11276 | 0.56228 | -0.05288 |
| Tricarboxylate transport protein, mitochondrial | Q8JZU2 | 0.624012 | 0.067916 |

| | | | |
|---|---|---|---|
| 2,4-dienoyl-CoA reductase, mitochondrial | Q9CQ62 | 0.025373 | 0.31177 |
| Mitochondrial dicarboxylate carrier | Q9QZD8 | 0.277011 | 0.160198 |
| B-cell receptor-associated protein 31 | Q61335 | 0.034798 | 0.214459 |
| Niban-like protein 1 | Q8R1F1 | 0.03893 | -0.25244 |
| Perilipin-1 | Q8CGN5 | 0.011598 | 0.406772 |
| Peroxiredoxin-1 | P35700 | 0.046589 | 0.144835 |
| ATP synthase subunit alpha, mitochondrial | Q03265 | 0.07404 | 0.142492 |
| Bcl-2-like protein 13 | P59017 | 0.101902 | 0.324215 |
| Forkhead box protein N4 | Q8K3Q3 | 0.575476 | -0.06177 |
| Peptidyl-prolyl cis-trans isomerase C | P30412 | 0.265079 | -0.09856 |
| Host cell factor 1;HCF N-terminal chain 1;HCF N-terminal chain 2 | Q61191 | 0.110326 | 0.221681 |
| Perilipin-4 | O88492 | 0.84723 | -0.02118 |
| Dynamin-like 120 kDa protein, mitochondrial;Dynamin-like 120 | P58281 | 0.01632 | 0.437367 |
| Caprin-1 | Q60865 | 0.062893 | -0.21411 |
| Delta-1-pyrroline-5-carboxylate synthase;Glutamate 5-kinase;Ga | Q9Z110 | 0.1927 | -0.17059 |
| Fructose-bisphosphate aldolase A | P05064 | 0.21007 | 0.151765 |
| Alpha-actinin-1 | Q7TPR4 | 0.259609 | -0.09935 |
| Antithrombin-III | P32261 | 0.040755 | -0.21216 |
| 5-AMP-activated protein kinase subunit gamma-1 | O54950 | 0.042442 | 0.272931 |
| LETM1 and EF-hand domain-containing protein 1, mitochondrial | Q9Z2I0 | 0.047261 | 0.350932 |
| Dihydrolipoyl dehydrogenase, mitochondrial | O08749 | 0.056175 | 0.176683 |
| Long-chain specific acyl-CoA dehydrogenase, mitochondrial | P51174 | 0.062199 | 0.213272 |
| Acetyl-CoA acetyltransferase, mitochondrial | Q8QZT1 | 0.083133 | 0.129276 |
| ADP/ATP translocase 1 | P48962 | 0.156058 | 0.130982 |
| Transaldolase | Q93092 | 0.327432 | 0.108697 |
| Delta-aminolevulinic acid dehydratase | P10518 | 0.051455 | 0.294288 |
| Tax1-binding protein 3 | Q9DBG9 | 0.514308 | 0.090606 |
| Peptidyl-prolyl cis-trans isomerase-like 1 | Q9D0W5 | 0.049411 | -0.43928 |
| Pyruvate dehydrogenase protein X component, mitochondrial | Q8BKZ9 | 0.071124 | 0.211572 |
| NAD(P) transhydrogenase, mitochondrial | Q61941 | 0.279275 | -0.08917 |
| Mitochondrial import inner membrane translocase subunit TIM4 | O35857 | 0.320628 | 0.112029 |
| VPS10 domain-containing receptor SorCS2 | Q9EPR5 | 0.72099 | 0.055688 |
| Prolyl 4-hydroxylase subunit alpha-2 | Q60716 | 0.904673 | 0.012942 |
| 40S ribosomal protein S24 | P62849 | 0.17526 | -0.17538 |
| Calponin-2 | Q08093 | 0.399156 | 0.068913 |
| Myosin light chain 1/3, skeletal muscle isoform;Myosin light cha | P05977 | 0.658612 | 0.051043 |
| Calreticulin | P14211 | 0.070887 | 0.178299 |
| Lipoxygenase homology domain-containing protein 1 | C8YR32 | 0.347815 | 0.183449 |
| Branched-chain-amino-acid aminotransferase, mitochondrial | O35855 | 0.012252 | 0.658241 |
| C-type mannose receptor 2 | Q64449 | 0.280147 | -0.11128 |
| Lysosomal alpha-mannosidase | O09159 | 0.075654 | -0.14898 |
| OTU domain-containing protein 6A | Q6IE21 | 0.042438 | -0.37466 |
| Carnitine O-palmitoyltransferase 2, mitochondrial | P52825 | 0.296075 | 0.112772 |
| MICOS complex subunit Mic60 | Q8CAQ8 | 0.138775 | 0.205414 |
| ATP synthase subunit delta, mitochondrial | Q9D3D9 | 0.160648 | 0.185005 |
| Serine/threonine-protein kinase OSR1 | Q6P9R2 | 0.235589 | -0.12804 |
| Lipoprotein lipase | P11152 | 0.508748 | -0.08292 |
| Dihydropyrimidinase-related protein 3 | Q62188 | 0.599422 | -0.04723 |
| Isocitrate dehydrogenase [NAD] subunit gamma 1, mitochondria | P70404 | 0.799106 | 0.029389 |
| Lamina-associated polypeptide 2, isoforms beta/delta/epsilon/g | Q61029 | 0.050111 | -0.20397 |

| | | | |
|---|---|---|---|
| Hydroxyacyl-coenzyme A dehydrogenase, mitochondrial | Q61425 | 0.075476 | 0.202052 |
| S-methyl-5-thioadenosine phosphorylase | Q9CQ65 | 0.081046 | -0.1669 |
| Succinyl-CoA ligase [ADP/GDP-forming] subunit alpha, mitochon | Q9WUM5 | 0.360958 | 0.087229 |
| Myristoylated alanine-rich C-kinase substrate | P26645 | 0.370561 | 0.074711 |
| Protein disulfide-isomerase A4 | P08003 | 0.109656 | 0.114594 |
| LIM and SH3 domain protein 1 | Q61792 | 0.231737 | -0.10178 |
| Filamin-A | Q8BTM8 | 0.845538 | 0.014341 |
| ADP-ribose pyrophosphatase, mitochondrial;Transient receptor | Q8BVU5 | 0.955332 | 0.006913 |
| Sideroflexin-1 | Q99JR1 | 0.289142 | -0.11571 |
| Pyruvate dehydrogenase E1 component subunit alpha, somatic | P35486 | 0.90663 | -0.00868 |
| Metalloproteinase inhibitor 2 | P25785 | 0.10467 | -0.23158 |
| 78 kDa glucose-regulated protein | P20029 | 0.557475 | 0.040963 |
| Cartilage-associated protein | Q9CYD3 | 0.049152 | -0.18069 |
| Cytochrome b5 type B | Q9CQX2 | 0.051214 | 0.191788 |
| Lipoma-preferred partner homolog | Q8BFW7 | 0.08168 | -0.19242 |
| ADP/ATP translocase 2;ADP/ATP translocase 2, N-terminally pro | P51881 | 0.297001 | 0.138281 |
| Proteasome subunit beta type-6 | Q60692 | 0.320557 | 0.122637 |
| Tripeptidyl-peptidase 1 | O89023 | 0.538719 | -0.05476 |
| GTPase KRas;GTPase KRas, N-terminally processed;GTPase HRas | P32883 | 0.651375 | -0.06733 |
| Tropomyosin alpha-1 chain | P58771 | 0.922997 | -0.01071 |
| Actin, cytoplasmic 2;Actin, cytoplasmic 2, N-terminally processe | P63260 | 0.118562 | -0.46443 |
| Nodal modulator 1 | Q6GQT9 | 0.149016 | 0.335048 |
| Epoxide hydrolase 1 | Q9D379 | 0.044337 | 0.154995 |
| 116 kDa U5 small nuclear ribonucleoprotein component | O08810 | 0.097575 | -0.15147 |
| Myoferlin | Q69ZN7 | 0.601872 | -0.04933 |
| Tubulin alpha-1B chain;Tubulin alpha-4A chain | P05213 | 0.210083 | 0.13188 |
| Microsomal glutathione S-transferase 3 | Q9CPU4 | 0.740724 | 0.084007 |
| Aldehyde dehydrogenase, dimeric NADP-preferring | P47739 | 0.383702 | -0.10702 |
| Fatty acid synthase;[Acyl-carrier-protein] S-acetyltransferase;[A | P19096 | 0.445295 | 0.080944 |
| Casein kinase II subunit alpha | Q60737 | 0.966022 | -0.00433 |
| Translocator protein | P50637 | 0.03206 | 0.482953 |
| cAMP-dependent protein kinase type II-beta regulatory subunit | P31324 | 0.194696 | 0.146175 |
| Protein S100-A6 | P14069 | 0.28928 | 0.317548 |
| Argininosuccinate synthase | P16460 | 0.459448 | -0.12041 |
| Hypoxanthine-guanine phosphoribosyltransferase | P00493 | 0.703972 | 0.081318 |
| Insulin-like growth factor-binding protein 4 | P47879 | 0.001335 | -0.67718 |
| Tropomyosin alpha-3 chain | P21107 | 0.097978 | -0.1362 |
| 26S proteasome non-ATPase regulatory subunit 14 | O35593 | 0.481826 | 0.061991 |
| Zyxin | Q62523 | 0.208193 | 0.129864 |
| Pyruvate dehydrogenase E1 component subunit beta, mitochon | Q9D051 | 0.224337 | 0.195799 |
| 3-hydroxyisobutyrate dehydrogenase, mitochondrial | Q99L13 | 0.387718 | 0.123863 |
| 60S ribosomal protein L27a | P14115 | 0.060611 | -0.29292 |
| Fumarate hydratase, mitochondrial | P97807 | 0.287885 | 0.085688 |
| Annexin A1 | P10107 | 0.871238 | -0.01986 |
| 4-trimethylaminobutyraldehyde dehydrogenase | Q9JLJ2 | 0.54799 | 0.040346 |
| Vacuolar protein sorting-associated protein VTA1 homolog | Q9CR26 | 0.31221 | -0.25324 |
| Catalase | P24270 | 0.429115 | 0.069525 |
| L-lactate dehydrogenase B chain | P16125 | 0.67895 | 0.070868 |
| Alcohol dehydrogenase class 4 mu/sigma chain | Q64437 | 0.698481 | 0.029105 |
| Rho-related GTP-binding protein RhoC | Q62159 | 0.881259 | 0.01084 |

| | | | |
|---|---|---|---|
| ATP synthase subunit beta, mitochondrial | P56480 | 0.203383 | 0.101701 |
| Dynactin subunit 2 | Q99KJ8 | 0.704843 | -0.04614 |
| Actin, alpha cardiac muscle 1;Actin, aortic smooth muscle;Actin, | P68033 | 0.945215 | 0.007678 |
| Sorting nexin-3 | O70492 | 0.034724 | -0.23075 |
| Cystatin-B | Q62426 | 0.383317 | -0.07577 |
| Collagen alpha-1(I) chain | P11087 | 0.560611 | -0.04543 |
| ATP synthase subunit e, mitochondrial | Q06185 | 0.213378 | 0.165353 |
| Eukaryotic translation initiation factor 4E | P63073 | 0.01741 | 0.260036 |
| Biglycan | P28653 | 0.076986 | -0.15871 |
| Cell division control protein 42 homolog | P60766 | 0.191659 | -0.10201 |
| Lysophospholipid acyltransferase 5 | Q91V01 | 0.108263 | 0.305876 |
| RNA-binding motif protein, X-linked 2 | Q8R0F5 | 0.483424 | -0.10358 |
| Glycogen phosphorylase, brain form | Q8CI94 | 0.253692 | -0.08144 |
| Protein SEC13 homolog | Q9D1M0 | 0.004459 | 0.345537 |
| Thy-1 membrane glycoprotein | P01831 | 0.092351 | -0.14651 |
| Elongation factor Tu, mitochondrial | Q8BFR5 | 0.153434 | 0.117716 |
| WD repeat-containing protein 1 | O88342 | 0.283946 | -0.09201 |
| Phosphate carrier protein, mitochondrial | Q8VEM8 | 0.156279 | 0.202997 |
| Transgelin | P37804 | 0.592905 | -0.07389 |
| Prostaglandin E synthase 3 | Q9R0Q7 | 0.016353 | 0.349359 |
| Sorting nexin-5 | Q9D8U8 | 0.040925 | -0.19377 |
| Plexin-B2 | B2RXS4 | 0.955075 | -0.00644 |
| Pyridoxal kinase | Q8K183 | 0.249008 | -0.10784 |
| Dihydrolipoyllysine-residue succinyltransferase component of 2- | Q9D2G2 | 0.309493 | 0.323337 |
| Phosphomannomutase 2 | Q9Z2M7 | 0.435473 | -0.11549 |
| Src substrate cortactin | Q60598 | 0.45086 | -0.05553 |
| 40S ribosomal protein S10 | P63325 | 0.45102 | 0.0726 |
| Vacuolar protein sorting-associated protein 35 | Q9EQH3 | 0.233732 | -0.11013 |
| Filamin-B | Q80X90 | 0.48781 | -0.07195 |
| Myeloid-associated differentiation marker | O35682 | 0.007161 | 6.364119 |
| ATP synthase F(0) complex subunit B1, mitochondrial | Q9CQQ7 | 0.501305 | 0.082163 |
| Glutaredoxin-related protein 5, mitochondrial | Q80Y14 | 0.966084 | 0.007189 |
| Calcium uniporter protein, mitochondrial | Q3UMR5 | 0.10102 | -0.25858 |
| Macrophage-capping protein | P24452 | 0.561649 | -0.09102 |
| Endoplasmic reticulum aminopeptidase 1 | Q9EQH2 | 0.648704 | -0.80664 |
| Calcyclin-binding protein | Q9CXW3 | 0.925907 | 0.013895 |
| Thioredoxin-like protein 1 | Q8CDN6 | 0.173 | -0.13813 |
| ATP-dependent RNA helicase A | O70133 | 0.061459 | -0.14304 |
| Superoxide dismutase [Cu-Zn] | P08228 | 0.537858 | 0.075108 |
| Rho GTPase-activating protein 1 | Q5FWK3 | 0.038866 | -0.30692 |
| Guanine nucleotide-binding protein subunit alpha-13 | P27601 | 0.587268 | 0.086518 |
| Filamin-C | Q8VHX6 | 0.979609 | -0.00247 |
| Voltage-dependent anion-selective channel protein 3 | Q60931 | 0.199474 | 0.09957 |
| Non-specific lipid-transfer protein | P32020 | 0.341155 | 0.208566 |
| Kinesin-1 heavy chain;Kinesin heavy chain isoform 5C;Kinesin he | Q61768 | 0.666586 | 0.049895 |
| Carbonyl reductase [NADPH] 2 | P08074 | 0.125566 | 0.152708 |
| AP-2 complex subunit beta | Q9DBG3 | 0.320772 | -0.07275 |
| Band 4.1-like protein 2 | O70318 | 0.609897 | -0.03623 |
| Myosin-10 | Q61879 | 0.672584 | -0.02713 |
| PDZ and LIM domain protein 5 | Q8CI51 | 0.86147 | -0.01601 |

| | | | |
|---|---|---|---|
| COP9 signalosome complex subunit 8 | Q8VBV7 | 0.040474 | 0.644188 |
| Fibulin-2 | P37889 | 0.442667 | -0.12236 |
| Dihydrolipoyllysine-residue acetyltransferase component of pyr | Q8BMF4 | 0.707938 | -0.03104 |
| Protein disulfide-isomerase | P09103 | 0.097145 | 0.116443 |
| F-actin-capping protein subunit beta | P47757 | 0.499805 | 0.070323 |
| Peptidyl-prolyl cis-trans isomerase FKBP10 | Q61576 | 0.064361 | 0.257235 |
| Equilibrative nucleoside transporter 1 | Q9JIM1 | 0.134745 | -0.18111 |
| Catenin alpha-1 | P26231 | 0.125543 | 0.118994 |
| Ezrin | P26040 | 0.55199 | -0.05109 |
| RNA-binding protein 3 | O89086 | 0.098882 | -0.1702 |
| Spectrin beta chain, non-erythrocytic 1 | Q62261 | 0.770983 | 0.02361 |
| Cytochrome b-c1 complex subunit 1, mitochondrial | Q9CZ13 | 0.2019 | 0.122334 |
| U1 small nuclear ribonucleoprotein A | Q62189 | 0.296315 | -0.18468 |
| Endosialin | Q91V98 | 0.462926 | 0.103085 |
| Coatomer subunit gamma-1 | Q9QZE5 | 0.558169 | -0.07757 |
| Large proline-rich protein BAG6 | Q9Z1R2 | 0.662963 | -0.13798 |
| Ornithine aminotransferase, mitochondrial | P29758 | 0.746843 | -0.03656 |
| Mitochondrial intermembrane space import and assembly prote | Q8VEA4 | 0.791168 | 0.104369 |
| Myb-binding protein 1A | Q7TPV4 | 0.979927 | 0.003193 |
| LIM domain and actin-binding protein 1 | Q9ERG0 | 0.399805 | -0.08598 |
| Matrix metalloproteinase-14 | P53690 | 0.879683 | 0.010531 |
| Early endosome antigen 1 | Q8BL66 | 0.262639 | -0.10012 |
| Mitochondrial import inner membrane translocase subunit Tim2 | Q9WTQ8 | 0.016468 | 6.264415 |
| Sideroflexin-3 | Q91V61 | 0.273386 | -0.16053 |
| F-actin-capping protein subunit alpha-1 | P47753 | 0.312955 | -0.11279 |
| Procollagen-lysine,2-oxoglutarate 5-dioxygenase 3 | Q9R0E1 | 0.584402 | -0.0635 |
| Vascular cell adhesion protein 1 | P29533 | 0.400286 | 0.085864 |
| Prostaglandin E synthase 2;Prostaglandin E synthase 2 truncatec | Q8BWM0 | 0.017638 | 5.672039 |
| Coiled-coil-helix-coiled-coil-helix domain-containing protein 2 | Q9D1L0 | 0.026869 | 0.271093 |
| Mitochondrial fission 1 protein | Q9CQ92 | 0.232353 | 0.119306 |
| Rho GDP-dissociation inhibitor 1 | Q99PT1 | 0.576934 | -0.06899 |
| Septin-7 | O55131 | 0.960886 | -0.00384 |
| Coatomer subunit gamma-2 | Q9QXK3 | 0.114434 | -0.47424 |
| Small acidic protein | Q9R0P4 | 0.499156 | 0.146397 |
| Peroxiredoxin-5, mitochondrial | P99029 | 0.191842 | 0.164081 |
| Actin-related protein 2/3 complex subunit 5 | Q9CPW4 | 0.328751 | -0.19052 |
| Ras GTPase-activating protein-binding protein 2 | P97379 | 0.229971 | 0.215398 |
| CAD protein;Glutamine-dependent carbamoyl-phosphate synth | B2RQC6 | 0.84275 | 0.404887 |
| Gelsolin | P13020 | 0.725378 | -0.02658 |
| GTP-binding nuclear protein Ran;GTP-binding nuclear protein Ra | P62827 | 0.761826 | -0.02868 |
| Calpain-6 | O35646 | 0.101292 | -0.18528 |
| 3-ketoacyl-CoA thiolase, mitochondrial | Q8BWT1 | 0.215972 | 0.134401 |
| Prolyl endopeptidase | Q9QUR6 | 0.075902 | 0.200611 |
| ATP synthase-coupling factor 6, mitochondrial | P97450 | 0.568125 | -0.06743 |
| Sodium/potassium-transporting ATPase subunit alpha-1 | Q8VDN2 | 0.379217 | -0.06377 |
| EH domain-containing protein 4 | Q9EQP2 | 0.004368 | -0.47056 |
| Transcription elongation factor B polypeptide 2 | P62869 | 0.335894 | 0.17935 |
| Myosin light polypeptide 6 | Q60605 | 0.578337 | 0.069196 |
| Tubulin beta-5 chain | P99024 | 0.984321 | -0.002 |
| Serine/threonine-protein kinase PAK 2;PAK-2p27;PAK-2p34 | Q8CIN4 | 0.265338 | -0.22833 |

| | | | |
|---|---|---|---|
| Phosphatidylinositol 5-phosphate 4-kinase type-2 gamma | Q91XU3 | 0.060897 | -0.23481 |
| NADP-dependent malic enzyme | P06801 | 0.165744 | 0.581869 |
| von Willebrand factor A domain-containing protein 5A | Q99KC8 | 0.485815 | 0.064416 |
| cAMP-dependent protein kinase type I-alpha regulatory subunit | Q9DBC7 | 0.561668 | 0.217066 |
| Beta-hexosaminidase subunit beta | P20060 | 0.978371 | -0.0039 |
| High mobility group protein B2 | P30681 | 0.03595 | -0.44313 |
| Nicotinamide N-methyltransferase | O55239 | 0.198363 | 0.140011 |
| ATP synthase subunit O, mitochondrial | Q9DB20 | 0.170935 | 0.205342 |
| Carbonyl reductase [NADPH] 3 | Q8K354 | 0.260017 | 0.164545 |
| Heterogeneous nuclear ribonucleoprotein F;Heterogeneous nuc | Q9Z2X1 | 0.302314 | -0.10231 |
| 60S ribosomal protein L24 | Q8BP67 | 0.426821 | 0.068713 |
| DnaJ homolog subfamily A member 2 | Q9QYJ0 | 0.007006 | -0.31075 |
| Acetoacetyl-CoA synthetase | Q9D2R0 | 0.038716 | 0.703355 |
| U6 snRNA-associated Sm-like protein LSm4 | Q9QXA5 | 0.096889 | -0.18526 |
| Vasodilator-stimulated phosphoprotein | P70460 | 0.001035 | 0.318067 |
| Dipeptidase 1 | P31428 | 0.919864 | 0.012715 |
| Reticulocalbin-3 | Q8BH97 | 0.889772 | -0.02075 |
| 14-3-3 protein beta/alpha;14-3-3 protein beta/alpha, N-termina | Q9CQV8 | 0.222394 | 0.133092 |
| Spectrin alpha chain, non-erythrocytic 1 | P16546 | 0.458614 | 0.053458 |
| Serine hydroxymethyltransferase | Q9CZN7 | 0.023028 | 0.181972 |
| Transitional endoplasmic reticulum ATPase | Q01853 | 0.111364 | 0.156584 |
| Adipocyte plasma membrane-associated protein | Q9D7N9 | 0.137468 | 0.247347 |
| Histone H1.2 | P15864 | 0.817117 | -0.01921 |
| 60S ribosomal protein L18a | P62717 | 0.862962 | 0.019538 |
| Acetyl-coenzyme A synthetase, cytoplasmic | Q9QXG4 | 0.030188 | 5.411604 |
| Leucine-rich repeat-containing protein 59 | Q922Q8 | 0.598968 | 0.13813 |
| Short/branched chain specific acyl-CoA dehydrogenase, mitoch | Q9DBL1 | 0.026992 | 0.436688 |
| Mitogen-activated protein kinase 1 | P63085 | 0.514167 | -0.10554 |
| Eukaryotic translation initiation factor 3 subunit K | Q9DBZ5 | 0.014125 | -0.59899 |
| ATP-citrate synthase | Q91V92 | 0.149971 | 0.197653 |
| Drebrin | Q9QXS6 | 0.454349 | -0.13656 |
| NADH-cytochrome b5 reductase 3;NADH-cytochrome b5 reduct | Q9DCN2 | 0.984823 | -0.00164 |
| Heat shock 70 kDa protein 1A;Heat shock 70 kDa protein 1B | Q61696 | 0.271786 | 0.116597 |
| Small nuclear ribonucleoprotein-associated protein B;Small nucl | P27048 | 0.350029 | 0.084445 |
| Inorganic pyrophosphatase 2, mitochondrial | Q91VM9 | 0.37816 | 0.112161 |
| 40S ribosomal protein S16 | P14131 | 0.750865 | -0.02274 |
| ADP-ribosylation factor 6 | P62331 | 0.437838 | 0.071344 |
| Cathepsin B;Cathepsin B light chain;Cathepsin B heavy chain | P10605 | 0.412646 | -0.10794 |
| Serine/arginine-rich splicing factor 2 | Q62093 | 0.142709 | -0.13586 |
| Ras-related protein Rab-22A | P35285 | 0.052908 | -0.32298 |
| Collagen alpha-1(VIII) chain;Vastatin | Q00780 | 0.763714 | -0.03616 |
| Alpha-actinin-4 | P57780 | 0.935341 | 0.005183 |
| Cytochrome c oxidase subunit 6C | Q9CPQ1 | 0.866379 | -0.01728 |
| Citrate lyase subunit beta-like protein, mitochondrial | Q8R4N0 | 0.324392 | 0.17731 |
| 1,2-dihydroxy-3-keto-5-methylthiopentene dioxygenase | Q99JT9 | 0.664342 | -0.05265 |
| Phenylalanine--tRNA ligase alpha subunit | Q8C0C7 | 0.717266 | 0.065455 |
| Histone H1.5 | P43276 | 0.750633 | 0.034069 |
| Granulins;Acrogranin;Granulin-1;Granulin-2;Granulin-3;Granulin | P28798 | 0.937141 | -0.01839 |
| Extended synaptotagmin-1 | Q3U7R1 | 0.748413 | -0.03273 |
| Versican core protein | Q62059 | 0.085406 | -0.3779 |

| | | | |
|---|---|---|---|
| Cytochrome c oxidase subunit 2 | P00405 | 0.175587 | 0.25838 |
| Transportin-1;Transportin-2 | Q8BFY9 | 0.097944 | -0.28522 |
| Glutathione S-transferase Mu 2 | P15626 | 0.723946 | -0.04939 |
| Plastin-3 | Q99K51 | 0.774823 | -0.02406 |
| AP-3 complex subunit delta-1 | O54774 | 0.003496 | 0.389243 |
| Methionine--tRNA ligase, cytoplasmic | Q68FL6 | 0.037325 | -0.33952 |
| Cytochrome c oxidase subunit NDUFA4 | Q62425 | 0.127291 | 0.134852 |
| Aflatoxin B1 aldehyde reductase member 2 | Q8CG76 | 0.128519 | 0.260985 |
| Probable ATP-dependent RNA helicase DDX6 | P54823 | 0.166868 | 0.223434 |
| Perilipin-3 | Q9DBG5 | 0.813867 | -0.01651 |
| Heat shock cognate 71 kDa protein | P63017 | 0.262729 | 0.097245 |
| CCA tRNA nucleotidyltransferase 1, mitochondrial | Q8K1J6 | 0.616305 | 0.048353 |
| Sorting nexin-6;Sorting nexin-6, N-terminally processed | Q6P8X1 | 0.274317 | -0.13551 |
| Calnexin | P35564 | 0.176598 | 0.198273 |
| Dihydroorotate dehydrogenase (quinone), mitochondrial | O35435 | 0.030565 | 5.784747 |
| Leucine-rich repeat-containing protein 47 | Q505F5 | 0.103354 | 0.31892 |
| Ribosome-binding protein 1 | Q99PL5 | 0.821431 | -0.01771 |
| Enoyl-CoA delta isomerase 1, mitochondrial | P42125 | 0.054664 | 0.202709 |
| Dehydrogenase/reductase SDR family member 4 | Q99LB2 | 0.039598 | 0.376128 |
| Sulfide:quinone oxidoreductase, mitochondrial | Q9R112 | 0.175759 | 0.168881 |
| Cytosolic non-specific dipeptidase | Q9D1A2 | 0.738353 | -0.03423 |
| Alanine--tRNA ligase, cytoplasmic | Q8BGQ7 | 0.390073 | -0.11501 |
| Profilin-2 | Q9JJV2 | 0.773112 | 0.043391 |
| Succinate dehydrogenase [ubiquinone] flavoprotein subunit, mi | Q8K2B3 | 0.0429 | 0.188294 |
| Retinoid-inducible serine carboxypeptidase | Q920A5 | 0.49011 | 0.052845 |
| Splicing factor 3A subunit 3 | Q9D554 | 0.610118 | 0.066751 |
| RNA-binding motif, single-stranded-interacting protein 1;RNA-bi | Q91W59 | 0.778881 | 0.040194 |
| Succinyl-CoA ligase [GDP-forming] subunit beta, mitochondrial | Q9Z2I8 | 0.093417 | 0.203114 |
| Talin-1 | P26039 | 0.941881 | 0.005062 |
| Elongation factor 1-gamma | Q9D8N0 | 0.075849 | 0.129416 |
| NADH dehydrogenase [ubiquinone] 1 subunit C2 | Q9CQ54 | 0.111795 | 0.346592 |
| Acylamino-acid-releasing enzyme | Q8R146 | 0.40945 | -0.12438 |
| AP2-associated protein kinase 1 | Q3UHJ0 | 0.625777 | -0.04861 |
| Galactokinase | Q9R0N0 | 0.097209 | -0.29011 |
| Serine/arginine-rich splicing factor 3 | P84104 | 0.174045 | -0.19607 |
| Mesencephalic astrocyte-derived neurotrophic factor | Q9CXI5 | 0.179999 | -0.22387 |
| 40S ribosomal protein S19 | Q9CZX8 | 0.215082 | 0.133045 |
| Endophilin-A2 | Q62419 | 0.255742 | -0.11069 |
| Isovaleryl-CoA dehydrogenase, mitochondrial | Q9JHI5 | 0.303796 | 0.089319 |
| Interleukin enhancer-binding factor 3 | Q9Z1X4 | 0.583552 | -0.0517 |
| Periostin | Q62009 | 0.852143 | -0.02007 |
| Glucose-6-phosphate isomerase | P06745 | 0.908191 | 0.009702 |
| Gamma-interferon-inducible lysosomal thiol reductase | Q9ESY9 | 0.959834 | -0.01043 |
| Phosphoribosyl pyrophosphate synthase-associated protein 1 | Q9D0M1 | 0.041164 | -0.32845 |
| 26S proteasome non-ATPase regulatory subunit 12 | Q9D8W5 | 0.630311 | 0.089934 |
| 40S ribosomal protein S29 | P62274 | 0.02925 | 4.927409 |
| Ubiquitin-conjugating enzyme E2 N | P61089 | 0.355409 | 0.065845 |
| Synaptonemal complex protein SC65 | Q8K2B0 | 0.462259 | -0.21354 |
| Rho GDP-dissociation inhibitor 2 | Q61599 | 0.959018 | -0.00569 |
| Pleiotrophin | P63089 | 0.171482 | 0.184219 |

| | | | |
|---|---|---|---|
| Amine oxidase [flavin-containing] B | Q8BW75 | 0.328586 | 0.107017 |
| Methylmalonate-semialdehyde dehydrogenase [acylating], mitc | Q9EQ20 | 0.330122 | 0.138443 |
| Mitochondrial import inner membrane translocase subunit Tim1 | P62075 | 0.452692 | 0.212598 |
| Density-regulated protein | Q9CQJ6 | 0.818638 | 0.028686 |
| Transmembrane protein 43 | Q9DBS1 | 0.930248 | -0.00678 |
| Syntaxin-7 | O70439 | 0.942218 | -0.01305 |
| NADH dehydrogenase [ubiquinone] iron-sulfur protein 8, mitocl | Q8K3J1 | 0.09827 | 0.398812 |
| Cytochrome c oxidase subunit 7A2, mitochondrial | P48771 | 0.426554 | 0.192661 |
| | C0HKD9 | 0.021827 | 5.539847 |
| Basigin | P18572 | 0.035806 | 0.238233 |
| UDP-N-acetylhexosamine pyrophosphorylase-like protein 1 | Q3TW96 | 0.187947 | -0.10818 |
| Arylsulfatase B | P50429 | 0.592587 | -0.04503 |
| Lysosome-associated membrane glycoprotein 2 | P17047 | 0.596089 | 0.050418 |
| NSFL1 cofactor p47 | Q9CZ44 | 0.637695 | -0.0619 |
| Septin-11 | Q8C1B7 | 0.941913 | 0.008092 |
| Myosin regulatory light chain 12B;Myosin regulatory light polyp | Q3THE2 | 0.280581 | 0.161227 |
| Actin, cytoplasmic 1;Actin, cytoplasmic 1, N-terminally processe | P60710 | 0.839889 | 0.022329 |
| Malate dehydrogenase, cytoplasmic | P14152 | 0.653134 | 0.049191 |
| Paralemmin-1 | Q9Z0P4 | 0.725094 | 0.104254 |
| Lamin-B1 | P14733 | 0.894372 | 0.013418 |
| Serine/threonine-protein phosphatase 2B catalytic subunit alph | P63328 | 0.40754 | -0.16994 |
| Platelet-activating factor acetylhydrolase IB subunit gamma | Q61205 | 0.822939 | 0.021734 |
| Mitochondrial import receptor subunit TOM70 | Q9CZW5 | 0.285077 | 0.114805 |
| Cytochrome c oxidase subunit 4 isoform 1, mitochondrial | P19783 | 0.559165 | 0.08746 |
| Phosphatidylethanolamine-binding protein 1;Hippocampal choli | P70296 | 0.878249 | -0.01303 |
| Ferritin light chain 1;Ferritin light chain 2 | P29391 | 0.294636 | -0.17797 |
| Receptor expression-enhancing protein 5 | Q60870 | 0.567016 | 0.145183 |
| Golgi resident protein GCP60 | Q8BMP6 | 0.031445 | -0.32923 |
| Small nuclear ribonucleoprotein E | P62305 | 0.03551 | -0.31308 |
| Cytoplasmic dynein 1 light intermediate chain 2 | Q6PDL0 | 0.078711 | -0.30153 |
| Aspartate aminotransferase, mitochondrial | P05202 | 0.042217 | 0.229305 |
| 26S proteasome non-ATPase regulatory subunit 13 | Q9WVJ2 | 0.127201 | 0.339108 |
| Neurolysin, mitochondrial | Q91YP2 | 0.17443 | -0.24071 |
| Carboxypeptidase Q | Q9WVJ3 | 0.209378 | -0.15997 |
| ES1 protein homolog, mitochondrial | Q9D172 | 0.695932 | 0.046944 |
| Synaptic vesicle membrane protein VAT-1 homolog | Q62465 | 0.778969 | -0.04487 |
| CD44 antigen | P15379 | 0.056482 | -0.42152 |
| 1-acylglycerol-3-phosphate O-acyltransferase ABHD5 | Q9DBL9 | 0.062232 | 4.026904 |
| Asparagine synthetase [glutamine-hydrolyzing] | Q61024 | 0.086197 | 0.190796 |
| ELAV-like protein 1 | P70372 | 0.094826 | -0.26408 |
| Dolichyl-diphosphooligosaccharide--protein glycosyltransferase | O54734 | 0.153177 | -0.20903 |
| Phospholipid scramblase 3 | Q9JIZ9 | 0.230971 | -0.23924 |
| Dynamin-1-like protein | Q8K1M6 | 0.260983 | 0.138226 |
| Short-chain specific acyl-CoA dehydrogenase, mitochondrial | Q07417 | 0.535404 | 0.113467 |
| ADP-ribosylation factor 5 | P84084 | 0.99981 | 2.18E-05 |
| Succinate dehydrogenase [ubiquinone] iron-sulfur subunit, mito | Q9CQA3 | 0.070983 | 0.241234 |
| Flotillin-2 | Q60634 | 0.249206 | 0.152817 |
| Immunoglobulin superfamily containing leucine-rich repeat prot | Q6GU68 | 0.065465 | 0.546664 |
| Neutral amino acid transporter B(0) | P51912 | 0.255613 | 0.180258 |
| Serpin B6 | Q60854 | 0.279941 | 0.096076 |

| | | | |
|---|---|---|---|
| Alpha-galactosidase A | P51569 | 0.477284 | -0.08037 |
| Phosphoglucomutase-1 | Q9D0F9 | 0.118023 | 0.17647 |
| Trifunctional enzyme subunit alpha, mitochondrial;Long-chain e | Q8BMS1 | 0.429216 | 0.09855 |
| Neutral alpha-glucosidase AB | Q8BHN3 | 0.388474 | 0.064315 |
| High mobility group protein B1 | P63158 | 0.878407 | -0.01564 |
| Nucleolar transcription factor 1 | P25976 | 0.300023 | -0.22189 |
| MICOS complex subunit Mic19 | Q9CRB9 | 0.556437 | 0.114688 |
| 40S ribosomal protein S13 | P62301 | 0.886287 | 0.012929 |
| PDZ and LIM domain protein 2 | Q8R1G6 | 0.216661 | -0.21521 |
| 26S proteasome non-ATPase regulatory subunit 9 | Q9CR00 | 0.818882 | -0.03442 |
| Spartin | Q8R1X6 | 0.057464 | 0.285256 |
| Cyclin-dependent kinase 4 inhibitor C | Q60772 | 0.079098 | -0.36288 |
| 6-phosphogluconate dehydrogenase, decarboxylating | Q9DCD0 | 0.169511 | 0.197854 |
| Heterogeneous nuclear ribonucleoprotein D0 | Q60668 | 0.691532 | 0.026025 |
| Glutathione S-transferase theta-1 | Q64471 | 0.901425 | -0.0213 |
| Prolow-density lipoprotein receptor-related protein 1;Low-dens | Q91ZX7 | 0.987946 | 0.000978 |
| Lysosome-associated membrane glycoprotein 1 | P11438 | 0.706468 | 0.027363 |
| Programmed cell death 6-interacting protein | Q9WU78 | 0.872441 | -0.02396 |
| ADP-ribosylation factor-like protein 1 | P61211 | 0.097181 | 0.332934 |
| Glutathione peroxidase 1 | P11352 | 0.947319 | 0.012652 |
| Serine/threonine-protein phosphatase PP1-beta catalytic subun | P62141 | 0.451982 | -0.14342 |
| Calponin-3 | Q9DAW9 | 0.835092 | -0.02476 |
| Fermitin family homolog 2 | Q8CIB5 | 0.360909 | -0.08785 |
| Dynein light chain 1, cytoplasmic | P63168 | 0.405089 | 0.062087 |
| Neural cell adhesion molecule 1 | P13595 | 0.218718 | 0.110016 |
| Enoyl-CoA delta isomerase 2, mitochondrial | Q9WUR2 | 0.24907 | 0.192803 |
| ATP-dependent (S)-NAD(P)H-hydrate dehydratase | Q9CZ42 | 0.970048 | -0.00412 |
| Heat shock protein 75 kDa, mitochondrial | Q9CQN1 | 0.300513 | 0.154674 |
| Insulin-like growth factor 2 mRNA-binding protein 3 | Q9CPN8 | 0.385113 | -0.05603 |
| Nuclease-sensitive element-binding protein 1 | P62960 | 0.474675 | 0.233931 |
| Selenide, water dikinase 1 | Q8BH69 | 0.528855 | -1.37453 |
| TBC1 domain family member 9B | Q5SVR0 | 0.251396 | -0.20532 |
| Mitochondrial import inner membrane translocase subunit Tim1 | Q9Z0V7 | 0.79348 | -0.06608 |
| NEDD8-activating enzyme E1 catalytic subunit | Q8C878 | 0.744282 | -0.60798 |
| Dynein light chain Tctex-type 1 | P51807 | 0.027318 | 0.271439 |
| Glycylpeptide N-tetradecanoyltransferase 1 | O70310 | 0.041253 | 0.277111 |
| U6 snRNA-associated Sm-like protein LSm3 | P62311 | 0.020508 | 0.504925 |
| Epididymis-specific alpha-mannosidase | O54782 | 0.862004 | -0.0237 |
| Tubulin beta-6 chain | Q922F4 | 0.46241 | 0.081997 |
| Translocation protein SEC62 | Q8BU14 | 0.221409 | 0.120572 |
| Glycerol-3-phosphate dehydrogenase, mitochondrial | Q64521 | 0.421744 | 0.166308 |
| Follistatin-related protein 1 | Q62356 | 0.891616 | -0.02018 |
| Beta-glucuronidase | P12265 | 0.914387 | 0.007178 |
| Tapasin | Q9R233 | 0.045269 | 0.492111 |
| 2-oxoglutarate dehydrogenase, mitochondrial | Q60597 | 0.636885 | 0.046844 |
| Protein canopy homolog 4 | Q8BQ47 | 0.260174 | -0.39646 |
| Septin-2 | P42208 | 0.909715 | 0.014316 |
| Zinc-binding alcohol dehydrogenase domain-containing protein | Q8BGC4 | 0.555738 | 0.093173 |
| Acylphosphatase-1 | P56376 | 0.342065 | 0.128583 |
| Adenylyl cyclase-associated protein 1 | P40124 | 0.182171 | -0.12092 |

| | | | |
|---|---|---|---|
| Syntaxin-12 | Q9ER00 | 0.34724 | 0.25456 |
| Isocitrate dehydrogenase [NADP] cytoplasmic | O88844 | 0.325435 | 0.118783 |
| Lysosomal protective protein;Lysosomal protective protein 32 kl | P16675 | 0.855445 | -0.02779 |
| Glutathione S-transferase theta-2 | Q99L20 | 0.642917 | 0.06681 |
| Protein S100-A4 | P07091 | 0.843364 | 0.035455 |
| N-acetylgalactosamine-6-sulfatase | Q571E4 | 0.609586 | -0.05575 |
| Adenylate kinase 2, mitochondrial;Adenylate kinase 2, mitochor | Q9WTP6 | 0.188763 | 0.20649 |
| Palladin | Q9ET54 | 0.307792 | -0.32979 |
| HIV Tat-specific factor 1 homolog | Q8BGC0 | 0.328604 | 0.330044 |
| Transketolase | P40142 | 0.34809 | 0.09337 |
| Ubiquitin-conjugating enzyme E2 L3 | P68037 | 0.440348 | 0.072685 |
| NADH dehydrogenase [ubiquinone] 1 beta subcomplex subunit | O09111 | 0.781809 | 0.043489 |
| Forkhead box protein K1 | P42128 | 0.426919 | -0.12862 |
| Band 4.1-like protein 1 | Q9Z2H5 | 0.53031 | 0.362926 |
| Dihydropyrimidinase-related protein 2 | O08553 | 0.443098 | 0.06434 |
| 14-3-3 protein zeta/delta | P63101 | 0.830234 | -0.01456 |
| Far upstream element-binding protein 2 | Q3U0V1 | 0.86882 | 0.01974 |
| Rab GDP dissociation inhibitor beta | Q61598 | 0.605077 | -0.06742 |
| Brain acid soluble protein 1 | Q91XV3 | 0.231692 | 0.183461 |
| Fatty aldehyde dehydrogenase | P47740 | 0.533364 | -0.11841 |
| Lysine--tRNA ligase | Q99MN1 | 0.628763 | 0.034499 |
| 4F2 cell-surface antigen heavy chain | P10852 | 0.812541 | 0.027411 |
| Electron transfer flavoprotein subunit beta | Q9DCW4 | 0.22277 | 0.220803 |
| Zinc transporter SLC39A7 | Q31125 | 0.041464 | 0.259323 |
| Adenosine kinase | P55264 | 0.524711 | -0.08227 |
| Methylmalonyl-CoA mutase, mitochondrial | P16332 | 0.260876 | -0.26015 |
| Actin-related protein 2/3 complex subunit 2 | Q9CVB6 | 0.294933 | -0.16217 |
| Stromal cell-derived factor 1 | P40224 | 0.352518 | -0.14471 |
| Alpha-enolase | P17182 | 0.688238 | -0.04467 |
| 60S ribosomal protein L7a | P12970 | 0.890309 | 0.012974 |
| Threonine--tRNA ligase, cytoplasmic | Q9D0R2 | 0.517478 | 0.050455 |
| 26S proteasome non-ATPase regulatory subunit 7 | P26516 | 0.686914 | 0.039877 |
| Purine nucleoside phosphorylase | P23492 | 0.794906 | -0.03923 |
| S-phase kinase-associated protein 1 | Q9WTX5 | 0.831575 | 0.023642 |
| Basement membrane-specific heparan sulfate proteoglycan cor | Q05793 | 0.950164 | -0.00649 |
| Bcl-2-associated transcription factor 1 | Q8K019 | 0.987145 | -0.00185 |
| Beta-actin-like protein 2 | Q8BFZ3 | 0.066302 | 0.30001 |
| Cytoplasmic FMR1-interacting protein 1;Cytoplasmic FMR1-inte | Q7TMB8 | 0.445054 | -0.15111 |
| Emerin | O08579 | 0.469818 | -0.07906 |
| Nidogen-1 | P10493 | 0.467566 | -0.17991 |
| Collagen alpha-1(V) chain | O88207 | 0.963836 | 0.004021 |
| NADH dehydrogenase [ubiquinone] 1 beta subcomplex subunit | Q9DCS9 | 0.397153 | 0.451932 |
| Heterogeneous nuclear ribonucleoprotein Q | Q7TMK9 | 0.072759 | 0.17055 |
| Hypoxia up-regulated protein 1 | Q9JKR6 | 0.193488 | 0.145873 |
| UV excision repair protein RAD23 homolog A | P54726 | 0.653321 | 0.15443 |
| Protein enabled homolog | Q03173 | 0.951097 | 0.0103 |
| Peroxiredoxin-6 | O08709 | 0.822919 | 0.024966 |
| Ubiquilin-1 | Q8R317 | 0.775638 | -0.09693 |
| Monoacylglycerol lipase ABHD12 | Q99LR1 | 0.433396 | -0.1748 |
| Staphylococcal nuclease domain-containing protein 1 | Q78PY7 | 0.498479 | -0.06862 |

| | | | |
|---|---|---|---|
| Eukaryotic translation initiation factor 3 subunit I | Q9QZD9 | 0.058398 | 0.273115 |
| Proteasome assembly chaperone 1 | Q9JK23 | 0.772178 | -0.09086 |
| Insulin-like growth factor 2 mRNA-binding protein 2 | Q5SF07 | 0.857682 | -0.0162 |
| Moesin | P26041 | 0.977862 | -0.00214 |
| Protein transport protein Sec31A | Q3UPL0 | 0.207044 | -0.0949 |
| UDP-glucose 6-dehydrogenase | O70475 | 0.835892 | 0.026609 |
| DNA damage-binding protein 1 | Q3U1J4 | 0.025162 | 0.300645 |
| Prolyl 4-hydroxylase subunit alpha-1 | Q60715 | 0.706021 | 0.042803 |
| GMP synthase [glutamine-hydrolyzing] | Q3THK7 | 0.333773 | -0.17387 |
| Prohibitin-2 | O35129 | 0.931693 | 0.006815 |
| Coactosin-like protein | Q9CQI6 | 0.961183 | -0.00589 |
| Phosphoenolpyruvate carboxykinase [GTP], mitochondrial | Q8BH04 | 0.96617 | -0.00572 |
| Protein deglycase DJ-1 | Q99LX0 | 0.691295 | -0.02769 |
| Ribosyldihydronicotinamide dehydrogenase [quinone] | Q9JI75 | 0.986807 | -0.00277 |
| Liprin-beta-1 | Q8C8U0 | 0.253068 | -0.2684 |
| ATPase Asna1 | O54984 | 0.375052 | -0.1434 |
| N-acetylgalactosamine kinase | Q68FH4 | 0.736682 | -0.0475 |
| Endoplasmin | P08113 | 0.220543 | 0.108878 |
| U6 snRNA-associated Sm-like protein LSm8 | Q6ZWM4 | 0.39441 | 0.147489 |
| Ubiquitin thioesterase OTUB1 | Q7TQI3 | 0.905349 | -0.01416 |
| Copper transport protein ATOX1 | O08997 | 0.804291 | -0.02817 |
| Proliferating cell nuclear antigen | P17918 | 0.637338 | -0.05437 |
| Growth arrest-specific protein 1 | Q01721 | 0.307098 | 0.128756 |
| Phosphoglycerate mutase 1 | Q9DBJ1 | 0.380657 | -0.07563 |
| Proteasome subunit alpha type-4 | Q9R1P0 | 0.490568 | 0.127639 |
| Phosphatidylinositol-binding clathrin assembly protein | Q7M6Y3 | 0.536661 | 0.186709 |
| Mitochondrial carrier homolog 2 | Q791V5 | 0.648937 | -0.13088 |
| 60S ribosomal protein L17 | Q9CPR4 | 0.124415 | 0.197677 |
| Procollagen C-endopeptidase enhancer 1 | Q61398 | 0.666312 | -0.04909 |
| Prostaglandin E synthase | Q9JM51 | 0.61819 | 0.243157 |
| Poly(ADP-ribose) glycohydrolase ARH3 | Q8CG72 | 0.116075 | -0.27317 |
| UPF0505 protein C16orf62 homolog | Q8BWQ6 | 0.19673 | 0.488108 |
| TAR DNA-binding protein 43 | Q921F2 | 0.203855 | -0.09847 |
| Tensin-3 | Q5SSZ5 | 0.298423 | 0.096893 |
| Mannosyl-oligosaccharide glucosidase | Q80UM7 | 0.491187 | 0.070614 |
| Proteasome activator complex subunit 2 | P97372 | 0.535257 | -0.09884 |
| Alcohol dehydrogenase [NADP(+)] | Q9JII6 | 0.558925 | -0.07791 |
| Ribonuclease UK114 | P52760 | 0.763182 | 0.035602 |
| Signal transducer and activator of transcription 5B;Signal transd | P42232 | 0.25272 | -0.14766 |
| Immunoglobulin-binding protein 1;Immunoglobulin-binding pro | Q61249 | 0.493381 | 0.180103 |
| ATP synthase subunit d, mitochondrial | Q9DCX2 | 0.591113 | 0.060156 |
| Heterogeneous nuclear ribonucleoprotein K | P61979 | 0.762525 | 0.021523 |
| SPRY domain-containing protein 4 | Q91WK1 | 0.748035 | -0.14956 |
| Proteasome subunit alpha type-6 | Q9QUM9 | 0.046914 | 0.257568 |
| Ragulator complex protein LAMTOR1 | Q9CQ22 | 0.137732 | 3.110822 |
| Cold shock domain-containing protein E1 | Q91W50 | 0.554127 | 0.219305 |
| 60S ribosomal protein L18 | P35980 | 0.193941 | 0.120125 |
| Nucleosome assembly protein 1-like 1 | P28656 | 0.109023 | 0.24016 |
| Ubiquitin-conjugating enzyme E2 H | P62257 | 0.589229 | -0.07904 |
| General transcription factor II-I | Q9ESZ8 | 0.487062 | -0.12199 |

| | | | |
|---|---|---|---|
| Beta-galactosidase | P23780 | 0.892034 | -0.01646 |
| Probable ubiquitin carboxyl-terminal hydrolase FAF-X | P70398 | 0.941132 | 0.019787 |
| Nucleobindin-1 | Q02819 | 0.551796 | 0.046678 |
| Histone H1.3 | P43277 | 0.445038 | 0.1179 |
| E3 ubiquitin-protein ligase UBR4 | A2AN08 | 0.84893 | 0.067925 |
| Poly(rC)-binding protein 1 | P60335 | 0.596783 | 0.055796 |
| Eukaryotic peptide chain release factor GTP-binding subunit ERF | Q8R050 | 0.046036 | 0.39257 |
| Septin-6 | Q9R1T4 | 0.816763 | -0.05709 |
| Adapter molecule crk | Q64010 | 0.75727 | -0.04055 |
| Sorting nexin-9 | Q91VH2 | 0.120885 | 0.204808 |
| 60S ribosomal protein L13 | P47963 | 0.459588 | 0.149846 |
| Glucosylceramidase | P17439 | 0.096931 | -0.21639 |
| Bifunctional glutamate/proline--tRNA ligase;Glutamate--tRNA lig | Q8CGC7 | 0.958479 | 0.003709 |
| Apoptosis-inducing factor 1, mitochondrial | Q9Z0X1 | 0.540032 | 0.156333 |
| Phenylalanine--tRNA ligase beta subunit | Q9WUA2 | 0.612861 | 0.050874 |
| Serine/threonine-protein kinase 3;Serine/threonine-protein kina | Q9JI10 | 0.642052 | 0.057282 |
| 2-oxoisovalerate dehydrogenase subunit alpha, mitochondrial | P50136 | 0.645291 | 0.119555 |
| 14-3-3 protein sigma | O70456 | 0.766803 | 0.031818 |
| Glutathione S-transferase Mu 1 | P10649 | 0.841891 | 0.029779 |
| Succinyl-CoA:3-ketoacid coenzyme A transferase 1, mitochondri | Q9D0K2 | 0.141962 | 0.117244 |
| Low molecular weight phosphotyrosine protein phosphatase | Q9D358 | 0.866324 | 0.021953 |
| Monofunctional C1-tetrahydrofolate synthase, mitochondrial | Q3V3R1 | 0.359874 | 0.156618 |
| Elongation factor Ts, mitochondrial | Q9CZR8 | 0.114956 | -0.41341 |
| UDP-glucose:glycoprotein glucosyltransferase 1 | Q6P5E4 | 0.099353 | 0.23711 |
| Isoleucine--tRNA ligase, mitochondrial | Q8BIJ6 | 0.535658 | 0.119139 |
| Histone H2B type 1-P;Histone H2B type 1-K;Histone H2B type 1- | Q8CGP2 | 0.726446 | 0.039569 |
| 26S protease regulatory subunit 6A | O88685 | 0.804387 | 0.039395 |
| SH3 domain-binding glutamic acid-rich-like protein 3 | Q91VW3 | 0.541534 | -0.10354 |
| Trans-L-3-hydroxyproline dehydratase | Q9CXA2 | 0.299065 | 0.156278 |
| ADP-ribosylation factor 1;ADP-ribosylation factor 3;ADP-ribosyla | P84078 | 0.408836 | 0.086627 |
| ATP-dependent RNA helicase DDX1 | Q91VR5 | 0.554038 | 0.076578 |
| COP9 signalosome complex subunit 3 | O88543 | 0.606218 | -0.12371 |
| Major facilitator superfamily domain-containing protein 1 | Q9DC37 | 0.62929 | 0.147208 |
| Guanine nucleotide-binding protein G(I)/G(S)/G(O) subunit gam | Q9DAS9 | 0.65348 | -0.03317 |
| | Q9QX15 | 0.199399 | 0.197713 |
| 60S ribosomal protein L35a | O55142 | 0.271432 | -0.19717 |
| Coatomer subunit zeta-1 | P61924 | 0.432473 | -0.41718 |
| Annexin A5 | P48036 | 0.500111 | 0.084793 |
| 40S ribosomal protein S21 | Q9CQR2 | 0.567698 | -0.04672 |
| Cullin-associated NEDD8-dissociated protein 1 | Q6ZQ38 | 0.972876 | 0.002296 |
| 60S ribosomal protein L19 | P84099 | 0.324085 | 0.099751 |
| Putative RNA-binding protein Luc7-like 2 | Q7TNC4 | 0.392446 | 0.097412 |
| 26S proteasome non-ATPase regulatory subunit 1 | Q3TXS7 | 0.492109 | 0.073895 |
| Cytochrome c oxidase subunit 7C, mitochondrial | P17665 | 0.586669 | 0.242656 |
| Malectin | Q6ZQI3 | 0.859149 | 0.011737 |
| NHP2-like protein 1;NHP2-like protein 1, N-terminally processed | Q9D0T1 | 0.646085 | -0.0983 |
| Peptidyl-prolyl cis-trans isomerase B | P24369 | 0.806026 | 0.028459 |
| ATP synthase subunit f, mitochondrial | P56135 | 0.630761 | 0.071158 |
| Leucine-rich PPR motif-containing protein, mitochondrial | Q6PB66 | 0.157738 | 0.146009 |
| Matrin-3 | Q8K310 | 0.395077 | -0.16083 |

| | | | |
|---|---|---|---|
| Dipeptidyl peptidase 3 | Q99KK7 | 0.418087 | 0.064438 |
| 39S ribosomal protein L12, mitochondrial | Q9DB15 | 0.980432 | 0.004753 |
| Cell division cycle and apoptosis regulator protein 1 | Q8CH18 | 0.006431 | 1.025811 |
| Golgi reassembly-stacking protein 2 | Q99JX3 | 0.178619 | 0.330483 |
| V-type proton ATPase subunit H | Q8BVE3 | 0.969502 | -0.00332 |
| S-adenosylmethionine synthase isoform type-2 | Q3THS6 | 0.328734 | -0.18033 |
| Myosin-9 | Q8VDD5 | 0.633226 | 0.048141 |
| 26S proteasome non-ATPase regulatory subunit 4 | O35226 | 0.818983 | 0.032722 |
| EH domain-containing protein 2 | Q8BH64 | 0.296337 | 0.088663 |
| Asparagine--tRNA ligase, cytoplasmic | Q8BP47 | 0.192481 | 0.376035 |
| Annexin A4 | P97429 | 0.287261 | 0.076047 |
| 26S protease regulatory subunit 6B | P54775 | 0.322865 | 0.098066 |
| Cytochrome b-c1 complex subunit 9 | Q8R1I1 | 0.268598 | -0.1702 |
| PC4 and SFRS1-interacting protein | Q99JF8 | 0.303332 | -0.2408 |
| Myosin-14 | Q6URW6 | 0.763296 | -0.07415 |
| Platelet-activating factor acetylhydrolase IB subunit beta | Q61206 | 0.258896 | 0.088715 |
| Acyl-protein thioesterase 2 | Q9WTL7 | 0.440359 | -0.14229 |
| Heat shock 70 kDa protein 4 | Q61316 | 0.622335 | 0.0419 |
| Superoxide dismutase [Mn], mitochondrial | P09671 | 0.629152 | -0.06519 |
| Perilipin-2 | P43883 | 0.262659 | 0.135833 |
| Dephospho-CoA kinase domain-containing protein | Q8BHC4 | 0.805137 | -0.05466 |
| Cytoplasmic dynein 1 light intermediate chain 1 | Q8R1Q8 | 0.975847 | 0.003833 |
| YTH domain-containing family protein 1;YTH domain-containing | P59326 | 0.043124 | -0.58966 |
| Heme-binding protein 1 | Q9R257 | 0.758314 | 0.031226 |
| 14-3-3 protein epsilon | P62259 | 0.989335 | 0.002899 |
| Thioredoxin domain-containing protein 17 | Q9CQM5 | 0.025538 | 0.431884 |
| Coiled-coil domain-containing protein 186 | Q8C9S4 | 0.133353 | 3.115046 |
| Zinc finger protein 512 | Q69Z99 | 0.145312 | 0.632387 |
| Galectin-9 | O08573 | 0.281624 | 0.193289 |
| Calpain-2 catalytic subunit | O08529 | 0.349461 | 0.178959 |
| Surfeit locus protein 4 | Q64310 | 0.447167 | 0.137919 |
| UPF0687 protein C20orf27 homolog | Q9D1K7 | 0.491929 | -0.21078 |
| Arginine--tRNA ligase, cytoplasmic | Q9D0I9 | 0.522021 | -0.04728 |
| Collagen alpha-1(VI) chain | Q04857 | 0.618362 | 0.119029 |
| Cystatin-C | P21460 | 0.656548 | -0.04635 |
| Complement C1q tumor necrosis factor-related protein 5 | Q8K479 | 0.960734 | -0.00839 |
| L-lactate dehydrogenase A chain | P06151 | 0.988515 | 0.001502 |
| Actin-related protein 2/3 complex subunit 1B | Q9WV32 | 0.199667 | 0.152136 |
| Sister chromatid cohesion protein PDS5 homolog B | Q4VA53 | 0.72989 | -0.77594 |
| Protein SET | Q9EQU5 | 0.673982 | 0.042563 |
| Ubiquitin-conjugating enzyme E2 variant 2 | Q9D2M8 | 0.749338 | 0.110127 |
| Heterogeneous nuclear ribonucleoprotein H;Heterogeneous nuc | O35737 | 0.200994 | 0.124772 |
| Protein transport protein Sec61 subunit beta | Q9CQS8 | 0.257112 | -0.34021 |
| Alpha/beta hydrolase domain-containing protein 14B | Q8VCR7 | 0.500362 | -0.09478 |
| Histone H3.3;Histone H3.2;Histone H3.1;Histone H3.3C | P84244 | 0.700355 | 0.074612 |
| Metalloreductase STEAP3 | Q8CI59 | 0.757238 | 0.026737 |
| 60S ribosomal protein L7 | P14148 | 0.868146 | 0.013282 |
| Translational activator of cytochrome c oxidase 1 | Q8K0Z7 | 0.04319 | 0.416131 |
| Bis(5-nucleosyl)-tetraphosphatase [asymmetrical] | P56380 | 0.952783 | 0.01225 |
| Probable glutathione peroxidase 8 | Q9D7B7 | 0.958027 | 0.015473 |

| | | | |
|---|---|---|---|
| Niemann-Pick C1 protein | O35604 | 0.190532 | 2.836757 |
| Clathrin light chain B | Q6IRU5 | 0.292324 | -0.14061 |
| Double-strand break repair protein MRE11A | Q61216 | 0.153408 | 0.338554 |
| Synaptosomal-associated protein 23 | O09044 | 0.877693 | 0.03422 |
| Tyrosine-protein phosphatase non-receptor type 11 | P35235 | 0.770319 | -0.0371 |
| Oligosaccharyltransferase complex subunit OSTC | Q78XF5 | 0.866399 | 0.043431 |
| SH3 domain-binding protein 1 | P55194 | 0.407461 | -0.50492 |
| ELKS/Rab6-interacting/CAST family member 1 | Q99MI1 | 0.616201 | 0.273494 |
| Ribonuclease inhibitor | Q91VI7 | 0.643893 | -0.04541 |
| Small ubiquitin-related modifier 2 | P61957 | 0.922969 | -0.00985 |
| NEDD8 | P29595 | 0.169938 | 0.390965 |
| H(+)/Cl(-) exchange transporter 7 | O70496 | 0.239029 | 0.163335 |
| MICOS complex subunit Mic25 | Q91VN4 | 0.86675 | -0.02499 |
| Transmembrane emp24 domain-containing protein 10 | Q9D1D4 | 0.577152 | -0.17353 |
| Protein transport protein Sec23A | Q01405 | 0.110384 | -0.14347 |
| Microtubule-associated protein RP/EB family member 1 | Q61166 | 0.679328 | 0.096045 |
| F-box only protein 6 | Q9QZN4 | 0.264473 | 0.188017 |
| Tubulin--tyrosine ligase-like protein 12 | Q3UDE2 | 0.42004 | -0.14711 |
| Electron transfer flavoprotein-ubiquinone oxidoreductase, mito | Q921G7 | 0.789595 | 0.05483 |
| RNA-binding protein 39 | Q8VH51 | 0.837654 | -0.0311 |
| Protein CREG1 | O88668 | 0.419845 | 0.148145 |
| Programmed cell death protein 4 | Q61823 | 0.385684 | -0.16212 |
| Sorbitol dehydrogenase | Q64442 | 0.691454 | 0.076268 |
| Myosin light chain kinase, smooth muscle;Myosin light chain kin | Q6PDN3 | 0.72416 | -0.08887 |
| Annexin A3 | O35639 | 0.740447 | 0.103062 |
| Ras-related protein Rab-21 | P35282 | 0.089508 | -0.33337 |
| Histone H1.4 | P43274 | 0.734898 | 0.03088 |
| Multifunctional methyltransferase subunit TRM112-like protein | Q9DCG9 | 0.093577 | -0.8252 |
| Ras-related protein Rab-2A;Ras-related protein Rab-2B | P53994 | 0.565671 | 0.077323 |
| Secernin-1 | Q9CZC8 | 0.342713 | 0.218374 |
| Guanine nucleotide exchange factor for Rab-3A | Q8VDV3 | 0.354809 | 0.102467 |
| 26S protease regulatory subunit 4 | P62192 | 0.487444 | -0.05212 |
| Chromobox protein homolog 3 | P23198 | 0.134589 | 0.125889 |
| Phosphoglycerate kinase 1 | P09411 | 0.746899 | 0.021506 |
| Cofilin-1 | P18760 | 0.983041 | 0.002125 |
| 40S ribosomal protein S3 | P62908 | 0.935886 | -0.01113 |
| Metaxin-1 | P47802 | 0.343454 | -0.23745 |
| Trafficking protein particle complex subunit 6B | Q9D289 | 0.625816 | 0.301043 |
| Cytoplasmic dynein 1 heavy chain 1 | Q9JHU4 | 0.92489 | -0.0097 |
| Scaffold attachment factor B1 | D3YXK2 | 0.060463 | 0.380785 |
| Mitochondrial 2-oxoglutarate/malate carrier protein | Q9CR62 | 0.172105 | 0.151758 |
| Suppressor of G2 allele of SKP1 homolog | Q9CX34 | 0.325407 | 0.07782 |
| Polyadenylate-binding protein 1 | P29341 | 0.484305 | -0.07144 |
| Cytochrome c oxidase subunit 5A, mitochondrial | P12787 | 0.635019 | 0.13449 |
| Major prion protein | P04925 | 0.010472 | -0.3114 |
| Ras GTPase-activating-like protein IQGAP1 | Q9JKF1 | 0.574841 | 0.089601 |
| Septin-9 | Q8OUG5 | 0.876325 | -0.02025 |
| Transmembrane protein 165 | P52875 | 0.467667 | 0.1518 |
| Thrombospondin-1 | P35441 | 0.561273 | 0.075271 |
| Pyrroline-5-carboxylate reductase 3 | Q9DCC4 | 0.208633 | 0.224372 |

| | | | |
|---|---|---|---|
| | B1AVD1 | 0.463873 | -0.24832 |
| Glutaminase kidney isoform, mitochondrial | D3Z7P3 | 0.709745 | -0.05598 |
| 40S ribosomal protein S20 | P60867 | 0.40196 | 0.110273 |
| Serine/threonine-protein phosphatase 2A 65 kDa regulatory sub | Q76MZ3 | 0.420219 | 0.0916 |
| Prefoldin subunit 3 | P61759 | 0.503046 | 0.244398 |
| Nucleolin | P09405 | 0.554299 | 0.115571 |
| Dynamin-2 | P39054 | 0.504712 | -0.06923 |
| Elongation factor 1-beta | O70251 | 0.82255 | 0.015061 |
| Integrin alpha-V;Integrin alpha-V heavy chain;Integrin alpha-V li | P43406 | 0.957319 | 0.008486 |
| Tetraspanin-6 | O70401 | 0.777304 | 0.027315 |
| Signal recognition particle subunit SRP68 | Q8BMA6 | 0.97734 | -0.00466 |
| Serine/threonine-protein kinase DCLK1 | Q9JLM8 | 0.245714 | 0.129308 |
| Cleavage and polyadenylation specificity factor subunit 5 | Q9CQF3 | 0.904742 | -0.03932 |
| Protein transport protein Sec31B | Q3TZ89 | 0.673953 | 0.069173 |
| N-acetyl-D-glucosamine kinase | Q9QZ08 | 0.144184 | -0.29321 |
| Heterogeneous nuclear ribonucleoproteins A2/B1 | O88569 | 0.857301 | 0.045614 |
| Bifunctional purine biosynthesis protein PURH;Phosphoribosylar | Q9CWJ9 | 0.89723 | -0.01879 |
| Protein LSM12 homolog | Q9D0R8 | 0.103897 | -0.44324 |
| Actin-related protein 2/3 complex subunit 4 | P59999 | 0.301749 | 0.146413 |
| 26S protease regulatory subunit 7 | P46471 | 0.359041 | 0.126312 |
| Serine/arginine-rich splicing factor 7 | Q8BL97 | 0.748049 | -0.04519 |
| 5-oxoprolinase | Q8K010 | 0.584203 | -0.22558 |
| Methylthioribulose-1-phosphate dehydratase | Q9WVQ5 | 0.599296 | 0.124943 |
| Isocitrate dehydrogenase [NADP], mitochondrial | P54071 | 0.175878 | 0.118063 |
| Eukaryotic translation initiation factor 1A, X-chromosomal;Eukar | Q8BMJ3 | 0.57176 | -0.05917 |
| Proteasome subunit beta type-5 | O55234 | 0.756665 | 0.034738 |
| Sorcin | Q6P069 | 0.7735 | 0.029452 |
| cAMP-dependent protein kinase catalytic subunit alpha;cAMP-d | P05132 | 0.134721 | -0.18805 |
| Alcohol dehydrogenase class-3 | P28474 | 0.558769 | 0.06437 |
| Coronin-1B | Q9WUM3 | 0.746548 | 0.031619 |
| Caspase-8;Caspase-8 subunit p18;Caspase-8 subunit p10 | O89110 | 0.147708 | -0.70997 |
| Catechol O-methyltransferase | O88587 | 0.408652 | 0.124769 |
| U1 small nuclear ribonucleoprotein 70 kDa | Q62376 | 0.537745 | -0.05795 |
| Spermidine synthase | Q64674 | 0.582263 | -0.06887 |
| Dehydrogenase/reductase SDR family member 1 | Q99L04 | 0.974993 | -0.01391 |
| Cytochrome b5 | P56395 | 0.5912 | 0.070027 |
| Ubiquitin-like modifier-activating enzyme 5 | Q8VE47 | 0.482879 | 0.212012 |
| Beta-enolase | P21550 | 0.31705 | -0.19423 |
| T-complex protein 1 subunit alpha | P11983 | 0.613585 | 0.062587 |
| Cytosolic acyl coenzyme A thioester hydrolase | Q91V12 | 0.378627 | 0.130143 |
| Protein WWC2 | Q6NXJ0 | 0.254812 | 2.823636 |
| H/ACA ribonucleoprotein complex subunit 2 | Q9CRB2 | 0.837361 | 0.062099 |
| Voltage-dependent anion-selective channel protein 1 | Q60932 | 0.001363 | 0.290083 |
| Aldehyde dehydrogenase, mitochondrial | P47738 | 0.237726 | 0.10073 |
| Thymidylate kinase | P97930 | 0.348372 | -0.17349 |
| Enhancer of rudimentary homolog | P84089 | 0.575994 | -0.07161 |
| Phosphoglucomutase-2 | Q7TSV4 | 0.021829 | 0.563738 |
| Golgi-associated plant pathogenesis-related protein 1 | Q9CYL5 | 0.254068 | 0.198546 |
| Aminoacyl tRNA synthase complex-interacting multifunctional p | P31230 | 0.54616 | 0.085562 |
| AP-2 complex subunit alpha-1 | P17426 | 0.671944 | -0.11023 |

| | | | |
|---|---|---|---|
| Isochorismatase domain-containing protein 1 | Q91V64 | 0.191939 | -0.27099 |
| Peptidyl-prolyl cis-trans isomerase FKBP1A | P26883 | 0.643931 | -0.07889 |
| Aspartate--tRNA ligase, cytoplasmic | Q922B2 | 0.589575 | -0.09132 |
| GTP-binding protein SAR1b;GTP-binding protein SAR1a | Q9CQC9 | 0.279687 | 0.200453 |
| Glypican-4;Secreted glypican-4 | P51655 | 0.639881 | -0.10495 |
| SRA stem-loop-interacting RNA-binding protein, mitochondrial | Q9D8T7 | 0.855338 | -0.05784 |
| Ferritin heavy chain;Ferritin heavy chain, N-terminally processec | P09528 | 0.108913 | 0.191902 |
| Eukaryotic translation initiation factor 4H | Q9WUK2 | 0.33861 | 0.115053 |
| | Q8BML9 | 0.914712 | 0.024774 |
| Peripheral plasma membrane protein CASK | O70589 | 0.682275 | -0.13107 |
| Coatomer subunit epsilon | O89079 | 0.925798 | -0.04312 |
| Cytochrome b-c1 complex subunit Rieske, mitochondrial;Cytoch | Q9CR68 | 0.8842 | 0.026681 |
| Transcription factor A, mitochondrial | P40630 | 0.943042 | -0.02813 |
| AH receptor-interacting protein | O08915 | 0.415147 | -0.19374 |
| Acyl-CoA dehydrogenase family member 9, mitochondrial | Q8JZN5 | 0.677658 | 0.102757 |
| CD63 antigen | P41731 | 0.01251 | 0.64035 |
| Regulator of microtubule dynamics protein 3 | Q3UJU9 | 0.226172 | 3.149215 |
| Actin-related protein 3 | Q99JY9 | 0.791181 | -0.02934 |
| RNA-binding protein 25 | B2RY56 | 0.798661 | -0.06265 |
| BAG family molecular chaperone regulator 3 | Q9JLV1 | 0.011071 | -0.32169 |
| Protein tweety homolog 3 | Q6P5F7 | 0.063698 | -0.29094 |
| Alpha-aminoadipic semialdehyde dehydrogenase | Q9DBF1 | 0.302375 | 0.073559 |
| Dihydropteridine reductase | Q8BVI4 | 0.38305 | 0.166408 |
| Integrin alpha-5;Integrin alpha-5 heavy chain;Integrin alpha-5 lig | P11688 | 0.413161 | -0.11035 |
| Protein canopy homolog 2 | Q9QXT0 | 0.559858 | -0.08739 |
| Serine/threonine-protein phosphatase 2A catalytic subunit alph | P63330 | 0.566541 | 0.239319 |
| Histone H2A.J;Histone H2A type 1-H;Histone H2A type 1-K;Histo | Q8R1M2 | 0.61448 | 0.05053 |
| Chloride intracellular channel protein 1 | Q9Z1Q5 | 0.625838 | 0.062132 |
| Lactoylglutathione lyase | Q9CPU0 | 0.676108 | 0.064619 |
| Macrophage migration inhibitory factor | P34884 | 0.694348 | 0.033146 |
| Aspartate aminotransferase, cytoplasmic | P05201 | 0.99354 | -0.0009 |
| Serine protease HTRA2, mitochondrial | Q9JIY5 | 0.357352 | -0.29024 |
| Serum deprivation-response protein | Q63918 | 0.535539 | 0.301085 |
| Cyclin-dependent kinase inhibitor 2A | P51480 | 0.339517 | 0.254795 |
| Eukaryotic translation initiation factor 3 subunit B | Q8JZQ9 | 0.428804 | -0.14204 |
| Arylsulfatase A | P50428 | 0.445805 | 0.074337 |
| 14 kDa phosphohistidine phosphatase | Q9DAK9 | 0.52988 | -0.09673 |
| Very-long-chain enoyl-CoA reductase | Q9CY27 | 0.578539 | 0.113842 |
| N-acetylglucosamine-6-sulfatase | Q8BFR4 | 0.58273 | 0.039859 |
| Guanine nucleotide-binding protein G(o) subunit alpha | P18872 | 0.594009 | -0.06914 |
| Aldose reductase | P45376 | 0.646131 | -0.0321 |
| T-complex protein 1 subunit gamma | P80318 | 0.70276 | 0.035221 |
| AP-2 complex subunit alpha-2 | P17427 | 0.852205 | -0.01549 |
| Integrin-linked kinase-associated serine/threonine phosphatase | Q8R0F6 | 0.875361 | -0.08767 |
| SPARC | P07214 | 0.903164 | 0.020426 |
| Stress-induced-phosphoprotein 1 | Q60864 | 0.903982 | 0.015315 |
| Tubulin alpha-1A chain;Tubulin alpha-3 chain | P68369 | 0.061218 | 0.434899 |
| Multifunctional protein ADE2;Phosphoribosylaminoimidazole-su | Q9DCL9 | 0.921747 | -0.01358 |
| Vigilin | Q8VDJ3 | 0.588056 | 0.041697 |
| UTP--glucose-1-phosphate uridylyltransferase | Q91ZJ5 | 0.81468 | -0.0233 |

| | | | |
|---|---|---|---|
| Astrocytic phosphoprotein PEA-15 | Q62048 | 0.239994 | 0.496991 |
| ADP-ribosylation factor-like protein 8A | Q8VEH3 | 0.271897 | 0.163165 |
| Proteasome activator complex subunit 1 | P97371 | 0.825977 | 0.023312 |
| 60S ribosomal protein L30 | P62889 | 0.830714 | -0.0255 |
| BAG family molecular chaperone regulator 5 | Q8CI32 | 0.405373 | 0.198865 |
| Pre-mRNA-processing factor 6 | Q91YR7 | 0.553907 | 0.221889 |
| 60S ribosomal protein L22 | P67984 | 0.573331 | -0.12592 |
| Interleukin enhancer-binding factor 2 | Q9CXY6 | 0.896169 | -0.04979 |
| PRA1 family protein 3 | Q8R5J9 | 0.935328 | -0.02445 |
| COP9 signalosome complex subunit 4 | O88544 | 0.857169 | -0.03934 |
| Glutathione S-transferase Mu 5 | P48774 | 0.950263 | 0.034245 |
| HCLS1-binding protein 3 | Q3TC93 | 0.23125 | 0.304105 |
| Glutamine--fructose-6-phosphate aminotransferase [isomerizing | P47856 | 0.172174 | -0.10963 |
| Ester hydrolase C11orf54 homolog | Q91V76 | 0.018122 | -0.26523 |
| Tyrosine--tRNA ligase, cytoplasmic;Tyrosine--tRNA ligase, cytopl | Q91WQ3 | 0.749941 | -0.03292 |
| Cytoplasmic dynein 1 intermediate chain 2 | O88487 | 0.104231 | 0.231485 |
| EMILIN-1 | Q99K41 | 0.183994 | -0.19403 |
| Histone H4 | P62806 | 0.578877 | 0.150742 |
| RAC-alpha serine/threonine-protein kinase | P31750 | 0.084335 | 0.199553 |
| 60S ribosomal protein L6 | P47911 | 0.620172 | 0.033946 |
| Tryptophan--tRNA ligase, cytoplasmic;T1-TrpRS;T2-TrpRS | P32921 | 0.624899 | -0.14418 |
| Endoplasmic reticulum resident protein 29 | P57759 | 0.116089 | 0.186803 |
| [Protein ADP-ribosylarginine] hydrolase | P54923 | 0.774928 | 0.051743 |
| Elongation factor 1-alpha 1;Elongation factor 1-alpha 2 | P10126 | 0.787938 | 0.021099 |
| Ras-related protein Rab-5A | Q9CQD1 | 0.954516 | -0.0437 |
| Guanine nucleotide-binding protein G(i) subunit alpha-2;Guanin | P08752 | 0.656652 | -0.07015 |
| Ras-related protein Rab-7a | P51150 | 0.420609 | -0.11369 |
| Serine/threonine-protein kinase N1 | P70268 | 0.553495 | -0.1256 |
| Eukaryotic translation initiation factor 3 subunit H | Q91WK2 | 0.224684 | 2.459649 |
| COP9 signalosome complex subunit 2 | P61202 | 0.031286 | 0.328006 |
| Lysosome membrane protein 2 | O35114 | 0.612538 | -0.04144 |
| Acidic leucine-rich nuclear phosphoprotein 32 family member E | P97822 | 0.337459 | 0.334838 |
| Eukaryotic initiation factor 4A-III;Eukaryotic initiation factor 4A-I | Q91VC3 | 0.247759 | -0.18888 |
| Putative phospholipase B-like 2;Putative phospholipase B-like 2 | Q3TCN2 | 0.376447 | 0.097727 |
| Flotillin-1 | O08917 | 0.770845 | 0.05148 |
| ATP-binding cassette sub-family F member 1 | Q6P542 | 0.621653 | 0.094662 |
| Proteasome subunit beta type-7 | P70195 | 0.980271 | -0.00416 |
| Vacuolar protein sorting-associated protein 4B | P46467 | 0.043655 | -0.38637 |
| Aminopeptidase B | Q8VCT3 | 0.402659 | -0.24497 |
| 40S ribosomal protein S8 | P62242 | 0.90888 | -0.0106 |
| Pyrroline-5-carboxylate reductase 2;Pyrroline-5-carboxylate red | Q922Q4 | 0.528889 | -0.13305 |
| Glyoxalase domain-containing protein 4 | Q9CPV4 | 0.31807 | 0.084815 |
| Enoyl-CoA hydratase domain-containing protein 2, mitochondria | Q3TLP5 | 0.670164 | -0.0585 |
| E3 SUMO-protein ligase RanBP2 | Q9ERU9 | 0.268842 | 0.253211 |
| Protein FAM136A | Q9CR98 | 0.692274 | -0.06366 |
| Endophilin-B1 | Q9JK48 | 0.921684 | -0.04331 |
| Pituitary tumor-transforming gene 1 protein-interacting protein | Q8R143 | 0.175092 | 0.331597 |
| Prelamin-A/C;Lamin-A/C | P48678 | 0.530535 | 0.078812 |
| Ras-related protein Rab-10 | P61027 | 0.644667 | -0.0767 |
| RNA-binding protein 10 | Q99KG3 | 0.197427 | -0.30132 |

| | | | |
|---|---|---|---|
| Signal recognition particle 14 kDa protein;Signal recognition par | P16254 | 0.687074 | 0.061097 |
| Serine/arginine-rich splicing factor 5 | O35326 | 0.279617 | 0.229028 |
| Treacle protein | O08784 | 0.462015 | -0.11495 |
| Coiled-coil domain-containing protein 22 | Q9JIG7 | 0.409136 | 0.189499 |
| Eukaryotic translation initiation factor 3 subunit G | Q9Z1D1 | 0.438926 | 0.264946 |
| Constitutive coactivator of PPAR-gamma-like protein 1 | Q6A0A9 | 0.458111 | -0.05688 |
| Collagen alpha-1(IV) chain;Arresten | P02463 | 0.772703 | 0.036659 |
| Small glutamine-rich tetratricopeptide repeat-containing protei | Q8BJU0 | 0.296579 | 0.088457 |
| Peroxisomal multifunctional enzyme type 2;(3R)-hydroxyacyl-Co | P51660 | 0.871892 | 0.02253 |
| Poly(rC)-binding protein 2;Poly(rC)-binding protein 3 | Q61990 | 0.573895 | 0.06813 |
| Serpin H1 | P19324 | 0.101832 | 0.151796 |
| Alpha-1B adrenergic receptor | P97717 | 0.256037 | -0.26587 |
| Eukaryotic translation initiation factor 2 subunit 2 | Q99L45 | 0.497202 | 0.193273 |
| Phosphoserine aminotransferase | Q99K85 | 0.64943 | -0.05919 |
| CTP synthase 2 | P70303 | 0.964932 | -0.00606 |
| 26S proteasome non-ATPase regulatory subunit 2 | Q8VDM4 | 0.095768 | 0.153757 |
| Type 2 phosphatidylinositol 4,5-bisphosphate 4-phosphatase | Q9CZX7 | 0.245488 | 2.445423 |
| Valacyclovir hydrolase | Q8R164 | 0.262643 | 0.136127 |
| GDP-L-fucose synthase | P23591 | 0.286072 | 0.118669 |
| 60S ribosomal protein L23 | P62830 | 0.524104 | -0.06517 |
| Heterogeneous nuclear ribonucleoprotein U | Q8VEK3 | 0.529147 | 0.056063 |
| U5 small nuclear ribonucleoprotein 200 kDa helicase | Q6P4T2 | 0.854139 | 0.021869 |
| Vimentin | P20152 | 0.831114 | 0.01698 |
| Inosine triphosphate pyrophosphatase | Q9D892 | 0.870214 | -0.0377 |
| Latent-transforming growth factor beta-binding protein 3 | Q61810 | 0.758322 | 0.345441 |
| T-complex protein 1 subunit zeta | P80317 | 0.855809 | -0.02182 |
| Protein phosphatase 1 regulatory subunit 12A | Q9DBR7 | 0.7697 | -0.04245 |
| Lactadherin | P21956 | 0.210359 | -0.15596 |
| Collagen alpha-2(V) chain | Q3U962 | 0.562854 | -0.09082 |
| Guanine nucleotide-binding protein subunit beta-2-like 1;Guani | P68040 | 0.684007 | 0.03976 |
| Mitotic checkpoint protein BUB3 | Q9WVA3 | 0.897699 | -0.02361 |
| Alpha-soluble NSF attachment protein | Q9DB05 | 0.136815 | 0.366511 |
| DnaJ homolog subfamily B member 11 | Q99KV1 | 0.875209 | 0.025838 |
| Aspartyl/asparaginyl beta-hydroxylase | Q8BSY0 | 0.332349 | 0.176388 |
| Ran-specific GTPase-activating protein | P34022 | 0.110314 | 0.13182 |
| Mitochondrial 10-formyltetrahydrofolate dehydrogenase | Q8K009 | 0.593048 | 0.078268 |
| Protein FAM98B | Q80VD1 | 0.388592 | 0.386533 |
| AP-3 complex subunit sigma-1 | Q9DCR2 | 0.256992 | 0.162807 |
| Ras-related protein Rab-8A | P55258 | 0.575174 | 0.091384 |
| ATP synthase F(0) complex subunit C1, mitochondrial;ATP synth | Q9CR84 | 0.799351 | 0.107255 |
| Proteasome subunit beta type-3 | Q9R1P1 | 0.987055 | 0.00127 |
| Cation-dependent mannose-6-phosphate receptor | P24668 | 0.267703 | -0.11795 |
| AP-3 complex subunit mu-1;AP-3 complex subunit mu-2 | Q9JKC8 | 0.302976 | 0.179869 |
| Proteasome subunit alpha type-1 | Q9R1P4 | 0.480959 | 0.062856 |
| Propionyl-CoA carboxylase beta chain, mitochondrial | Q99MN9 | 0.662261 | -0.06706 |
| Protein kinase C delta-binding protein | Q91VJ2 | 0.827286 | 0.027195 |
| Heat shock-related 70 kDa protein 2 | P17156 | 0.654353 | -0.09525 |
| Exocyst complex component 4 | O35382 | 0.668836 | 0.144687 |
| Succinyl-CoA ligase [ADP-forming] subunit beta, mitochondrial | Q9Z2I9 | 0.721741 | 0.04379 |
| Hepatoma-derived growth factor-related protein 2 | Q3UMU9 | 0.868712 | -0.12323 |

| | | | |
|---|---|---|---|
| Protein jagunal homolog 1 | Q5XKN4 | 0.897891 | -0.03593 |
| Serine/arginine repetitive matrix protein 1 | Q52KI8 | 0.355663 | -0.5951 |
| Cell division cycle 5-like protein | Q6A068 | 0.502526 | -0.17423 |
| SH3 and multiple ankyrin repeat domains protein 3 | Q4ACU6 | 0.790482 | 0.152192 |
| Leucyl-cystinyl aminopeptidase | Q8C129 | 0.036275 | 0.27462 |
| Large neutral amino acids transporter small subunit 1 | Q9Z127 | 0.484941 | -0.12222 |
| Caveolin-1 | P49817 | 0.601555 | 0.045125 |
| Ras-related protein Rab-11A;Ras-related protein Rab-11B | P62492 | 0.72925 | 0.11039 |
| Eukaryotic initiation factor 4A-II;Eukaryotic initiation factor 4A-II | P10630 | 0.662268 | -0.08855 |
| Sorting and assembly machinery component 50 homolog | Q8BGH2 | 0.585955 | 0.097934 |
| UPF0317 protein C14orf159 homolog, mitochondrial | Q8BH86 | 0.297008 | 0.146479 |
| U2 small nuclear ribonucleoprotein A | P57784 | 0.507166 | -0.31337 |
| Eukaryotic translation initiation factor 2A;Eukaryotic translation | Q8BJW6 | 0.478013 | -0.27508 |
| Major vault protein | Q9EQK5 | 0.554482 | -0.06507 |
| Na(+)/H(+) exchange regulatory cofactor NHE-RF2 | Q9JHL1 | 0.984474 | -0.00865 |
| Death-associated protein 1 | Q91XC8 | 0.606411 | 0.151917 |
| tRNA-splicing ligase RtcB homolog | Q99LF4 | 0.886706 | 0.02377 |
| Triosephosphate isomerase | P17751 | 0.774061 | -0.03428 |
| Inverted formin-2 | Q0GNC1 | 0.941986 | 0.023955 |
| Histone H2A.V;Histone H2A.Z | Q3THW5 | 0.230303 | 0.211158 |
| Vesicle-associated membrane protein 3 | P63024 | 0.390689 | -0.20025 |
| T-complex protein 1 subunit eta | P80313 | 0.900314 | -0.01715 |
| Arf-GAP with Rho-GAP domain, ANK repeat and PH domain-cont | Q4LDD4 | 0.581256 | 0.11068 |
| DDRGK domain-containing protein 1 | Q80WW9 | 0.714387 | 0.235157 |
| Vesicle-fusing ATPase | P46460 | 0.671593 | -0.11419 |
| Acyl-coenzyme A thioesterase 13;Acyl-coenzyme A thioesterase | Q9CQR4 | 0.132974 | 0.199823 |
| U4/U6.U5 tri-snRNP-associated protein 2 | Q3TIX9 | 0.384932 | 1.962371 |
| Protein Noxp20 | Q9D281 | 0.095994 | 0.461498 |
| Syntaxin-binding protein 3 | Q60770 | 0.390588 | 0.232904 |
| Coatomer subunit alpha;Xenin;Proxenin | Q8CIE6 | 0.565402 | 0.073505 |
| C-1-tetrahydrofolate synthase, cytoplasmic;Methylenetetrahydr | Q922D8 | 0.703273 | -0.03543 |
| Proteasome subunit beta type-4 | P99026 | 0.36746 | 0.164386 |
| Drebrin-like protein | Q62418 | 0.426045 | -0.11114 |
| Hsp90 co-chaperone Cdc37;Hsp90 co-chaperone Cdc37, N-term | Q61081 | 0.593625 | 0.13474 |
| Tubulin-specific chaperone A | P48428 | 0.632393 | -0.06279 |
| Prostaglandin reductase 2 | Q8VDQ1 | 0.757892 | -0.0557 |
| Nascent polypeptide-associated complex subunit alpha;Nascent | Q60817 | 0.820501 | 0.036262 |
| Proteasome subunit alpha type-7 | Q9Z2U0 | 0.865464 | 0.025238 |
| Prosaposin | Q61207 | 0.726847 | 0.026558 |
| Mitogen-activated protein kinase 3 | Q63844 | 0.035166 | 0.458535 |
| SUMO-activating enzyme subunit 2 | Q9Z1F9 | 0.267437 | -3.51486 |
| Reticulon-3 | Q9ES97 | 0.301391 | -0.13843 |
| 60S acidic ribosomal protein P2 | P99027 | 0.457923 | 0.073594 |
| Cysteine sulfinic acid decarboxylase | Q9DBE0 | 0.510612 | 0.146876 |
| Peptidyl-prolyl cis-trans isomerase FKBP4;Peptidyl-prolyl cis-trar | P30416 | 0.685051 | 0.038171 |
| RNA-binding protein 14 | Q8C2Q3 | 0.918035 | -0.0157 |
| Choline transporter-like protein 2 | Q8BY89 | 0.676384 | 0.071211 |
| Microtubule-associated proteins 1A/1B light chain 3B | Q9CQV6 | 0.345207 | 0.301557 |
| DAZ-associated protein 1 | Q9JII5 | 0.501795 | 0.082746 |
| Translocon-associated protein subunit delta | Q62186 | 0.556446 | -0.07894 |

| | | | |
|---|---|---|---|
| N(G),N(G)-dimethylarginine dimethylaminohydrolase 2 | Q99LD8 | 0.792958 | -0.04438 |
| Splicing factor U2AF 35 kDa subunit | Q9D883 | 0.238169 | 0.361237 |
| Transient receptor potential cation channel subfamily V membe | Q8K424 | 0.124492 | -0.51556 |
| V-type proton ATPase subunit G 1 | Q9CR51 | 0.502841 | -0.17914 |
| Costars family protein ABRACL | Q4KML4 | 0.807286 | -0.05078 |
| Protein transport protein Sec61 subunit alpha isoform 1;Protein | P61620 | 0.563331 | 0.086851 |
| BH3-interacting domain death agonist;BH3-interacting domain d | P70444 | 0.468784 | 0.246163 |
| Guanine nucleotide-binding protein-like 1 | P36916 | 0.242821 | -0.62563 |
| Transcription factor BTF3 homolog 4 | Q9CQH7 | 0.712719 | 0.154977 |
| Ras-related protein Rab-5B | P61021 | 0.702228 | -0.06603 |
| Protein DEK | Q7TNV0 | 0.647556 | 0.140672 |
| 60S ribosomal protein L34 | Q9D1R9 | 0.720687 | 0.098971 |
| RNA binding protein fox-1 homolog 2 | Q8BP71 | 0.76624 | 0.058228 |
| AP-3 complex subunit beta-1 | Q9Z1T1 | 0.266987 | 0.137497 |
| Endoplasmic reticulum-Golgi intermediate compartment protein | Q9DC16 | 0.368127 | -0.10587 |
| Fascin | Q61553 | 0.783364 | 0.036332 |
| Ras-related protein Rap-1b;Ras-related protein Rap-1A | Q99JI6 | 0.015578 | 0.248442 |
| AP-1 complex subunit gamma-1 | P22892 | 0.178856 | 0.276079 |
| RNA-binding protein 8A | Q9CWZ3 | 0.493997 | 0.208957 |
| V-type proton ATPase catalytic subunit A | P50516 | 0.746497 | -0.02692 |
| ADP-ribosylation factor 4 | P61750 | 0.906998 | -0.01999 |
| Protein IMPACT | O55091 | 0.43225 | -0.1996 |
| Transcriptional activator GLI3;Transcriptional repressor GLI3R | Q61602 | 0.706012 | 0.162488 |
| RNA binding motif protein, X-linked-like-1;RNA-binding motif pr | Q91VM5 | 0.825851 | 0.039593 |
| 14-3-3 protein theta | P68254 | 0.725342 | 0.060818 |
| EGF-containing fibulin-like extracellular matrix protein 2 | Q9WVJ9 | 0.430174 | -0.07328 |
| 14-3-3 protein eta | P68510 | 0.986772 | 0.00435 |
| Splicing factor 1 | Q64213 | 0.042634 | 0.317937 |
| | E9Q7G0 | 0.809375 | -0.10578 |
| Pyridoxal-dependent decarboxylase domain-containing protein | Q99K01 | 0.203411 | -0.63586 |
| SWI/SNF-related matrix-associated actin-dependent regulator o | Q91ZW3 | 0.234409 | 0.409792 |
| Ganglioside-induced differentiation-associated protein 2 | Q9DBL2 | 0.297944 | 2.3434 |
| 28 kDa heat- and acid-stable phosphoprotein | Q3UHX2 | 0.799076 | 0.055321 |
| Vesicle-trafficking protein SEC22b | O08547 | 0.895766 | 0.025432 |
| Protein S100-A11 | P50543 | 0.701105 | 0.257889 |
| Zinc finger CCCH domain-containing protein 4 | Q6ZPZ3 | 0.654886 | 0.238363 |
| Annexin A6 | P14824 | 0.823083 | 0.03019 |
| Alpha-2-macroglobulin receptor-associated protein | P55302 | 0.580099 | 0.05091 |
| Non-histone chromosomal protein HMG-17 | P09602 | 0.61582 | 0.307948 |
| CD166 antigen | Q61490 | 0.972881 | 0.006196 |
| PAX3- and PAX7-binding protein 1 | P58501 | 0.778379 | 0.050105 |
| Ras-related protein Rab-5C | P35278 | 0.651911 | 0.099799 |
| Cation-independent mannose-6-phosphate receptor | Q07113 | 0.589383 | -0.11562 |
| RNA-binding protein FUS | P56959 | 0.889884 | -0.03042 |
| Ragulator complex protein LAMTOR3 | O88653 | 0.488086 | -0.13714 |
| Procollagen-lysine,2-oxoglutarate 5-dioxygenase 1 | Q9R0E2 | 0.828099 | 0.025365 |
| Ras-related protein Rab-1B | Q9D1G1 | 0.915208 | -0.01638 |
| COP9 signalosome complex subunit 6 | O88545 | 0.510336 | 0.072878 |
| Ras-related protein Rab-14 | Q91V41 | 0.607215 | -0.0687 |
| Enhancer of mRNA-decapping protein 4 | Q3UJB9 | 0.071601 | -0.73872 |

| | | | |
|---|---|---|---|
| Acidic leucine-rich nuclear phosphoprotein 32 family member A | O35381 | 0.274073 | 0.157909 |
| Serine/arginine-rich splicing factor 1 | Q6PDM2 | 0.583163 | -0.06425 |
| Enoyl-CoA hydratase, mitochondrial | Q8BH95 | 0.164722 | 0.167015 |
| Cytochrome c oxidase assembly factor 6 homolog | Q8BGD8 | 0.609498 | 0.072482 |
| Golgi apparatus protein 1 | Q61543 | 0.993534 | 0.000831 |
| 26S proteasome non-ATPase regulatory subunit 6 | Q99JI4 | 0.063871 | 0.360612 |
| 26S protease regulatory subunit 8 | P62196 | 0.441338 | 0.120763 |
| 60S ribosomal protein L3 | P27659 | 0.607857 | 0.042049 |
| Glyceraldehyde-3-phosphate dehydrogenase | P16858 | 0.652161 | -0.04265 |
| Microtubule-associated protein 4 | P27546 | 0.662041 | 0.05364 |
| Transportin-3 | Q6P2B1 | 0.20473 | -0.34502 |
| Platelet-activating factor acetylhydrolase IB subunit alpha | P63005 | 0.744078 | 0.062828 |
| Heat shock protein HSP 90-alpha | P07901 | 0.871359 | 0.018961 |
| Mammalian ependymin-related protein 1 | Q99M71 | 0.145541 | 0.189186 |
| Protein disulfide-isomerase TMX3 | Q8BXZ1 | 0.189359 | 0.551729 |
| Protein canopy homolog 3 | Q9DAU1 | 0.275877 | -0.40569 |
| Far upstream element-binding protein 1 | Q91WJ8 | 0.655773 | 0.185176 |
| 40S ribosomal protein SA | P14206 | 0.075192 | 0.214873 |
| NADH dehydrogenase [ubiquinone] 1 alpha subcomplex subunit | Q7TMF3 | 0.708515 | -0.15482 |
| Glutathione reductase, mitochondrial | P47791 | 0.179232 | 0.241746 |
| GDP-mannose 4,6 dehydratase | Q8K0C9 | 0.592437 | 0.065266 |
| Septin-8 | Q8CHH9 | 0.823274 | 0.082152 |
| Vinculin | Q64727 | 0.603862 | 0.041282 |
| Elongation factor 2 | P58252 | 0.288713 | 0.09099 |
| Ubiquilin-2 | Q9QZM0 | 0.802937 | -0.03581 |
| Aspartyl aminopeptidase | Q9Z2W0 | 0.767593 | -0.03595 |
| Sarcoplasmic/endoplasmic reticulum calcium ATPase 2 | O55143 | 0.432022 | -0.07135 |
| SWI/SNF complex subunit SMARCC2;SWI/SNF complex subunit S | Q6PDG5 | 0.505442 | -0.20403 |
| Talin-2 | Q71LX4 | 0.655028 | 0.122611 |
| H-2 class I histocompatibility antigen, D-B alpha chain;H-2 class I | P01899 | 0.006106 | 0.258686 |
| Aldose reductase-related protein 2 | P45377 | 0.676372 | -0.03025 |
| Ubiquitin-40S ribosomal protein S27a;Ubiquitin;40S ribosomal p | P62983 | 0.905184 | -0.02934 |
| Translationally-controlled tumor protein | P63028 | 0.857634 | 0.072428 |
| A-kinase anchor protein 12 | Q9WTQ5 | 0.185962 | 0.277708 |
| Partitioning defective 3 homolog | Q99NH2 | 0.838496 | 0.155455 |
| EH domain-binding protein 1 | Q69ZW3 | 0.128757 | 0.56207 |
| Eukaryotic translation initiation factor 3 subunit E | P60229 | 0.993423 | 0.001421 |
| 40S ribosomal protein S3a | P97351 | 0.307533 | 0.074989 |
| Thioredoxin domain-containing protein 5 | Q91W90 | 0.334103 | 0.159871 |
| Apoptosis regulator BAX | Q07813 | 0.346836 | 0.345861 |
| Transforming protein RhoA | Q9QUI0 | 0.8808 | 0.046242 |
| Malignant T-cell-amplified sequence 1;Malignant T-cell-amplifie | Q9DB27 | 0.346641 | 2.198254 |
| 60S ribosomal protein L37a | P61514 | 0.563493 | 0.061526 |
| 40S ribosomal protein S7 | P62082 | 0.634159 | 0.086411 |
| Thioredoxin domain-containing protein 12 | Q9CQU0 | 0.708373 | 0.053852 |
| | Q99J77 | 0.998227 | 0.000301 |
| Alpha-2-macroglobulin;Alpha-2-macroglobulin 165 kDa subunit; | Q61838 | 0.770755 | -0.06674 |
| Nucleophosmin | Q61937 | 0.070711 | 0.146588 |
| Group XV phospholipase A2 | Q8VEB4 | 0.090948 | 0.151941 |
| 40S ribosomal protein S15a | P62245 | 0.221497 | 0.108961 |

| | | | |
|---|---|---|---|
| Importin-5 | Q8BKC5 | 0.341211 | 0.141751 |
| Protein disulfide-isomerase A5 | Q921X9 | 0.57201 | 0.115509 |
| Aconitate hydratase, mitochondrial | Q99KI0 | 0.610424 | 0.034706 |
| Charged multivesicular body protein 1a | Q921W0 | 0.69901 | 0.203346 |
| Quinone oxidoreductase | P47199 | 0.823068 | -0.02223 |
| PRKC apoptosis WT1 regulator protein | Q925B0 | 0.724242 | 0.15064 |
| 1-phosphatidylinositol 4,5-bisphosphate phosphodiesterase delt | Q8R3B1 | 0.252269 | -0.18275 |
| AP-2 complex subunit mu | P84091 | 0.7593 | -0.02096 |
| Acid ceramidase;Acid ceramidase subunit alpha;Acid ceramidase | Q9WV54 | 0.077892 | 0.477168 |
| Glia maturation factor beta | Q9CQI3 | 0.821759 | 0.017452 |
| Cadherin-13 | Q9WTR5 | 0.212838 | 0.303237 |
| Pre-mRNA 3-end-processing factor FIP1 | Q9D824 | 0.903948 | -0.02189 |
| Maleylacetoacetate isomerase | Q9WVL0 | 0.265376 | -0.39324 |
| Structural maintenance of chromosomes protein 3 | Q9CW03 | 0.441913 | 1.780243 |
| GDP-fucose protein O-fucosyltransferase 2 | Q8VHI3 | 0.929348 | 0.021068 |
| Latexin | P70202 | 0.261236 | 0.10856 |
| Integrin-linked protein kinase | O55222 | 0.362258 | 0.237405 |
| Eukaryotic initiation factor 4A-I | P60843 | 0.748343 | 0.063111 |
| Transient receptor potential cation channel subfamily V membe | Q9WTR1 | 0.795422 | -0.05756 |
| RNA-binding protein 7 | Q9CQT2 | 0.81827 | -0.02342 |
| Puromycin-sensitive aminopeptidase | Q11011 | 0.834165 | -0.01592 |
| Erythrocyte band 7 integral membrane protein | P54116 | 0.676547 | 0.049331 |
| Cytochrome b-c1 complex subunit 2, mitochondrial | Q9DB77 | 0.293904 | 0.130156 |
| Transcription elongation factor B polypeptide 1 | P83940 | 0.702154 | -0.0381 |
| Destrin | Q9R0P5 | 0.583767 | 0.053859 |
| V-type proton ATPase subunit B, brain isoform | P62814 | 0.540544 | 0.076599 |
| Pleckstrin homology domain-containing family O member 2 | Q8K124 | 0.03382 | 0.411622 |
| ADP-dependent glucokinase | Q8VDL4 | 0.615798 | 0.157007 |
| 60S ribosomal protein L28 | P41105 | 0.700058 | 0.049656 |
| Collagen alpha-2(VI) chain | Q02788 | 0.40897 | -0.06176 |
| Glucosamine-6-phosphate isomerase 1 | O88958 | 0.882302 | 0.043233 |
| Coronin-1C | Q9WUM4 | 0.960707 | -0.00681 |
| Eukaryotic translation initiation factor 3 subunit D | O70194 | 0.891538 | 0.026853 |
| Proteolipid protein 2 | Q9R1Q7 | 0.494179 | 0.078425 |
| NADPH--cytochrome P450 reductase | P37040 | 0.65022 | 0.067948 |
| Peptidyl-prolyl cis-trans isomerase FKBP3 | Q62446 | 0.942185 | 0.010965 |
| LEM domain-containing protein 2 | Q6DVA0 | 0.608347 | -0.25088 |
| Alpha-methylacyl-CoA racemase | O09174 | 0.098406 | 0.332041 |
| S-formylglutathione hydrolase | Q9R0P3 | 0.300931 | 0.098595 |
| V-type proton ATPase subunit C 1 | Q9Z1G3 | 0.395485 | 0.096674 |
| KH domain-containing, RNA-binding, signal transduction-associa | Q60749 | 0.911292 | -0.07499 |
| Isochorismatase domain-containing protein 2A, mitochondrial | P85094 | 0.610051 | -0.16536 |
| Trimethyllysine dioxygenase, mitochondrial | Q91ZE0 | 0.727176 | 0.065638 |
| Erlin-2 | Q8BFZ9 | 0.74335 | -0.03286 |
| Protein BRICK1 | Q91VR8 | 0.766559 | 0.097377 |
| Putative N-acetylglucosamine-6-phosphate deacetylase | Q8JZV7 | 0.175801 | 0.17249 |
| Luc7-like protein 3 | Q5SUF2 | 0.667496 | 0.301406 |
| T-complex protein 1 subunit theta | P42932 | 0.696874 | 0.024661 |
| Annexin A2 | P07356 | 0.929083 | 0.009399 |
| Osteopetrosis-associated transmembrane protein 1 | Q8BGT0 | 0.50346 | 0.253941 |

| | | | |
|---|---|---|---|
| Hematological and neurological expressed 1 protein;Hematolog | P97825 | 0.803681 | 0.061826 |
| Importin subunit alpha-4 | O35344 | 0.936549 | 0.017984 |
| Acidic leucine-rich nuclear phosphoprotein 32 family member B | Q9EST5 | 0.38793 | 0.32437 |
| Eukaryotic translation initiation factor 4 gamma 1 | Q6NZJ6 | 0.444977 | 0.093491 |
| Small nuclear ribonucleoprotein Sm D2 | P62317 | 0.853872 | 0.034818 |
| Alpha-centractin | P61164 | 0.731783 | 0.079914 |
| Mitochondrial import receptor subunit TOM40 homolog | Q9QYA2 | 0.999914 | -2E-05 |
| E3 ubiquitin-protein ligase RNF5 | O35445 | 0.803451 | -0.72674 |
| Copine-1 | Q8C166 | 0.06941 | 0.217275 |
| Eukaryotic translation initiation factor 2 subunit 1 | Q6ZWX6 | 0.202647 | 0.279575 |
| Cytoskeleton-associated protein 5 | A2AGT5 | 0.556848 | -0.1287 |
| Glutathione S-transferase A4 | P24472 | 0.930693 | 0.010104 |
| Apolipoprotein O | Q9DCZ4 | 0.16976 | 0.230073 |
| Heterochromatin protein 1-binding protein 3 | Q3TEA8 | 0.082914 | 0.152073 |
| Calumenin | O35887 | 0.128952 | 0.115776 |
| Cytoskeleton-associated protein 4 | Q8BMK4 | 0.260925 | 0.115895 |
| Inositol monophosphatase 1 | O55023 | 0.313095 | 0.133224 |
| Eukaryotic translation initiation factor 6 | O55135 | 0.330268 | -0.16019 |
| Membrane-associated progesterone receptor component 2 | Q80UU9 | 0.33384 | -0.20994 |
| Sialate O-acetylesterase;Sialate O-acetylesterase small subunit;S | P70665 | 0.350393 | 0.477539 |
| Trifunctional purine biosynthetic protein adenosine-3;Phosphori | Q64737 | 0.363589 | -0.18725 |
| Elongation factor 1-delta | P57776 | 0.376197 | 0.090548 |
| Importin subunit beta-1 | P70168 | 0.475788 | 0.086741 |
| DNA-(apurinic or apyrimidinic site) lyase;DNA-(apurinic or apyrir | P28352 | 0.49935 | 0.110385 |
| Dipeptidyl peptidase 2 | Q9ET22 | 0.505999 | 0.078996 |
| Sepiapterin reductase | Q64105 | 0.603952 | 0.072981 |
| 6.8 kDa mitochondrial proteolipid | P56379 | 0.624843 | -0.0914 |
| Clathrin heavy chain 1 | Q68FD5 | 0.627515 | 0.039954 |
| Sorting nexin-12 | O70493 | 0.822143 | -0.04301 |
| Protein YIPF5 | Q9EQQ2 | 0.839385 | -0.57309 |
| Biliverdin reductase A | Q9CY64 | 0.488574 | -0.06029 |
| La-related protein 1 | Q6ZQ58 | 0.572349 | -0.10783 |
| Inositol 1,4,5-trisphosphate receptor type 3 | P70227 | 0.609054 | 0.125 |
| Protein disulfide-isomerase A3 | P27773 | 0.62466 | 0.04509 |
| 10 kDa heat shock protein, mitochondrial | Q64433 | 0.703227 | 0.049308 |
| H/ACA ribonucleoprotein complex subunit 4 | Q9ESX5 | 0.230441 | 0.248817 |
| Tripeptidyl-peptidase 2 | Q64514 | 0.920668 | 0.023718 |
| Glutaredoxin-1 | Q9QUH0 | 0.53487 | 0.080338 |
| Nucleoprotein TPR | F6ZDS4 | 0.651696 | -0.23805 |
| Programmed cell death protein 5 | P56812 | 0.893656 | 0.066535 |
| ATP-dependent Clp protease proteolytic subunit, mitochondrial | O88696 | 0.954271 | 0.008343 |
| Vacuolar protein sorting-associated protein 26A | P40336 | 0.596041 | 0.070525 |
| Cytosol aminopeptidase | Q9CPY7 | 0.742606 | 0.050337 |
| Disks large homolog 1 | Q811D0 | 0.754379 | 0.069364 |
| Reticulocalbin-2 | Q8BP92 | 0.795379 | 0.073024 |
| Protein S100-A10 | P08207 | 0.927045 | -0.00964 |
| Unconventional myosin-Ib | P46735 | 0.428452 | 0.161806 |
| 60S ribosomal protein L9 | P51410 | 0.440332 | 0.094612 |
| Annexin A11 | P97384 | 0.635056 | 0.062303 |
| THO complex subunit 4;Aly/REF export factor 2 | O08583 | 0.659255 | -0.05345 |

| | | | |
|---|---|---|---|
| MARCKS-related protein | P28667 | 0.081628 | 0.378788 |
| 60S ribosomal protein L12 | P35979 | 0.40292 | -0.10813 |
| Anoctamin-6 | Q6P9J9 | 0.326858 | 0.084417 |
| 40S ribosomal protein S26 | P62855 | 0.766939 | -0.05654 |
| Flavin reductase (NADPH) | Q923D2 | 0.968218 | 0.006465 |
| Nucleolar protein 58 | Q6DFW4 | 0.173363 | 0.185138 |
| Mitochondrial carrier homolog 1 | Q791T5 | 0.998621 | -0.00063 |
| UBX domain-containing protein 1 | Q922Y1 | 0.492426 | -0.14289 |
| Tubulin beta-2B chain;Tubulin beta-2A chain | Q9CWF2 | 0.708502 | 0.050799 |
| SUMO-conjugating enzyme UBC9 | P63280 | 0.87731 | 0.046339 |
| Presequence protease, mitochondrial | Q8K411 | 0.071501 | 0.187851 |
| COP9 signalosome complex subunit 7a | Q9CZ04 | 0.50815 | -0.24711 |
| Signal recognition particle 54 kDa protein | P14576 | 0.88883 | 0.0607 |
| Cyclic AMP-dependent transcription factor ATF-7 | Q8R0S1 | 0.746442 | 0.140002 |
| Dynamin-1 | P39053 | 0.798315 | -0.02909 |
| Insulin-like growth factor 2 mRNA-binding protein 1 | O88477 | 0.405572 | 0.13296 |
| Clathrin light chain A | O08585 | 0.50907 | 0.105358 |
| Guanine nucleotide-binding protein G(I)/G(S)/G(T) subunit beta- | P62880 | 0.316844 | -0.08323 |
| Death domain-containing membrane protein NRADD | Q8CJ26 | 0.644187 | -0.0817 |
| Radixin | P26043 | 0.692657 | -0.03222 |
| Ubiquitin-conjugating enzyme E2 variant 1 | Q9CZY3 | 0.828612 | -0.05113 |
| N(G),N(G)-dimethylarginine dimethylaminohydrolase 1 | Q9CWS0 | 0.831434 | -0.04141 |
| Transmembrane protein 263 | Q9DAM7 | 0.980349 | -0.00382 |
| Actin-related protein 2 | P61161 | 0.444444 | -0.13029 |
| Membrane primary amine oxidase | O70423 | 0.233846 | 0.228186 |
| Tissue factor | P20352 | 0.32163 | 0.355084 |
| 3-mercaptopyruvate sulfurtransferase | Q99J99 | 0.677007 | 0.072712 |
| Histone deacetylase complex subunit SAP18 | O55128 | 0.742056 | 0.129096 |
| 60S ribosomal protein L5 | P47962 | 0.300439 | 0.074571 |
| Adenylosuccinate lyase | P54822 | 0.527128 | 0.066752 |
| Guanine nucleotide-binding protein G(s) subunit alpha isoforms | P63094 | 0.548033 | -0.05933 |
| 40S ribosomal protein S11 | P62281 | 0.792645 | -0.03132 |
| 40S ribosomal protein S2 | P25444 | 0.810914 | 0.017271 |
| SAFB-like transcription modulator | Q8CH25 | 0.911492 | 0.305232 |
| 40S ribosomal protein S27 | Q6ZWU9 | 0.933897 | -0.01187 |
| Alpha-parvin | Q9EPC1 | 0.402423 | 0.205246 |
| Ras-related protein Rab-1A | P62821 | 0.137373 | 0.135973 |
| Eukaryotic translation initiation factor 2 subunit 3, X-linked;Euka | Q9Z0N1 | 0.942438 | -0.00789 |
| Protein lunapark | Q7TQ95 | 0.491841 | 0.103393 |
| Dual specificity mitogen-activated protein kinase kinase 1;Dual s | P31938 | 0.771862 | 0.050536 |
| Splicing factor U2AF 65 kDa subunit | P26369 | 0.976929 | 0.007288 |
| 60S ribosomal protein L36a | P83882 | 0.838347 | 0.044427 |
| Calpain-1 catalytic subunit | O35350 | 0.108769 | 0.221575 |
| Ubiquitin fusion degradation protein 1 homolog | P70362 | 0.23855 | -0.4883 |
| Ras-related C3 botulinum toxin substrate 1;Ras-related C3 botul | P63001 | 0.285255 | 0.112998 |
| Amine oxidase [flavin-containing] A | Q64133 | 0.598987 | -0.10483 |
| 60S ribosomal protein L31 | P62900 | 0.917022 | 0.009922 |
| Prolyl 3-hydroxylase 1 | Q3V1T4 | 0.294459 | 0.289441 |
| Alpha-N-acetylgalactosaminidase | Q9QWR8 | 0.355347 | 0.154165 |
| E3 ubiquitin-protein ligase NEDD4-like | Q8CFI0 | 0.568429 | 0.123326 |

| | | | |
|---|---|---|---|
| 40S ribosomal protein S9 | Q6ZWN5 | 0.738026 | 0.079222 |
| Ubiquitin carboxyl-terminal hydrolase isozyme L5 | Q9WUP7 | 0.901775 | 0.02754 |
| Splicing factor 3A subunit 1 | Q8K4Z5 | 0.709149 | 0.071674 |
| Protein quaking | Q9QYS9 | 0.221468 | 0.192816 |
| Uncharacterized protein FLJ45252 homolog | Q6PIU9 | 0.84402 | 0.033401 |
| T-complex protein 1 subunit epsilon | P80316 | 0.788865 | 0.038621 |
| Ubiquitin-like modifier-activating enzyme 1 | Q02053 | 0.799601 | 0.016507 |
| ATP-dependent 6-phosphofructokinase, platelet type | Q9WUA3 | 0.154811 | -0.19454 |
| Prenylcysteine oxidase | Q9CQF9 | 0.575113 | 0.070562 |
| SNW domain-containing protein 1 | Q9CSN1 | 0.115444 | 0.241462 |
| Calcium-binding mitochondrial carrier protein SCaMC-1 | Q8BMD8 | 0.616209 | 0.060906 |
| Nuclear factor NF-kappa-B p100 subunit;Nuclear factor NF-kapp | Q9WTK5 | 0.765338 | 0.062831 |
| Methionine aminopeptidase 2 | O08663 | 0.940991 | 0.018375 |
| Ras suppressor protein 1 | Q01730 | 0.731568 | -0.06385 |
| Heat shock protein HSP 90-beta | P11499 | 0.5197 | 0.090547 |
| Lon protease homolog, mitochondrial | Q8CGK3 | 0.395262 | -0.12431 |
| Actin-related protein 2/3 complex subunit 3 | Q9JM76 | 0.558352 | 0.097022 |
| Supervillin | Q8K4L3 | 0.571819 | 0.098914 |
| Glypican-1;Secreted glypican-1 | Q9QZF2 | 0.753999 | 0.070655 |
| 3-hydroxyacyl-CoA dehydrogenase type-2 | O08756 | 0.850885 | 0.015587 |
| Prothymosin alpha;Prothymosin alpha, N-terminally processed;T | P26350 | 0.898233 | 0.026631 |
| Sec1 family domain-containing protein 1 | Q8BRF7 | 0.95588 | -0.015 |
| Lysosomal acid phosphatase | P24638 | 0.049453 | -0.23009 |
| Nuclear ubiquitous casein and cyclin-dependent kinase substrat | Q80XU3 | 0.563378 | 0.165528 |
| Catenin beta-1 | Q02248 | 0.644727 | 0.095332 |
| Heterogeneous nuclear ribonucleoprotein U-like protein 2 | Q00PI9 | 0.78191 | 0.032212 |
| Guanine nucleotide-binding protein G(I)/G(S)/G(T) subunit beta- | Q61011 | 0.953364 | 0.16538 |
| Microtubule-associated protein 1S;MAP1S heavy chain;MAP1S l | Q8C052 | 0.761071 | -0.06064 |
| Histone deacetylase 1;Histone deacetylase 2 | O09106 | 0.204049 | 0.254188 |
| Reticulocalbin-1 | Q05186 | 0.179527 | 0.103881 |
| ATPase family AAA domain-containing protein 1 | Q9D5T0 | 0.569538 | 0.37103 |
| Histone-binding protein RBBP4;Histone-binding protein RBBP7 | Q60972 | 0.611453 | 0.111641 |
| Ragulator complex protein LAMTOR5 | Q9D1L9 | 0.644115 | 0.079119 |
| G-protein coupled receptor 124 | Q91ZV8 | 0.690061 | -0.33962 |
| Beta-hexosaminidase subunit alpha | P29416 | 0.704263 | 0.047374 |
| Aldehyde dehydrogenase, cytosolic 1 | O35945 | 0.88213 | 0.0253 |
| T-complex protein 1 subunit delta | P80315 | 0.8864 | 0.019611 |
| Tubulin beta-4B chain;Tubulin beta-4A chain | P68372 | 0.921382 | 0.013944 |
| Lupus La protein homolog | P32067 | 0.936592 | 0.022659 |
| Histidine triad nucleotide-binding protein 2, mitochondrial | Q9D0S9 | 0.943148 | -0.01656 |
| Dynein light chain roadblock-type 1 | P62627 | 0.240032 | 0.241238 |
| Transmembrane emp24 domain-containing protein 2 | Q9R0Q3 | 0.587348 | 0.174455 |
| PTB domain-containing engulfment adapter protein 1 | Q8K2A1 | 0.952522 | -0.00654 |
| 40S ribosomal protein S28 | P62858 | 0.775833 | -0.05971 |
| Protein CDV3 | Q4VAA2 | 0.949631 | 0.007592 |
| Collagen alpha-2(IV) chain;Canstatin | P08122 | 0.997287 | -0.00027 |
| Plasma membrane calcium-transporting ATPase 1;Calcium-trans | G5E829 | 0.574183 | 0.159684 |
| Serine--tRNA ligase, cytoplasmic | P26638 | 0.966309 | -0.00789 |
| STIP1 homology and U box-containing protein 1 | Q9WUD1 | 0.8017 | 0.036499 |
| Plectin | Q9QXS1 | 0.908761 | 0.010124 |

| | | | |
|---|---|---|---|
| Histone H2A type 2-C;Histone H2A type 2-A;Histone H2A type 2- | Q64523 | 0.227042 | 0.101763 |
| Exportin-1 | Q6P5F9 | 0.087619 | 0.359738 |
| Dolichyl-diphosphooligosaccharide--protein glycosyltransferase | P46978 | 0.260794 | 0.179653 |
| Platelet-derived growth factor receptor beta | P05622 | 0.435982 | 0.092997 |
| Signal peptide peptidase-like 2A | Q9JJF9 | 0.582962 | -0.28336 |
| UMP-CMP kinase | Q9DBP5 | 0.61407 | -0.03839 |
| Delta-sarcoglycan | P82347 | 0.67392 | 0.125007 |
| Adenylate kinase isoenzyme 1 | Q9R0Y5 | 0.687276 | 0.046972 |
| Non-histone chromosomal protein HMG-14 | P18608 | 0.72204 | -0.04859 |
| NADH dehydrogenase [ubiquinone] 1 alpha subcomplex subunit | Q9DC69 | 0.760103 | 0.033869 |
| Four and a half LIM domains protein 2 | O70433 | 0.771681 | -0.09806 |
| Pre-mRNA-splicing factor ATP-dependent RNA helicase DHX15 | O35286 | 0.265171 | 0.158849 |
| Fumarylacetoacetase | P35505 | 0.301558 | 0.14955 |
| Casein kinase II subunit alpha | O54833 | 0.373281 | -0.36313 |
| Small nuclear ribonucleoprotein Sm D3 | P62320 | 0.53388 | -0.14069 |
| Splicing factor, proline- and glutamine-rich | Q8VIJ6 | 0.56388 | 0.041638 |
| UPF0160 protein MYG1, mitochondrial | Q9JK81 | 0.573686 | -0.12125 |
| SH3 domain-binding glutamic acid-rich-like protein | Q9JJU8 | 0.604906 | -0.0637 |
| Dynactin subunit 1 | O08788 | 0.867761 | -0.04527 |
| Nicastrin | P57716 | 0.67619 | -0.05972 |
| Eukaryotic translation initiation factor 3 subunit C | Q8R1B4 | 0.77812 | 0.026957 |
| NADH-ubiquinone oxidoreductase 75 kDa subunit, mitochondria | Q91VD9 | 0.56346 | 0.084623 |
| Kinectin | Q61595 | 0.834811 | 0.190599 |
| Heme oxygenase 1 | P14901 | 0.579102 | 0.074726 |
| Heterogeneous nuclear ribonucleoproteins C1/C2 | Q9Z204 | 0.683719 | 0.068876 |
| ATP-dependent 6-phosphofructokinase, liver type | P12382 | 0.813766 | -0.03526 |
| Apoptotic chromatin condensation inducer in the nucleus | Q9JIX8 | 0.972643 | -0.00851 |
| Catenin delta-1 | P30999 | 0.833101 | 0.044025 |
| Periodic tryptophan protein 2 homolog | Q8BU03 | 0.844367 | 0.023499 |
| AP-1 complex subunit sigma-1A | P61967 | 0.166563 | 0.671585 |
| Protein arginine N-methyltransferase 1 | Q9JIF0 | 0.467742 | -0.21654 |
| Xaa-Pro aminopeptidase 1 | Q6P1B1 | 0.248152 | 0.203474 |
| Alpha-adducin | Q9QYC0 | 0.736999 | 0.060765 |
| Leiomodin-2 | Q3UHZ5 | 0.737998 | 0.061815 |
| Ras GTPase-activating protein-binding protein 1 | P97855 | 0.49963 | -0.10973 |
| ADP-ribosylation factor-like protein 3 | Q9WUL7 | 0.763521 | 0.074714 |
| E3 ubiquitin-protein ligase HUWE1 | Q7TMY8 | 0.872884 | -0.07573 |
| 60S ribosomal protein L15 | Q9CZM2 | 0.555515 | 0.061438 |
| NADH-cytochrome b5 reductase 1 | Q9DB73 | 0.201407 | 0.145572 |
| Ubiquitin carboxyl-terminal hydrolase isozyme L3;Ubiquitin carb | Q9JKB1 | 0.59932 | 0.080735 |
| Myeloid-derived growth factor | Q9CPT4 | 0.813098 | 0.02563 |
| Glutathione S-transferase P 1 | P19157 | 0.94143 | 0.011143 |
| Propionyl-CoA carboxylase alpha chain, mitochondrial | Q91ZA3 | 0.950929 | -0.01167 |
| Argininosuccinate lyase | Q91YI0 | 0.95158 | -0.01284 |
| Activated RNA polymerase II transcriptional coactivator p15 | P11031 | 0.974733 | 0.004613 |
| 26S proteasome non-ATPase regulatory subunit 11 | Q8BG32 | 0.988871 | -0.002 |
| Tyrosine-protein phosphatase non-receptor type 23 | Q6PB44 | 0.85224 | -0.02424 |
| Crk-like protein | P47941 | 0.974518 | 0.009969 |
| Voltage-dependent anion-selective channel protein 2 | Q60930 | 0.016451 | 0.250866 |
| 40S ribosomal protein S14 | P62264 | 0.281808 | -0.10381 |

| | | | |
|---|---|---|---|
| Bcl-2 homologous antagonist/killer | Q08734 | 0.28627 | -0.41699 |
| Peptidyl-prolyl cis-trans isomerase FKBP9 | Q9Z247 | 0.555956 | 0.122026 |
| 40S ribosomal protein S5;40S ribosomal protein S5, N-terminally | P97461 | 0.981223 | 0.002389 |
| 26S protease regulatory subunit 10B | P62334 | 0.903907 | 0.025548 |
| Thioredoxin | P10639 | 0.438449 | 0.09481 |
| Beta-2-syntrophin | Q61235 | 0.321184 | -0.14519 |
| 6-phosphogluconolactonase | Q9CQ60 | 0.427716 | -0.14668 |
| Beta-centractin | Q8R5C5 | 0.639695 | -0.14453 |
| Unconventional myosin-Ic | Q9WTI7 | 0.347744 | 0.11087 |
| Cytochrome c oxidase subunit 6B1 | P56391 | 0.302034 | -0.20953 |
| MAP7 domain-containing protein 1 | A2AJI0 | 0.617786 | 0.100644 |
| Heterogeneous nuclear ribonucleoprotein L | Q8R081 | 0.802612 | 0.042646 |
| Transgelin-2 | Q9WVA4 | 0.574475 | 0.05584 |
| GTP:AMP phosphotransferase AK3, mitochondrial | Q9WTP7 | 0.816964 | -0.02835 |
| Dynein light chain 2, cytoplasmic | Q9D0M5 | 0.418541 | -0.27107 |
| 60S ribosomal protein L14 | Q9CR57 | 0.75206 | -0.05916 |
| Nuclear pore complex protein Nup93 | Q8BJ71 | 0.816012 | -0.03283 |
| Rho-related GTP-binding protein RhoG | P84096 | 0.910985 | -0.05115 |
| Cullin-7 | Q8VE73 | 0.911698 | -0.03855 |
| Insulin-degrading enzyme | Q9JHR7 | 0.983837 | 0.005837 |
| Proteasome subunit alpha type-5 | Q9Z2U1 | 0.454443 | -0.06791 |
| Glycine--tRNA ligase | Q9CZD3 | 0.961835 | 0.004807 |
| Serine/arginine-rich splicing factor 6 | Q3TWW8 | 0.320368 | -0.15056 |
| CAP-Gly domain-containing linker protein 2 | Q9Z0H8 | 0.183092 | -0.43605 |
| Transmembrane protein 106B | Q80X71 | 0.751368 | 0.030934 |
| Protein-lysine 6-oxidase | P28301 | 0.900003 | -0.02748 |
| Acyl-coenzyme A thioesterase 9, mitochondrial;Acyl-coenzyme A | Q9R0X4 | 0.270247 | -0.16517 |
| Valine--tRNA ligase | Q9Z1Q9 | 0.645742 | 0.061132 |
| Carbonyl reductase [NADPH] 1 | P48758 | 0.737825 | 0.053826 |
| Estradiol 17-beta-dehydrogenase 11 | Q9EQ06 | 0.373789 | 0.207504 |
| N(4)-(beta-N-acetylglucosaminyl)-L-asparaginase;Glycosylaspara | Q64191 | 0.732651 | -0.07131 |
| Probable ATP-dependent RNA helicase DDX5 | Q61656 | 0.96139 | 0.005925 |
| Sphingosine-1-phosphate lyase 1 | Q8R0X7 | 0.583466 | 0.073312 |
| Neudesin | Q9CQ45 | 0.229709 | 0.206174 |
| SUN domain-containing protein 2 | Q8BJS4 | 0.343944 | 0.17003 |
| Proliferation-associated protein 2G4 | P50580 | 0.439592 | 0.103155 |
| Vacuolar protein sorting-associated protein 29 | Q9QZ88 | 0.470481 | -0.08124 |
| Transcriptional activator protein Pur-alpha | P42669 | 0.472257 | 0.110802 |
| Calcium-binding mitochondrial carrier protein Aralar1 | Q8BH59 | 0.575175 | 0.093083 |
| Endoplasmic reticulum resident protein 44 | Q9D1Q6 | 0.582731 | 0.077569 |
| Procollagen galactosyltransferase 1 | Q8K297 | 0.88014 | 0.038857 |
| Translocon-associated protein subunit gamma | Q9DCF9 | 0.912815 | 0.031706 |
| U6 snRNA-associated Sm-like protein LSm2 | O35900 | 0.209553 | 0.492572 |
| Cathepsin D | P18242 | 0.544211 | -0.10934 |
| Ras-related protein Rab-18 | P35293 | 0.632766 | 0.099653 |
| Collagen alpha-1(III) chain | P08121 | 0.736325 | 0.024039 |
| rRNA 2-O-methyltransferase fibrillarin | P35550 | 0.238927 | 0.142734 |
| Protein NipSnap homolog 2 | O55126 | 0.397977 | 0.096145 |
| Secretory carrier-associated membrane protein 3 | O35609 | 0.415962 | -0.13962 |
| Heterogeneous nuclear ribonucleoprotein A3 | Q8BG05 | 0.616133 | 0.111683 |

| | | | |
|---|---|---|---|
| FERM, RhoGEF and pleckstrin domain-containing protein 1 | F8VPU2 | 0.625757 | -0.20302 |
| BolA-like protein 1 | Q9D8S9 | 0.703965 | 0.115538 |
| Parathymosin | Q9D0J8 | 0.937926 | -0.01809 |
| RuvB-like 1 | P60122 | 0.553512 | -0.14075 |
| 60S ribosomal protein L36 | P47964 | 0.095298 | 0.296033 |
| Cytochrome c oxidase subunit 5B, mitochondrial | P19536 | 0.75018 | -0.04188 |
| Plasminogen activator inhibitor 1 RNA-binding protein | Q9CY58 | 0.677745 | -0.09645 |
| Heterogeneous nuclear ribonucleoprotein A1;Heterogeneous nu | P49312 | 0.684504 | 0.031073 |
| D-3-phosphoglycerate dehydrogenase | Q61753 | 0.87362 | 0.012675 |
| Basic leucine zipper and W2 domain-containing protein 1 | Q9CQC6 | 0.094587 | 0.380832 |
| Myotrophin | P62774 | 0.420207 | 2.360977 |
| ATP-dependent RNA helicase DDX3X;ATP-dependent RNA helica | Q62167 | 0.700838 | -0.03823 |
| Diablo homolog, mitochondrial | Q9JIQ3 | 0.848613 | 0.049832 |
| Transmembrane and coiled-coil domain-containing protein 1 | Q921L3 | 0.019883 | -0.64658 |
| Isoleucine--tRNA ligase, cytoplasmic | Q8BU30 | 0.488913 | 0.078358 |
| Core histone macro-H2A.1 | Q9QZQ8 | 0.62193 | -0.05599 |
| Tumor protein D54 | Q9CYZ2 | 0.367445 | 0.12926 |
| Extended synaptotagmin-2 | Q3TZZ7 | 0.149853 | -0.2891 |
| Rab GDP dissociation inhibitor alpha | P50396 | 0.009416 | 0.34971 |
| Membrane-associated progesterone receptor component 1 | O55022 | 0.127773 | 0.270559 |
| Glucose-6-phosphate 1-dehydrogenase X | Q00612 | 0.340581 | -0.09369 |
| Stathmin | P54227 | 0.391602 | 0.194807 |
| Poly(U)-binding-splicing factor PUF60 | Q3UEB3 | 0.634351 | -0.12234 |
| Putative hydrolase RBBP9 | O88851 | 0.703061 | 0.065697 |
| 60S ribosomal protein L8 | P62918 | 0.828329 | -0.02052 |
| Charged multivesicular body protein 4b | Q9D8B3 | 0.893706 | -0.03159 |
| Hepatoma-derived growth factor | P51859 | 0.716405 | 0.054183 |
| Actin filament-associated protein 1 | Q80YS6 | 0.111643 | 0.492161 |
| Ras-related GTP-binding protein C | Q99K70 | 0.891783 | -0.02339 |
| Fragile X mental retardation syndrome-related protein 1 | Q61584 | 0.515196 | -0.09361 |
| CDGSH iron-sulfur domain-containing protein 1 | Q91WS0 | 0.890621 | -0.0369 |
| ERO1-like protein alpha | Q8R180 | 0.184294 | 0.178208 |
| ADP-ribosylation factor-like protein 8B | Q9CQW2 | 0.592827 | 0.068959 |
| Serine/threonine-protein phosphatase 2A activator | P58389 | 0.617947 | 0.079615 |
| Vesicle-associated membrane protein-associated protein A | Q9WV55 | 0.639756 | 0.032494 |
| Pigment epithelium-derived factor | P97298 | 0.707738 | 0.245594 |
| Adenine phosphoribosyltransferase | P08030 | 0.749781 | 0.058655 |
| Serrate RNA effector molecule homolog | Q99MR6 | 0.845746 | 0.057099 |
| Annexin A7 | Q07076 | 0.879752 | 0.013778 |
| ATP synthase subunit g, mitochondrial | Q9CPQ8 | 0.885239 | -0.03124 |
| EH domain-containing protein 1 | Q9WVK4 | 0.863066 | -0.01362 |
| Up-regulated during skeletal muscle growth protein 5 | Q78IK2 | 0.334996 | 0.205295 |
| Regulator of nonsense transcripts 1 | Q9EPU0 | 0.554819 | 0.09335 |
| Transcriptional activator protein Pur-beta | O35295 | 0.54857 | 0.074513 |
| Polymerase I and transcript release factor | O54724 | 0.685225 | 0.042969 |
| Dolichyl-diphosphooligosaccharide--protein glycosyltransferase | Q9DBG6 | 0.816964 | 0.020634 |
| Proteasome subunit alpha type-2 | P49722 | 0.466826 | -0.07654 |
| MICOS complex subunit Mic27 | Q78IK4 | 0.58286 | -0.05253 |
| Histone H2B type 1-C/E/G;Histone H2B type 2-B;Histone H2B ty | Q6ZWY9 | 0.864278 | 0.060263 |
| Single-stranded DNA-binding protein, mitochondrial | Q9CYR0 | 0.031579 | 0.460061 |

| | | | |
|---|---|---|---|
| NADH dehydrogenase [ubiquinone] iron-sulfur protein 3, mitocł | Q9DCT2 | 0.086817 | -0.44115 |
| Inactive tyrosine-protein kinase 7 | Q8BKG3 | 0.303706 | -0.17974 |
| Cathepsin L1;Cathepsin L1 heavy chain;Cathepsin L1 light chain | P06797 | 0.349598 | 0.233251 |
| Septin-5 | Q9Z2Q6 | 0.389242 | 0.214333 |
| Ubiquitin carboxyl-terminal hydrolase 14 | Q9JMA1 | 0.393004 | 0.189106 |
| Caspase-3;Caspase-3 subunit p17;Caspase-3 subunit p12 | P70677 | 0.417623 | 0.153641 |
| WW domain-binding protein 11 | Q923D5 | 0.498235 | 0.359505 |
| Collagen alpha-2(I) chain | Q01149 | 0.691121 | 0.064169 |
| Reticulon-4 | Q99P72 | 0.723128 | 0.032393 |
| Protein LYRIC | Q80WJ7 | 0.818695 | 0.037489 |
| UPF0568 protein C14orf166 homolog | Q9CQE8 | 0.852378 | 0.018056 |
| 60S ribosomal protein L23a | P62751 | 0.983443 | 0.001557 |
| Atlastin-3 | Q91YH5 | 0.671882 | 0.035229 |
| TSC22 domain family protein 3 | Q9Z2S7 | 0.124307 | -0.54357 |
| Tropomyosin alpha-4 chain | Q6IRU2 | 0.539746 | 0.161505 |
| Chromodomain-helicase-DNA-binding protein 4;Chromodomain | Q6PDQ2 | 0.233646 | 0.268404 |
| Phosphatidylinositide phosphatase SAC1 | Q9EP69 | 0.287946 | 0.120894 |
| Palmitoyl-protein thioesterase 1 | O88531 | 0.44865 | 0.113387 |
| NADH dehydrogenase [ubiquinone] 1 alpha subcomplex subunit | Q9Z1P6 | 0.924645 | -0.02666 |
| RNA-binding protein EWS | Q61545 | 0.368936 | 0.127774 |
| Leucine--tRNA ligase, cytoplasmic | Q8BMJ2 | 0.380558 | -0.07771 |
| Eukaryotic translation initiation factor 3 subunit L | Q8QZY1 | 0.448591 | 0.148972 |
| Obg-like ATPase 1 | Q9CZ30 | 0.490859 | 0.057392 |
| Eukaryotic translation initiation factor 4B | Q8BGD9 | 0.960575 | -0.01314 |
| 60S ribosomal protein L10;60S ribosomal protein L10-like | Q6ZWV3 | 0.982662 | -0.00374 |
| Band 4.1-like protein 3;Band 4.1-like protein 3, N-terminally pro | Q9WV92 | 0.36068 | 0.146156 |
| Glutamate dehydrogenase 1, mitochondrial | P26443 | 0.450294 | 0.090019 |
| Torsin-1A-interacting protein 1 | Q921T2 | 0.432569 | 0.106544 |
| Bifunctional ATP-dependent dihydroxyacetone kinase/FAD-AMF | Q8VC30 | 0.494851 | 2.007592 |
| T-complex protein 1 subunit beta | P80314 | 0.624416 | 0.045109 |
| Dolichyl-diphosphooligosaccharide--protein glycosyltransferase | Q91YQ5 | 0.851379 | 0.025504 |
| 40S ribosomal protein S12 | P63323 | 0.651983 | 0.033109 |
| V-type proton ATPase subunit d 1 | P51863 | 0.88978 | 0.013186 |
| Prefoldin subunit 6 | Q03958 | 0.127557 | -0.24656 |
| Dynactin subunit 4 | Q8CBY8 | 0.159468 | 0.42258 |
| Eukaryotic translation initiation factor 3 subunit F | Q9DCH4 | 0.66757 | -0.13239 |
| Galectin-3 | P16110 | 0.996094 | -0.0005 |
| E3 ubiquitin-protein ligase TRIM23 | Q8BGX0 | 0.748078 | -0.08988 |
| Calpastatin | P51125 | 0.775119 | 0.030744 |
| Xanthine dehydrogenase/oxidase;Xanthine dehydrogenase;Xant | Q00519 | 0.823509 | 0.02355 |
| Golgi phosphoprotein 3 | Q9CRA5 | 0.494559 | 0.180961 |
| Histidine--tRNA ligase, cytoplasmic;Probable histidine--tRNA liga | Q61035 | 0.569191 | 0.097871 |
| Calcium/calmodulin-dependent protein kinase type II subunit de | Q6PHZ2 | 0.80618 | -0.02826 |
| RNA-binding protein Raly | Q64012 | 0.821466 | -0.03846 |
| 40S ribosomal protein S6 | P62754 | 0.728639 | -0.02423 |
| Charged multivesicular body protein 2b | Q8BJF9 | 0.390787 | 0.184194 |
| Transmembrane 9 superfamily member 2 | P58021 | 0.505163 | -0.1647 |
| Heterogeneous nuclear ribonucleoprotein A0 | Q9CX86 | 0.29416 | 0.138564 |
| Ras-related protein Rab-6A;Ras-related protein Rab-6B | P35279 | 0.32355 | -0.16285 |
| Small nuclear ribonucleoprotein F | P62307 | 0.5688 | -0.06263 |

| | | | |
|---|---|---|---|
| Small nuclear ribonucleoprotein G | P62309 | 0.882997 | 0.034688 |
| Peptidyl-prolyl cis-trans isomerase A;Peptidyl-prolyl cis-trans iso | P17742 | 0.959791 | -0.00738 |
| Galectin-1 | P16045 | 0.494241 | 0.046247 |
| Synaptophysin-like protein 1 | O09117 | 0.248467 | 0.287611 |
| 60S ribosomal protein L27 | P61358 | 0.263826 | 0.131538 |
| Splicing factor 3B subunit 3 | Q921M3 | 0.286689 | 0.16153 |
| Cullin-5 | Q9D5V5 | 0.354473 | 2.941298 |
| Lipid phosphate phosphohydrolase 3 | Q99JY8 | 0.450026 | -0.12366 |
| Nucleolar protein 56 | Q9D6Z1 | 0.471993 | -0.1332 |
| Transmembrane emp24 domain-containing protein 4 | Q8R1V4 | 0.472606 | 0.39183 |
| V-type proton ATPase 16 kDa proteolipid subunit | P63082 | 0.51979 | -0.15326 |
| Dehydrogenase/reductase SDR family member 7 | Q9CXR1 | 0.580096 | 0.137716 |
| 3-ketoacyl-CoA thiolase A, peroxisomal;3-ketoacyl-CoA thiolase | Q921H8 | 0.683606 | -0.06239 |
| V-type proton ATPase 116 kDa subunit a isoform 1 | Q9Z1G4 | 0.693348 | -0.03347 |
| Pyruvate kinase PKM | P52480 | 0.726143 | 0.030785 |
| Protein-methionine sulfoxide oxidase MICAL1 | Q8VDP3 | 0.878476 | -0.02273 |
| Matrix-remodeling-associated protein 7 | Q9CZH7 | 0.884171 | -0.03215 |
| High mobility group protein HMGI-C | P52927 | 0.906447 | -0.05183 |
| V-type proton ATPase subunit E 1 | P50518 | 0.91552 | -0.00911 |
| Coatomer subunit beta | O55029 | 0.947908 | 0.008426 |
| Protein disulfide-isomerase A6 | Q922R8 | 0.976872 | -0.00448 |
| 60S ribosomal protein L29 | P47915 | 0.998484 | 0.00059 |
| 2-amino-3-ketobutyrate coenzyme A ligase, mitochondrial | O88986 | 0.637272 | 0.120953 |
| 40S ribosomal protein S17 | P63276 | 0.396826 | -0.13244 |
| F-actin-capping protein subunit alpha-2 | P47754 | 0.869429 | 0.024955 |
| Importin subunit alpha-3 | O35343 | 0.936263 | -0.01791 |
| Proline synthase co-transcribed bacterial homolog protein | Q9Z2Y8 | 0.249513 | -0.31841 |
| Acyl-coenzyme A thioesterase 2, mitochondrial;Acyl-coenzyme / | Q9QYR9 | 0.250902 | 0.227343 |
| UBX domain-containing protein 4 | Q8VCH8 | 0.368287 | -0.45056 |
| RuvB-like 2 | Q9WTM5 | 0.676884 | 0.082204 |
| Thioredoxin-related transmembrane protein 1 | Q8VBT0 | 0.693319 | -0.10895 |
| AP-1 complex subunit beta-1 | O35643 | 0.708299 | -0.06867 |
| Nuclear transport factor 2 | P61971 | 0.937107 | 0.021331 |
| DNA topoisomerase 1 | Q04750 | 0.318662 | 0.179903 |
| Microtubule-associated protein 1A;MAP1A heavy chain;MAP1 li | Q9QYR6 | 0.391633 | 0.308501 |
| Ubiquitin carboxyl-terminal hydrolase 5 | P56399 | 0.430448 | 0.060816 |
| 60S ribosomal protein L4 | Q9D8E6 | 0.619723 | 0.053224 |
| PEST proteolytic signal-containing nuclear protein | Q6P8I4 | 0.632554 | 0.113968 |
| Lysosomal alpha-glucosidase | P70699 | 0.772364 | 0.030496 |
| Guanine nucleotide-binding protein G(I)/G(S)/G(T) subunit beta- | P62874 | 0.546387 | 0.079579 |
| 40S ribosomal protein S15 | P62843 | 0.213922 | 0.141889 |
| Probable ATP-dependent RNA helicase DDX17 | Q501J6 | 0.282557 | 0.098765 |
| NEDD8-conjugating enzyme Ubc12 | P61082 | 0.451116 | -0.07702 |
| Histone H1.0;Histone H1.0, N-terminally processed | P10922 | 0.682249 | 0.084876 |
| 60S ribosomal protein L32 | P62911 | 0.094691 | -0.16042 |
| 60S ribosomal protein L26 | P61255 | 0.431646 | -0.0746 |
| Coatomer subunit beta | Q9JIF7 | 0.091742 | 0.147385 |
| 60S acidic ribosomal protein P0 | P14869 | 0.147302 | 0.170766 |
| 40S ribosomal protein S18 | P62270 | 0.815678 | -0.03 |
| Heterogeneous nuclear ribonucleoprotein L-like | Q921F4 | 0.056395 | 0.165293 |

| | | | |
|---|---|---|---|
| LIM and senescent cell antigen-like-containing domain protein 1 | Q99JW4 | 0.534718 | 0.113981 |
| Inosine-5-monophosphate dehydrogenase 2 | P24547 | 0.64919 | 0.048715 |
| Coatomer subunit delta | Q5XJY5 | 0.900738 | 0.010897 |
| Peroxiredoxin-2 | Q61171 | 0.763547 | 0.0257 |
| Transcription elongation factor A protein 1 | P10711 | 0.932463 | 0.011943 |
| Complement C3;Complement C3 beta chain;C3-beta-c;Complem | P01027 | 0.977043 | -0.00476 |
| Protein CutA | Q9CQ89 | 0.161188 | -0.71258 |
| Phosphatidylinositol transfer protein beta isoform | P53811 | 0.629431 | -0.12642 |
| ATPase inhibitor, mitochondrial | O35143 | 0.823198 | 0.089446 |
| ADP-ribosylation factor GTPase-activating protein 1 | Q9EPJ9 | 0.65709 | -0.26476 |
| NEDD8-activating enzyme E1 regulatory subunit | Q8VBW6 | 0.278374 | -2.34746 |
| Glutaredoxin-3 | Q9CQM9 | 0.385744 | 0.346811 |
| Katanin p60 ATPase-containing subunit A-like 2;Spermatogenesi | Q9D3R6 | 0.451945 | -0.1073 |
| 60S ribosomal protein L13a | P19253 | 0.50684 | 0.062472 |
| Sequestosome-1 | Q64337 | 0.056376 | 0.515834 |
| Neuroplastin | P97300 | 0.090283 | 0.191128 |
| 14-3-3 protein gamma;14-3-3 protein gamma, N-terminally proc | P61982 | 0.997543 | -0.00037 |
| Acylglycerol kinase, mitochondrial | Q9ESW4 | 0.999267 | -0.00017 |
| Importin-4 | Q8VI75 | 0.866506 | -0.07861 |
| 40S ribosomal protein S4, X isoform | P62702 | 0.223262 | 0.129851 |
| 60S ribosomal protein L10a | P53026 | 0.863797 | 0.023244 |
| Heterogeneous nuclear ribonucleoprotein A/B | Q99020 | 0.387397 | 0.068848 |
| 60S ribosomal protein L11 | Q9CXW4 | 0.850145 | 0.02061 |
| Proteasome subunit alpha type-3 | O70435 | 0.988627 | -0.00237 |
| Eukaryotic translation initiation factor 3 subunit A | P23116 | 0.532262 | 0.053719 |
| Gamma-glutamyl hydrolase | Q9Z0L8 | 0.634615 | 0.050201 |
| Omega-amidase NIT2 | Q9JHW2 | 0.782663 | -0.02711 |
| 60S ribosomal protein L35 | Q6ZWV7 | 0.888565 | 0.024012 |
| ADP-ribosylation factor GTPase-activating protein 2 | Q99K28 | 0.51261 | -0.24447 |
| Ribose-5-phosphate isomerase | P47968 | 0.603795 | 0.087221 |
| Peptidyl-prolyl cis-trans isomerase FKBP7 | O54998 | 0.146598 | 0.15429 |
| Hexokinase-1 | P17710 | 0.173589 | 0.129603 |
| Pre-mRNA-processing-splicing factor 8 | Q99PV0 | 0.1768 | 0.255708 |
| 40S ribosomal protein S23 | P62267 | 0.7579 | 0.027349 |
| Vesicle-associated membrane protein 8 | O70404 | 0.815406 | -0.01616 |
| Ras-related GTP-binding protein A;Ras-related GTP-binding prot | Q80X95 | 0.948968 | 0.011065 |
| V-type proton ATPase subunit D | P57746 | 0.174801 | 0.319727 |
| Barrier-to-autointegration factor;Barrier-to-autointegration fact | O54962 | 0.346289 | 0.111901 |
| Transformer-2 protein homolog beta | P62996 | 0.859137 | -0.0195 |
| SAP domain-containing ribonucleoprotein | Q9D1J3 | 0.970676 | 0.008768 |
| 60S ribosomal protein L38 | Q9JJI8 | 0.980154 | -0.00487 |
| Serine-threonine kinase receptor-associated protein | Q9Z1Z2 | 0.183252 | -0.23781 |
| | E9PVA8 | 0.221305 | 0.281557 |
| 26S proteasome non-ATPase regulatory subunit 8 | Q9CX56 | 0.2517 | 0.142671 |
| Phosphoglycolate phosphatase | Q8CHP8 | 0.295133 | -0.18213 |
| Twinfilin-1 | Q91YR1 | 0.426181 | 0.13355 |
| Nucleoside diphosphate kinase A | P15532 | 0.509788 | 0.076537 |
| Peptidyl-prolyl cis-trans isomerase D | Q9CR16 | 0.58398 | 0.051908 |
| Metallothionein-1 | P02802 | 0.701837 | -0.15265 |
| Chloride intracellular channel protein 4 | Q9QYB1 | 0.716282 | 0.084853 |

| | | | |
|---|---|---|---|
| Transmembrane protein 109 | Q3UBX0 | 0.996126 | -0.00074 |
| Histidine triad nucleotide-binding protein 1 | P70349 | 0.209239 | 0.148799 |
| Spliceosome RNA helicase Ddx39b | Q9Z1N5 | 0.398377 | 0.13869 |
| 182 kDa tankyrase-1-binding protein | P58871 | 0.439655 | 0.106263 |
| Carnitine O-acetyltransferase | P47934 | 0.241725 | -0.21218 |
| Ubiquitin-associated protein 2-like | Q80X50 | 0.83395 | -0.03209 |
| Sorting nexin-1 | Q9WV80 | 0.327848 | -0.20104 |
| Adenosylhomocysteinase | P50247 | 0.356612 | 0.109902 |
| Nuclear migration protein nudC | O35685 | 0.613159 | 0.17847 |
| Nardilysin | Q8BHG1 | 0.665835 | -0.12951 |
| General vesicular transport factor p115 | Q9Z1Z0 | 0.718537 | 0.027596 |
| Hormone-sensitive lipase | P54310 | 0.568513 | -0.25713 |
| 26S proteasome non-ATPase regulatory subunit 3 | P14685 | 0.789483 | -0.02937 |
| Integrin beta-1 | P09055 | 0.218505 | 0.106807 |
| NAD(P)H dehydrogenase [quinone] 1 | Q64669 | 0.246058 | 0.106637 |
| Leukotriene A-4 hydrolase | P24527 | 0.310019 | 0.105758 |
| Thimet oligopeptidase | Q8C1A5 | 0.351117 | 0.153297 |
| Low-density lipoprotein receptor-related protein 8 | Q924X6 | 0.358706 | -0.19524 |
| Ubiquitin-like modifier-activating enzyme 6 | Q8C7R4 | 0.441633 | 0.140597 |
| Carnitine O-palmitoyltransferase 1, liver isoform | P97742 | 0.988405 | -0.0054 |
| P2X purinoceptor 4 | Q9JJX6 | 0.379618 | -0.16882 |
| 40S ribosomal protein S25 | P62852 | 0.745325 | 0.033422 |
| Nicotinamide phosphoribosyltransferase | Q99KQ4 | 0.51726 | -0.07265 |
| Syntaxin-4 | P70452 | 0.082849 | -0.25797 |
| Eukaryotic translation initiation factor 5A-2;Eukaryotic translatic | Q8BGY2 | 0.29795 | 0.083071 |
| Casein kinase II subunit beta | P67871 | 0.298411 | 0.132664 |
| Epididymal secretory protein E1 | Q9Z0J0 | 0.495343 | 0.099522 |
| SUN domain-containing protein 1 | Q9D666 | 0.477472 | -0.33763 |
| Vesicle-associated membrane protein-associated protein B | Q9QY76 | 0.856078 | 0.032435 |
| Serine/arginine repetitive matrix protein 2 | Q8BTI8 | 0.241092 | 0.130952 |
| NADH dehydrogenase [ubiquinone] iron-sulfur protein 6, mitocł | P52503 | 0.399221 | -0.14587 |
| Leukocyte elastase inhibitor A | Q9D154 | 0.412586 | 0.138232 |
| Transmembrane 9 superfamily member 3 | Q9ET30 | 0.190502 | 0.23814 |
| KIF1-binding protein | Q6ZPU9 | 0.516562 | -0.26136 |
| NAD(P)H-hydrate epimerase | Q8K4Z3 | 0.063725 | 0.218381 |
| E3 ubiquitin-protein ligase NEDD4 | P46935 | 0.08473 | 0.15932 |
| Aminoacyl tRNA synthase complex-interacting multifunctional p | Q8R010 | 0.240799 | 0.327408 |
| Sorting nexin-2 | Q9CWK8 | 0.07656 | 0.268423 |
| Glutathione S-transferase omega-1 | O09131 | 0.638895 | 0.046025 |
| Cullin-4B;Cullin-4A | A2A432 | 0.502567 | -0.08626 |
| Cullin-associated NEDD8-dissociated protein 2 | Q6ZQ73 | 0.911545 | 0.039514 |
| Hsc70-interacting protein | Q99L47 | 0.32955 | 0.119814 |
| Serine/threonine-protein phosphatase 2A 56 kDa regulatory sub | Q60996 | 0.4489 | -0.158 |
| Ankycorbin | Q9EP71 | 0.865061 | -0.06239 |
| Tropomodulin-3 | Q9JHJ0 | 0.988806 | 0.002299 |
| CD81 antigen | P35762 | 0.567673 | -0.1322 |
| 26S proteasome non-ATPase regulatory subunit 5 | Q8BJY1 | 0.806313 | 0.030496 |
| Protein eva-1 homolog B | Q8K2Y3 | 0.113126 | -0.37059 |
| Acyl-protein thioesterase 1 | P97823 | 0.419509 | -0.10563 |
| Protein VAC14 homolog | Q80WQ2 | 0.422482 | 0.111144 |

| | | | |
|---|---|---|---|
| Stomatin-like protein 2, mitochondrial | Q99JB2 | 0.581857 | -0.14214 |
| Protein ERGIC-53 | Q9D0F3 | 0.606032 | -0.07601 |
| H-2 class I histocompatibility antigen, L-D alpha chain;H-2 class I | P01897 | 0.68785 | 0.138826 |
| Prefoldin subunit 2 | O70591 | 0.71707 | 0.070601 |
| Ubiquitin-fold modifier 1 | P61961 | 0.257127 | -0.21648 |
| Vesicle-associated membrane protein 7 | P70280 | 0.795627 | 0.103562 |
| Non-POU domain-containing octamer-binding protein | Q99K48 | 0.218467 | 0.122222 |
| Secretory carrier-associated membrane protein 2 | Q9ERN0 | 0.828869 | -0.04693 |
| 60S ribosomal protein L21 | O09167 | 0.944866 | 0.007974 |
| Acetyl-CoA acetyltransferase, cytosolic | Q8CAY6 | 0.107299 | 0.206417 |
| Methylcrotonoyl-CoA carboxylase beta chain, mitochondrial | Q3ULD5 | 0.538798 | 0.202128 |
| Polypyrimidine tract-binding protein 1 | P17225 | 0.286071 | 0.100171 |
| Nucleoside diphosphate kinase B | Q01768 | 0.631217 | 0.041686 |
| Peroxiredoxin-4 | O08807 | 0.957688 | 0.009378 |
| Transcription intermediary factor 1-beta | Q62318 | 0.291936 | 0.186299 |
| NADH dehydrogenase [ubiquinone] flavoprotein 2, mitochondri | Q9D6J6 | 0.434594 | -0.2072 |
| 60S ribosomal protein L22-like 1 | Q9D7S7 | 0.52735 | -0.08775 |
| Estradiol 17-beta-dehydrogenase 8 | P50171 | 0.737333 | -0.04924 |
| Proteasome subunit beta type-1 | O09061 | 0.61614 | 0.048227 |
| 28S ribosomal protein S36, mitochondrial | Q9CQX8 | 0.88116 | 0.05148 |
| Guanine nucleotide-binding protein G(q) subunit alpha | P21279 | 0.001729 | 0.439692 |
| Peptidyl-prolyl cis-trans isomerase FKBP5;Peptidyl-prolyl cis-trar | Q64378 | 0.654313 | -0.09577 |
| 40S ribosomal protein S30 | P62862 | 0.825326 | 0.027362 |
| MOB kinase activator 1A;MOB kinase activator 1B | Q921Y0 | 0.061791 | 0.717032 |
| Osteoclast-stimulating factor 1 | Q62422 | 0.118671 | 0.311264 |
| Hydroxyacylglutathione hydrolase, mitochondrial | Q99KB8 | 0.665346 | -0.25518 |
| Glucosidase 2 subunit beta | O08795 | 0.279213 | 0.128843 |
| | E9Q5C9 | 0.882688 | -0.01918 |
| mRNA cap guanine-N7 methyltransferase | Q9D0L8 | 0.915586 | -0.04022 |
| 60S acidic ribosomal protein P1 | P47955 | 0.751959 | -0.16146 |
| 72 kDa type IV collagenase;PEX | P33434 | 0.8171 | -0.08856 |
| m7GpppX diphosphatase | Q9DAR7 | 0.932649 | 0.024077 |
| DnaJ homolog subfamily A member 1 | P63037 | 0.941784 | 0.028278 |
| Bleomycin hydrolase | Q8R016 | 0.81354 | 0.037362 |
| Heterogeneous nuclear ribonucleoprotein M | Q9D0E1 | 0.922598 | -0.01523 |

| Rosi-Vehicle | | Rosi-TPhP | | Normalized Log2 Intensity | | | | | | | | |
|---|---|---|---|---|---|---|---|---|---|---|---|---|
| P.VALUE.1 | LOGFC.1 | P.VALUE.2 | LOGFC.2 | TPHP_1.1 | TPHP_2.1 | TPHP_3.1 | VEHICLE_1.1 | VEHICLE_2.1 | VEHICLE_3.1 | ROSI_1.1 | ROSI_2.1 | ROSI_3.1 |
| 3.43E-08 | 2.814867 | 1.15E-07 | 2.403696 | 12.5 | 12.7 | 12.6 | 12.3 | 12.2 | 12.1 | 14.9 | 15.3 | 14.8 |
| 1.83E-06 | 2.141319 | 3.16E-05 | 1.452584 | 12.7 | 12.7 | 12.5 | 11.7 | 12.2 | 11.9 | 14.2 | 14.3 | 13.7 |
| 1.46E-06 | 2.55564 | 1.81E-05 | 1.818467 | 12.6 | 12.5 | 12.3 | 11.5 | 12.1 | 11.5 | 14.5 | 14.4 | 13.9 |
| 2.03E-06 | 2.865055 | 1.24E-05 | 2.241971 | 11.2 | 10.9 | 10.8 | 10.1 | 10.8 | 10.2 | 13.5 | 13.4 | 12.9 |
| 1.65E-06 | 2.056136 | 8.79E-06 | 1.641672 | 15.4 | 15.1 | 14.9 | 14.7 | 14.8 | 14.7 | 17 | 16.9 | 16.4 |
| 2.27E-06 | 2.407481 | 1.05E-05 | 1.957104 | 15.9 | 15.9 | 15.7 | 15.1 | 15.4 | 15.7 | 17.9 | 18.1 | 17.4 |
| 3.87E-06 | 1.542501 | 3.99E-05 | 1.119144 | 14 | 13.8 | 13.8 | 13.3 | 13.5 | 13.6 | 15.2 | 15.1 | 14.7 |
| 4.21E-06 | 1.252376 | 3.68E-05 | 0.929617 | 16 | 15.8 | 15.8 | 15.4 | 15.6 | 15.6 | 16.9 | 16.9 | 16.6 |
| 3.69E-06 | 1.733593 | 1.42E-05 | 1.442418 | 14.7 | 14.8 | 14.5 | 14.2 | 14.5 | 14.4 | 16.3 | 16.2 | 15.8 |
| 9.3E-06 | 1.342944 | 0.000584 | 0.738723 | 10 | 10.2 | 10 | 9.54 | 9.34 | 9.57 | 10.8 | 11.1 | 10.6 |
| 1.38E-05 | 0.967463 | 5.41E-05 | 0.799445 | 14.1 | 14.1 | 13.9 | 13.9 | 13.8 | 13.9 | 14.8 | 15 | 14.6 |
| 1.65E-05 | 0.716931 | 5.24E-05 | 0.610208 | 13.5 | 13.5 | 13.4 | 13.4 | 13.3 | 13.4 | 14.2 | 14.1 | 14 |
| 2.19E-05 | 1.004703 | 4.78E-05 | 0.900175 | 10.6 | 10.6 | 10.5 | 10.4 | 10.6 | 10.4 | 11.5 | 11.6 | 11.3 |
| 2.79E-05 | 2.669345 | 0.00017 | 2.060173 | 9.95 | 10.3 | 10 | 10 | 8.7 | 9.7 | 12.1 | 12.4 | 12 |
| 2.61E-05 | 1.531105 | 0.000133 | 1.213693 | 12.6 | 12.6 | 12.4 | 12.3 | 12.3 | 12 | 14 | 13.8 | 13.3 |
| 4.4E-05 | 0.808859 | 0.00306 | 0.419663 | 12.6 | 12.8 | 12.8 | 12.2 | 12.3 | 12.4 | 13.1 | 13.2 | 13 |
| 6.7E-05 | 1.037488 | 0.00055 | 0.758769 | 11.9 | 11.7 | 11.7 | 11.5 | 11.6 | 11.3 | 12.7 | 12.6 | 12.2 |
| 9.64E-05 | 0.560068 | 0.003024 | 0.325744 | 15.7 | 15.8 | 15.9 | 15.5 | 15.6 | 15.7 | 16.2 | 16.2 | 16.1 |
| 9.46E-05 | 0.70689 | 0.00125 | 0.476418 | 12.1 | 12.3 | 12.3 | 12.1 | 11.8 | 12 | 12.7 | 12.8 | 12.6 |
| 8.99E-05 | 0.525268 | 0.000495 | 0.40722 | 16.4 | 16.5 | 16.5 | 16.4 | 16.3 | 16.4 | 16.9 | 16.9 | 16.9 |
| 0.000107 | 0.64207 | 0.000733 | 0.479921 | 15.7 | 15.6 | 15.6 | 15.5 | 15.4 | 15.5 | 16.2 | 16.2 | 16 |
| 0.000115 | 0.676716 | 0.000728 | 0.511568 | 14.2 | 14 | 14.1 | 14 | 14 | 13.8 | 14.6 | 14.7 | 14.5 |
| 0.00014 | 0.66545 | 0.034282 | 0.247355 | 11.8 | 11.6 | 11.7 | 11.3 | 11.4 | 11.1 | 12 | 12 | 11.9 |
| 0.000148 | 0.428382 | 0.003988 | 0.252526 | 14.7 | 14.7 | 14.7 | 14.6 | 14.5 | 14.5 | 14.9 | 15 | 14.9 |
| 0.000135 | 0.66248 | 0.001968 | 0.43592 | 16.6 | 16.5 | 16.5 | 16.2 | 16.4 | 16.3 | 17 | 17 | 16.8 |
| 0.000144 | 0.828895 | 0.002019 | 0.547935 | 14 | 14.2 | 14.2 | 14 | 13.6 | 13.9 | 14.7 | 14.7 | 14.6 |
| 0.000163 | 0.745552 | 0.00194 | 0.505518 | 14.5 | 14.4 | 14.4 | 14.1 | 14.3 | 14.2 | 14.9 | 15.1 | 14.8 |
| 0.00016 | 0.659108 | 0.001836 | 0.44996 | 16.3 | 16.4 | 16.4 | 16.2 | 16.2 | 16.2 | 16.9 | 17 | 16.6 |
| 0.000157 | 0.62684 | 0.000324 | 0.562625 | 13.4 | 13.3 | 13.4 | 13.4 | 13.3 | 13.2 | 14 | 14.1 | 13.8 |
| 0.000159 | 0.641035 | 0.000243 | 0.602037 | 12.6 | 12.5 | 12.7 | 12.6 | 12.6 | 12.5 | 13.3 | 13.3 | 13.1 |
| 0.000188 | 0.554345 | 0.00516 | 0.322788 | 13.2 | 13.2 | 13.3 | 13 | 13.1 | 12.8 | 13.5 | 13.5 | 13.5 |
| 0.000192 | 0.543761 | 0.003781 | 0.3361 | 15.1 | 15.2 | 15.2 | 15 | 14.9 | 15 | 15.4 | 15.6 | 15.6 |
| 0.000187 | 0.592586 | 0.003095 | 0.37811 | 13.8 | 13.7 | 13.6 | 13.5 | 13.6 | 13.4 | 14.2 | 14.1 | 14 |
| 0.000228 | 1.259827 | 0.001968 | 0.895592 | 13.5 | 13.9 | 14 | 13.7 | 13.1 | 13.6 | 14.5 | 14.9 | 14.7 |
| 0.00022 | 1.657735 | 0.00179 | 1.19146 | 10.6 | 11.7 | 11.5 | 10.7 | 10.8 | 10.8 | 12.5 | 12.3 | 12.5 |
| 0.000232 | 0.585255 | 0.000437 | 0.531575 | 11.9 | 11.9 | 11.8 | 11.8 | 11.6 | 12 | 12.3 | 12.4 | 12.4 |
| 0.000261 | -0.60394 | 0.00057 | -0.53578 | 13.8 | 13.6 | 13.7 | 13.7 | 13.8 | 13.8 | 13.3 | 13 | 13.1 |
| 0.000279 | 0.827566 | 0.002249 | 0.592881 | 15.2 | 15 | 14.9 | 14.7 | 15.1 | 14.7 | 15.7 | 15.7 | 15.5 |
| 0.000288 | -0.74397 | 0.006594 | -0.44075 | 10.3 | 10.2 | 10.2 | 10.6 | 10.4 | 10.6 | 9.58 | 10 | 9.87 |
| 0.000316 | 0.731426 | 0.000652 | 0.653924 | 14.4 | 14.7 | 14.8 | 14.7 | 14.4 | 14.6 | 15.2 | 15.4 | 15.3 |
| 0.000305 | -0.5361 | 0.0006 | -0.48321 | 16.7 | 16.6 | 16.5 | 16.6 | 16.7 | 16.6 | 16.1 | 16 | 16.2 |
| 0.000316 | 0.812313 | 0.000558 | 0.744397 | 13.1 | 13.5 | 13.4 | 13.4 | 13.1 | 13.3 | 14.2 | 14.2 | 14 |

| | | | | | | | | | | | | |
|---|---|---|---|---|---|---|---|---|---|---|---|---|
| 0.000331 | 0.685419 | 0.011162 | 0.373649 | 14 | 13.7 | 13.8 | 13.6 | 13.6 | 13.3 | 14.3 | 14.2 | 14.1 |
| 0.000335 | 0.829189 | 0.001307 | 0.668991 | 12.6 | 12.9 | 12.8 | 12.8 | 12.6 | 12.4 | 13.6 | 13.5 | 13.2 |
| 0.000356 | 0.503927 | 0.009109 | 0.289468 | 13 | 13.1 | 13.1 | 12.7 | 12.8 | 13 | 13.3 | 13.4 | 13.3 |
| 0.000362 | -0.60911 | 0.008372 | -0.35667 | 12.3 | 12.2 | 12.2 | 12.4 | 12.6 | 12.5 | 11.8 | 12 | 11.9 |
| 0.000378 | 0.737306 | 0.029265 | 0.330533 | 12.6 | 12.6 | 12.3 | 12.2 | 11.9 | 12 | 12.9 | 12.9 | 12.7 |
| 0.000386 | 0.363155 | 0.007692 | 0.21832 | 16.4 | 16.5 | 16.5 | 16.3 | 16.3 | 16.3 | 16.7 | 16.7 | 16.7 |
| 0.000393 | 0.408103 | 0.005149 | 0.265611 | 16.7 | 16.6 | 16.6 | 16.5 | 16.5 | 16.5 | 16.9 | 17 | 16.9 |
| 0.000615 | 0.96383 | 0.006702 | 0.639616 | 11.4 | 11.6 | 11.5 | 11.5 | 10.8 | 11.3 | 12.1 | 12.2 | 12.1 |
| 0.000667 | -0.5749 | 0.00135 | -0.51313 | 12.6 | 12.5 | 12.6 | 12.5 | 12.5 | 12.8 | 12.2 | 12 | 12 |
| 0.000705 | -0.44257 | 0.003193 | -0.34402 | 14.6 | 14.3 | 14.5 | 14.6 | 14.6 | 14.5 | 14.1 | 14.1 | 14.1 |
| 0.00076 | 0.655763 | 0.008068 | 0.434082 | 9.87 | 9.93 | 10.1 | 9.53 | 9.79 | 9.95 | 10.3 | 10.4 | 10.5 |
| 0.00078 | 0.564129 | 0.000617 | 0.585307 | 15.3 | 15 | 15 | 15 | 15.2 | 15.2 | 15.7 | 15.8 | 15.6 |
| 0.000861 | 0.749383 | 0.062213 | 0.312016 | 11.4 | 11 | 11.4 | 10.8 | 10.9 | 10.8 | 11.8 | 11.4 | 11.5 |
| 0.000819 | -0.52014 | 0.015147 | -0.30603 | 11 | 10.8 | 10.9 | 11.1 | 11.1 | 11.2 | 10.7 | 10.7 | 10.5 |
| 0.000831 | -0.62845 | 0.005266 | -0.45786 | 14.7 | 14.3 | 14.3 | 14.5 | 14.6 | 14.7 | 14.1 | 13.8 | 14 |
| 0.000866 | 0.579787 | 0.005083 | 0.428021 | 16.2 | 16 | 15.9 | 15.9 | 16 | 15.9 | 16.6 | 16.5 | 16.3 |
| 0.000838 | -0.42862 | 0.003977 | -0.32927 | 16.5 | 16.3 | 16.3 | 16.4 | 16.5 | 16.5 | 16 | 16 | 16.1 |
| 0.000953 | -0.44532 | 0.02807 | -0.23316 | 10.4 | 10.2 | 10.2 | 10.5 | 10.5 | 10.5 | 9.95 | 10.1 | 10.1 |
| 0.001001 | 0.574529 | 0.028691 | 0.301597 | 11.5 | 11.3 | 11.6 | 11.1 | 11.1 | 11.3 | 11.7 | 11.8 | 11.7 |
| 0.000965 | 0.764996 | 0.024653 | 0.414063 | 11.4 | 10.9 | 11.1 | 10.9 | 10.8 | 10.6 | 11.4 | 11.7 | 11.5 |
| 0.000963 | 0.404529 | 0.020734 | 0.227845 | 15.1 | 15.2 | 15.1 | 14.9 | 14.8 | 15.1 | 15.3 | 15.4 | 15.3 |
| 0.001017 | 0.49873 | 0.020074 | 0.285459 | 14.5 | 14.3 | 14.3 | 14 | 14.3 | 14.1 | 14.7 | 14.7 | 14.6 |
| 0.001014 | 0.33111 | 0.015072 | 0.201834 | 14.9 | 14.9 | 14.9 | 14.8 | 14.9 | 14.8 | 15.2 | 15.2 | 15.1 |
| 0.000999 | 0.425479 | 0.008034 | 0.294497 | 16.4 | 16.2 | 16.1 | 16.1 | 16.2 | 16.1 | 16.6 | 16.6 | 16.5 |
| 0.000975 | 0.532629 | 0.003737 | 0.423932 | 14 | 13.8 | 14 | 13.9 | 14 | 13.7 | 14.4 | 14.5 | 14.3 |
| 0.00109 | 0.644114 | 0.026391 | 0.349826 | 11.9 | 12 | 12 | 11.8 | 11.7 | 11.6 | 12.6 | 12.3 | 12.1 |
| 0.001115 | 0.664336 | 0.002666 | 0.57373 | 11.6 | 12.1 | 12 | 11.8 | 11.7 | 12 | 12.5 | 12.5 | 12.5 |
| 0.001228 | -0.93234 | 0.031862 | -0.49306 | 13.7 | 13.5 | 13.5 | 13.8 | 13.9 | 14.3 | 13.3 | 13.1 | 12.8 |
| 0.00119 | 0.502322 | 0.021382 | 0.290751 | 11.6 | 11.8 | 11.6 | 11.6 | 11.3 | 11.5 | 11.8 | 12.1 | 11.9 |
| 0.001236 | -0.37775 | 0.00576 | -0.28858 | 15.3 | 15.2 | 15.2 | 15.3 | 15.3 | 15.4 | 15 | 15 | 14.9 |
| 0.00121 | 0.522117 | 0.004872 | 0.410088 | 13.1 | 13 | 13.1 | 12.8 | 12.9 | 13.2 | 13.6 | 13.4 | 13.5 |
| 0.001196 | -0.74462 | 0.000764 | -0.80031 | 11.1 | 10.9 | 10.8 | 10.7 | 11 | 10.9 | 9.93 | 10 | 10.4 |
| 0.00128 | -0.51225 | 0.001099 | -0.52519 | 13.6 | 13.5 | 13.2 | 13.4 | 13.3 | 13.5 | 13 | 12.9 | 12.8 |
| 0.001356 | -0.57064 | 0.010272 | -0.39526 | 11.9 | 11.8 | 11.8 | 12 | 12.2 | 11.9 | 11.4 | 11.6 | 11.3 |
| 0.001362 | -0.37455 | 0.000472 | -0.44346 | 13.8 | 13.9 | 13.8 | 13.8 | 13.8 | 13.7 | 13.3 | 13.4 | 13.4 |
| 0.001329 | -0.54096 | 0.000761 | -0.592 | 10.8 | 10.6 | 10.4 | 10.5 | 10.5 | 10.6 | 9.98 | 9.85 | 10.1 |
| 0.001581 | 0.403502 | 0.030349 | 0.225204 | 16.6 | 16.8 | 16.8 | 16.6 | 16.5 | 16.5 | 16.9 | 17 | 16.9 |
| 0.001649 | 0.862349 | 0.006322 | 0.6789 | 11.3 | 10.9 | 10.8 | 11 | 10.8 | 10.6 | 11.9 | 11.5 | 11.6 |
| 0.001688 | 0.964662 | 0.189483 | 0.296421 | 12.8 | 12.5 | 12.1 | 12.2 | 11.8 | 11.5 | 12.9 | 12.9 | 12.6 |
| 0.001696 | -0.44819 | 0.00822 | -0.33691 | 13.2 | 13.5 | 13.4 | 13.4 | 13.5 | 13.5 | 13 | 13 | 13.1 |
| 0.001816 | -0.33617 | 0.033624 | -0.1872 | 15.5 | 15.4 | 15.4 | 15.6 | 15.6 | 15.6 | 15.2 | 15.2 | 15.3 |
| 0.002035 | -0.70114 | 0.068667 | -0.32648 | 16.1 | 15.5 | 15.6 | 16.1 | 16 | 16.2 | 15.4 | 15.5 | 15.3 |
| 0.002046 | 0.455943 | 0.00958 | 0.343172 | 12.3 | 12.2 | 12.1 | 12.1 | 12 | 12.3 | 12.5 | 12.7 | 12.5 |
| 0.002207 | 0.556893 | 0.022997 | 0.351479 | 15.5 | 15.8 | 15.9 | 15.6 | 15.3 | 15.6 | 16.1 | 16.1 | 16 |
| 0.002127 | 0.536839 | 0.018968 | 0.351834 | 13.2 | 13.2 | 13.1 | 13 | 13.2 | 12.7 | 13.5 | 13.5 | 13.4 |
| 0.002191 | -0.44589 | 0.013198 | -0.31785 | 11.1 | 11.3 | 11.3 | 11.4 | 11.3 | 11.5 | 10.8 | 10.9 | 11 |
| 0.002206 | 0.53523 | 0.000925 | 0.618147 | 13 | 12.8 | 12.9 | 12.8 | 13 | 13 | 13.7 | 13.5 | 13.3 |
| 0.00218 | -0.38642 | 0.004549 | -0.33919 | 15.5 | 15.3 | 15.4 | 15.4 | 15.5 | 15.4 | 15 | 15 | 15.1 |
| 0.00216 | 0.500583 | 0.00306 | 0.471194 | 11.4 | 11.2 | 11.5 | 11.4 | 11.2 | 11.5 | 11.8 | 11.9 | 11.8 |
| 0.002421 | -0.38771 | 0.071449 | -0.18373 | 13.3 | 13.5 | 13.4 | 13.5 | 13.5 | 13.7 | 13.2 | 13.2 | 13.1 |

| | | | | | | | | | | | | |
|---|---|---|---|---|---|---|---|---|---|---|---|---|
| 0.002348 | 0.436095 | 0.04573 | 0.234043 | 13.7 | 13.9 | 13.7 | 13.5 | 13.6 | 13.6 | 14.1 | 14.1 | 13.8 |
| 0.002322 | -0.36924 | 0.04204 | -0.20234 | 12.4 | 12.4 | 12.5 | 12.7 | 12.7 | 12.5 | 12.3 | 12.1 | 12.3 |
| 0.0024 | 0.395995 | 0.009206 | 0.308766 | 13.6 | 13.9 | 13.7 | 13.6 | 13.6 | 13.7 | 14.1 | 14.1 | 13.9 |
| 0.00239 | -0.34667 | 0.000732 | -0.42138 | 15.5 | 15.5 | 15.5 | 15.5 | 15.5 | 15.3 | 15.1 | 15 | 15.2 |
| 0.002595 | 0.275948 | 0.035289 | 0.161355 | 16.9 | 16.9 | 16.9 | 16.7 | 16.8 | 16.8 | 17.1 | 17.1 | 17 |
| 0.002599 | -0.34104 | 0.016302 | -0.23926 | 16.1 | 16.1 | 16.1 | 16.2 | 16.1 | 16.3 | 15.8 | 15.9 | 15.9 |
| 0.002578 | -0.3098 | 0.001981 | -0.32414 | 18.1 | 18.1 | 18.1 | 18.2 | 18 | 18.1 | 17.8 | 17.7 | 17.8 |
| 0.002547 | 0.521102 | 0.00275 | 0.514189 | 11.6 | 11.9 | 11.9 | 11.9 | 11.6 | 11.9 | 12.2 | 12.4 | 12.3 |
| 0.002724 | 0.438489 | 0.00066 | 0.554202 | 11.2 | 11.1 | 11.1 | 11.2 | 11.3 | 11.3 | 11.7 | 11.9 | 11.5 |
| 0.002699 | 0.309361 | 0.002301 | 0.318045 | 12.9 | 12.8 | 13 | 12.9 | 12.9 | 13 | 13.2 | 13.2 | 13.2 |
| 0.002871 | -0.53878 | 0.041598 | -0.30719 | 12.5 | 12 | 12.1 | 12.4 | 12.5 | 12.4 | 12 | 11.9 | 11.8 |
| 0.002892 | 0.285075 | 0.006739 | 0.244113 | 17.5 | 17.5 | 17.5 | 17.4 | 17.5 | 17.4 | 17.8 | 17.8 | 17.6 |
| 0.003381 | -0.32216 | 0.106873 | -0.14148 | 11.6 | 11.5 | 11.4 | 11.7 | 11.7 | 11.7 | 11.4 | 11.3 | 11.4 |
| 0.003246 | 0.348432 | 0.098006 | 0.156644 | 13.5 | 13.7 | 13.6 | 13.5 | 13.4 | 13.4 | 13.8 | 13.7 | 13.8 |
| 0.00324 | -0.40265 | 0.061435 | -0.21022 | 13.9 | 14.2 | 13.9 | 14.3 | 14.1 | 14.2 | 13.8 | 13.7 | 13.9 |
| 0.00331 | 0.515258 | 0.016348 | 0.376977 | 15.9 | 15.6 | 15.6 | 15.4 | 15.7 | 15.5 | 16.1 | 16.1 | 16 |
| 0.003371 | 0.479879 | 0.015322 | 0.357242 | 12.1 | 12.2 | 12.1 | 12.2 | 12.1 | 11.8 | 12.5 | 12.6 | 12.4 |
| 0.003271 | -0.35544 | 0.008023 | -0.30068 | 15.3 | 15.2 | 15.2 | 15.2 | 15.2 | 15.4 | 15 | 14.9 | 14.8 |
| 0.003341 | -0.59582 | 0.006407 | -0.52848 | 12.2 | 11.7 | 11.7 | 12 | 12 | 11.8 | 11.4 | 11.2 | 11.4 |
| 0.003397 | -0.44478 | 0.003891 | -0.43407 | 15.9 | 16 | 16 | 16.1 | 16 | 15.9 | 15.4 | 15.4 | 15.7 |
| 0.003541 | -1.08986 | 0.046608 | -0.62543 | 9.19 | 8.35 | 8.43 | 9 | 9.16 | 9.2 | 8.13 | 8.38 | 7.58 |
| 0.003632 | 0.857216 | 0.037911 | 0.522168 | 10.1 | 10.3 | 10.4 | 9.69 | 9.81 | 10.3 | 10.6 | 10.6 | 11.2 |
| 0.003805 | -0.26302 | 0.000218 | -0.41802 | 14.9 | 14.9 | 14.9 | 14.8 | 14.7 | 14.8 | 14.5 | 14.5 | 14.5 |
| 0.003785 | -0.32729 | 0.061299 | -0.17581 | 12.5 | 12.3 | 12.4 | 12.6 | 12.6 | 12.5 | 12.3 | 12.2 | 12.2 |
| 0.003883 | -0.36719 | 0.008343 | -0.31785 | 16 | 15.8 | 15.8 | 15.8 | 16 | 15.9 | 15.6 | 15.5 | 15.6 |
| 0.004119 | 0.38661 | 0.030418 | 0.25473 | 17 | 16.9 | 16.8 | 16.8 | 16.9 | 16.7 | 17.1 | 17.3 | 17.2 |
| 0.004184 | 0.977921 | 0.006764 | 0.893914 | 10.7 | 9.98 | 10.5 | 9.82 | 10.4 | 10.6 | 11.4 | 11.3 | 11.1 |
| 0.004542 | -0.45602 | 0.017277 | -0.34899 | 13.7 | 13.6 | 13.5 | 13.6 | 13.6 | 13.9 | 13.4 | 13.1 | 13.3 |
| 0.004614 | 0.39486 | 0.014699 | 0.313917 | 16 | 15.8 | 15.8 | 15.7 | 15.8 | 15.7 | 16.2 | 16.2 | 16 |
| 0.004582 | 0.386294 | 0.004312 | 0.390622 | 11 | 10.8 | 11 | 11 | 10.9 | 10.9 | 11.5 | 11.3 | 11.2 |
| 0.004774 | 0.722832 | 0.232898 | 0.239878 | 13.3 | 13.8 | 13.6 | 13.4 | 12.8 | 13.1 | 13.6 | 13.9 | 13.9 |
| 0.004893 | 0.399732 | 0.039962 | 0.253557 | 13.6 | 13.8 | 13.7 | 13.6 | 13.4 | 13.6 | 14.1 | 14 | 13.8 |
| 0.005006 | 1.078889 | 0.026582 | 0.761341 | 12.5 | 13.4 | 13.6 | 13 | 12.9 | 12.8 | 13.8 | 13.8 | 14.3 |
| 0.004998 | -0.59791 | 0.015417 | -0.47749 | 12.1 | 11.9 | 11.8 | 12 | 12.2 | 12 | 11.4 | 11.8 | 11.2 |
| 0.005047 | 0.79549 | 0.008846 | 0.714172 | 10.4 | 10.7 | 10.5 | 10.6 | 10.7 | 9.98 | 11.3 | 11.4 | 11 |
| 0.005154 | 0.534946 | 2.72E-05 | 1.212131 | 8.07 | 8.02 | 8.01 | 8.99 | 8.59 | 8.55 | 9.41 | 9.06 | 9.26 |
| 0.005209 | -0.27796 | 0.087126 | -0.14176 | 14.4 | 14.5 | 14.5 | 14.6 | 14.6 | 14.6 | 14.2 | 14.4 | 14.3 |
| 0.005196 | 0.321981 | 0.015147 | 0.25999 | 11.6 | 11.6 | 11.5 | 11.5 | 11.6 | 11.4 | 11.9 | 11.9 | 11.8 |
| 0.005638 | -0.35758 | 0.000939 | -0.48745 | 14.1 | 13.9 | 14 | 13.9 | 14 | 13.7 | 13.5 | 13.5 | 13.6 |
| 0.00566 | 0.560099 | 0.04029 | 0.3643 | 14 | 13.7 | 13.8 | 13.4 | 13.8 | 13.8 | 14.3 | 14.3 | 14 |
| 0.005622 | 0.511804 | 0.021499 | 0.38794 | 14.1 | 13.6 | 13.7 | 13.7 | 13.8 | 13.6 | 14.3 | 14.3 | 14.1 |
| 0.005741 | -0.50332 | 0.155088 | -0.2104 | 12.8 | 12.7 | 12.6 | 12.9 | 13.1 | 13 | 12.7 | 12.6 | 12.2 |
| 0.005772 | 0.282587 | 0.031233 | 0.196899 | 14.5 | 14.4 | 14.4 | 14.3 | 14.4 | 14.4 | 14.7 | 14.7 | 14.6 |
| 0.005872 | -0.44439 | 0.007452 | -0.42453 | 17.1 | 16.8 | 16.8 | 16.9 | 17.1 | 16.8 | 16.5 | 16.4 | 16.5 |
| 0.006025 | 0.239694 | 0.015029 | 0.199348 | 13.8 | 13.9 | 13.9 | 13.8 | 13.8 | 13.8 | 14 | 14.1 | 14 |
| 0.00614 | -0.87129 | 0.030462 | -0.61805 | 11.5 | 11.8 | 11 | 11.7 | 11.8 | 11.5 | 10.8 | 10.5 | 11.2 |
| 0.006292 | 0.308466 | 0.021465 | 0.238942 | 14.2 | 14.4 | 14.4 | 14.2 | 14.3 | 14.3 | 14.6 | 14.6 | 14.4 |
| 0.006257 | 0.610666 | 0.011651 | 0.539798 | 14.5 | 14 | 14 | 14 | 14.2 | 14.1 | 14.8 | 14.8 | 14.5 |
| 0.006169 | -0.26888 | 0.003533 | -0.29799 | 13.6 | 13.7 | 13.7 | 13.7 | 13.6 | 13.7 | 13.3 | 13.4 | 13.5 |
| 0.006226 | -0.26038 | 0.005012 | -0.27122 | 15.2 | 15.1 | 15.1 | 15.1 | 15.1 | 15.2 | 14.9 | 14.8 | 14.9 |

| | | | | | | | | | | | | |
|---|---|---|---|---|---|---|---|---|---|---|---|---|
| 0.006487 | 0.269483 | 0.051991 | 0.167782 | 16.1 | 15.9 | 15.9 | 15.8 | 15.9 | 15.8 | 16.1 | 16.1 | 16.1 |
| 0.006525 | -0.43139 | 0.011528 | -0.38525 | 13.6 | 13.5 | 13.5 | 13.5 | 13.6 | 13.7 | 13.4 | 13 | 13 |
| 0.006669 | -0.39587 | 0.006031 | -0.40355 | 14.3 | 14.6 | 14.6 | 14.5 | 14.6 | 14.5 | 14.2 | 14 | 14.1 |
| 0.006728 | -0.33033 | 0.303643 | -0.09957 | 13.2 | 13.1 | 13.1 | 13.4 | 13.5 | 13.3 | 13.1 | 12.9 | 13.1 |
| 0.006857 | -0.29855 | 0.026988 | -0.22278 | 15 | 14.7 | 14.8 | 14.8 | 14.9 | 14.9 | 14.6 | 14.5 | 14.6 |
| 0.007023 | -0.27076 | 0.017139 | -0.22534 | 17.9 | 18.1 | 18 | 18.1 | 18 | 18 | 17.7 | 17.7 | 17.9 |
| 0.007259 | 0.43934 | 0.055774 | 0.273987 | 13 | 12.7 | 12.8 | 12.5 | 12.8 | 12.6 | 13.1 | 13.2 | 13 |
| 0.007396 | 0.310411 | 0.576973 | 0.050375 | 12.6 | 12.4 | 12.4 | 12.3 | 12.1 | 12.3 | 12.5 | 12.5 | 12.5 |
| 0.007348 | -0.27993 | 0.159427 | -0.12122 | 13.2 | 13.2 | 13.2 | 13.4 | 13.4 | 13.3 | 13 | 13.1 | 13.2 |
| 0.007494 | -0.2553 | 0.064774 | -0.15329 | 14.4 | 14.4 | 14.4 | 14.5 | 14.6 | 14.5 | 14.3 | 14.2 | 14.3 |
| 0.007668 | 0.600797 | 0.119453 | 0.294922 | 11.6 | 11.3 | 11.9 | 11.5 | 11.2 | 11.1 | 12 | 11.7 | 12 |
| 0.007628 | -0.50188 | 0.022698 | -0.3983 | 14.7 | 14.9 | 15.1 | 14.9 | 14.8 | 15.2 | 14.6 | 14.4 | 14.5 |
| 0.007816 | -0.2345 | 0.049968 | -0.15307 | 15.1 | 15.1 | 15 | 15.2 | 15.1 | 15.2 | 14.9 | 14.9 | 14.8 |
| 0.007966 | -0.30954 | 7.94E-05 | -0.65508 | 10.6 | 10.6 | 10.8 | 10.3 | 10.3 | 10.2 | 9.89 | 10.1 | 10 |
| 0.007889 | -0.27062 | 0.143883 | -0.12411 | 13 | 12.8 | 12.9 | 13 | 13.1 | 13 | 12.7 | 12.8 | 12.8 |
| 0.008032 | 0.262764 | 0.088109 | 0.145047 | 15 | 14.9 | 14.9 | 14.8 | 14.9 | 14.7 | 15.1 | 15.1 | 15 |
| 0.008015 | -0.28235 | 0.045245 | -0.19034 | 15.2 | 15 | 15 | 15.1 | 15.2 | 15.2 | 14.9 | 14.8 | 14.9 |
| 0.008501 | 0.451621 | 0.091816 | 0.248624 | 14.4 | 14.1 | 14.1 | 14 | 14.3 | 13.9 | 14.5 | 14.6 | 14.4 |
| 0.00849 | -0.46277 | 0.019382 | -0.38888 | 10.8 | 10.8 | 10.9 | 11 | 10.6 | 11.1 | 10.4 | 10.5 | 10.4 |
| 0.008744 | 0.398009 | 0.682884 | 0.04865 | 12.1 | 12.2 | 12.2 | 11.7 | 11.7 | 12.1 | 12.2 | 12.2 | 12.2 |
| 0.008747 | -0.275 | 0.336046 | -0.08123 | 13.1 | 13 | 13.2 | 13.3 | 13.3 | 13.3 | 12.9 | 13.1 | 13.1 |
| 0.008643 | -0.38515 | 0.009405 | -0.37871 | 12.8 | 12.7 | 12.5 | 12.8 | 12.7 | 12.6 | 12.2 | 12.3 | 12.4 |
| 0.008953 | -0.29907 | 0.058986 | -0.19123 | 13.7 | 13.8 | 13.9 | 14 | 13.8 | 13.9 | 13.6 | 13.6 | 13.6 |
| 0.008965 | 1.027254 | 0.046226 | 0.703918 | 13.2 | 14.1 | 14.1 | 13.9 | 12.9 | 13.6 | 14.4 | 14.6 | 14.5 |
| 0.008951 | -0.48531 | 0.030675 | -0.36982 | 12.6 | 12.3 | 12.2 | 12.4 | 12.5 | 12.6 | 12 | 11.8 | 12.2 |
| 0.008974 | -0.24155 | 0.029448 | -0.18603 | 15.1 | 15.1 | 15.1 | 15.1 | 15.2 | 15.1 | 14.9 | 14.9 | 14.9 |
| 0.009175 | 0.314536 | 0.030134 | 0.241937 | 14 | 14.3 | 14.3 | 14.2 | 14 | 14.2 | 14.4 | 14.5 | 14.5 |
| 0.009318 | -0.2924 | 0.065982 | -0.18227 | 14.5 | 14.4 | 14.5 | 14.5 | 14.7 | 14.5 | 14.3 | 14.3 | 14.2 |
| 0.009532 | -0.33707 | 0.028383 | -0.26512 | 16.6 | 16.3 | 16.3 | 16.4 | 16.6 | 16.4 | 16.2 | 16.1 | 16.1 |
| 0.009756 | 5.986716 | 0.836022 | -0.3774 | 8.35 | 9.56 | 9.32 | 0.13 | 7.43 | 0.58 | 7.55 | 9.01 | 9.54 |
| 0.0097 | 0.396121 | 0.027905 | 0.313957 | 14.1 | 14.2 | 14.3 | 14.3 | 13.9 | 14.1 | 14.4 | 14.5 | 14.6 |
| 0.00983 | 0.555053 | 0.010482 | 0.547864 | 12 | 12.5 | 12.4 | 12.3 | 12 | 12.5 | 12.8 | 12.8 | 12.9 |
| 0.009898 | -0.47096 | 0.166293 | -0.21238 | 12.2 | 11.6 | 12 | 12.2 | 12.1 | 12.3 | 11.8 | 11.7 | 11.6 |
| 0.010223 | -0.50498 | 0.025344 | -0.41397 | 14.6 | 14.2 | 14.2 | 14.3 | 14.7 | 14.3 | 14.1 | 13.9 | 13.8 |
| 0.010213 | -5.72806 | 0.020696 | -4.92142 | 7.79 | 6.38 | 7.1 | 7.23 | 8.1 | 8.35 | -0.2 | -0.1 | 6.81 |
| 0.010479 | 0.484069 | 0.01207 | 0.470175 | 9.91 | 10.1 | 10.4 | 9.94 | 10.2 | 10.2 | 10.7 | 10.5 | 10.6 |
| 0.01057 | -0.30767 | 0.103826 | -0.16954 | 12.1 | 11.9 | 11.9 | 12 | 12 | 12.2 | 11.7 | 11.9 | 11.8 |
| 0.010876 | -0.21769 | 0.2885 | -0.07466 | 14.3 | 14.3 | 14.2 | 14.5 | 14.5 | 14.4 | 14.2 | 14.3 | 14.2 |
| 0.011064 | 0.385756 | 0.029177 | 0.310648 | 14.7 | 15 | 15.2 | 14.9 | 14.8 | 15 | 15.2 | 15.2 | 15.4 |
| 0.011152 | -0.40911 | 0.433736 | -0.10219 | 13 | 12.7 | 12.7 | 13 | 13.1 | 13.2 | 12.8 | 12.6 | 12.6 |
| 0.011269 | -0.50388 | 0.005004 | -0.5904 | 9.23 | 9.01 | 9.18 | 8.83 | 9.13 | 9.2 | 8.82 | 8.53 | 8.3 |
| 0.011294 | -0.30828 | 0.011742 | -0.30581 | 16.3 | 16 | 16.1 | 16.1 | 16.2 | 16.1 | 15.8 | 15.7 | 15.9 |
| 0.011413 | -0.23357 | 0.001661 | -0.33314 | 15.3 | 15.2 | 15.2 | 15.1 | 15.2 | 15.2 | 14.9 | 15 | 14.9 |
| 0.011587 | 0.674465 | 0.054126 | 0.465899 | 11.9 | 11.7 | 11.7 | 11.2 | 12 | 11.4 | 12.2 | 12.4 | 12 |
| 0.011677 | -0.36476 | 0.006114 | -0.41465 | 11.6 | 11.5 | 11.5 | 11.4 | 11.7 | 11.4 | 11.2 | 10.9 | 11.2 |
| 0.011843 | 0.290525 | 0.161147 | 0.137817 | 12.1 | 12.1 | 12.1 | 12 | 11.8 | 12.1 | 12.3 | 12.4 | 12.2 |
| 0.011928 | -0.22359 | 0.059962 | -0.15083 | 14.6 | 14.5 | 14.5 | 14.6 | 14.6 | 14.6 | 14.4 | 14.5 | 14.4 |
| 0.012117 | -0.22138 | 0.026979 | -0.18515 | 14.8 | 14.9 | 14.9 | 14.9 | 14.9 | 15 | 14.7 | 14.7 | 14.7 |
| 0.012078 | -0.20084 | 0.023349 | -0.17371 | 15.8 | 15.8 | 15.8 | 15.8 | 15.8 | 15.8 | 15.6 | 15.6 | 15.7 |
| 0.012118 | -0.28824 | 0.015849 | -0.27223 | 13.7 | 13.6 | 13.6 | 13.7 | 13.5 | 13.8 | 13.5 | 13.3 | 13.4 |

| | | | | | | | | | | | | |
|---|---|---|---|---|---|---|---|---|---|---|---|---|
| 0.01232 | 0.850132 | 0.456192 | 0.205944 | 10.6 | 11.2 | 11.3 | 10.5 | 10.7 | 9.97 | 11 | 11.4 | 11.3 |
| 0.012363 | -0.48931 | 0.042201 | -0.36695 | 15.3 | 15.7 | 15.7 | 15.8 | 15.4 | 15.8 | 15.1 | 15.1 | 15.3 |
| 0.012685 | 0.256863 | 0.007201 | 0.287899 | 13.3 | 13.3 | 13.4 | 13.4 | 13.4 | 13.4 | 13.5 | 13.7 | 13.6 |
| 0.013187 | 0.197156 | 0.228677 | 0.080713 | 17.2 | 17.3 | 17.2 | 17.1 | 17.1 | 17.1 | 17.3 | 17.3 | 17.3 |
| 0.013167 | -0.31718 | 0.004759 | -0.3875 | 13.5 | 13.4 | 13.4 | 13.2 | 13.5 | 13.3 | 13.1 | 12.9 | 13.1 |
| 0.013323 | 0.380503 | 0.333286 | 0.123269 | 12.3 | 12.2 | 12.2 | 12.2 | 11.7 | 12.1 | 12.5 | 12.3 | 12.4 |
| 0.013423 | -0.34508 | 0.170325 | -0.16397 | 10.6 | 10.6 | 10.8 | 10.6 | 10.9 | 11 | 10.5 | 10.6 | 10.4 |
| 0.013804 | -0.21919 | 0.001316 | -0.33819 | 13 | 12.9 | 13 | 12.8 | 12.8 | 12.9 | 12.7 | 12.6 | 12.5 |
| 0.01391 | -0.25931 | 0.035738 | -0.20822 | 14.7 | 14.6 | 14.5 | 14.5 | 14.7 | 14.7 | 14.4 | 14.3 | 14.5 |
| 0.014102 | -0.28586 | 0.239748 | -0.11566 | 13.9 | 14 | 14 | 14.2 | 14 | 14.2 | 13.9 | 13.8 | 13.9 |
| 0.014274 | -0.24573 | 0.009158 | -0.26934 | 15.8 | 15.7 | 15.6 | 15.7 | 15.8 | 15.6 | 15.5 | 15.4 | 15.5 |
| 0.014826 | 0.273209 | 0.124955 | 0.150875 | 13.7 | 13.5 | 13.7 | 13.6 | 13.5 | 13.5 | 13.9 | 13.8 | 13.7 |
| 0.014887 | -0.51301 | 0.082654 | -0.32833 | 12.5 | 12.3 | 12 | 12.3 | 12.7 | 12.3 | 12.1 | 11.9 | 11.7 |
| 0.014692 | -0.41638 | 0.004821 | -0.51947 | 11.6 | 11.2 | 11.4 | 11.1 | 11.2 | 11.5 | 10.9 | 10.8 | 10.9 |
| 0.014718 | -0.39527 | 0.037315 | -0.31769 | 12.9 | 12.8 | 12.7 | 13 | 12.8 | 12.8 | 12.2 | 12.6 | 12.6 |
| 0.014712 | -0.94959 | 0.02922 | -0.81161 | 9.09 | 8.62 | 8.66 | 9.01 | 8.71 | 9.05 | 8.06 | 8.58 | 7.29 |
| 0.014442 | -0.34205 | 0.023922 | -0.30548 | 12.2 | 12 | 12.2 | 12 | 12.3 | 12.2 | 12 | 11.8 | 11.7 |
| 0.014534 | 1.1901 | 0.021962 | 1.085731 | 9.89 | 11.3 | 11.4 | 10.6 | 10.6 | 11.1 | 12 | 11.9 | 11.9 |
| 0.014873 | -0.38215 | 0.014307 | -0.38535 | 11.8 | 11.4 | 11.8 | 11.7 | 11.6 | 11.8 | 11.4 | 11.3 | 11.2 |
| 0.015261 | -0.29856 | 0.059607 | -0.21258 | 14.4 | 14.3 | 14.2 | 14.3 | 14.5 | 14.3 | 14 | 14.1 | 14.2 |
| 0.015289 | -0.20812 | 0.012116 | -0.21865 | 14.7 | 14.6 | 14.6 | 14.6 | 14.7 | 14.6 | 14.4 | 14.4 | 14.4 |
| 0.015388 | -0.25599 | 0.09785 | -0.15587 | 12.9 | 12.7 | 12.9 | 12.9 | 12.9 | 13 | 12.7 | 12.6 | 12.7 |
| 0.015644 | 6.334971 | 0.973534 | 0.070557 | 9.22 | 9.54 | 9.17 | 0.11 | 0.25 | 8.78 | 8.99 | 9.87 | 9.29 |
| 0.015922 | -0.41748 | 0.096914 | -0.25695 | 13.9 | 13.7 | 13.4 | 13.7 | 13.8 | 13.9 | 13.4 | 13.4 | 13.4 |
| 0.015912 | -0.32044 | 0.083155 | -0.20765 | 12.6 | 12.3 | 12.3 | 12.4 | 12.6 | 12.6 | 12.2 | 12.3 | 12.1 |
| 0.016066 | -0.34039 | 0.038332 | -0.27689 | 12.6 | 12.5 | 12.7 | 12.5 | 12.6 | 12.9 | 12.4 | 12.4 | 12.2 |
| 0.016176 | -0.29473 | 0.004477 | -0.38059 | 13.1 | 12.7 | 12.8 | 12.8 | 12.7 | 12.8 | 12.5 | 12.4 | 12.5 |
| 0.01628 | 5.772949 | 0.958836 | 0.100909 | 8.96 | 9.01 | 7.23 | 0.05 | 0.25 | 7.89 | 8.38 | 8.51 | 8.62 |
| 0.016759 | 0.302124 | 0.763816 | 0.031031 | 11.4 | 11.7 | 11.6 | 11.4 | 11.2 | 11.3 | 11.5 | 11.6 | 11.6 |
| 0.01674 | 0.279246 | 0.121711 | 0.159941 | 12.6 | 12.6 | 12.8 | 12.6 | 12.5 | 12.6 | 12.8 | 13 | 12.9 |
| 0.016721 | -0.35925 | 0.040663 | -0.29026 | 15.6 | 15.3 | 15.2 | 15.4 | 15.6 | 15.3 | 15.1 | 15 | 15.1 |
| 0.016912 | -0.2294 | 0.018252 | -0.22556 | 15.1 | 15.2 | 15.1 | 15.2 | 15 | 15.2 | 14.9 | 14.9 | 14.9 |
| 0.017089 | -0.80533 | 0.250836 | -0.33109 | 9.11 | 9.18 | 9.25 | 9.54 | 10.3 | 9.12 | 8.88 | 8.64 | 9.02 |
| 0.017171 | -0.62257 | 0.006095 | -0.76896 | 9.87 | 10.2 | 9.9 | 10 | 10.1 | 9.41 | 9.08 | 9.49 | 9.1 |
| 0.017439 | 0.344939 | 0.154866 | 0.180858 | 16.4 | 16.1 | 16.1 | 15.9 | 16.2 | 16 | 16.5 | 16.4 | 16.3 |
| 0.017458 | -0.54919 | 0.08638 | -0.35867 | 10.9 | 10.8 | 10.6 | 10.8 | 11 | 11 | 10.8 | 10.1 | 10.2 |
| 0.017754 | 0.494865 | 0.130518 | 0.279467 | 10.5 | 11.2 | 11 | 10.6 | 10.7 | 10.8 | 11.1 | 11.3 | 11.1 |
| 0.017757 | -5.90528 | 0.013067 | -6.31017 | 9.84 | 8.44 | 8.13 | 8 | 8.37 | 8.83 | -0.2 | -0.2 | 7.89 |
| 0.017893 | -0.21795 | 0.031157 | -0.19137 | 16.9 | 17 | 17 | 17 | 16.9 | 17 | 16.7 | 16.8 | 16.8 |
| 0.018002 | -0.27251 | 0.02902 | -0.24383 | 14.8 | 14.6 | 14.6 | 14.6 | 14.8 | 14.7 | 14.5 | 14.3 | 14.4 |
| 0.01838 | -0.2962 | 0.299351 | -0.11092 | 12.6 | 12.3 | 12.5 | 12.6 | 12.8 | 12.5 | 12.3 | 12.3 | 12.4 |
| 0.018397 | 0.296286 | 0.144452 | 0.161885 | 15.3 | 15 | 15.1 | 14.9 | 15.1 | 15 | 15.4 | 15.3 | 15.2 |
| 0.018909 | 0.28934 | 0.392766 | 0.088729 | 11 | 11.1 | 11.2 | 10.9 | 10.8 | 11 | 11.1 | 11.1 | 11.3 |
| 0.018848 | 0.333998 | 0.007819 | 0.401424 | 12.6 | 12.5 | 12.5 | 12.5 | 12.8 | 12.6 | 13.1 | 12.9 | 12.8 |
| 0.019346 | -0.20078 | 0.081049 | -0.13701 | 13.7 | 13.7 | 13.7 | 13.8 | 13.8 | 13.7 | 13.6 | 13.6 | 13.5 |
| 0.019917 | -0.34636 | 0.327139 | 0.124196 | 11 | 11.2 | 11 | 11.5 | 11.3 | 11.7 | 11.3 | 11.1 | 11.1 |
| 0.019772 | 0.510761 | 0.095596 | 0.331412 | 10.8 | 10.7 | 10.6 | 10.8 | 10.5 | 10.3 | 10.8 | 11.4 | 11.1 |
| 0.019804 | -0.34831 | 0.008389 | -0.41751 | 15 | 14.7 | 14.7 | 14.6 | 14.9 | 14.7 | 14.4 | 14.3 | 14.5 |
| 0.019883 | -0.28716 | 0.020504 | -0.28517 | 17 | 16.8 | 16.7 | 16.8 | 16.9 | 16.7 | 16.6 | 16.5 | 16.5 |
| 0.02022 | -0.5529 | 0.127384 | -0.32457 | 13 | 12.3 | 12.5 | 12.6 | 12.9 | 12.9 | 12.4 | 12.4 | 12 |

| | | | | | | | | | | | | |
|---|---|---|---|---|---|---|---|---|---|---|---|---|
| 0.020616 | -0.31043 | 0.50151 | -0.07562 | 10.4 | 10.7 | 10.5 | 10.7 | 10.9 | 10.6 | 10.4 | 10.5 | 10.4 |
| 0.020798 | 1.098751 | 0.21238 | 0.516882 | 10.3 | 11.2 | 10.3 | 9.57 | 10.8 | 9.57 | 11.1 | 11.3 | 10.9 |
| 0.020834 | -0.25408 | 0.007078 | -0.3185 | 13.3 | 13.2 | 13.1 | 13 | 13.2 | 13.2 | 12.9 | 12.9 | 12.9 |
| 0.020817 | -1.03365 | 0.008471 | -1.25071 | 13 | 12.8 | 12.6 | 12.6 | 12.3 | 12.9 | 12.5 | 11.3 | 11 |
| 0.020513 | -0.40346 | 0.021403 | -0.39956 | 15.6 | 15.2 | 15.3 | 15.2 | 15.6 | 15.3 | 15.1 | 14.9 | 14.9 |
| 0.020932 | -0.50506 | 0.733105 | -0.06193 | 10.5 | 10.4 | 10.4 | 10.7 | 11.1 | 10.9 | 10.8 | 10.3 | 10.1 |
| 0.021229 | 0.286215 | 0.181252 | 0.146205 | 12.6 | 12.8 | 12.8 | 12.8 | 12.5 | 12.5 | 12.8 | 13 | 12.9 |
| 0.021712 | 0.389285 | 0.214629 | 0.183943 | 14.5 | 14.5 | 14.4 | 14 | 14.5 | 14.3 | 14.7 | 14.8 | 14.5 |
| 0.021704 | 0.38732 | 0.139651 | 0.222776 | 13.3 | 13 | 13.1 | 12.8 | 13.2 | 12.9 | 13.5 | 13.3 | 13.2 |
| 0.02162 | -0.26507 | 0.117892 | -0.16276 | 14 | 13.7 | 13.8 | 13.9 | 14.1 | 13.9 | 13.7 | 13.7 | 13.7 |
| 0.022048 | 0.233486 | 0.080108 | 0.164773 | 13.9 | 13.9 | 13.9 | 13.9 | 13.8 | 13.8 | 13.9 | 14.2 | 14 |
| 0.022456 | -0.24318 | 0.453946 | 0.067565 | 11.3 | 11.2 | 11.3 | 11.6 | 11.5 | 11.7 | 11.2 | 11.4 | 11.4 |
| 0.022342 | 0.805019 | 0.729404 | 0.101664 | 9.71 | 9.81 | 9.74 | 9.74 | 8.57 | 8.85 | 9.86 | 9.58 | 10.1 |
| 0.022386 | -0.2788 | 0.369483 | -0.09355 | 12.6 | 12.5 | 12.7 | 12.8 | 12.7 | 12.9 | 12.6 | 12.5 | 12.4 |
| 0.02278 | -0.17736 | 5.44E-05 | -0.49543 | 13 | 13 | 13 | 12.7 | 12.7 | 12.6 | 12.5 | 12.5 | 12.5 |
| 0.02302 | -0.34507 | 0.019662 | -0.35779 | 13 | 13 | 13.1 | 13 | 12.8 | 13.2 | 12.5 | 12.8 | 12.6 |
| 0.023151 | -0.40812 | 0.028813 | -0.38737 | 13.4 | 13.6 | 13.8 | 13.8 | 13.3 | 13.7 | 13.2 | 13.1 | 13.3 |
| 0.023338 | 0.282201 | 0.176671 | 0.149109 | 14.1 | 13.9 | 13.8 | 13.7 | 13.9 | 13.7 | 14 | 14.2 | 14 |
| 0.023353 | -0.19272 | 0.007358 | -0.24618 | 15.6 | 15.7 | 15.6 | 15.7 | 15.6 | 15.5 | 15.4 | 15.4 | 15.4 |
| 0.02396 | 0.180291 | 0.979874 | -0.00168 | 13.6 | 13.6 | 13.7 | 13.4 | 13.4 | 13.4 | 13.6 | 13.6 | 13.6 |
| 0.023973 | 0.243737 | 0.347835 | 0.087153 | 16.2 | 16.4 | 16.4 | 16.3 | 16.1 | 16.3 | 16.4 | 16.5 | 16.4 |
| 0.024023 | 0.417331 | 0.289143 | 0.169984 | 11.9 | 11.7 | 11.7 | 11.2 | 11.7 | 11.6 | 12.1 | 11.8 | 11.9 |
| 0.023928 | -0.22446 | 0.034525 | -0.20524 | 16.8 | 16.5 | 16.6 | 16.7 | 16.7 | 16.7 | 16.4 | 16.4 | 16.5 |
| 0.023679 | -0.30682 | 0.018068 | -0.32635 | 14.5 | 14.1 | 14.1 | 14.1 | 14.3 | 14.2 | 14 | 13.9 | 13.9 |
| 0.024624 | 5.684165 | 0.897487 | 0.272562 | 8.84 | 8.97 | 8.69 | 9.12 | 0.49 | 0.66 | 8.73 | 9 | 9.59 |
| 0.024555 | 0.700029 | 0.057103 | 0.561899 | 13.2 | 14 | 14 | 13.8 | 13.2 | 13.7 | 14 | 14.5 | 14.4 |
| 0.024765 | 0.445749 | 0.956481 | 0.009061 | 11.2 | 11.2 | 10.9 | 10.5 | 10.7 | 10.8 | 11.3 | 11.3 | 10.8 |
| 0.02483 | -0.42788 | 0.071166 | -0.32233 | 14.5 | 14.1 | 13.9 | 14.1 | 14.3 | 14.4 | 13.9 | 13.8 | 13.8 |
| 0.025105 | -0.5264 | 0.713359 | 0.072589 | 10.8 | 10.2 | 10.5 | 11.1 | 11.4 | 10.8 | 10.7 | 10.5 | 10.4 |
| 0.025361 | 0.341284 | 0.280257 | 0.143631 | 13.5 | 13.7 | 13.7 | 13.3 | 13.4 | 13.6 | 13.9 | 13.9 | 13.6 |
| 0.025431 | -0.47781 | 0.085544 | -0.34124 | 11.4 | 11.9 | 11.9 | 11.9 | 11.6 | 12.1 | 11.2 | 11.6 | 11.4 |
| 0.025391 | -0.22959 | 0.026164 | -0.22796 | 15.1 | 14.9 | 14.9 | 14.9 | 15.1 | 15 | 14.8 | 14.7 | 14.7 |
| 0.025703 | -0.27095 | 0.004557 | -0.38754 | 12.9 | 12.8 | 12.9 | 12.9 | 12.8 | 12.6 | 12.4 | 12.5 | 12.6 |
| 0.025731 | -0.23337 | 0.005937 | -0.31781 | 12.5 | 12.4 | 12.4 | 12.3 | 12.5 | 12.3 | 12.1 | 12.2 | 12.1 |
| 0.02571 | 0.329843 | 0.108246 | 0.217682 | 11.4 | 11.4 | 11.3 | 11.5 | 11.2 | 11.1 | 11.6 | 11.4 | 11.7 |
| 0.026151 | -0.18921 | 0.043374 | -0.16646 | 15 | 14.9 | 14.9 | 15 | 14.9 | 14.9 | 14.7 | 14.8 | 14.8 |
| 0.026365 | -0.23831 | 0.007809 | -0.30966 | 13.7 | 13.9 | 13.9 | 13.9 | 13.7 | 13.7 | 13.4 | 13.6 | 13.6 |
| 0.02667 | -0.33971 | 0.101092 | -0.23177 | 13.7 | 13.6 | 13.4 | 13.6 | 13.5 | 13.9 | 13.4 | 13.2 | 13.3 |
| 0.026815 | -0.22674 | 0.308107 | -0.09089 | 14.3 | 14.4 | 14.5 | 14.6 | 14.5 | 14.5 | 14.2 | 14.4 | 14.3 |
| 0.027143 | -0.38437 | 0.676653 | -0.06139 | 10.2 | 10 | 10.2 | 10.6 | 10.5 | 10.3 | 10.1 | 9.84 | 10.3 |
| 0.0273 | -0.31458 | 0.044083 | -0.27842 | 13 | 13 | 12.8 | 12.8 | 13.1 | 12.9 | 12.5 | 12.7 | 12.7 |
| 0.027368 | -0.16744 | 0.02408 | -0.17262 | 15.8 | 15.7 | 15.7 | 15.7 | 15.8 | 15.8 | 15.6 | 15.5 | 15.6 |
| 0.027613 | 0.26834 | 0.021182 | 0.285618 | 13.3 | 13 | 13 | 13 | 13.2 | 13.1 | 13.4 | 13.4 | 13.3 |
| 0.027738 | 0.454839 | 0.139251 | 0.27753 | 11.3 | 11 | 11.2 | 11.3 | 10.6 | 11.1 | 11.6 | 11.3 | 11.4 |
| 0.028145 | -0.31393 | 0.056407 | -0.26128 | 12.3 | 11.9 | 12 | 12.1 | 12.2 | 12.1 | 11.9 | 11.9 | 11.6 |
| 0.027875 | -0.46966 | 0.015774 | -0.53511 | 10.7 | 10.4 | 10.4 | 10 | 10.6 | 10.7 | 9.92 | 9.98 | 10 |
| 0.028094 | -0.2783 | 0.017039 | -0.31237 | 15.3 | 14.9 | 15.2 | 15 | 15.2 | 15.1 | 14.9 | 14.8 | 14.9 |
| 0.02796 | 0.607988 | 0.024686 | 0.626378 | 12.5 | 13.4 | 13.3 | 13.2 | 12.8 | 13.2 | 13.7 | 13.7 | 13.7 |
| 0.028331 | 0.264378 | 0.017101 | 0.297111 | 11.7 | 11.4 | 11.5 | 11.6 | 11.6 | 11.5 | 12 | 11.7 | 11.8 |
| 0.028621 | -0.51402 | 0.498946 | -0.13612 | 10.8 | 10.8 | 10.8 | 10.8 | 11.5 | 11.2 | 10.4 | 10.6 | 10.9 |

| | | | | | | | | | | | | |
|---|---|---|---|---|---|---|---|---|---|---|---|---|
| 0.029042 | 0.462805 | 0.273286 | 0.204424 | 11.3 | 11.8 | 11.8 | 11.6 | 11.1 | 11.3 | 11.7 | 11.9 | 12 |
| 0.029863 | -0.40224 | 0.463506 | -0.11701 | 12 | 12.1 | 12.1 | 12.2 | 12.6 | 12.2 | 12.2 | 11.9 | 11.7 |
| 0.029863 | -0.35733 | 0.052533 | -0.30794 | 13.1 | 13.2 | 13.1 | 13.1 | 13.4 | 12.9 | 12.7 | 12.8 | 12.8 |
| 0.029677 | -0.21555 | 0.046846 | -0.19149 | 14.7 | 14.6 | 14.6 | 14.7 | 14.7 | 14.5 | 14.5 | 14.4 | 14.5 |
| 0.030235 | 0.249365 | 0.177566 | -0.13988 | 11.4 | 11.5 | 11.3 | 11.1 | 10.9 | 10.9 | 11.1 | 11.3 | 11.3 |
| 0.030263 | -0.358 | 0.895032 | -0.01848 | 10.9 | 10.8 | 10.6 | 11 | 11.3 | 10.9 | 10.8 | 10.6 | 10.8 |
| 0.03027 | 0.208747 | 0.377877 | 0.073895 | 14.2 | 14 | 14 | 13.9 | 14.1 | 13.9 | 14.2 | 14.1 | 14.1 |
| 0.030124 | 0.405998 | 0.373277 | 0.145013 | 12.2 | 12.3 | 12.1 | 12.1 | 12.1 | 11.6 | 12.4 | 12.5 | 12.2 |
| 0.030592 | 0.386128 | 0.300077 | 0.162694 | 12.5 | 12.4 | 12.5 | 12.3 | 11.9 | 12.4 | 12.5 | 12.8 | 12.4 |
| 0.030552 | -0.17866 | 0.044468 | -0.16215 | 15.2 | 15.3 | 15.2 | 15.2 | 15.3 | 15.2 | 15.1 | 15 | 15.1 |
| 0.030973 | 0.211572 | 0.194549 | 0.114327 | 17.6 | 17.5 | 17.5 | 17.4 | 17.5 | 17.4 | 17.7 | 17.7 | 17.5 |
| 0.03099 | 0.243307 | 0.069344 | 0.194953 | 13.6 | 13.8 | 13.7 | 13.7 | 13.7 | 13.6 | 13.7 | 14 | 13.9 |
| 0.031246 | -0.30201 | 0.187473 | -0.1665 | 12.5 | 12.7 | 12.9 | 12.8 | 12.9 | 12.9 | 12.4 | 12.6 | 12.6 |
| 0.031374 | 0.349365 | 0.291181 | 0.151092 | 15.2 | 15.6 | 15.7 | 15.4 | 15.1 | 15.4 | 15.6 | 15.7 | 15.6 |
| 0.031477 | 5.742718 | 0.98521 | -0.04203 | 9.55 | 9.11 | 8.75 | 0.1 | 9.47 | 0.49 | 8.94 | 9.21 | 9.14 |
| 0.031622 | 0.451914 | 0.464631 | 0.132994 | 10.4 | 11 | 11 | 10.5 | 10.2 | 10.6 | 10.9 | 10.8 | 11 |
| 0.031906 | -0.19771 | 0.045731 | -0.18001 | 16.3 | 16.5 | 16.5 | 16.5 | 16.3 | 16.5 | 16.2 | 16.2 | 16.3 |
| 0.032237 | 0.233425 | 0.7413 | 0.030716 | 13.9 | 13.7 | 13.7 | 13.6 | 13.6 | 13.5 | 13.8 | 13.9 | 13.7 |
| 0.032627 | 0.395294 | 0.903169 | 0.019166 | 11 | 10.7 | 11 | 10.6 | 10.7 | 10.3 | 11.2 | 10.7 | 10.8 |
| 0.032817 | 0.293605 | 0.304282 | 0.124724 | 12.1 | 12.1 | 12.2 | 12 | 12.1 | 11.7 | 12.3 | 12.1 | 12.3 |
| 0.032768 | -0.25602 | 0.05592 | -0.22179 | 15.6 | 15.3 | 15.3 | 15.4 | 15.6 | 15.4 | 15.2 | 15.2 | 15.2 |
| 0.033435 | -0.32561 | 0.135133 | -0.21059 | 15.4 | 15 | 15 | 15.1 | 15.4 | 15.2 | 15 | 14.8 | 15 |
| 0.033638 | -0.37389 | 0.021309 | -0.41728 | 10.7 | 10.7 | 10.4 | 10.4 | 10.7 | 10.6 | 10 | 10.1 | 10.4 |
| 0.033826 | 0.200312 | 0.881446 | 0.012017 | 15.1 | 15.2 | 15.1 | 15.1 | 15 | 14.8 | 15.1 | 15.2 | 15.1 |
| 0.034495 | -0.18652 | 0.011548 | -0.23936 | 14.6 | 14.5 | 14.6 | 14.5 | 14.5 | 14.5 | 14.3 | 14.4 | 14.3 |
| 0.034428 | -0.32138 | 0.015352 | -0.38813 | 12.5 | 12.5 | 12.1 | 12.1 | 12.3 | 12.5 | 12 | 12 | 11.9 |
| 0.034753 | -0.35275 | 0.02227 | -0.39295 | 12.5 | 12.2 | 12 | 12.2 | 12.3 | 12.1 | 12 | 11.7 | 11.8 |
| 0.035237 | 0.270352 | 0.545319 | 0.067237 | 14.5 | 14.4 | 14.4 | 14.1 | 14.5 | 14.1 | 14.5 | 14.5 | 14.5 |
| 0.035149 | -0.17101 | 0.031305 | -0.17607 | 16.2 | 16.2 | 16.3 | 16.3 | 16.2 | 16.3 | 16 | 16 | 16.1 |
| 0.035971 | 0.159981 | 0.642287 | 0.030564 | 15.9 | 15.9 | 15.9 | 15.8 | 15.8 | 15.7 | 16 | 15.9 | 15.9 |
| 0.035826 | 0.489596 | 0.481332 | 0.143004 | 11 | 11.2 | 11.1 | 10.7 | 11.2 | 10.4 | 11.1 | 11.4 | 11.3 |
| 0.035682 | -0.3616 | 0.135912 | -0.23723 | 12.2 | 12.2 | 11.9 | 12 | 12.5 | 12.1 | 11.9 | 11.8 | 11.9 |
| 0.035889 | -0.24228 | 0.078582 | -0.19368 | 11.9 | 11.8 | 11.7 | 11.8 | 11.9 | 11.9 | 11.7 | 11.5 | 11.6 |
| 0.036614 | -0.38766 | 0.544599 | -0.09754 | 10.8 | 10.5 | 10.7 | 10.7 | 11 | 11.2 | 10.8 | 10.5 | 10.5 |
| 0.036822 | -0.32954 | 0.339162 | -0.13347 | 11.9 | 11.9 | 11.9 | 11.9 | 12 | 12.4 | 11.8 | 11.6 | 11.8 |
| 0.036952 | -0.38179 | 0.330009 | -0.15793 | 10.9 | 10.8 | 10.4 | 10.8 | 11.1 | 10.8 | 10.5 | 10.6 | 10.6 |
| 0.036757 | 0.247979 | 0.278173 | 0.114934 | 13.8 | 13.8 | 13.7 | 13.8 | 13.5 | 13.6 | 13.7 | 13.9 | 13.9 |
| 0.036929 | -0.22669 | 0.235494 | -0.116 | 10.9 | 10.9 | 11.1 | 11 | 11.1 | 11.1 | 10.8 | 10.9 | 10.8 |
| 0.036388 | 0.204592 | 0.194114 | 0.115272 | 13.7 | 13.8 | 13.7 | 13.6 | 13.7 | 13.7 | 13.9 | 14 | 13.7 |
| 0.036824 | -0.22727 | 0.088974 | -0.17557 | 13.3 | 13.2 | 13 | 13.1 | 13.3 | 13.2 | 13 | 13 | 13 |
| 0.036577 | -0.26229 | 0.049252 | -0.24222 | 14.6 | 14.3 | 14.3 | 14.3 | 14.5 | 14.4 | 14.2 | 14.1 | 14.2 |
| 0.037028 | 0.204422 | 0.044504 | 0.19472 | 14.9 | 15.1 | 15.1 | 15.1 | 14.9 | 15.1 | 15.2 | 15.3 | 15.3 |
| 0.037159 | 0.503084 | 0.034298 | 0.513517 | 10.1 | 9.81 | 9.86 | 10.3 | 10 | 9.48 | 10.4 | 10.6 | 10.3 |
| 0.037397 | -0.33682 | 0.951962 | -0.00837 | 10.7 | 10.8 | 11 | 10.9 | 11.2 | 11.4 | 10.9 | 10.7 | 10.9 |
| 0.037305 | 0.44983 | 0.080831 | 0.359896 | 10.3 | 10.2 | 9.96 | 9.96 | 9.88 | 10.4 | 10.6 | 10.7 | 10.2 |
| 0.037647 | 4.623878 | 0.873862 | -0.30353 | 7.22 | 7.71 | 8.36 | -0 | 7.93 | 0.59 | 7.65 | 7.79 | 6.93 |
| 0.037822 | -0.16732 | 0.008642 | -0.23316 | 14 | 14 | 14 | 13.8 | 14 | 13.9 | 13.8 | 13.8 | 13.7 |
| 0.037857 | -0.68933 | 0.124174 | -0.4758 | 10.6 | 10.9 | 9.93 | 10.3 | 10.9 | 10.9 | 9.97 | 9.81 | 10.2 |
| 0.03872 | -0.26607 | 0.042024 | -0.26038 | 14.3 | 14 | 14.1 | 14.1 | 14.3 | 14.1 | 14 | 13.8 | 13.9 |
| 0.03983 | -0.30132 | 0.004323 | -0.48554 | 14.4 | 14.5 | 14.5 | 14.5 | 14.1 | 14.3 | 13.9 | 13.9 | 14.2 |

| | | | | | | | | | | | | |
|---|---|---|---|---|---|---|---|---|---|---|---|---|
| 0.039351 | 0.25372 | 0.191902 | 0.146702 | 11.8 | 11.9 | 11.8 | 11.6 | 11.8 | 11.9 | 12.1 | 11.9 | 12 |
| 0.039461 | 0.329094 | 0.19143 | 0.190651 | 14.3 | 14.1 | 14 | 13.8 | 14.1 | 14 | 14.4 | 14.3 | 14.1 |
| 0.039452 | 0.664124 | 0.132667 | 0.451526 | 10.3 | 11.3 | 10.6 | 10.8 | 10.1 | 10.6 | 11.1 | 11.1 | 11.2 |
| 0.039671 | 0.29817 | 0.057235 | 0.269484 | 10.5 | 10.6 | 10.7 | 10.6 | 10.3 | 10.8 | 10.9 | 11 | 10.8 |
| 0.039799 | 0.184858 | 0.034622 | 0.191643 | 14.1 | 14.1 | 14 | 14 | 14.2 | 14 | 14.3 | 14.3 | 14.2 |
| 0.039578 | -0.4301 | 0.044428 | -0.41705 | 11.6 | 11.7 | 11.1 | 11.5 | 11.5 | 11.5 | 11.1 | 10.9 | 11.2 |
| 0.040326 | 0.521616 | 0.579636 | 0.122804 | 10.7 | 10.6 | 10.7 | 10.6 | 9.7 | 10.4 | 10.8 | 10.8 | 10.8 |
| 0.040292 | -0.56395 | 0.011315 | -0.75662 | 11.9 | 11.2 | 11.1 | 11.3 | 11.2 | 11.1 | 10.8 | 10.8 | 10.3 |
| 0.041683 | 4.721416 | 0.684576 | -0.81843 | 8.76 | 8.73 | 9.14 | 8.7 | 0.55 | 0.76 | 7.5 | 7.86 | 8.81 |
| 0.041717 | 0.228912 | 0.923599 | -0.00932 | 12.5 | 12.6 | 12.7 | 12.3 | 12.4 | 12.3 | 12.5 | 12.5 | 12.7 |
| 0.041274 | -0.18291 | 0.348765 | -0.07474 | 15 | 14.9 | 15 | 15 | 15.1 | 15.1 | 14.9 | 15 | 14.8 |
| 0.041509 | -0.19661 | 0.098073 | -0.15158 | 14.2 | 14.3 | 14.4 | 14.5 | 14.3 | 14.3 | 14.2 | 14.2 | 14.2 |
| 0.041742 | -0.22196 | 0.017935 | -0.27238 | 15.6 | 15.5 | 15.6 | 15.5 | 15.7 | 15.4 | 15.3 | 15.3 | 15.3 |
| 0.041947 | -0.30695 | 0.089316 | -0.24504 | 14.1 | 13.7 | 13.6 | 13.7 | 14 | 13.9 | 13.6 | 13.5 | 13.6 |
| 0.041946 | -0.26123 | 0.037348 | -0.26932 | 14.8 | 14.5 | 14.8 | 14.7 | 14.8 | 14.5 | 14.5 | 14.3 | 14.4 |
| 0.04266 | -0.33632 | 0.007493 | -0.49755 | 12 | 11.7 | 11.8 | 11.9 | 11.8 | 11.4 | 11.3 | 11.3 | 11.4 |
| 0.042589 | -0.2586 | 0.030868 | -0.28093 | 19.9 | 19.6 | 19.6 | 19.6 | 19.8 | 19.6 | 19.4 | 19.4 | 19.5 |
| 0.043151 | 0.253825 | 0.08872 | 0.204634 | 16 | 16.1 | 15.9 | 15.9 | 15.9 | 16.1 | 16.3 | 16.3 | 16 |
| 0.042956 | 0.690155 | 0.075443 | 0.585901 | 9.34 | 8.52 | 9.58 | 8.66 | 9.32 | 9.15 | 9.91 | 9.57 | 9.71 |
| 0.043067 | -0.23594 | 0.034853 | -0.24936 | 15.1 | 15.2 | 15 | 15.1 | 15 | 15.2 | 15 | 14.8 | 14.8 |
| 0.044893 | -0.46307 | 0.170414 | -0.29313 | 11.5 | 11.3 | 10.9 | 11.2 | 11.5 | 11.5 | 10.7 | 11 | 11.1 |
| 0.045059 | -0.22377 | 0.031537 | -0.24551 | 12.2 | 12.4 | 12.4 | 12.3 | 12.2 | 12.4 | 12 | 12.2 | 12 |
| 0.045203 | 0.238166 | 0.253356 | 0.123361 | 12.7 | 12.8 | 12.6 | 12.4 | 12.6 | 12.7 | 12.9 | 12.8 | 12.7 |
| 0.045386 | 0.340925 | 0.115926 | 0.253464 | 13.9 | 13.6 | 13.4 | 13.4 | 13.8 | 13.4 | 13.9 | 13.9 | 13.8 |
| 0.045859 | -0.19527 | 0.058574 | -0.18223 | 15.4 | 15.6 | 15.4 | 15.4 | 15.6 | 15.5 | 15.4 | 15.3 | 15.3 |
| 0.046002 | -0.37537 | 0.248839 | -0.1974 | 11.1 | 11 | 11.2 | 11.2 | 11.6 | 11.1 | 11.1 | 10.7 | 10.9 |
| 0.046829 | 0.572774 | 0.117307 | 0.427591 | 12.6 | 13.2 | 13.3 | 13 | 12.5 | 13.2 | 13.3 | 13.6 | 13.5 |
| 0.04741 | -0.29573 | 0.79697 | 0.033505 | 9.43 | 9.57 | 9.49 | 10 | 9.87 | 9.61 | 9.49 | 9.72 | 9.38 |
| 0.047422 | -0.28996 | 0.856145 | 0.023124 | 11.6 | 11.3 | 11.3 | 11.6 | 11.9 | 11.6 | 11.4 | 11.3 | 11.5 |
| 0.048037 | -0.34924 | 0.757496 | -0.04771 | 10.6 | 10.3 | 10.2 | 10.6 | 10.9 | 10.5 | 10.4 | 10.3 | 10.3 |
| 0.048712 | 0.220548 | 0.928554 | -0.00876 | 14.4 | 14.4 | 14.3 | 14.1 | 14.3 | 14 | 14.3 | 14.4 | 14.3 |
| 0.048819 | 0.463189 | 0.550176 | 0.124081 | 11 | 10.9 | 11.4 | 11 | 10.5 | 10.7 | 11.4 | 11.1 | 11.1 |
| 0.048278 | -0.37672 | 0.423707 | -0.13601 | 11 | 11 | 10.5 | 11 | 11.2 | 11 | 10.9 | 10.6 | 10.6 |
| 0.048676 | -0.27279 | 0.363555 | -0.11283 | 14 | 14 | 13.6 | 14 | 14.1 | 14 | 13.9 | 13.6 | 13.8 |
| 0.048418 | 0.27039 | 0.090494 | 0.223446 | 12.5 | 12.6 | 12.8 | 12.7 | 12.3 | 12.6 | 12.8 | 12.9 | 12.8 |
| 0.048844 | -0.35994 | 0.076502 | -0.31507 | 15.8 | 15.3 | 15.4 | 15.4 | 15.8 | 15.5 | 15.2 | 15.1 | 15.2 |
| 0.049687 | -0.43715 | 0.936024 | -0.01562 | 9.41 | 9.14 | 9.2 | 9.64 | 9.52 | 9.86 | 9.31 | 9.56 | 8.83 |
| 0.050039 | 4.288227 | 0.891391 | 0.261322 | 10 | 8.38 | 8.65 | -0 | 7.39 | 7.62 | 9.76 | 9.47 | 8.61 |
| 0.049955 | 0.225181 | 0.732969 | 0.034385 | 13.3 | 13.4 | 13.3 | 13.3 | 13.1 | 13.1 | 13.5 | 13.2 | 13.4 |
| 0.049273 | -0.32313 | 0.682421 | -0.05905 | 13.3 | 12.8 | 12.8 | 13.4 | 13.2 | 13 | 12.9 | 12.9 | 12.9 |
| 0.049561 | -0.30666 | 0.481526 | -0.09764 | 13 | 12.8 | 12.8 | 13.1 | 13.2 | 13 | 13 | 12.9 | 12.5 |
| 0.049274 | -0.42817 | 0.33567 | -0.18894 | 11 | 10.5 | 10.7 | 10.6 | 11.2 | 11.1 | 10.6 | 10.5 | 10.4 |
| 0.049603 | 0.26471 | 0.300482 | 0.126484 | 12.7 | 12.8 | 12.8 | 12.7 | 12.4 | 12.8 | 12.8 | 12.9 | 13 |
| 0.049108 | 0.407295 | 0.132608 | 0.293828 | 12.4 | 12.3 | 12.2 | 12.2 | 12.5 | 11.8 | 12.6 | 12.7 | 12.5 |
| 0.049891 | -0.20621 | 0.049872 | -0.20623 | 13.6 | 13.6 | 13.7 | 13.6 | 13.5 | 13.8 | 13.4 | 13.4 | 13.5 |
| 0.0509 | 0.2661 | 0.835067 | 0.024866 | 12.9 | 13 | 13.1 | 12.8 | 12.6 | 12.9 | 13.2 | 13.1 | 12.8 |
| 0.051168 | 0.282388 | 0.32273 | 0.12957 | 13.1 | 13.5 | 13.5 | 13.3 | 13.1 | 13.3 | 13.5 | 13.4 | 13.6 |
| 0.051921 | 0.584959 | 0.884585 | 0.038295 | 9.15 | 9.77 | 9.8 | 8.75 | 9.27 | 9.06 | 9.61 | 10 | 9.21 |
| 0.051874 | 0.336736 | 0.318738 | 0.156478 | 11.3 | 11.4 | 11.5 | 11.3 | 10.9 | 11.3 | 11.4 | 11.7 | 11.5 |
| 0.052002 | -0.18953 | 0.008993 | -0.2856 | 14.9 | 14.7 | 14.8 | 14.6 | 14.8 | 14.7 | 14.5 | 14.4 | 14.5 |

| | | | | | | | | | | | | |
|---|---|---|---|---|---|---|---|---|---|---|---|---|
| 0.052171 | -0.24636 | 0.162656 | -0.16599 | 12.6 | 12.7 | 12.8 | 12.7 | 12.8 | 12.9 | 12.4 | 12.7 | 12.6 |
| 0.052848 | 0.229072 | 0.615191 | 0.052602 | 15.2 | 15 | 14.9 | 14.8 | 14.9 | 14.9 | 15.2 | 15.1 | 15 |
| 0.052935 | 0.269339 | 0.187388 | 0.170789 | 15.7 | 15.4 | 15.3 | 15.2 | 15.6 | 15.3 | 15.7 | 15.6 | 15.6 |
| 0.054394 | 0.159238 | 0.21551 | 0.094923 | 15 | 15.1 | 15.1 | 15 | 15 | 15 | 15.1 | 15.2 | 15.2 |
| 0.054457 | -0.22366 | 0.069519 | -0.20802 | 16.3 | 16.5 | 16.4 | 16.5 | 16.3 | 16.5 | 16.1 | 16.2 | 16.3 |
| 0.054763 | -0.45112 | 0.285445 | -0.22923 | 11.6 | 11.2 | 11.1 | 11.4 | 11.7 | 11.4 | 11.5 | 10.8 | 10.9 |
| 0.05489 | 0.420947 | 0.140391 | 0.306259 | 12.7 | 13.3 | 13.4 | 13.1 | 12.8 | 13.2 | 13.3 | 13.5 | 13.5 |
| 0.055277 | 0.196929 | 0.069448 | 0.184 | 13.9 | 14.1 | 14.1 | 14.2 | 14 | 14 | 14.2 | 14.3 | 14.2 |
| 0.056505 | -0.35809 | 0.398901 | -0.14288 | 13.1 | 12.4 | 12.5 | 12.9 | 12.9 | 12.8 | 12.5 | 12.5 | 12.6 |
| 0.056565 | -0.32489 | 0.081507 | -0.29047 | 12 | 12.1 | 12 | 12.1 | 12.3 | 11.9 | 12 | 11.6 | 11.6 |
| 0.05682 | -0.28619 | 0.002239 | -0.57145 | 11.6 | 11.4 | 11.2 | 11.1 | 11.1 | 11.1 | 10.7 | 10.8 | 11.1 |
| 0.057446 | -0.40015 | 0.841069 | -0.03728 | 10.3 | 10 | 10.3 | 10.3 | 10.6 | 10.8 | 10.3 | 10.3 | 9.8 |
| 0.057235 | 0.291269 | 0.495692 | 0.093415 | 13.5 | 13.3 | 13.5 | 13.4 | 13.4 | 12.9 | 13.6 | 13.5 | 13.5 |
| 0.056994 | -0.14096 | 0.030189 | -0.16699 | 15.4 | 15.3 | 15.4 | 15.3 | 15.3 | 15.3 | 15.2 | 15.1 | 15.2 |
| 0.057545 | -0.37028 | 0.070116 | -0.34898 | 12.1 | 12.1 | 11.8 | 12 | 12.4 | 11.7 | 11.6 | 11.7 | 11.7 |
| 0.057507 | -0.13948 | 0.056137 | -0.14046 | 16 | 16.1 | 16 | 16 | 16 | 16 | 15.8 | 15.8 | 15.9 |
| 0.058379 | 0.155143 | 0.106402 | 0.12778 | 15.1 | 15.2 | 15.2 | 15.2 | 15.2 | 15.1 | 15.3 | 15.3 | 15.3 |
| 0.058821 | -0.31918 | 0.075984 | -0.29522 | 14.5 | 14 | 13.9 | 14.1 | 14.3 | 14.1 | 13.9 | 13.9 | 13.8 |
| 0.059314 | 0.38965 | 0.756683 | 0.056717 | 11.1 | 11.4 | 11.7 | 11.4 | 10.9 | 10.9 | 11.6 | 11.3 | 11.4 |
| 0.059217 | -0.40906 | 0.053289 | -0.42172 | 12.3 | 11.8 | 11.7 | 11.7 | 12.2 | 11.9 | 11.6 | 11.5 | 11.5 |
| 0.060104 | -0.39794 | 0.198575 | -0.25452 | 13 | 13.3 | 13.4 | 13.6 | 13 | 13.5 | 13 | 13.1 | 12.8 |
| 0.060522 | -0.25211 | 0.08433 | -0.22735 | 14.3 | 14.6 | 14.7 | 14.6 | 14.5 | 14.5 | 14.2 | 14.3 | 14.4 |
| 0.060829 | -0.19813 | 0.258681 | -0.11027 | 13.7 | 13.5 | 13.4 | 13.6 | 13.6 | 13.6 | 13.4 | 13.6 | 13.3 |
| 0.061435 | -0.15395 | 0.015961 | -0.21603 | 14.1 | 14.2 | 14.2 | 14.1 | 14 | 14.1 | 13.8 | 13.9 | 14 |
| 0.062979 | -0.17819 | 0.008333 | -0.2882 | 15 | 15.2 | 15.2 | 15.1 | 15 | 15 | 14.8 | 14.8 | 14.9 |
| 0.063181 | 0.335041 | 0.385741 | 0.142238 | 11 | 11 | 10.9 | 10.6 | 11 | 10.7 | 11.4 | 11.1 | 10.9 |
| 0.063679 | -0.2295 | 0.067607 | -0.22538 | 11.1 | 11.1 | 10.9 | 11.1 | 10.8 | 11.1 | 10.8 | 10.9 | 10.6 |
| 0.064142 | 0.300531 | 0.327055 | 0.145857 | 13.3 | 13.7 | 13.7 | 13.6 | 13.3 | 13.3 | 13.7 | 13.8 | 13.7 |
| 0.064125 | -0.13121 | 0.25288 | -0.07518 | 14.4 | 14.4 | 14.4 | 14.5 | 14.5 | 14.5 | 14.3 | 14.4 | 14.4 |
| 0.06436 | 0.670344 | 0.199641 | 0.436414 | 14.2 | 15.3 | 15.3 | 15.1 | 14.3 | 14.7 | 15.4 | 15.3 | 15.4 |
| 0.064378 | -4.48837 | 0.174699 | -3.11384 | 8.07 | 8.39 | 9.44 | 8.45 | 10.9 | 10.7 | 0.12 | 9.34 | 7.11 |
| 0.064731 | -0.35611 | 0.390204 | -0.15079 | 9.73 | 9.47 | 9.9 | 9.92 | 9.67 | 10.1 | 9.57 | 9.74 | 9.34 |
| 0.064621 | -0.52425 | 0.097994 | -0.45817 | 11.6 | 11 | 12 | 11.7 | 11.7 | 11.5 | 11.4 | 10.9 | 10.9 |
| 0.06504 | -3.85884 | 0.109223 | -3.25086 | 8.02 | 8.49 | 7.8 | 8.03 | 8.98 | 9.11 | -0.1 | 7.98 | 6.63 |
| 0.065734 | 0.214361 | 0.585108 | -0.05708 | 11.1 | 11.3 | 11.2 | 10.9 | 10.9 | 11.1 | 11.3 | 11 | 11.2 |
| 0.065699 | 0.242911 | 0.7712 | -0.0342 | 11.6 | 11.6 | 11.2 | 11.1 | 11.3 | 11.1 | 11.4 | 11.5 | 11.4 |
| 0.065929 | 0.372501 | 0.469859 | -0.13242 | 11.2 | 10.5 | 10.8 | 10.3 | 10.3 | 10.4 | 10.6 | 10.9 | 10.6 |
| 0.066166 | -0.28155 | 0.087241 | -0.25785 | 14.1 | 13.6 | 13.9 | 13.8 | 14 | 13.9 | 13.5 | 13.7 | 13.6 |
| 0.066763 | -0.22583 | 0.02035 | -0.30783 | 12.5 | 12.4 | 12.2 | 12.4 | 12.3 | 12.2 | 12.1 | 11.9 | 12.1 |
| 0.067315 | 0.192634 | 0.450553 | 0.072062 | 10.7 | 10.8 | 10.6 | 10.6 | 10.5 | 10.7 | 10.9 | 10.7 | 10.8 |
| 0.067208 | 0.416624 | 0.238532 | 0.250316 | 13.1 | 13.6 | 13.7 | 13.4 | 12.9 | 13.5 | 13.6 | 13.7 | 13.8 |
| 0.06728 | -0.30435 | 0.083565 | -0.28417 | 13.3 | 13 | 13.2 | 13.2 | 12.9 | 13.4 | 12.9 | 12.8 | 12.9 |
| 0.067534 | -0.13709 | 0.056889 | -0.14427 | 14.7 | 14.7 | 14.7 | 14.6 | 14.7 | 14.7 | 14.6 | 14.5 | 14.6 |
| 0.067729 | 0.438201 | 0.801079 | -0.05391 | 10.1 | 10.5 | 10.7 | 9.5 | 10.2 | 10.1 | 10.4 | 10.4 | 10.3 |
| 0.068051 | 0.201839 | 0.143741 | 0.154995 | 13.2 | 12.9 | 13.2 | 13 | 13 | 13.2 | 13.2 | 13.2 | 13.3 |
| 0.068825 | -0.68862 | 0.397728 | -0.29216 | 12.7 | 11.3 | 11.7 | 12.3 | 12.5 | 12.1 | 11.7 | 11.7 | 11.5 |
| 0.069171 | -0.25732 | 0.057725 | -0.27164 | 14.7 | 14.5 | 14.6 | 14.6 | 14.8 | 14.4 | 14.4 | 14.2 | 14.3 |
| 0.069535 | 0.318184 | 0.176354 | 0.225011 | 12 | 11.9 | 11.9 | 11.6 | 12 | 11.8 | 12.2 | 12.3 | 11.9 |
| 0.07042 | 0.266149 | 0.311435 | 0.137566 | 11.5 | 11.5 | 11.8 | 11.7 | 11.3 | 11.4 | 11.8 | 11.7 | 11.8 |
| 0.07119 | -0.17233 | 0.551378 | -0.05141 | 14.3 | 14.2 | 14.3 | 14.4 | 14.6 | 14.3 | 14.3 | 14.3 | 14.2 |

| | | | | | | | | | | | | |
|---|---|---|---|---|---|---|---|---|---|---|---|---|
| 0.071063 | 0.531435 | 0.309123 | 0.276875 | 9.91 | 9.84 | 10.1 | 10.4 | 9.43 | 9.35 | 10.4 | 10.1 | 10.2 |
| 0.071565 | 0.23584 | 0.332078 | 0.117057 | 15.7 | 15.9 | 15.9 | 15.8 | 15.5 | 15.8 | 15.8 | 16 | 16.1 |
| 0.072127 | -0.30673 | 0.096304 | -0.27894 | 14.7 | 14.3 | 14.2 | 14.3 | 14.7 | 14.4 | 14.2 | 14.2 | 14.1 |
| 0.073018 | 0.286853 | 0.151826 | 0.220043 | 12.6 | 12.4 | 12.6 | 12.3 | 12.7 | 12.4 | 12.9 | 12.7 | 12.6 |
| 0.073699 | -0.35827 | 0.053761 | -0.39373 | 16.8 | 17.2 | 17.2 | 17.2 | 16.7 | 17.2 | 16.6 | 16.6 | 16.8 |
| 0.074514 | -0.21562 | 0.166616 | -0.15986 | 11.7 | 11.8 | 11.7 | 11.9 | 11.7 | 11.8 | 11.7 | 11.5 | 11.5 |
| 0.076082 | 0.293082 | 0.563126 | 0.086592 | 13.6 | 13.4 | 13.4 | 13.3 | 13.4 | 13.1 | 13.8 | 13.5 | 13.3 |
| 0.075968 | -0.61803 | 0.368884 | -0.28825 | 10.1 | 10.2 | 10.2 | 10.9 | 9.71 | 10.8 | 10.1 | 9.53 | 9.89 |
| 0.076347 | 0.646584 | 0.347586 | 0.31654 | 10.7 | 11.9 | 11.5 | 11.2 | 10.7 | 11.1 | 11.2 | 11.9 | 11.8 |
| 0.076343 | 0.190939 | 0.328105 | 0.097569 | 15.8 | 15.6 | 15.6 | 15.5 | 15.6 | 15.5 | 15.8 | 15.8 | 15.6 |
| 0.076328 | -0.18253 | 0.021844 | -0.25522 | 12.2 | 12.4 | 12.1 | 12.3 | 12.1 | 12.1 | 11.9 | 12 | 12 |
| 0.075578 | 0.310638 | 0.117126 | 0.267148 | 11.4 | 11.1 | 11.5 | 11.4 | 11.3 | 11.2 | 11.6 | 11.8 | 11.4 |
| 0.076935 | -0.31254 | 0.265896 | -0.18391 | 11 | 11.4 | 11.5 | 11.6 | 11.2 | 11.5 | 11.2 | 11.2 | 11.1 |
| 0.077153 | 1.124507 | 0.206416 | 0.761581 | 12.6 | 14.2 | 14.2 | 13.9 | 12.2 | 13.8 | 14.2 | 14.5 | 14.6 |
| 0.077808 | -0.1616 | 0.022435 | -0.22594 | 14.9 | 14.9 | 14.9 | 15 | 14.7 | 14.8 | 14.7 | 14.7 | 14.7 |
| 0.077936 | -0.13321 | 0.1094 | -0.11864 | 16.1 | 16.1 | 16.1 | 16.2 | 16.1 | 16.1 | 15.9 | 16 | 16 |
| 0.078771 | -0.23373 | 0.060541 | -0.25347 | 14.7 | 14.5 | 14.4 | 14.4 | 14.7 | 14.5 | 14.3 | 14.2 | 14.4 |
| 0.078946 | -0.25271 | 0.17782 | -0.18529 | 14.8 | 14.6 | 14.5 | 14.6 | 14.9 | 14.5 | 14.5 | 14.5 | 14.4 |
| 0.07973 | 0.284804 | 0.494654 | 0.101343 | 15.5 | 15.9 | 15.9 | 15.7 | 15.4 | 15.6 | 15.8 | 15.8 | 16 |
| 0.079637 | -0.36595 | 0.211085 | -0.24754 | 12.3 | 12.3 | 11.9 | 12.1 | 12.7 | 12.1 | 12.1 | 11.9 | 11.8 |
| 0.079601 | 0.138109 | 0.170112 | 0.10361 | 14.3 | 14.3 | 14.3 | 14.2 | 14.3 | 14.3 | 14.4 | 14.5 | 14.4 |
| 0.080022 | -0.22467 | 0.054805 | -0.25208 | 14.6 | 14.4 | 14.4 | 14.3 | 14.6 | 14.4 | 14.3 | 14.1 | 14.2 |
| 0.080529 | 0.33452 | 0.51488 | 0.113717 | 15 | 14.6 | 14.4 | 14.3 | 14.4 | 14.6 | 15 | 14.7 | 14.7 |
| 0.08117 | 0.213019 | 0.675304 | -0.0463 | 11.7 | 11.7 | 11.7 | 11.5 | 11.2 | 11.5 | 11.5 | 11.6 | 11.8 |
| 0.081965 | -0.24651 | 0.221248 | -0.16425 | 12.6 | 12.5 | 12.5 | 12.6 | 12.5 | 12.7 | 12.6 | 12.3 | 12.2 |
| 0.082476 | 0.427359 | 0.012937 | 0.687509 | 10 | 10.5 | 10.2 | 10.6 | 10.2 | 10.7 | 10.6 | 11.1 | 11.1 |
| 0.082251 | -0.28789 | 0.410103 | -0.12572 | 13.4 | 13 | 13.4 | 13.3 | 13.6 | 13.5 | 13.2 | 13.3 | 13 |
| 0.082502 | -0.29144 | 0.346225 | -0.14673 | 11.8 | 11.5 | 11.6 | 11.7 | 12.1 | 11.6 | 11.6 | 11.5 | 11.4 |
| 0.082757 | -0.21327 | 0.155411 | -0.1686 | 17.4 | 17.2 | 17.1 | 17.2 | 17.5 | 17.2 | 17.2 | 17 | 17.1 |
| 0.083549 | -0.18054 | 0.06709 | -0.19351 | 14.6 | 14.6 | 14.5 | 14.5 | 14.7 | 14.5 | 14.4 | 14.5 | 14.3 |
| 0.084548 | -0.14701 | 0.029722 | -0.19747 | 13.9 | 13.9 | 13.9 | 13.8 | 13.8 | 14 | 13.6 | 13.8 | 13.7 |
| 0.084438 | 0.188297 | 0.158895 | 0.148419 | 12.5 | 12.2 | 12.3 | 12.3 | 12.3 | 12.3 | 12.5 | 12.5 | 12.4 |
| 0.084629 | -0.28794 | 0.127425 | -0.24871 | 15.4 | 15.6 | 15.6 | 15.7 | 15.3 | 15.7 | 15.1 | 15.3 | 15.4 |
| 0.08483 | -0.21207 | 0.060446 | -0.23572 | 12.6 | 12.7 | 12.3 | 12.4 | 12.6 | 12.6 | 12.3 | 12.3 | 12.3 |
| 0.084191 | -0.19899 | 0.092944 | -0.1925 | 13.8 | 14 | 13.9 | 14 | 13.8 | 13.9 | 13.6 | 13.7 | 13.8 |
| 0.084889 | -0.21909 | 0.087083 | -0.21724 | 12.6 | 12.7 | 12.7 | 12.8 | 12.5 | 12.8 | 12.3 | 12.6 | 12.5 |
| 0.088535 | 0.273575 | 0.855729 | -0.02643 | 11 | 11.1 | 10.8 | 10.6 | 10.7 | 10.6 | 10.8 | 10.8 | 11.2 |
| 0.088725 | -0.36517 | 0.288331 | -0.21406 | 11.6 | 11.3 | 11.4 | 11.6 | 11.9 | 11.2 | 11.1 | 11.3 | 11.3 |
| 0.089589 | -0.20163 | 0.273595 | -0.12257 | 12.4 | 12.2 | 12.1 | 12.1 | 12.3 | 12.4 | 12.2 | 12.1 | 12 |
| 0.089879 | -0.45591 | 0.275899 | -0.27601 | 11.2 | 10.7 | 10.5 | 10.9 | 11.2 | 10.9 | 10.9 | 10.6 | 10.1 |
| 0.090358 | -0.16588 | 0.084099 | -0.1699 | 13.7 | 13.9 | 13.7 | 13.7 | 13.7 | 13.9 | 13.6 | 13.6 | 13.6 |
| 0.09084 | 0.97277 | 0.332818 | 0.520838 | 8.22 | 9.42 | 10 | 9.67 | 8.43 | 8.2 | 9.89 | 9.64 | 9.69 |
| 0.091161 | 0.158473 | 0.887048 | -0.01208 | 13.1 | 13.2 | 13 | 13 | 12.9 | 12.9 | 13.1 | 13.2 | 13 |
| 0.091683 | 0.197071 | 0.631312 | 0.051198 | 13.5 | 13.7 | 13.7 | 13.7 | 13.4 | 13.5 | 13.7 | 13.8 | 13.7 |
| 0.091342 | 0.636516 | 0.18394 | 0.482086 | 9.79 | 9.47 | 8.82 | 8.94 | 9.68 | 9 | 9.32 | 10.2 | 10 |
| 0.091564 | -0.31273 | 0.083091 | -0.32303 | 11.4 | 11.9 | 11.7 | 11.7 | 11.8 | 11.5 | 11.6 | 11.4 | 11.1 |
| 0.092093 | -0.20701 | 0.064505 | -0.23197 | 13.8 | 13.6 | 13.6 | 13.5 | 13.7 | 13.8 | 13.5 | 13.5 | 13.4 |
| 0.092798 | -0.6288 | 0.144981 | -0.53187 | 8.65 | 9.22 | 8.86 | 8.71 | 9.37 | 8.94 | 7.76 | 9.05 | 8.32 |
| 0.093723 | -0.40378 | 0.311593 | -0.22898 | 11.4 | 11.1 | 11.6 | 11.6 | 11.5 | 11.5 | 11 | 11.5 | 10.8 |
| 0.093621 | -0.1845 | 0.265779 | -0.11588 | 14.8 | 14.5 | 14.5 | 14.7 | 14.7 | 14.7 | 14.5 | 14.5 | 14.4 |

| | | | | | | | | | | | | |
|---|---|---|---|---|---|---|---|---|---|---|---|---|
| 0.094344 | 0.234692 | 0.763499 | -0.03842 | 12.1 | 12.4 | 12.5 | 12.1 | 11.8 | 12.1 | 12.3 | 12.2 | 12.3 |
| 0.094723 | 0.576078 | 0.059718 | 0.666935 | 9.16 | 10.2 | 9.48 | 9.72 | 9.23 | 10.2 | 10.3 | 10.3 | 10.3 |
| 0.094569 | -0.16621 | 0.125303 | -0.15001 | 14.1 | 14.1 | 13.9 | 14 | 14.1 | 14.1 | 14 | 13.8 | 13.9 |
| 0.094224 | -0.14249 | 0.098439 | -0.14035 | 16.9 | 16.9 | 17 | 17 | 16.9 | 16.9 | 16.7 | 16.8 | 16.9 |
| 0.095087 | -0.13095 | 0.615838 | -0.03604 | 13.8 | 13.8 | 13.8 | 13.8 | 13.9 | 13.9 | 13.7 | 13.7 | 13.8 |
| 0.096186 | -0.23489 | 0.069219 | -0.26149 | 13.7 | 13.6 | 13.6 | 13.5 | 13.9 | 13.5 | 13.3 | 13.4 | 13.5 |
| 0.096477 | 0.205623 | 0.408801 | -0.09502 | 13.3 | 13.5 | 13.6 | 13.1 | 13.1 | 13.2 | 13.2 | 13.4 | 13.5 |
| 0.09722 | -0.20603 | 0.05322 | -0.24884 | 15 | 14.7 | 14.8 | 14.7 | 15 | 14.7 | 14.6 | 14.6 | 14.6 |
| 0.099138 | -0.31578 | 0.425489 | -0.1419 | 12.3 | 12.2 | 11.6 | 12.3 | 12.2 | 12.1 | 11.9 | 11.9 | 11.9 |
| 0.100404 | 0.143402 | 0.114862 | 0.136587 | 15.2 | 15.3 | 15.1 | 15.2 | 15.1 | 15.3 | 15.3 | 15.4 | 15.4 |
| 0.100764 | -0.21821 | 0.108772 | -0.21231 | 12.2 | 12.2 | 12 | 12.2 | 12.3 | 11.9 | 11.8 | 11.9 | 12 |
| 0.100595 | -0.24306 | 0.10753 | -0.23734 | 14.8 | 14.4 | 14.4 | 14.5 | 14.7 | 14.3 | 14.3 | 14.2 | 14.2 |
| 0.101208 | -0.12479 | 0.187291 | -0.0971 | 14.6 | 14.6 | 14.7 | 14.7 | 14.7 | 14.7 | 14.6 | 14.6 | 14.5 |
| 0.102054 | -0.29999 | 0.104742 | -0.29722 | 11.4 | 11 | 11.4 | 11.1 | 11.5 | 11.1 | 10.9 | 11 | 11 |
| 0.103733 | -0.4005 | 0.560916 | -0.1321 | 12.7 | 12.3 | 12 | 12.2 | 12.9 | 12.7 | 12.3 | 12.1 | 12.2 |
| 0.103757 | -0.28065 | 0.394851 | -0.13725 | 10.8 | 11.2 | 10.8 | 10.9 | 11.2 | 11.2 | 10.7 | 10.7 | 11 |
| 0.103714 | -0.25097 | 0.17472 | -0.20347 | 12.8 | 12.6 | 12.4 | 12.4 | 12.7 | 12.7 | 12.4 | 12.3 | 12.4 |
| 0.10462 | 0.150091 | 0.62816 | 0.041213 | 16.7 | 16.6 | 16.5 | 16.5 | 16.6 | 16.4 | 16.7 | 16.6 | 16.6 |
| 0.104726 | 0.300289 | 0.37843 | 0.1528 | 10.1 | 10.4 | 10.2 | 10.1 | 9.76 | 10.3 | 10.6 | 10.4 | 10.2 |
| 0.106688 | -0.21018 | 0.128465 | -0.19602 | 13.4 | 13.1 | 12.9 | 13.2 | 13.1 | 13.1 | 13 | 12.8 | 13 |
| 0.107028 | -0.20008 | 0.157281 | -0.1719 | 13.3 | 13.2 | 13.1 | 13.2 | 13.3 | 13.3 | 13.2 | 13 | 13 |
| 0.107332 | -0.20164 | 0.22155 | -0.14727 | 12.1 | 12.2 | 12.3 | 12.3 | 12.1 | 12.4 | 11.9 | 12 | 12.2 |
| 0.10803 | -0.21382 | 0.020164 | -0.34257 | 12.2 | 12.6 | 12.5 | 12.2 | 12.4 | 12.3 | 12.1 | 12.1 | 12.1 |
| 0.10771 | -0.14786 | 0.401401 | -0.07223 | 16.1 | 16 | 16 | 16.1 | 16.2 | 16 | 16 | 15.9 | 16 |
| 0.108426 | 0.319752 | 0.308719 | 0.192113 | 11.5 | 11.1 | 11.2 | 11.1 | 11.5 | 10.8 | 11.4 | 11.5 | 11.4 |
| 0.108342 | 0.52369 | 0.277841 | 0.336981 | 13.3 | 14 | 14 | 13.9 | 13 | 13.8 | 13.9 | 14.2 | 14.3 |
| 0.107828 | 0.502072 | 0.052079 | 0.632954 | 9.09 | 8.45 | 9.05 | 9.15 | 9.43 | 8.4 | 9.58 | 9.35 | 9.56 |
| 0.108965 | 0.207743 | 0.932365 | 0.010066 | 14.4 | 14.6 | 14.5 | 14.5 | 14.1 | 14.3 | 14.5 | 14.5 | 14.4 |
| 0.108952 | -0.19822 | 0.211505 | -0.14912 | 15.2 | 15 | 14.9 | 15 | 15.2 | 15.1 | 14.9 | 14.7 | 15 |
| 0.109559 | 0.845862 | 0.235147 | 0.602705 | 12.9 | 14.3 | 14.3 | 14 | 12.8 | 14 | 14.1 | 14.6 | 14.6 |
| 0.111415 | -0.27735 | 0.979101 | -0.00418 | 10.8 | 11.3 | 11.1 | 11.5 | 11.3 | 11.2 | 11.1 | 10.9 | 11.2 |
| 0.111606 | 0.620166 | 0.712794 | 0.132057 | 9.77 | 9.1 | 9.07 | 9.53 | 8.09 | 8.85 | 9.5 | 9.46 | 9.37 |
| 0.111852 | -0.12723 | 0.696329 | -0.02876 | 14.7 | 14.8 | 14.8 | 14.9 | 14.8 | 14.9 | 14.7 | 14.7 | 14.8 |
| 0.111936 | -0.1558 | 0.02023 | -0.25269 | 12.8 | 12.8 | 12.8 | 12.8 | 12.6 | 12.7 | 12.6 | 12.4 | 12.5 |
| 0.111868 | -0.17514 | 0.036805 | -0.24576 | 13.4 | 13.2 | 13.1 | 13.1 | 13.2 | 13.3 | 13 | 13 | 13 |
| 0.111173 | -0.27367 | 0.285265 | -0.17483 | 12.9 | 12.4 | 12.5 | 12.5 | 12.7 | 12.8 | 12.4 | 12.3 | 12.6 |
| 0.111904 | -0.22862 | 0.272366 | -0.15071 | 15 | 14.7 | 14.5 | 14.7 | 15 | 14.7 | 14.7 | 14.6 | 14.5 |
| 0.111663 | -0.20453 | 0.069235 | -0.24013 | 12.9 | 12.9 | 12.7 | 12.6 | 13 | 12.8 | 12.6 | 12.6 | 12.5 |
| 0.112704 | -0.21362 | 0.596598 | -0.06597 | 11.1 | 11.4 | 11.5 | 11.5 | 11.6 | 11.3 | 11.3 | 11.2 | 11.2 |
| 0.11307 | 0.447288 | 0.318294 | 0.267185 | 12.5 | 13.2 | 13.3 | 13 | 12.7 | 12.8 | 13 | 13.5 | 13.4 |
| 0.112874 | 0.191771 | 0.256055 | 0.131615 | 14.4 | 14.3 | 14.2 | 14.1 | 14.4 | 14.2 | 14.5 | 14.4 | 14.3 |
| 0.112835 | -0.12282 | 0.069877 | -0.14435 | 15.5 | 15.6 | 15.6 | 15.6 | 15.5 | 15.6 | 15.4 | 15.4 | 15.4 |
| 0.114098 | -0.79918 | 0.187162 | -0.64962 | 13.4 | 12 | 12.2 | 12.4 | 13.5 | 12.3 | 11.7 | 12.4 | 11.6 |
| 0.114552 | 0.194094 | 0.577787 | -0.06347 | 14 | 13.7 | 13.6 | 13.5 | 13.6 | 13.4 | 13.7 | 13.7 | 13.7 |
| 0.114572 | 3.340887 | 0.905767 | 0.230065 | 8.85 | 8.17 | 9.74 | 8.77 | 7.99 | 0.67 | 8.76 | 9.31 | 9.39 |
| 0.115002 | 0.629181 | 0.282127 | 0.409877 | 9.94 | 11.2 | 11.2 | 10.9 | 10.1 | 10.7 | 11.1 | 11.2 | 11.2 |
| 0.116005 | 0.149408 | 0.738188 | 0.029282 | 13.6 | 13.5 | 13.5 | 13.4 | 13.4 | 13.5 | 13.7 | 13.6 | 13.4 |
| 0.117228 | 0.233803 | 0.96302 | -0.00636 | 13.4 | 13.2 | 13.4 | 12.9 | 13.2 | 13.3 | 13.6 | 13.3 | 13.3 |
| 0.117476 | -0.24697 | 0.26667 | -0.16793 | 12.7 | 12.3 | 12.2 | 12.4 | 12.6 | 12.4 | 12.5 | 12.1 | 12.1 |
| 0.118946 | -0.29285 | 0.337583 | -0.17086 | 10.7 | 10.6 | 10.4 | 10.6 | 10.9 | 10.5 | 10.5 | 10.6 | 10.1 |

| | | | | | | | | | | | | |
|---|---|---|---|---|---|---|---|---|---|---|---|---|
| 0.118939 | -0.20555 | 0.146787 | -0.18909 | 14 | 13.8 | 13.6 | 13.6 | 13.9 | 13.8 | 13.7 | 13.5 | 13.6 |
| 0.118686 | -0.45471 | 0.105726 | -0.47449 | 9.34 | 8.49 | 8.41 | 8.71 | 9.06 | 8.41 | 8.29 | 8.2 | 8.32 |
| 0.119429 | 0.13124 | 0.293376 | 0.084562 | 14.4 | 14.5 | 14.3 | 14.4 | 14.3 | 14.4 | 14.5 | 14.5 | 14.4 |
| 0.120269 | -0.2556 | 0.034919 | -0.3735 | 13.9 | 13.5 | 13.7 | 13.3 | 13.8 | 13.6 | 13.4 | 13.3 | 13.3 |
| 0.120063 | -0.60186 | 0.088969 | -0.66979 | 14.4 | 13.1 | 13.2 | 13.2 | 13.9 | 13.3 | 12.8 | 12.8 | 12.9 |
| 0.120779 | -0.17615 | 0.051822 | -0.23195 | 15.4 | 15.2 | 15.2 | 15.1 | 15.3 | 15.2 | 15.1 | 14.9 | 15 |
| 0.121625 | 0.287731 | 0.545066 | -0.10484 | 10.9 | 11.2 | 11.2 | 10.6 | 10.6 | 11 | 10.8 | 11.3 | 10.9 |
| 0.121561 | -0.41365 | 0.173676 | -0.35656 | 10.4 | 9.57 | 9.6 | 9.77 | 9.87 | 10.1 | 9.7 | 9.57 | 9.24 |
| 0.12283 | -0.21907 | 0.197212 | -0.17852 | 12.2 | 12.2 | 12.1 | 12.2 | 12.3 | 12.2 | 12.2 | 12.1 | 11.7 |
| 0.123408 | 0.203197 | 0.98942 | -0.00161 | 13.3 | 13.7 | 13.4 | 13.3 | 13.4 | 13.1 | 13.5 | 13.4 | 13.5 |
| 0.123201 | 0.332831 | 0.370653 | 0.182985 | 13.4 | 14 | 13.8 | 13.9 | 13.3 | 13.5 | 13.8 | 13.9 | 14 |
| 0.124094 | -0.19771 | 0.874836 | 0.018679 | 11.1 | 11.3 | 11.5 | 11.4 | 11.6 | 11.5 | 11.2 | 11.2 | 11.4 |
| 0.12394 | -0.11893 | 0.114271 | -0.12264 | 15.2 | 15.1 | 15.1 | 15.1 | 15.2 | 15.1 | 15.1 | 14.9 | 15 |
| 0.124839 | 0.419382 | 0.31313 | 0.26305 | 14.6 | 15.5 | 15.4 | 15.3 | 14.7 | 15.1 | 15.4 | 15.5 | 15.4 |
| 0.125486 | -0.16561 | 0.05597 | -0.21649 | 13.2 | 13 | 13 | 13.1 | 13 | 13 | 13 | 12.8 | 12.8 |
| 0.127299 | 0.202133 | 0.257114 | 0.144851 | 9.5 | 9.37 | 9.63 | 9.39 | 9.53 | 9.4 | 9.69 | 9.82 | 9.43 |
| 0.127436 | 0.425899 | 0.25567 | 0.306344 | 9.3 | 9.95 | 8.99 | 9.57 | 9.23 | 9.07 | 9.51 | 9.73 | 9.92 |
| 0.127496 | 0.176644 | 0.200573 | 0.144825 | 14.8 | 14.7 | 14.6 | 14.7 | 14.8 | 14.5 | 14.7 | 14.9 | 14.9 |
| 0.128152 | -0.24569 | 0.093714 | -0.27547 | 14.3 | 14 | 13.9 | 14 | 14.3 | 13.9 | 13.8 | 13.7 | 13.9 |
| 0.128478 | 0.122026 | 0.94856 | 0.004783 | 14.9 | 15 | 15 | 14.8 | 14.9 | 14.9 | 15 | 15 | 15 |
| 0.129515 | -0.21383 | 0.099512 | -0.23578 | 12.2 | 12.1 | 12.2 | 12.3 | 12.3 | 11.8 | 11.8 | 12 | 12 |
| 0.130218 | -0.27226 | 0.029233 | -0.42888 | 13.6 | 13.4 | 13.1 | 13.2 | 13.4 | 13 | 12.9 | 13 | 12.9 |
| 0.130571 | -0.39372 | 0.934825 | 0.019686 | 9.92 | 10.6 | 9.92 | 10.3 | 10.4 | 11 | 9.92 | 10.3 | 10.3 |
| 0.131622 | 0.21355 | 0.857373 | -0.02356 | 12.7 | 12.3 | 12.5 | 12.3 | 12.4 | 12.1 | 12.5 | 12.5 | 12.4 |
| 0.131489 | 0.309752 | 0.330975 | 0.190613 | 11.9 | 11.2 | 11.7 | 11.7 | 11.4 | 11.4 | 12 | 11.7 | 11.7 |
| 0.131373 | -0.18382 | 0.075597 | -0.22339 | 17.9 | 17.5 | 17.6 | 17.6 | 17.8 | 17.6 | 17.6 | 17.4 | 17.4 |
| 0.132299 | 0.258322 | 0.193113 | 0.218928 | 10.8 | 11.2 | 10.9 | 11.1 | 11.1 | 10.7 | 11.2 | 11.1 | 11.3 |
| 0.132601 | -0.27231 | 0.329256 | -0.16877 | 12.1 | 12.3 | 12.2 | 12.4 | 12 | 12.4 | 11.9 | 11.9 | 12.3 |
| 0.132858 | 0.235688 | 0.587817 | 0.079409 | 11.4 | 11.6 | 11.6 | 11 | 11.5 | 11.5 | 11.5 | 11.7 | 11.7 |
| 0.133633 | -0.16612 | 0.035002 | -0.25275 | 13.5 | 13.4 | 13.6 | 13.5 | 13.5 | 13.3 | 13.2 | 13.3 | 13.3 |
| 0.133697 | 0.207269 | 0.322927 | 0.130691 | 12.6 | 12.9 | 12.7 | 12.7 | 12.5 | 12.8 | 12.9 | 13 | 12.8 |
| 0.133797 | -0.38535 | 0.289693 | -0.26164 | 12.6 | 11.9 | 12 | 12.3 | 12.7 | 11.9 | 11.8 | 11.9 | 12 |
| 0.13313 | 0.491174 | 0.275022 | 0.343967 | 11 | 11.5 | 11.7 | 11.8 | 10.6 | 11.3 | 11.6 | 11.8 | 11.8 |
| 0.134007 | -0.11882 | 0.263408 | -0.08565 | 14.9 | 14.9 | 14.9 | 15 | 14.9 | 14.9 | 14.8 | 14.8 | 14.8 |
| 0.134466 | 0.235451 | 0.795961 | 0.037738 | 11.9 | 12.3 | 12.3 | 11.8 | 12.2 | 12 | 12.2 | 12.1 | 12.4 |
| 0.134525 | -0.27812 | 0.640426 | -0.08095 | 10.8 | 10.9 | 10.7 | 11.1 | 11.3 | 10.6 | 10.8 | 10.7 | 10.7 |
| 0.134803 | -0.84073 | 0.425761 | -0.42355 | 12.1 | 10.1 | 10.6 | 10.9 | 12 | 11.2 | 10.7 | 10.5 | 10.4 |
| 0.13545 | -0.19964 | 0.045837 | -0.28443 | 16.4 | 16.1 | 16 | 16 | 16.2 | 16.1 | 15.9 | 15.9 | 15.9 |
| 0.135159 | -0.13047 | 0.316785 | -0.08375 | 13.1 | 13.1 | 13 | 13.2 | 13.1 | 13 | 13 | 13 | 13 |
| 0.135449 | -0.10889 | 0.128497 | -0.11118 | 14.9 | 14.9 | 14.9 | 14.8 | 15 | 14.9 | 14.8 | 14.8 | 14.8 |
| 0.137283 | -0.15693 | 0.027307 | -0.25668 | 13.2 | 13 | 13.2 | 13 | 13.2 | 12.9 | 12.8 | 12.9 | 12.9 |
| 0.137119 | 0.17826 | 0.474532 | 0.080848 | 14.1 | 14.4 | 14.4 | 14.3 | 14.1 | 14.3 | 14.3 | 14.4 | 14.4 |
| 0.13678 | 0.170008 | 0.376641 | 0.096113 | 12.3 | 12.4 | 12.3 | 12.1 | 12.4 | 12.2 | 12.5 | 12.5 | 12.3 |
| 0.136674 | 0.709865 | 0.307517 | 0.467209 | 12.2 | 13.5 | 13.4 | 13.2 | 12.2 | 13.1 | 13 | 13.7 | 13.9 |
| 0.137339 | -0.10593 | 0.104114 | -0.11766 | 13.8 | 13.7 | 13.7 | 13.7 | 13.7 | 13.7 | 13.6 | 13.6 | 13.6 |
| 0.137943 | -0.34015 | 0.274597 | -0.24186 | 13.7 | 13 | 13 | 13.1 | 13.6 | 13.3 | 13.1 | 12.9 | 12.9 |
| 0.138094 | -0.18503 | 0.094001 | -0.21349 | 14.5 | 14.5 | 14.4 | 14.3 | 14.7 | 14.4 | 14.3 | 14.4 | 14.1 |
| 0.138419 | 0.234765 | 0.284145 | 0.163607 | 11.7 | 11.6 | 11.4 | 11.2 | 11.6 | 11.7 | 11.7 | 11.8 | 11.7 |
| 0.138984 | 0.153964 | 0.934314 | 0.007955 | 13.9 | 13.8 | 13.7 | 13.7 | 13.8 | 13.6 | 13.9 | 13.9 | 13.7 |
| 0.139155 | -0.29428 | 0.477141 | -0.13345 | 12 | 11.4 | 11.2 | 11.7 | 11.7 | 11.7 | 11.4 | 11.4 | 11.3 |

| | | | | | | | | | | | | |
|---|---|---|---|---|---|---|---|---|---|---|---|---|
| 0.140279 | -0.12371 | 0.037825 | -0.18815 | 14.9 | 14.7 | 14.9 | 14.8 | 14.8 | 14.8 | 14.7 | 14.7 | 14.7 |
| 0.140518 | 0.307783 | 0.145926 | 0.30303 | 11.3 | 10.9 | 11.4 | 11.1 | 11.3 | 11.2 | 11.4 | 11.9 | 11.3 |
| 0.141127 | 0.455813 | 0.075648 | -0.57 | 10.2 | 9.49 | 10.2 | 8.45 | 9.49 | 8.81 | 9.36 | 9.41 | 9.36 |
| 0.141238 | 0.366024 | 0.877772 | 0.035541 | 10.8 | 10.2 | 11.1 | 10.5 | 10.5 | 10.1 | 10.7 | 10.7 | 10.8 |
| 0.141883 | -0.13735 | 0.150473 | -0.13403 | 12.5 | 12.3 | 12.4 | 12.3 | 12.5 | 12.3 | 12.3 | 12.3 | 12.2 |
| 0.142242 | -0.28251 | 0.571735 | -0.10218 | 11.9 | 11.3 | 11.6 | 11.6 | 11.8 | 12 | 11.7 | 11.4 | 11.4 |
| 0.142775 | -0.15799 | 0.067015 | -0.20613 | 18.5 | 18.7 | 18.7 | 18.7 | 18.4 | 18.6 | 18.3 | 18.4 | 18.5 |
| 0.143069 | 0.225055 | 0.202469 | 0.192333 | 12.7 | 12.9 | 12.9 | 12.7 | 12.7 | 13.1 | 12.9 | 13.1 | 13.1 |
| 0.14335 | 0.128881 | 0.625763 | 0.040217 | 13.6 | 13.6 | 13.6 | 13.5 | 13.6 | 13.5 | 13.7 | 13.6 | 13.5 |
| 0.1437 | 0.428426 | 0.847565 | 0.05239 | 12.4 | 13 | 13 | 12.8 | 11.9 | 12.7 | 12.7 | 12.9 | 13 |
| 0.144084 | 0.108097 | 0.643521 | 0.03205 | 14.9 | 14.8 | 14.8 | 14.7 | 14.7 | 14.8 | 14.8 | 14.9 | 14.8 |
| 0.144011 | -0.15089 | 0.028556 | -0.24895 | 13 | 13.1 | 13.1 | 12.8 | 13.1 | 13 | 12.8 | 12.9 | 12.8 |
| 0.145494 | -0.23105 | 0.681781 | -0.06085 | 11.6 | 11.3 | 11.3 | 11.4 | 11.6 | 11.7 | 11.6 | 11.4 | 11.1 |
| 0.145057 | -0.35377 | 0.619537 | -0.11297 | 10.5 | 10.2 | 9.85 | 9.98 | 10.6 | 10.6 | 10.1 | 10.2 | 9.86 |
| 0.14526 | -0.38464 | 0.228719 | -0.31049 | 12 | 11.6 | 11.6 | 11.9 | 12.2 | 11.2 | 11.5 | 11.3 | 11.3 |
| 0.146331 | -0.11752 | 0.022655 | -0.20624 | 14 | 14 | 14.1 | 14 | 13.9 | 14 | 13.9 | 13.8 | 13.8 |
| 0.147168 | -0.28166 | 0.449478 | -0.13936 | 11.9 | 11.2 | 11.8 | 11.8 | 11.7 | 11.9 | 11.6 | 11.4 | 11.5 |
| 0.146691 | 0.131668 | 0.304695 | 0.089768 | 15.3 | 15.4 | 15.4 | 15.4 | 15.2 | 15.4 | 15.4 | 15.4 | 15.4 |
| 0.147091 | -0.20877 | 0.301283 | -0.14358 | 12.7 | 12.3 | 12.4 | 12.4 | 12.6 | 12.5 | 12.3 | 12.4 | 12.1 |
| 0.147637 | -0.18082 | 0.023227 | -0.31665 | 12.5 | 12.6 | 12.2 | 12.3 | 12.3 | 12.4 | 12.2 | 12 | 12.2 |
| 0.147855 | 0.343927 | 0.100625 | 0.398591 | 9.91 | 10.8 | 10.4 | 10.6 | 10.2 | 10.5 | 10.7 | 10.8 | 10.8 |
| 0.149251 | -0.19614 | 0.142463 | -0.19997 | 12.2 | 11.9 | 11.9 | 11.9 | 12.2 | 11.9 | 11.7 | 11.8 | 11.9 |
| 0.150406 | -0.38993 | 0.438297 | 0.199737 | 9.39 | 9.1 | 8.96 | 9.43 | 9.64 | 10.1 | 9.79 | 9.17 | 9.07 |
| 0.150892 | -0.156 | 0.093258 | -0.18723 | 13 | 12.8 | 12.8 | 12.7 | 13 | 12.9 | 12.7 | 12.6 | 12.7 |
| 0.15093 | -0.33445 | 0.147866 | -0.33735 | 14.5 | 13.8 | 13.8 | 13.9 | 14.3 | 13.8 | 13.8 | 13.6 | 13.7 |
| 0.154671 | 0.247273 | 0.274083 | -0.18461 | 10.7 | 10.8 | 11 | 10.3 | 10.4 | 10.5 | 10.3 | 10.9 | 10.7 |
| 0.154046 | 2.935131 | 0.92537 | -0.17992 | 8.23 | 8.52 | 9.33 | 8.72 | 0.47 | 7.53 | 8.5 | 8.28 | 8.75 |
| 0.154524 | 0.616213 | 0.968039 | -0.01617 | 8.41 | 8.92 | 9.44 | 7.83 | 8.01 | 9.03 | 8.85 | 8.5 | 9.37 |
| 0.153623 | -0.26409 | 0.026139 | -0.45738 | 10.7 | 10.8 | 10.9 | 10.9 | 10.3 | 10.6 | 10.5 | 10.4 | 10.2 |
| 0.154781 | 0.283227 | 0.578368 | 0.104268 | 13.3 | 13.9 | 13.8 | 13.6 | 13.3 | 13.7 | 13.7 | 13.8 | 13.8 |
| 0.154348 | -0.27174 | 0.045436 | -0.40966 | 11.4 | 11.3 | 11.2 | 11.2 | 11.1 | 11.3 | 11.3 | 10.7 | 10.7 |
| 0.152595 | -0.46327 | 0.413521 | -0.25248 | 12.2 | 11.6 | 11.2 | 11.5 | 12.1 | 12.1 | 11.8 | 11.3 | 11.2 |
| 0.154191 | -0.11128 | 0.391347 | -0.064 | 14.3 | 14.3 | 14.3 | 14.4 | 14.3 | 14.3 | 14.2 | 14.3 | 14.2 |
| 0.1521 | 0.364152 | 0.317345 | 0.245123 | 15.1 | 15.8 | 15.8 | 15.6 | 15 | 15.7 | 15.8 | 15.8 | 15.8 |
| 0.152591 | -0.1589 | 0.294976 | -0.11255 | 13.4 | 13.3 | 13.1 | 13.5 | 13.3 | 13.3 | 13.2 | 13.2 | 13 |
| 0.153305 | -0.26104 | 0.165265 | -0.25265 | 11.6 | 11.4 | 11.6 | 11.8 | 11.4 | 11.3 | 11.3 | 11 | 11.4 |
| 0.153147 | -0.15987 | 0.149806 | -0.16137 | 15.6 | 15.3 | 15.4 | 15.4 | 15.5 | 15.3 | 15.4 | 15.2 | 15.3 |
| 0.15573 | -0.17059 | 0.018284 | -0.32273 | 13.1 | 12.9 | 12.9 | 12.7 | 12.7 | 13 | 12.8 | 12.6 | 12.6 |
| 0.15567 | -3.40619 | 0.260431 | -2.63025 | 9.36 | 8.14 | 7.52 | 7.5 | 10.2 | 9.65 | -0 | 8.53 | 8.63 |
| 0.156465 | -0.15271 | 0.080766 | -0.19527 | 13.2 | 13.2 | 13.2 | 13.2 | 13.1 | 13.1 | 12.8 | 13.1 | 13.1 |
| 0.156247 | 0.521875 | 0.251742 | 0.411748 | 9.27 | 10.7 | 10.3 | 9.95 | 9.98 | 10 | 10.8 | 10.4 | 10.3 |
| 0.157258 | 0.139781 | 0.870878 | 0.015009 | 13.7 | 13.7 | 13.7 | 13.6 | 13.4 | 13.6 | 13.6 | 13.7 | 13.8 |
| 0.158429 | -0.43394 | 0.74514 | -0.09373 | 10.9 | 10.6 | 10.1 | 10.5 | 11.1 | 11 | 10.9 | 10.2 | 10.2 |
| 0.158602 | -0.20905 | 0.419581 | -0.11428 | 13.2 | 13.6 | 13.4 | 13.6 | 13.3 | 13.6 | 13.2 | 13.3 | 13.4 |
| 0.158483 | -0.2913 | 0.086714 | -0.36592 | 17.4 | 16.8 | 16.9 | 16.8 | 17.3 | 16.8 | 16.7 | 16.6 | 16.6 |
| 0.158004 | -0.13036 | 0.097575 | -0.1571 | 13.8 | 13.6 | 13.5 | 13.5 | 13.7 | 13.6 | 13.5 | 13.5 | 13.4 |
| 0.157638 | -0.12098 | 0.121981 | -0.13426 | 15 | 15 | 14.9 | 14.9 | 15 | 15 | 14.9 | 14.9 | 14.8 |
| 0.159122 | 0.268723 | 0.418663 | -0.14741 | 9.76 | 9.92 | 9.62 | 9.39 | 9.57 | 9.1 | 9.69 | 9.85 | 9.31 |
| 0.159595 | 0.311409 | 0.174614 | 0.299159 | 11.1 | 11.7 | 11.9 | 11.4 | 11.4 | 11.8 | 11.8 | 11.8 | 11.9 |
| 0.16019 | -0.44183 | 0.147793 | -0.4573 | 10.7 | 9.55 | 10.2 | 9.95 | 10.3 | 10.2 | 10 | 9.55 | 9.47 |

| | | | | | | | | | | | | |
|---|---|---|---|---|---|---|---|---|---|---|---|---|
| 0.161006 | 3.064457 | 0.911311 | 0.227701 | 8.49 | 9.44 | 8.23 | 8.97 | 0.44 | 8.25 | 9.39 | 8.53 | 8.92 |
| 0.161148 | -0.19275 | 0.686739 | -0.05214 | 10.7 | 11.1 | 11 | 11 | 11 | 11.2 | 10.9 | 10.8 | 11 |
| 0.161598 | 0.331025 | 0.972834 | -0.00753 | 11.5 | 12.3 | 12.2 | 11.7 | 11.4 | 11.9 | 11.9 | 12.2 | 12 |
| 0.162017 | -0.33229 | 0.127831 | -0.36651 | 12.4 | 12.5 | 12.3 | 12.5 | 12 | 12.5 | 11.7 | 11.9 | 12.4 |
| 0.1674 | -0.1868 | 0.258133 | -0.1497 | 12.5 | 12.3 | 12.2 | 12.5 | 12.5 | 12.2 | 12.3 | 12.3 | 12 |
| 0.167542 | 0.380033 | 0.215578 | 0.336603 | 8.78 | 9.35 | 8.57 | 9.2 | 8.95 | 8.42 | 9.42 | 9.08 | 9.2 |
| 0.168477 | -0.87532 | 0.539157 | -0.37039 | 8.89 | 9.51 | 8.82 | 8.83 | 9.91 | 10 | 9.79 | 7.47 | 8.85 |
| 0.168626 | 0.794906 | 0.349529 | 0.521412 | 7.28 | 8.63 | 8.68 | 7.92 | 6.99 | 8.85 | 8.68 | 9.04 | 8.44 |
| 0.16976 | -0.14285 | 0.333096 | -0.09745 | 14.2 | 14.1 | 13.9 | 14.1 | 14.2 | 14.1 | 14 | 14 | 14 |
| 0.170686 | -0.14885 | 0.197402 | -0.139 | 13.7 | 13.7 | 13.9 | 13.7 | 13.9 | 13.7 | 13.5 | 13.6 | 13.8 |
| 0.171998 | 0.388841 | 0.993655 | -0.00212 | 9.22 | 9.69 | 9.74 | 8.66 | 9.41 | 9.4 | 9.22 | 9.55 | 9.87 |
| 0.172507 | 0.192402 | 0.826382 | 0.029067 | 11.4 | 11.7 | 11.9 | 11.5 | 11.6 | 11.4 | 11.6 | 11.7 | 11.8 |
| 0.172299 | 0.216541 | 0.133211 | 0.241535 | 11.8 | 12.3 | 12.3 | 12.3 | 12.1 | 12.2 | 12.4 | 12.4 | 12.4 |
| 0.172913 | -0.4474 | 0.386018 | -0.27388 | 13.9 | 12.8 | 13.1 | 13.1 | 13.9 | 13.3 | 13.2 | 13 | 12.8 |
| 0.17565 | -0.11868 | 0.763861 | 0.024794 | 13.9 | 14.1 | 14 | 14.1 | 14.3 | 14.2 | 14.1 | 14.1 | 14 |
| 0.176512 | -0.33243 | 0.092664 | -0.42848 | 9.77 | 9.44 | 9.61 | 8.99 | 9.73 | 9.82 | 9.06 | 9.06 | 9.42 |
| 0.177042 | 0.232062 | 0.785559 | 0.044046 | 11.5 | 11.3 | 11.5 | 11.4 | 11.4 | 11 | 11.8 | 11.5 | 11.3 |
| 0.177311 | -0.25628 | 0.545681 | -0.10917 | 11.2 | 11.4 | 11.1 | 11.1 | 11.2 | 11.8 | 11.1 | 11.2 | 11.1 |
| 0.177445 | 0.294277 | 0.263049 | 0.239447 | 12.4 | 11.6 | 12.1 | 11.8 | 12.2 | 11.9 | 12.4 | 12.2 | 12.1 |
| 0.177659 | -0.21745 | 0.240755 | -0.18635 | 11.1 | 11 | 11 | 10.9 | 11.1 | 11.1 | 11.2 | 10.7 | 10.7 |
| 0.178023 | 0.257495 | 0.547617 | 0.10935 | 12.3 | 12.8 | 12.7 | 12.6 | 12.2 | 12.6 | 12.6 | 12.6 | 12.9 |
| 0.179543 | -0.26011 | 0.594218 | -0.09799 | 11 | 11.4 | 11.3 | 11.2 | 11.4 | 11.5 | 11 | 10.8 | 11.4 |
| 0.181771 | 0.271354 | 0.323495 | 0.195087 | 12 | 12.7 | 12.5 | 12.5 | 12.1 | 12.4 | 12.7 | 12.5 | 12.7 |
| 0.181769 | -0.356 | 0.304144 | -0.26713 | 10.7 | 11 | 10.8 | 10.6 | 10.8 | 11.3 | 10.3 | 10.3 | 11 |
| 0.181577 | -0.44025 | 0.108811 | -0.54332 | 10.8 | 10.5 | 10 | 10.1 | 10.6 | 10.3 | 10.2 | 10.2 | 9.23 |
| 0.182717 | -0.25155 | 0.647402 | 0.08182 | 10.7 | 10.2 | 10.2 | 10.6 | 11 | 10.5 | 10.5 | 10.4 | 10.5 |
| 0.183302 | -0.12845 | 0.108615 | -0.15933 | 13.5 | 13.2 | 13.3 | 13.3 | 13.3 | 13.3 | 13.2 | 13.1 | 13.2 |
| 0.18611 | -0.62668 | 0.658627 | 0.19852 | 10.7 | 9.18 | 10.2 | 10.9 | 10.7 | 10.9 | 10.8 | 10.4 | 9.44 |
| 0.186161 | 0.186854 | 0.420844 | 0.109531 | 13.5 | 13.9 | 13.8 | 13.8 | 13.4 | 13.8 | 13.9 | 13.9 | 13.9 |
| 0.187306 | -0.31242 | 0.040285 | -0.53079 | 11.5 | 11.8 | 11.5 | 11.4 | 11.2 | 11.6 | 11.3 | 10.6 | 11.3 |
| 0.187174 | 0.1506 | 0.656673 | 0.048133 | 10.2 | 10.5 | 10.2 | 10.3 | 10.2 | 10.2 | 10.4 | 10.3 | 10.3 |
| 0.187482 | -0.10332 | 0.495007 | -0.05119 | 12.8 | 12.8 | 12.7 | 12.8 | 12.8 | 12.9 | 12.8 | 12.7 | 12.7 |
| 0.188234 | -0.10875 | 0.014848 | -0.23464 | 12.8 | 12.7 | 12.8 | 12.7 | 12.7 | 12.5 | 12.5 | 12.5 | 12.6 |
| 0.188414 | -0.09268 | 0.114622 | -0.11419 | 16.4 | 16.3 | 16.3 | 16.3 | 16.3 | 16.3 | 16.3 | 16.2 | 16.2 |
| 0.187996 | -0.13968 | 0.182088 | -0.1418 | 15.5 | 15.7 | 15.6 | 15.7 | 15.5 | 15.6 | 15.5 | 15.4 | 15.5 |
| 0.188921 | -0.1929 | 0.212945 | -0.18176 | 15.3 | 14.9 | 14.8 | 14.9 | 15.1 | 14.9 | 14.8 | 14.8 | 14.8 |
| 0.190488 | -0.33753 | 0.682278 | -0.10008 | 11.3 | 10.6 | 10.5 | 10.7 | 11 | 11.3 | 10.7 | 10.5 | 10.9 |
| 0.190244 | 0.850821 | 0.381749 | 0.549778 | 9.78 | 11.1 | 11.2 | 11.3 | 9.18 | 10.7 | 10.8 | 11 | 11.8 |
| 0.190394 | -0.14288 | 0.219001 | -0.13318 | 16.9 | 16.6 | 16.6 | 16.6 | 16.8 | 16.6 | 16.6 | 16.5 | 16.5 |
| 0.19094 | 0.248683 | 0.469229 | -0.1321 | 12 | 11.9 | 11.9 | 11.6 | 11.1 | 11.9 | 11.7 | 11.7 | 11.9 |
| 0.192716 | 0.143922 | 0.940092 | -0.00784 | 13.1 | 13 | 13.1 | 13 | 13.1 | 12.7 | 13 | 13.1 | 13.2 |
| 0.192787 | -0.1057 | 0.0391 | -0.18352 | 12.1 | 12.2 | 12.2 | 12.1 | 12.2 | 12 | 12 | 12 | 12.1 |
| 0.192262 | -0.13887 | 0.508438 | -0.06743 | 15 | 14.7 | 14.8 | 14.9 | 15 | 14.8 | 14.9 | 14.7 | 14.7 |
| 0.193144 | 0.387792 | 0.380167 | 0.253302 | 10.7 | 10.3 | 10.5 | 9.81 | 10.4 | 11 | 11 | 10.7 | 10.6 |
| 0.193807 | -0.13228 | 0.091284 | 0.179119 | 11.5 | 11.6 | 11.6 | 12 | 11.7 | 11.9 | 11.6 | 11.8 | 11.7 |
| 0.193987 | -0.21751 | 0.08057 | -0.30711 | 14.1 | 14 | 13.9 | 13.8 | 14.3 | 13.6 | 13.7 | 13.8 | 13.6 |
| 0.195569 | -0.17819 | 0.245886 | -0.15794 | 15.4 | 15.8 | 15.7 | 15.7 | 15.5 | 15.7 | 15.4 | 15.4 | 15.7 |
| 0.19669 | -0.28065 | 0.062142 | -0.43245 | 9.64 | 9.3 | 9.69 | 9.32 | 9.12 | 9.73 | 9.27 | 8.81 | 9.25 |
| 0.196513 | -0.17518 | 0.079047 | -0.25045 | 12.3 | 12 | 12 | 12 | 12.1 | 11.9 | 11.8 | 11.7 | 12 |
| 0.197065 | -0.23086 | 0.024445 | -0.45523 | 11.8 | 11.8 | 11.8 | 11.8 | 11.6 | 11.3 | 11.1 | 11.4 | 11.6 |

| | | | | | | | | | | | | |
|---|---|---|---|---|---|---|---|---|---|---|---|---|
| 0.198202 | -0.45336 | 0.543023 | -0.20504 | 9.81 | 9.82 | 8.61 | 9.72 | 9.4 | 9.86 | 9.29 | 9.39 | 8.95 |
| 0.198446 | -0.20374 | 0.338671 | -0.14776 | 10.9 | 10.8 | 11.2 | 11.2 | 11.1 | 10.7 | 10.8 | 10.8 | 10.8 |
| 0.200389 | 0.174009 | 0.622671 | 0.063736 | 13.6 | 13.3 | 13.2 | 13.2 | 13.5 | 13.1 | 13.5 | 13.5 | 13.4 |
| 0.200287 | 0.150606 | 0.598986 | 0.059006 | 13.7 | 14.1 | 13.9 | 13.8 | 13.8 | 13.8 | 14 | 14 | 13.9 |
| 0.19959 | 0.487805 | 0.504728 | 0.243407 | 10.2 | 11.1 | 11.3 | 10.9 | 9.92 | 11 | 11 | 11 | 11.2 |
| 0.199818 | 0.261977 | 0.45692 | 0.146406 | 15.9 | 16.5 | 16.4 | 16.3 | 15.9 | 16.2 | 16.2 | 16.5 | 16.6 |
| 0.200895 | -0.13829 | 0.505735 | -0.06906 | 13.5 | 13.4 | 13.3 | 13.4 | 13.7 | 13.4 | 13.4 | 13.3 | 13.4 |
| 0.201056 | -0.09065 | 0.143047 | -0.10571 | 14.3 | 14.3 | 14.2 | 14.2 | 14.3 | 14.3 | 14.2 | 14.1 | 14.2 |
| 0.202254 | -0.21377 | 0.186714 | -0.22225 | 13.9 | 14.2 | 14.2 | 14.3 | 13.8 | 14.2 | 13.9 | 13.8 | 14 |
| 0.202654 | -0.12973 | 0.131673 | -0.15704 | 12.3 | 12.3 | 12 | 12.2 | 12.2 | 12.1 | 12 | 12.1 | 12.1 |
| 0.203154 | -0.22066 | 0.211885 | -0.216 | 10.8 | 11.2 | 10.9 | 11 | 11 | 10.9 | 10.4 | 10.8 | 11 |
| 0.205336 | -0.14234 | 0.030471 | -0.27165 | 12.4 | 12.4 | 12.3 | 12.2 | 12.4 | 12.1 | 12.1 | 12.1 | 12 |
| 0.205472 | -0.43922 | 0.244882 | -0.39991 | 11 | 10.9 | 10.7 | 10.7 | 10.4 | 11.6 | 10.7 | 10.8 | 9.94 |
| 0.205931 | -0.21836 | 0.107701 | -0.28753 | 10.6 | 10.1 | 10.6 | 10.2 | 10.6 | 10.4 | 10.1 | 10.2 | 10.2 |
| 0.207728 | -0.24816 | 0.809536 | 0.045052 | 12.6 | 11.9 | 12.5 | 12.5 | 12.8 | 12.5 | 12.3 | 12.6 | 12.2 |
| 0.208913 | 0.336104 | 0.271326 | 0.290491 | 15.8 | 16.4 | 16.5 | 16.4 | 15.7 | 16.4 | 16.4 | 16.6 | 16.6 |
| 0.210136 | -0.19228 | 0.253748 | -0.17349 | 14.6 | 14.3 | 14.2 | 14.3 | 14.6 | 14.2 | 14.2 | 14.2 | 14.1 |
| 0.210916 | -0.32826 | 0.646215 | 0.114973 | 11.1 | 11.1 | 11.3 | 11.8 | 11 | 12.1 | 11.2 | 11.4 | 11.4 |
| 0.210719 | -0.18055 | 0.039427 | -0.32696 | 12.4 | 12.3 | 12 | 11.9 | 12.2 | 12.2 | 12 | 11.8 | 11.9 |
| 0.211938 | -0.17621 | 0.04859 | -0.30253 | 12.2 | 12.1 | 12.4 | 12.3 | 12.2 | 11.9 | 12 | 12.1 | 11.8 |
| 0.212713 | -0.18426 | 0.336463 | -0.13907 | 12.1 | 11.9 | 11.8 | 12 | 12.2 | 11.8 | 11.6 | 11.9 | 11.9 |
| 0.213315 | -0.53547 | 0.456091 | -0.30989 | 9.56 | 9.16 | 8.5 | 8.65 | 9.38 | 9.87 | 9.15 | 8.23 | 8.92 |
| 0.213284 | 0.309186 | 0.443308 | 0.184243 | 11.7 | 12.1 | 12.2 | 12.2 | 11.4 | 12.1 | 12.4 | 12 | 12.2 |
| 0.2149 | -0.10707 | 0.022394 | -0.22514 | 15.7 | 15.5 | 15.7 | 15.5 | 15.6 | 15.6 | 15.5 | 15.4 | 15.4 |
| 0.215343 | -0.13518 | 0.470645 | -0.07601 | 12.8 | 12.7 | 12.8 | 12.9 | 12.8 | 12.7 | 12.8 | 12.8 | 12.5 |
| 0.214636 | 0.146175 | 0.334012 | 0.111436 | 12.8 | 12.8 | 13.1 | 12.8 | 13 | 12.8 | 13.1 | 12.9 | 13 |
| 0.215126 | -0.13333 | 0.139071 | -0.16279 | 14.1 | 14.3 | 14.3 | 14.3 | 14.1 | 14.3 | 13.9 | 14.1 | 14.2 |
| 0.216214 | -0.15167 | 0.755368 | 0.036382 | 11.6 | 11.5 | 11.4 | 11.6 | 11.6 | 11.9 | 11.6 | 11.5 | 11.6 |
| 0.216321 | -0.14178 | 0.087065 | -0.20615 | 13.5 | 13.3 | 13.5 | 13.4 | 13.2 | 13.6 | 13.2 | 13.2 | 13.3 |
| 0.216247 | 0.127032 | 0.342539 | 0.095412 | 14.2 | 14.4 | 14.4 | 14.5 | 14.2 | 14.3 | 14.5 | 14.5 | 14.3 |
| 0.218392 | -0.59148 | 0.79555 | 0.118493 | 8.63 | 7.97 | 8.71 | 9.3 | 9.45 | 8.7 | 8.92 | 9.22 | 7.54 |
| 0.218012 | -0.19144 | 0.058598 | -0.31621 | 11.5 | 11.3 | 11.2 | 10.9 | 11.5 | 11.3 | 11.1 | 11.1 | 11 |
| 0.218151 | -0.12041 | 0.507365 | -0.06247 | 12.3 | 12.4 | 12.5 | 12.4 | 12.6 | 12.4 | 12.3 | 12.4 | 12.3 |
| 0.217834 | -0.16086 | 0.465955 | -0.09199 | 12.6 | 12.9 | 12.6 | 12.9 | 12.6 | 12.9 | 12.5 | 12.6 | 12.6 |
| 0.219273 | -0.5738 | 0.229584 | -0.55989 | 9.92 | 9.96 | 10.2 | 10.2 | 9.47 | 10.4 | 8.45 | 10.1 | 9.83 |
| 0.219639 | 0.166875 | 0.461467 | 0.096848 | 13.4 | 13.1 | 12.9 | 13 | 13.2 | 13 | 13.3 | 13.1 | 13.2 |
| 0.22028 | 0.383468 | 0.568221 | 0.171455 | 10.4 | 10.2 | 9.49 | 9.76 | 9.91 | 9.82 | 10.6 | 10.3 | 9.69 |
| 0.222194 | -0.24102 | 0.803492 | -0.04678 | 11 | 11.1 | 10.5 | 11.1 | 11.1 | 11 | 11.1 | 10.7 | 10.7 |
| 0.222458 | -0.15775 | 0.102138 | -0.22034 | 14.3 | 14 | 13.9 | 14 | 14.1 | 13.9 | 13.9 | 13.8 | 13.8 |
| 0.222897 | 0.18464 | 0.706395 | 0.054497 | 12.8 | 12.5 | 12.4 | 12.5 | 12.4 | 12.4 | 12.4 | 12.9 | 12.6 |
| 0.22503 | 3.02676 | 0.93178 | 0.203124 | 8.63 | 10 | 10.2 | 9.63 | 0.48 | 10.3 | 9.68 | 9.68 | 10.1 |
| 0.225505 | 0.385016 | 0.302579 | 0.322917 | 9.04 | 9.71 | 9.8 | 8.9 | 9.57 | 9.9 | 10.1 | 9.65 | 9.79 |
| 0.225885 | 0.078691 | 0.008 | -0.21139 | 16.2 | 16.2 | 16.1 | 15.9 | 15.9 | 15.8 | 16 | 15.9 | 15.9 |
| 0.227235 | 0.103248 | 0.975311 | 0.002518 | 15.4 | 15.4 | 15.3 | 15.2 | 15.3 | 15.2 | 15.4 | 15.4 | 15.3 |
| 0.227892 | -0.22755 | 0.764049 | -0.05406 | 10.8 | 11 | 11 | 11.3 | 11.4 | 10.7 | 10.9 | 10.7 | 11 |
| 0.227698 | -0.16064 | 0.489311 | -0.08903 | 13.1 | 12.8 | 12.8 | 12.9 | 13.2 | 12.9 | 12.9 | 12.7 | 12.9 |
| 0.228883 | 0.257775 | 0.16076 | -0.30596 | 11.3 | 11.4 | 11 | 10.4 | 10.9 | 10.7 | 10.7 | 10.8 | 11.3 |
| 0.228992 | 0.21048 | 0.942877 | 0.011934 | 11.6 | 11 | 11.3 | 10.9 | 11.3 | 11.1 | 11.4 | 11.2 | 11.3 |
| 0.228411 | -0.17725 | 0.089512 | -0.26281 | 12.8 | 12.4 | 12.5 | 12.4 | 12.6 | 12.4 | 12.2 | 12.2 | 12.4 |
| 0.229287 | -0.3268 | 0.413204 | -0.21657 | 10.2 | 9.69 | 10.4 | 10.1 | 10.3 | 10.3 | 10.2 | 10.1 | 9.36 |

| | | | | | | | | | | | | |
|---|---|---|---|---|---|---|---|---|---|---|---|---|
| 0.229622 | -0.24725 | 0.903573 | 0.023745 | 8.86 | 9.1 | 8.95 | 8.95 | 9.36 | 9.41 | 9.36 | 8.71 | 8.91 |
| 0.230162 | -0.2136 | 0.436111 | -0.13471 | 12.5 | 12.3 | 11.9 | 12.1 | 12.4 | 12.3 | 12.2 | 12.1 | 11.9 |
| 0.230625 | -0.21106 | 0.482358 | -0.11974 | 14.2 | 13.7 | 13.7 | 13.9 | 14.2 | 13.7 | 13.8 | 13.7 | 13.6 |
| 0.231232 | 0.224037 | 0.894762 | 0.023584 | 11.8 | 11.7 | 11.5 | 11.2 | 11.9 | 11.4 | 11.7 | 11.6 | 11.8 |
| 0.231698 | -0.27966 | 0.441866 | -0.17471 | 11.4 | 10.6 | 11 | 10.9 | 11.4 | 11 | 10.9 | 10.8 | 10.8 |
| 0.231746 | 0.397969 | 0.176711 | 0.455809 | 10.3 | 10.2 | 10.3 | 10.4 | 9.67 | 10.9 | 10.3 | 10.8 | 11.1 |
| 0.232131 | 0.137413 | 0.621691 | -0.05449 | 12.3 | 12.1 | 12.2 | 11.8 | 12.1 | 12.1 | 12.2 | 12.2 | 12 |
| 0.23346 | 0.145771 | 0.792584 | 0.030718 | 13.5 | 13.8 | 13.8 | 13.7 | 13.5 | 13.6 | 13.6 | 13.8 | 13.8 |
| 0.234447 | 0.288548 | 0.273515 | 0.263774 | 10.2 | 10.8 | 11 | 10.7 | 10.7 | 10.6 | 10.7 | 11.2 | 11 |
| 0.235218 | -0.39658 | 0.414882 | -0.2655 | 8.64 | 9.4 | 9.41 | 9.04 | 9.3 | 9.5 | 9.15 | 8.26 | 9.24 |
| 0.234968 | -0.57682 | 0.268712 | -0.53371 | 11.7 | 11.5 | 12.1 | 11.2 | 11.2 | 13 | 11.3 | 11.1 | 11.2 |
| 0.235955 | 0.227625 | 0.290642 | 0.200944 | 11.3 | 11.6 | 11.2 | 11.3 | 11.1 | 11.6 | 11.7 | 11.7 | 11.3 |
| 0.237651 | -0.48763 | 0.263437 | -0.4595 | 8.73 | 8.9 | 8.67 | 8.1 | 9.02 | 9.28 | 8.73 | 8.65 | 7.55 |
| 0.239252 | -0.28684 | 0.690378 | -0.0931 | 10.2 | 10.7 | 10.9 | 10.3 | 11 | 11.1 | 10.5 | 10.4 | 10.6 |
| 0.239072 | 0.303393 | 0.42437 | 0.200636 | 11.7 | 11 | 10.8 | 11.1 | 10.8 | 11.2 | 11.4 | 11.4 | 11.3 |
| 0.240413 | 0.252216 | 0.087137 | -0.38813 | 9.95 | 10.3 | 10.1 | 9.93 | 9.2 | 9.28 | 9.71 | 9.56 | 9.89 |
| 0.240658 | 3.043793 | 0.966025 | -0.10542 | 9.85 | 9.75 | 9.53 | -0.1 | 9.76 | 10.1 | 9.42 | 9.71 | 9.69 |
| 0.240174 | 0.136136 | 0.161747 | 0.165481 | 14.1 | 14 | 13.7 | 13.9 | 14 | 14.1 | 14.1 | 14.1 | 14.1 |
| 0.241619 | 0.300717 | 0.165205 | 0.363364 | 10.6 | 10.9 | 11.3 | 11.2 | 10.6 | 11.1 | 11 | 11.4 | 11.5 |
| 0.245479 | -0.12209 | 0.074989 | 0.199604 | 11.2 | 11.3 | 11 | 11.4 | 11.6 | 11.5 | 11.4 | 11.3 | 11.5 |
| 0.242642 | -0.17035 | 0.397654 | 0.12059 | 14.7 | 14.9 | 15 | 15.3 | 15 | 15.2 | 14.9 | 15 | 15.2 |
| 0.245488 | -0.08365 | 0.046681 | -0.15721 | 14.3 | 14.3 | 14.4 | 14.2 | 14.2 | 14.3 | 14.1 | 14.2 | 14.2 |
| 0.243316 | 0.227231 | 0.744421 | 0.060823 | 13.1 | 12.8 | 12.8 | 12.7 | 13.1 | 12.4 | 13 | 13.1 | 12.9 |
| 0.244845 | -0.16055 | 0.704703 | -0.0502 | 11.8 | 12.3 | 12.1 | 12.2 | 12.1 | 12.2 | 11.9 | 12.1 | 12 |
| 0.243773 | -0.18091 | 0.533299 | -0.09352 | 12.9 | 12.4 | 12.4 | 12.7 | 12.7 | 12.6 | 12.4 | 12.6 | 12.4 |
| 0.245543 | 0.502132 | 0.529995 | 0.262813 | 13.4 | 14.4 | 14.3 | 14.3 | 12.9 | 14.1 | 14.2 | 14.3 | 14.3 |
| 0.244172 | -0.12133 | 0.11314 | -0.17186 | 14.4 | 14.5 | 14.6 | 14.5 | 14.5 | 14.4 | 14.5 | 14.3 | 14.3 |
| 0.242556 | 0.154778 | 0.471421 | 0.092647 | 14.5 | 14.7 | 14.7 | 14.7 | 14.3 | 14.7 | 14.7 | 14.8 | 14.7 |
| 0.243832 | -0.18772 | 0.12953 | -0.25234 | 10.9 | 11 | 10.7 | 10.5 | 11.1 | 10.8 | 10.6 | 10.7 | 10.5 |
| 0.243294 | -0.10257 | 0.134376 | -0.13571 | 15.6 | 15.4 | 15.4 | 15.5 | 15.5 | 15.4 | 15.3 | 15.3 | 15.4 |
| 0.245549 | -0.13516 | 0.248438 | -0.13426 | 13.7 | 13.5 | 13.4 | 13.6 | 13.6 | 13.4 | 13.4 | 13.4 | 13.5 |
| 0.246236 | -0.37196 | 0.790159 | -0.08172 | 9.67 | 8.99 | 9.11 | 9.74 | 9.45 | 9.45 | 9.58 | 8.51 | 9.43 |
| 0.246614 | 0.581855 | 0.562772 | 0.28077 | 12 | 13.2 | 13.2 | 13.1 | 11.5 | 12.9 | 13 | 13.1 | 13.2 |
| 0.24913 | -0.31188 | 0.054273 | -0.56668 | 9.58 | 9.98 | 9.14 | 9.69 | 9.32 | 8.93 | 8.82 | 9.15 | 9.04 |
| 0.251037 | -0.21102 | 0.696211 | -0.06897 | 13.1 | 12.6 | 12.5 | 12.8 | 12.8 | 13 | 12.9 | 12.5 | 12.5 |
| 0.249365 | -0.11523 | 0.075641 | -0.18956 | 14.9 | 14.7 | 14.8 | 14.6 | 14.9 | 14.7 | 14.6 | 14.6 | 14.6 |
| 0.24903 | -0.18327 | 0.573077 | -0.08654 | 12.1 | 12.4 | 12.4 | 12.6 | 12.2 | 12.4 | 12.3 | 12.1 | 12.3 |
| 0.249282 | 0.244485 | 0.525133 | 0.130644 | 11.3 | 11.3 | 11.3 | 11.5 | 10.7 | 11.4 | 11.4 | 11.5 | 11.4 |
| 0.25058 | -0.08632 | 0.108102 | -0.12617 | 15.5 | 15.5 | 15.5 | 15.4 | 15.6 | 15.5 | 15.3 | 15.4 | 15.5 |
| 0.250271 | -0.15453 | 0.512446 | -0.08539 | 12.3 | 12.6 | 12.4 | 12.2 | 12.6 | 12.6 | 12.3 | 12.3 | 12.4 |
| 0.249865 | 0.083575 | 0.124546 | 0.115675 | 15.4 | 15.4 | 15.3 | 15.4 | 15.5 | 15.4 | 15.5 | 15.5 | 15.5 |
| 0.250764 | -0.11035 | 0.141287 | -0.14557 | 14.8 | 14.6 | 14.7 | 14.8 | 14.7 | 14.6 | 14.6 | 14.6 | 14.6 |
| 0.251631 | -0.09955 | 0.327109 | -0.08406 | 13.7 | 13.8 | 13.7 | 13.6 | 13.8 | 13.7 | 13.7 | 13.6 | 13.6 |
| 0.251645 | -0.6696 | 0.314093 | -0.58193 | 8.8 | 8.59 | 9.4 | 9.41 | 7.75 | 9.89 | 8.63 | 7.76 | 8.64 |
| 0.25112 | -0.20144 | 0.210221 | -0.22186 | 12.9 | 12.7 | 12.3 | 12.5 | 12.7 | 12.6 | 12.4 | 12.2 | 12.7 |
| 0.250251 | 0.152628 | 0.297045 | 0.137314 | 15.2 | 15.5 | 15.5 | 15.5 | 15.2 | 15.4 | 15.5 | 15.6 | 15.6 |
| 0.252371 | -0.24605 | 0.009401 | -0.68095 | 11.8 | 11.8 | 11.6 | 11.3 | 11.5 | 11.1 | 10.8 | 11 | 11.5 |
| 0.253571 | -0.16493 | 0.291573 | -0.15136 | 12.8 | 13 | 12.8 | 13.1 | 12.6 | 12.9 | 12.6 | 12.6 | 12.8 |
| 0.256639 | -0.09035 | 0.11235 | -0.13204 | 14.6 | 14.6 | 14.5 | 14.5 | 14.6 | 14.5 | 14.4 | 14.4 | 14.4 |
| 0.257758 | -0.11731 | 0.357277 | -0.09401 | 14.8 | 14.6 | 14.7 | 14.6 | 14.9 | 14.7 | 14.6 | 14.6 | 14.6 |

| | | | | | | | | | | | | |
|---|---|---|---|---|---|---|---|---|---|---|---|---|
| 0.259251 | 0.475401 | 0.957329 | -0.02159 | 10.2 | 10.9 | 11 | 11 | 10 | 9.51 | 10.7 | 10.4 | 10.8 |
| 0.259413 | 0.167858 | 0.973732 | 0.004693 | 13.1 | 12.7 | 12.8 | 12.5 | 12.9 | 12.6 | 12.9 | 12.8 | 12.8 |
| 0.259342 | 0.124702 | 0.352347 | 0.10139 | 13.3 | 13.5 | 13.6 | 13.6 | 13.3 | 13.5 | 13.5 | 13.6 | 13.7 |
| 0.259902 | -0.14013 | 0.35017 | -0.11463 | 14.8 | 14.5 | 14.5 | 14.6 | 14.8 | 14.6 | 14.6 | 14.4 | 14.6 |
| 0.260879 | 0.27405 | 0.748258 | 0.075184 | 10.1 | 9.51 | 9.88 | 9.51 | 9.86 | 9.54 | 9.45 | 10.1 | 10.1 |
| 0.261962 | 0.433798 | 0.57148 | 0.21191 | 12.8 | 13.9 | 14 | 13.8 | 12.7 | 13.5 | 13.7 | 13.8 | 13.9 |
| 0.261907 | -0.25908 | 0.551996 | -0.13316 | 12 | 11.7 | 11.4 | 11.4 | 12.1 | 12 | 11.8 | 11.4 | 11.5 |
| 0.261602 | -0.44691 | 0.314365 | -0.39712 | 10.1 | 10.4 | 11 | 10.2 | 10.7 | 10.8 | 10.7 | 9.4 | 10.2 |
| 0.261774 | -0.35295 | 0.29366 | -0.3285 | 9.71 | 10.6 | 10.4 | 9.94 | 10.1 | 10.8 | 10.1 | 9.74 | 9.87 |
| 0.263434 | -0.25483 | 0.338739 | -0.21549 | 11.9 | 11.3 | 11.5 | 11.5 | 12 | 11.3 | 11.3 | 11.3 | 11.5 |
| 0.26357 | -0.64045 | 0.240926 | -0.67469 | 9.76 | 9.17 | 8.54 | 8.95 | 10.2 | 8.21 | 7.92 | 8.67 | 8.87 |
| 0.265514 | 0.280957 | 0.92378 | -0.02315 | 10.7 | 11.2 | 11.7 | 10.8 | 11.1 | 10.7 | 11.3 | 11.2 | 11 |
| 0.265993 | -0.08739 | 0.768487 | 0.022235 | 12.4 | 12.4 | 12.3 | 12.4 | 12.5 | 12.6 | 12.4 | 12.4 | 12.5 |
| 0.267004 | 0.106462 | 0.003269 | 0.371694 | 12.6 | 12.6 | 12.6 | 12.8 | 12.8 | 12.9 | 12.9 | 13.1 | 12.9 |
| 0.270365 | -0.11826 | 0.417492 | -0.08534 | 14.2 | 13.9 | 14 | 13.9 | 14.1 | 14.1 | 14 | 13.9 | 13.9 |
| 0.271287 | 0.149081 | 0.531717 | -0.0824 | 12.6 | 12.9 | 12.6 | 12.7 | 12.5 | 12.3 | 12.7 | 12.6 | 12.6 |
| 0.270977 | -0.15771 | 0.792072 | 0.036322 | 11.1 | 10.8 | 11.3 | 11.1 | 11.4 | 11.2 | 11.2 | 11 | 11.1 |
| 0.271853 | -0.30776 | 0.117108 | -0.4585 | 18.6 | 17.9 | 18 | 17.8 | 18.5 | 17.7 | 17.9 | 17.7 | 17.5 |
| 0.272885 | 0.119078 | 0.44895 | -0.08048 | 12.4 | 12.8 | 12.7 | 12.4 | 12.5 | 12.3 | 12.5 | 12.6 | 12.6 |
| 0.273013 | -0.07758 | 0.129395 | -0.11152 | 15.1 | 15 | 15 | 15 | 15.1 | 15 | 15 | 14.9 | 14.9 |
| 0.274198 | -0.33318 | 0.52395 | -0.189 | 14.1 | 13.2 | 13.4 | 13.3 | 14.3 | 13.6 | 13.5 | 13.3 | 13.3 |
| 0.275953 | 0.123827 | 0.568234 | -0.06298 | 13.3 | 13.2 | 13.1 | 12.9 | 13 | 13.1 | 13.2 | 13.2 | 13 |
| 0.276307 | -0.20457 | 0.181583 | -0.25631 | 12 | 11.5 | 11.9 | 11.8 | 11.9 | 11.6 | 11.4 | 11.8 | 11.4 |
| 0.27638 | 0.088685 | 0.39915 | 0.067586 | 17.7 | 17.6 | 17.6 | 17.6 | 17.7 | 17.5 | 17.6 | 17.7 | 17.7 |
| 0.27661 | 0.867784 | 0.254983 | 0.911483 | 7.61 | 10.3 | 10.7 | 9.9 | 8.91 | 9.88 | 10.1 | 10.5 | 10.8 |
| 0.277014 | -0.17741 | 0.500432 | -0.10726 | 13.7 | 13.5 | 13.3 | 13.4 | 13.9 | 13.4 | 13.4 | 13.4 | 13.4 |
| 0.27913 | -0.15555 | 0.762518 | -0.04186 | 14.8 | 14.4 | 14.3 | 14.5 | 14.7 | 14.6 | 14.6 | 14.4 | 14.4 |
| 0.280202 | 0.235384 | 0.11396 | 0.360987 | 10.5 | 10.4 | 10.7 | 10.8 | 10.2 | 11 | 10.8 | 10.8 | 11 |
| 0.280627 | 2.162297 | 0.877411 | -0.29735 | 7.95 | 8.71 | 8.57 | 8.57 | 8.47 | 0.81 | 7.4 | 8.26 | 8.67 |
| 0.281826 | -0.14476 | 0.005656 | -0.47276 | 11.1 | 10.9 | 10.8 | 10.4 | 10.6 | 10.8 | 10.4 | 10.4 | 10.5 |
| 0.281859 | 0.090819 | 0.1317 | 0.132262 | 11.8 | 11.9 | 11.9 | 11.8 | 12 | 11.9 | 12.1 | 12 | 11.9 |
| 0.282849 | 0.377916 | 0.898736 | 0.043078 | 9.76 | 10.7 | 10.4 | 10.4 | 9.32 | 10.2 | 10.5 | 10 | 10.4 |
| 0.283561 | -0.17412 | 0.924697 | 0.014761 | 11 | 10.8 | 10.5 | 11.1 | 11 | 10.8 | 10.8 | 10.6 | 10.9 |
| 0.283924 | -0.1199 | 0.071012 | -0.21763 | 14.2 | 14 | 14 | 14.1 | 13.9 | 13.8 | 13.9 | 13.8 | 13.8 |
| 0.283673 | 0.196499 | 0.42084 | 0.145019 | 12.5 | 13.2 | 12.9 | 12.8 | 12.7 | 12.8 | 12.8 | 13.1 | 13.1 |
| 0.284798 | -0.21158 | 0.135973 | -0.30624 | 10.1 | 10.3 | 10.6 | 10.2 | 10.2 | 10.3 | 9.71 | 10.4 | 10 |
| 0.285059 | 0.187139 | 0.275265 | 0.191303 | 12.1 | 12.4 | 12.3 | 12.4 | 11.9 | 12.5 | 12.4 | 12.6 | 12.4 |
| 0.28581 | -0.18419 | 0.244857 | 0.202172 | 11.6 | 11.1 | 11.6 | 12 | 11.7 | 11.8 | 11.7 | 11.8 | 11.5 |
| 0.286144 | -0.31694 | 0.801615 | -0.07197 | 11.8 | 12.8 | 12.6 | 12.8 | 12.3 | 13 | 12.3 | 12.2 | 12.6 |
| 0.285869 | -0.10267 | 0.335035 | -0.09207 | 13.7 | 13.7 | 13.5 | 13.6 | 13.7 | 13.6 | 13.6 | 13.5 | 13.4 |
| 0.28671 | -0.23087 | 0.641382 | -0.09782 | 10.7 | 10 | 10.2 | 10.2 | 10.7 | 10.5 | 10.4 | 10.2 | 10.1 |
| 0.28819 | 0.090632 | 0.943531 | 0.005817 | 13.4 | 13.4 | 13.5 | 13.2 | 13.3 | 13.4 | 13.4 | 13.5 | 13.3 |
| 0.289565 | 0.150221 | 0.153981 | 0.208726 | 11.2 | 11.1 | 11.4 | 11.2 | 11.4 | 11.2 | 11.2 | 11.6 | 11.4 |
| 0.29071 | 0.241065 | 0.955905 | -0.01215 | 10.5 | 11.3 | 11 | 11 | 10.7 | 10.3 | 11 | 10.8 | 10.9 |
| 0.29049 | -0.17568 | 0.490792 | -0.11203 | 10.5 | 10.6 | 10.5 | 10.5 | 10.9 | 10.4 | 10.5 | 10.5 | 10.3 |
| 0.290185 | 0.483923 | 0.252304 | 0.527234 | 11.3 | 12.5 | 12.5 | 12.6 | 11.3 | 12.5 | 12.4 | 12.6 | 12.9 |
| 0.291859 | 0.251319 | 0.727643 | -0.08028 | 14.6 | 15.4 | 15.3 | 15 | 14.5 | 14.8 | 14.9 | 15 | 15.2 |
| 0.293167 | 0.135349 | 0.650688 | 0.056537 | 18 | 18.3 | 18.3 | 18.3 | 17.9 | 18.3 | 18.3 | 18.3 | 18.3 |
| 0.293637 | 0.18004 | 0.147961 | 0.256737 | 10.6 | 10.5 | 10.5 | 10.7 | 10.8 | 10.3 | 11.1 | 10.6 | 10.8 |
| 0.294362 | 0.24041 | 0.035742 | 0.541729 | 8.55 | 8.91 | 9.02 | 9.13 | 8.93 | 9.32 | 9.6 | 9.59 | 8.92 |

| | | | | | | | | | | | | |
|---|---|---|---|---|---|---|---|---|---|---|---|---|
| 0.294656 | -0.16411 | 0.16266 | -0.2252 | 10.8 | 11.1 | 10.9 | 11.1 | 10.7 | 10.8 | 10.7 | 10.9 | 10.6 |
| 0.295041 | 0.221261 | 0.969562 | -0.00777 | 10.8 | 11.4 | 11.5 | 11 | 10.8 | 11.3 | 11.1 | 11.4 | 11.2 |
| 0.295575 | -0.16659 | 0.737406 | -0.05164 | 11 | 10.6 | 10.9 | 10.9 | 11 | 10.9 | 10.9 | 10.5 | 10.9 |
| 0.295965 | 0.243348 | 0.810658 | 0.053849 | 13.2 | 13.7 | 13.6 | 13.6 | 12.9 | 13.5 | 13.3 | 13.6 | 13.7 |
| 0.2967 | 0.363258 | 0.770091 | 0.098312 | 12 | 12.7 | 12.9 | 12.6 | 11.6 | 12.6 | 12.4 | 12.6 | 12.8 |
| 0.297507 | -0.08134 | 0.745965 | -0.02446 | 12.6 | 12.7 | 12.7 | 12.7 | 12.7 | 12.8 | 12.7 | 12.7 | 12.6 |
| 0.298196 | 0.136607 | 0.439013 | 0.099948 | 13.4 | 13.6 | 13.7 | 13.7 | 13.3 | 13.6 | 13.6 | 13.6 | 13.8 |
| 0.299319 | 0.087919 | 0.994747 | -0.00054 | 14.4 | 14.2 | 14.2 | 14.1 | 14.2 | 14.2 | 14.3 | 14.2 | 14.2 |
| 0.299867 | 0.150116 | 0.373607 | 0.127586 | 12.8 | 12.8 | 12.7 | 12.8 | 13 | 12.5 | 13.1 | 12.8 | 12.8 |
| 0.300425 | -0.12875 | 0.129261 | -0.19688 | 14.7 | 14.4 | 14.4 | 14.3 | 14.6 | 14.3 | 14.3 | 14.3 | 14.2 |
| 0.302531 | 0.090404 | 0.475501 | -0.06139 | 16.1 | 16.1 | 16.2 | 16.1 | 15.9 | 15.9 | 16 | 16.1 | 16.1 |
| 0.303616 | -0.23888 | 0.904139 | 0.02699 | 9.16 | 8.51 | 8.46 | 8.9 | 9.17 | 8.88 | 8.77 | 9 | 8.45 |
| 0.30342 | 0.299039 | 0.707238 | 0.105766 | 10.9 | 11.7 | 11.6 | 11.3 | 10.8 | 11.6 | 11.7 | 11.3 | 11.5 |
| 0.302676 | -0.1382 | 0.546189 | -0.07901 | 15.5 | 15.2 | 15.3 | 15.3 | 15.6 | 15.3 | 15.3 | 15.3 | 15.2 |
| 0.303 | 0.147089 | 0.2849 | 0.153145 | 11.7 | 12 | 11.6 | 11.7 | 12 | 11.6 | 11.9 | 12 | 11.9 |
| 0.306608 | 0.088819 | 0.447516 | -0.06494 | 14.1 | 13.9 | 13.9 | 13.8 | 13.9 | 13.8 | 13.9 | 13.9 | 14 |
| 0.306728 | 2.129689 | 0.875331 | -0.31573 | 8 | 9.61 | 7.89 | 9.51 | 7.83 | 0.83 | 7.86 | 8.17 | 8.53 |
| 0.305889 | 0.123544 | 0.913967 | -0.01258 | 12.4 | 12.3 | 12.3 | 12.1 | 12.3 | 12.2 | 12.5 | 12.4 | 12.1 |
| 0.305309 | 0.113692 | 0.962908 | -0.00498 | 11 | 11.2 | 11 | 11 | 10.9 | 11 | 11.2 | 10.9 | 11.1 |
| 0.305218 | -0.10724 | 0.678551 | -0.04207 | 14.2 | 13.9 | 13.9 | 14.1 | 14 | 14.1 | 13.9 | 14 | 13.9 |
| 0.306444 | -0.09316 | 0.118626 | -0.14922 | 15.7 | 15.4 | 15.4 | 15.4 | 15.4 | 15.5 | 15.4 | 15.3 | 15.3 |
| 0.30684 | 0.125818 | 0.393406 | 0.103948 | 11.2 | 11 | 11 | 11 | 11.3 | 10.9 | 11.2 | 11.2 | 11.1 |
| 0.30866 | -0.08383 | 0.227586 | -0.10081 | 18.9 | 19 | 19 | 18.9 | 18.8 | 19 | 18.8 | 18.8 | 18.9 |
| 0.308331 | 0.243309 | 0.24451 | 0.281011 | 10.5 | 10.2 | 10.2 | 10.5 | 9.86 | 10.8 | 10.6 | 10.8 | 10.4 |
| 0.30941 | 1.178288 | 0.464872 | 0.832847 | 7.42 | 10.5 | 10.6 | 9.92 | 6.99 | 10.5 | 10 | 10.3 | 10.6 |
| 0.309745 | -0.12622 | 0.395809 | -0.1044 | 14.5 | 14.4 | 14.2 | 14.3 | 14.6 | 14.2 | 14.3 | 14.3 | 14.1 |
| 0.310202 | -0.15194 | 0.457267 | -0.1095 | 12.6 | 12.8 | 12.8 | 12.8 | 12.5 | 12.9 | 12.5 | 12.5 | 12.8 |
| 0.310837 | -0.12384 | 0.786001 | 0.032116 | 11.1 | 11.1 | 11.3 | 11.4 | 11.3 | 11.2 | 11.1 | 11.4 | 11.1 |
| 0.31215 | -0.16242 | 0.646918 | -0.07159 | 12.7 | 13 | 13 | 13.2 | 12.7 | 13.1 | 12.7 | 12.9 | 13 |
| 0.313031 | -0.10147 | 0.172641 | -0.14123 | 14.8 | 14.5 | 14.5 | 14.6 | 14.6 | 14.5 | 14.5 | 14.5 | 14.5 |
| 0.313151 | -0.19164 | 0.372837 | -0.16803 | 11.4 | 11.3 | 11.5 | 11.2 | 11.2 | 11.9 | 11.2 | 11.2 | 11.2 |
| 0.314202 | 0.237782 | 0.576792 | -0.12873 | 11.4 | 11.4 | 10.6 | 10.9 | 10.7 | 10.8 | 11.3 | 10.8 | 11 |
| 0.314006 | -0.17133 | 0.251501 | -0.19717 | 12.2 | 12 | 11.7 | 11.8 | 12.2 | 11.9 | 11.9 | 11.8 | 11.7 |
| 0.316267 | 0.182755 | 0.971179 | 0.006367 | 12.2 | 12.6 | 12.7 | 12.6 | 12.1 | 12.2 | 12.4 | 12.5 | 12.5 |
| 0.317755 | 0.078123 | 0.484684 | -0.0537 | 14.5 | 14.6 | 14.4 | 14.4 | 14.4 | 14.3 | 14.5 | 14.4 | 14.4 |
| 0.319303 | -0.14941 | 0.144635 | -0.22768 | 14.1 | 13.6 | 13.7 | 13.6 | 13.9 | 13.7 | 13.7 | 13.5 | 13.5 |
| 0.319959 | 0.449653 | 0.885273 | 0.06312 | 10 | 9.74 | 9.59 | 9.46 | 9.07 | 9.67 | 10.7 | 9.94 | 8.87 |
| 0.321092 | 0.141041 | 0.874293 | -0.02177 | 11.9 | 12.2 | 12.1 | 12.1 | 11.6 | 11.9 | 12 | 12.1 | 12 |
| 0.321337 | -0.16544 | 0.13977 | -0.25683 | 12.5 | 12.2 | 11.9 | 11.9 | 12.2 | 12.2 | 12 | 12 | 11.8 |
| 0.321537 | 0.43157 | 0.449935 | 0.324315 | 10.8 | 12 | 12 | 11.8 | 10.7 | 11.8 | 11.7 | 11.9 | 12.1 |
| 0.32067 | 0.080359 | 0.327892 | 0.079089 | 13.8 | 14 | 14 | 14 | 13.8 | 14 | 14 | 14 | 14 |
| 0.324958 | -0.10381 | 0.889912 | 0.014134 | 12.5 | 12.4 | 12.4 | 12.3 | 12.6 | 12.7 | 12.5 | 12.4 | 12.4 |
| 0.324242 | -0.17161 | 0.064097 | -0.35148 | 10.1 | 9.9 | 10.4 | 10 | 10.1 | 9.75 | 9.76 | 10 | 9.59 |
| 0.324909 | 0.089228 | 0.764327 | 0.026372 | 15.2 | 15.4 | 15.3 | 15.3 | 15.2 | 15.3 | 15.2 | 15.4 | 15.3 |
| 0.324168 | 0.155455 | 0.171348 | 0.222515 | 11.1 | 11 | 11 | 11.1 | 10.9 | 11.4 | 11.5 | 11 | 11.3 |
| 0.32549 | -0.12645 | 0.238993 | -0.15364 | 12.1 | 12.1 | 12 | 12 | 12 | 12 | 12.1 | 11.9 | 11.6 |
| 0.326374 | -0.21429 | 0.577188 | -0.11904 | 12.1 | 12.1 | 11.3 | 12 | 12 | 11.8 | 11.7 | 11.6 | 11.8 |
| 0.329286 | -0.33875 | 0.176784 | -0.48343 | 10.2 | 9.78 | 9.87 | 9.9 | 10.1 | 9.35 | 10.2 | 9.19 | 9 |
| 0.328957 | 0.123462 | 0.52105 | 0.079672 | 14.1 | 13.8 | 13.9 | 13.7 | 14 | 13.8 | 14 | 14 | 13.9 |
| 0.330112 | 0.749173 | 0.26186 | 0.872403 | 8.06 | 10 | 10.4 | 10.3 | 8.27 | 10.3 | 10.3 | 10.4 | 10.5 |

| | | | | | | | | | | | | |
|---|---|---|---|---|---|---|---|---|---|---|---|---|
| 0.329779 | 0.28164 | 0.27576 | 0.317565 | 11.3 | 12 | 12 | 12.1 | 11.3 | 12 | 12 | 12.1 | 12.1 |
| 0.330953 | -0.62835 | 0.957633 | -0.03325 | 8.89 | 7.77 | 9.56 | 9.78 | 9.94 | 8.28 | 8.04 | 9.43 | 8.64 |
| 0.331485 | -0.25663 | 0.748305 | -0.0824 | 9.17 | 8.63 | 9.17 | 9.61 | 8.79 | 9.08 | 8.79 | 9.22 | 8.71 |
| 0.331293 | 0.573305 | 0.469107 | 0.421113 | 13 | 14.3 | 14.2 | 14.4 | 12.5 | 14.2 | 13.8 | 14.4 | 14.5 |
| 0.33546 | 0.111707 | 0.173691 | -0.16291 | 13.6 | 13.9 | 13.6 | 13.4 | 13.3 | 13.5 | 13.4 | 13.6 | 13.5 |
| 0.335673 | -0.17104 | 0.777315 | -0.04882 | 10.8 | 11.1 | 10.6 | 10.7 | 11.1 | 11.1 | 10.9 | 10.7 | 10.8 |
| 0.335128 | -0.08518 | 0.155665 | -0.13031 | 15 | 15.1 | 15 | 14.9 | 15 | 15.1 | 15 | 14.9 | 14.8 |
| 0.335813 | 0.315514 | 0.524154 | 0.205123 | 13.7 | 14.6 | 14.7 | 14.5 | 13.7 | 14.6 | 14.4 | 14.5 | 14.7 |
| 0.336341 | -0.19986 | 0.584402 | -0.1113 | 12.3 | 11.8 | 12.2 | 12 | 12.6 | 12 | 12 | 11.9 | 12.1 |
| 0.336815 | 0.176414 | 0.661349 | 0.07848 | 10.5 | 11 | 11 | 10.9 | 10.7 | 10.7 | 10.7 | 10.9 | 11.2 |
| 0.337712 | 0.133878 | 0.925842 | -0.0126 | 11.4 | 11.3 | 11.2 | 11.2 | 11 | 11.3 | 11.5 | 11.3 | 11.1 |
| 0.337451 | -0.46071 | 0.752425 | -0.14733 | 9.68 | 9.8 | 9.02 | 9.97 | 10.7 | 8.8 | 9.01 | 9.61 | 9.44 |
| 0.3395 | -0.37554 | 0.792596 | -0.10046 | 9.74 | 9.33 | 9.26 | 9.58 | 9.99 | 9.58 | 9.84 | 9.78 | 8.41 |
| 0.33932 | -0.10724 | 0.699773 | -0.04217 | 14.8 | 14.6 | 14.6 | 14.5 | 14.9 | 14.7 | 14.6 | 14.6 | 14.6 |
| 0.339127 | -0.43818 | 0.348188 | -0.42953 | 9.52 | 9.96 | 9.49 | 9.32 | 9.69 | 9.98 | 8.14 | 9.7 | 9.84 |
| 0.342517 | 0.286023 | 0.648614 | 0.134106 | 12 | 12.9 | 12.8 | 12.7 | 11.9 | 12.6 | 12.6 | 12.7 | 12.8 |
| 0.343239 | -0.16292 | 0.281757 | -0.18669 | 12.2 | 11.8 | 11.8 | 11.6 | 12.1 | 12 | 11.8 | 11.7 | 11.8 |
| 0.343879 | 0.116209 | 0.229101 | 0.150491 | 17 | 17.2 | 17.2 | 17.3 | 17 | 17.3 | 17.3 | 17.3 | 17.3 |
| 0.345359 | -0.32011 | 0.312614 | -0.34407 | 9.72 | 9.19 | 9.29 | 9.98 | 9.04 | 9.1 | 9.42 | 8.52 | 9.22 |
| 0.346442 | 0.1626 | 0.772615 | -0.04856 | 13.7 | 13.5 | 13.6 | 13.4 | 13.8 | 13.1 | 13.7 | 13.6 | 13.4 |
| 0.346624 | -0.22069 | 0.928445 | -0.02044 | 10.1 | 10.8 | 10.8 | 10.6 | 10.7 | 11 | 10.4 | 10.8 | 10.5 |
| 0.34666 | -0.13271 | 0.409236 | -0.11556 | 14.4 | 14 | 14.1 | 14.2 | 14.4 | 14.1 | 14.1 | 14 | 14.1 |
| 0.348122 | -0.19201 | 0.155138 | -0.30269 | 13.4 | 14 | 14 | 13.9 | 13.4 | 13.6 | 13.3 | 13.4 | 13.7 |
| 0.348436 | 0.617961 | 0.554244 | 0.382804 | 11.2 | 13.1 | 13.1 | 12.9 | 11.1 | 12.6 | 12.5 | 12.8 | 13.2 |
| 0.349655 | -0.25794 | 0.594837 | -0.14375 | 11.2 | 10.6 | 10.5 | 10.9 | 11.3 | 10.4 | 10.4 | 10.8 | 10.6 |
| 0.350716 | 0.118264 | 0.513499 | -0.08156 | 11.7 | 11.8 | 11.9 | 11.4 | 11.8 | 11.6 | 11.8 | 11.7 | 11.6 |
| 0.351162 | 2.114044 | 0.94508 | 0.151673 | 7.94 | 9.2 | 7.8 | 9.05 | 0.49 | 9.51 | 8.27 | 8.11 | 9.01 |
| 0.353567 | 0.240569 | 0.39216 | -0.22093 | 11.7 | 11.1 | 10.7 | 10.9 | 10.5 | 10.8 | 11.1 | 11.1 | 10.7 |
| 0.35323 | -0.25291 | 0.095425 | -0.48582 | 9 | 9.17 | 9.49 | 9.12 | 9.41 | 8.44 | 8.82 | 8.92 | 8.47 |
| 0.353811 | -0.12073 | 0.15231 | -0.19423 | 14.7 | 14.6 | 14.5 | 14.4 | 14.8 | 14.4 | 14.4 | 14.4 | 14.3 |
| 0.354453 | -0.08824 | 0.572424 | -0.05281 | 13.8 | 13.5 | 13.5 | 13.7 | 13.6 | 13.6 | 13.6 | 13.5 | 13.6 |
| 0.356365 | 0.168451 | 0.981721 | 0.004065 | 11.8 | 11.9 | 11.9 | 12 | 11.8 | 11.4 | 11.8 | 12.1 | 11.7 |
| 0.357782 | -0.12933 | 0.894139 | -0.0182 | 14.3 | 14.6 | 14.7 | 14.8 | 14.5 | 14.8 | 14.6 | 14.5 | 14.6 |
| 0.35778 | 0.236697 | 0.685172 | 0.101957 | 13 | 13.8 | 13.7 | 13.6 | 13.1 | 13.5 | 13.4 | 13.7 | 13.8 |
| 0.356118 | -0.12373 | 0.642346 | -0.06093 | 13.5 | 13.2 | 13.2 | 13.2 | 13.5 | 13.4 | 13.4 | 13.1 | 13.2 |
| 0.357431 | 0.170587 | 0.231553 | 0.226283 | 11 | 11.5 | 11.6 | 11.7 | 11.3 | 11.3 | 11.7 | 11.5 | 11.6 |
| 0.356072 | 0.15156 | 0.477431 | 0.115297 | 13.7 | 14 | 14.1 | 14 | 13.6 | 14.1 | 13.9 | 14.1 | 14.2 |
| 0.357398 | 0.140963 | 0.445788 | 0.115726 | 13.5 | 13.1 | 13 | 13.1 | 13.4 | 13 | 13.3 | 13.3 | 13.4 |
| 0.359317 | 0.071417 | 0.558124 | 0.044859 | 15.3 | 15.3 | 15.3 | 15.2 | 15.3 | 15.3 | 15.4 | 15.3 | 15.3 |
| 0.363913 | 0.17381 | 0.15382 | -0.28473 | 10.7 | 10.3 | 10.4 | 9.7 | 10.1 | 10.3 | 10 | 10.1 | 10.4 |
| 0.363553 | -2.84001 | 0.82456 | 0.674852 | 7.41 | 8.02 | -0.1 | 8.79 | 9.03 | 8.03 | 8.53 | 9.33 | -0.5 |
| 0.363175 | -0.1208 | 0.89149 | 0.017631 | 10.9 | 10.9 | 10.9 | 10.9 | 10.9 | 11.3 | 11.1 | 10.8 | 11 |
| 0.363319 | 0.090984 | 0.858331 | 0.017389 | 14.3 | 14.4 | 14.2 | 14.2 | 14.4 | 14.1 | 14.3 | 14.3 | 14.4 |
| 0.364149 | 0.205321 | 0.790993 | 0.058445 | 14 | 14.6 | 14.4 | 14.5 | 13.8 | 14.2 | 14.1 | 14.5 | 14.5 |
| 0.363744 | -0.08742 | 0.20416 | -0.12559 | 13.9 | 13.8 | 13.8 | 13.8 | 13.9 | 13.6 | 13.6 | 13.7 | 13.8 |
| 0.363484 | -0.14257 | 0.416095 | -0.12687 | 12 | 11.8 | 12.1 | 11.8 | 12.3 | 11.9 | 11.9 | 11.7 | 12 |
| 0.365467 | -0.15782 | 0.201602 | -0.22904 | 12.6 | 12.4 | 12.2 | 12 | 12.5 | 12.5 | 12.2 | 12.2 | 12.1 |
| 0.366884 | 0.287508 | 0.963847 | -0.01405 | 11 | 10.4 | 10.3 | 10.7 | 10.5 | 9.64 | 10.5 | 10.5 | 10.7 |
| 0.367857 | 0.112308 | 0.807881 | 0.029561 | 10.4 | 10.3 | 10.2 | 10.2 | 10.3 | 10.1 | 10.5 | 10.3 | 10.2 |
| 0.367813 | 0.122809 | 0.155902 | 0.201749 | 11.6 | 11.4 | 11.6 | 11.9 | 11.5 | 11.4 | 11.7 | 11.8 | 11.8 |

| | | | | | | | | | | | | |
|---|---|---|---|---|---|---|---|---|---|---|---|---|
| 0.369547 | -0.15556 | 0.515836 | -0.11118 | 14.1 | 14.5 | 14.5 | 14.6 | 14.1 | 14.6 | 14.2 | 14.3 | 14.4 |
| 0.370391 | -0.2688 | 0.057268 | -0.63004 | 9.85 | 9.94 | 9.72 | 9.86 | 8.73 | 9.84 | 9.23 | 9.04 | 9.35 |
| 0.371613 | -0.28399 | 0.462324 | 0.231578 | 8.19 | 9.06 | 9.41 | 9.2 | 9.72 | 9.29 | 9.34 | 9.04 | 8.97 |
| 0.371511 | -0.2418 | 0.812281 | -0.06266 | 11.8 | 11.2 | 11.2 | 11.5 | 12 | 11.2 | 11.6 | 11.3 | 11 |
| 0.371483 | 0.190784 | 0.26509 | 0.241565 | 10.3 | 10.5 | 10.9 | 10.9 | 10.4 | 10.5 | 11.1 | 10.7 | 10.5 |
| 0.372709 | 0.136087 | 0.741309 | 0.049236 | 12.8 | 13 | 13.1 | 13.1 | 12.6 | 13 | 13 | 12.9 | 13.2 |
| 0.376514 | -0.30314 | 0.128626 | -0.5493 | 10.7 | 10.3 | 10.3 | 10.4 | 10.6 | 9.57 | 10.4 | 9.46 | 9.82 |
| 0.379496 | -0.46125 | 0.748656 | 0.164381 | 8.42 | 9.08 | 10 | 9.15 | 9.98 | 10.3 | 9.08 | 8.88 | 10.1 |
| 0.379646 | 0.377868 | 0.59818 | 0.222891 | 8.63 | 8.69 | 9.28 | 7.74 | 9.05 | 9.34 | 8.74 | 9.26 | 9.27 |
| 0.380283 | -0.15479 | 0.60875 | -0.08876 | 10.7 | 11 | 11.1 | 11.2 | 11 | 10.7 | 10.9 | 10.7 | 10.9 |
| 0.380955 | -0.27477 | 0.198838 | -0.41544 | 9.99 | 9.68 | 10.2 | 9.74 | 9.5 | 10.3 | 10 | 9.6 | 9.02 |
| 0.382876 | 0.246807 | 0.595218 | 0.147836 | 13.5 | 14.2 | 14.2 | 14.3 | 13.3 | 14 | 14 | 14.1 | 14.2 |
| 0.382612 | -0.17496 | 0.253397 | -0.23319 | 13.6 | 14.2 | 13.9 | 13.9 | 13.6 | 14.1 | 14 | 13.5 | 13.5 |
| 0.383501 | -0.10621 | 0.067772 | -0.2437 | 12.6 | 12.5 | 12.6 | 12.6 | 12.3 | 12.3 | 12.2 | 12.4 | 12.2 |
| 0.384667 | -0.10208 | 0.97354 | 0.003796 | 13 | 12.8 | 13 | 13 | 13.1 | 13.1 | 13.1 | 12.8 | 13 |
| 0.385654 | -0.11731 | 0.263986 | -0.15364 | 15.9 | 16.3 | 16.2 | 16.2 | 15.9 | 16.2 | 15.9 | 15.9 | 16.1 |
| 0.386131 | 0.074079 | 0.063522 | -0.17436 | 13.8 | 14 | 13.9 | 13.7 | 13.6 | 13.7 | 13.8 | 13.7 | 13.7 |
| 0.387528 | 0.17107 | 0.590074 | -0.10501 | 12.2 | 12.7 | 12.6 | 12.4 | 12 | 12.3 | 12.2 | 12.4 | 12.6 |
| 0.388182 | -0.26613 | 0.142382 | -0.47509 | 10.3 | 9.93 | 10 | 9.97 | 9.55 | 10.1 | 8.97 | 10.2 | 9.69 |
| 0.387294 | -0.07359 | 0.577872 | -0.04667 | 15.5 | 15.3 | 15.4 | 15.5 | 15.4 | 15.5 | 15.4 | 15.3 | 15.4 |
| 0.387986 | -0.15145 | 0.450935 | -0.13146 | 11.2 | 10.7 | 11 | 11.2 | 11 | 10.8 | 11.1 | 10.9 | 10.6 |
| 0.389605 | -0.21958 | 0.936027 | -0.01998 | 12 | 12.6 | 12.9 | 12.9 | 12.3 | 12.8 | 12.4 | 12.7 | 12.3 |
| 0.390352 | -0.37758 | 0.230401 | -0.54007 | 10.2 | 11.1 | 11.4 | 11.2 | 9.9 | 11.1 | 10.1 | 10.2 | 10.8 |
| 0.3903 | 0.158252 | 0.514996 | 0.118658 | 15.6 | 16 | 16.1 | 16 | 15.6 | 16 | 15.9 | 16 | 16.1 |
| 0.391736 | -0.15137 | 0.240313 | -0.21219 | 14 | 13.6 | 13.6 | 13.7 | 14 | 13.4 | 13.4 | 13.6 | 13.5 |
| 0.393048 | -0.07962 | 0.944363 | -0.00635 | 10.9 | 10.9 | 11.2 | 11.2 | 11.1 | 11 | 11 | 11 | 11 |
| 0.392996 | -0.22972 | 0.384529 | -0.23407 | 13.4 | 12.8 | 12.5 | 12.8 | 13.3 | 12.7 | 12.5 | 12.6 | 12.9 |
| 0.395726 | 0.11813 | 0.167907 | -0.19981 | 11 | 11.1 | 11 | 10.8 | 10.7 | 10.7 | 11.1 | 10.6 | 10.9 |
| 0.395957 | 0.380686 | 0.285024 | 0.486464 | 9.5 | 10.8 | 11 | 10.8 | 9.9 | 10.9 | 10.7 | 10.9 | 11.2 |
| 0.398324 | -0.40927 | 0.634483 | 0.226585 | 9.7 | 8.49 | 9.1 | 9.7 | 10.4 | 9.05 | 9.63 | 9.7 | 8.63 |
| 0.397389 | 0.284732 | 0.704677 | -0.12506 | 9.99 | 10.2 | 10.9 | 10.5 | 10.1 | 9.38 | 10.4 | 10.3 | 10.1 |
| 0.397284 | 1.881939 | 0.831835 | -0.46146 | 8.3 | 9.08 | 9.46 | 9.28 | 9.74 | 0.79 | 8.05 | 8.73 | 8.67 |
| 0.398139 | 0.18773 | 0.546435 | 0.132409 | 12.7 | 13.4 | 13.2 | 13.3 | 12.7 | 13.2 | 13.1 | 13.2 | 13.4 |
| 0.397834 | 0.168061 | 0.470113 | 0.142629 | 14.4 | 15 | 15 | 14.9 | 14.4 | 14.9 | 14.8 | 14.9 | 15 |
| 0.398821 | 0.577872 | 0.634796 | 0.319983 | 12.2 | 14.4 | 14.2 | 14 | 12.3 | 13.8 | 13.5 | 14 | 14.2 |
| 0.400048 | 0.456593 | 0.682047 | 0.218229 | 11.4 | 12.9 | 12.9 | 12.6 | 11.2 | 12.7 | 12.5 | 12.5 | 12.9 |
| 0.40182 | -0.11574 | 0.296872 | -0.14593 | 16.7 | 16.4 | 16.3 | 16.4 | 16.6 | 16.3 | 16.4 | 16.3 | 16.2 |
| 0.402369 | 0.078103 | 0.765946 | 0.027192 | 13.6 | 13.9 | 13.8 | 13.7 | 13.7 | 13.7 | 13.8 | 13.8 | 13.7 |
| 0.403099 | 0.520989 | 0.727337 | 0.213041 | 11.1 | 12.9 | 12.8 | 12.6 | 10.8 | 12.5 | 12.2 | 12.6 | 12.6 |
| 0.402786 | -0.15616 | 0.385256 | -0.16236 | 11.1 | 11.2 | 11.2 | 11.3 | 10.8 | 11.4 | 10.9 | 11.2 | 10.9 |
| 0.403935 | 0.151738 | 0.571242 | 0.101633 | 9.83 | 9.68 | 9.99 | 9.98 | 9.88 | 9.49 | 9.87 | 10.2 | 9.72 |
| 0.404415 | 0.187564 | 0.691129 | 0.087765 | 12.9 | 13.5 | 13.6 | 13.5 | 12.9 | 13.3 | 13.3 | 13.5 | 13.4 |
| 0.405337 | 0.180712 | 0.188055 | 0.296334 | 14.8 | 15.3 | 15.1 | 15.4 | 14.8 | 15.4 | 15.2 | 15.3 | 15.5 |
| 0.405173 | -0.18712 | 0.482756 | -0.1567 | 13.4 | 12.9 | 12.5 | 12.8 | 13.1 | 12.9 | 12.7 | 12.7 | 13 |
| 0.407087 | -0.16513 | 0.885692 | -0.02799 | 11 | 11 | 10.4 | 10.8 | 10.9 | 11.2 | 10.7 | 11 | 10.6 |
| 0.407002 | 0.098991 | 0.533263 | 0.073625 | 13.2 | 13.6 | 13.6 | 13.6 | 13.3 | 13.5 | 13.6 | 13.6 | 13.5 |
| 0.406937 | 0.130551 | 0.353493 | 0.146931 | 12.2 | 12 | 12.4 | 11.9 | 12.5 | 12.2 | 12.4 | 12.3 | 12.3 |
| 0.4078 | 0.092428 | 0.857932 | 0.01955 | 12.4 | 12.4 | 12.4 | 12.2 | 12.4 | 12.3 | 12.5 | 12.2 | 12.5 |
| 0.408632 | -0.11202 | 0.744547 | -0.04331 | 13.3 | 13.4 | 12.9 | 13.3 | 13.3 | 13.2 | 13.3 | 13.1 | 13.1 |
| 0.411211 | -0.30788 | 0.259868 | 0.430839 | 8.25 | 9.32 | 8.21 | 9.26 | 9.28 | 9.46 | 9.57 | 9.06 | 8.45 |

| | | | | | | | | | | | | |
|---|---|---|---|---|---|---|---|---|---|---|---|---|
| 0.410509 | 0.116722 | 0.767016 | -0.04119 | 14.3 | 14.6 | 14.5 | 14.5 | 14 | 14.4 | 14.4 | 14.5 | 14.5 |
| 0.410951 | -0.0975 | 0.774784 | -0.03325 | 13.2 | 13.5 | 13.3 | 13.5 | 13.2 | 13.6 | 13.4 | 13.2 | 13.4 |
| 0.412453 | 0.094321 | 0.523868 | -0.07269 | 12.4 | 12 | 12.2 | 12 | 12 | 12 | 12.1 | 12.1 | 12 |
| 0.412275 | 0.117985 | 0.747175 | 0.045502 | 11.7 | 11.7 | 11.6 | 11.5 | 11.9 | 11.3 | 11.6 | 11.8 | 11.7 |
| 0.413104 | -0.08588 | 0.408797 | -0.08671 | 13.7 | 13.9 | 13.8 | 13.8 | 13.7 | 13.9 | 13.6 | 13.7 | 13.8 |
| 0.41555 | 0.143732 | 0.231532 | -0.21688 | 11.2 | 10.8 | 11.2 | 10.4 | 10.8 | 10.9 | 10.8 | 10.8 | 10.9 |
| 0.415693 | 0.127965 | 0.962637 | 0.007202 | 13.4 | 13.6 | 13.7 | 13.6 | 13.1 | 13.7 | 13.6 | 13.6 | 13.6 |
| 0.414397 | 0.067805 | 0.751966 | 0.025756 | 14.8 | 14.7 | 14.6 | 14.6 | 14.7 | 14.6 | 14.7 | 14.7 | 14.6 |
| 0.415355 | 0.078291 | 0.221434 | 0.120946 | 17.3 | 17.1 | 17 | 17.1 | 17.3 | 17.2 | 17.3 | 17.3 | 17.2 |
| 0.414573 | -0.10175 | 0.225524 | -0.15539 | 15 | 15.2 | 15.3 | 15.2 | 14.9 | 15.3 | 14.9 | 15 | 15.2 |
| 0.41657 | 0.213899 | 0.056066 | 0.558914 | 9.07 | 8.6 | 8.38 | 9.33 | 9.15 | 8.61 | 9.11 | 9.05 | 9.57 |
| 0.417772 | -0.15888 | 0.267568 | -0.22171 | 12.7 | 12.5 | 12.2 | 12.4 | 12.7 | 12.1 | 12.2 | 12.4 | 12.2 |
| 0.417899 | -0.0969 | 0.337143 | -0.11586 | 15.1 | 14.9 | 14.9 | 15 | 15.1 | 14.8 | 15 | 14.9 | 14.7 |
| 0.421108 | -0.09937 | 0.039646 | -0.28856 | 12 | 12 | 11.9 | 11.7 | 11.8 | 11.9 | 11.9 | 11.7 | 11.5 |
| 0.420411 | -0.32637 | 0.052163 | -0.8781 | 11.1 | 9.56 | 10.8 | 9.98 | 9.74 | 10.1 | 9.7 | 9.19 | 9.94 |
| 0.421261 | -0.29392 | 0.755368 | 0.111764 | 9.12 | 9.09 | 9.87 | 9.64 | 10.1 | 9.6 | 9.71 | 9.97 | 8.74 |
| 0.421103 | 0.339343 | 0.70999 | 0.154167 | 8.07 | 8.61 | 8.12 | 8.42 | 7.15 | 8.68 | 7.99 | 8.75 | 8.53 |
| 0.423413 | 0.088484 | 0.262882 | -0.12639 | 14.2 | 14 | 14.1 | 13.7 | 14 | 13.9 | 14 | 14 | 13.9 |
| 0.423431 | -0.33719 | 0.660246 | -0.18237 | 9.84 | 9.84 | 9.83 | 9.89 | 9.86 | 10.2 | 10.5 | 9.89 | 8.63 |
| 0.42536 | 0.137799 | 0.543973 | -0.10395 | 12 | 11.8 | 11.8 | 11.6 | 11.9 | 11.4 | 12.1 | 11.6 | 11.7 |
| 0.426396 | -0.09812 | 0.200782 | -0.16339 | 11.4 | 11.4 | 11.5 | 11.6 | 11.2 | 11.5 | 11.4 | 11.2 | 11.3 |
| 0.428055 | 0.297297 | 0.56241 | 0.215146 | 13.4 | 14.3 | 14.3 | 14.2 | 13.2 | 14.3 | 14 | 14.2 | 14.4 |
| 0.428632 | -0.06375 | 0.207145 | -0.10503 | 16.7 | 16.8 | 16.7 | 16.7 | 16.6 | 16.8 | 16.6 | 16.6 | 16.6 |
| 0.429861 | 0.066547 | 0.767772 | -0.02444 | 16.7 | 16.6 | 16.5 | 16.6 | 16.6 | 16.4 | 16.6 | 16.6 | 16.6 |
| 0.430178 | 0.115391 | 0.308159 | 0.151206 | 12.8 | 12.5 | 12.7 | 12.6 | 12.9 | 12.5 | 12.8 | 12.9 | 12.7 |
| 0.430761 | -0.09759 | 0.614324 | -0.06164 | 12.8 | 12.8 | 12.6 | 12.6 | 12.9 | 12.9 | 12.8 | 12.5 | 12.7 |
| 0.431834 | -0.07138 | 0.999727 | -3E-05 | 14.6 | 14.5 | 14.4 | 14.5 | 14.6 | 14.7 | 14.5 | 14.5 | 14.5 |
| 0.432964 | -0.24164 | 0.900872 | -0.03761 | 10 | 11.2 | 11.1 | 10.9 | 10.8 | 11.2 | 10.6 | 10.7 | 10.9 |
| 0.432654 | 0.218397 | 0.726374 | 0.095786 | 12.9 | 13.5 | 13.5 | 13.5 | 12.6 | 13.4 | 13.2 | 13.4 | 13.5 |
| 0.433881 | 0.057351 | 0.020506 | -0.20133 | 13.1 | 13.1 | 13.2 | 13 | 12.8 | 12.9 | 13 | 12.9 | 12.9 |
| 0.435275 | 0.057376 | 0.24542 | 0.087624 | 13.4 | 13.3 | 13.3 | 13.4 | 13.3 | 13.3 | 13.4 | 13.5 | 13.4 |
| 0.435148 | -0.19614 | 0.504526 | -0.1668 | 15.2 | 14.5 | 14.6 | 14.7 | 15.2 | 14.5 | 14.8 | 14.5 | 14.5 |
| 0.435824 | 0.320827 | 0.543031 | 0.248399 | 12.9 | 14.2 | 14.1 | 14.1 | 13 | 13.9 | 13.8 | 14 | 14.2 |
| 0.437117 | -0.15683 | 0.053721 | -0.43454 | 12.3 | 12.2 | 12.3 | 12 | 12.2 | 11.7 | 12.2 | 11.5 | 11.8 |
| 0.437374 | 0.603741 | 0.56065 | 0.448286 | 10.3 | 12.8 | 12.5 | 12.3 | 10.5 | 12.3 | 11.9 | 12.4 | 12.6 |
| 0.437987 | 0.270479 | 0.404572 | -0.29159 | 9.79 | 10.8 | 10.8 | 10 | 10.2 | 9.51 | 10.2 | 10.6 | 9.85 |
| 0.440199 | -0.13575 | 0.435617 | -0.13717 | 12.8 | 12.5 | 12.3 | 12.3 | 12.8 | 12.5 | 12.4 | 12.4 | 12.3 |
| 0.441346 | 0.055718 | 0.786369 | -0.01927 | 15.1 | 15 | 15.1 | 15.1 | 15 | 15 | 15.1 | 15.1 | 15.1 |
| 0.444636 | -0.12502 | 0.104805 | -0.28489 | 15.2 | 14.6 | 14.9 | 14.7 | 14.9 | 14.6 | 14.6 | 14.6 | 14.7 |
| 0.445954 | -0.27731 | 0.109837 | -0.62317 | 10.3 | 10.6 | 10.4 | 9.67 | 9.84 | 10.8 | 9.48 | 9.53 | 10.4 |
| 0.445915 | -0.23955 | 0.366965 | -0.28579 | 10.7 | 11 | 10.9 | 10.6 | 10.3 | 11.5 | 10.8 | 10.6 | 10.3 |
| 0.448576 | 1.751269 | 0.84386 | -0.44699 | 9.28 | 8.64 | 9.27 | 9.88 | 9.95 | 0.78 | 8.17 | 8.45 | 9.24 |
| 0.447957 | 0.08149 | 0.849829 | 0.019964 | 13.5 | 13.5 | 13.6 | 13.5 | 13.3 | 13.6 | 13.6 | 13.4 | 13.6 |
| 0.448824 | 0.139156 | 0.770386 | 0.052746 | 14.8 | 15.1 | 15.1 | 15.1 | 14.5 | 15.1 | 14.9 | 15 | 15.2 |
| 0.447195 | 0.111072 | 0.691222 | 0.05722 | 11.3 | 11.2 | 11 | 11.3 | 11.1 | 10.8 | 11.3 | 11.3 | 11 |
| 0.448903 | -0.1046 | 0.447648 | -0.1049 | 11.3 | 11 | 10.9 | 11 | 11 | 11.1 | 10.7 | 11 | 11.1 |
| 0.449781 | -0.17599 | 0.634998 | -0.10925 | 10.7 | 10.2 | 9.97 | 10 | 10.6 | 10.3 | 9.98 | 10.1 | 10.4 |
| 0.453391 | -0.05531 | 0.021008 | -0.2019 | 15.1 | 15 | 15 | 14.9 | 14.9 | 14.9 | 14.9 | 14.8 | 14.8 |
| 0.452444 | 0.062338 | 0.289375 | -0.0896 | 13.4 | 13.6 | 13.5 | 13.3 | 13.4 | 13.4 | 13.4 | 13.4 | 13.5 |
| 0.452628 | 0.064827 | 0.605501 | -0.04413 | 13.8 | 13.9 | 13.8 | 13.7 | 13.7 | 13.8 | 13.8 | 13.9 | 13.7 |

| | | | | | | | | | | | | |
|---|---|---|---|---|---|---|---|---|---|---|---|---|
| 0.452946 | -0.11047 | 0.109886 | -0.25223 | 11.2 | 11.3 | 11.4 | 11.3 | 11.3 | 10.9 | 11.1 | 11.2 | 10.9 |
| 0.451646 | -0.15515 | 0.2052 | -0.27066 | 12.1 | 11.7 | 11.9 | 11.8 | 12 | 11.5 | 12 | 11.5 | 11.3 |
| 0.453548 | 0.051585 | 0.803021 | 0.016879 | 16 | 15.9 | 15.8 | 15.9 | 15.9 | 15.9 | 15.9 | 15.9 | 15.9 |
| 0.453184 | 0.400062 | 0.708276 | 0.196716 | 12.2 | 13.6 | 13.4 | 13.4 | 11.8 | 13.4 | 13 | 13.3 | 13.5 |
| 0.451633 | -0.07615 | 0.590564 | -0.05392 | 13.2 | 13.2 | 13.3 | 13.4 | 13.1 | 13.3 | 13.1 | 13.2 | 13.2 |
| 0.454685 | 0.32397 | 0.685192 | 0.17333 | 11.2 | 12.7 | 12.3 | 12.4 | 11.4 | 11.9 | 11.9 | 12.3 | 12.6 |
| 0.455645 | -0.11604 | 0.664101 | 0.066708 | 11.2 | 11.6 | 11.3 | 11.5 | 11.6 | 11.5 | 11.2 | 11.3 | 11.7 |
| 0.456194 | -0.05178 | 0.653544 | -0.03082 | 13.5 | 13.5 | 13.6 | 13.6 | 13.5 | 13.5 | 13.5 | 13.5 | 13.5 |
| 0.457738 | -0.18377 | 0.02339 | -0.66094 | 11 | 11.1 | 10.9 | 10.9 | 10.2 | 10.5 | 10.6 | 10.5 | 9.87 |
| 0.457852 | -0.05849 | 0.340981 | -0.07594 | 13 | 12.9 | 12.8 | 12.8 | 12.9 | 12.9 | 12.9 | 12.8 | 12.7 |
| 0.458991 | 0.174121 | 0.579827 | -0.12912 | 11.3 | 11.2 | 10.7 | 10.7 | 10.9 | 10.8 | 11.3 | 11 | 10.6 |
| 0.460333 | 0.136345 | 0.394451 | 0.158234 | 10.6 | 10.1 | 10.4 | 10.3 | 10.6 | 10.2 | 10.7 | 10.2 | 10.6 |
| 0.46382 | -0.25247 | 0.679213 | 0.140764 | 10.8 | 10.5 | 10 | 10.5 | 11.3 | 10.7 | 10.8 | 11 | 9.97 |
| 0.463889 | 1.692555 | 0.969176 | -0.08769 | 8.17 | 8.81 | 8.09 | 9.39 | 0.48 | 9.86 | 8.46 | 7.71 | 8.63 |
| 0.464139 | -0.17708 | 0.414763 | -0.19815 | 9.91 | 10.3 | 10 | 9.77 | 10.6 | 9.84 | 10.1 | 9.87 | 9.63 |
| 0.466532 | 0.068612 | 0.667966 | -0.03995 | 12.5 | 12.3 | 12.5 | 12.3 | 12.2 | 12.5 | 12.4 | 12.4 | 12.4 |
| 0.467666 | 0.187306 | 0.843436 | -0.0501 | 12.9 | 13.4 | 13.4 | 13.3 | 12.4 | 13.2 | 13.1 | 13.2 | 13.2 |
| 0.467694 | -0.14493 | 0.30582 | -0.20804 | 14.9 | 14.3 | 14.4 | 14.3 | 14.9 | 14.3 | 14.3 | 14.4 | 14.3 |
| 0.465555 | -0.16458 | 0.631645 | -0.10702 | 13.7 | 13.8 | 13.7 | 14.1 | 13.2 | 14 | 13.8 | 13.5 | 13.6 |
| 0.466719 | 0.075398 | 0.346038 | 0.09882 | 11.7 | 11.6 | 11.7 | 11.7 | 11.9 | 11.6 | 11.7 | 11.7 | 11.9 |
| 0.466999 | -0.05621 | 0.598947 | -0.0403 | 14 | 13.8 | 13.8 | 13.9 | 13.9 | 13.8 | 13.9 | 13.8 | 13.9 |
| 0.470055 | -0.08644 | 0.268051 | -0.13577 | 13.2 | 12.9 | 12.9 | 12.8 | 13 | 13 | 12.9 | 12.8 | 13 |
| 0.470944 | 0.087636 | 0.722967 | -0.04252 | 14.4 | 14.1 | 14.1 | 13.9 | 14.2 | 13.9 | 14.2 | 14.1 | 14.1 |
| 0.471845 | -0.07258 | 0.728972 | -0.03449 | 13.4 | 13.2 | 13.3 | 13.3 | 13.5 | 13.3 | 13.3 | 13.4 | 13.2 |
| 0.472349 | -0.07117 | 0.222114 | -0.12503 | 13.8 | 13.8 | 13.7 | 13.8 | 13.8 | 13.5 | 13.6 | 13.7 | 13.7 |
| 0.473065 | 0.090234 | 0.912183 | 0.013636 | 13.7 | 13.4 | 13.5 | 13.3 | 13.7 | 13.4 | 13.5 | 13.7 | 13.5 |
| 0.474345 | -0.12044 | 0.010883 | -0.53206 | 11.1 | 11 | 11 | 10.8 | 10.3 | 10.7 | 10.4 | 10.8 | 10.3 |
| 0.47438 | -0.22581 | 0.23923 | -0.38281 | 9.67 | 8.74 | 8.72 | 8.44 | 9.1 | 9.11 | 8.93 | 8.64 | 8.4 |
| 0.474711 | 0.093279 | 0.734744 | 0.043623 | 14.7 | 15 | 15 | 15 | 14.6 | 14.9 | 15 | 15 | 14.9 |
| 0.476152 | -0.05298 | 0.904395 | 0.008783 | 15.4 | 15.5 | 15.5 | 15.5 | 15.5 | 15.6 | 15.5 | 15.5 | 15.5 |
| 0.476215 | -0.21148 | 0.394389 | -0.25471 | 11 | 10.7 | 10.7 | 10.9 | 10.9 | 10.3 | 9.92 | 11 | 10.6 |
| 0.479 | -0.09946 | 0.508667 | -0.09265 | 13.1 | 13.3 | 13.5 | 13.5 | 13.1 | 13.3 | 13.1 | 13.2 | 13.4 |
| 0.479866 | 0.14142 | 0.564862 | 0.114566 | 13.6 | 14 | 14 | 14.1 | 13.5 | 14 | 13.8 | 14 | 14.1 |
| 0.481637 | -0.08079 | 0.184373 | -0.15922 | 12 | 12 | 12 | 11.9 | 11.7 | 12.1 | 11.9 | 11.7 | 11.9 |
| 0.481197 | 0.106593 | 0.79555 | 0.038645 | 13.2 | 12.7 | 12.9 | 12.7 | 13 | 12.8 | 13 | 13 | 12.8 |
| 0.480769 | -0.10839 | 0.439102 | -0.11936 | 11.4 | 11.1 | 11 | 11.1 | 11.3 | 11 | 11 | 11.2 | 10.9 |
| 0.483357 | 0.345834 | 0.240729 | 0.596717 | 10.1 | 10.7 | 11.2 | 11.4 | 9.84 | 11.5 | 11 | 11.2 | 11.6 |
| 0.485209 | 0.129684 | 0.286797 | -0.20236 | 11.6 | 11.5 | 11.2 | 10.9 | 11.1 | 11.2 | 10.9 | 11.4 | 11.4 |
| 0.485134 | -0.06522 | 0.10385 | -0.16381 | 13.7 | 13.8 | 13.8 | 13.5 | 13.8 | 13.6 | 13.7 | 13.6 | 13.5 |
| 0.486396 | 0.078552 | 0.870453 | -0.01812 | 13.1 | 12.9 | 12.8 | 12.7 | 13 | 12.7 | 13 | 12.9 | 12.9 |
| 0.487312 | 0.475063 | 0.423748 | 0.550058 | 9.99 | 11.9 | 11.9 | 12 | 10.1 | 12 | 11.7 | 11.9 | 11.9 |
| 0.488943 | 0.226034 | 0.244904 | 0.39139 | 8.38 | 8.55 | 8.64 | 8.53 | 8.61 | 8.93 | 8.93 | 9.61 | 8.21 |
| 0.48856 | -0.1319 | 0.308811 | -0.19753 | 11.4 | 11.2 | 10.8 | 10.9 | 11.3 | 11 | 11.1 | 11 | 10.7 |
| 0.488323 | 0.07044 | 0.318049 | 0.103304 | 12.2 | 12.3 | 12.2 | 12.3 | 12.4 | 12.1 | 12.4 | 12.2 | 12.3 |
| 0.490055 | 0.229392 | 0.688059 | 0.132015 | 10.7 | 10.9 | 11 | 11.3 | 9.94 | 11 | 10.9 | 11.1 | 10.9 |
| 0.491932 | -0.08355 | 0.058886 | -0.25604 | 12.4 | 12.8 | 12.6 | 12.6 | 12.3 | 12.4 | 12.3 | 12.4 | 12.4 |
| 0.491293 | 0.48765 | 0.789877 | 0.186245 | 7.11 | 9.15 | 9.44 | 8.47 | 7.24 | 9.08 | 8.48 | 8.56 | 9.22 |
| 0.490804 | -0.0441 | 0.293087 | -0.06876 | 15.3 | 15.3 | 15.2 | 15.3 | 15.3 | 15.3 | 15.2 | 15.2 | 15.2 |
| 0.491937 | -0.07373 | 0.440351 | -0.08313 | 17.9 | 17.6 | 17.7 | 17.7 | 17.9 | 17.7 | 17.7 | 17.7 | 17.7 |
| 0.494119 | 0.259703 | 0.987704 | 0.005762 | 10.2 | 9.97 | 10.1 | 9.08 | 10.5 | 9.93 | 9.98 | 10.6 | 9.75 |

| | | | | | | | | | | | | |
|---|---|---|---|---|---|---|---|---|---|---|---|---|
| 0.494437 | 0.172284 | 0.658402 | 0.110458 | 9.96 | 9.81 | 9.81 | 9.84 | 9.93 | 9.62 | 10.5 | 9.97 | 9.43 |
| 0.495263 | 0.156472 | 0.544781 | 0.138488 | 10.1 | 10.1 | 10.5 | 10.4 | 9.84 | 10.4 | 10.1 | 10.3 | 10.7 |
| 0.496841 | -0.25282 | 0.143309 | -0.57719 | 9.61 | 10.1 | 10.2 | 10 | 10.1 | 8.78 | 9.63 | 9.24 | 9.32 |
| 0.49723 | 0.082745 | 0.92867 | -0.01075 | 13.1 | 13.2 | 12.9 | 12.9 | 12.9 | 13.2 | 13.1 | 13.2 | 12.9 |
| 0.498307 | 0.129866 | 0.617681 | 0.095048 | 10.5 | 11.2 | 11 | 10.9 | 10.9 | 10.7 | 11.1 | 11.1 | 10.9 |
| 0.500137 | 0.159016 | 0.734387 | 0.079101 | 13.9 | 14.6 | 14.5 | 14.6 | 14 | 14.4 | 14.2 | 14.5 | 14.6 |
| 0.499714 | -0.12756 | 0.49978 | -0.12754 | 10.6 | 11.3 | 11.1 | 11.1 | 10.9 | 11 | 10.7 | 10.9 | 10.9 |
| 0.502446 | 1.984169 | 0.365564 | 2.710905 | 7.81 | -0.2 | 7.96 | 8.61 | 0.17 | 9.02 | 7.39 | 8.15 | 8.23 |
| 0.502943 | 0.072547 | 0.199685 | -0.14473 | 11.8 | 11.8 | 11.7 | 11.4 | 11.6 | 11.7 | 11.8 | 11.6 | 11.5 |
| 0.504013 | 0.140805 | 0.510017 | -0.13877 | 13.1 | 12.7 | 12.5 | 12.2 | 12.8 | 12.4 | 12.8 | 12.5 | 12.6 |
| 0.50461 | -0.14666 | 0.933899 | -0.01796 | 12.6 | 12.1 | 12.2 | 12.1 | 12.8 | 12.6 | 12.4 | 12.4 | 12.1 |
| 0.505313 | -0.07854 | 0.45394 | -0.08865 | 13.9 | 13.6 | 13.7 | 13.7 | 13.9 | 13.6 | 13.8 | 13.6 | 13.5 |
| 0.507273 | 0.105708 | 0.437851 | -0.12437 | 10.2 | 10.6 | 10.4 | 10.2 | 9.99 | 10.3 | 10.5 | 10.2 | 10.1 |
| 0.510482 | -0.05275 | 0.02863 | -0.20483 | 13.7 | 13.5 | 13.6 | 13.4 | 13.5 | 13.5 | 13.4 | 13.4 | 13.4 |
| 0.514776 | 0.046539 | 0.340484 | -0.06924 | 15.8 | 15.9 | 15.9 | 15.8 | 15.8 | 15.7 | 15.8 | 15.7 | 15.8 |
| 0.512925 | -0.06555 | 0.09522 | -0.18144 | 16 | 16.2 | 16.3 | 16.1 | 15.9 | 16.1 | 15.9 | 16 | 16.1 |
| 0.512339 | 0.084808 | 0.705652 | -0.04842 | 11.3 | 11.5 | 11 | 11.2 | 11.1 | 11.2 | 11.1 | 11.2 | 11.3 |
| 0.509629 | -0.10661 | 0.737609 | 0.053577 | 12.2 | 11.6 | 11.8 | 11.9 | 12.2 | 12.1 | 12 | 11.8 | 12 |
| 0.508484 | 0.141218 | 0.124096 | 0.351157 | 12.1 | 11.5 | 11.4 | 11.6 | 12.1 | 11.9 | 11.9 | 12.2 | 12 |
| 0.51221 | 0.330131 | 0.767189 | -0.14741 | 9.26 | 8.77 | 9.07 | 9.17 | 9.21 | 7.3 | 8.94 | 9.15 | 8.58 |
| 0.509967 | -0.134 | 0.790915 | 0.053257 | 11.7 | 11.2 | 10.9 | 11.5 | 11.2 | 11.6 | 11.3 | 11.3 | 11.3 |
| 0.510029 | 0.066611 | 0.810506 | -0.02394 | 14.5 | 14.4 | 14.3 | 14.2 | 14.5 | 14.3 | 14.4 | 14.3 | 14.4 |
| 0.512585 | 0.079436 | 0.951293 | -0.0073 | 14.4 | 14.7 | 14.7 | 14.6 | 14.4 | 14.5 | 14.5 | 14.6 | 14.7 |
| 0.513491 | 0.106667 | 0.981565 | -0.00372 | 12.9 | 13.1 | 13.2 | 13.3 | 12.7 | 12.9 | 13.1 | 13.1 | 13 |
| 0.508913 | -0.07844 | 0.203085 | -0.15743 | 14.3 | 14.1 | 14 | 14.1 | 14.2 | 13.8 | 14.1 | 13.9 | 13.9 |
| 0.514986 | -0.09209 | 0.257142 | -0.16507 | 13.5 | 13.1 | 13.3 | 13.1 | 13.3 | 13.5 | 13.2 | 13.1 | 13.2 |
| 0.515115 | 0.12242 | 0.268684 | 0.213817 | 12 | 11.7 | 11.9 | 11.7 | 12.2 | 12 | 12.4 | 12 | 11.8 |
| 0.513283 | 0.054181 | 0.861883 | 0.014227 | 16.7 | 16.9 | 16.9 | 16.9 | 16.7 | 16.9 | 16.9 | 16.9 | 16.9 |
| 0.514254 | 0.126395 | 0.387242 | 0.169408 | 13.5 | 13.8 | 14 | 14.1 | 13.5 | 13.9 | 13.9 | 14 | 14 |
| 0.510532 | 1.885399 | 0.395542 | 2.458486 | 7.82 | 8.01 | 0.25 | 9.25 | 0.4 | 8.16 | 6.74 | 8.11 | 8.62 |
| 0.516509 | -0.05636 | 0.963411 | 0.003929 | 12.8 | 12.6 | 12.7 | 12.6 | 12.8 | 12.8 | 12.7 | 12.8 | 12.6 |
| 0.517607 | 0.123999 | 0.241671 | 0.231833 | 9.84 | 10.3 | 10.3 | 10.6 | 10.2 | 9.99 | 10.2 | 10.4 | 10.5 |
| 0.517191 | -0.15915 | 0.26125 | -0.28415 | 10.4 | 11.3 | 11.1 | 10.7 | 10.6 | 11.1 | 10.6 | 10.5 | 10.8 |
| 0.516873 | -0.0601 | 0.269671 | -0.10519 | 17 | 16.8 | 16.8 | 16.8 | 16.9 | 16.8 | 16.8 | 16.8 | 16.7 |
| 0.518472 | 0.084345 | 0.786111 | 0.035037 | 16.1 | 15.7 | 15.8 | 15.7 | 15.9 | 15.8 | 16 | 15.9 | 15.8 |
| 0.5191 | 0.129154 | 0.54949 | -0.11966 | 11 | 11.3 | 11.6 | 11.3 | 11.1 | 10.7 | 11.3 | 11.2 | 11 |
| 0.520664 | -0.15507 | 0.460956 | -0.17879 | 11.8 | 11.7 | 11.1 | 11.6 | 11.8 | 11.1 | 11.5 | 11.1 | 11.4 |
| 0.5222 | -0.08291 | 0.224475 | -0.16325 | 14.1 | 13.9 | 13.7 | 13.6 | 13.9 | 13.9 | 13.8 | 13.7 | 13.7 |
| 0.522576 | -0.33953 | 0.846554 | -0.10148 | 9.83 | 10.2 | 10.1 | 10.4 | 9.46 | 10.9 | 8.84 | 10.4 | 10.5 |
| 0.523224 | 0.321954 | 0.610744 | 0.255418 | 11.2 | 12.8 | 12.7 | 12.7 | 11.5 | 12.4 | 12.2 | 12.6 | 12.7 |
| 0.52321 | -0.09416 | 0.488154 | -0.1025 | 13.6 | 13.4 | 13.2 | 13.3 | 13.6 | 13.3 | 13.2 | 13.4 | 13.4 |
| 0.524475 | -0.08503 | 0.258458 | -0.15556 | 13 | 12.9 | 12.7 | 12.6 | 12.8 | 12.9 | 12.6 | 12.6 | 12.9 |
| 0.527033 | 0.097919 | 0.756143 | 0.047582 | 15.2 | 14.7 | 14.8 | 14.8 | 15.1 | 14.7 | 15 | 14.9 | 15 |
| 0.527258 | 0.141599 | 0.744629 | 0.072235 | 11.7 | 12.2 | 12.3 | 11.7 | 11.9 | 12.4 | 12.3 | 12.1 | 12.1 |
| 0.526662 | -0.1803 | 0.379751 | -0.25332 | 9.26 | 9 | 9.56 | 8.62 | 9.6 | 9.37 | 9.24 | 8.72 | 9.09 |
| 0.526895 | -0.06753 | 0.586225 | -0.05789 | 15.1 | 15.4 | 15.2 | 15.2 | 15.2 | 15.3 | 15.1 | 15 | 15.3 |
| 0.528009 | 0.127948 | 0.865726 | -0.03386 | 11.1 | 10.9 | 11.5 | 11.3 | 10.8 | 11 | 10.9 | 11.3 | 11.2 |
| 0.530491 | 0.076383 | 0.879514 | -0.01823 | 13.1 | 12.9 | 12.8 | 12.8 | 12.9 | 12.8 | 13.1 | 12.8 | 12.9 |
| 0.530254 | -0.08286 | 0.283942 | -0.14516 | 9.14 | 9.37 | 9.42 | 9.12 | 9.48 | 9.14 | 9.05 | 9.2 | 9.24 |
| 0.531545 | -0.07635 | 0.849379 | -0.0229 | 12.9 | 12.9 | 13.2 | 13 | 13 | 13.2 | 12.9 | 13 | 13.1 |

| | | | | | | | | | | | | |
|---|---|---|---|---|---|---|---|---|---|---|---|---|
| 0.532864 | -0.1237 | 0.029836 | -0.50249 | 12.6 | 12.6 | 12.8 | 12.5 | 12.3 | 12 | 12.4 | 11.8 | 12.3 |
| 0.532671 | -0.07981 | 0.822811 | 0.028317 | 14.2 | 14.3 | 14.6 | 14.6 | 14.3 | 14.5 | 14.3 | 14.4 | 14.4 |
| 0.534381 | 0.052455 | 0.702693 | -0.03196 | 12.7 | 12.9 | 12.8 | 12.7 | 12.7 | 12.7 | 12.9 | 12.7 | 12.8 |
| 0.535279 | -0.11946 | 0.741696 | -0.06292 | 11.6 | 12.1 | 12.2 | 12.2 | 11.7 | 12.1 | 12.1 | 11.8 | 11.9 |
| 0.535618 | -0.10182 | 0.510736 | -0.10828 | 12.6 | 12.7 | 12.5 | 12.6 | 12.9 | 12.3 | 12.7 | 12.4 | 12.4 |
| 0.53677 | 0.07984 | 0.419542 | -0.1053 | 11.5 | 11.8 | 11.7 | 11.5 | 11.5 | 11.4 | 11.6 | 11.3 | 11.7 |
| 0.536809 | 0.229868 | 0.535714 | 0.230503 | 11.2 | 12.4 | 12.4 | 12.4 | 11.5 | 12.2 | 12 | 12.3 | 12.4 |
| 0.537712 | -0.12789 | 0.941617 | 0.015007 | 11.9 | 11.7 | 11.2 | 11.6 | 12 | 11.6 | 11.7 | 11.7 | 11.5 |
| 0.5389 | 0.084167 | 0.805502 | 0.033368 | 11.8 | 11.6 | 12 | 11.7 | 11.9 | 11.6 | 11.8 | 11.8 | 11.9 |
| 0.539758 | -0.18626 | 0.44697 | -0.2326 | 12.3 | 11.7 | 11.5 | 11.3 | 12.2 | 11.9 | 12 | 11.4 | 11.4 |
| 0.543168 | 0.057304 | 0.186472 | -0.13055 | 14.1 | 14.3 | 14.1 | 13.8 | 14 | 14.1 | 14 | 14 | 14.1 |
| 0.544097 | -0.226 | 0.954252 | 0.02111 | 10.1 | 9.78 | 8.97 | 9.28 | 10.1 | 10.3 | 9.81 | 9.34 | 9.81 |
| 0.546258 | 0.265078 | 0.640134 | 0.204379 | 10.4 | 11.6 | 11.6 | 11.5 | 10.4 | 11.4 | 11.1 | 11.3 | 11.7 |
| 0.547791 | 0.262534 | 0.77701 | 0.122531 | 11.4 | 11.9 | 12.2 | 12.1 | 10.6 | 12.2 | 11.7 | 11.9 | 12.1 |
| 0.548564 | -0.06898 | 0.726548 | -0.03989 | 14.5 | 14.8 | 14.8 | 14.8 | 14.6 | 14.7 | 14.6 | 14.6 | 14.7 |
| 0.551125 | -0.0942 | 0.17238 | -0.22717 | 11.6 | 11.7 | 11.6 | 11.3 | 11.8 | 11.5 | 11.2 | 11.5 | 11.6 |
| 0.551126 | 0.094851 | 0.94673 | -0.01051 | 13.2 | 13.6 | 13.5 | 13.5 | 13 | 13.5 | 13.4 | 13.3 | 13.6 |
| 0.55308 | -0.04824 | 0.665231 | 0.034994 | 14.4 | 14.5 | 14.4 | 14.6 | 14.5 | 14.4 | 14.4 | 14.5 | 14.5 |
| 0.552721 | 0.105514 | 0.303816 | 0.187211 | 10.7 | 11.3 | 10.9 | 11.2 | 10.8 | 11.2 | 11.2 | 11.1 | 11.3 |
| 0.552327 | -0.04877 | 0.838518 | -0.01655 | 13.1 | 13.1 | 13 | 13 | 13.1 | 13.2 | 13 | 13 | 13.1 |
| 0.553001 | -0.14158 | 0.702708 | -0.09045 | 11.8 | 12.4 | 12.5 | 12.4 | 12.1 | 12.4 | 11.8 | 12.4 | 12.2 |
| 0.556132 | -0.11569 | 0.703589 | -0.07427 | 11.3 | 12 | 11.8 | 11.9 | 11.5 | 11.8 | 11.5 | 11.6 | 11.7 |
| 0.556315 | 0.092376 | 0.540487 | 0.096199 | 10.2 | 10.6 | 10 | 10.2 | 10.4 | 10.2 | 10.3 | 10.5 | 10.3 |
| 0.557513 | -0.09912 | 0.852132 | 0.031171 | 13.5 | 13.1 | 13.1 | 13.3 | 13.6 | 13.3 | 13.6 | 13.2 | 13.1 |
| 0.559555 | 0.107783 | 0.515686 | -0.1204 | 12.3 | 12.2 | 12.2 | 12.4 | 11.6 | 12.1 | 12.3 | 12.1 | 12.1 |
| 0.559861 | 0.204353 | 0.665531 | -0.15073 | 8.98 | 9.3 | 9.34 | 9.46 | 8.34 | 8.76 | 9.54 | 8.49 | 9.15 |
| 0.558948 | -0.10259 | 0.328149 | -0.17531 | 12.9 | 12.7 | 12.4 | 12.5 | 12.8 | 12.3 | 12.6 | 12.5 | 12.3 |
| 0.55849 | 0.231313 | 0.794124 | 0.102217 | 10 | 10.3 | 10.1 | 10.4 | 9.13 | 10.5 | 10.6 | 10.4 | 9.71 |
| 0.562487 | 0.04067 | 0.628096 | -0.0339 | 14.7 | 14.8 | 14.7 | 14.6 | 14.7 | 14.7 | 14.7 | 14.7 | 14.7 |
| 0.56308 | -0.0609 | 0.242064 | -0.12766 | 12.6 | 12.6 | 12.4 | 12.4 | 12.5 | 12.5 | 12.5 | 12.6 | 12.3 |
| 0.563212 | -0.05704 | 0.9813 | 0.002287 | 12.8 | 12.6 | 12.6 | 12.6 | 12.7 | 12.8 | 12.6 | 12.6 | 12.7 |
| 0.563021 | -0.06953 | 0.748714 | -0.03821 | 14.9 | 14.7 | 14.6 | 14.7 | 14.9 | 14.7 | 14.7 | 14.8 | 14.6 |
| 0.56104 | -0.04236 | 0.418232 | -0.05963 | 15.8 | 15.8 | 15.8 | 15.8 | 15.8 | 15.7 | 15.7 | 15.8 | 15.7 |
| 0.563022 | 1.605529 | 0.638206 | 1.300297 | 0.43 | 8.05 | 9.01 | 7.54 | 0.52 | 8.51 | 6.86 | 6.72 | 7.81 |
| 0.564844 | -0.08329 | 0.620551 | -0.07142 | 14.1 | 13.8 | 13.7 | 13.8 | 14 | 13.8 | 13.8 | 13.9 | 13.7 |
| 0.565565 | -0.13908 | 0.176664 | -0.34433 | 10.4 | 10.5 | 10.3 | 9.89 | 9.86 | 10.8 | 10.1 | 10 | 9.98 |
| 0.567855 | 0.048987 | 0.3213 | -0.08699 | 14.2 | 14.1 | 14.2 | 14 | 14.2 | 13.9 | 14.1 | 14.1 | 14 |
| 0.567587 | 0.063107 | 0.521396 | 0.070992 | 14.4 | 14.2 | 14.2 | 14.3 | 14.4 | 14.1 | 14.4 | 14.3 | 14.3 |
| 0.569959 | -0.08502 | 0.226059 | -0.18841 | 11.8 | 12 | 11.9 | 11.8 | 11.9 | 11.6 | 12 | 11.6 | 11.5 |
| 0.570241 | -0.09975 | 0.398657 | -0.15029 | 11.1 | 11.1 | 11.2 | 10.8 | 11.4 | 10.9 | 10.7 | 11.1 | 11.1 |
| 0.5717 | 0.144058 | 0.59092 | 0.13677 | 10.7 | 11.4 | 11.4 | 11.2 | 10.7 | 11.6 | 11.3 | 11.4 | 11.1 |
| 0.572291 | 0.124128 | 0.715214 | 0.079702 | 12.7 | 13.5 | 13.2 | 13.3 | 12.8 | 13.2 | 13.1 | 13.3 | 13.3 |
| 0.575427 | 0.071545 | 0.256023 | -0.15003 | 12.6 | 12.7 | 12.7 | 12.5 | 12.2 | 12.6 | 12.5 | 12.4 | 12.6 |
| 0.576026 | 0.22328 | 0.100864 | 0.711583 | 8.7 | 9.53 | 9.44 | 10.2 | 10.1 | 8.88 | 10.3 | 9.8 | 9.67 |
| 0.575947 | -0.05749 | 0.122635 | -0.17049 | 14.8 | 14.6 | 14.6 | 14.4 | 14.7 | 14.6 | 14.6 | 14.5 | 14.5 |
| 0.576708 | -0.11141 | 0.973425 | -0.00658 | 13.5 | 12.8 | 13 | 12.9 | 13.3 | 13.3 | 13.2 | 12.9 | 13.2 |
| 0.579787 | -0.05326 | 0.513018 | -0.06318 | 13.7 | 13.5 | 13.4 | 13.4 | 13.6 | 13.6 | 13.5 | 13.5 | 13.4 |
| 0.582541 | 0.147676 | 0.597439 | -0.14176 | 13 | 13.6 | 13.5 | 13.5 | 12.6 | 13.1 | 13.1 | 13.2 | 13.4 |
| 0.58194 | -0.09012 | 0.159043 | -0.24429 | 13.2 | 12.7 | 13 | 12.9 | 13 | 12.5 | 12.7 | 12.7 | 12.8 |
| 0.581092 | -0.11923 | 0.27607 | -0.24255 | 11 | 10.9 | 11 | 10.8 | 10.7 | 11 | 10.9 | 10.2 | 11.1 |

| | | | | | | | | | | | | |
|---|---|---|---|---|---|---|---|---|---|---|---|---|
| 0.582194 | 0.131159 | 0.826066 | 0.051938 | 15.3 | 15.9 | 15.8 | 15.9 | 15.1 | 15.7 | 15.5 | 15.8 | 15.8 |
| 0.582065 | 0.124009 | 0.667296 | 0.096469 | 10.6 | 10.4 | 10.4 | 10.4 | 10.7 | 10.3 | 10.7 | 10.9 | 10.1 |
| 0.585645 | -0.10532 | 0.367972 | -0.17699 | 13.2 | 12.7 | 12.4 | 12.6 | 12.9 | 12.6 | 12.5 | 12.6 | 12.6 |
| 0.586711 | 0.082272 | 0.468616 | -0.11054 | 9.55 | 9.56 | 9.34 | 9.11 | 9.27 | 9.5 | 9.62 | 9.18 | 9.32 |
| 0.586719 | 0.093085 | 0.7259 | 0.059684 | 12.1 | 12.2 | 11.9 | 12 | 11.8 | 12.2 | 11.8 | 12.3 | 12.2 |
| 0.587248 | -0.07888 | 0.424178 | -0.1175 | 15.5 | 15.1 | 15.1 | 15.1 | 15.4 | 15.1 | 15.3 | 15.2 | 15 |
| 0.587988 | -0.0355 | 0.432456 | -0.05201 | 16.4 | 16.4 | 16.3 | 16.4 | 16.4 | 16.3 | 16.3 | 16.3 | 16.4 |
| 0.590713 | 0.069264 | 0.06601 | 0.263806 | 10.9 | 11.2 | 10.7 | 11 | 11.2 | 11.2 | 11.3 | 11.1 | 11.2 |
| 0.590632 | -0.06766 | 0.285772 | -0.13822 | 13.6 | 13.3 | 13.6 | 13.5 | 13.5 | 13.3 | 13.5 | 13.3 | 13.3 |
| 0.593381 | -0.0759 | 0.049085 | -0.31737 | 11.8 | 11.7 | 11.7 | 11.3 | 11.6 | 11.6 | 11.2 | 11.6 | 11.5 |
| 0.593219 | -0.06498 | 0.31286 | -0.12589 | 13.8 | 14 | 14.1 | 14 | 13.7 | 14 | 13.8 | 13.9 | 13.9 |
| 0.59367 | 0.113054 | 0.811226 | 0.050223 | 12.7 | 13.3 | 13.4 | 13.2 | 12.7 | 13.1 | 13.1 | 13.1 | 13.3 |
| 0.593106 | -0.1339 | 0.544523 | -0.15228 | 9.45 | 9.7 | 9.64 | 9.64 | 9.67 | 9.43 | 9.74 | 8.83 | 9.77 |
| 0.594394 | -0.09967 | 0.846992 | -0.03582 | 11.5 | 12.1 | 11.8 | 12.1 | 11.8 | 11.6 | 11.8 | 11.6 | 11.8 |
| 0.596345 | 0.074156 | 0.90596 | -0.01639 | 17.7 | 17.4 | 17.4 | 17.3 | 17.7 | 17.3 | 17.6 | 17.5 | 17.4 |
| 0.596951 | -0.07617 | 0.736771 | 0.048144 | 14.5 | 14.1 | 14.1 | 14.2 | 14.3 | 14.5 | 14.4 | 14.2 | 14.1 |
| 0.600584 | 0.086588 | 0.949234 | -0.01043 | 13.3 | 13.6 | 13.6 | 13.5 | 13 | 13.7 | 13.5 | 13.5 | 13.5 |
| 0.601481 | -0.09123 | 0.290269 | -0.19014 | 11.4 | 11 | 11.1 | 10.8 | 11.4 | 11.1 | 11 | 11 | 11 |
| 0.600215 | -0.11887 | 0.409841 | -0.18953 | 11.3 | 10.9 | 10.7 | 10.8 | 11.3 | 10.6 | 10.9 | 10.8 | 10.6 |
| 0.601165 | 0.043692 | 0.735322 | 0.028105 | 12.2 | 12.1 | 12.2 | 12.1 | 12.2 | 12.2 | 12.3 | 12.2 | 12.1 |
| 0.601475 | -0.1097 | 0.518345 | -0.13633 | 15.2 | 15.8 | 15.7 | 15.8 | 15.2 | 15.6 | 15.4 | 15.4 | 15.5 |
| 0.603018 | 0.142283 | 0.566138 | 0.157283 | 12.7 | 13.4 | 13.4 | 13.5 | 12.7 | 13.4 | 13.4 | 13.2 | 13.4 |
| 0.603886 | 0.053554 | 0.021557 | 0.283646 | 11.5 | 11.7 | 11.5 | 11.6 | 11.8 | 11.9 | 11.8 | 11.9 | 11.8 |
| 0.606599 | 0.147167 | 0.948323 | -0.01836 | 9.97 | 10.9 | 10.3 | 10.4 | 10.4 | 9.87 | 10.1 | 10.4 | 10.7 |
| 0.606541 | -0.10667 | 0.340046 | -0.202 | 13.3 | 13.5 | 13.5 | 13.6 | 12.9 | 13.6 | 13.1 | 13.1 | 13.4 |
| 0.606754 | 0.060274 | 0.809309 | 0.028062 | 13.1 | 12.9 | 12.8 | 12.9 | 13 | 12.7 | 13 | 12.9 | 12.8 |
| 0.606531 | 1.469363 | 0.647006 | 1.303983 | 0.41 | 8.79 | 8.56 | 8.04 | 0.55 | 8.67 | 7.12 | 6.41 | 8.15 |
| 0.608176 | 0.10283 | 0.421152 | 0.163469 | 11.8 | 12.5 | 12.1 | 12.4 | 12.1 | 12 | 12.1 | 12.5 | 12.2 |
| 0.608794 | 0.09778 | 0.419175 | -0.15641 | 11.3 | 11.4 | 11.2 | 11.2 | 11.3 | 10.7 | 11.4 | 11 | 11 |
| 0.611737 | 0.037252 | 0.372701 | -0.06663 | 14 | 14.1 | 14 | 14 | 14 | 13.8 | 14 | 13.9 | 14 |
| 0.616191 | 0.326211 | 0.944635 | -0.04482 | 9.49 | 7.7 | 9.74 | 7.7 | 9.68 | 8.43 | 8.94 | 9.11 | 8.75 |
| 0.614105 | 0.110796 | 0.996906 | -0.00084 | 11.5 | 12.2 | 12.5 | 11.9 | 12 | 11.9 | 11.9 | 12.1 | 12.1 |
| 0.616801 | 0.085809 | 0.968617 | 0.00669 | 10.7 | 10.2 | 10.5 | 10.2 | 10.7 | 10.2 | 10.5 | 10.5 | 10.3 |
| 0.613819 | 0.431167 | 0.375635 | 0.770785 | 9.8 | 12.6 | 12.6 | 12.5 | 10.8 | 12.7 | 12.1 | 12.3 | 12.9 |
| 0.612928 | -0.06338 | 0.384775 | -0.11075 | 16.1 | 15.8 | 15.8 | 15.8 | 16 | 15.7 | 15.8 | 15.7 | 15.8 |
| 0.61595 | 0.086258 | 0.72186 | 0.060958 | 10.3 | 10.5 | 10.8 | 10.4 | 10.7 | 10.4 | 10.7 | 10.8 | 10.3 |
| 0.61483 | -0.06962 | 0.521211 | -0.08923 | 15.1 | 14.9 | 14.9 | 14.9 | 15.2 | 14.7 | 14.9 | 14.9 | 14.8 |
| 0.61666 | -0.0713 | 0.550937 | -0.08525 | 14.6 | 14.2 | 14.3 | 14.2 | 14.6 | 14.3 | 14.4 | 14.3 | 14.2 |
| 0.614465 | 0.144675 | 0.67021 | 0.122016 | 12.7 | 13.7 | 13.6 | 13.6 | 12.9 | 13.4 | 13.3 | 13.5 | 13.6 |
| 0.61705 | -0.11711 | 0.667011 | -0.10054 | 9.79 | 9.01 | 8.93 | 9.24 | 9.27 | 9.27 | 9.18 | 9.34 | 8.91 |
| 0.617793 | -0.09856 | 0.111913 | -0.33979 | 11.3 | 11.4 | 11.4 | 10.9 | 11.3 | 11.2 | 11.5 | 11 | 10.7 |
| 0.619549 | -0.15935 | 0.311178 | -0.3338 | 10.2 | 10.1 | 9.72 | 10.5 | 9.43 | 9.57 | 9.42 | 10.1 | 9.49 |
| 0.61973 | 0.054989 | 0.57948 | 0.061533 | 12.1 | 11.9 | 11.8 | 11.9 | 11.8 | 12 | 11.8 | 12 | 12.1 |
| 0.621697 | -0.10404 | 0.832366 | -0.04433 | 11.5 | 11.9 | 12.2 | 12.1 | 11.6 | 12.1 | 11.9 | 11.9 | 11.8 |
| 0.621684 | -0.05979 | 0.578953 | -0.06738 | 13.4 | 13.1 | 13.2 | 13.1 | 13.4 | 13.2 | 13.3 | 13.1 | 13.1 |
| 0.620967 | -0.0401 | 0.623311 | -0.03983 | 12.5 | 12.4 | 12.6 | 12.5 | 12.4 | 12.6 | 12.4 | 12.5 | 12.4 |
| 0.62255 | -0.13954 | 0.304571 | -0.29922 | 9.91 | 10.9 | 10.5 | 10.2 | 10.1 | 10.5 | 10.4 | 10.3 | 9.72 |
| 0.623443 | 0.092459 | 0.594646 | 0.100349 | 13.1 | 12.6 | 12.8 | 12.5 | 13 | 13 | 13 | 12.9 | 13 |
| 0.624494 | -0.07156 | 0.464318 | -0.10806 | 12.2 | 12.5 | 12.5 | 12.6 | 12.1 | 12.3 | 12.3 | 12.2 | 12.4 |
| 0.624055 | -0.04363 | 0.547602 | -0.05376 | 18.9 | 19 | 19 | 19 | 18.9 | 19 | 18.8 | 19 | 19 |

| | | | | | | | | | | | | |
|---|---|---|---|---|---|---|---|---|---|---|---|---|
| 0.627687 | -0.03918 | 0.107809 | -0.14094 | 17.3 | 17.1 | 17.1 | 17.1 | 17.1 | 17 | 17.1 | 17 | 17 |
| 0.629254 | 0.092604 | 0.186179 | -0.26713 | 12.6 | 12.4 | 12.3 | 12.2 | 12.3 | 11.6 | 12.1 | 12.1 | 12.2 |
| 0.633505 | 0.073517 | 0.494739 | -0.10614 | 12.5 | 12.7 | 12.8 | 12.6 | 12.3 | 12.6 | 12.4 | 12.5 | 12.8 |
| 0.629464 | 0.056871 | 0.758149 | -0.03613 | 13 | 13.1 | 13.2 | 13.2 | 12.8 | 13 | 13 | 13.1 | 13.1 |
| 0.630771 | 0.247457 | 0.315363 | 0.530815 | 11.4 | 12.9 | 13.1 | 13.3 | 11.9 | 13 | 12.8 | 13.1 | 13.1 |
| 0.632693 | 0.036351 | 0.337196 | 0.074738 | 13.4 | 13.4 | 13.4 | 13.4 | 13.5 | 13.4 | 13.5 | 13.5 | 13.4 |
| 0.633157 | 0.142039 | 0.954011 | 0.017033 | 11.6 | 12.4 | 12.4 | 12.3 | 11.5 | 12.3 | 12.2 | 12.1 | 12.3 |
| 0.630242 | -0.0563 | 0.38568 | -0.10327 | 13.6 | 13.5 | 13.5 | 13.6 | 13.6 | 13.3 | 13.4 | 13.3 | 13.6 |
| 0.632689 | -0.06551 | 0.901057 | -0.01692 | 13.6 | 14 | 14 | 14.2 | 13.8 | 13.8 | 13.9 | 13.8 | 13.9 |
| 0.63176 | -0.0534 | 0.43926 | -0.08727 | 12.9 | 12.7 | 12.8 | 12.9 | 12.5 | 12.8 | 12.7 | 12.8 | 12.6 |
| 0.632468 | 0.162417 | 0.448416 | 0.260482 | 8.93 | 9.85 | 9.43 | 9.02 | 9.82 | 9.66 | 10.2 | 9.45 | 9.35 |
| 0.637949 | 0.064784 | 0.497922 | -0.09407 | 13.2 | 13 | 12.9 | 12.7 | 13.2 | 12.8 | 12.9 | 12.9 | 13 |
| 0.638528 | 0.066064 | 0.55448 | -0.08349 | 12.9 | 12.8 | 12.5 | 12.6 | 12.5 | 12.8 | 12.8 | 12.7 | 12.5 |
| 0.637426 | 0.188522 | 0.190176 | 0.55165 | 8.15 | 9.02 | 9.65 | 9.74 | 9.08 | 9.1 | 9.09 | 9.55 | 9.84 |
| 0.638376 | -0.10569 | 0.875504 | 0.034999 | 10.6 | 10.2 | 10.3 | 10 | 10.6 | 10.8 | 10.4 | 10.5 | 10.3 |
| 0.639204 | -0.0337 | 0.308098 | -0.07533 | 15.1 | 15.1 | 15 | 15 | 15 | 15 | 14.9 | 15 | 15 |
| 0.637269 | -0.10129 | 0.925441 | 0.019959 | 10.7 | 11 | 11.2 | 11.4 | 10.6 | 11.3 | 10.8 | 11.1 | 11 |
| 0.6363 | -0.05814 | 0.963636 | 0.005564 | 12.9 | 12.6 | 12.8 | 12.8 | 13.1 | 12.7 | 12.8 | 12.7 | 12.8 |
| 0.639261 | 0.128267 | 0.528465 | 0.173536 | 11.7 | 12.7 | 12.5 | 12.6 | 12 | 12.4 | 12.5 | 12.3 | 12.5 |
| 0.640486 | -0.06687 | 0.959858 | -0.00715 | 10.5 | 10.5 | 10.3 | 10.6 | 10.5 | 10.4 | 10.1 | 10.4 | 10.7 |
| 0.641251 | 0.044785 | 0.851948 | 0.017828 | 14 | 13.9 | 13.9 | 13.9 | 14 | 13.8 | 14 | 14 | 13.8 |
| 0.642207 | 0.067786 | 0.907496 | -0.01684 | 11.3 | 11.7 | 11.6 | 11.2 | 11.4 | 11.7 | 11.6 | 11.5 | 11.4 |
| 0.643866 | 0.424974 | 0.797766 | 0.234375 | 7.08 | 8.45 | 8.32 | 6.28 | 7.06 | 9.94 | 8.03 | 7.76 | 8.77 |
| 0.648406 | 0.06122 | 0.919336 | -0.01351 | 12.2 | 12.5 | 12.5 | 12.5 | 12.3 | 12.1 | 12.3 | 12.5 | 12.3 |
| 0.648042 | 0.077287 | 0.960106 | 0.00841 | 14.3 | 14.6 | 14.6 | 14.6 | 14.2 | 14.5 | 14.3 | 14.6 | 14.7 |
| 0.647014 | -0.06889 | 0.822157 | -0.03363 | 12.6 | 13 | 13 | 13 | 12.7 | 13.1 | 12.8 | 12.8 | 12.9 |
| 0.647968 | 0.114139 | 0.624053 | 0.122652 | 11.5 | 12.2 | 12.3 | 12.1 | 11.7 | 12.3 | 12.2 | 12 | 12.3 |
| 0.652258 | -0.09465 | 0.51235 | -0.13868 | 12.6 | 13.2 | 13.3 | 13.1 | 12.8 | 13 | 12.6 | 12.9 | 13.1 |
| 0.653217 | 0.05408 | 0.798536 | 0.030581 | 12.5 | 12.3 | 12.2 | 12.4 | 12.5 | 12.1 | 12.3 | 12.4 | 12.4 |
| 0.659108 | 0.201857 | 0.317881 | -0.46973 | 10.2 | 8.48 | 10.3 | 8.79 | 8.94 | 9.19 | 8.9 | 9.31 | 9.31 |
| 0.66042 | -0.12953 | 0.767106 | 0.087008 | 10.6 | 11.6 | 11.6 | 11.6 | 11.3 | 11.5 | 11.2 | 11.2 | 11.7 |
| 0.662146 | 0.074033 | 0.450712 | -0.12944 | 11.7 | 11.5 | 11.4 | 11.3 | 11 | 11.6 | 11.2 | 11.5 | 11.5 |
| 0.663338 | -0.07897 | 0.447149 | -0.13973 | 14.3 | 14.8 | 14.8 | 14.6 | 14.3 | 14.7 | 14.4 | 14.4 | 14.6 |
| 0.663327 | 0.080657 | 0.918509 | 0.018842 | 10.9 | 11.4 | 11.2 | 11.3 | 11.1 | 11 | 10.9 | 11.4 | 11.4 |
| 0.666616 | 0.069386 | 0.282031 | 0.179116 | 12.2 | 12.4 | 12.7 | 12.8 | 12.3 | 12.6 | 12.7 | 12.6 | 12.6 |
| 0.667074 | 0.107167 | 0.89577 | 0.032453 | 13.3 | 13.9 | 13.9 | 13.9 | 13.3 | 13.7 | 13.5 | 13.6 | 14 |
| 0.666836 | 0.204853 | 0.557559 | 0.280579 | 8.97 | 10 | 10.5 | 9.89 | 9.23 | 10.6 | 10.1 | 9.89 | 10.4 |
| 0.669981 | 0.044166 | 0.866953 | -0.01727 | 14.7 | 14.8 | 14.8 | 14.9 | 14.6 | 14.7 | 14.6 | 14.8 | 14.9 |
| 0.672047 | 0.045891 | 0.368106 | -0.09968 | 11.9 | 12.2 | 12.2 | 12.1 | 11.9 | 11.8 | 12 | 11.9 | 12.1 |
| 0.67407 | 0.064412 | 0.914651 | -0.01632 | 11 | 10.9 | 11.4 | 11 | 10.8 | 11.2 | 11.1 | 11 | 11.1 |
| 0.673447 | 0.045872 | 0.851771 | 0.020242 | 12.1 | 12.3 | 12.4 | 12.2 | 12.2 | 12.3 | 12.1 | 12.4 | 12.3 |
| 0.67142 | -0.06471 | 0.619803 | -0.07586 | 16.2 | 15.8 | 15.6 | 15.7 | 16 | 15.8 | 15.8 | 15.8 | 15.8 |
| 0.673628 | 0.080309 | 0.630184 | 0.091975 | 11.2 | 11.5 | 11 | 11.4 | 11.4 | 11 | 11.1 | 11.6 | 11.3 |
| 0.671947 | 0.090061 | 0.629124 | 0.102899 | 12.7 | 13.1 | 13.1 | 13.2 | 12.6 | 13.1 | 12.8 | 13.1 | 13.3 |
| 0.674198 | 0.061591 | 0.697099 | 0.056978 | 12.8 | 12.8 | 12.4 | 12.6 | 12.9 | 12.6 | 12.9 | 12.7 | 12.7 |
| 0.673268 | 0.060839 | 0.663302 | 0.062839 | 12.6 | 12.2 | 12.2 | 12.3 | 12.4 | 12.2 | 12.3 | 12.6 | 12.2 |
| 0.674927 | -0.05483 | 0.814227 | -0.0306 | 12.2 | 12.3 | 12.4 | 12.3 | 12.3 | 12.5 | 12.2 | 12.1 | 12.5 |
| 0.675255 | 0.131521 | 0.698369 | 0.121552 | 10.4 | 9.34 | 10.2 | 10 | 9.58 | 10.3 | 9.99 | 9.98 | 10.4 |
| 0.680397 | 0.035274 | 0.031525 | -0.21559 | 15.6 | 15.6 | 15.5 | 15.3 | 15.4 | 15.1 | 15.3 | 15.3 | 15.3 |
| 0.679822 | -0.03849 | 0.488198 | 0.065325 | 14.4 | 14.3 | 14.2 | 14.4 | 14.3 | 14.4 | 14.4 | 14.5 | 14.2 |

| | | | | | | | | | | | | |
|---|---|---|---|---|---|---|---|---|---|---|---|---|
| 0.680161 | 0.155969 | 0.15535 | 0.572958 | 9.07 | 9.9 | 10.4 | 9.89 | 10.5 | 10.3 | 10.8 | 10.3 | 9.91 |
| 0.679918 | 0.084986 | 0.85664 | -0.03704 | 15.3 | 15.9 | 15.8 | 15.8 | 15.2 | 15.7 | 15.6 | 15.6 | 15.8 |
| 0.680335 | 0.042058 | 0.697388 | 0.039669 | 14.3 | 14.4 | 14.1 | 14.4 | 14.1 | 14.3 | 14.3 | 14.3 | 14.3 |
| 0.6813 | 0.087362 | 0.770737 | 0.061814 | 12 | 11.6 | 11.9 | 11.8 | 11.6 | 12.2 | 12.1 | 12.1 | 11.6 |
| 0.68582 | 0.048772 | 0.7024 | -0.04604 | 15.1 | 15.4 | 15.3 | 15.3 | 15 | 15.2 | 15.2 | 15.2 | 15.3 |
| 0.687418 | -0.05726 | 0.539612 | 0.087932 | 11.8 | 11.7 | 12.2 | 12 | 12.3 | 12 | 12.1 | 12 | 11.9 |
| 0.686642 | -0.07344 | 0.687417 | 0.073244 | 12.5 | 12.7 | 12.6 | 12.5 | 13.1 | 12.5 | 12.9 | 12.5 | 12.6 |
| 0.687023 | 0.124156 | 0.392433 | 0.268689 | 11.9 | 12.9 | 12.8 | 13.1 | 12.2 | 12.7 | 12.5 | 12.8 | 13 |
| 0.689605 | -0.04601 | 0.196065 | -0.15688 | 15.3 | 15 | 15.1 | 14.9 | 15.1 | 15.1 | 15 | 14.9 | 15 |
| 0.690205 | 0.078454 | 0.1682 | 0.287983 | 12.2 | 12 | 11.5 | 11.9 | 12.1 | 12.2 | 12.4 | 12.1 | 12 |
| 0.692873 | -0.07941 | 0.380467 | -0.18006 | 10.8 | 10.6 | 9.98 | 10.3 | 10.4 | 10.3 | 10.2 | 10.2 | 10.4 |
| 0.693747 | 0.067391 | 0.88452 | 0.024745 | 15.1 | 15.6 | 15.6 | 15.6 | 15.2 | 15.5 | 15.3 | 15.5 | 15.6 |
| 0.695504 | -0.03871 | 0.350872 | -0.09455 | 16.1 | 16.3 | 16.3 | 16.2 | 16.1 | 16.2 | 16 | 16.1 | 16.2 |
| 0.696163 | -0.04799 | 0.872479 | -0.01964 | 13.5 | 13.3 | 13.5 | 13.3 | 13.5 | 13.7 | 13.4 | 13.5 | 13.4 |
| 0.697826 | 0.127894 | 0.245064 | 0.398968 | 8.98 | 9.19 | 8.79 | 9.18 | 9.23 | 9.36 | 10.1 | 9.35 | 8.68 |
| 0.700246 | 0.072217 | 0.488579 | 0.131382 | 13.9 | 13.3 | 13.4 | 13.6 | 13.9 | 13.4 | 13.8 | 13.7 | 13.6 |
| 0.702758 | -0.05401 | 0.880557 | -0.02118 | 11.6 | 11.4 | 11.3 | 11.5 | 11.2 | 11.7 | 11.4 | 11.4 | 11.4 |
| 0.706667 | 0.172924 | 0.626904 | 0.224071 | 9.9 | 11.2 | 11 | 11.2 | 9.87 | 11.2 | 10.8 | 10.8 | 11.2 |
| 0.70722 | 0.13114 | 0.628051 | 0.169693 | 10.3 | 10.9 | 11.4 | 11.3 | 10.3 | 11.1 | 10.9 | 11.2 | 11.1 |
| 0.706987 | -0.10882 | 0.692206 | -0.11466 | 9.78 | 9.93 | 9.75 | 9.38 | 10.1 | 9.97 | 9.6 | 10.3 | 9.23 |
| 0.70801 | -0.03352 | 0.700887 | 0.034389 | 13.4 | 13.3 | 13.2 | 13.3 | 13.4 | 13.4 | 13.4 | 13.2 | 13.4 |
| 0.708526 | -0.03773 | 0.673804 | -0.04253 | 14.6 | 14.5 | 14.4 | 14.6 | 14.3 | 14.6 | 14.5 | 14.5 | 14.5 |
| 0.711518 | -0.05443 | 0.517686 | 0.096128 | 13.2 | 13.7 | 13.5 | 13.8 | 13.4 | 13.7 | 13.6 | 13.5 | 13.7 |
| 0.712345 | 0.114161 | 0.103416 | 0.550208 | 7.26 | 7.02 | 7.98 | 7.71 | 7.72 | 8.14 | 8.43 | 7.54 | 7.94 |
| 0.714978 | -0.03569 | 0.500134 | -0.06663 | 12.4 | 12.5 | 12.4 | 12.4 | 12.6 | 12.3 | 12.3 | 12.4 | 12.4 |
| 0.715233 | -0.08012 | 0.810107 | -0.05263 | 11.4 | 12.1 | 12 | 12 | 11.5 | 12.1 | 11.7 | 11.7 | 11.9 |
| 0.715923 | 0.05254 | 0.157353 | 0.217712 | 11.6 | 11.6 | 11.7 | 12 | 11.8 | 11.5 | 11.7 | 11.7 | 12.1 |
| 0.717291 | 0.048022 | 0.920934 | -0.01311 | 12.6 | 12.3 | 12.8 | 12.6 | 12.5 | 12.6 | 12.5 | 12.6 | 12.7 |
| 0.716777 | 0.058372 | 0.977358 | 0.004547 | 12.3 | 12.3 | 12.2 | 12 | 12.4 | 12.2 | 12.5 | 12.3 | 12 |
| 0.720027 | 0.081759 | 0.583657 | -0.12574 | 12 | 11.5 | 11.9 | 11.6 | 11.9 | 11.3 | 11.7 | 12 | 11.4 |
| 0.720551 | -0.0747 | 0.986979 | -0.00339 | 10.1 | 10.1 | 9.77 | 10.1 | 9.7 | 10.5 | 9.91 | 9.96 | 10.2 |
| 0.720528 | -0.04397 | 0.68514 | -0.04989 | 14.2 | 14.5 | 14.5 | 14.5 | 14.2 | 14.4 | 14.2 | 14.3 | 14.4 |
| 0.721382 | 0.0474 | 0.844978 | -0.02591 | 12 | 11.6 | 12 | 12 | 11.8 | 11.6 | 11.8 | 11.8 | 11.9 |
| 0.723092 | -0.05814 | 0.13431 | -0.26431 | 12.8 | 12.5 | 12.7 | 12.2 | 12.6 | 12.5 | 12.5 | 12.1 | 12.6 |
| 0.72588 | 0.061355 | 0.538119 | -0.10868 | 10.7 | 11.2 | 11.1 | 11.1 | 10.9 | 10.6 | 10.8 | 11 | 10.9 |
| 0.726782 | 0.045889 | 0.663525 | -0.05727 | 13.1 | 12.8 | 12.7 | 12.8 | 12.9 | 12.6 | 12.8 | 12.9 | 12.8 |
| 0.727006 | -0.03877 | 0.702383 | 0.042471 | 14.2 | 14.1 | 14.2 | 14.1 | 14.4 | 14.2 | 14.3 | 14.1 | 14.1 |
| 0.723914 | 0.053731 | 0.707664 | -0.05707 | 12.5 | 12.9 | 12.7 | 12.8 | 12.5 | 12.3 | 12.7 | 12.5 | 12.7 |
| 0.726256 | -0.05777 | 0.371626 | -0.15086 | 12.9 | 12.6 | 12.6 | 12.9 | 12.6 | 12.3 | 12.5 | 12.6 | 12.6 |
| 0.726217 | -0.04914 | 0.377261 | -0.12671 | 13.3 | 13 | 13.3 | 13 | 13.4 | 13 | 13.1 | 13 | 13.1 |
| 0.72587 | -0.09066 | 0.617934 | -0.12951 | 11.7 | 11.3 | 12 | 11.9 | 11.9 | 11.1 | 11.6 | 11.5 | 11.5 |
| 0.724263 | -0.10254 | 0.645175 | -0.13425 | 10.7 | 9.74 | 10.8 | 10.3 | 10.4 | 10.5 | 10.6 | 10.1 | 10.3 |
| 0.729274 | -0.12925 | 0.12349 | -0.62182 | 10.7 | 9.73 | 10.3 | 10.3 | 9.51 | 9.36 | 9.68 | 10.1 | 9.08 |
| 0.72874 | 0.062006 | 0.350236 | 0.17135 | 15.5 | 14.9 | 14.9 | 15 | 15.4 | 15.2 | 15.4 | 15.2 | 15.2 |
| 0.728479 | 0.072135 | 0.894279 | -0.02752 | 13.4 | 14 | 14.1 | 13.8 | 13.4 | 13.9 | 13.8 | 13.7 | 13.9 |
| 0.729951 | -0.02465 | 0.50022 | -0.04869 | 17.1 | 17 | 16.9 | 17 | 17 | 16.9 | 16.9 | 17 | 16.9 |
| 0.734199 | 0.0394 | 0.383626 | -0.10333 | 13.3 | 13.4 | 13.7 | 13.5 | 13.3 | 13.3 | 13.4 | 13.5 | 13.3 |
| 0.732927 | 0.038034 | 0.603989 | -0.05811 | 12.7 | 12.4 | 12.7 | 12.4 | 12.6 | 12.5 | 12.6 | 12.4 | 12.6 |
| 0.732196 | -0.05766 | 0.628034 | 0.081961 | 10.9 | 11.3 | 11.5 | 11.4 | 11.3 | 11.5 | 11.1 | 11.3 | 11.5 |
| 0.734399 | 0.075234 | 0.869053 | -0.03645 | 15 | 15.6 | 15.6 | 15.4 | 14.9 | 15.5 | 15.3 | 15.2 | 15.5 |

| | | | | | | | | | | | | |
|---|---|---|---|---|---|---|---|---|---|---|---|---|
| 0.731815 | -0.14208 | 0.882769 | 0.060937 | 8.73 | 9.31 | 9.17 | 9.31 | 9.77 | 8.74 | 9.26 | 9.81 | 8.32 |
| 0.73325 | -0.10353 | 0.476674 | -0.21907 | 10.4 | 11.2 | 11.4 | 11.1 | 10.4 | 11.2 | 10.7 | 10.9 | 10.8 |
| 0.734518 | 0.07906 | 0.677485 | 0.097149 | 14.2 | 14.9 | 14.8 | 14.8 | 14.2 | 14.8 | 14.6 | 14.7 | 14.8 |
| 0.736263 | -0.07936 | 0.794182 | 0.06138 | 11.7 | 11.1 | 11.2 | 11.6 | 11.7 | 11.1 | 11.6 | 11.4 | 11.1 |
| 0.738626 | 0.053971 | 0.160397 | -0.24206 | 11 | 11.2 | 10.7 | 10.8 | 10.7 | 10.5 | 10.5 | 10.9 | 10.8 |
| 0.738601 | 0.043911 | 0.518733 | 0.085794 | 14.1 | 13.6 | 13.7 | 13.8 | 14 | 13.8 | 14 | 13.9 | 13.9 |
| 0.739226 | -0.0771 | 0.933195 | 0.019349 | 12.1 | 13 | 12.8 | 12.8 | 12.5 | 12.9 | 12.6 | 12.6 | 12.7 |
| 0.740599 | -0.02527 | 0.466792 | -0.05634 | 14.7 | 14.8 | 14.9 | 14.7 | 14.8 | 14.7 | 14.7 | 14.8 | 14.7 |
| 0.740117 | -0.02651 | 0.625419 | -0.03918 | 15.7 | 15.6 | 15.6 | 15.5 | 15.7 | 15.6 | 15.6 | 15.6 | 15.6 |
| 0.74224 | 0.068252 | 0.158069 | -0.31258 | 11.8 | 11.3 | 11.4 | 11.2 | 11.3 | 10.7 | 11.1 | 11.4 | 10.9 |
| 0.742992 | -0.94289 | 0.26874 | -3.30386 | 9.08 | 8.49 | 8.1 | 8.3 | 9.13 | 1.17 | 8.55 | -0.6 | 7.8 |
| 0.74274 | 0.032618 | 0.481661 | 0.070848 | 14.5 | 14.7 | 14.7 | 14.7 | 14.6 | 14.8 | 14.7 | 14.8 | 14.6 |
| 0.744229 | 0.085371 | 0.891645 | 0.03554 | 11.4 | 12.3 | 12.2 | 12 | 11.6 | 12.2 | 11.7 | 12.2 | 12.1 |
| 0.747311 | -0.07401 | 0.033104 | 0.572567 | 9.8 | 9.1 | 9.38 | 10.3 | 9.69 | 10.2 | 9.97 | 10.2 | 9.82 |
| 0.748301 | -0.03587 | 0.321393 | -0.11423 | 12.3 | 11.9 | 12 | 12 | 12.1 | 12 | 11.9 | 12.1 | 11.9 |
| 0.748092 | 0.036291 | 0.422676 | 0.092278 | 11.9 | 12.1 | 12.3 | 12.1 | 12.1 | 12.1 | 12.3 | 12.1 | 12.1 |
| 0.749118 | 0.044772 | 0.549643 | -0.08449 | 12.6 | 12.7 | 12.6 | 12.5 | 12.3 | 12.7 | 12.7 | 12.4 | 12.4 |
| 0.751401 | -0.05942 | 0.241047 | 0.229681 | 11.4 | 11.3 | 11.8 | 12 | 11.6 | 11.7 | 12 | 11.8 | 11.4 |
| 0.756867 | -0.0328 | 0.005945 | -0.38251 | 12.5 | 12.3 | 12.3 | 12 | 12.2 | 11.9 | 12.1 | 11.9 | 11.9 |
| 0.756963 | 0.050913 | 0.204921 | -0.21965 | 15.1 | 14.7 | 15 | 14.7 | 14.9 | 14.4 | 14.7 | 14.7 | 14.7 |
| 0.75559 | 0.029769 | 0.218749 | 0.123463 | 14.1 | 13.9 | 14 | 14.1 | 14.2 | 14 | 14.1 | 14.2 | 14 |
| 0.753834 | 0.069783 | 0.576978 | -0.12502 | 11.4 | 11.9 | 11.8 | 11.7 | 11 | 11.7 | 11.4 | 11.6 | 11.7 |
| 0.755324 | 0.079809 | 0.437802 | 0.202148 | 11.2 | 10.7 | 10.5 | 10.7 | 11.4 | 10.8 | 11.2 | 10.9 | 10.9 |
| 0.756332 | -0.05339 | 0.494303 | -0.11909 | 11.2 | 10.8 | 10.6 | 10.6 | 10.9 | 11 | 10.7 | 10.8 | 10.7 |
| 0.754339 | -0.02967 | 0.922934 | -0.00914 | 14.4 | 14.1 | 14.2 | 14.2 | 14.4 | 14.2 | 14.2 | 14.3 | 14.2 |
| 0.754388 | -0.07417 | 0.857153 | -0.04258 | 11.6 | 11.9 | 11.3 | 11.8 | 11.9 | 11.1 | 11.6 | 11.5 | 11.6 |
| 0.758511 | -0.0458 | 0.507104 | -0.09998 | 14.3 | 13.9 | 14 | 13.9 | 14.3 | 13.9 | 13.9 | 14 | 13.9 |
| 0.759307 | -0.08712 | 0.068659 | -0.57928 | 10.7 | 9.97 | 9.64 | 9.45 | 9.98 | 9.43 | 9.64 | 9.37 | 9.58 |
| 0.759699 | 0.052715 | 0.659872 | 0.076102 | 11.3 | 11.1 | 10.9 | 11.3 | 11.3 | 10.7 | 11.2 | 11.1 | 11.2 |
| 0.761932 | -0.04309 | 0.722764 | 0.050522 | 10.5 | 10.7 | 10.5 | 10.6 | 10.9 | 10.5 | 10.5 | 10.6 | 10.8 |
| 0.762615 | 0.081235 | 0.66157 | 0.11813 | 10.9 | 11.2 | 10.8 | 10.8 | 11.5 | 10.8 | 11.4 | 11.3 | 10.6 |
| 0.767721 | 0.037428 | 0.283971 | -0.14078 | 11.4 | 11.6 | 11.7 | 11.5 | 11.5 | 11.2 | 11.3 | 11.4 | 11.6 |
| 0.764492 | 0.038379 | 0.811121 | -0.03058 | 13.8 | 14.2 | 13.9 | 13.9 | 13.8 | 14 | 13.9 | 14.1 | 13.8 |
| 0.765748 | 0.047303 | 0.838471 | -0.03231 | 12 | 11.6 | 11.6 | 11.4 | 11.9 | 11.6 | 11.6 | 11.7 | 11.8 |
| 0.765635 | -0.0206 | 0.449812 | -0.0531 | 14.6 | 14.7 | 14.7 | 14.6 | 14.7 | 14.6 | 14.7 | 14.6 | 14.5 |
| 0.764484 | -0.19593 | 0.504733 | -0.44152 | 10.2 | 9.78 | 9.42 | 10.4 | 10 | 8.29 | 9.92 | 9.92 | 8.27 |
| 0.767184 | 0.054423 | 0.981581 | -0.00423 | 12.6 | 13.3 | 13.2 | 13.1 | 12.7 | 13.1 | 13 | 13.1 | 13.1 |
| 0.767598 | 0.086878 | 0.919108 | 0.029779 | 13.5 | 14.3 | 14.1 | 14.3 | 13.4 | 14.1 | 13.7 | 13.9 | 14.3 |
| 0.76656 | -0.0271 | 0.655486 | -0.04087 | 13.6 | 13.5 | 13.5 | 13.6 | 13.5 | 13.4 | 13.4 | 13.4 | 13.6 |
| 0.763989 | 0.065146 | 0.658015 | 0.096389 | 12 | 11.6 | 11.4 | 11.9 | 11.8 | 11.4 | 11.7 | 12 | 11.6 |
| 0.77072 | -0.02305 | 0.904827 | -0.00944 | 14.4 | 14.4 | 14.4 | 14.4 | 14.3 | 14.5 | 14.4 | 14.4 | 14.4 |
| 0.772211 | 0.059893 | 0.488066 | -0.1454 | 11.8 | 11.4 | 11.6 | 11.1 | 11.4 | 11.7 | 11.8 | 11.4 | 11.2 |
| 0.772199 | -0.04535 | 0.386722 | -0.1387 | 12 | 11.7 | 11.9 | 12.1 | 11.5 | 11.9 | 11.9 | 11.8 | 11.7 |
| 0.773009 | -0.03549 | 0.382354 | -0.11001 | 11.4 | 11.4 | 11.3 | 11.6 | 11.3 | 11.1 | 11.3 | 11.2 | 11.3 |
| 0.778081 | 0.02979 | 0.90056 | -0.01318 | 15.3 | 15.2 | 15.1 | 15.2 | 15.3 | 15 | 15.3 | 15.2 | 15.1 |
| 0.778146 | 0.025139 | 0.959632 | 0.004505 | 12.3 | 12.3 | 12.2 | 12.2 | 12.4 | 12.2 | 12.2 | 12.4 | 12.3 |
| 0.779487 | -0.029 | 0.64777 | 0.047536 | 13 | 12.8 | 12.8 | 13.1 | 12.9 | 12.8 | 12.9 | 12.9 | 13 |
| 0.780625 | 0.026439 | 0.414804 | 0.078973 | 12.8 | 12.8 | 12.7 | 12.9 | 12.7 | 12.8 | 12.7 | 12.9 | 12.8 |
| 0.780868 | 0.098237 | 0.914176 | 0.037974 | 13.5 | 14.2 | 14.2 | 14.2 | 13.2 | 14.2 | 13.6 | 14.1 | 14.3 |
| 0.783503 | 0.050069 | 0.04897 | -0.40999 | 13 | 12.8 | 12.7 | 12.1 | 12.6 | 12.4 | 12.6 | 12.5 | 12.1 |

| | | | | | | | | | | | | |
|---|---|---|---|---|---|---|---|---|---|---|---|---|
| 0.78508 | -0.06361 | 0.13328 | 0.37754 | 9.82 | 9.14 | 9.44 | 9.61 | 10 | 10.1 | 10 | 10 | 9.49 |
| 0.787432 | -0.04557 | 0.435562 | 0.134173 | 11.3 | 11.1 | 11 | 11.6 | 11.2 | 11.1 | 11 | 11.5 | 11.4 |
| 0.781932 | -0.06718 | 0.236757 | -0.30043 | 11.6 | 11.1 | 11.7 | 11.5 | 11.3 | 10.9 | 11.6 | 10.9 | 11.1 |
| 0.786703 | 0.065849 | 0.545731 | -0.14848 | 11.3 | 11.7 | 11.4 | 11.4 | 10.7 | 11.7 | 11.5 | 11.2 | 11.4 |
| 0.785137 | 0.059028 | 0.551764 | -0.13008 | 9.91 | 9.78 | 10.1 | 10 | 9.95 | 9.24 | 9.64 | 9.97 | 9.78 |
| 0.788192 | -0.04992 | 0.29047 | -0.20356 | 11.5 | 11.9 | 11.8 | 11.9 | 11.2 | 11.7 | 11.6 | 11.4 | 11.6 |
| 0.788585 | -0.14048 | 0.352886 | -0.49999 | 8.33 | 9.07 | 9.21 | 9.55 | 8.36 | 7.62 | 8.8 | 8.58 | 7.74 |
| 0.783811 | -0.04418 | 0.506367 | -0.10835 | 17.5 | 18 | 17.9 | 17.9 | 17.5 | 17.8 | 17.6 | 17.7 | 17.8 |
| 0.787565 | -0.02459 | 0.536648 | -0.05699 | 13.7 | 13.9 | 13.9 | 13.8 | 13.8 | 13.8 | 13.8 | 13.8 | 13.8 |
| 0.786058 | 0.044423 | 0.96612 | 0.006935 | 11.5 | 11 | 11.5 | 11.2 | 11.5 | 11.2 | 11.3 | 11.5 | 11.4 |
| 0.788 | 0.026125 | 0.933658 | 0.00807 | 13.2 | 13 | 13.1 | 13 | 13 | 13.2 | 13.1 | 13.2 | 13.1 |
| 0.788636 | -0.02017 | 0.772832 | -0.02172 | 14.5 | 14.4 | 14.3 | 14.3 | 14.4 | 14.4 | 14.4 | 14.3 | 14.3 |
| 0.789453 | 0.022128 | 0.874188 | -0.0131 | 13.9 | 13.9 | 14.1 | 13.9 | 14 | 14 | 14 | 14 | 13.9 |
| 0.79087 | -0.08664 | 0.186711 | 0.456922 | 10.7 | 10.5 | 9.42 | 10.4 | 10.7 | 11 | 10.8 | 10.5 | 10.6 |
| 0.7908 | -0.06915 | 0.387196 | -0.23065 | 15.8 | 16.5 | 16.5 | 16.4 | 15.7 | 16.2 | 15.9 | 16 | 16.3 |
| 0.792788 | 0.056528 | 0.338738 | -0.21188 | 11.8 | 11.4 | 11.5 | 10.8 | 11.7 | 11.4 | 11.4 | 11.4 | 11.2 |
| 0.792061 | -0.02893 | 0.196211 | -0.14982 | 10.8 | 10.6 | 10.5 | 10.6 | 10.3 | 10.5 | 10.4 | 10.5 | 10.4 |
| 0.792522 | 0.038696 | 0.613879 | -0.07469 | 12.3 | 12.7 | 12.5 | 12.4 | 12.5 | 12.3 | 12.4 | 12.6 | 12.2 |
| 0.795252 | -0.07329 | 0.868692 | -0.04663 | 9.96 | 9.65 | 9.25 | 10 | 9.07 | 9.87 | 9.79 | 9.45 | 9.47 |
| 0.798805 | -0.03535 | 0.259015 | -0.16312 | 12.4 | 12.3 | 12.4 | 12 | 12.5 | 12.2 | 12.1 | 12.2 | 12.3 |
| 0.797099 | -0.02225 | 0.526263 | 0.055462 | 12.7 | 12.8 | 12.8 | 12.9 | 12.9 | 12.8 | 12.7 | 12.9 | 12.9 |
| 0.796433 | 0.049849 | 0.610316 | -0.09912 | 12.8 | 12.2 | 12.4 | 12.4 | 12.6 | 12 | 12.3 | 12.4 | 12.4 |
| 0.798122 | -0.02101 | 0.35299 | -0.07841 | 12.4 | 12.5 | 12.4 | 12.4 | 12.2 | 12.4 | 12.3 | 12.4 | 12.3 |
| 0.799156 | 0.067773 | 0.761518 | 0.080912 | 15.2 | 15.9 | 15.8 | 15.9 | 15.2 | 15.9 | 15.5 | 15.7 | 15.9 |
| 0.799072 | 0.043891 | 0.782451 | 0.04763 | 12.9 | 12.5 | 12.5 | 12.7 | 12.9 | 12.4 | 12.8 | 12.5 | 12.7 |
| 0.800222 | 0.039415 | 0.498887 | -0.10674 | 13.1 | 13.6 | 13.6 | 13.4 | 13.2 | 13.2 | 13.3 | 13.3 | 13.4 |
| 0.800303 | 0.02965 | 0.608929 | -0.06037 | 15.7 | 15.5 | 15.5 | 15.3 | 15.6 | 15.5 | 15.5 | 15.6 | 15.4 |
| 0.802589 | -0.03331 | 0.309828 | -0.13985 | 11.9 | 11.9 | 11.6 | 11.9 | 11.6 | 11.6 | 11.7 | 11.8 | 11.5 |
| 0.80181 | -0.72821 | 0.358459 | -2.7358 | 8.2 | 7.65 | 7.27 | -0.2 | 9.02 | 8.32 | -0.1 | 7.37 | 7.64 |
| 0.801342 | 0.023044 | 0.809586 | -0.02206 | 14.9 | 14.7 | 14.8 | 14.7 | 14.9 | 14.8 | 14.9 | 14.7 | 14.7 |
| 0.802177 | 0.034135 | 0.949391 | 0.008631 | 15.4 | 15.8 | 15.8 | 15.8 | 15.5 | 15.7 | 15.7 | 15.7 | 15.8 |
| 0.803582 | 0.018172 | 0.837889 | -0.01494 | 14.4 | 14.4 | 14.4 | 14.4 | 14.5 | 14.3 | 14.5 | 14.4 | 14.4 |
| 0.80399 | 0.023648 | 0.912421 | 0.010462 | 14.1 | 14 | 14 | 13.9 | 14.2 | 13.9 | 14 | 14 | 14.1 |
| 0.808082 | 0.036348 | 0.087052 | 0.282909 | 10.2 | 10.3 | 10.1 | 10.7 | 10.3 | 10.4 | 10.4 | 10.8 | 10.4 |
| 0.808085 | 0.068293 | 0.229464 | -0.35429 | 10.1 | 10.4 | 10.1 | 10.3 | 9.9 | 9.14 | 9.82 | 9.63 | 10.1 |
| 0.807708 | 0.074706 | 0.505432 | 0.207095 | 12 | 13 | 12.8 | 12.9 | 12.2 | 13 | 12.6 | 12.7 | 13 |
| 0.807444 | -0.02496 | 0.811215 | -0.02446 | 15.1 | 15.3 | 15.3 | 15.2 | 15.2 | 15.4 | 15.1 | 15.2 | 15.3 |
| 0.809832 | -0.06724 | 0.935312 | 0.022638 | 9.23 | 9.56 | 9.99 | 10.2 | 9.44 | 9.41 | 9.45 | 9.9 | 9.5 |
| 0.809735 | -0.02589 | 0.601071 | -0.05663 | 12.8 | 12.9 | 12.9 | 12.8 | 12.9 | 12.8 | 12.6 | 12.8 | 13 |
| 0.809704 | 0.025434 | 0.985742 | 0.001884 | 13.6 | 13.9 | 13.9 | 13.7 | 13.8 | 13.8 | 13.8 | 13.9 | 13.8 |
| 0.812648 | -0.06192 | 0.36504 | -0.24288 | 11.4 | 11 | 11 | 11.2 | 11.2 | 10.5 | 11.3 | 10.6 | 10.8 |
| 0.812312 | -0.04046 | 0.425967 | -0.13833 | 12.3 | 12.1 | 11.9 | 11.8 | 12.2 | 11.9 | 11.7 | 12.2 | 12 |
| 0.813005 | -0.02724 | 0.992933 | 0.001018 | 13.8 | 13.7 | 13.5 | 13.7 | 13.9 | 13.6 | 13.8 | 13.7 | 13.6 |
| 0.811871 | -0.04058 | 0.99009 | -0.00211 | 9.74 | 9.94 | 9.9 | 10.2 | 9.62 | 9.9 | 9.74 | 10.1 | 9.72 |
| 0.814788 | -0.01633 | 0.909547 | 0.007907 | 15 | 15 | 15 | 15.1 | 15 | 15.1 | 15 | 15 | 15.1 |
| 0.816127 | -0.04876 | 0.284508 | -0.23296 | 11.3 | 10.5 | 10.7 | 10.8 | 10.4 | 10.7 | 10.6 | 10.8 | 10.4 |
| 0.816241 | -0.05668 | 0.659377 | 0.108017 | 9.9 | 10.4 | 10.4 | 10.8 | 9.93 | 10.5 | 10.4 | 10.5 | 10.1 |
| 0.819765 | -0.02904 | 0.211716 | -0.1676 | 13.7 | 13.9 | 13.9 | 13.8 | 13.4 | 13.9 | 13.7 | 13.6 | 13.7 |
| 0.818856 | -0.03661 | 0.438274 | 0.126239 | 11.7 | 11.9 | 11.7 | 11.6 | 12.1 | 12.1 | 12.1 | 11.9 | 11.7 |
| 0.819399 | -0.02486 | 0.729352 | 0.037763 | 13.2 | 13.1 | 13.1 | 13.3 | 13.1 | 13.2 | 13 | 13.2 | 13.3 |

| | | | | | | | | | | | | |
|---|---|---|---|---|---|---|---|---|---|---|---|---|
| 0.81872 | -0.05408 | 0.707592 | -0.08876 | 12.4 | 13 | 12.7 | 12.8 | 12.2 | 13 | 12.4 | 12.7 | 12.7 |
| 0.820148 | -0.03333 | 0.859396 | -0.02595 | 18.2 | 17.8 | 17.8 | 17.7 | 18.2 | 17.8 | 17.9 | 17.9 | 17.9 |
| 0.821065 | 0.015088 | 0.642278 | -0.03116 | 17.9 | 17.8 | 17.8 | 17.8 | 17.8 | 17.7 | 17.8 | 17.8 | 17.8 |
| 0.829628 | 0.05132 | 0.336266 | -0.23629 | 10.3 | 10.6 | 10.7 | 10.3 | 9.79 | 10.6 | 10.4 | 10 | 10.5 |
| 0.822452 | 0.025356 | 0.360258 | -0.10618 | 14.2 | 13.9 | 14 | 14 | 13.9 | 13.9 | 14 | 13.9 | 13.8 |
| 0.831936 | 0.031001 | 0.383157 | -0.13053 | 12.6 | 12.6 | 12.4 | 12.6 | 12.1 | 12.4 | 12.3 | 12.4 | 12.4 |
| 0.830599 | 0.661066 | 0.467845 | -2.28023 | 8.54 | 8.63 | 6.9 | -0.3 | 7.58 | 7.96 | 8.82 | 8.96 | -0.5 |
| 0.831272 | -0.03426 | 0.581607 | 0.089391 | 11.6 | 11.1 | 11.2 | 11.7 | 11.2 | 11.5 | 11.5 | 11.4 | 11.4 |
| 0.826299 | -0.04 | 0.61166 | 0.093199 | 12.3 | 12.5 | 12.7 | 12.7 | 12.3 | 12.9 | 12.5 | 12.6 | 12.7 |
| 0.825894 | -0.11812 | 0.355473 | -0.50995 | 9.5 | 8.53 | 9.26 | 9.41 | 9.18 | 7.53 | 8.01 | 8.88 | 8.87 |
| 0.831783 | 0.049948 | 0.398232 | 0.203207 | 11.6 | 11.5 | 11.5 | 12.1 | 11.7 | 11.2 | 11.6 | 12 | 11.6 |
| 0.826796 | -0.05399 | 0.445487 | -0.19171 | 10.7 | 10.3 | 10.8 | 10.5 | 10.2 | 10.7 | 10 | 10.4 | 10.8 |
| 0.825709 | 0.033579 | 0.533806 | 0.095971 | 12.9 | 12.3 | 12.5 | 12.7 | 12.7 | 12.5 | 12.7 | 12.7 | 12.6 |
| 0.8269 | -0.01849 | 0.859374 | 0.014973 | 13.4 | 13.3 | 13.2 | 13.2 | 13.4 | 13.4 | 13.3 | 13.3 | 13.2 |
| 0.827453 | -0.01911 | 0.572797 | -0.04989 | 17 | 16.8 | 16.8 | 16.8 | 17 | 16.8 | 16.9 | 16.8 | 16.8 |
| 0.826604 | 0.032594 | 0.710851 | 0.055326 | 10.7 | 11.1 | 11 | 11.1 | 10.7 | 11.1 | 11.1 | 11 | 10.9 |
| 0.832451 | 0.046721 | 0.721735 | 0.078868 | 11.4 | 11.8 | 12.1 | 12.1 | 11.4 | 11.8 | 11.7 | 11.8 | 12 |
| 0.829153 | 0.095282 | 0.739541 | 0.147116 | 14.4 | 15.8 | 15.7 | 15.7 | 14.6 | 15.8 | 15.4 | 15.4 | 15.6 |
| 0.829487 | 0.018515 | 0.748437 | 0.027624 | 14 | 14 | 14 | 13.9 | 14.2 | 14 | 14 | 14.1 | 14 |
| 0.823909 | -0.02874 | 0.77378 | -0.03717 | 12.5 | 12.6 | 12.5 | 12.5 | 12.6 | 12.4 | 12.7 | 12.4 | 12.3 |
| 0.830994 | 0.033066 | 0.808573 | 0.037551 | 14.5 | 13.9 | 14 | 14.1 | 14.3 | 14.1 | 14.4 | 14.2 | 14 |
| 0.830611 | -0.06655 | 0.829137 | -0.06714 | 14.4 | 15.4 | 15.3 | 15.3 | 14.5 | 15.3 | 14.8 | 15 | 15 |
| 0.83342 | -0.05363 | 0.499548 | -0.17458 | 11.5 | 11.3 | 11.4 | 11.2 | 10.9 | 11.7 | 11.1 | 11.6 | 10.9 |
| 0.835684 | 0.031687 | 0.299642 | 0.16413 | 12.5 | 12.7 | 12.6 | 12.7 | 12.5 | 13 | 12.5 | 12.8 | 12.9 |
| 0.835967 | -0.03145 | 0.711254 | -0.0564 | 13.1 | 12.6 | 12.7 | 12.7 | 12.9 | 12.8 | 12.9 | 12.7 | 12.6 |
| 0.838069 | 0.045818 | 0.776514 | 0.063725 | 11.7 | 12.3 | 12.4 | 12.5 | 11.8 | 12.1 | 12.1 | 12.2 | 12.3 |
| 0.83993 | -0.05345 | 0.331557 | 0.264957 | 9.7 | 8.85 | 9.2 | 9.19 | 9.99 | 9.53 | 9.64 | 9.38 | 9.53 |
| 0.840509 | 0.038145 | 0.332966 | -0.1892 | 13.1 | 12.8 | 12.6 | 12.4 | 13 | 12.4 | 12.8 | 12.6 | 12.5 |
| 0.84147 | 0.097581 | 0.279656 | 0.548142 | 9.86 | 8.14 | 10.1 | 9.81 | 9.44 | 10.2 | 9.84 | 9.99 | 9.91 |
| 0.844391 | 0.038543 | 0.824109 | -0.04366 | 11.7 | 11.6 | 11.3 | 11.3 | 11.8 | 11.2 | 11.3 | 11.7 | 11.5 |
| 0.844981 | 0.053806 | 0.558288 | 0.162758 | 11.9 | 11.4 | 11.3 | 11.3 | 12.1 | 11.7 | 12.1 | 11.6 | 11.5 |
| 0.843584 | -0.03609 | 0.858603 | 0.032576 | 12.7 | 13.2 | 13 | 13.3 | 12.7 | 13.1 | 13.1 | 12.9 | 13 |
| 0.845083 | -0.05288 | 0.784179 | -0.07421 | 12 | 11.6 | 11.2 | 11.2 | 11.7 | 11.9 | 11.3 | 11.9 | 11.4 |
| 0.850074 | 0.033006 | 0.410422 | -0.1469 | 11.8 | 11.7 | 12 | 11.6 | 12 | 11.3 | 11.7 | 11.7 | 11.6 |
| 0.847782 | -0.06751 | 0.301935 | -0.37601 | 9.07 | 9.73 | 8.86 | 9.11 | 8.71 | 8.91 | 9.4 | 8.98 | 8.14 |
| 0.847858 | 0.014508 | 0.544798 | -0.04631 | 14.7 | 14.7 | 14.8 | 14.7 | 14.7 | 14.7 | 14.6 | 14.6 | 14.8 |
| 0.849719 | -0.02018 | 0.496888 | -0.0734 | 14.4 | 14.2 | 14.2 | 14.4 | 14.3 | 14.1 | 14.2 | 14.3 | 14.1 |
| 0.849588 | -0.04491 | 0.508079 | -0.15888 | 10.5 | 10.9 | 11.1 | 10.9 | 10.2 | 11 | 10.8 | 10.6 | 10.6 |
| 0.848402 | 0.02012 | 0.921395 | -0.01038 | 13.9 | 14 | 14.2 | 14.1 | 14.1 | 13.9 | 14 | 14.1 | 14 |
| 0.851443 | -0.02443 | 0.434344 | -0.10401 | 13.1 | 13.1 | 13.4 | 13.3 | 13.2 | 12.9 | 13.1 | 13 | 13.3 |
| 0.852868 | -0.02011 | 0.161903 | -0.16199 | 13 | 12.8 | 12.9 | 12.8 | 12.9 | 12.6 | 12.8 | 12.7 | 12.7 |
| 0.852238 | -0.01648 | 0.215862 | -0.11524 | 14.6 | 14.5 | 14.5 | 14.5 | 14.5 | 14.3 | 14.4 | 14.4 | 14.5 |
| 0.853292 | 0.018562 | 0.354474 | 0.095583 | 9.94 | 10.2 | 10.2 | 10.2 | 10.2 | 10.2 | 10.1 | 10.3 | 10.3 |
| 0.854268 | 0.037908 | 0.820084 | -0.04697 | 15.1 | 15.6 | 15.7 | 15.6 | 15 | 15.5 | 15.3 | 15.4 | 15.5 |
| 0.857791 | -0.01563 | 0.125435 | 0.144794 | 12.6 | 12.6 | 12.5 | 12.9 | 12.6 | 12.8 | 12.7 | 12.7 | 12.8 |
| 0.857753 | -0.01667 | 0.538114 | 0.057934 | 14.6 | 14.3 | 14.3 | 14.4 | 14.6 | 14.5 | 14.5 | 14.5 | 14.4 |
| 0.86141 | -0.01384 | 0.06952 | -0.16122 | 13.6 | 13.7 | 13.8 | 13.6 | 13.6 | 13.6 | 13.6 | 13.5 | 13.6 |
| 0.861877 | 0.019092 | 0.191848 | -0.15167 | 14.6 | 14.3 | 14.3 | 14.2 | 14.3 | 14.2 | 14.3 | 14.3 | 14.1 |
| 0.861642 | -0.02241 | 0.952901 | 0.007588 | 14.7 | 14.3 | 14.4 | 14.4 | 14.7 | 14.4 | 14.5 | 14.4 | 14.5 |
| 0.863613 | 0.013115 | 0.074177 | -0.15218 | 12.2 | 12.3 | 12.3 | 12.1 | 12 | 12.1 | 12.1 | 12.1 | 12 |

| | | | | | | | | | | | | |
|---|---|---|---|---|---|---|---|---|---|---|---|---|
| 0.863702 | -0.03114 | 0.43288 | -0.14512 | 9.83 | 9.18 | 9.2 | 9.22 | 9.35 | 9.3 | 9.39 | 9.29 | 9.1 |
| 0.863559 | -0.01829 | 0.534005 | -0.06701 | 13.2 | 13.1 | 13.1 | 13.3 | 13.2 | 12.9 | 13.1 | 13 | 13.1 |
| 0.865593 | 0.014794 | 0.964406 | 0.003897 | 14.1 | 14.3 | 14.3 | 14.3 | 14.1 | 14.3 | 14.2 | 14.2 | 14.2 |
| 0.8662 | -0.01436 | 0.64054 | -0.04006 | 16.3 | 16.1 | 16.1 | 16.1 | 16.2 | 16.1 | 16.1 | 16.1 | 16.1 |
| 0.867879 | -0.02346 | 0.802013 | -0.0354 | 12.1 | 12 | 11.8 | 12 | 12.2 | 11.7 | 11.9 | 12.1 | 11.8 |
| 0.869119 | 0.027274 | 0.846598 | 0.032033 | 13.2 | 13.3 | 13.3 | 13.3 | 12.9 | 13.6 | 13.4 | 13.3 | 13.3 |
| 0.87401 | -0.07545 | 0.204669 | 0.637137 | 10.3 | 8.39 | 9.5 | 9.61 | 10.6 | 10.1 | 10.2 | 9.9 | 10 |
| 0.874032 | -0.04124 | 0.744055 | 0.085172 | 10.6 | 10.4 | 10.3 | 10.3 | 11.2 | 10.2 | 10.5 | 10.3 | 10.8 |
| 0.875049 | 0.062897 | 0.947043 | -0.02655 | 12.7 | 13.4 | 13.3 | 13.5 | 12.2 | 13.5 | 12.8 | 13 | 13.5 |
| 0.87581 | 0.092664 | 0.551106 | 0.357423 | 7.57 | 9.38 | 9.29 | 9.62 | 9.47 | 7.95 | 9.24 | 8.94 | 9.14 |
| 0.87749 | 0.320973 | 0.222847 | 2.668434 | 0.36 | 8.48 | 8.99 | 8.47 | 7.67 | 8.73 | 8.5 | 8.59 | 8.75 |
| 0.877971 | 0.059892 | 0.469549 | -0.28692 | 9 | 9.32 | 9.23 | 9.52 | 8.05 | 8.94 | 8.44 | 9.42 | 8.82 |
| 0.877204 | -0.02164 | 0.545459 | 0.085663 | 12.8 | 12.4 | 12.3 | 12.6 | 12.8 | 12.5 | 12.7 | 12.5 | 12.6 |
| 0.878811 | 0.014149 | 0.605528 | -0.04832 | 14.2 | 14.3 | 14.2 | 14.3 | 14 | 14.2 | 14.1 | 14.3 | 14.1 |
| 0.882727 | 0.035134 | 0.071337 | -0.4807 | 10 | 9.74 | 9.74 | 9.66 | 9.51 | 8.8 | 9.23 | 9.23 | 9.62 |
| 0.882388 | -0.01512 | 0.071379 | -0.20625 | 12.6 | 12.5 | 12.4 | 12.2 | 12.5 | 12.3 | 12.3 | 12.3 | 12.4 |
| 0.882385 | -0.01785 | 0.884811 | -0.01748 | 15.5 | 15.2 | 15.1 | 15.2 | 15.4 | 15.2 | 15.2 | 15.2 | 15.4 |
| 0.884088 | -0.0265 | 0.884812 | -0.02634 | 11.9 | 11.9 | 11.7 | 12 | 12.1 | 11.6 | 12.2 | 11.7 | 11.7 |
| 0.88641 | -0.06678 | 0.979807 | 0.011825 | 12.2 | 13.6 | 13.3 | 13.6 | 12.3 | 13.5 | 12.7 | 13.1 | 13.4 |
| 0.887197 | 0.014385 | 0.274045 | -0.11547 | 14.7 | 14.6 | 14.4 | 14.4 | 14.6 | 14.4 | 14.5 | 14.5 | 14.4 |
| 0.888543 | -0.01898 | 0.755848 | -0.04223 | 13.9 | 13.6 | 13.7 | 13.5 | 14 | 13.7 | 13.8 | 13.7 | 13.7 |
| 0.890359 | 0.010712 | 0.462317 | -0.05814 | 14.5 | 14.5 | 14.5 | 14.5 | 14.5 | 14.4 | 14.4 | 14.4 | 14.6 |
| 0.890768 | 0.014973 | 0.958738 | -0.00564 | 14.1 | 13.8 | 13.7 | 13.8 | 13.9 | 13.9 | 13.8 | 13.9 | 14 |
| 0.891011 | 0.022789 | 0.879779 | 0.025158 | 15.4 | 15.8 | 15.7 | 15.7 | 15.4 | 15.8 | 15.4 | 15.7 | 15.8 |
| 0.893509 | -0.01137 | 0.45191 | -0.06509 | 14.7 | 14.6 | 14.7 | 14.7 | 14.4 | 14.6 | 14.6 | 14.6 | 14.6 |
| 0.893348 | 0.014063 | 0.731328 | -0.03614 | 11.4 | 11.5 | 11.7 | 11.5 | 11.3 | 11.6 | 11.6 | 11.5 | 11.5 |
| 0.892687 | -0.01323 | 0.887493 | 0.013879 | 11.8 | 11.8 | 11.9 | 11.8 | 11.9 | 11.9 | 11.7 | 11.9 | 12 |
| 0.893049 | 0.023038 | 0.995465 | -0.00097 | 11 | 10.8 | 11 | 10.7 | 11.1 | 11 | 11.3 | 10.9 | 10.7 |
| 0.894155 | 0.04899 | 0.434716 | 0.293465 | 9.85 | 10.9 | 11 | 11.2 | 10.2 | 11.1 | 10.8 | 10.8 | 11.1 |
| 0.894735 | 0.02205 | 0.697051 | -0.06517 | 8.62 | 8.68 | 8.62 | 8.31 | 8.45 | 8.9 | 8.59 | 8.38 | 8.76 |
| 0.897686 | 0.012717 | 0.178334 | -0.14157 | 13.3 | 13.4 | 13.4 | 13.4 | 13.1 | 13.2 | 13.3 | 13.1 | 13.3 |
| 0.896673 | -0.01161 | 0.142345 | -0.14122 | 14.1 | 13.9 | 14 | 13.8 | 13.9 | 13.9 | 13.9 | 13.8 | 13.9 |
| 0.898175 | -0.02277 | 0.145452 | -0.27848 | 11.8 | 11.5 | 11.8 | 11.1 | 11.5 | 11.7 | 11.4 | 11.5 | 11.3 |
| 0.897826 | -0.01136 | 0.663618 | -0.03871 | 14.2 | 14.1 | 14 | 14.2 | 14 | 14.1 | 14.1 | 14.1 | 14 |
| 0.897976 | 0.008862 | 0.718404 | 0.025019 | 13 | 13 | 13.1 | 13.1 | 13.1 | 13.1 | 13.1 | 13.2 | 13.1 |
| 0.901077 | 0.021498 | 0.951879 | 0.010433 | 11.1 | 10.9 | 10.8 | 10.7 | 11.1 | 10.9 | 11.3 | 10.8 | 10.8 |
| 0.901968 | 0.027259 | 0.210186 | -0.29247 | 10.2 | 10.6 | 10.5 | 10.3 | 10.4 | 9.68 | 9.99 | 10.1 | 10.4 |
| 0.903693 | 0.013956 | 0.406559 | -0.09795 | 12.3 | 12.3 | 12.2 | 12.3 | 12 | 12.1 | 12 | 12.2 | 12.3 |
| 0.90313 | 0.013363 | 0.765297 | 0.03286 | 13.3 | 13.7 | 13.6 | 13.5 | 13.5 | 13.6 | 13.5 | 13.5 | 13.6 |
| 0.904415 | -0.02866 | 0.875421 | -0.03742 | 12.1 | 11.3 | 11.5 | 11.5 | 12 | 11.4 | 11.5 | 11.7 | 11.5 |
| 0.905016 | 0.023363 | 0.885374 | 0.028232 | 11.2 | 11.3 | 11.1 | 10.8 | 11.4 | 11.4 | 11.3 | 11.4 | 11 |
| 0.907193 | -0.01961 | 0.217989 | 0.218193 | 12.1 | 11.7 | 11.8 | 12.1 | 12.2 | 12 | 12.4 | 12 | 11.8 |
| 0.907074 | -0.02554 | 0.18606 | -0.3071 | 10.9 | 10.6 | 10.5 | 10.7 | 10.5 | 9.99 | 10.5 | 10.6 | 10 |
| 0.912466 | 0.013088 | 0.294269 | -0.12958 | 12.4 | 12.2 | 12.3 | 12.2 | 11.9 | 12.3 | 12.2 | 12.1 | 12.2 |
| 0.912107 | 0.018506 | 0.252296 | 0.200636 | 11.8 | 12.2 | 12.3 | 12.3 | 12.1 | 12.4 | 12 | 12.4 | 12.5 |
| 0.910544 | -0.01846 | 0.367971 | -0.15201 | 12.5 | 12.5 | 12.2 | 12.5 | 12.5 | 11.9 | 12.3 | 12.3 | 12.2 |
| 0.912177 | 0.012625 | 0.580364 | -0.06391 | 12.5 | 12.3 | 12.5 | 12.4 | 12.3 | 12.5 | 12.5 | 12.5 | 12.2 |
| 0.908657 | -0.01077 | 0.510397 | -0.06268 | 13.8 | 13.8 | 13.8 | 13.8 | 13.6 | 13.9 | 13.8 | 13.7 | 13.8 |
| 0.91019 | 0.044773 | 0.621651 | 0.19742 | 10.3 | 11.8 | 11.7 | 11.6 | 11 | 11.7 | 11.4 | 11.6 | 11.5 |
| 0.9101 | -0.02626 | 0.635217 | -0.11111 | 10.7 | 10.8 | 10.2 | 10.9 | 10.3 | 10.2 | 10.6 | 10.4 | 10.3 |

| | | | | | | | | | | | | |
|---|---|---|---|---|---|---|---|---|---|---|---|---|
| 0.91143 | 0.017025 | 0.907586 | 0.017768 | 12 | 12.5 | 12.3 | 12.3 | 12.4 | 12.1 | 12.3 | 12.2 | 12.4 |
| 0.913924 | 0.01214 | 0.245117 | -0.13666 | 14 | 13.7 | 14 | 13.6 | 13.8 | 13.8 | 13.9 | 13.8 | 13.6 |
| 0.913714 | -0.01738 | 0.344903 | -0.15607 | 14.9 | 15.3 | 15.2 | 15.1 | 14.7 | 15.1 | 14.9 | 14.9 | 15.1 |
| 0.914296 | -0.01451 | 0.382484 | -0.12077 | 12.3 | 12.4 | 12.2 | 12.1 | 12.1 | 12.4 | 12.2 | 12 | 12.4 |
| 0.914992 | -0.01847 | 0.281645 | 0.193712 | 11.9 | 11.4 | 11.4 | 11.7 | 12 | 11.7 | 11.7 | 12 | 11.6 |
| 0.91624 | 0.016081 | 0.753435 | 0.048172 | 13 | 13.4 | 13.4 | 13.5 | 13 | 13.4 | 13.2 | 13.3 | 13.4 |
| 0.919527 | -0.0201 | 0.375691 | 0.180943 | 9.56 | 9.3 | 9.95 | 9.67 | 9.61 | 10.1 | 9.9 | 9.74 | 9.7 |
| 0.918961 | -0.01179 | 0.310368 | -0.12169 | 13.4 | 13.5 | 13.4 | 13.1 | 13.6 | 13.4 | 13.4 | 13.3 | 13.3 |
| 0.918361 | 0.035884 | 0.685355 | -0.14259 | 9.65 | 9.34 | 9.84 | 9.75 | 9.97 | 8.57 | 9.3 | 9.57 | 9.52 |
| 0.919164 | -0.03025 | 0.740009 | 0.099256 | 10.9 | 11.1 | 10.6 | 11.5 | 10.3 | 11.2 | 11 | 10.8 | 11.1 |
| 0.917767 | -0.00787 | 0.644123 | -0.03547 | 12.7 | 12.6 | 12.6 | 12.6 | 12.6 | 12.5 | 12.6 | 12.5 | 12.5 |
| 0.920588 | 0.044476 | 0.505289 | 0.301606 | 9.98 | 8.41 | 9.55 | 9.16 | 10.2 | 9.37 | 9.58 | 9.99 | 9.26 |
| 0.920762 | 0.01092 | 0.714752 | 0.040291 | 13.5 | 13.8 | 13.7 | 13.8 | 13.6 | 13.7 | 13.6 | 13.7 | 13.8 |
| 0.924568 | 0.007805 | 0.250824 | -0.099 | 14.3 | 14.4 | 14.2 | 14.1 | 14.2 | 14.2 | 14.2 | 14.1 | 14.1 |
| 0.924382 | -0.00834 | 0.214194 | -0.11497 | 13.7 | 13.6 | 13.6 | 13.4 | 13.6 | 13.6 | 13.6 | 13.5 | 13.5 |
| 0.92235 | -0.00981 | 0.270336 | -0.11557 | 14.6 | 14.5 | 14.4 | 14.5 | 14.5 | 14.2 | 14.4 | 14.4 | 14.3 |
| 0.92362 | -0.01531 | 0.307938 | -0.16861 | 12.2 | 11.9 | 11.8 | 11.5 | 12 | 11.9 | 11.7 | 11.9 | 11.8 |
| 0.924207 | 0.019672 | 0.315117 | 0.214911 | 10.4 | 9.76 | 10.1 | 10.3 | 10.4 | 10 | 10.5 | 9.94 | 10.4 |
| 0.924159 | -0.01705 | 0.390611 | -0.15765 | 10.1 | 10.8 | 10.6 | 10.1 | 10.4 | 10.6 | 10.3 | 10.3 | 10.4 |
| 0.926814 | -0.03412 | 0.938359 | -0.02873 | 11.1 | 9.81 | 10.5 | 10.6 | 11 | 9.9 | 10.5 | 10.4 | 10.4 |
| 0.930187 | -0.0164 | 0.425477 | 0.152414 | 12.6 | 13.1 | 13.1 | 13.3 | 12.8 | 13.2 | 13.2 | 12.9 | 13.2 |
| 0.929776 | -0.00904 | 0.68052 | -0.04246 | 14 | 14 | 13.8 | 13.9 | 13.8 | 13.9 | 13.9 | 13.9 | 13.7 |
| 0.932034 | -0.00944 | 0.571816 | 0.063213 | 12.8 | 12.7 | 12.5 | 12.7 | 12.9 | 12.6 | 12.8 | 12.7 | 12.6 |
| 0.933878 | -0.01112 | 0.094475 | 0.24685 | 10.9 | 11.1 | 11.3 | 11.6 | 11.3 | 11.2 | 11.3 | 11.3 | 11.5 |
| 0.933643 | -0.00641 | 0.264876 | -0.08948 | 13.9 | 13.8 | 13.8 | 13.8 | 13.8 | 13.7 | 13.8 | 13.7 | 13.7 |
| 0.933193 | -0.01031 | 0.265082 | -0.14297 | 12 | 12.3 | 12.2 | 12.1 | 11.8 | 12.1 | 11.9 | 12.1 | 12.1 |
| 0.934659 | -0.01178 | 0.447463 | -0.1113 | 15.3 | 15 | 15 | 14.8 | 15.3 | 14.9 | 15.1 | 15 | 14.9 |
| 0.935649 | 0.037733 | 0.431462 | 0.375368 | 8.53 | 8.57 | 6.92 | 8.5 | 8.31 | 8.22 | 8.32 | 7.89 | 8.94 |
| 0.936059 | -0.01433 | 0.79393 | -0.04677 | 12.8 | 12.4 | 12.3 | 12.2 | 12.8 | 12.4 | 12.5 | 12.5 | 12.4 |
| 0.939944 | 0.008032 | 0.268675 | -0.12292 | 12.8 | 13 | 13 | 12.8 | 12.9 | 12.6 | 12.8 | 12.8 | 12.7 |
| 0.939587 | 0.012804 | 0.361163 | 0.158673 | 10.2 | 10.5 | 10.1 | 10.5 | 10.7 | 10.1 | 10.3 | 10.4 | 10.6 |
| 0.939739 | -0.01247 | 0.37368 | -0.1507 | 13 | 12.6 | 12.4 | 12.5 | 12.5 | 12.6 | 12.6 | 12.7 | 12.3 |
| 0.94109 | -0.01268 | 0.170427 | -0.25082 | 11.9 | 12.3 | 12.2 | 12.2 | 11.7 | 11.8 | 12.1 | 11.9 | 11.7 |
| 0.946833 | 0.026493 | 0.476277 | 0.287857 | 10.1 | 11 | 9.84 | 9.76 | 11 | 10.9 | 10.5 | 10.6 | 10.6 |
| 0.948904 | 0.006695 | 0.070577 | -0.21169 | 13 | 13 | 13.1 | 13 | 12.8 | 12.8 | 12.7 | 12.9 | 13 |
| 0.948684 | -0.00537 | 0.076502 | -0.16469 | 14.5 | 14.5 | 14.5 | 14.4 | 14.3 | 14.3 | 14.2 | 14.5 | 14.3 |
| 0.949393 | -0.0169 | 0.219477 | -0.34431 | 10.3 | 10.5 | 10.2 | 10 | 9.45 | 10.6 | 9.87 | 9.96 | 10.2 |
| 0.951329 | -0.0083 | 0.069474 | -0.27672 | 12.2 | 12.4 | 12.5 | 12 | 12 | 12.4 | 12.2 | 12 | 12.1 |
| 0.952566 | 0.005792 | 0.681141 | -0.04023 | 14.4 | 14.6 | 14.6 | 14.6 | 14.4 | 14.4 | 14.4 | 14.5 | 14.6 |
| 0.953298 | -0.00742 | 0.539035 | 0.078841 | 10.6 | 11 | 10.8 | 10.8 | 10.9 | 11 | 11 | 10.7 | 11 |
| 0.953835 | 0.020585 | 0.957546 | -0.01893 | 10.1 | 10.8 | 10.8 | 10.9 | 9.78 | 10.9 | 10.3 | 10.4 | 10.9 |
| 0.955781 | -0.0066 | 0.305285 | -0.12641 | 14.7 | 14.5 | 14.4 | 14.3 | 14.6 | 14.3 | 14.5 | 14.4 | 14.3 |
| 0.956133 | -0.01126 | 0.480732 | 0.146746 | 11.6 | 11.4 | 11.1 | 11.5 | 11.8 | 11.1 | 11.3 | 11.6 | 11.6 |
| 0.956266 | -0.02012 | 0.908284 | 0.042272 | 10.5 | 11.6 | 11.5 | 11.4 | 10.9 | 11.5 | 10.6 | 11.5 | 11.6 |
| 0.958776 | 0.008472 | 0.969957 | 0.006173 | 11.4 | 11.4 | 11.7 | 11.2 | 11.7 | 11.6 | 11.5 | 11.6 | 11.4 |
| 0.961131 | 0.011151 | 0.53624 | 0.143348 | 12.1 | 12.4 | 11.9 | 12.4 | 12.6 | 11.7 | 12.2 | 12.3 | 12.3 |
| 0.964808 | 0.005477 | 0.840477 | -0.02502 | 12.7 | 12.4 | 12.7 | 12.5 | 12.4 | 12.8 | 12.6 | 12.5 | 12.6 |
| 0.968086 | 0.008581 | 0.106249 | 0.379172 | 10.6 | 10 | 10.5 | 10.9 | 10.4 | 11 | 10.9 | 10.5 | 10.8 |
| 0.967223 | -0.00526 | 0.442088 | 0.100369 | 12 | 11.7 | 12.1 | 12 | 12.1 | 12 | 12 | 12.1 | 11.9 |
| 0.969185 | 0.005237 | 0.443789 | -0.10591 | 11.4 | 11.3 | 11.5 | 11.2 | 11.2 | 11.4 | 11.5 | 11 | 11.3 |

| | | | | | | | | | | | | |
|---|---|---|---|---|---|---|---|---|---|---|---|---|
| 0.96854 | 0.010077 | 0.555965 | 0.152217 | 9.19 | 9.31 | 9.38 | 9.38 | 9.67 | 9.25 | 8.86 | 9.91 | 9.56 |
| 0.968748 | -0.00572 | 0.632985 | 0.070289 | 12.2 | 12.6 | 12.7 | 12.6 | 12.4 | 12.7 | 12.5 | 12.6 | 12.7 |
| 0.966279 | 0.014529 | 0.718764 | -0.1243 | 9.49 | 9.03 | 9 | 9 | 8.38 | 9.74 | 8.78 | 9.01 | 9.37 |
| 0.968006 | -0.00778 | 0.687757 | -0.07838 | 9.01 | 8.64 | 9.18 | 8.69 | 8.89 | 9.03 | 9.12 | 8.55 | 8.91 |
| 0.96997 | 0.006883 | 0.243544 | 0.22336 | 9.8 | 9.85 | 9.84 | 9.74 | 9.89 | 10.5 | 10.1 | 10 | 10.1 |
| 0.971875 | 0.014066 | 0.822789 | -0.0895 | 10.8 | 10.1 | 11 | 10.8 | 11.1 | 9.61 | 10.6 | 10.5 | 10.4 |
| 0.974177 | -0.00305 | 0.208274 | -0.12527 | 13.7 | 13.7 | 13.9 | 13.7 | 13.5 | 13.7 | 13.7 | 13.6 | 13.6 |
| 0.973732 | -0.00714 | 0.854518 | 0.039795 | 11.9 | 12.4 | 12.5 | 12.5 | 12 | 12.4 | 12.1 | 12.2 | 12.6 |
| 0.973349 | 0.003852 | 0.971477 | -0.00412 | 14.1 | 13.9 | 14 | 13.9 | 14.1 | 13.9 | 14.2 | 13.9 | 13.9 |
| 0.975506 | 0.003598 | 0.112588 | -0.20282 | 12.3 | 12.3 | 11.9 | 11.9 | 11.9 | 12 | 12 | 11.9 | 11.9 |
| 0.975675 | 0.009901 | 0.558396 | -0.19223 | 10.8 | 9.91 | 10.5 | 10.3 | 10.4 | 9.88 | 10.8 | 9.88 | 9.97 |
| 0.977447 | 0.002554 | 0.297645 | -0.09762 | 14.1 | 14.2 | 14.2 | 14.2 | 14 | 14 | 13.9 | 14.1 | 14.1 |
| 0.977887 | 0.002389 | 0.650622 | -0.0393 | 15.1 | 15 | 14.9 | 15 | 15 | 14.9 | 15 | 15 | 15 |
| 0.978119 | -0.00485 | 0.935863 | -0.01423 | 13.6 | 13.2 | 13.3 | 13.2 | 13.6 | 13.2 | 13.6 | 13.4 | 13.1 |
| 0.980853 | 0.004085 | 0.301898 | -0.18221 | 14.2 | 13.7 | 13.7 | 13.6 | 13.9 | 13.4 | 13.7 | 13.6 | 13.6 |
| 0.979246 | -0.00676 | 0.449143 | 0.200439 | 10.6 | 11.1 | 10.9 | 11.1 | 10.5 | 11.5 | 10.8 | 11.3 | 11 |
| 0.980664 | -0.00332 | 0.542695 | 0.084434 | 11.8 | 12 | 11.8 | 12.1 | 12.1 | 11.6 | 11.9 | 12 | 11.9 |
| 0.980696 | -0.00354 | 0.755463 | 0.045699 | 11.2 | 10.9 | 10.6 | 11 | 11.1 | 10.8 | 10.9 | 11 | 11 |
| 0.981853 | -0.00217 | 0.600641 | -0.0504 | 12.7 | 12.4 | 12.6 | 12.5 | 12.5 | 12.5 | 12.6 | 12.4 | 12.4 |
| 0.982725 | 0.007451 | 0.898238 | -0.04403 | 10.3 | 9.7 | 9.91 | 9.56 | 10.4 | 9.76 | 10.5 | 9.34 | 9.99 |
| 0.984127 | -0.00194 | 0.001684 | -0.44163 | 11.8 | 11.7 | 11.6 | 11.3 | 11.3 | 11.3 | 11.1 | 11.4 | 11.3 |
| 0.984564 | -0.00411 | 0.668067 | 0.091659 | 12.6 | 13 | 12.9 | 13.2 | 12.5 | 13.2 | 12.8 | 12.9 | 13.1 |
| 0.984371 | -0.00242 | 0.810187 | -0.02979 | 12.1 | 12.1 | 12 | 12.2 | 12 | 11.9 | 12.1 | 12.2 | 11.9 |
| 0.985675 | 0.006102 | 0.063586 | -0.71093 | 10 | 10.6 | 10.4 | 10.3 | 9.55 | 9.04 | 9.61 | 9.36 | 9.98 |
| 0.987185 | 0.002946 | 0.121681 | -0.30832 | 10.2 | 10.4 | 10.7 | 9.79 | 10.4 | 10.2 | 10 | 10.3 | 10 |
| 0.988165 | -0.0087 | 0.675962 | 0.24648 | 8.42 | 7.1 | 9.1 | 8.97 | 8.97 | 7.43 | 8.18 | 8.57 | 8.6 |
| 0.992082 | 0.001135 | 0.283155 | -0.12771 | 13.2 | 13.1 | 13.2 | 13 | 13.2 | 12.8 | 13 | 13.1 | 13 |
| 0.994219 | -0.00094 | 0.888398 | 0.018239 | 11.7 | 11.8 | 11.7 | 11.7 | 11.7 | 11.9 | 12 | 11.7 | 11.6 |
| 0.993893 | 0.002903 | 0.909524 | 0.043126 | 8.7 | 9.82 | 9.46 | 9.08 | 9.28 | 9.75 | 9.98 | 9.28 | 8.85 |
| 0.995241 | 0.003037 | 0.747483 | 0.164496 | 10.1 | 10.9 | 9.24 | 10.6 | 10.7 | 9.45 | 9.82 | 10.3 | 10.7 |
| 0.997881 | -0.00102 | 0.819152 | 0.087541 | 12.8 | 13.7 | 13.9 | 14 | 12.8 | 13.8 | 13.3 | 13.5 | 13.8 |
| 0.996869 | -0.00112 | 0.929533 | -0.02519 | 12.2 | 13 | 12.7 | 12.9 | 12.1 | 12.8 | 12.5 | 12.5 | 12.7 |
| 0.997395 | 0.001264 | 0.944381 | -0.02701 | 11.8 | 11.6 | 10.6 | 11.4 | 11.1 | 11.4 | 11.8 | 10.7 | 11.5 |
| 0.999708 | -5.8E-05 | 0.813258 | -0.03742 | 14.1 | 14.5 | 14.5 | 14.5 | 14.1 | 14.4 | 14.3 | 14.3 | 14.4 |
| 0.999648 | 6.92E-05 | 0.922248 | 0.015296 | 15.7 | 16.1 | 16.1 | 16.2 | 15.7 | 16.1 | 15.9 | 16 | 16.1 |
